# Re-annotation of the *Theileria parva* genome refines 53% of the proteome and uncovers essential components of N-glycosylation, a conserved pathway in many organisms

**DOI:** 10.1101/749366

**Authors:** Kyle Tretina, Roger Pelle, Joshua Orvis, Hanzel T. Gotia, Olukemi O. Ifeonu, Priti Kumari, Nicholas C. Palmateer, Shaikh B.A. Iqbal, Lindsay Fry, Vishvanath M. Nene, Claudia Daubenberger, Richard P. Bishop, Joana C. Silva

## Abstract

**Background:** The apicomplexan parasite *Theileria parva* causes a livestock disease called East coast fever (ECF), with millions of animals are at risk in sub-Saharan East and Southern Africa, the geographic distribution of *T. parva*. Over a million bovines die each year of ECF, with a tremendous economic burden to pastoralists in endemic countries. Comprehensive, accurate parasite genome annotation can facilitate the discovery of novel chemotherapeutic targets for disease treatment, as well as elucidate the biology of the parasite. However, genome annotation remains a significant challenge because of limitations in the quality and quantity of the data being used to inform the location and function of protein-coding genes and, when RNA data are used, the underlying biological complexity of the processes involved in gene expression. Here, we apply our recently published RNAseq dataset derived from the schizont life-cycle stage of *T. parva* to update structural and functional gene annotations across the entire nuclear genome.

**Results:** The re-annotation effort lead to evidence-supported updates in over half of all protein-coding sequence (CDS) predictions, including exon changes, gene merges and gene splitting, an increase in average CDS length of approximately 50 base pairs, and the identification of 128 new genes. Among the new genes identified were those involved in N-glycosylation, a process previously thought not to exist in this organism and a potentially new chemotherapeutic target pathway for treating ECF. Alternatively-spliced genes were identified, and antisense and multi-gene family transcription were extensively characterized.

**Conclusions:** The process of re-annotation led to novel insights into the organization and expression profiles of protein-coding sequences in this parasite, and uncovered a minimal N-glycosylation pathway that changes our current understanding of the evolution of this post-translation modification in apicomplexan parasites.

## Background

East Coast fever (ECF) in eastern, central, and southern Africa causes an estimated loss of over 1 million heads of cattle yearly, with an annual economic loss that surpasses $300 million USD, impacting mainly smallholder farmers [1]. Cattle are the most valuable possession of smallholder farmers in this region, as they are a source of milk, meat and hides, provide manure and traction in mixed crop-livestock systems, and revenue derived from livestock pays for school fees and dowries [2, 3]. ECF is a tick-transmitted disease caused by the apicomplexan parasite *Theileria parva*. Lymphocytes infected with *T. parva* proliferate in the regional lymph node draining the tick bite site, and then metastasize into various lymphoid and non-lymphoid organs, and induce a severe inflammatory reaction that leads to respiratory failure and death of susceptible cattle, which typically die within three to four weeks of infection [4–7]. *T. parva* control is vital to food security in this region of the world, which is plagued by a range of other infectious diseases of humans and their livestock.

Efficacious and affordable chemotherapeutics and vaccines are essential tools in the effective control of infectious disease agents [8, 9]. A reliable structural annotation of the genome, consisting at minimum of the correct location of all protein-coding sequences (CDSs), enables the identification, prioritization and experimental screening of potential vaccine and drug targets [10–12]. The accurate identification of the complete proteome can greatly enhance microbiological studies, and reveals metabolic processes unique to pathogens [13]. In turn, a better understanding of the biology of *T. parva* transmission, colonization and pathogenesis may ultimately reveal novel targets for pathogen control [14]. Currently, much like for other apicomplexan parasites [15, 16], knowledge on the functional role of genomic sequences outside of *T. parva* CDSs is sparse, and many gene models containing only CDSs are supported by little or no experimental evidence. RNAseq data, generated through deep sequencing of cDNA using next generation sequencing technologies, can provide an extraordinary level of insight into gene structure and regulation [12, 17]. Here, we used the first high-coverage RNAseq data for this species [18] to improve existing gene models through the identification of start and stop codons, primary intron splice sites and untranslated regions (UTRs). While RNAseq data exists in publicly available databases for other, closely related pathogens, such as *Theileria annulata* and *Babesia bovis*, recent systematic re-annotation efforts for these genomes have yet to be published. This new gene model annotation brought to light several new insights into gene expression in this gene-dense eukaryote, and led to the discovery of several new prospective chemotherapeutic targets for treating ECF.

## Results

### The annotation of the *Theileria parva* genome is significantly improved, revealing a higher gene density than previously thought

The nuclear genome of the reference *T. parva* Muguga isolate consists of four linear chromosomes which are currently assembled into eight contigs (Supplementary Table S1, Additional File 1): chromosomes 1 and 2 are assembled into a single contig each, chromosome 3 is in four contigs and chromosome 4 in two [19]. The new genome annotation was based on this assembly and on extensive RNAseq data (Supplementary Figure S1, Additional File 1). We performed a comprehensive revision of the entire *T. parva* genome annotation, including automated structural annotation and a double-pass manual curation of each locus (see Methods).

The re-annotation process resulted in the discovery of 128 new genes, 274 adjacent gene models were merged, 157 gene models were split, and 38 genes were replaced by new genes encoded in the reverse orientation (Figure 1). In addition, exon boundaries have been corrected in over a thousand genes. Overall, 83% of all nuclear genes in the original annotation were altered in some way, with changes made on every contig. This resulted in significant alterations to the predicted proteome, with 53% of the nuclear proteins in the original annotation having altered amino acid sequences in the new annotation, a remarkable ∼50 bp increase in average CDS length, a reduction of the average length of intergenic regions by close to 100 bp and the assignment of an additional 200,000 base pairs (or 2.4% of the genome), previously classified as intergenic or intronic sequences, to the proteome. This results in a genome that is denser than previously thought, with an overall increase in the coding fraction of the genome from 68% to 71%, more closely resembling *T. annulata* Ankara, which has a coding fraction of 72.9% (Supplementary Table S2, Additional File 1). In fact, *T. parva* has the densest genome out of the indicated genomes investigated, with one protein-coding gene every ∼2,100 bp (Supplementary Table S2, Additional File 1).

**Figure 1.**
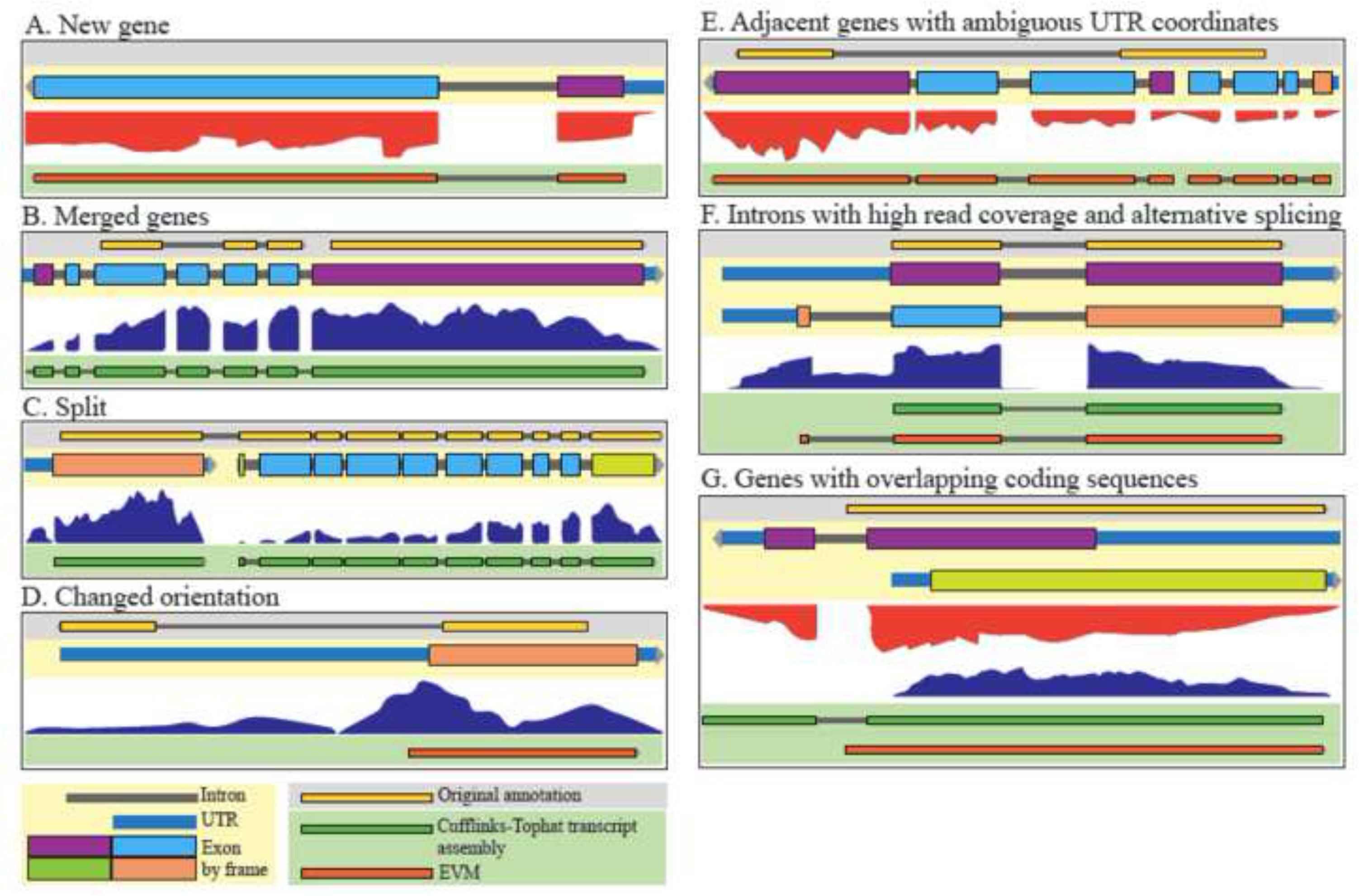
Manual gene model curation examples. Several tracks are shown: updated gene model (beige background), original (2005) gene annotation (grey background), RNAseq data (white background), transcript assembly (dark green, on green background), and EVM predictions (orange, on green background). (A) A new gene discovered on the basis of RNAseq data (TpMuguga_03g02005). (B) A case where two genes in the 2005 annotation merge in the new annotation on the basis of RNAseq read coverage (TpMuguga_04g02435). (C) A case where a gene in the 2005 annotation has been split into two genes in the new annotation (TpMuguga_04g02190 and TpMuguga_04g02185). (D) A case where a gene has been reversed in orientation on the basis of RNAseq data (TpMuguga_02g02095). (E) A case where overlapping genes led to ambiguity in UTR coordinates, and so the UTRs were not defined in this intergenic region (TpMuguga_01g00527 and TpMuguga_01g00528). (F) A case of a single gene where alternative splicing exists (as seen by significant read coverage in at least one intronic region), but there is one most prevalent isoform (TpMuguga_03g00622). (G) A case of two genes that overlap by coding sequences. Coding exons are colored by reading frame (TpMuguga_05g00017 and TpMuguga_05g00018).

Several lines of evidence suggest that this annotation represents a very significant improvement of the *T. parva* proteome relative to the original annotation. First, there was an increase in the proportion of proteins with at least one PFAM domain in the new proteome compared to the original proteome, implying that the new annotation captures functional elements that were previously missed (Figure 2a). Given the close evolutionary relationship and near complete synteny between *T. parva* and *T. annulata* [20], their respective proteomes are expected to be very similar. Indeed, a comparison of the two predicted proteomes results in 52 additional reciprocal best hits and protein length differences between orthologs in *T. parva* and *T. annulata* also decreased significantly (Figure 2b). It is likely that some of the most significant differences between the *T. parva* and *T. annulata* proteomes, in particular the 25% fewer protein-coding genes and much longer CDSs in the latter, represent annotation errors in the *T. annulata* genome that will be corrected upon revision with more recently accumulated evidence. The total number of non-canonical splice sites in the genome increased from 0.15% to 0.36% of all introns, but the sequence diversity of non-canonical splice sites decreased from eight non-canonical splice donor and acceptor site combinations to only a single splice site pair – GC/AG donor and acceptor dinucleotides, recognized by the U2-type spliceosome [21] (Figure 2c). The new annotation is also considerably more consistent with the RNAseq data, with a larger number of introns, a higher proportion of which is supported by at least one RNAseq read (Figure 2d). A total of 118 introns from the original genome annotation have been removed, due to contradicting RNAseq evidence.

**Figure 2.**
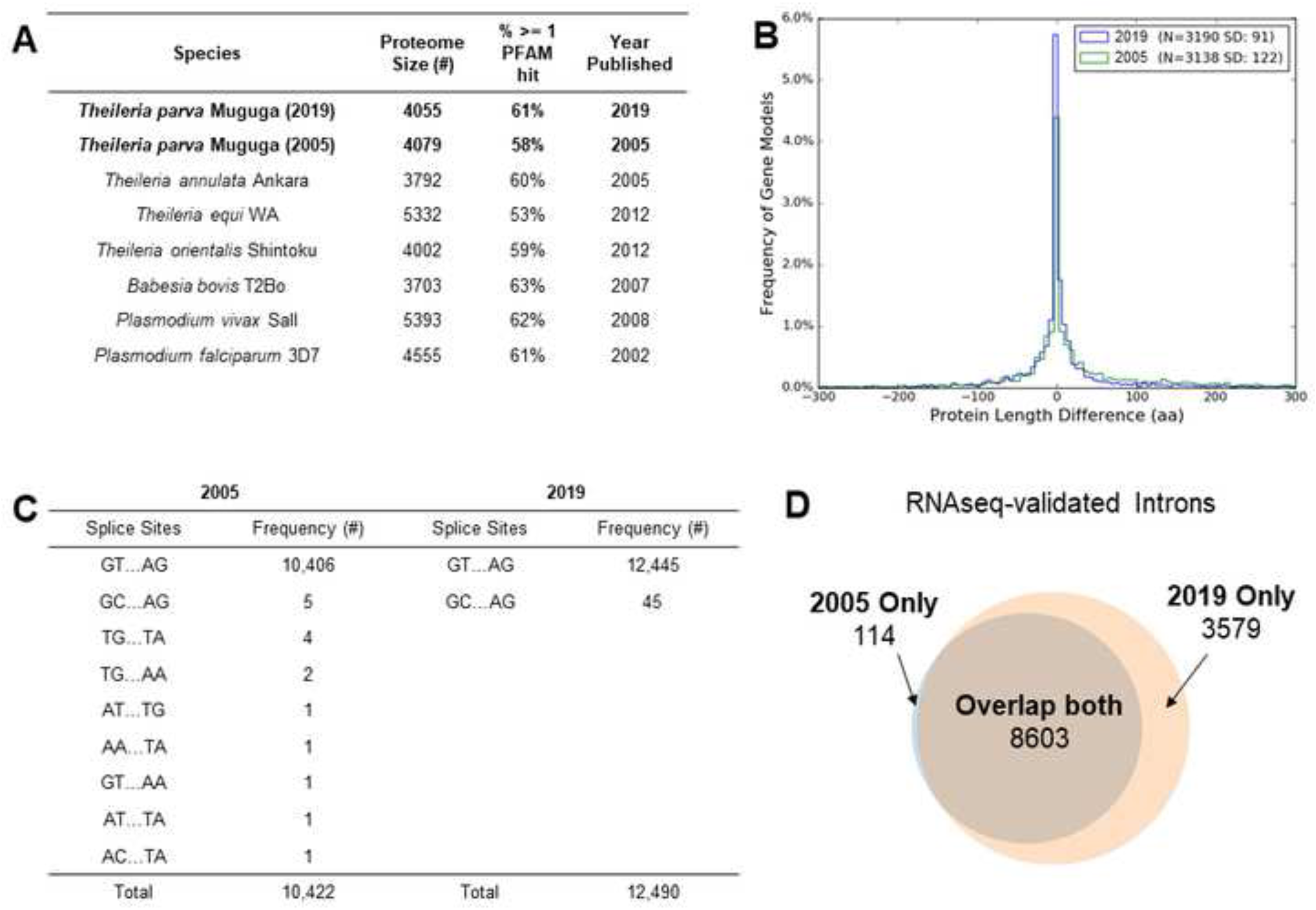
Comparative metrics of original and new *T. parva* annotations. (A) The percentage of proteins with at least one PFAM domain found by Hidden Markov Model searches of the predicted proteomes of the new *T. parva* Muguga annotation was 2% higher than those in the 2005 annotation, implying that the new annotation captures functional elements that were previously missed. (B) The new *T. parva* Muguga annotation has more reciprocal best-hit orthologs (N) with *T. annulata* Ankara than the 2005 *T. parva* Muguga annotation. The variation in protein length (SD) between *T. parva* and *T. annulata* ortholog pairs is greatly reduced in the new relative to the original *T. parva* annotation. Only nuclear genes were used for this analysis. The x-axis was limited to the range −300 to +300 for easy visual interpretation. (C) The number of canonical GT/AG intron splice sites increased and the number of non-canonical intron splice site combinations decreased in the new *T. parva* Muguga annotation compared to the 2005 annotation. (D) The number and proportion of introns validated by at least one RNAseq read increased in the new *T. parva* Muguga annotation compared to the 2005 annotation. These lines of evidence suggest that the new annotation is more accurate, and also considerably more consistent with the RNAseq data, as expected.

The tremendous power of RNAseq to inform on gene and isoform structure in this species revealed a significant amount of transcriptome diversity and complexity. First, the proportion of loci (defined here as a continuous genomic region encoding the length of a CDS, intervening introns, and flanking UTRs) that appear to overlap an adjacent locus increased from 2% to 29% in the new annotation. In many of these instances, read coverage, coding potential, and other evidence support the presence of adjacent genes with overlapping UTRs (Supplemental Figure S2a). In 130 cases, the overlap includes not only UTRs but also CDSs (Supplemental Figure S2b). Secondly, there are many instances of overlapping loci in which the respective CDSs are encoded in the same strand; in these cases, no UTRs were defined in the intervening intergenic region, since their exact boundaries could not be determined (Additional File 2). Finally, during manual curation, we observed many instances of potential alternative splicing, the clearest of which were the cases of well-supported introns where RNAseq coverage was nevertheless significantly higher than zero (Supplemental Figure S3; Figure 1f). In fact, we identified 872 introns, in 490 expressed genes (with average read coverage > 0), where the read coverage was at least equal to the mean read coverage for the coding sequences of the respective gene (Supplemental Figure S3b,c; Additional File 3), instances that are only possible to detect when read coverage varies considerably across the gene, which is not uncommon (e.g, Figure 1c,e). In these cases, only the most prevalent isoform was annotated (Figure 1f). Finally, despite its power, RNAseq evidence is not sufficient to resolve the structure of all loci; when the evidence did not clearly favor one gene model over another, the gene model in the original annotation was maintained by default. Interestingly, the vast majority of the genes appear to have only one or, sometimes, two most prevalent isoforms, as has been proposed for *Plasmodium* [22], although this was not defined quantitatively here. The median length of the annotated mRNA reported here is ∼1,500 bp, and the maximum length >15,000 bp (Supplementary Figure S4, Additional File 1).

### Most genes are transcribed during the schizont stage of the *Theileria parva* life-cycle, and antisense transcription is widespread

We sequenced cDNA generated from polyA-enriched total RNA collected from a *T. parva*-infected, schizont-transformed bovine cell line (see Methods section). A total of 8.3×10^7^ paired-end reads were obtained with an Illumina HiSeq 2000 platform, 70.04% of which mapped to the *T. parva* reference genome (Supplementary Table S1, Additional File 1). RNAseq provided a complete and quantitative view of transcription revealing that most of the genome of this parasite is transcribed during the schizont stage of its life cycle (Supplementary Figures S3, S5, Additional File 1). We found that 4011 of all 4054 (98%) predicted protein-coding parasite genes are transcribed at the schizont stage, and 12,172 of all 12,296 introns are supported by RNAseq reads (Figure 2d). We found evidence of expression for almost all of the known humoral and cellular immunity antigens (Supplementary Table S3, Additional File 1). In fact, Tp9, one of those antigens, is among the 15 most highly expressed genes in our dataset (Supplementary Table S4, Additional File 1). Interestingly, its ortholog in *T. annulata* has been hypothesized to contribute to schizont-induced host cell transformation [23].

As has recently been suggested from *in silico* analyses [18], transcription in *T. parva* occurs from diverse kinds of promoters, with many instances of adjacent loci overlapping on the same or opposite strands. In fact, of the 4,085 predicted protein-coding nuclear genes, only 74 had an estimated reads per kilobase of transcript per million reads (RPKM) of zero and an additional 154 had RPKM<1. Interestingly, of the 74 genes with an RPKM of zero, most are hypothetical, with no predicted functional annotation, and without any high-confidence orthologs (Supplementary Table S5, Additional File 1). Since tRNAs are not polyadenylated, they were not found in our RNAseq dataset (Materials and Methods). Annotated protein-coding genes lacking RNAseq evidence are mostly orthologs of *Plasmodium falciparum* apicoplast proteins with mid-blood stage expression [24, 25], *T. parva* repeat (*Tpr*) family proteins, or DUF529 domain-containing proteins (Supplementary Table S5, Additional File 1). These data are consistent with a study published in 2005, which used MPSS to estimate expression levels of *T. parva* genes in the schizont stage of the parasite [26], as well as a more recent study comparing gene expression between the schizont and the sporozoite/sporoblast stages [27]. The expression levels in the sense strand for each gene, as quantified by RPKM, when log-transformed, followed a unimodal distribution similar to a normal distribution (Figure 3a).

**Figure 3.**
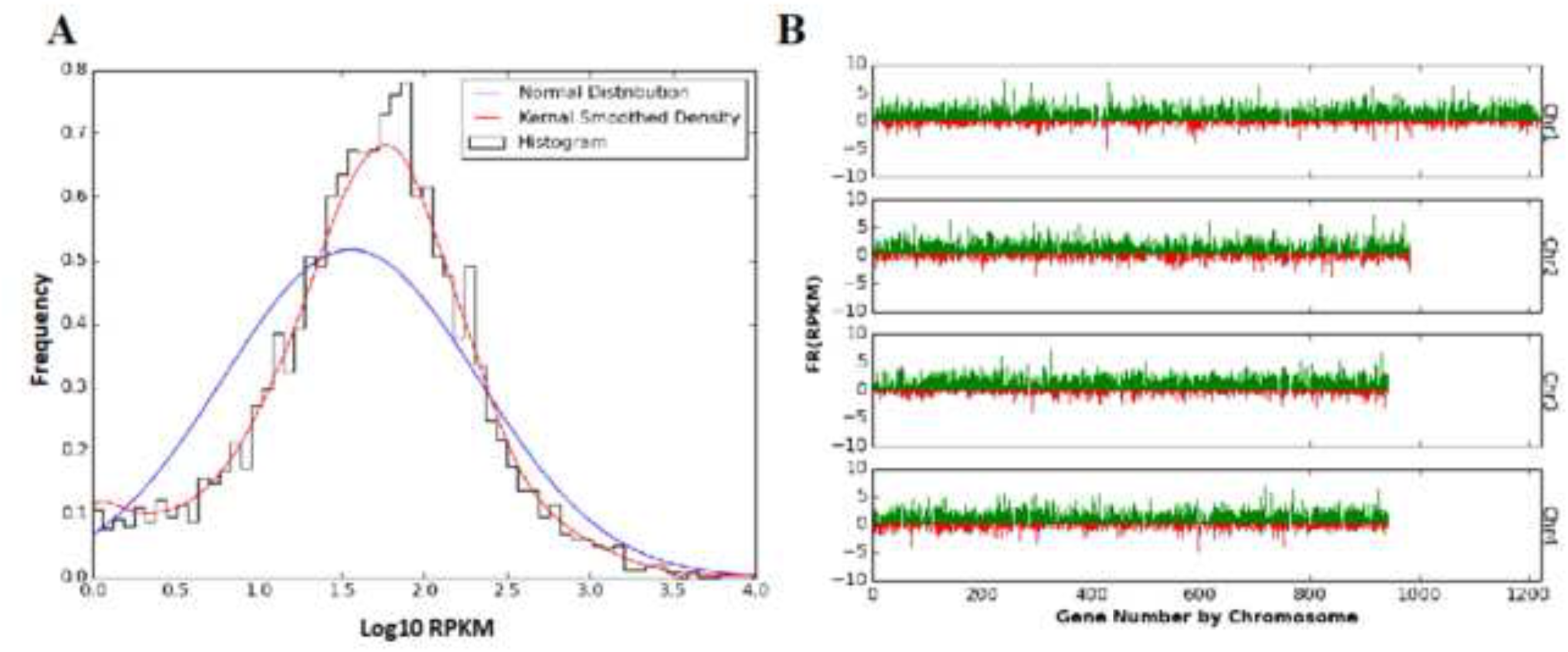
Distribution of RNAseq RPKM values for *T. parva* Muguga genes. (A) A histogram of sense RPKM values after logarithmic transformation of the data. Frequencies on the y-axis correspond to probability density. The blue line shows a normal distribution around the same median, while the red line shows a more reliable fixed-width, Gaussian, kernel-smoothed estimate of the probability density. (B) The sense (green) and antisense (red) reads per kilobase transcript per million reads (RPKM) after fourth-root transformation of the data. Genes are sorted by position on the chromosome for all four nuclear chromosomes of *T. parva* Muguga.

### *T. parva* multi-gene families show variable expression levels

Large gene families are known to play a role in the pathogenesis of protozoan infections, perhaps the most well-known being the *var* gene family in *P. falciparum*. These genes encode proteins that are essential for the sequestration of infected red blood cells, a critical biological feature determining severe malaria pathology of *P. falciparum* [28]. Using the OrthoMCL algorithm as described previously [19], we clustered paralogs in this genome, identifying changes in the size of several of the largest *T. parva* gene families (Supplementary Table S6, Additional File 1), and finding variable patterns in their levels of expression (Supplementary Figure S6, Additional File 1). The roles of most of these gene families are not known. For example, the *Tpr* (*T. parva* repeat) gene family has been suggested to be rapidly evolving and expressed as protein in the piroplasm stages [19]. This is consistent with our findings, which show *Tpr* genes not to be highly expressed in the schizont (Supplementary Figure S6, Additional File 1) or the sporoblast (Supplementary Figure S7, Additional File 1) stages [27, 29]. Interestingly, in that same dataset, we find a significant up-regulation of subtelomeric variable secreted protein gene (SVSP) family genes in the sporozoite stages relative to both the sporoblast and schizont stages, suggesting that they may be important for invasion or the establishment of infection in the vertebrate host (Supplementary Figure S7, Additional File 1) [30].

This new *T. parva* genome annotation not only improved our resolution of the gene models of multi-gene family members and other transformation factors (Supplementary Figure S8, Additional File 1) [31], but also uncovered 128 genes that were not present in the original annotation.

### A mechanism of core N-glycosylation is now predicted in *T. parva*

Among the 128 newly identified genes, one was annotated as a potential Alg14 ortholog, an important part of a glycosyltransferase complex in many organisms that add a N-acetylglucosamine (GlcNAc) to the N-glycan precursor. N-glycosylation is an important type of protein post-translation modification, during which a sugar is linked to the nitrogen of specific amino acid residues, a process that occurs in the membrane of the endoplasmic reticulum and is critical for both the structure and function of many eukaryotic proteins. N-glycosylation is a ubiquitous protein modification process, but the glycans being transferred differ among the domains of life [32]. However, in apicomplexan parasites that infect red blood cells, there appears to be a selection against long N-glycan chains [33]. *Theileria* parasites were previously believed to not add N-acetylglucosamine to their glycan precursors, since sequence similarity searches did not identify the necessary enzymes. While the study by Samuelson and Robbins [33] did not discover any Alg enzymes, we find that *T. parva* has Alg7 (*Tp*Alg7; TpMuguga_01g00118), Alg13 (*Tp*Alg13; TpMuguga_02g00515), and Alg14 (*Tp*Alg14; TpMuguga_01g02045) homologs, which show differential mRNA-level expression between the sporozoite and schizont life cycle stages (Supplementary Figure S9, Additional File 1). In fact, the structure of each of these *Theileria* proteins can be predicted *ab initio* with high confidence (Supplementary Table S7, Additional File 1) and have predicted secondary structural characteristics very similar to their homologs in *Saccharomyces cerevisiae* (Figure 4a). However, the structure of the *Tp*Alg7-encoding locus was altered as a result of the re-annotation effort and *Tp*Alg14 is the product of a newly identified gene, which might have prevented the original identification of the pathway. Therefore, *Theileria* parasites likely have a minimal N-glycosylation system. Interestingly, we can find Alg14 orthologs by blastp search in *T. orientalis* (TOT_010000184), *T. equi* (BEWA_032670), but not in *T. annulata*. Using the adjacent gene, EngB, as a marker, a look at the *T. annulata* genomic region that is syntenic to *Tp*Alg14 revealed that *T. annulata* has a hypothetical gene annotated on the opposite strand (Figure 4b), which could be an incorrect annotation. A tblastn search of the *T. annulata* genome using *Tp*Alg14 led to the discovery of a nucleotide sequence which translated results in an alignment with E-value of 7×10^−15^ and 70% identity over the length of the protein, suggesting the existence of an *T. annulata* Alg14 ortholog (*Ta*Alg14). In fact, the gene model that was at the *Tp*Alg14 locus in the original annotation, TP01_0196, was likely a result of an incorrect annotation transfer from *T. annulata* (or vice-versa), since TP01_0196 shared 52% identity with the gene annotated on the opposite strand at the putative *Ta*Alg14 locus (E-value 4×10^−131^). Since previous studies have used *T. annulata* as a model *Theileria* parasite, this could be the reason that N-glycosylation was not discovered in this parasite genus.

**Figure 4.**
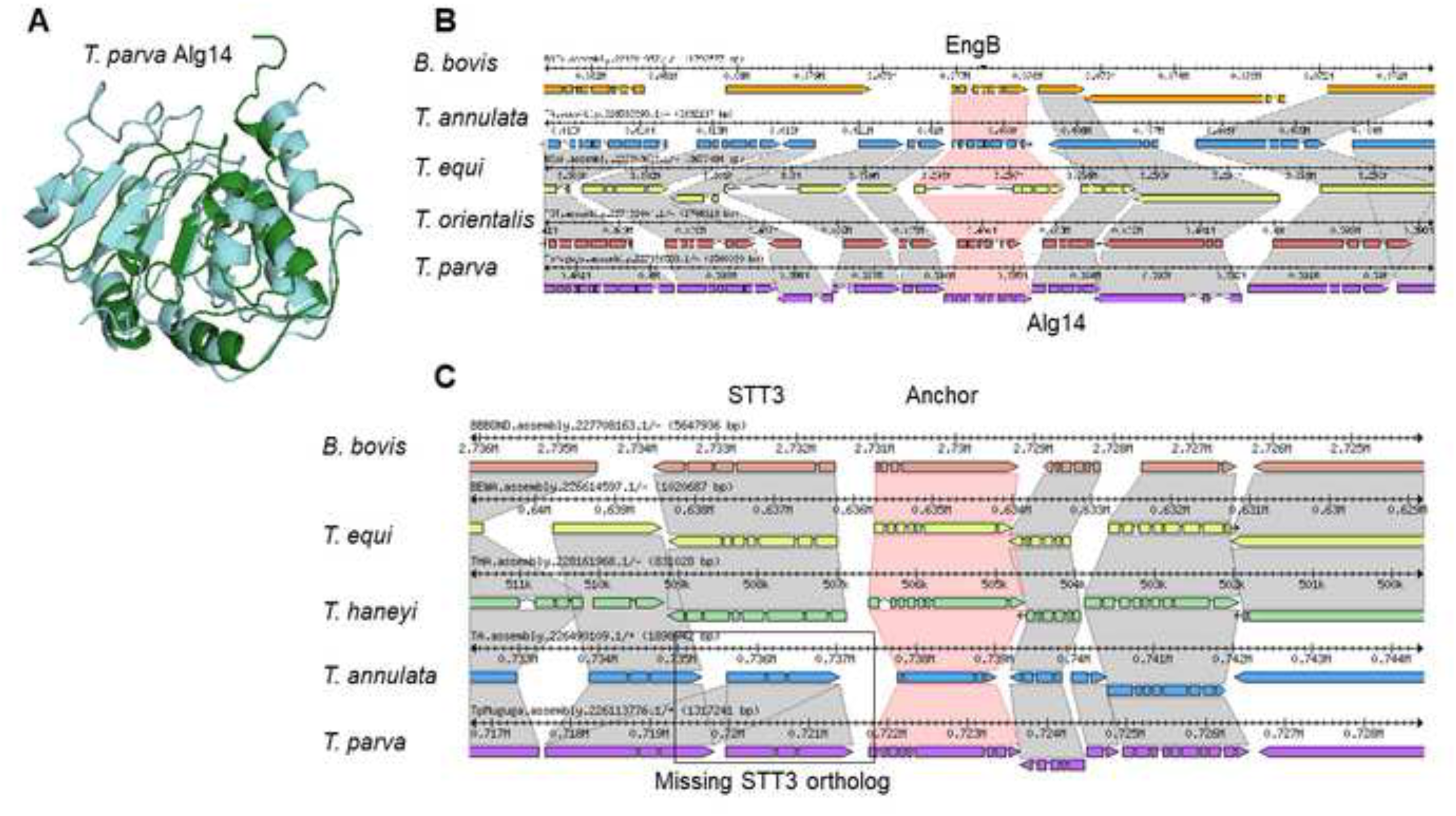
The uncovered *Theileria parva* Alg14 shows a similar predicted structure to the empirically determined *Saccharomyces cerevisiae* Alg14 protein structure, and is syntenic in multiple piroplasms. (A) A Phyre2 prediction of *T. parva* Alg14 (*Tp*Alg14; green; TpMuguga_01g02045) and the Protein Database (http://www.rcsb.org/) [77] nuclear magnetic resonance structure of *Saccharomyces cerevisiae* Alg14 (*Sc*Alg14; teal; PDB 2JZC) were aligned in MacPyMol (https://pymol.org/2/) [78]. (B) Shown are the syntenic regions around Alg14 orthologs (synteny in grey), using the adjacent gene EngB as an anchor (synteny in red) in the Sybil software package [79]. (C) Shown are the syntenic regions around STT3 orthologs (synteny in grey), using a *B. bovis* STT3-adjacent gene (<u>BBOV_II000210</u>) as an anchor (synteny in red) in the Sybil software package.

While the presence of N-glycans in *Plasmodium* parasite proteins was initially controversial [34], more recent work provided evidence of short N-glycans on the exterior of *P. falciparum* schizonts and trophozoites [35]. As a key difference, *Plasmodium* parasites have a clear ortholog of the oligosaccharyl transferase STT3 (EC 2.4.99.18, PF3D7_1116600 in *P. falciparum* 3D7), which catalyzes the transfer of GlcNAc and GlcNAc_2_ to asparagine residues in nascent proteins, and recent work has identified several other proteins in this protein complex in *Plasmodium* genomes [34]. No such ortholog was found in *T. parva* Muguga or *T. annulata* Ankara by blastp or tblastn searches with the *Plasmodium* protein. Since there are STT3 orthologs in *T. equi* and *T. haneyi* (Figure 4c), as well as *Cytauxoon felis*, it appears that the absence of STT3 in *T. parva* and *T. annulata* represents evolutionary loss of STT3 orthologs in this lineage. This means that while lipid precursor N-glycosylation does likely occur at the ER in these two species, the canonical mechanism of N-glycan precursor transfer to proteins is apparently absent.

## Discussion

The re-annotation of the *T. parva* genome has resulted in significant improvement to the accuracy of gene models, showing that this genome is even more gene-dense than previously thought, with the addition of 2.4% of the genome to CDSs as well as the discovery of additional overlapping genes. Multi-gene families appear to have played a prominent role in the evolution of the lineage leading to *T. parva* and *T. annulata* [36], implying a role for these genes in host-pathogen interactions. These genes have diversified and/or expanded in copy number, possibly as an adaptation to a particular niche, since the high density of the genome is strongly suggestive of selection against non-functional DNA. We now have a clearer picture of the structure, copy number, and relative expression level of these genes. In addition, a recently generated sporozoite and sporoblast datasets opens up new opportunities to study differential gene expression throughout other stages of the parasite life cycle [27].

The model of transcription that emerges from these recent studies is one of ubiquitous transcription of most genes in the schizont stage, but with a wide range of expression levels [18, 26, 27], suggesting that there are likely important *cis* regulatory motifs that control the level of expression or mRNA stability [18, 37]. Transcription can also arise from potential bidirectional and cryptic promoters with highly prevalent antisense transcription. It remains to be determined if sense and anti-sense transcripts are generated in the same or different cells in culture, an issue that may be addressed with single-cell RNAseq. Due to the short-read nature of our sequencing platform, we were only able to accurately annotate the most prevalent isoform of each gene. The sequencing of full-length transcripts, for example with Pacific Biosciences sequencing technology, would provide a more comprehensive description of the *T. parva* transcriptome, including alternatively spliced variants and the boundaries of overlapping transcripts.

In yeast and humans, antisense transcription, defined by the existence of non-coding RNA encoded on the DNA strand opposite to, and overlapping with, that encoding the mRNA, is rare compared to sense transcription [38]. In *T. parva*, however, antisense transcription is highly prevalent throughout the genome (Figure 3b), as has been found in *P. falciparum*, where antisense transcription is synthesized largely by RNA polymerase II [39, 40] and can alter the expression of multigene family members by regulating the packaging of these loci into chromatin [41]. Most of the antisense transcripts seem to completely overlap with their sense counterparts, although the functional relevance of this observation has yet to be determined.

The discovery of evidence that N-glycosylation may occur in *Theileria* parasites could open up novel treatment options against *Theileria* infections. N-glycosylation is thought to be important for *Toxoplasma gondii* invasion, growth, and motility [42–44]. While the results are somewhat confounded by a lack of inhibitor specificity, treatment with the N-glycosylation inhibitor tunicamycin results in parasites with abnormal endoplasmic reticulum, malformed nuclei, and impaired secretory organelles [45]. While once controversial due to differences in analytical methods, parasite life-cycle stages, and host contamination, *P. falciparum* is now thought to have N-glycosylated proteins, although this is not as frequent a mechanism of protein modification as glycosylphosphatidylinositol [46]. This work has been supported by bioinformatic analyses, finding that *P. falciparum* contains glycosyltransferases (albeit few) [47]. Early work using N-glycosylation inhibitors has shown strong *in vitro* growth inhibition of *Plasmodium* asexual blood stages [48–52], but the function of N-glycosylation of apicomplexan parasite proteins is a topic that requires further study. Importantly, the lack of an STT3 ortholog in *T. parva*, if true, would suggest that protein-targeted N-glycosylation does not occur in this parasite (as does in *Plasmodium*), and may only occur on the ER and potentially the surface of the parasite. Even though cytoplasmic N-glycosyltransferases have been found in bacteria, they have not been found in eukaryotes, and their presence in *T. parva* seems unlikely. The absence of a N-glycan protein transfer system is largely supported by genome-wide searches for the enrichment of N-glycan acceptor sites in *T. annulata* [53]. While N-glycosylation is often touted as an ‘essential’ protein modification in eukaryotes [54], the absence of an STT3 ortholog in some *Theileria* species suggests that this process may be critical as a lipid, rather than protein, modification. This does not diminish the potential relevance of N-glycosylation in these parasites. Regardless of whether these short N-glycans provoke host immune responses or play a homeostatic role in parasite protein folding, they could be important therapeutic targets. Finally, given the possibility that glycans encode immunological ‘self’, ‘non-self’ or ‘damage’ identities [55], it is tempting to speculate that the absence of proteinaceous N-glycans in *Theileria* species could represent an evolutionary adaptation to immune evasion in a parasite lineage that resides free in the host cytoplasmic environment.

## Conclusions

This study emphasizes the critical interplay between genome annotations and our knowledge of pathogen biology. The significant improvement of the *T. parva* Muguga reference genome gene annotation will facilitate numerous studies of this parasite, and has already given better resolution to genome-wide patterns of gene transcription, including antisense transcription and transcription of multi-gene families. The better the resolution at which we understand gene structure and expression, the more accurately we can characterize and study gene function, novel druggable pathways suitable for interventions and, ultimately, the biology of the pathogen in its different host organisms. For example, the discovery of N-glycosylation precursors in some *Theileria* parasites in the absence of a protein transfer system opens up new questions about the role of lipid N-glycosylation precursors in eukaryote biology as well as the potential evolutionary reasons why protein N-glycosylation would be lost in this apicomplexan lineage.

## Methods

### 1) RNA sequencing and genome annotation

An RNA sample was obtained from the reference *T. parva* isolate (Muguga) from the haploid schizont stage of the parasite life cycle, which proliferates in host lymphocytes. The extraction method included complement lysis of schizont-infected host lymphocytes, DNase digestion of contaminating host DNA and differential centrifugation to enrich for schizonts [26, 56]. PolyA-enriched RNA was sequenced using Illumina sequencing technology, to produce strand-specific RNAseq data. RNAseq reads were aligned with TopHat and RPKM values calculated using HTseq [57].

### 2) Genome re-annotation

For the re-annotation of the *Theileria parva* genome, a number of evidence tracks were generated and loaded into the genome browser JBrowse [58] for manual curation using the WebApollo plugin [59]. RNAseq reads were aligned to the genome with TopHat [60], a splice-aware alignment tool (Supplementary Table S1, Additional File 1). These alignments were used to generate strand-specific read alignment coverage glyphs and XY plots for visualization in WebApollo. TopHat alignment also yields a file of all reported splice junctions using segmented mapping and coverage information, which is useful for curating intron splice sites. RNAseq reads were also assembled into transcripts using CuffLinks [61] and mapped to the genome with TopHat. We also generated two genome-dependent Trinity/PASA [62] transcriptome assemblies (one reference annotation-dependent and one independent of the reference annotation), as well as one completely *de novo* Trinity transcriptome assembly. A variety of other proteome data were aligned to the genome with AAT [63] and used as evidence tracks, including previously generated *Theileria annulata* mass spectrometry data [64], and all non-*Theileria* apicomplexan proteins from NCBI’s RefSeq.

In order to assess gene prediction accuracy before the manual curation phase, a set of 342 high-confidence *T. parva* gene models were selected from the current reference annotation on the basis of two criteria: (1) RNAseq reads must cover each exon in the gene, (2) Trinity *de novo* assembled transcripts and read coverage must be concordant with the presence or absence of any introns in the gene model. Out of these 342 genes, 50 were randomly selected as a validation set and the remaining 292 were used as a training gene set for gene prediction software. The exon distribution of the validation set closely resembles that of the training set (Supplementary Table S8, Additional File 1).

Multiple gene prediction software tools were used and then assessed by the accuracy with which they predict the validation set using an in-house script. These included: *i*) Augustus [65], using RNAseq reads, the *T. parva* training gene set, or no evidence; *ii*) Semi-HMM-based Nucleic Acid Parser (SNAP) [66] and Glimmer [67] were trained with the *T. parva* training set; *iii*) Fgenesh [68] used a pre-existing training set of *Plasmodium* genes from its website; *iv*) the *ab initio* predictor GeneMark-ES [69]. Finally, gene models were selected with the consensus predictor Evidence Modeler (EVM) [70], using 57, differently-weighted combinations of the other evidence, while maximizing prediction accuracy (Supplementary Figure S8, Additional File 1). Based on their performance in comparison with the validation set, only the top four EVM predictions were loaded as evidence tracks for use in manual curation (Supplementary Figure S9, Additional File 1). tRNA and rRNA predictions were generated using tRNAscan-SE [71] and RNAmmer [72] and loaded as evidence tracks, along with the original *T. parva* Muguga annotation (Supplementary Figure S10, Additional File 1). A genome-wide, double-pass, manual curation of all gene models was completed, weighing the RNAseq evidence over the evidence from alignments with homologs from other species and the gene prediction programs. The annotation assignments were allocated in 50 kb segments, with different annotators doing adjacent segments, as well as altering the annotator for the first and second pass in order to reduce annotator bias.

Functional annotation of the *T. parva* proteome consisted of HMM3 searches of the complete proteome against our custom HMM collection that includes TIGRFams [73], Pfams [74], as well as custom-built HMMs [75] and RAPSearch2 searches against UniRef100 (with a cutoff of 1×10^−10^). In addition, a TMHMM search which was used to assign “putative integral membrane protein” to proteins with 3 of more helical spans (assuming there were no other hits to the previous searches). These searches were synthesized using Attributor (https://github.com/jorvis/Attributor) to generate the final annotation based on the different evidence sources to assign gene product names, EC numbers, GO terms and gene symbols to genes, conservatively where possible.

### 3) Multi-gene family clustering

Genes were clustered with OrthoMCL, using an inflation value of 4 and a BLAST p-value cutoff of 10^−5^, as previously done [19]. All individual conserved domain searches were done using NCBI’s Conserved Domain Database version 3.11 [76] with 45,746 PSSMs, with an E-value threshold of 0.01 and a composition based statistics adjustment. HMM searches of the entire PFAM database were done using default settings.

## List of abbreviations

ECF: East Coast fever
UTR: Untranslated Regions
CDS: Coding Sequence
RPKM: reads per kilobase million
Tpr: *T. parva* repeat family
SVSP: subtelomeric variable secreted protein
SNAP: Semi-HMM-based Nucleic Acid Parser

## Declarations

### Ethics approval and consent to participate

Not applicable.

### Consent for publication

Not applicable.

### Availability of data and materials

The *T. parva* Muguga re-annotation is publicly available in GenBank and can be visualized at the following online link (http://jbrowse.igs.umaryland.edu/t_parva/), as well as under the NCBI BioProject PRJNA16138.

### Authors’ contributions

RP isolated *T. parva* schizont RNA for RNAseq using differential centrifugation and standard kits as described. JO, OOI, and PK built and maintained the JBrowse instance for manual curation using the WebApollo plugin, as well as the functional annotation pipeline. KT, JO, HTG, OOI, PK, and SBAI generated alignment tracks to assist the annotation. KT, HTG, OOI, NCP, SBAI, JCS completed manual re-annotation of the genome. KT performed all other analyses. JCS, RPB, CD, VMN, and LF conceived the study design. KT and JCS wrote the manuscript. All authors critically reviewed and approved the manuscript.

## Acknowledgements

We would like to thank Donald P. Knowles for his kind and helpful feedback and support for this project.

