## Additional Data 1 for "Re-annotation of the *Theileria parva* genome refines 53% of the proteome and uncovers essential components of N-glycosylation, a conserved pathway in many organisms"

### Supplemental Information

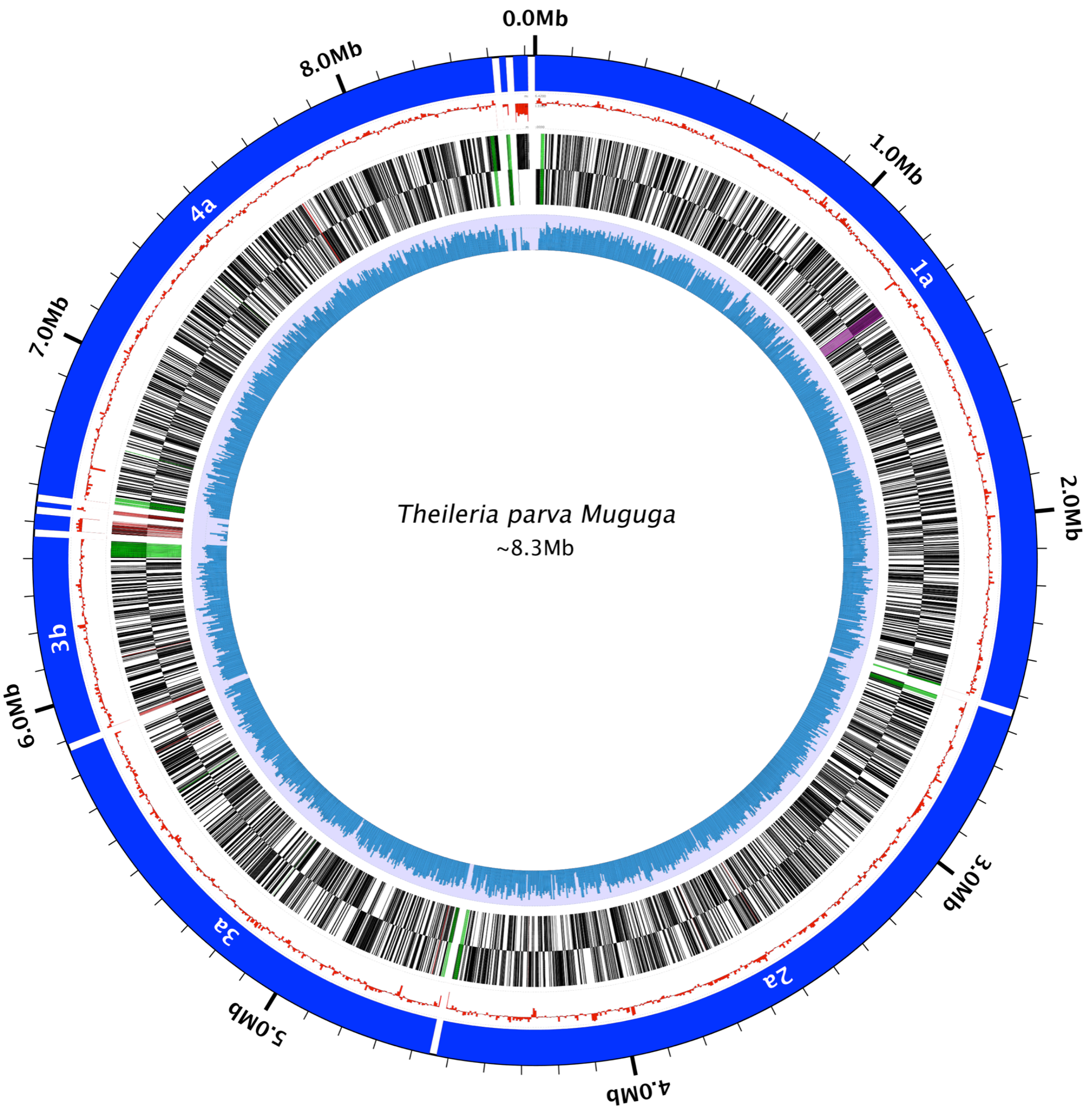


**Supplemental Figure S1. Architecture of the *Theileria parva* Muguga genome and associated new structural annotation and RNAseq expression data.**

Data are shown for each of the 4 *T. parva* nuclear and one mitochondrial chromosomes. The five concentric rings, from outer- to innermost, represent (i) the boundaries of each contig (blue), with the last and most AT-rich contig representing the mitochondrion, (ii) the deviation from the average GC percentage (red), (iii) the genes on the forward strand (green = SVSP family genes, red = Tpr genes, purple = TpHN genes), (iv) the genes on the reverse strand (same colors as forward), and (v) RNAseq coverage depth (blue, on light purple background). The figure was generated using the Circleator software. The apicoplast genome was not re-annotated since only three of its 70 currently annotated genes had representative RNAseq coverage, and their existing annotation was consistent with the RNAseq data.

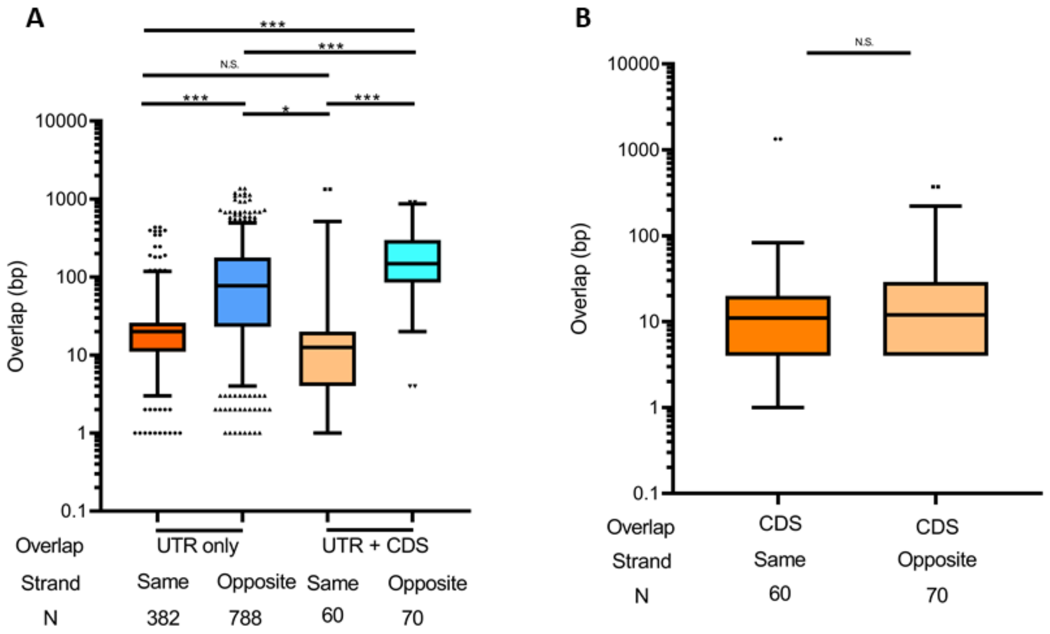


**Supplemental Figure S2. Updated gene annotation efforts in the *Theileria parva* genome reveal the existence of many genes that overlap adjacent genes at either UTR or CDS sequences.** (A) Boxplots of the distribution of the total overlap length by genes that overlap by only the untranslated regions (UTR), or where the overlap includes a protein coding sequence (CDS) on either the same or opposite strands. (B) Shown are boxplots for the distribution of CDS length overlap in adjacent genes. Distributions were compared with a one-way ANOVA using alpha = 0.05. Significance is shown as follows: * $0.05>p\geq0.01$; ** $0.01>p\geq0.001$; *** $p<0.001$. Bar plot lines represent the mean and 5-95% percentiles from the mean.

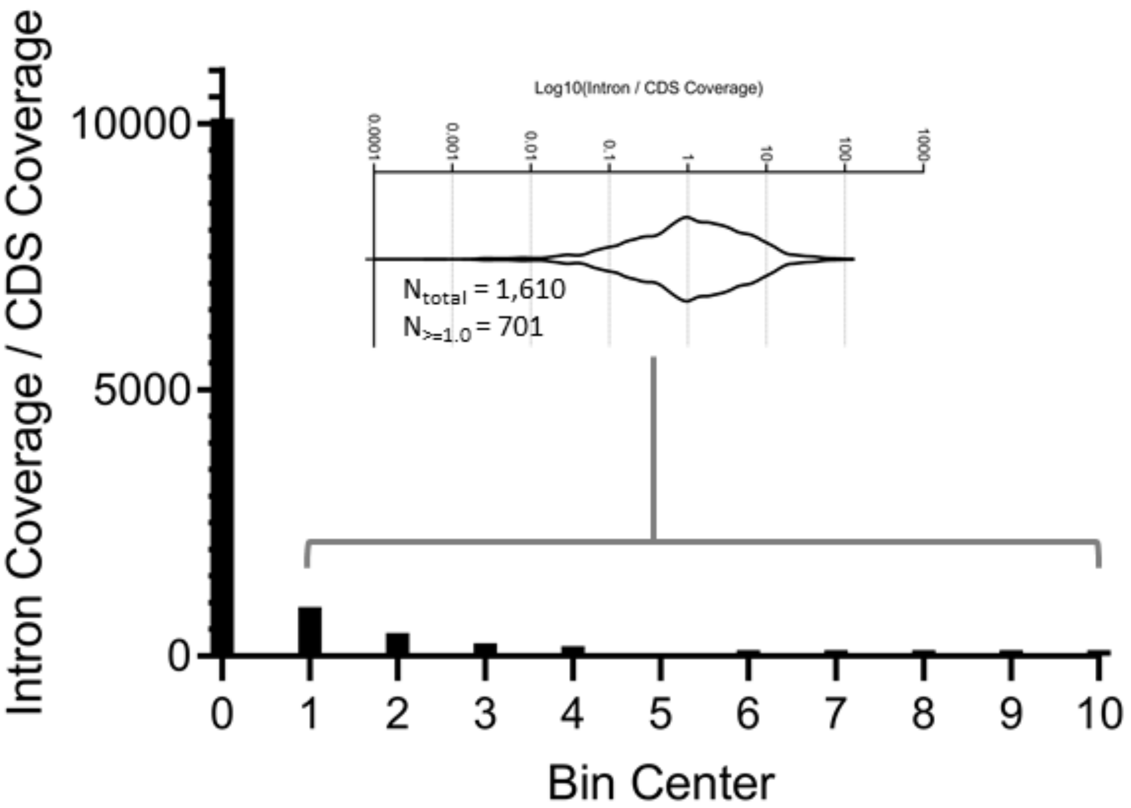


**Supplemental Figure S3. RNA-seq reads that map to introns in the *Theileria parva* Muguga genome support the existence of genes where a subset of introns are not spliced.** Read coverage per intron and protein coding sequence (CDSs) was calculated using HT-seq and their ratio plotted as a frequency histogram. While most introns had a coverage of zero (hence an intron_coverage/CDS_coverage ratio of zero), 1,610 introns had some read coverage, 701 of which had an intron_coverage/CDS_coverage ratio above or equal to 1.0. These 1,610 introns correspond to 744 genes.

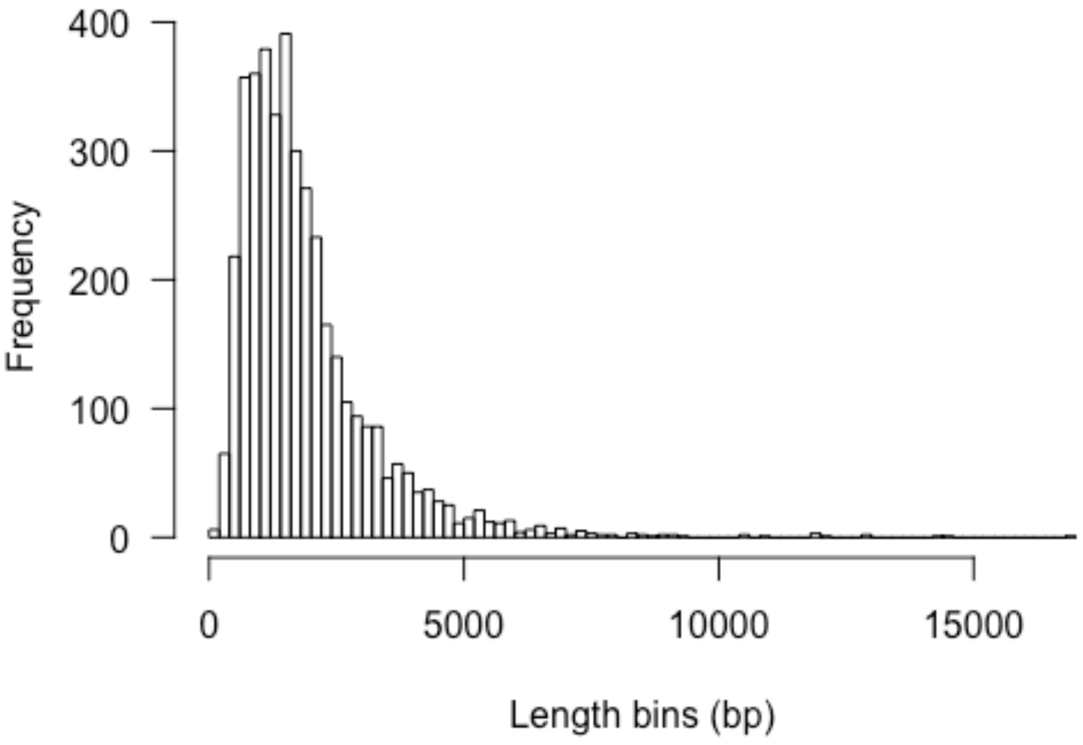


**Supplemental Figure S4. Histogram of mRNA length across all annotated transcripts in the current *T. parva* Muguga annotation.**

All mRNAs were used to make this graph, including those for which no untranslated region was annotated.

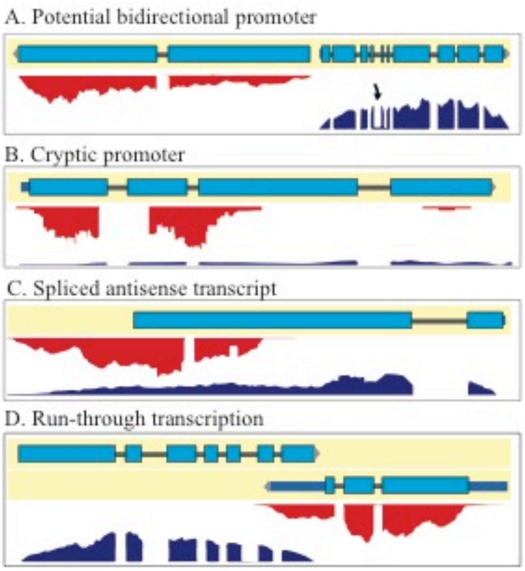


**Supplementary Figure S5. Types of transcription initiated in *Theileria parva*.**

Transcription in *T. parva* Muguga results from potential bidirectional (A) and cryptic promoters (B). Antisense transcription can be spliced (C), and may result in run-through transcription overlapping another gene (D). The model of transcription that emerges from these data is one of ubiquitous sense transcription of most genes in the schizont stage, but with a wide range of expression levels. Transcription can arise from potential bidirectional and cryptic promoters with highly prevalent antisense transcription. Blue boxes indicate exons, red glyphs indicate reverse strand-aligned RNA sequencing reads, and blue glyphs indicate forward strand-aligned RNA sequencing reads.

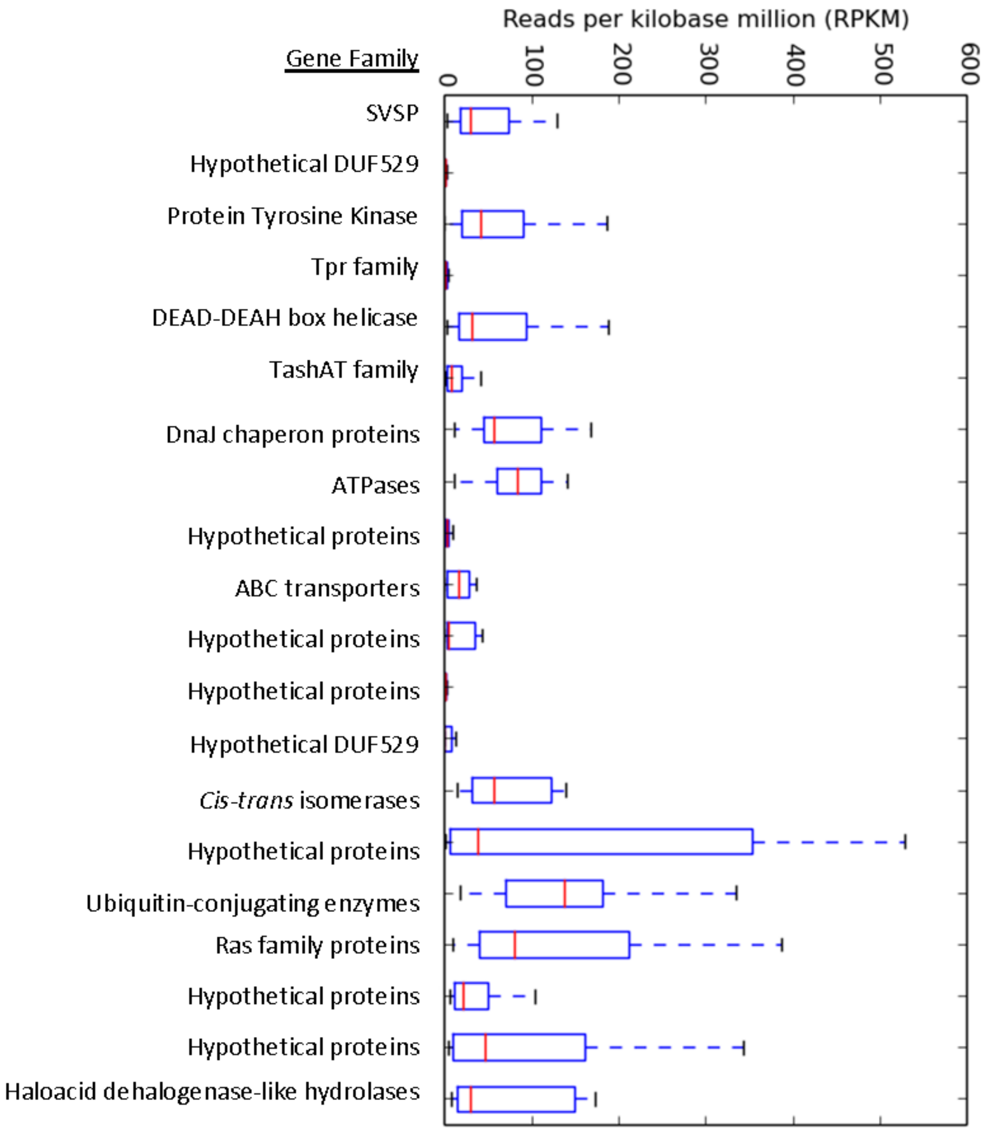


**Supplemental Figure S6. The RPKM distribution of the top 20 gene families in *T. parva* Muguga schizont RNAseq data.**

Gene families are sorted in order of decreasing size (top to bottom) on the left y-axis.

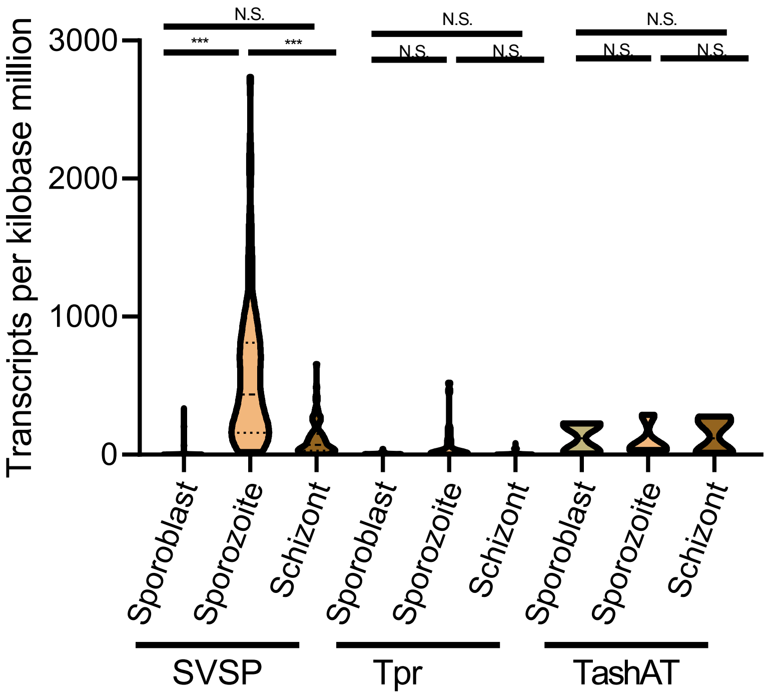


**Supplemental Figure S7. The expression distribution of three selected gene families in *T. parva* Muguga (SVSP = subtelomeric variable secreted protein gene family; Tpr = *T. parva* repeat gene family; TashAT = *Theileria annulata* schizont AT hook gene family).**

The data used in this plot was taken from a recent paper looking at gene expression in three life-cycle stages of *T. parva* (1). Within-family comparisons were tested using an ordinary one-way ANOVA (N.S. = not significant; *** = $p<0.001$).

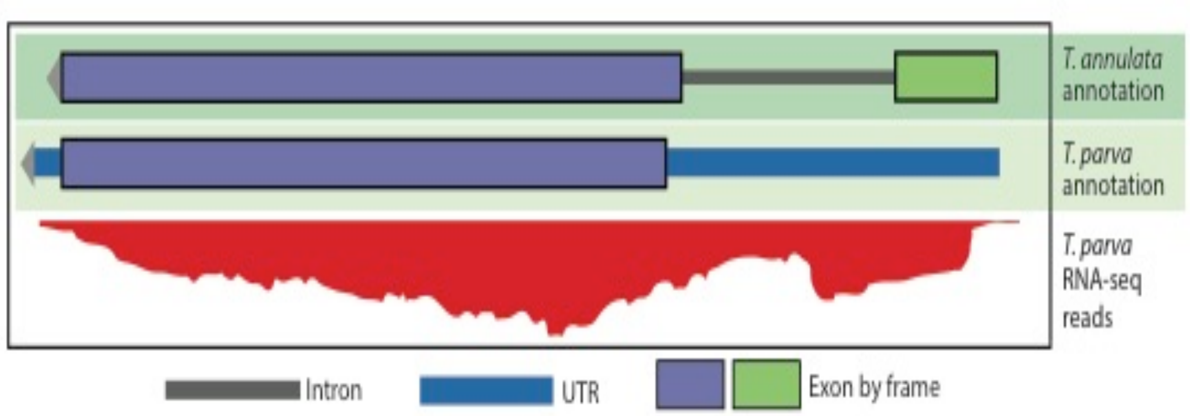


```
TA18945             MIRNFLNFLW NNTSLRFLII NTIIFAMDKV RCAHLLLKHT GSRNPVNRNT GMAVTRTKEE AVSEMKGYLE MLRKSDNLDQ EFRRLATAKS ECSSARKGGD
BEWA_033600         ---------- ---------- ---MSSSEKI RCAHILLKHT ESRNPVNRRT NQKVFRTKDE ANREIADILV RLKEAVNLGH EFKRIATDIS ECSSAKNGGD
TOT_010000107       ---------- ---------- ------MDKV RCAHLLLKHR DSRNPVNRNT GERVTRTKEE AVAELRSLLE ALRKSDRPDH EFRRLATAKS ECSSAKNGGD
TpMuguga_01g00114   ---------- ---------- ------MDKV RCAHLLLKHT GSRNPVNRNT GMAVTRTKEE AVNEMKGYLE MLRKSENLDQ EFRRLATAKS ECSSARKGGD
                                                  :*: ****:****   *******.*  . * ***:* *  *: . *   *::: . .: **:*:**  * *****::***

TA18945             LGFFDRNTMQ KPFTEASFKL EVNEISDLVE TDSGVHLIYR IA*
BEWA_033600         LGYFDRFTMQ KPFTEAAFRL KVDEISDIVD TDSGIHIIYR IA*
TOT_010000107       LGFFDRNTMQ KPFTEASFKL RVNEISDLVE TDSGVHLIYR IS*
TpMuguga_01g00114   LGFFDRNTMQ KPFTEASFNL KVNEISDLVE TDSGIHLIYR IA*
                    **:*** *** ******:*.* .*:****:*: ****:*:*** *:
```

**Supplementary Figure S8.** ***Theileria*** **PIN1 is an example of a parasite-secreted protein that plays a role in host transformation.** Shown is an alignment of the recently described *T. annulata* gene (2), *TaPIN1* (TA18945), with orthologs in *T. equi*, *T. orientalis*, and *T. parva*. *TaPIN1* has an extended N-terminal sequence that contains a signal peptide, as such it is the only putative protein in this ortholog cluster with a predicted secretion signal. The others are either predicted to localize to the mitochondria (TpMuguga_01g00114, TOT_010000107), or a localization prediction was not possible with TargetP default settings (BEWA_033600). However, the RNAseq evidence for the TpMuguga_01g00114 locus suggests that alternative splicing may occur that includes an N-terminal exon, which may contain a secretion signal that could alter the localization of TpMuguga_01g00114. This splice form of the gene is not prevalent in the RNAseq sample analyzed, but it is possible that further investigation with long-read sequencing technology will reveal more about the structure of this gene.

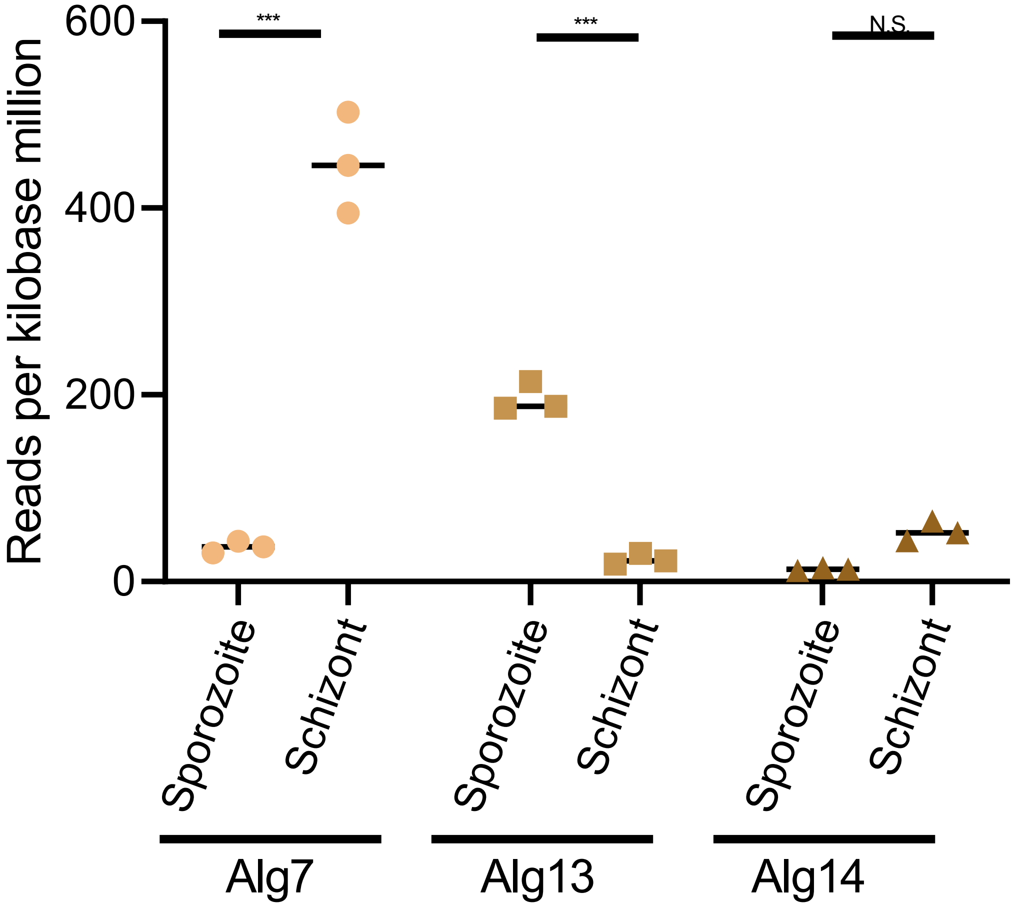


**Supplemental Figure S9. The expression N-glycosylation pathway components in the sporozoite and schizont life cycle stages of *T. parva* Muguga.**

The data used in this analysis was taken from a recent paper looking at gene expression in three life-cycle stages of *T. parva* (1). Between-stage comparisons were tested using a two-way ANOVA (N.S. = not significant; *** = $p<0.001$).

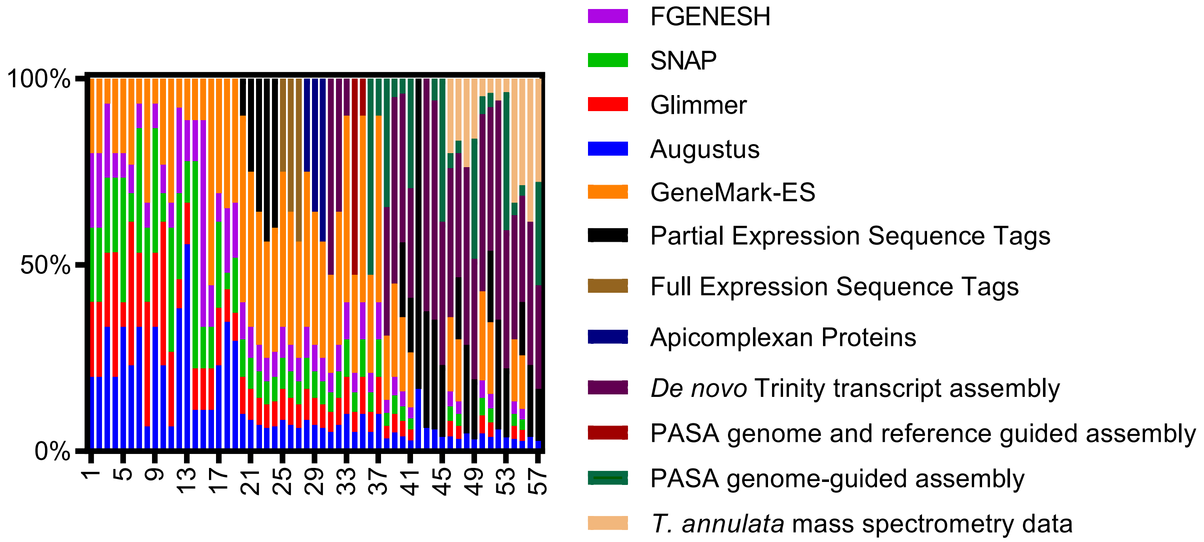


**Supplemental Figure S10. A representation of the relative weights of each evidence in each EVM prediction tested.**

Augustus: the *ab initio* predictor AUGUSTUS (3); TA-MS: *Theileria annulata* mass spectrometry data (4); PASA genome-guided assembly: a gene prediction by the Program to Assemble Spliced Alignments using the reference annotation to guide the assembly (5); PASA genome-guided, reference-guided assembly: a gene prediction using PASA with the assembly guided by both the reference genome and the reference annotation; *de novo* Trinity transcript assembly: an RNAseq transcript assembly using the RNAseq data generated in the present study, using the Trinity program (6); Protein alignments: alignments of all non-*T. parva* apicomplexan proteins available in GenBank; Full ESTs: all full-length *T. parva* EST data in GenBank; Partial ESTs: all partial-length *T. parva* EST data in GenBank; GeneMark-ES: gene model predictions using the GeneMark-ES gene prediction software (7); FGENESH: gene model predictions using FGENESH (8); SNAP: gene model predictions using SNAP (9); Glimmer: gene model predictions using Glimmer (10). Each EVM prediction (x-axis) was evaluated and weights were changed to compensate for the kinds of errors (over-merged/split genes, missed genes, etc.) in the previous prediction, relative to a validation gene set.

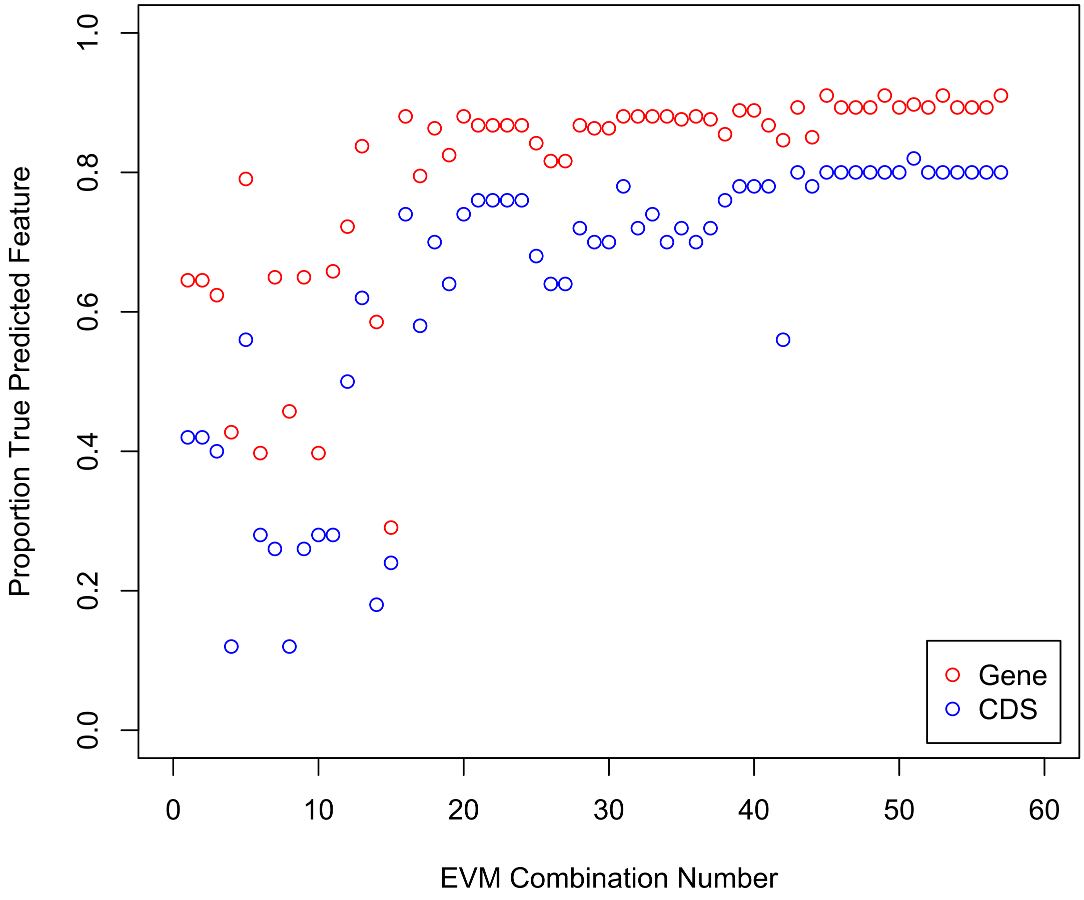


**Supplemental Figure S11. The percentage of validated genes, or coding exons correctly predicted by EVM with each evidence combination.**

To assess improvement in gene predictions, we tested the ability of each EVM prediction to correctly predict the locations of exons/introns in the high-quality, independent, validation set of ~300 *T. parva* genes. As the relative weights in each EVM combination were tweaked, there was an increase in the number of true-predictions (perfectly predicted) both at the level of the whole CDS (blue) and individual exons (red).

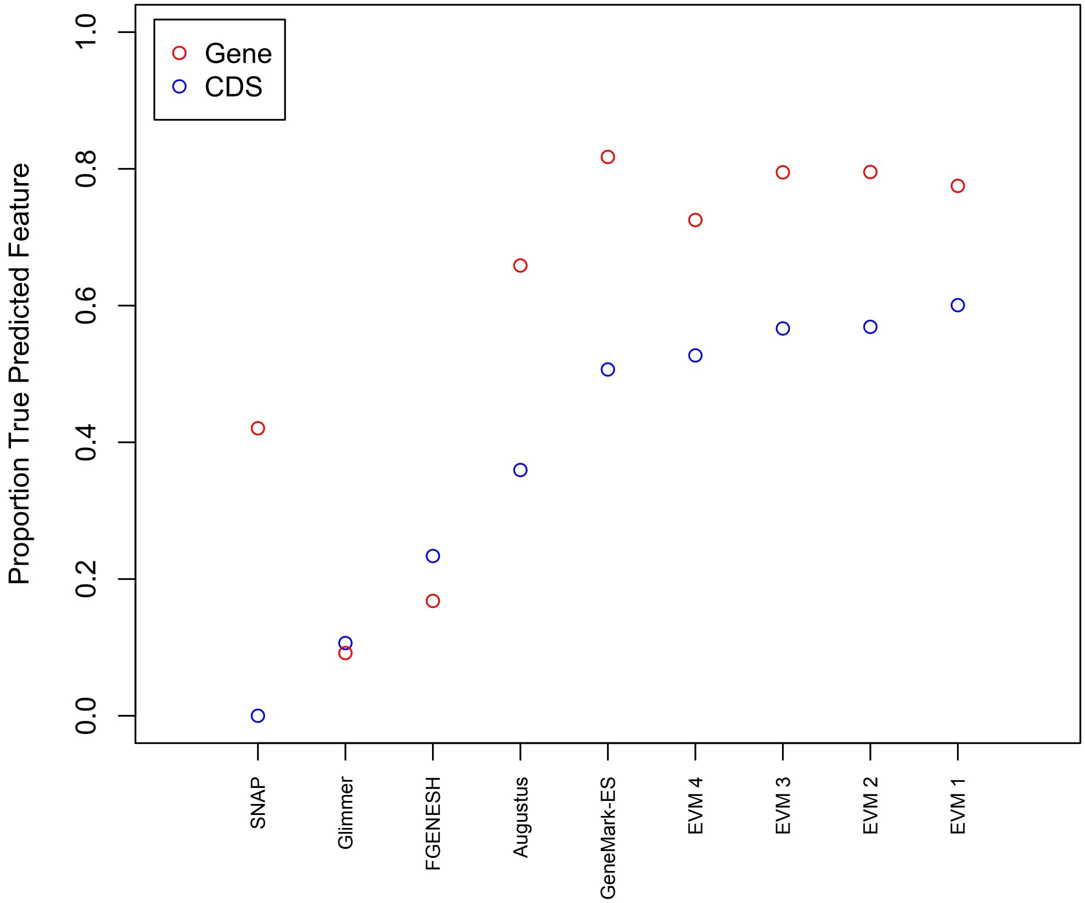


**Supplemental Figure S112. A comparison of the prediction accuracy of each gene predictor used in this study.**

Only the top four EVM predictions are shown. Given the unusual nature of the *T. parva* genome (and piroplasms in general), which is small, gene-dense, and AT-rich, we wanted to retrospectively determine which gene prediction programs most accurately predicted the final *T. parva* genome annotation. While the top four EVM predictions performed the best (EVM 1-4), GeneMark-ES true-predicted accuracy was comparable at both the complete CDS (blue) and exon (red) levels.

**Supplemental Table S1.** RNAseq read counts, length and GC content of each *T. parva* chromosome

| Chromosome | GenBank Locus ID | Reads Mapped | Coding Genes 2005 | Coding Genes New | Length (bp) | GC content (%) |
|---|---|---|---|---|---|---|
| 1 | AAGK01000001 | 4778302 | 1221 | 1200 | 2540030 | 34.16 |
| 2 | AAGK01000002 | 3971696 | 958 | 970 | 1971884 | 34.38 |
| 3 | AAGK01000005 | 1619758 | 623 | 616 | 1317241 | 33.87 |
| 3 | AAGK01000006 | 2072524 | 281 | 276 | 570487 | 33.63 |
| 3 | AAGK01000007 | 2238 | 18 | 18 | 41585 | 38.74 |
| 3 | AAGK01000008 | 264 | 6 | 6 | 13275 | 40.90 |
| 4 | AAGK01000004 | 2738272 | 921 | 915 | 1835834 | 33.95 |
| 4 | AAGK01000003 | 11408 | 18 | 10 | 17691 | 31.04 |
| Apicoplast | AAGK01000009 | 7834 | 44 | 44 | 39579 | 19.48 |

Out of the 21704856 total mapped reads, 15,202,296 (70.04%) mapped to *T. parva*, and 6,502,560 (29.96%) mapped to *B. taurus* with TopHat.

**Supplemental Table S2.** A comparison of genome characteristics of *T. parva* Muguga to several other piroplasms and *Plasmodium falciparum* 3D7.

| Features | This Paper | Tp | Ta | Te | Bb | Pf |
|---|---|---|---|---|---|---|
| Size (Mbp) | 8.3 | 8.3 | 8.35 | 11.6 | 8.3 | 22.8 |
| Number of chromosomes | 4 | 4 | 4 | 4 | 4 | 14 |
| Total G+C composition (%) | 34.1 | 34.1 | 32.5 | 39.5 | 41.8 | 19.4 |
| Number of nuclear coding genes | 4011 | 4035 | 3087 | 5330 | 3670 | 5368 |
| Average CDS length (bp) | 1457 | 1407 | 1600 | 1472 | 1514 | 2283 |
| Percent Genes with introns | 73.9 | 73.6 | 70.6 | 52.4 | 61.5 | 53.9 |
| Mean length of intergenic region (bp) | 307 | 405 | 396 | 550 | 589 | 1694 |
| G+C composition of intergenic regions | 30.9 | 26.2 | 24.1 | 39.3 | 37 | 13.8 |
| G+C composition of exons | 36.5 | 37.6 | 35.7 | 39.8 | 44 | 23.7 |
| G+C composition of introns | 24.7 | 25.4 | 24.4 | 37.6 | 35.9 | 13.6 |
| Percent Coding | 71.0 | 68.4 | 72.9 | 69.3 | 70.3 | 52.6 |
| Gene Density* | 2081 | 2057 | 2195 | 2185 | 2228 | 4338 |

All numbers except column "This paper" are from Kappmeyer *et al.,* 2012 (11); *Gene density = genome size / number of protein-coding genes; Pf = *Plasmodium falciparum* strain 3D7; This Paper = *Theileria parva* strain Muguga (new annotation); Tp = *Theileria parva* strain Muguga (2005 annotation); Ta = *Theileria annulata* strain Ankara; Bb = *Babesia bovis* strain T2Bo; Te = *Theileria equi* strain WA. Mean intergenic length calculated as the distance between coding sequences of adjacent genes.

**Supplemental Table S3.** A list of the expression levels (RPKM = reads per kilobase of transcript per million reads) of known *T. parva* antigens. Note: p32 and p67 are humoral antigens.

| Gene Name | 2005 ID | Locus Tag | RPKM | Antigen | |
|---|---|---|---|---|---|
| | | | | CD4 | CD8 |
| PIM | TP04_0051 | TpMuguga_04g00051 | 7545 | + | + |
| Tp7 | TP02_0244 | TpMuguga_02g00244 | 4358 | + | + |
| Tp9 | TP02_0895 | TpMuguga_02g00895 | 2974 | + | + |
| Tp12 | TP01_1091 | TpMuguga_01g01091 | 2619 | - | + |
| - | TP01_1182 | TpMuguga_01g01182 | 2252 | + | - |
| Tp8 | TP02_0140 | TpMuguga_02g00140 | 1869 | - | + |
| p104 | TP04_0437 | TpMuguga_04g00437 | 1024 | + | - |
| - | TP03_0655 | TpMuguga_03g00655 | 846 | + | - |
| N10 | TP01_1074 | TpMuguga_01g01074 | 781 | + | - |
| Tp2 | TP01_0056 | TpMuguga_01g00056 | 376 | + | + |
| Tp6 | TP01_0188 | TpMuguga_01g00188 | 317 | - | + |
| p150 | TP03_0861 | TpMuguga_03g00861 | 314 | + | - |
| p32 | TP01_1056 | TpMuguga_01g01056 | 209 | ? | ? |
| - | TP01_1225 | TpMuguga_01g01225 | 196 | + | - |
| Tp1 | TP03_0849 | TpMuguga_03g00849 | 174 | + | + |
| Tp4 | TP03_0210 | TpMuguga_03g00210 | 131 | - | + |
| N36 | TP04_0916 | TpMuguga_04g00916 | 130 | + | - |

| | | | | | |
|---|---|---|---|---|---|
| Tp5 | TP02_0767 | TpMuguga_02g00767 | 111 | - | + |
| Tp10 | TP04_0772 | TpMuguga_04g00772 | 108 | - | + |
| - | TP03_0263 | TpMuguga_03g00263 | 105 | + | + |
| - | TP01_1078 | TpMuguga_01g01078 | 43 | + | - |
| - | TP04_0917 | TpMuguga_04g00917 | 38 | + | - |
| - | TP02_0958 | TpMuguga_02g00958 | 32 | + | - |
| N43 | TP01_1077 | TpMuguga_01g01077 | 17 | + | - |
| - | TP04_0164 | TpMuguga_04g00164 | 16 | - | + |
| - | TP01_1081 | TpMuguga_01g01081 | 11 | + | - |
| p67 | TP03_0287 | TpMuguga_03g00287 | 9 | + | + |
| Tp3 | TP01_0868 | TpMuguga_01g00868 | 7 | - | + |
| - | TP01_1082 | TpMuguga_01g01082 | 7 | + | - |
| - | TP04_0683 | TpMuguga_04g00683 | 2 | + | - |
| N60 | TP01_0726 | TpMuguga_01g00726 | 1 | + | - |
| - | TP02_0123 | TpMuguga_02g00123 | 1 | + | + |
| - | TP02_0243 | TpMuguga_02g00243 | 1 | + | - |
| - | TP04_0752 | TpMuguga_04g00752 | 1 | + | - |
| - | TP02_0010 | TpMuguga_02g00010 | 0 | + | - |

**Supplemental Table S4.** A list of the highest-expressed genes in the *T. parva* schizont RNAseq dataset.

| Contig | Locus Tag | Product_Name | RPKM | 2005 Annotation ID |
|---|---|---|---|---|
| 1 | TpMuguga_01g01234 | hypothetical protein | 26824 | TP01_1234 |
| 3 | TpMuguga_03g00931 | hypothetical protein | 26823 | TP03_0931 |
| 1 | TpMuguga_01g01236 | hypothetical protein | 12454 | TP01_1236 |
| 3 | TpMuguga_03g00933 | hypothetical protein | 12453 | TP03_0933 |
| 2 | TpMuguga_02g00148 | Heat shock 70 kDa protein (12) | 11644 | TP02_0148 |
| 4 | TpMuguga_04g00383 | Glyceraldehyde 3-phosphate dehydrogenase NAD binding domain | 8698 | TP04_0383 |
| 4 | TpMuguga_04g00051 | Polymorphic Immunodominant Molecule (13) | 7545 | TP04_0051 |
| 1 | TpMuguga_01g00726 | Elongation factor Tu GTP binding domain | 6406 | TP01_0726 |
| 2 | TpMuguga_02g00244 | Heat shock protein 90 (14) | 4358 | TP02_0244 |
| 2 | TpMuguga_02g00903 | Actin | 4183 | TP02_0903 |
| 1 | TpMuguga_01g00244 | Cyclophilin type peptidyl-prolyl cis-trans isomerase/CLD | 3322 | TP01_0244 |
| 2 | TpMuguga_02g00895 | Tp9 (15) | 2974 | TP02_0895 |

| | | | | |
|---|---|---|---|---|
| 3 | TpMuguga_03g00152 | Core histone H2A/H2B/H3/H4 | 2852 | TP03_0152 |
| 4 | TpMuguga_04g00616 | hypothetical protein | 2628 | TP04_0616 |
| 1 | TpMuguga_01g01091 | hypothetical protein | 2619 | TP01_1091 |
| 1 | TpMuguga_01g01235 | hypothetical protein | 2522 | TP01_1235 |
| 3 | TpMuguga_03g00932 | hypothetical protein | 2521 | TP03_0932 |
| 3 | TpMuguga_03g00419 | Thioredoxin | 2477 | TP03_0419 |
| 4 | TpMuguga_04g00050 | Ribosomal protein S19 | 2364 | TP04_0050 |
| 1 | TpMuguga_01g01182 | lactate/malate dehydrogenase alpha/beta C-terminal domain | 2252 | TP01_1182 |

**Supplemental Table S5.** A table of key *T. parva* genes with reads per kilobase of transcript per million reads of zero (*Tp* = *Theileria parva* Muguga; *Pf* = *Plasmodium falciparum* 3D7).

| **Product Category** | ***Tp* Locus Tag (*Pf* Ortholog)** |
|---|---|
| hypothetical protein | TpMuguga_01g00436, TpMuguga_01g00462, TpMuguga_01g00561, TpMuguga_01g00693, TpMuguga_01g00764<br><br>TpMuguga_02g00473, TpMuguga_02g00502, TpMuguga_02g00618, TpMuguga_02g00855, TpMuguga_02g00876,<br><br>TpMuguga_03g00435<br><br>TpMuguga_04g00101, TpMuguga_04g00276, TpMuguga_04g00479, TpMuguga_04g00480, TpMuguga_04g00585, TpMuguga_04g00591, TpMuguga_04g00782, TpMuguga_04g00839, TpMuguga_04g00840, TpMuguga_04g00841, TpMuguga_04g02360<br><br>TpMuguga_05g00009, TpMuguga_05g00010, TpMuguga_05g00027, |

| | |
|---|---|
| | TpMuguga_05g00008(PFC10_API0036), TpMuguga_05g00001(PFC10_API0055), TpMuguga_05g00025(PFC10_API0020) |
| *Tpr* family protein | TpMuguga_03g00917, TpMuguga_02g00564 |
| Domain of unknown function DUF529 | TpMuguga_01g00685, TpMuguga_04g00145 |

**Supplemental Table S6.** A description of the top 20 largest multi-gene families defined by OrthoMCL in *T. parva* Muguga and their conservation in *T. annulata* (Ta), *T. orientalis* (To), and *T. equi* (Te), as defined by Jaccard-filtered clusters of orthologous genes.

| Family Number | Number of Proteins | Name | Ta | To | Te |
|---|---|---|---|---|---|
| 1 | 91 | SVSP family | 50 | 0 | 0 |
| 2 | 49 | Hypothetical FAINT-domain containing proteins | 46 | 16 | 0 |
| 3 | 36 | Protein tyrosine kinases | 80 | 0 | 0 |
| 4 | 17 | *Tpr* family | 13 | 0 | 0 |
| 5 | 13 | DEAD/DEAH box helicases | 18 | 32 | 41 |
| 6 | 12 | TashAT family | 13 | 12 | 13 |
| 7 | 10 | DnaJ chaperon proteins | 10 | 5 | 25 |
| 8 | 8 | ATPases | 7 | 7 | 9 |
| 9 | 8 | Hypothetical proteins | 8 | 7 | 8 |
| 10 | 8 | ABC transporters | 8 | 7 | 7 |
| 11 | 8 | Hypothetical proteins | 13 | 5 | 0 |
| 12 | 8 | Hypothetical proteins | 14 | 4 | 0 |
| 13 | 7 | Hypothetical FAINT-domain containing proteins | 8 | 5 | 107 |
| 14 | 7 | Cyclophilin type peptidyl-prolyl cis-trans isomerases | 8 | 9 | 8 |

| | | | | | |
|---|---|---|---|---|---|
| 15 | 7 | Secreted hypothetical proteins | 7 | 7 | 7 |
| 16 | 7 | Ubiquitin-conjugating enzymes | 9 | 6 | 3 |
| 17 | 7 | Ras family | 8 | 4 | 1 |
| 18 | 6 | Hypothetical proteins | 5 | 7 | 58 |
| 19 | 6 | Secreted FAINT-domain containing | 7 | 7 | 7 |
| 20 | 6 | Secreted haloacid dehalogenase-like hydrolases | 6 | 6 | 6 |

**Supplemental Table S7.** Summary of the top-ranked Phyre2 hits for each proposed Alg homolog discussed in this study.

| ***T. parva* Locus Tag** | **Homolog** | **Phyre2 Homolog** | **Phyre2 Confidence (%)** | **Resolution (Angstroms)** | **Identity (%)** | **Residues modelled at >90% confidence** |
|---|---|---|---|---|---|---|
| TpMuguga_01g02045 | Alg14 | *Pseudomonas aeroginosa* MurG | 98 | 1.9 | 17 | 62 |
| TpMuguga_01g00118 | Alg7 | *Homo sapiens* dpagt1 | 100 | 3.2 | 34 | 100 |
| TpMuguga_02g00515 | Alg13 | *Saccharomyces cerevisiae* alg13 | 100 | UNK | 26 | 94 |

**Supplemental Table S8.** The exon distribution of the validation and training sets used for gene prediction.

| **Exon #** | **validation** | **training** |
|---|---|---|
| 1 | 12 | 163 |
| 2 | 8 | 74 |
| 3 | 5 | 26 |
| 4 | 3 | 2 |
| 5 | 4 | 5 |
| 6 | 3 | 5 |
| 7 | 3 | 3 |
| 8 | 4 | 5 |
| 9 | 1 | 1 |
| 10 | 3 | 3 |
| 11 | 1 | 1 |
| 12 | 2 | 1 |
| 13 | 1 | 1 |
| 14 | 0 | 1 |
| 15 | 0 | 1 |
| **Total** | 50 | 292 |

diversity within parasite isolates used in a live vaccine against Theileria parva. *Int J Parasitol*, **46**, 495-506.
