## Additional Data 2 for "Re-annotation of the *Theileria parva* genome refines 53% of the proteome and uncovers essential components of N-glycosylation, a conserved pathway in many organisms"

**Additional file 2: Pairs of consecutive genes with overlap in UTR only or both UTR and CDS.**

| Gene_ID | Overlapping_Gene | Bp_Overlap_Total | Bp_Overlap_CDS | Same_Strand_Test |
|---|---|---|---|---|
| TpMuguga_01g00018 | TpMuguga_01g00019 | 17 | 0 | TRUE |
| TpMuguga_01g00019 | TpMuguga_01g00018 | 17 | 0 | TRUE |
| TpMuguga_01g02180 | TpMuguga_01g02190 | 4 | 4 | TRUE |
| TpMuguga_01g02190 | TpMuguga_01g02180 | 4 | 4 | TRUE |
| TpMuguga_01g00026 | TpMuguga_01g00027 | 719 | 0 | FALSE |
| TpMuguga_01g00028 | TpMuguga_01g00027 | 81 | 0 | FALSE |
| TpMuguga_01g00027 | TpMuguga_01g00026 | 719 | 0 | FALSE |
| TpMuguga_01g00027 | TpMuguga_01g00028 | 81 | 0 | FALSE |
| TpMuguga_01g00030 | TpMuguga_01g00029 | 24 | 0 | TRUE |
| TpMuguga_01g00029 | TpMuguga_01g00030 | 24 | 0 | TRUE |
| TpMuguga_01g02330 | TpMuguga_01g00031 | 34 | 12 | FALSE |
| TpMuguga_01g00031 | TpMuguga_01g02330 | 34 | 12 | FALSE |
| TpMuguga_01g00040 | TpMuguga_01g00039 | 5 | 0 | TRUE |
| TpMuguga_01g00039 | TpMuguga_01g00040 | 5 | 0 | TRUE |
| TpMuguga_01g00042 | TpMuguga_01g00041 | 201 | 0 | FALSE |
| TpMuguga_01g00041 | TpMuguga_01g00042 | 201 | 0 | FALSE |
| TpMuguga_01g00044 | TpMuguga_01g00043 | 6 | 0 | TRUE |
| TpMuguga_01g00043 | TpMuguga_01g00044 | 6 | 0 | TRUE |
| TpMuguga_01g00048 | TpMuguga_01g00047 | 20 | 0 | FALSE |
| TpMuguga_01g00047 | TpMuguga_01g00048 | 20 | 0 | FALSE |
| TpMuguga_01g02170 | TpMuguga_01g00051 | 86 | 4 | FALSE |
| TpMuguga_01g00051 | TpMuguga_01g02170 | 86 | 4 | FALSE |
| TpMuguga_01g02230 | TpMuguga_01g00053 | 449 | 0 | FALSE |
| TpMuguga_01g00053 | TpMuguga_01g02230 | 449 | 0 | FALSE |
| TpMuguga_01g00055 | TpMuguga_01g00054 | 24 | 0 | TRUE |
| TpMuguga_01g00054 | TpMuguga_01g00055 | 24 | 0 | TRUE |
| TpMuguga_01g00060 | TpMuguga_01g00061 | 80 | 0 | FALSE |
| TpMuguga_01g00061 | TpMuguga_01g00060 | 80 | 0 | FALSE |
| TpMuguga_01g02310 | TpMuguga_01g00071 | 86 | 0 | FALSE |
| TpMuguga_01g00071 | TpMuguga_01g02310 | 86 | 0 | FALSE |
| TpMuguga_01g00075 | TpMuguga_01g00076 | 338 | 0 | FALSE |
| TpMuguga_01g00076 | TpMuguga_01g00075 | 338 | 0 | FALSE |
| TpMuguga_01g00077 | TpMuguga_01g00078 | 333 | 48 | FALSE |
| TpMuguga_01g00078 | TpMuguga_01g00077 | 333 | 48 | FALSE |
| TpMuguga_01g00083 | TpMuguga_01g00084 | 20 | 0 | TRUE |
| TpMuguga_01g00084 | TpMuguga_01g00083 | 20 | 0 | TRUE |
| TpMuguga_01g00086 | TpMuguga_01g00085 | 102 | 0 | FALSE |
| TpMuguga_01g00086 | TpMuguga_01g00087 | 266 | 0 | FALSE |
| TpMuguga_01g00085 | TpMuguga_01g00086 | 102 | 0 | FALSE |
| TpMuguga_01g00085 | TpMuguga_01g01239 | 15 | 0 | TRUE |
| TpMuguga_01g01239 | TpMuguga_01g00085 | 15 | 0 | TRUE |
| TpMuguga_01g00087 | TpMuguga_01g00086 | 266 | 0 | FALSE |
| TpMuguga_01g00098 | TpMuguga_01g00097 | 32 | 0 | TRUE |
| TpMuguga_01g00097 | TpMuguga_01g00098 | 32 | 0 | TRUE |
| TpMuguga_01g00102 | TpMuguga_01g00101 | 11 | 0 | TRUE |
| TpMuguga_01g00101 | TpMuguga_01g00102 | 11 | 0 | TRUE |
| TpMuguga_01g00103 | TpMuguga_01g00104 | 7 | 0 | TRUE |
| TpMuguga_01g00104 | TpMuguga_01g00103 | 7 | 0 | TRUE |
| TpMuguga_01g00107 | TpMuguga_01g02240 | 231 | 0 | FALSE |
| TpMuguga_01g02240 | TpMuguga_01g00107 | 231 | 0 | FALSE |
| TpMuguga_01g00113 | TpMuguga_01g00112 | 42 | 0 | FALSE |
| TpMuguga_01g00112 | TpMuguga_01g00113 | 42 | 0 | FALSE |
| TpMuguga_01g02110 | TpMuguga_01g00115 | 15 | 0 | FALSE |
| TpMuguga_01g00115 | TpMuguga_01g02110 | 15 | 0 | FALSE |
| TpMuguga_01g02070 | TpMuguga_01g00119 | 20 | 0 | TRUE |
| TpMuguga_01g00119 | TpMuguga_01g02070 | 20 | 0 | TRUE |
| TpMuguga_01g00124 | TpMuguga_01g01240 | 54 | 0 | TRUE |
| TpMuguga_01g01240 | TpMuguga_01g00124 | 54 | 0 | TRUE |
| TpMuguga_01g00125 | TpMuguga_01g01241 | 6 | 0 | TRUE |
| TpMuguga_01g00125 | TpMuguga_01g00126 | 89 | 0 | FALSE |
| TpMuguga_01g01241 | TpMuguga_01g00125 | 6 | 0 | TRUE |
| TpMuguga_01g00126 | TpMuguga_01g00125 | 89 | 0 | FALSE |
| TpMuguga_01g00127 | TpMuguga_01g00128 | 44 | 0 | FALSE |
| TpMuguga_01g00128 | TpMuguga_01g00127 | 44 | 0 | FALSE |
| TpMuguga_01g00129 | TpMuguga_01g00130 | 125 | 0 | FALSE |
| TpMuguga_01g00130 | TpMuguga_01g00129 | 125 | 0 | FALSE |

| | | | | |
|---|---|---|---|---|
| TpMuguga_01g00137 | TpMuguga_01g00136 | 5 | 0 | FALSE |
| TpMuguga_01g00136 | TpMuguga_01g00137 | 5 | 0 | FALSE |
| TpMuguga_01g00143 | TpMuguga_01g02465 | 15 | 0 | FALSE |
| TpMuguga_01g02465 | TpMuguga_01g00143 | 15 | 0 | FALSE |
| TpMuguga_01g00149 | TpMuguga_01g00150 | 99 | 0 | FALSE |
| TpMuguga_01g02590 | TpMuguga_01g00150 | 4 | 4 | FALSE |
| TpMuguga_01g00150 | TpMuguga_01g00149 | 99 | 0 | FALSE |
| TpMuguga_01g00150 | TpMuguga_01g02590 | 4 | 4 | FALSE |
| TpMuguga_01g00151 | TpMuguga_01g02625 | 43 | 0 | FALSE |
| TpMuguga_01g02625 | TpMuguga_01g00151 | 43 | 0 | FALSE |
| TpMuguga_01g00160 | TpMuguga_01g00159 | 121 | 0 | FALSE |
| TpMuguga_01g00160 | TpMuguga_01g00161 | 11 | 0 | FALSE |
| TpMuguga_01g00159 | TpMuguga_01g00160 | 121 | 0 | FALSE |
| TpMuguga_01g00161 | TpMuguga_01g00160 | 11 | 0 | FALSE |
| TpMuguga_01g00162 | TpMuguga_01g00163 | 11 | 0 | TRUE |
| TpMuguga_01g00163 | TpMuguga_01g00162 | 11 | 0 | TRUE |
| TpMuguga_01g00173 | TpMuguga_01g00172 | 45 | 0 | FALSE |
| TpMuguga_01g00172 | TpMuguga_01g00173 | 45 | 0 | FALSE |
| TpMuguga_01g00180 | TpMuguga_01g00181 | 3 | 0 | TRUE |
| TpMuguga_01g00181 | TpMuguga_01g00180 | 3 | 0 | TRUE |
| TpMuguga_01g00183 | TpMuguga_01g00182 | 136 | 0 | FALSE |
| TpMuguga_01g00182 | TpMuguga_01g00183 | 136 | 0 | FALSE |
| TpMuguga_01g00190 | TpMuguga_01g00189 | 79 | 0 | FALSE |
| TpMuguga_01g00189 | TpMuguga_01g00190 | 79 | 0 | FALSE |
| TpMuguga_01g00192 | TpMuguga_01g00193 | 22 | 0 | TRUE |
| TpMuguga_01g00193 | TpMuguga_01g00192 | 22 | 0 | TRUE |
| TpMuguga_01g02045 | TpMuguga_01g00196 | 82 | 23 | FALSE |
| TpMuguga_01g00196 | TpMuguga_01g02045 | 82 | 23 | FALSE |
| TpMuguga_01g00199 | TpMuguga_01g00200 | 9 | 0 | FALSE |
| TpMuguga_01g00200 | TpMuguga_01g00199 | 9 | 0 | FALSE |
| TpMuguga_01g02055 | TpMuguga_01g00206 | 149 | 0 | FALSE |
| TpMuguga_01g00206 | TpMuguga_01g02055 | 149 | 0 | FALSE |
| TpMuguga_01g00209 | TpMuguga_01g00210 | 4 | 0 | FALSE |
| TpMuguga_01g00210 | TpMuguga_01g00209 | 4 | 0 | FALSE |
| TpMuguga_01g00224 | TpMuguga_01g00223 | 10 | 0 | FALSE |
| TpMuguga_01g00223 | TpMuguga_01g00224 | 10 | 0 | FALSE |
| TpMuguga_01g00244 | TpMuguga_01g00245 | 6 | 0 | FALSE |
| TpMuguga_01g00245 | TpMuguga_01g00244 | 6 | 0 | FALSE |
| TpMuguga_01g00249 | TpMuguga_01g00250 | 77 | 0 | FALSE |
| TpMuguga_01g00250 | TpMuguga_01g00249 | 77 | 0 | FALSE |
| TpMuguga_01g00266 | TpMuguga_01g00267 | 30 | 0 | FALSE |
| TpMuguga_01g00267 | TpMuguga_01g00266 | 30 | 0 | FALSE |
| TpMuguga_01g02345 | TpMuguga_01g00272 | 171 | 0 | FALSE |
| TpMuguga_01g00272 | TpMuguga_01g02345 | 171 | 0 | FALSE |
| TpMuguga_01g00277 | TpMuguga_01g00276 | 23 | 0 | TRUE |
| TpMuguga_01g00277 | TpMuguga_01g00278 | 45 | 0 | FALSE |
| TpMuguga_01g00276 | TpMuguga_01g00277 | 23 | 0 | TRUE |
| TpMuguga_01g00278 | TpMuguga_01g00277 | 45 | 0 | FALSE |
| TpMuguga_01g00284 | TpMuguga_01g00285 | 172 | 0 | FALSE |
| TpMuguga_01g00285 | TpMuguga_01g00284 | 172 | 0 | FALSE |
| TpMuguga_01g00290 | TpMuguga_01g00289 | 25 | 0 | FALSE |
| TpMuguga_01g00289 | TpMuguga_01g00290 | 25 | 0 | FALSE |
| TpMuguga_01g00293 | TpMuguga_01g00292 | 117 | 0 | FALSE |
| TpMuguga_01g00292 | TpMuguga_01g00293 | 117 | 0 | FALSE |
| TpMuguga_01g00292 | TpMuguga_01g00291 | 65 | 0 | FALSE |
| TpMuguga_01g00291 | TpMuguga_01g00292 | 65 | 0 | FALSE |
| TpMuguga_01g02155 | TpMuguga_01g02685 | 15 | 0 | FALSE |
| TpMuguga_01g02685 | TpMuguga_01g02155 | 15 | 0 | FALSE |
| TpMuguga_01g00304 | TpMuguga_01g00303 | 26 | 0 | TRUE |
| TpMuguga_01g00303 | TpMuguga_01g00304 | 26 | 0 | TRUE |
| TpMuguga_01g02445 | TpMuguga_01g00305 | 1 | 0 | FALSE |
| TpMuguga_01g02445 | TpMuguga_01g00308 | 8 | 0 | TRUE |
| TpMuguga_01g00305 | TpMuguga_01g02445 | 1 | 0 | FALSE |
| TpMuguga_01g00308 | TpMuguga_01g02445 | 8 | 0 | TRUE |
| TpMuguga_01g00308 | TpMuguga_01g00309 | 140 | 0 | FALSE |
| TpMuguga_01g02410 | TpMuguga_01g00309 | 55 | 0 | FALSE |
| TpMuguga_01g00309 | TpMuguga_01g00308 | 140 | 0 | FALSE |
| TpMuguga_01g00309 | TpMuguga_01g02410 | 55 | 0 | FALSE |
| TpMuguga_01g00315 | TpMuguga_01g00314 | 262 | 0 | FALSE |
| TpMuguga_01g00315 | TpMuguga_01g00316 | 21 | 0 | FALSE |
| TpMuguga_01g00314 | TpMuguga_01g00315 | 262 | 0 | FALSE |

| | | | | |
|---|---|---|---|---|
| TpMuguga_01g00316 | TpMuguga_01g00315 | 21 | 0 | FALSE |
| TpMuguga_01g02145 | TpMuguga_01g02285 | 332 | 0 | FALSE |
| TpMuguga_01g00317 | TpMuguga_01g02285 | 139 | 0 | FALSE |
| TpMuguga_01g02285 | TpMuguga_01g02145 | 332 | 0 | FALSE |
| TpMuguga_01g02285 | TpMuguga_01g00317 | 139 | 0 | FALSE |
| TpMuguga_01g00326 | TpMuguga_01g00327 | 8 | 0 | TRUE |
| TpMuguga_01g00327 | TpMuguga_01g00326 | 8 | 0 | TRUE |
| TpMuguga_01g00327 | TpMuguga_01g00329 | 68 | 0 | TRUE |
| TpMuguga_01g00329 | TpMuguga_01g00327 | 68 | 0 | TRUE |
| TpMuguga_01g00331 | TpMuguga_01g00332 | 186 | 0 | FALSE |
| TpMuguga_01g00332 | TpMuguga_01g00331 | 186 | 0 | FALSE |
| TpMuguga_01g00337 | TpMuguga_01g00338 | 461 | 0 | FALSE |
| TpMuguga_01g00338 | TpMuguga_01g00337 | 461 | 0 | FALSE |
| TpMuguga_01g00346 | TpMuguga_01g00345 | 542 | 0 | FALSE |
| TpMuguga_01g00345 | TpMuguga_01g00346 | 542 | 0 | FALSE |
| TpMuguga_01g00350 | TpMuguga_01g00349 | 49 | 0 | FALSE |
| TpMuguga_01g00349 | TpMuguga_01g00350 | 49 | 0 | FALSE |
| TpMuguga_01g00363 | TpMuguga_01g00362 | 2 | 0 | FALSE |
| TpMuguga_01g00362 | TpMuguga_01g00363 | 2 | 0 | FALSE |
| TpMuguga_01g00377 | TpMuguga_01g00376 | 45 | 0 | FALSE |
| TpMuguga_01g00376 | TpMuguga_01g00377 | 45 | 0 | FALSE |
| TpMuguga_01g00382 | TpMuguga_01g02460 | 11 | 0 | TRUE |
| TpMuguga_01g02460 | TpMuguga_01g00382 | 11 | 0 | TRUE |
| TpMuguga_01g00385 | TpMuguga_01g00386 | 168 | 0 | FALSE |
| TpMuguga_01g00386 | TpMuguga_01g00385 | 168 | 0 | FALSE |
| TpMuguga_01g00393 | TpMuguga_01g00394 | 3 | 0 | FALSE |
| TpMuguga_01g00394 | TpMuguga_01g00393 | 3 | 0 | FALSE |
| TpMuguga_01g00396 | TpMuguga_01g00395 | 161 | 0 | FALSE |
| TpMuguga_01g00395 | TpMuguga_01g00396 | 161 | 0 | FALSE |
| TpMuguga_01g00398 | TpMuguga_01g00399 | 24 | 0 | TRUE |
| TpMuguga_01g00399 | TpMuguga_01g00398 | 24 | 0 | TRUE |
| TpMuguga_01g00403 | TpMuguga_01g00404 | 26 | 0 | TRUE |
| TpMuguga_01g00404 | TpMuguga_01g00403 | 26 | 0 | TRUE |
| TpMuguga_01g00422 | TpMuguga_01g00421 | 31 | 0 | TRUE |
| TpMuguga_01g00422 | TpMuguga_01g02800 | 16 | 0 | TRUE |
| TpMuguga_01g00421 | TpMuguga_01g00422 | 31 | 0 | TRUE |
| TpMuguga_01g02800 | TpMuguga_01g00422 | 16 | 0 | TRUE |
| TpMuguga_01g00426 | TpMuguga_01g00425 | 47 | 0 | FALSE |
| TpMuguga_01g00425 | TpMuguga_01g00426 | 47 | 0 | FALSE |
| TpMuguga_01g00431 | TpMuguga_01g00432 | 27 | 0 | TRUE |
| TpMuguga_01g00432 | TpMuguga_01g00431 | 27 | 0 | TRUE |
| TpMuguga_01g02100 | TpMuguga_01g00440 | 37 | 11 | FALSE |
| TpMuguga_01g00440 | TpMuguga_01g02100 | 37 | 11 | FALSE |
| TpMuguga_01g00440 | TpMuguga_01g00439 | 15 | 0 | TRUE |
| TpMuguga_01g00439 | TpMuguga_01g00440 | 15 | 0 | TRUE |
| TpMuguga_01g00445 | TpMuguga_01g01232 | 178 | 0 | FALSE |
| TpMuguga_01g01232 | TpMuguga_01g00445 | 178 | 0 | FALSE |
| TpMuguga_01g01233 | TpMuguga_01g01234 | 108 | 0 | FALSE |
| TpMuguga_01g01234 | TpMuguga_01g01233 | 108 | 0 | FALSE |
| TpMuguga_01g00473 | TpMuguga_01g00474 | 19 | 0 | TRUE |
| TpMuguga_01g00474 | TpMuguga_01g00473 | 19 | 0 | TRUE |
| TpMuguga_01g00481 | TpMuguga_01g00482 | 103 | 0 | FALSE |
| TpMuguga_01g00482 | TpMuguga_01g00481 | 103 | 0 | FALSE |
| TpMuguga_01g00505 | TpMuguga_01g00506 | 43 | 0 | FALSE |
| TpMuguga_01g00506 | TpMuguga_01g00505 | 43 | 0 | FALSE |
| TpMuguga_01g00507 | TpMuguga_01g00508 | 98 | 0 | FALSE |
| TpMuguga_01g00508 | TpMuguga_01g00507 | 98 | 0 | FALSE |
| TpMuguga_01g00509 | TpMuguga_01g02850 | 91 | 0 | FALSE |
| TpMuguga_01g02850 | TpMuguga_01g00509 | 91 | 0 | FALSE |
| TpMuguga_01g00512 | TpMuguga_01g00511 | 461 | 0 | FALSE |
| TpMuguga_01g00511 | TpMuguga_01g00512 | 461 | 0 | FALSE |
| TpMuguga_01g00520 | TpMuguga_01g00521 | 17 | 0 | TRUE |
| TpMuguga_01g00521 | TpMuguga_01g00520 | 17 | 0 | TRUE |
| TpMuguga_01g00525 | TpMuguga_01g02250 | 55 | 0 | FALSE |
| TpMuguga_01g02250 | TpMuguga_01g00525 | 55 | 0 | FALSE |
| TpMuguga_01g00532 | TpMuguga_01g00533 | 130 | 0 | FALSE |
| TpMuguga_01g00533 | TpMuguga_01g00532 | 130 | 0 | FALSE |
| TpMuguga_01g00557 | TpMuguga_01g00556 | 133 | 0 | FALSE |
| TpMuguga_01g00556 | TpMuguga_01g00557 | 133 | 0 | FALSE |
| TpMuguga_01g00556 | TpMuguga_01g00555 | 484 | 0 | FALSE |
| TpMuguga_01g00555 | TpMuguga_01g00556 | 484 | 0 | FALSE |

| | | | | |
|---|---|---|---|---|
| TpMuguga_01g00564 | TpMuguga_01g00563 | 30 | 0 | TRUE |
| TpMuguga_01g00563 | TpMuguga_01g00564 | 30 | 0 | TRUE |
| TpMuguga_01g00568 | TpMuguga_01g00567 | 436 | 0 | FALSE |
| TpMuguga_01g00567 | TpMuguga_01g00568 | 436 | 0 | FALSE |
| TpMuguga_01g00573 | TpMuguga_01g02580 | 7 | 0 | FALSE |
| TpMuguga_01g00573 | TpMuguga_01g00574 | 240 | 0 | FALSE |
| TpMuguga_01g02580 | TpMuguga_01g00573 | 7 | 0 | FALSE |
| TpMuguga_01g00574 | TpMuguga_01g00573 | 240 | 0 | FALSE |
| TpMuguga_01g01230 | TpMuguga_01g02360 | 104 | 0 | FALSE |
| TpMuguga_01g01230 | TpMuguga_01g01231 | 22 | 0 | TRUE |
| TpMuguga_01g02360 | TpMuguga_01g01230 | 104 | 0 | FALSE |
| TpMuguga_01g01231 | TpMuguga_01g01230 | 22 | 0 | TRUE |
| TpMuguga_01g02210 | TpMuguga_01g00581 | 10 | 0 | FALSE |
| TpMuguga_01g00581 | TpMuguga_01g02210 | 10 | 0 | FALSE |
| TpMuguga_01g00594 | TpMuguga_01g01244 | 84 | 0 | FALSE |
| TpMuguga_01g01244 | TpMuguga_01g00594 | 84 | 0 | FALSE |
| TpMuguga_01g00596 | TpMuguga_01g00597 | 267 | 0 | FALSE |
| TpMuguga_01g00597 | TpMuguga_01g00596 | 267 | 0 | FALSE |
| TpMuguga_01g02510 | TpMuguga_01g00601 | 8 | 0 | FALSE |
| TpMuguga_01g00601 | TpMuguga_01g02510 | 8 | 0 | FALSE |
| TpMuguga_01g00626 | TpMuguga_01g02825 | 275 | 0 | FALSE |
| TpMuguga_01g02825 | TpMuguga_01g00626 | 275 | 0 | FALSE |
| TpMuguga_01g02730 | TpMuguga_01g00632 | 70 | 0 | FALSE |
| TpMuguga_01g00632 | TpMuguga_01g02730 | 70 | 0 | FALSE |
| TpMuguga_01g00642 | TpMuguga_01g00643 | 19 | 0 | TRUE |
| TpMuguga_01g00643 | TpMuguga_01g00642 | 19 | 0 | TRUE |
| TpMuguga_01g00643 | TpMuguga_01g02450 | 9 | 0 | FALSE |
| TpMuguga_01g02450 | TpMuguga_01g00643 | 9 | 0 | FALSE |
| TpMuguga_01g00656 | TpMuguga_01g00657 | 121 | 0 | TRUE |
| TpMuguga_01g00657 | TpMuguga_01g00656 | 121 | 0 | TRUE |
| TpMuguga_01g00661 | TpMuguga_01g00660 | 18 | 0 | TRUE |
| TpMuguga_01g00661 | TpMuguga_01g00662 | 75 | 0 | FALSE |
| TpMuguga_01g00660 | TpMuguga_01g00661 | 18 | 0 | TRUE |
| TpMuguga_01g00662 | TpMuguga_01g00661 | 75 | 0 | FALSE |
| TpMuguga_01g00719 | TpMuguga_01g00718 | 126 | 17 | FALSE |
| TpMuguga_01g00718 | TpMuguga_01g00719 | 126 | 17 | FALSE |
| TpMuguga_01g02520 | TpMuguga_01g02725 | 10 | 0 | TRUE |
| TpMuguga_01g02725 | TpMuguga_01g02520 | 10 | 0 | TRUE |
| TpMuguga_01g02205 | TpMuguga_01g00730 | 22 | 0 | FALSE |
| TpMuguga_01g00730 | TpMuguga_01g02205 | 22 | 0 | FALSE |
| TpMuguga_01g00733 | TpMuguga_01g00734 | 9 | 0 | FALSE |
| TpMuguga_01g00734 | TpMuguga_01g00733 | 9 | 0 | FALSE |
| TpMuguga_01g02305 | TpMuguga_01g00736 | 219 | 0 | FALSE |
| TpMuguga_01g00736 | TpMuguga_01g02305 | 219 | 0 | FALSE |
| TpMuguga_01g00744 | TpMuguga_01g00745 | 198 | 0 | FALSE |
| TpMuguga_01g00745 | TpMuguga_01g00744 | 198 | 0 | FALSE |
| TpMuguga_01g00754 | TpMuguga_01g00753 | 2 | 0 | FALSE |
| TpMuguga_01g00753 | TpMuguga_01g00754 | 2 | 0 | FALSE |
| TpMuguga_01g02080 | TpMuguga_01g00765 | 1 | 0 | FALSE |
| TpMuguga_01g00765 | TpMuguga_01g02080 | 1 | 0 | FALSE |
| TpMuguga_01g00768 | TpMuguga_01g00769 | 31 | 0 | TRUE |
| TpMuguga_01g00769 | TpMuguga_01g00768 | 31 | 0 | TRUE |
| TpMuguga_01g00777 | TpMuguga_01g02795 | 8 | 0 | TRUE |
| TpMuguga_01g02795 | TpMuguga_01g00777 | 8 | 0 | TRUE |
| TpMuguga_01g02235 | TpMuguga_01g00781 | 48 | 0 | FALSE |
| TpMuguga_01g00781 | TpMuguga_01g02235 | 48 | 0 | FALSE |
| TpMuguga_01g00783 | TpMuguga_01g00784 | 15 | 0 | TRUE |
| TpMuguga_01g00784 | TpMuguga_01g00783 | 15 | 0 | TRUE |
| TpMuguga_01g02370 | TpMuguga_01g00785 | 495 | 0 | FALSE |
| TpMuguga_01g00785 | TpMuguga_01g02370 | 495 | 0 | FALSE |
| TpMuguga_01g00788 | TpMuguga_01g00789 | 78 | 0 | FALSE |
| TpMuguga_01g00789 | TpMuguga_01g00788 | 78 | 0 | FALSE |
| TpMuguga_01g00796 | TpMuguga_01g00795 | 78 | 0 | FALSE |
| TpMuguga_01g00795 | TpMuguga_01g00796 | 78 | 0 | FALSE |
| TpMuguga_01g00797 | TpMuguga_01g02565 | 73 | 0 | FALSE |
| TpMuguga_01g02565 | TpMuguga_01g00797 | 73 | 0 | FALSE |
| TpMuguga_01g00799 | TpMuguga_01g00800 | 30 | 0 | FALSE |
| TpMuguga_01g02340 | TpMuguga_01g00800 | 17 | 17 | TRUE |
| TpMuguga_01g00800 | TpMuguga_01g00799 | 30 | 0 | FALSE |
| TpMuguga_01g00800 | TpMuguga_01g02340 | 17 | 17 | TRUE |
| TpMuguga_01g00805 | TpMuguga_01g00806 | 485 | 0 | FALSE |

| | | | | |
|---|---|---|---|---|
| TpMuguga_01g00806 | TpMuguga_01g00805 | 485 | 0 | FALSE |
| TpMuguga_01g00816 | TpMuguga_01g00817 | 30 | 0 | TRUE |
| TpMuguga_01g00817 | TpMuguga_01g00816 | 30 | 0 | TRUE |
| TpMuguga_01g00821 | TpMuguga_01g00822 | 15 | 0 | FALSE |
| TpMuguga_01g00822 | TpMuguga_01g00821 | 15 | 0 | FALSE |
| TpMuguga_01g00826 | TpMuguga_01g02635 | 80 | 0 | FALSE |
| TpMuguga_01g02635 | TpMuguga_01g00826 | 80 | 0 | FALSE |
| TpMuguga_01g02770 | TpMuguga_01g00830 | 75 | 0 | FALSE |
| TpMuguga_01g00830 | TpMuguga_01g02770 | 75 | 0 | FALSE |
| TpMuguga_01g00842 | TpMuguga_01g00841 | 9 | 0 | TRUE |
| TpMuguga_01g00841 | TpMuguga_01g00842 | 9 | 0 | TRUE |
| TpMuguga_01g02550 | TpMuguga_01g02665 | 14 | 14 | TRUE |
| TpMuguga_01g02665 | TpMuguga_01g02550 | 14 | 14 | TRUE |
| TpMuguga_01g00845 | TpMuguga_01g00846 | 11 | 0 | TRUE |
| TpMuguga_01g00846 | TpMuguga_01g00845 | 11 | 0 | TRUE |
| TpMuguga_01g00855 | TpMuguga_01g00854 | 485 | 0 | FALSE |
| TpMuguga_01g00854 | TpMuguga_01g00855 | 485 | 0 | FALSE |
| TpMuguga_01g00858 | TpMuguga_01g02840 | 89 | 0 | FALSE |
| TpMuguga_01g00858 | TpMuguga_01g00859 | 7 | 0 | FALSE |
| TpMuguga_01g02840 | TpMuguga_01g00858 | 89 | 0 | FALSE |
| TpMuguga_01g00859 | TpMuguga_01g00858 | 7 | 0 | FALSE |
| TpMuguga_01g00864 | TpMuguga_01g00863 | 16 | 0 | FALSE |
| TpMuguga_01g00863 | TpMuguga_01g00864 | 16 | 0 | FALSE |
| TpMuguga_01g00877 | TpMuguga_01g00876 | 24 | 0 | TRUE |
| TpMuguga_01g00876 | TpMuguga_01g00877 | 24 | 0 | TRUE |
| TpMuguga_01g02560 | TpMuguga_01g00881 | 166 | 29 | FALSE |
| TpMuguga_01g00881 | TpMuguga_01g02560 | 166 | 29 | FALSE |
| TpMuguga_01g00884 | TpMuguga_01g00885 | 15 | 0 | TRUE |
| TpMuguga_01g00885 | TpMuguga_01g00884 | 15 | 0 | TRUE |
| TpMuguga_01g00891 | TpMuguga_01g00892 | 3 | 0 | FALSE |
| TpMuguga_01g00893 | TpMuguga_01g00892 | 9 | 0 | TRUE |
| TpMuguga_01g00892 | TpMuguga_01g00891 | 3 | 0 | FALSE |
| TpMuguga_01g00892 | TpMuguga_01g00893 | 9 | 0 | TRUE |
| TpMuguga_01g00895 | TpMuguga_01g00896 | 20 | 0 | FALSE |
| TpMuguga_01g00896 | TpMuguga_01g00895 | 20 | 0 | FALSE |
| TpMuguga_01g00908 | TpMuguga_01g00907 | 14 | 0 | TRUE |
| TpMuguga_01g00907 | TpMuguga_01g00908 | 14 | 0 | TRUE |
| TpMuguga_01g00914 | TpMuguga_01g00915 | 9 | 0 | FALSE |
| TpMuguga_01g00915 | TpMuguga_01g00914 | 9 | 0 | FALSE |
| TpMuguga_01g00921 | TpMuguga_01g00923 | 18 | 0 | TRUE |
| TpMuguga_01g00923 | TpMuguga_01g00921 | 18 | 0 | TRUE |
| TpMuguga_01g00942 | TpMuguga_01g00941 | 19 | 0 | FALSE |
| TpMuguga_01g00941 | TpMuguga_01g00942 | 19 | 0 | FALSE |
| TpMuguga_01g00951 | TpMuguga_01g00950 | 143 | 0 | FALSE |
| TpMuguga_01g00950 | TpMuguga_01g00951 | 143 | 0 | FALSE |
| TpMuguga_01g00955 | TpMuguga_01g00956 | 26 | 0 | TRUE |
| TpMuguga_01g02280 | TpMuguga_01g00952 | 5 | 0 | TRUE |
| TpMuguga_01g00952 | TpMuguga_01g02280 | 5 | 0 | TRUE |
| TpMuguga_01g00956 | TpMuguga_01g00955 | 26 | 0 | TRUE |
| TpMuguga_01g00962 | TpMuguga_01g02790 | 19 | 0 | FALSE |
| TpMuguga_01g00963 | TpMuguga_01g02790 | 36 | 0 | FALSE |
| TpMuguga_01g02790 | TpMuguga_01g00962 | 19 | 0 | FALSE |
| TpMuguga_01g02790 | TpMuguga_01g00963 | 36 | 0 | FALSE |
| TpMuguga_01g00967 | TpMuguga_01g00966 | 49 | 0 | TRUE |
| TpMuguga_01g00966 | TpMuguga_01g00967 | 49 | 0 | TRUE |
| TpMuguga_01g00969 | TpMuguga_01g00970 | 396 | 0 | FALSE |
| TpMuguga_01g00970 | TpMuguga_01g00969 | 396 | 0 | FALSE |
| TpMuguga_01g00979 | TpMuguga_01g00978 | 89 | 0 | FALSE |
| TpMuguga_01g00979 | TpMuguga_01g00980 | 39 | 0 | FALSE |
| TpMuguga_01g00978 | TpMuguga_01g00979 | 89 | 0 | FALSE |
| TpMuguga_01g00981 | TpMuguga_01g00980 | 34 | 0 | FALSE |
| TpMuguga_01g00980 | TpMuguga_01g00979 | 39 | 0 | FALSE |
| TpMuguga_01g00980 | TpMuguga_01g00981 | 34 | 0 | FALSE |
| TpMuguga_01g02675 | TpMuguga_01g02680 | 19 | 19 | TRUE |
| TpMuguga_01g02680 | TpMuguga_01g02675 | 19 | 19 | TRUE |
| TpMuguga_01g02680 | TpMuguga_01g00984 | 42 | 0 | TRUE |
| TpMuguga_01g00984 | TpMuguga_01g02680 | 42 | 0 | TRUE |
| TpMuguga_01g00994 | TpMuguga_01g00995 | 24 | 0 | TRUE |
| TpMuguga_01g00996 | TpMuguga_01g00995 | 198 | 0 | FALSE |
| TpMuguga_01g00996 | TpMuguga_01g00997 | 381 | 0 | FALSE |
| TpMuguga_01g00995 | TpMuguga_01g00994 | 24 | 0 | TRUE |

| | | | | |
|---|---|---|---|---|
| TpMuguga_01g00995 | TpMuguga_01g00996 | 198 | 0 | FALSE |
| TpMuguga_01g00997 | TpMuguga_01g00996 | 381 | 0 | FALSE |
| TpMuguga_01g00999 | TpMuguga_01g00998 | 5 | 0 | TRUE |
| TpMuguga_01g00998 | TpMuguga_01g00999 | 5 | 0 | TRUE |
| TpMuguga_01g01013 | TpMuguga_01g01012 | 23 | 0 | TRUE |
| TpMuguga_01g01012 | TpMuguga_01g01013 | 23 | 0 | TRUE |
| TpMuguga_01g01024 | TpMuguga_01g01023 | 20 | 0 | TRUE |
| TpMuguga_01g01024 | TpMuguga_01g01025 | 627 | 0 | FALSE |
| TpMuguga_01g01023 | TpMuguga_01g01024 | 20 | 0 | TRUE |
| TpMuguga_01g01025 | TpMuguga_01g01024 | 627 | 0 | FALSE |
| TpMuguga_01g01027 | TpMuguga_01g01026 | 13 | 0 | FALSE |
| TpMuguga_01g01026 | TpMuguga_01g01027 | 13 | 0 | FALSE |
| TpMuguga_01g01034 | TpMuguga_01g01033 | 39 | 0 | TRUE |
| TpMuguga_01g01033 | TpMuguga_01g01034 | 39 | 0 | TRUE |
| TpMuguga_01g01040 | TpMuguga_01g01041 | 24 | 0 | TRUE |
| TpMuguga_01g01041 | TpMuguga_01g01040 | 24 | 0 | TRUE |
| TpMuguga_01g01042 | TpMuguga_01g01043 | 144 | 0 | FALSE |
| TpMuguga_01g01043 | TpMuguga_01g01042 | 144 | 0 | FALSE |
| TpMuguga_01g02630 | TpMuguga_01g01052 | 24 | 0 | TRUE |
| TpMuguga_01g01052 | TpMuguga_01g02630 | 24 | 0 | TRUE |
| TpMuguga_01g01054 | TpMuguga_01g01053 | 9 | 0 | TRUE |
| TpMuguga_01g01053 | TpMuguga_01g01054 | 9 | 0 | TRUE |
| TpMuguga_01g01059 | TpMuguga_01g01060 | 19 | 0 | TRUE |
| TpMuguga_01g01060 | TpMuguga_01g01059 | 19 | 0 | TRUE |
| TpMuguga_01g01062 | TpMuguga_01g02105 | 10 | 0 | TRUE |
| TpMuguga_01g02105 | TpMuguga_01g01062 | 10 | 0 | TRUE |
| TpMuguga_01g01065 | TpMuguga_01g01064 | 148 | 22 | FALSE |
| TpMuguga_01g01064 | TpMuguga_01g01065 | 148 | 22 | FALSE |
| TpMuguga_01g01071 | TpMuguga_01g01070 | 32 | 0 | FALSE |
| TpMuguga_01g01071 | TpMuguga_01g01072 | 51 | 0 | TRUE |
| TpMuguga_01g01070 | TpMuguga_01g01071 | 32 | 0 | FALSE |
| TpMuguga_01g01072 | TpMuguga_01g01071 | 51 | 0 | TRUE |
| TpMuguga_01g02260 | TpMuguga_01g02290 | 4 | 4 | TRUE |
| TpMuguga_01g02290 | TpMuguga_01g02260 | 4 | 4 | TRUE |
| TpMuguga_01g01100 | TpMuguga_01g01101 | 3 | 0 | TRUE |
| TpMuguga_01g01101 | TpMuguga_01g01100 | 3 | 0 | TRUE |
| TpMuguga_01g02075 | TpMuguga_01g01112 | 168 | 0 | FALSE |
| TpMuguga_01g01112 | TpMuguga_01g02075 | 168 | 0 | FALSE |
| TpMuguga_01g01116 | TpMuguga_01g02805 | 14 | 0 | FALSE |
| TpMuguga_01g02805 | TpMuguga_01g01116 | 14 | 0 | FALSE |
| TpMuguga_01g02215 | TpMuguga_01g02870 | 3 | 0 | TRUE |
| TpMuguga_01g02870 | TpMuguga_01g02215 | 3 | 0 | TRUE |
| TpMuguga_01g01131 | TpMuguga_01g01132 | 433 | 0 | TRUE |
| TpMuguga_01g01132 | TpMuguga_01g01131 | 433 | 0 | TRUE |
| TpMuguga_01g01132 | TpMuguga_01g01133 | 50 | 0 | FALSE |
| TpMuguga_01g01134 | TpMuguga_01g01133 | 38 | 0 | TRUE |
| TpMuguga_01g01133 | TpMuguga_01g01132 | 50 | 0 | FALSE |
| TpMuguga_01g01133 | TpMuguga_01g01134 | 38 | 0 | TRUE |
| TpMuguga_01g01136 | TpMuguga_01g01135 | 19 | 0 | FALSE |
| TpMuguga_01g01135 | TpMuguga_01g01136 | 19 | 0 | FALSE |
| TpMuguga_01g01148 | TpMuguga_01g01149 | 208 | 0 | FALSE |
| TpMuguga_01g01149 | TpMuguga_01g01148 | 208 | 0 | FALSE |
| TpMuguga_01g01151 | TpMuguga_01g01150 | 5 | 0 | FALSE |
| TpMuguga_01g01150 | TpMuguga_01g01151 | 5 | 0 | FALSE |
| TpMuguga_01g01153 | TpMuguga_01g01152 | 113 | 0 | FALSE |
| TpMuguga_01g01153 | TpMuguga_01g02415 | 56 | 0 | TRUE |
| TpMuguga_01g01152 | TpMuguga_01g01153 | 113 | 0 | FALSE |
| TpMuguga_01g01156 | TpMuguga_01g02415 | 23 | 0 | TRUE |
| TpMuguga_01g02415 | TpMuguga_01g01153 | 56 | 0 | TRUE |
| TpMuguga_01g02415 | TpMuguga_01g01156 | 23 | 0 | TRUE |
| TpMuguga_01g02090 | TpMuguga_01g01180 | 94 | 0 | FALSE |
| TpMuguga_01g01180 | TpMuguga_01g02090 | 94 | 0 | FALSE |
| TpMuguga_01g01190 | TpMuguga_01g01191 | 64 | 0 | FALSE |
| TpMuguga_01g01191 | TpMuguga_01g01190 | 64 | 0 | FALSE |
| TpMuguga_01g01192 | TpMuguga_01g01248 | 1 | 0 | FALSE |
| TpMuguga_01g01248 | TpMuguga_01g01192 | 1 | 0 | FALSE |
| TpMuguga_01g01195 | TpMuguga_01g01194 | 114 | 0 | FALSE |
| TpMuguga_01g01194 | TpMuguga_01g01195 | 114 | 0 | FALSE |
| TpMuguga_01g01194 | TpMuguga_01g01193 | 42 | 0 | TRUE |
| TpMuguga_01g01193 | TpMuguga_01g01194 | 42 | 0 | TRUE |
| TpMuguga_01g01203 | TpMuguga_01g01204 | 139 | 0 | FALSE |

| | | | | |
|---|---|---|---|---|
| TpMuguga_01g01204 | TpMuguga_01g01203 | 139 | 0 | FALSE |
| TpMuguga_01g01210 | TpMuguga_01g01209 | 923 | 0 | FALSE |
| TpMuguga_01g01210 | TpMuguga_01g01211 | 24 | 0 | TRUE |
| TpMuguga_01g01209 | TpMuguga_01g01210 | 923 | 0 | FALSE |
| TpMuguga_01g01212 | TpMuguga_01g01211 | 17 | 0 | FALSE |
| TpMuguga_01g01211 | TpMuguga_01g01210 | 24 | 0 | TRUE |
| TpMuguga_01g01211 | TpMuguga_01g01212 | 17 | 0 | FALSE |
| TpMuguga_02g00003 | TpMuguga_02g02590 | 200 | 0 | FALSE |
| TpMuguga_02g00004 | TpMuguga_02g02590 | 601 | 135 | FALSE |
| TpMuguga_02g02590 | TpMuguga_02g00003 | 200 | 0 | FALSE |
| TpMuguga_02g02590 | TpMuguga_02g00004 | 601 | 135 | FALSE |
| TpMuguga_02g00008 | TpMuguga_02g00009 | 1 | 0 | TRUE |
| TpMuguga_02g00009 | TpMuguga_02g00008 | 1 | 0 | TRUE |
| TpMuguga_02g00026 | TpMuguga_02g02595 | 9 | 0 | FALSE |
| TpMuguga_02g02355 | TpMuguga_02g02595 | 9 | 0 | TRUE |
| TpMuguga_02g02595 | TpMuguga_02g00026 | 9 | 0 | FALSE |
| TpMuguga_02g02595 | TpMuguga_02g02355 | 9 | 0 | TRUE |
| TpMuguga_02g02165 | TpMuguga_02g02465 | 20 | 20 | TRUE |
| TpMuguga_02g00074 | TpMuguga_02g02465 | 157 | 0 | FALSE |
| TpMuguga_02g02465 | TpMuguga_02g02165 | 20 | 20 | TRUE |
| TpMuguga_02g02465 | TpMuguga_02g00074 | 157 | 0 | FALSE |
| TpMuguga_02g00088 | TpMuguga_02g00087 | 20 | 0 | TRUE |
| TpMuguga_02g00087 | TpMuguga_02g00088 | 20 | 0 | TRUE |
| TpMuguga_02g02195 | TpMuguga_02g00099 | 2 | 0 | FALSE |
| TpMuguga_02g02195 | TpMuguga_02g00100 | 20 | 0 | TRUE |
| TpMuguga_02g00099 | TpMuguga_02g02195 | 2 | 0 | FALSE |
| TpMuguga_02g00100 | TpMuguga_02g02195 | 20 | 0 | TRUE |
| TpMuguga_02g00100 | TpMuguga_02g00101 | 3 | 0 | FALSE |
| TpMuguga_02g00102 | TpMuguga_02g00101 | 41 | 0 | FALSE |
| TpMuguga_02g00101 | TpMuguga_02g00100 | 3 | 0 | FALSE |
| TpMuguga_02g00101 | TpMuguga_02g00102 | 41 | 0 | FALSE |
| TpMuguga_02g02005 | TpMuguga_02g00110 | 26 | 0 | TRUE |
| TpMuguga_02g00110 | TpMuguga_02g02005 | 26 | 0 | TRUE |
| TpMuguga_02g00113 | TpMuguga_02g00112 | 176 | 0 | FALSE |
| TpMuguga_02g00113 | TpMuguga_02g00114 | 21 | 0 | TRUE |
| TpMuguga_02g00112 | TpMuguga_02g00113 | 176 | 0 | FALSE |
| TpMuguga_02g00114 | TpMuguga_02g00113 | 21 | 0 | TRUE |
| TpMuguga_02g00115 | TpMuguga_02g00116 | 4 | 0 | FALSE |
| TpMuguga_02g00116 | TpMuguga_02g00115 | 4 | 0 | FALSE |
| TpMuguga_02g00118 | TpMuguga_02g00117 | 47 | 0 | TRUE |
| TpMuguga_02g00117 | TpMuguga_02g00118 | 47 | 0 | TRUE |
| TpMuguga_02g00120 | TpMuguga_02g00121 | 425 | 0 | FALSE |
| TpMuguga_02g00121 | TpMuguga_02g00120 | 425 | 0 | FALSE |
| TpMuguga_02g00124 | TpMuguga_02g00125 | 10 | 0 | FALSE |
| TpMuguga_02g00125 | TpMuguga_02g00124 | 10 | 0 | FALSE |
| TpMuguga_02g00126 | TpMuguga_02g00127 | 4 | 0 | FALSE |
| TpMuguga_02g00127 | TpMuguga_02g00126 | 4 | 0 | FALSE |
| TpMuguga_02g00132 | TpMuguga_02g02405 | 293 | 0 | FALSE |
| TpMuguga_02g02405 | TpMuguga_02g00132 | 293 | 0 | FALSE |
| TpMuguga_02g02405 | TpMuguga_02g02410 | 29 | 29 | TRUE |
| TpMuguga_02g02410 | TpMuguga_02g02405 | 29 | 29 | TRUE |
| TpMuguga_02g00135 | TpMuguga_02g00134 | 23 | 0 | FALSE |
| TpMuguga_02g00135 | TpMuguga_02g00136 | 16 | 0 | FALSE |
| TpMuguga_02g00134 | TpMuguga_02g00135 | 23 | 0 | FALSE |
| TpMuguga_02g00136 | TpMuguga_02g00135 | 16 | 0 | FALSE |
| TpMuguga_02g00138 | TpMuguga_02g00139 | 17 | 0 | TRUE |
| TpMuguga_02g00139 | TpMuguga_02g00138 | 17 | 0 | TRUE |
| TpMuguga_02g00146 | TpMuguga_02g00145 | 285 | 27 | FALSE |
| TpMuguga_02g00145 | TpMuguga_02g00146 | 285 | 27 | FALSE |
| TpMuguga_02g00149 | TpMuguga_02g02435 | 15 | 0 | TRUE |
| TpMuguga_02g02435 | TpMuguga_02g00149 | 15 | 0 | TRUE |
| TpMuguga_02g02255 | TpMuguga_02g00162 | 236 | 0 | FALSE |
| TpMuguga_02g00162 | TpMuguga_02g02255 | 236 | 0 | FALSE |
| TpMuguga_02g00167 | TpMuguga_02g00166 | 21 | 0 | FALSE |
| TpMuguga_02g00166 | TpMuguga_02g00167 | 21 | 0 | FALSE |
| TpMuguga_02g00180 | TpMuguga_02g00179 | 9 | 0 | TRUE |
| TpMuguga_02g00179 | TpMuguga_02g00180 | 9 | 0 | TRUE |
| TpMuguga_02g00181 | TpMuguga_02g00182 | 50 | 0 | FALSE |
| TpMuguga_02g00182 | TpMuguga_02g00181 | 50 | 0 | FALSE |
| TpMuguga_02g00188 | TpMuguga_02g00189 | 24 | 0 | TRUE |
| TpMuguga_02g00189 | TpMuguga_02g00188 | 24 | 0 | TRUE |

| | | | | |
|---|---|---|---|---|
| TpMuguga_02g00190 | TpMuguga_02g00191 | 43 | 0 | FALSE |
| TpMuguga_02g00191 | TpMuguga_02g00190 | 43 | 0 | FALSE |
| TpMuguga_02g00203 | TpMuguga_02g02125 | 183 | 0 | FALSE |
| TpMuguga_02g02125 | TpMuguga_02g00203 | 183 | 0 | FALSE |
| TpMuguga_02g02125 | TpMuguga_02g02345 | 10 | 0 | TRUE |
| TpMuguga_02g02345 | TpMuguga_02g02125 | 10 | 0 | TRUE |
| TpMuguga_02g00206 | TpMuguga_02g00205 | 362 | 0 | FALSE |
| TpMuguga_02g00205 | TpMuguga_02g00206 | 362 | 0 | FALSE |
| TpMuguga_02g00208 | TpMuguga_02g00207 | 16 | 0 | TRUE |
| TpMuguga_02g00207 | TpMuguga_02g00208 | 16 | 0 | TRUE |
| TpMuguga_02g02000 | TpMuguga_02g00229 | 32 | 0 | TRUE |
| TpMuguga_02g00229 | TpMuguga_02g02000 | 32 | 0 | TRUE |
| TpMuguga_02g00229 | TpMuguga_02g00230 | 79 | 0 | FALSE |
| TpMuguga_02g00230 | TpMuguga_02g00229 | 79 | 0 | FALSE |
| TpMuguga_02g00236 | TpMuguga_02g00237 | 57 | 0 | FALSE |
| TpMuguga_02g00237 | TpMuguga_02g00236 | 57 | 0 | FALSE |
| TpMuguga_02g00245 | TpMuguga_02g00246 | 21 | 0 | TRUE |
| TpMuguga_02g00246 | TpMuguga_02g00245 | 21 | 0 | TRUE |
| TpMuguga_02g02400 | TpMuguga_02g00252 | 24 | 0 | TRUE |
| TpMuguga_02g00252 | TpMuguga_02g02400 | 24 | 0 | TRUE |
| TpMuguga_02g00254 | TpMuguga_02g00253 | 134 | 0 | FALSE |
| TpMuguga_02g00253 | TpMuguga_02g00254 | 134 | 0 | FALSE |
| TpMuguga_02g00256 | TpMuguga_02g00257 | 215 | 0 | FALSE |
| TpMuguga_02g00257 | TpMuguga_02g00256 | 215 | 0 | FALSE |
| TpMuguga_02g00264 | TpMuguga_02g00263 | 28 | 0 | FALSE |
| TpMuguga_02g00263 | TpMuguga_02g00264 | 28 | 0 | FALSE |
| TpMuguga_02g00268 | TpMuguga_02g00269 | 22 | 0 | TRUE |
| TpMuguga_02g00269 | TpMuguga_02g00268 | 22 | 0 | TRUE |
| TpMuguga_02g00281 | TpMuguga_02g00282 | 41 | 0 | FALSE |
| TpMuguga_02g00282 | TpMuguga_02g00281 | 41 | 0 | FALSE |
| TpMuguga_02g00283 | TpMuguga_02g02445 | 68 | 0 | FALSE |
| TpMuguga_02g02445 | TpMuguga_02g00283 | 68 | 0 | FALSE |
| TpMuguga_02g00294 | TpMuguga_02g02130 | 536 | 0 | FALSE |
| TpMuguga_02g02130 | TpMuguga_02g00294 | 536 | 0 | FALSE |
| TpMuguga_02g02130 | TpMuguga_02g00297 | 24 | 0 | FALSE |
| TpMuguga_02g00297 | TpMuguga_02g02130 | 24 | 0 | FALSE |
| TpMuguga_02g00298 | TpMuguga_02g02580 | 681 | 0 | FALSE |
| TpMuguga_02g00299 | TpMuguga_02g02580 | 2 | 0 | FALSE |
| TpMuguga_02g02580 | TpMuguga_02g00298 | 681 | 0 | FALSE |
| TpMuguga_02g02580 | TpMuguga_02g00299 | 2 | 0 | FALSE |
| TpMuguga_02g00300 | TpMuguga_02g00301 | 351 | 0 | FALSE |
| TpMuguga_02g00301 | TpMuguga_02g00300 | 351 | 0 | FALSE |
| TpMuguga_02g00313 | TpMuguga_02g00314 | 7 | 0 | TRUE |
| TpMuguga_02g00314 | TpMuguga_02g00313 | 7 | 0 | TRUE |
| TpMuguga_02g00315 | TpMuguga_02g02670 | 107 | 33 | FALSE |
| TpMuguga_02g02670 | TpMuguga_02g00315 | 107 | 33 | FALSE |
| TpMuguga_02g02045 | TpMuguga_02g00324 | 136 | 0 | FALSE |
| TpMuguga_02g00324 | TpMuguga_02g02045 | 136 | 0 | FALSE |
| TpMuguga_02g02040 | TpMuguga_02g00326 | 12 | 0 | TRUE |
| TpMuguga_02g00326 | TpMuguga_02g02040 | 12 | 0 | TRUE |
| TpMuguga_02g00328 | TpMuguga_02g00329 | 12 | 0 | TRUE |
| TpMuguga_02g00329 | TpMuguga_02g00328 | 12 | 0 | TRUE |
| TpMuguga_02g00332 | TpMuguga_02g00331 | 24 | 0 | TRUE |
| TpMuguga_02g00331 | TpMuguga_02g00332 | 24 | 0 | TRUE |
| TpMuguga_02g00335 | TpMuguga_02g02550 | 80 | 0 | FALSE |
| TpMuguga_02g00336 | TpMuguga_02g02550 | 10 | 0 | FALSE |
| TpMuguga_02g00336 | TpMuguga_02g00337 | 209 | 0 | FALSE |
| TpMuguga_02g02550 | TpMuguga_02g00335 | 80 | 0 | FALSE |
| TpMuguga_02g02550 | TpMuguga_02g00336 | 10 | 0 | FALSE |
| TpMuguga_02g00337 | TpMuguga_02g00336 | 209 | 0 | FALSE |
| TpMuguga_02g00341 | TpMuguga_02g00342 | 52 | 0 | FALSE |
| TpMuguga_02g00343 | TpMuguga_02g00342 | 93 | 0 | FALSE |
| TpMuguga_02g00343 | TpMuguga_02g00344 | 18 | 0 | TRUE |
| TpMuguga_02g00342 | TpMuguga_02g00341 | 52 | 0 | FALSE |
| TpMuguga_02g00342 | TpMuguga_02g00343 | 93 | 0 | FALSE |
| TpMuguga_02g00344 | TpMuguga_02g00343 | 18 | 0 | TRUE |
| TpMuguga_02g02270 | TpMuguga_02g00345 | 190 | 0 | FALSE |
| TpMuguga_02g00345 | TpMuguga_02g02270 | 190 | 0 | FALSE |
| TpMuguga_02g00356 | TpMuguga_02g00355 | 30 | 0 | FALSE |
| TpMuguga_02g00355 | TpMuguga_02g00356 | 30 | 0 | FALSE |
| TpMuguga_02g00358 | TpMuguga_02g00357 | 62 | 0 | FALSE |

| | | | | |
|---|---|---|---|---|
| TpMuguga_02g00358 | TpMuguga_02g00359 | 18 | 0 | FALSE |
| TpMuguga_02g00357 | TpMuguga_02g00358 | 62 | 0 | FALSE |
| TpMuguga_02g00359 | TpMuguga_02g00358 | 18 | 0 | FALSE |
| TpMuguga_02g02160 | TpMuguga_02g00366 | 57 | 0 | FALSE |
| TpMuguga_02g00366 | TpMuguga_02g02160 | 57 | 0 | FALSE |
| TpMuguga_02g00375 | TpMuguga_02g00974 | 9 | 0 | FALSE |
| TpMuguga_02g00974 | TpMuguga_02g00375 | 9 | 0 | FALSE |
| TpMuguga_02g00974 | TpMuguga_02g00376 | 64 | 0 | FALSE |
| TpMuguga_02g00376 | TpMuguga_02g00974 | 64 | 0 | FALSE |
| TpMuguga_02g00380 | TpMuguga_02g00379 | 11 | 0 | FALSE |
| TpMuguga_02g00380 | TpMuguga_02g00965 | 24 | 0 | TRUE |
| TpMuguga_02g00379 | TpMuguga_02g00380 | 11 | 0 | FALSE |
| TpMuguga_02g00965 | TpMuguga_02g00380 | 24 | 0 | TRUE |
| TpMuguga_02g00389 | TpMuguga_02g00388 | 32 | 0 | FALSE |
| TpMuguga_02g00389 | TpMuguga_02g00387 | 118 | 0 | TRUE |
| TpMuguga_02g00388 | TpMuguga_02g00389 | 32 | 0 | FALSE |
| TpMuguga_02g00387 | TpMuguga_02g00389 | 118 | 0 | TRUE |
| TpMuguga_02g00394 | TpMuguga_02g00395 | 206 | 37 | FALSE |
| TpMuguga_02g00395 | TpMuguga_02g00394 | 206 | 37 | FALSE |
| TpMuguga_02g02175 | TpMuguga_02g00402 | 4 | 0 | TRUE |
| TpMuguga_02g00402 | TpMuguga_02g02175 | 4 | 0 | TRUE |
| TpMuguga_02g00402 | TpMuguga_02g00401 | 24 | 0 | TRUE |
| TpMuguga_02g00401 | TpMuguga_02g00402 | 24 | 0 | TRUE |
| TpMuguga_02g00406 | TpMuguga_02g00405 | 23 | 0 | TRUE |
| TpMuguga_02g00405 | TpMuguga_02g00406 | 23 | 0 | TRUE |
| TpMuguga_02g00413 | TpMuguga_02g00414 | 7 | 0 | FALSE |
| TpMuguga_02g00414 | TpMuguga_02g00413 | 7 | 0 | FALSE |
| TpMuguga_02g00421 | TpMuguga_02g00420 | 13 | 0 | TRUE |
| TpMuguga_02g00420 | TpMuguga_02g00421 | 13 | 0 | TRUE |
| TpMuguga_02g00427 | TpMuguga_02g00426 | 47 | 0 | FALSE |
| TpMuguga_02g00426 | TpMuguga_02g00427 | 47 | 0 | FALSE |
| TpMuguga_02g00433 | TpMuguga_02g00434 | 5 | 0 | TRUE |
| TpMuguga_02g00434 | TpMuguga_02g00433 | 5 | 0 | TRUE |
| TpMuguga_02g00435 | TpMuguga_02g00436 | 12 | 0 | TRUE |
| TpMuguga_02g00436 | TpMuguga_02g00435 | 12 | 0 | TRUE |
| TpMuguga_02g02115 | TpMuguga_02g00448 | 18 | 0 | FALSE |
| TpMuguga_02g00448 | TpMuguga_02g02115 | 18 | 0 | FALSE |
| TpMuguga_02g00450 | TpMuguga_02g00451 | 64 | 0 | FALSE |
| TpMuguga_02g00451 | TpMuguga_02g00450 | 64 | 0 | FALSE |
| TpMuguga_02g00463 | TpMuguga_02g02480 | 424 | 0 | FALSE |
| TpMuguga_02g02480 | TpMuguga_02g00463 | 424 | 0 | FALSE |
| TpMuguga_02g00466 | TpMuguga_02g00465 | 37 | 0 | TRUE |
| TpMuguga_02g00466 | TpMuguga_02g00467 | 11 | 0 | TRUE |
| TpMuguga_02g00465 | TpMuguga_02g00466 | 37 | 0 | TRUE |
| TpMuguga_02g00467 | TpMuguga_02g00466 | 11 | 0 | TRUE |
| TpMuguga_02g02390 | TpMuguga_02g00469 | 13 | 0 | TRUE |
| TpMuguga_02g00470 | TpMuguga_02g00471 | 249 | 0 | FALSE |
| TpMuguga_02g00469 | TpMuguga_02g02390 | 13 | 0 | TRUE |
| TpMuguga_02g00472 | TpMuguga_02g00471 | 17 | 0 | TRUE |
| TpMuguga_02g00471 | TpMuguga_02g00470 | 249 | 0 | FALSE |
| TpMuguga_02g00471 | TpMuguga_02g00472 | 17 | 0 | TRUE |
| TpMuguga_02g00480 | TpMuguga_02g00479 | 69 | 0 | FALSE |
| TpMuguga_02g00479 | TpMuguga_02g00480 | 69 | 0 | FALSE |
| TpMuguga_02g00500 | TpMuguga_02g00501 | 147 | 0 | FALSE |
| TpMuguga_02g00501 | TpMuguga_02g00500 | 147 | 0 | FALSE |
| TpMuguga_02g00505 | TpMuguga_02g00504 | 10 | 0 | TRUE |
| TpMuguga_02g00504 | TpMuguga_02g00505 | 10 | 0 | TRUE |
| TpMuguga_02g00512 | TpMuguga_02g00511 | 221 | 0 | FALSE |
| TpMuguga_02g00511 | TpMuguga_02g00512 | 221 | 0 | FALSE |
| TpMuguga_02g00515 | TpMuguga_02g00514 | 27 | 0 | FALSE |
| TpMuguga_02g00514 | TpMuguga_02g00515 | 27 | 0 | FALSE |
| TpMuguga_02g00518 | TpMuguga_02g00519 | 10 | 0 | TRUE |
| TpMuguga_02g00519 | TpMuguga_02g00518 | 10 | 0 | TRUE |
| TpMuguga_02g00528 | TpMuguga_02g00529 | 13 | 0 | FALSE |
| TpMuguga_02g00529 | TpMuguga_02g00528 | 13 | 0 | FALSE |
| TpMuguga_02g00532 | TpMuguga_02g00531 | 219 | 0 | FALSE |
| TpMuguga_02g00531 | TpMuguga_02g00532 | 219 | 0 | FALSE |
| TpMuguga_02g02190 | TpMuguga_02g02370 | 83 | 0 | FALSE |
| TpMuguga_02g02370 | TpMuguga_02g02190 | 83 | 0 | FALSE |
| TpMuguga_02g00540 | TpMuguga_02g00539 | 20 | 0 | FALSE |
| TpMuguga_02g00539 | TpMuguga_02g00540 | 20 | 0 | FALSE |

| | | | | |
|---|---|---|---|---|
| TpMuguga_02g02490 | TpMuguga_02g00547 | 199 | 0 | FALSE |
| TpMuguga_02g00547 | TpMuguga_02g02490 | 199 | 0 | FALSE |
| TpMuguga_02g00556 | TpMuguga_02g00555 | 45 | 0 | FALSE |
| TpMuguga_02g00555 | TpMuguga_02g00556 | 45 | 0 | FALSE |
| TpMuguga_02g00558 | TpMuguga_02g00559 | 48 | 0 | FALSE |
| TpMuguga_02g00559 | TpMuguga_02g00558 | 48 | 0 | FALSE |
| TpMuguga_02g00560 | TpMuguga_02g00561 | 24 | 0 | TRUE |
| TpMuguga_02g00561 | TpMuguga_02g00560 | 24 | 0 | TRUE |
| TpMuguga_02g00574 | TpMuguga_02g00575 | 199 | 0 | FALSE |
| TpMuguga_02g00575 | TpMuguga_02g00574 | 199 | 0 | FALSE |
| TpMuguga_02g00585 | TpMuguga_02g00584 | 11 | 0 | TRUE |
| TpMuguga_02g00584 | TpMuguga_02g00585 | 11 | 0 | TRUE |
| TpMuguga_02g02240 | TpMuguga_02g02450 | 108 | 83 | TRUE |
| TpMuguga_02g02450 | TpMuguga_02g02240 | 108 | 83 | TRUE |
| TpMuguga_02g00594 | TpMuguga_02g00596 | 2 | 0 | FALSE |
| TpMuguga_02g00596 | TpMuguga_02g00594 | 2 | 0 | FALSE |
| TpMuguga_02g00597 | TpMuguga_02g00595 | 104 | 0 | FALSE |
| TpMuguga_02g00595 | TpMuguga_02g00597 | 104 | 0 | FALSE |
| TpMuguga_02g00603 | TpMuguga_02g00604 | 268 | 0 | FALSE |
| TpMuguga_02g00604 | TpMuguga_02g00603 | 268 | 0 | FALSE |
| TpMuguga_02g00609 | TpMuguga_02g00610 | 54 | 0 | TRUE |
| TpMuguga_02g00610 | TpMuguga_02g00609 | 54 | 0 | TRUE |
| TpMuguga_02g00613 | TpMuguga_02g00612 | 22 | 0 | TRUE |
| TpMuguga_02g00612 | TpMuguga_02g00613 | 22 | 0 | TRUE |
| TpMuguga_02g02030 | TpMuguga_02g00614 | 291 | 0 | FALSE |
| TpMuguga_02g00614 | TpMuguga_02g02030 | 291 | 0 | FALSE |
| TpMuguga_02g02010 | TpMuguga_02g02615 | 24 | 0 | TRUE |
| TpMuguga_02g02615 | TpMuguga_02g02010 | 24 | 0 | TRUE |
| TpMuguga_02g00627 | TpMuguga_02g00626 | 14 | 0 | TRUE |
| TpMuguga_02g00626 | TpMuguga_02g00627 | 14 | 0 | TRUE |
| TpMuguga_02g00628 | TpMuguga_02g00629 | 25 | 0 | FALSE |
| TpMuguga_02g00629 | TpMuguga_02g00628 | 25 | 0 | FALSE |
| TpMuguga_02g00647 | TpMuguga_02g00646 | 24 | 0 | TRUE |
| TpMuguga_02g00646 | TpMuguga_02g00647 | 24 | 0 | TRUE |
| TpMuguga_02g00652 | TpMuguga_02g00651 | 2 | 0 | TRUE |
| TpMuguga_02g00651 | TpMuguga_02g00652 | 2 | 0 | TRUE |
| TpMuguga_02g00663 | TpMuguga_02g00664 | 23 | 0 | TRUE |
| TpMuguga_02g00664 | TpMuguga_02g00663 | 23 | 0 | TRUE |
| TpMuguga_02g00681 | TpMuguga_02g00680 | 38 | 0 | FALSE |
| TpMuguga_02g00680 | TpMuguga_02g00681 | 38 | 0 | FALSE |
| TpMuguga_02g00683 | TpMuguga_02g00682 | 70 | 0 | FALSE |
| TpMuguga_02g00682 | TpMuguga_02g00683 | 70 | 0 | FALSE |
| TpMuguga_02g00684 | TpMuguga_02g00685 | 128 | 0 | FALSE |
| TpMuguga_02g00685 | TpMuguga_02g00684 | 128 | 0 | FALSE |
| TpMuguga_02g00690 | TpMuguga_02g02650 | 13 | 0 | FALSE |
| TpMuguga_02g02650 | TpMuguga_02g00690 | 13 | 0 | FALSE |
| TpMuguga_02g02185 | TpMuguga_02g02260 | 6 | 0 | FALSE |
| TpMuguga_02g02260 | TpMuguga_02g02185 | 6 | 0 | FALSE |
| TpMuguga_02g02260 | TpMuguga_02g00696 | 7 | 0 | FALSE |
| TpMuguga_02g00696 | TpMuguga_02g02260 | 7 | 0 | FALSE |
| TpMuguga_02g00698 | TpMuguga_02g00699 | 122 | 20 | FALSE |
| TpMuguga_02g00699 | TpMuguga_02g00698 | 122 | 20 | FALSE |
| TpMuguga_02g02020 | TpMuguga_02g00709 | 33 | 0 | FALSE |
| TpMuguga_02g00709 | TpMuguga_02g02020 | 33 | 0 | FALSE |
| TpMuguga_02g00715 | TpMuguga_02g00716 | 157 | 0 | FALSE |
| TpMuguga_02g00716 | TpMuguga_02g00715 | 157 | 0 | FALSE |
| TpMuguga_02g00722 | TpMuguga_02g00723 | 23 | 0 | TRUE |
| TpMuguga_02g00723 | TpMuguga_02g00722 | 23 | 0 | TRUE |
| TpMuguga_02g00726 | TpMuguga_02g00727 | 17 | 0 | FALSE |
| TpMuguga_02g00727 | TpMuguga_02g00726 | 17 | 0 | FALSE |
| TpMuguga_02g00730 | TpMuguga_02g00732 | 93 | 0 | TRUE |
| TpMuguga_02g00732 | TpMuguga_02g00730 | 93 | 0 | TRUE |
| TpMuguga_02g02090 | TpMuguga_02g00740 | 178 | 0 | FALSE |
| TpMuguga_02g00740 | TpMuguga_02g02090 | 178 | 0 | FALSE |
| TpMuguga_02g00741 | TpMuguga_02g00742 | 298 | 0 | FALSE |
| TpMuguga_02g00742 | TpMuguga_02g00741 | 298 | 0 | FALSE |
| TpMuguga_02g00747 | TpMuguga_02g00745 | 35 | 0 | FALSE |
| TpMuguga_02g00747 | TpMuguga_02g00746 | 68 | 0 | FALSE |
| TpMuguga_02g00745 | TpMuguga_02g00747 | 35 | 0 | FALSE |
| TpMuguga_02g00746 | TpMuguga_02g00747 | 68 | 0 | FALSE |
| TpMuguga_02g00783 | TpMuguga_02g00784 | 164 | 0 | FALSE |

| | | | | |
|---|---|---|---|---|
| TpMuguga_02g00784 | TpMuguga_02g00783 | 164 | 0 | FALSE |
| TpMuguga_02g02320 | TpMuguga_02g00790 | 2 | 0 | TRUE |
| TpMuguga_02g00790 | TpMuguga_02g02320 | 2 | 0 | TRUE |
| TpMuguga_02g02625 | TpMuguga_02g00792 | 24 | 0 | TRUE |
| TpMuguga_02g00792 | TpMuguga_02g02625 | 24 | 0 | TRUE |
| TpMuguga_02g00800 | TpMuguga_02g00799 | 12 | 0 | TRUE |
| TpMuguga_02g00799 | TpMuguga_02g00800 | 12 | 0 | TRUE |
| TpMuguga_02g00836 | TpMuguga_02g00835 | 167 | 0 | FALSE |
| TpMuguga_02g00835 | TpMuguga_02g00836 | 167 | 0 | FALSE |
| TpMuguga_02g00843 | TpMuguga_02g00842 | 37 | 0 | FALSE |
| TpMuguga_02g00843 | TpMuguga_02g00844 | 78 | 0 | FALSE |
| TpMuguga_02g00842 | TpMuguga_02g00843 | 37 | 0 | FALSE |
| TpMuguga_02g00844 | TpMuguga_02g00843 | 78 | 0 | FALSE |
| TpMuguga_02g00844 | TpMuguga_02g02150 | 355 | 0 | FALSE |
| TpMuguga_02g00847 | TpMuguga_02g02150 | 85 | 11 | FALSE |
| TpMuguga_02g02150 | TpMuguga_02g00844 | 355 | 0 | FALSE |
| TpMuguga_02g02150 | TpMuguga_02g00847 | 85 | 11 | FALSE |
| TpMuguga_02g00861 | TpMuguga_02g00860 | 23 | 0 | FALSE |
| TpMuguga_02g00860 | TpMuguga_02g00861 | 23 | 0 | FALSE |
| TpMuguga_02g02145 | TpMuguga_02g00871 | 2 | 0 | TRUE |
| TpMuguga_02g00871 | TpMuguga_02g02145 | 2 | 0 | TRUE |
| TpMuguga_02g00873 | TpMuguga_02g00874 | 39 | 0 | FALSE |
| TpMuguga_02g00874 | TpMuguga_02g00873 | 39 | 0 | FALSE |
| TpMuguga_02g00878 | TpMuguga_02g00879 | 248 | 0 | FALSE |
| TpMuguga_02g00879 | TpMuguga_02g00878 | 248 | 0 | FALSE |
| TpMuguga_02g02085 | TpMuguga_02g00882 | 20 | 0 | FALSE |
| TpMuguga_02g00882 | TpMuguga_02g02085 | 20 | 0 | FALSE |
| TpMuguga_02g00887 | TpMuguga_02g00886 | 10 | 0 | TRUE |
| TpMuguga_02g00886 | TpMuguga_02g00887 | 10 | 0 | TRUE |
| TpMuguga_02g00896 | TpMuguga_02g00897 | 38 | 0 | FALSE |
| TpMuguga_02g00897 | TpMuguga_02g00896 | 38 | 0 | FALSE |
| TpMuguga_02g00900 | TpMuguga_02g00901 | 93 | 0 | FALSE |
| TpMuguga_02g00901 | TpMuguga_02g00900 | 93 | 0 | FALSE |
| TpMuguga_02g00904 | TpMuguga_02g00903 | 7 | 0 | FALSE |
| TpMuguga_02g00904 | TpMuguga_02g00905 | 386 | 0 | TRUE |
| TpMuguga_02g00903 | TpMuguga_02g00904 | 7 | 0 | FALSE |
| TpMuguga_02g00905 | TpMuguga_02g00904 | 386 | 0 | TRUE |
| TpMuguga_02g00938 | TpMuguga_02g00939 | 10 | 0 | FALSE |
| TpMuguga_02g00939 | TpMuguga_02g00938 | 10 | 0 | FALSE |
| TpMuguga_02g00941 | TpMuguga_02g00942 | 26 | 0 | FALSE |
| TpMuguga_02g00942 | TpMuguga_02g00941 | 26 | 0 | FALSE |
| TpMuguga_02g00964 | TpMuguga_02g00945 | 11 | 0 | TRUE |
| TpMuguga_02g00964 | TpMuguga_02g00948 | 177 | 13 | FALSE |
| TpMuguga_02g00945 | TpMuguga_02g00964 | 11 | 0 | TRUE |
| TpMuguga_02g00948 | TpMuguga_02g00964 | 177 | 13 | FALSE |
| TpMuguga_04g02000 | TpMuguga_04g02010 | 1086 | 0 | FALSE |
| TpMuguga_04g02010 | TpMuguga_04g02000 | 1086 | 0 | FALSE |
| TpMuguga_04g02555 | TpMuguga_04g02560 | 8 | 8 | TRUE |
| TpMuguga_04g02560 | TpMuguga_04g02555 | 8 | 8 | TRUE |
| TpMuguga_04g00026 | TpMuguga_04g00027 | 38 | 0 | FALSE |
| TpMuguga_04g00027 | TpMuguga_04g00026 | 38 | 0 | FALSE |
| TpMuguga_04g00032 | TpMuguga_04g00033 | 12 | 0 | FALSE |
| TpMuguga_04g00034 | TpMuguga_04g00033 | 162 | 0 | FALSE |
| TpMuguga_04g00033 | TpMuguga_04g00032 | 12 | 0 | FALSE |
| TpMuguga_04g00033 | TpMuguga_04g00034 | 162 | 0 | FALSE |
| TpMuguga_04g00038 | TpMuguga_04g00039 | 7 | 0 | TRUE |
| TpMuguga_04g00039 | TpMuguga_04g00038 | 7 | 0 | TRUE |
| TpMuguga_04g00056 | TpMuguga_04g00055 | 555 | 0 | FALSE |
| TpMuguga_04g00055 | TpMuguga_04g00056 | 555 | 0 | FALSE |
| TpMuguga_04g00064 | TpMuguga_04g00063 | 33 | 0 | TRUE |
| TpMuguga_04g00063 | TpMuguga_04g00064 | 33 | 0 | TRUE |
| TpMuguga_04g00074 | TpMuguga_04g00075 | 16 | 0 | TRUE |
| TpMuguga_04g00075 | TpMuguga_04g00074 | 16 | 0 | TRUE |
| TpMuguga_04g00089 | TpMuguga_04g02455 | 137 | 4 | FALSE |
| TpMuguga_04g02455 | TpMuguga_04g00089 | 137 | 4 | FALSE |
| TpMuguga_04g02385 | TpMuguga_04g00118 | 21 | 0 | FALSE |
| TpMuguga_04g00118 | TpMuguga_04g02385 | 21 | 0 | FALSE |
| TpMuguga_04g00138 | TpMuguga_04g00139 | 418 | 0 | FALSE |
| TpMuguga_04g00139 | TpMuguga_04g00138 | 418 | 0 | FALSE |
| TpMuguga_04g00167 | TpMuguga_04g02380 | 159 | 0 | FALSE |
| TpMuguga_04g02380 | TpMuguga_04g00167 | 159 | 0 | FALSE |

| | | | | |
|---|---|---|---|---|
| TpMuguga_04g00169 | TpMuguga_04g00168 | 676 | 0 | FALSE |
| TpMuguga_04g00169 | TpMuguga_04g00170 | 11 | 0 | FALSE |
| TpMuguga_04g00168 | TpMuguga_04g00169 | 676 | 0 | FALSE |
| TpMuguga_04g00170 | TpMuguga_04g00169 | 11 | 0 | FALSE |
| TpMuguga_04g00170 | TpMuguga_04g02210 | 120 | 0 | FALSE |
| TpMuguga_04g02210 | TpMuguga_04g00170 | 120 | 0 | FALSE |
| TpMuguga_04g00176 | TpMuguga_04g00177 | 7 | 0 | FALSE |
| TpMuguga_04g00177 | TpMuguga_04g00176 | 7 | 0 | FALSE |
| TpMuguga_04g00187 | TpMuguga_04g00186 | 446 | 0 | FALSE |
| TpMuguga_04g00186 | TpMuguga_04g00187 | 446 | 0 | FALSE |
| TpMuguga_04g00195 | TpMuguga_04g02065 | 28 | 0 | TRUE |
| TpMuguga_04g02065 | TpMuguga_04g00195 | 28 | 0 | TRUE |
| TpMuguga_04g00208 | TpMuguga_04g00207 | 119 | 0 | FALSE |
| TpMuguga_04g00208 | TpMuguga_04g00209 | 13 | 0 | FALSE |
| TpMuguga_04g00207 | TpMuguga_04g00208 | 119 | 0 | FALSE |
| TpMuguga_04g00209 | TpMuguga_04g00208 | 13 | 0 | FALSE |
| TpMuguga_04g00214 | TpMuguga_04g00215 | 181 | 0 | FALSE |
| TpMuguga_04g00215 | TpMuguga_04g00214 | 181 | 0 | FALSE |
| TpMuguga_04g00216 | TpMuguga_04g02175 | 1 | 0 | TRUE |
| TpMuguga_04g00217 | TpMuguga_04g02175 | 74 | 0 | FALSE |
| TpMuguga_04g02175 | TpMuguga_04g00216 | 1 | 0 | TRUE |
| TpMuguga_04g02175 | TpMuguga_04g00217 | 74 | 0 | FALSE |
| TpMuguga_04g02255 | TpMuguga_04g00225 | 459 | 0 | FALSE |
| TpMuguga_04g00225 | TpMuguga_04g02255 | 459 | 0 | FALSE |
| TpMuguga_04g00231 | TpMuguga_04g00230 | 37 | 0 | FALSE |
| TpMuguga_04g00230 | TpMuguga_04g00231 | 37 | 0 | FALSE |
| TpMuguga_04g00234 | TpMuguga_04g00235 | 57 | 25 | FALSE |
| TpMuguga_04g00235 | TpMuguga_04g00234 | 57 | 25 | FALSE |
| TpMuguga_04g00237 | TpMuguga_04g00236 | 237 | 0 | FALSE |
| TpMuguga_04g00236 | TpMuguga_04g00237 | 237 | 0 | FALSE |
| TpMuguga_04g00254 | TpMuguga_04g00253 | 12 | 0 | FALSE |
| TpMuguga_04g00253 | TpMuguga_04g00254 | 12 | 0 | FALSE |
| TpMuguga_04g00257 | TpMuguga_04g00258 | 500 | 0 | FALSE |
| TpMuguga_04g00258 | TpMuguga_04g00257 | 500 | 0 | FALSE |
| TpMuguga_04g00264 | TpMuguga_04g00265 | 138 | 0 | FALSE |
| TpMuguga_04g00265 | TpMuguga_04g00264 | 138 | 0 | FALSE |
| TpMuguga_04g00268 | TpMuguga_04g00269 | 34 | 0 | FALSE |
| TpMuguga_04g00269 | TpMuguga_04g00268 | 34 | 0 | FALSE |
| TpMuguga_04g00271 | TpMuguga_04g00270 | 396 | 0 | TRUE |
| TpMuguga_04g00270 | TpMuguga_04g00271 | 396 | 0 | TRUE |
| TpMuguga_04g00290 | TpMuguga_04g00291 | 1356 | 0 | FALSE |
| TpMuguga_04g00291 | TpMuguga_04g00290 | 1356 | 0 | FALSE |
| TpMuguga_04g00293 | TpMuguga_04g00292 | 142 | 0 | FALSE |
| TpMuguga_04g00292 | TpMuguga_04g00293 | 142 | 0 | FALSE |
| TpMuguga_04g00298 | TpMuguga_04g02200 | 221 | 0 | FALSE |
| TpMuguga_04g02200 | TpMuguga_04g00298 | 221 | 0 | FALSE |
| TpMuguga_04g00303 | TpMuguga_04g00304 | 8 | 0 | FALSE |
| TpMuguga_04g00304 | TpMuguga_04g00303 | 8 | 0 | FALSE |
| TpMuguga_04g02550 | TpMuguga_04g00307 | 37 | 0 | TRUE |
| TpMuguga_04g00307 | TpMuguga_04g02550 | 37 | 0 | TRUE |
| TpMuguga_04g00321 | TpMuguga_04g00322 | 95 | 0 | FALSE |
| TpMuguga_04g00322 | TpMuguga_04g00321 | 95 | 0 | FALSE |
| TpMuguga_04g00325 | TpMuguga_04g00326 | 194 | 0 | FALSE |
| TpMuguga_04g00326 | TpMuguga_04g00325 | 194 | 0 | FALSE |
| TpMuguga_04g00329 | TpMuguga_04g02715 | 30 | 0 | FALSE |
| TpMuguga_04g02195 | TpMuguga_04g02715 | 20 | 20 | TRUE |
| TpMuguga_04g02715 | TpMuguga_04g00329 | 30 | 0 | FALSE |
| TpMuguga_04g02715 | TpMuguga_04g02195 | 20 | 20 | TRUE |
| TpMuguga_04g02045 | TpMuguga_04g02240 | 1 | 1 | TRUE |
| TpMuguga_04g02240 | TpMuguga_04g02045 | 1 | 1 | TRUE |
| TpMuguga_04g00341 | TpMuguga_04g00340 | 67 | 0 | FALSE |
| TpMuguga_04g00340 | TpMuguga_04g00341 | 67 | 0 | FALSE |
| TpMuguga_04g00345 | TpMuguga_04g00346 | 1 | 0 | FALSE |
| TpMuguga_04g00346 | TpMuguga_04g00345 | 1 | 0 | FALSE |
| TpMuguga_04g00349 | TpMuguga_04g00350 | 167 | 0 | FALSE |
| TpMuguga_04g00350 | TpMuguga_04g00349 | 167 | 0 | FALSE |
| TpMuguga_04g02060 | TpMuguga_04g00354 | 16 | 0 | TRUE |
| TpMuguga_04g00354 | TpMuguga_04g02060 | 16 | 0 | TRUE |
| TpMuguga_04g00361 | TpMuguga_04g00360 | 22 | 0 | FALSE |
| TpMuguga_04g00360 | TpMuguga_04g00361 | 22 | 0 | FALSE |
| TpMuguga_04g00365 | TpMuguga_04g00366 | 25 | 0 | FALSE |

| | | | | |
|---|---|---|---|---|
| TpMuguga_04g00366 | TpMuguga_04g00365 | 25 | 0 | FALSE |
| TpMuguga_04g02085 | TpMuguga_04g00369 | 345 | 0 | TRUE |
| TpMuguga_04g00369 | TpMuguga_04g02085 | 345 | 0 | TRUE |
| TpMuguga_04g00375 | TpMuguga_04g02675 | 100 | 0 | FALSE |
| TpMuguga_04g02675 | TpMuguga_04g00375 | 100 | 0 | FALSE |
| TpMuguga_04g00386 | TpMuguga_04g00385 | 40 | 0 | FALSE |
| TpMuguga_04g00385 | TpMuguga_04g00386 | 40 | 0 | FALSE |
| TpMuguga_04g00401 | TpMuguga_04g02050 | 515 | 29 | TRUE |
| TpMuguga_04g02050 | TpMuguga_04g00401 | 515 | 29 | TRUE |
| TpMuguga_04g00415 | TpMuguga_04g00416 | 78 | 0 | FALSE |
| TpMuguga_04g00416 | TpMuguga_04g00415 | 78 | 0 | FALSE |
| TpMuguga_04g00429 | TpMuguga_04g00428 | 507 | 0 | FALSE |
| TpMuguga_04g00428 | TpMuguga_04g00429 | 507 | 0 | FALSE |
| TpMuguga_04g00439 | TpMuguga_04g00437 | 106 | 0 | FALSE |
| TpMuguga_04g00437 | TpMuguga_04g00439 | 106 | 0 | FALSE |
| TpMuguga_04g00443 | TpMuguga_04g00444 | 410 | 9 | FALSE |
| TpMuguga_04g00444 | TpMuguga_04g00443 | 410 | 9 | FALSE |
| TpMuguga_04g00447 | TpMuguga_04g02575 | 193 | 0 | FALSE |
| TpMuguga_04g02575 | TpMuguga_04g00447 | 193 | 0 | FALSE |
| TpMuguga_04g02505 | TpMuguga_04g02510 | 1 | 1 | TRUE |
| TpMuguga_04g02510 | TpMuguga_04g02505 | 1 | 1 | TRUE |
| TpMuguga_04g00475 | TpMuguga_04g00476 | 84 | 0 | FALSE |
| TpMuguga_04g00476 | TpMuguga_04g00475 | 84 | 0 | FALSE |
| TpMuguga_04g00483 | TpMuguga_04g00484 | 24 | 0 | TRUE |
| TpMuguga_04g00484 | TpMuguga_04g00483 | 24 | 0 | TRUE |
| TpMuguga_04g00485 | TpMuguga_04g00937 | 2 | 0 | FALSE |
| TpMuguga_04g00937 | TpMuguga_04g00485 | 2 | 0 | FALSE |
| TpMuguga_04g02215 | TpMuguga_04g00491 | 83 | 0 | TRUE |
| TpMuguga_04g00492 | TpMuguga_04g00491 | 9 | 0 | FALSE |
| TpMuguga_04g00491 | TpMuguga_04g02215 | 83 | 0 | TRUE |
| TpMuguga_04g00491 | TpMuguga_04g00492 | 9 | 0 | FALSE |
| TpMuguga_04g02225 | TpMuguga_04g00501 | 128 | 8 | FALSE |
| TpMuguga_04g00501 | TpMuguga_04g02225 | 128 | 8 | FALSE |
| TpMuguga_04g00514 | TpMuguga_04g00513 | 13 | 0 | FALSE |
| TpMuguga_04g00514 | TpMuguga_04g02410 | 30 | 0 | FALSE |
| TpMuguga_04g00513 | TpMuguga_04g00514 | 13 | 0 | FALSE |
| TpMuguga_04g00517 | TpMuguga_04g02410 | 158 | 0 | FALSE |
| TpMuguga_04g02410 | TpMuguga_04g00514 | 30 | 0 | FALSE |
| TpMuguga_04g02410 | TpMuguga_04g00517 | 158 | 0 | FALSE |
| TpMuguga_04g00527 | TpMuguga_04g00528 | 1 | 0 | TRUE |
| TpMuguga_04g00529 | TpMuguga_04g00528 | 11 | 0 | FALSE |
| TpMuguga_04g00528 | TpMuguga_04g00527 | 1 | 0 | TRUE |
| TpMuguga_04g00528 | TpMuguga_04g00529 | 11 | 0 | FALSE |
| TpMuguga_04g00538 | TpMuguga_04g00539 | 35 | 0 | FALSE |
| TpMuguga_04g00539 | TpMuguga_04g00538 | 35 | 0 | FALSE |
| TpMuguga_04g00543 | TpMuguga_04g00544 | 10 | 0 | FALSE |
| TpMuguga_04g00544 | TpMuguga_04g00543 | 10 | 0 | FALSE |
| TpMuguga_04g00547 | TpMuguga_04g00546 | 166 | 0 | FALSE |
| TpMuguga_04g00546 | TpMuguga_04g00547 | 166 | 0 | FALSE |
| TpMuguga_04g00550 | TpMuguga_04g00549 | 37 | 0 | FALSE |
| TpMuguga_04g00549 | TpMuguga_04g00550 | 37 | 0 | FALSE |
| TpMuguga_04g02365 | TpMuguga_04g00553 | 39 | 0 | FALSE |
| TpMuguga_04g00553 | TpMuguga_04g02365 | 39 | 0 | FALSE |
| TpMuguga_04g00557 | TpMuguga_04g00558 | 80 | 0 | FALSE |
| TpMuguga_04g00558 | TpMuguga_04g00557 | 80 | 0 | FALSE |
| TpMuguga_04g00558 | TpMuguga_04g00559 | 21 | 0 | TRUE |
| TpMuguga_04g00559 | TpMuguga_04g00558 | 21 | 0 | TRUE |
| TpMuguga_04g00563 | TpMuguga_04g02710 | 14 | 0 | FALSE |
| TpMuguga_04g00566 | TpMuguga_04g02710 | 18 | 0 | TRUE |
| TpMuguga_04g02710 | TpMuguga_04g00563 | 14 | 0 | FALSE |
| TpMuguga_04g02710 | TpMuguga_04g00566 | 18 | 0 | TRUE |
| TpMuguga_04g00572 | TpMuguga_04g00573 | 24 | 0 | TRUE |
| TpMuguga_04g00573 | TpMuguga_04g00572 | 24 | 0 | TRUE |
| TpMuguga_04g00577 | TpMuguga_04g00576 | 449 | 0 | FALSE |
| TpMuguga_04g00577 | TpMuguga_04g00578 | 40 | 0 | FALSE |
| TpMuguga_04g00576 | TpMuguga_04g00577 | 449 | 0 | FALSE |
| TpMuguga_04g00578 | TpMuguga_04g00577 | 40 | 0 | FALSE |
| TpMuguga_04g00578 | TpMuguga_04g00579 | 103 | 0 | FALSE |
| TpMuguga_04g00579 | TpMuguga_04g00578 | 103 | 0 | FALSE |
| TpMuguga_04g00579 | TpMuguga_04g00580 | 72 | 0 | FALSE |
| TpMuguga_04g00580 | TpMuguga_04g00579 | 72 | 0 | FALSE |

| | | | | |
|---|---|---|---|---|
| TpMuguga_04g00580 | TpMuguga_04g00581 | 122 | 0 | TRUE |
| TpMuguga_04g00581 | TpMuguga_04g00580 | 122 | 0 | TRUE |
| TpMuguga_04g02130 | TpMuguga_04g02270 | 171 | 0 | FALSE |
| TpMuguga_04g02270 | TpMuguga_04g02130 | 171 | 0 | FALSE |
| TpMuguga_04g00593 | TpMuguga_04g00594 | 19 | 0 | FALSE |
| TpMuguga_04g00594 | TpMuguga_04g00593 | 19 | 0 | FALSE |
| TpMuguga_04g00611 | TpMuguga_04g00610 | 740 | 29 | FALSE |
| TpMuguga_04g00610 | TpMuguga_04g00611 | 740 | 29 | FALSE |
| TpMuguga_04g00616 | TpMuguga_04g00615 | 43 | 0 | TRUE |
| TpMuguga_04g00615 | TpMuguga_04g00616 | 43 | 0 | TRUE |
| TpMuguga_04g02075 | TpMuguga_04g00623 | 171 | 0 | FALSE |
| TpMuguga_04g00623 | TpMuguga_04g02075 | 171 | 0 | FALSE |
| TpMuguga_04g00623 | TpMuguga_04g00624 | 36 | 0 | TRUE |
| TpMuguga_04g00624 | TpMuguga_04g00623 | 36 | 0 | TRUE |
| TpMuguga_04g00629 | TpMuguga_04g02545 | 53 | 0 | TRUE |
| TpMuguga_04g02545 | TpMuguga_04g00629 | 53 | 0 | TRUE |
| TpMuguga_04g00634 | TpMuguga_04g00633 | 16 | 0 | TRUE |
| TpMuguga_04g00633 | TpMuguga_04g00634 | 16 | 0 | TRUE |
| TpMuguga_04g00636 | TpMuguga_04g00637 | 11 | 0 | FALSE |
| TpMuguga_04g00637 | TpMuguga_04g00636 | 11 | 0 | FALSE |
| TpMuguga_04g00637 | TpMuguga_04g00638 | 298 | 0 | FALSE |
| TpMuguga_04g00639 | TpMuguga_04g00638 | 28 | 0 | FALSE |
| TpMuguga_04g00638 | TpMuguga_04g00637 | 298 | 0 | FALSE |
| TpMuguga_04g00638 | TpMuguga_04g00639 | 28 | 0 | FALSE |
| TpMuguga_04g00641 | TpMuguga_04g00640 | 978 | 0 | FALSE |
| TpMuguga_04g00640 | TpMuguga_04g00641 | 978 | 0 | FALSE |
| TpMuguga_04g02090 | TpMuguga_04g02330 | 137 | 0 | FALSE |
| TpMuguga_04g02330 | TpMuguga_04g02090 | 137 | 0 | FALSE |
| TpMuguga_04g00671 | TpMuguga_04g00670 | 6 | 0 | TRUE |
| TpMuguga_04g00671 | TpMuguga_04g00672 | 24 | 0 | TRUE |
| TpMuguga_04g00670 | TpMuguga_04g00671 | 6 | 0 | TRUE |
| TpMuguga_04g02160 | TpMuguga_04g00672 | 81 | 0 | TRUE |
| TpMuguga_04g00672 | TpMuguga_04g00671 | 24 | 0 | TRUE |
| TpMuguga_04g00672 | TpMuguga_04g02160 | 81 | 0 | TRUE |
| TpMuguga_04g00682 | TpMuguga_04g00681 | 21 | 0 | TRUE |
| TpMuguga_04g00681 | TpMuguga_04g00682 | 21 | 0 | TRUE |
| TpMuguga_04g00688 | TpMuguga_04g00687 | 16 | 0 | TRUE |
| TpMuguga_04g00687 | TpMuguga_04g00688 | 16 | 0 | TRUE |
| TpMuguga_04g00691 | TpMuguga_04g02620 | 49 | 0 | FALSE |
| TpMuguga_04g02620 | TpMuguga_04g00691 | 49 | 0 | FALSE |
| TpMuguga_04g02250 | TpMuguga_04g00693 | 7 | 0 | FALSE |
| TpMuguga_04g00693 | TpMuguga_04g02250 | 7 | 0 | FALSE |
| TpMuguga_04g00713 | TpMuguga_04g00712 | 54 | 0 | FALSE |
| TpMuguga_04g00712 | TpMuguga_04g00713 | 54 | 0 | FALSE |
| TpMuguga_04g00714 | TpMuguga_04g00715 | 231 | 0 | FALSE |
| TpMuguga_04g00715 | TpMuguga_04g00714 | 231 | 0 | FALSE |
| TpMuguga_04g00718 | TpMuguga_04g00717 | 39 | 0 | FALSE |
| TpMuguga_04g00717 | TpMuguga_04g00718 | 39 | 0 | FALSE |
| TpMuguga_04g00717 | TpMuguga_04g00716 | 27 | 0 | FALSE |
| TpMuguga_04g00716 | TpMuguga_04g00717 | 27 | 0 | FALSE |
| TpMuguga_04g00720 | TpMuguga_04g00722 | 20 | 4 | FALSE |
| TpMuguga_04g00722 | TpMuguga_04g00720 | 20 | 4 | FALSE |
| TpMuguga_04g00724 | TpMuguga_04g00725 | 283 | 0 | FALSE |
| TpMuguga_04g00725 | TpMuguga_04g00724 | 283 | 0 | FALSE |
| TpMuguga_04g00727 | TpMuguga_04g00726 | 57 | 0 | FALSE |
| TpMuguga_04g00727 | TpMuguga_04g00728 | 63 | 0 | FALSE |
| TpMuguga_04g00726 | TpMuguga_04g00727 | 57 | 0 | FALSE |
| TpMuguga_04g00728 | TpMuguga_04g00727 | 63 | 0 | FALSE |
| TpMuguga_04g00732 | TpMuguga_04g00731 | 47 | 0 | FALSE |
| TpMuguga_04g00731 | TpMuguga_04g00732 | 47 | 0 | FALSE |
| TpMuguga_04g00734 | TpMuguga_04g00735 | 104 | 0 | FALSE |
| TpMuguga_04g00735 | TpMuguga_04g00734 | 104 | 0 | FALSE |
| TpMuguga_04g00737 | TpMuguga_04g00736 | 195 | 0 | FALSE |
| TpMuguga_04g00737 | TpMuguga_04g00738 | 1 | 0 | TRUE |
| TpMuguga_04g00736 | TpMuguga_04g00737 | 195 | 0 | FALSE |
| TpMuguga_04g00738 | TpMuguga_04g00737 | 1 | 0 | TRUE |
| TpMuguga_04g00746 | TpMuguga_04g02665 | 140 | 0 | FALSE |
| TpMuguga_04g02665 | TpMuguga_04g00746 | 140 | 0 | FALSE |
| TpMuguga_04g00759 | TpMuguga_04g00758 | 485 | 4 | FALSE |
| TpMuguga_04g00758 | TpMuguga_04g00759 | 485 | 4 | FALSE |
| TpMuguga_04g00779 | TpMuguga_04g02460 | 235 | 0 | FALSE |

| | | | | |
|---|---|---|---|---|
| TpMuguga_04g02460 | TpMuguga_04g00779 | 235 | 0 | FALSE |
| TpMuguga_04g00780 | TpMuguga_04g00781 | 1335 | 1335 | TRUE |
| TpMuguga_04g02110 | TpMuguga_04g00781 | 28 | 0 | FALSE |
| TpMuguga_04g02110 | TpMuguga_04g00941 | 6 | 0 | TRUE |
| TpMuguga_04g00781 | TpMuguga_04g00780 | 1335 | 1335 | TRUE |
| TpMuguga_04g00781 | TpMuguga_04g02110 | 28 | 0 | FALSE |
| TpMuguga_04g00781 | TpMuguga_04g00941 | 72 | 0 | FALSE |
| TpMuguga_04g00941 | TpMuguga_04g02110 | 6 | 0 | TRUE |
| TpMuguga_04g00941 | TpMuguga_04g00781 | 72 | 0 | FALSE |
| TpMuguga_04g00786 | TpMuguga_04g00785 | 8 | 0 | FALSE |
| TpMuguga_04g00785 | TpMuguga_04g00786 | 8 | 0 | FALSE |
| TpMuguga_04g00790 | TpMuguga_04g00789 | 18 | 0 | TRUE |
| TpMuguga_04g00789 | TpMuguga_04g00790 | 18 | 0 | TRUE |
| TpMuguga_04g02610 | TpMuguga_04g00926 | 20 | 0 | TRUE |
| TpMuguga_04g00926 | TpMuguga_04g02610 | 20 | 0 | TRUE |
| TpMuguga_04g00794 | TpMuguga_04g00795 | 34 | 0 | FALSE |
| TpMuguga_04g00795 | TpMuguga_04g00794 | 34 | 0 | FALSE |
| TpMuguga_04g00798 | TpMuguga_04g00799 | 18 | 0 | FALSE |
| TpMuguga_04g00799 | TpMuguga_04g00798 | 18 | 0 | FALSE |
| TpMuguga_04g00801 | TpMuguga_04g00802 | 24 | 0 | TRUE |
| TpMuguga_04g00802 | TpMuguga_04g00801 | 24 | 0 | TRUE |
| TpMuguga_04g00808 | TpMuguga_04g00809 | 298 | 4 | FALSE |
| TpMuguga_04g00809 | TpMuguga_04g00808 | 298 | 4 | FALSE |
| TpMuguga_04g02125 | TpMuguga_04g00815 | 218 | 0 | FALSE |
| TpMuguga_04g00815 | TpMuguga_04g02125 | 218 | 0 | FALSE |
| TpMuguga_04g00817 | TpMuguga_04g00818 | 3 | 0 | FALSE |
| TpMuguga_04g00818 | TpMuguga_04g00817 | 3 | 0 | FALSE |
| TpMuguga_04g00819 | TpMuguga_04g00820 | 671 | 0 | FALSE |
| TpMuguga_04g00820 | TpMuguga_04g00819 | 671 | 0 | FALSE |
| TpMuguga_04g00822 | TpMuguga_04g00823 | 29 | 0 | TRUE |
| TpMuguga_04g00823 | TpMuguga_04g00822 | 29 | 0 | TRUE |
| TpMuguga_04g00827 | TpMuguga_04g00826 | 115 | 0 | FALSE |
| TpMuguga_04g00826 | TpMuguga_04g00827 | 115 | 0 | FALSE |
| TpMuguga_04g00830 | TpMuguga_04g00829 | 50 | 0 | FALSE |
| TpMuguga_04g00830 | TpMuguga_04g00831 | 19 | 0 | TRUE |
| TpMuguga_04g00829 | TpMuguga_04g00830 | 50 | 0 | FALSE |
| TpMuguga_04g00832 | TpMuguga_04g00831 | 37 | 0 | TRUE |
| TpMuguga_04g00831 | TpMuguga_04g00830 | 19 | 0 | TRUE |
| TpMuguga_04g00831 | TpMuguga_04g00832 | 37 | 0 | TRUE |
| TpMuguga_04g00942 | TpMuguga_04g00834 | 73 | 0 | FALSE |
| TpMuguga_04g00834 | TpMuguga_04g00942 | 73 | 0 | FALSE |
| TpMuguga_04g00838 | TpMuguga_04g00837 | 4 | 0 | TRUE |
| TpMuguga_04g00837 | TpMuguga_04g00838 | 4 | 0 | TRUE |
| TpMuguga_04g00847 | TpMuguga_04g00846 | 144 | 0 | FALSE |
| TpMuguga_04g00846 | TpMuguga_04g00847 | 144 | 0 | FALSE |
| TpMuguga_04g00849 | TpMuguga_04g00848 | 12 | 0 | FALSE |
| TpMuguga_04g00848 | TpMuguga_04g00849 | 12 | 0 | FALSE |
| TpMuguga_04g00851 | TpMuguga_04g00850 | 48 | 0 | FALSE |
| TpMuguga_04g00850 | TpMuguga_04g00851 | 48 | 0 | FALSE |
| TpMuguga_04g00854 | TpMuguga_04g02585 | 49 | 0 | FALSE |
| TpMuguga_04g00854 | TpMuguga_04g00855 | 10 | 0 | TRUE |
| TpMuguga_04g02585 | TpMuguga_04g00854 | 49 | 0 | FALSE |
| TpMuguga_04g00855 | TpMuguga_04g00854 | 10 | 0 | TRUE |
| TpMuguga_04g02265 | TpMuguga_04g00859 | 13 | 0 | TRUE |
| TpMuguga_04g02265 | TpMuguga_04g00862 | 4 | 0 | FALSE |
| TpMuguga_04g00859 | TpMuguga_04g02265 | 13 | 0 | TRUE |
| TpMuguga_04g00862 | TpMuguga_04g02265 | 4 | 0 | FALSE |
| TpMuguga_04g02340 | TpMuguga_04g02415 | 10 | 0 | TRUE |
| TpMuguga_04g02340 | TpMuguga_04g00866 | 92 | 0 | FALSE |
| TpMuguga_04g02415 | TpMuguga_04g02340 | 10 | 0 | TRUE |
| TpMuguga_04g00866 | TpMuguga_04g02340 | 92 | 0 | FALSE |
| TpMuguga_04g00867 | TpMuguga_04g00868 | 169 | 0 | FALSE |
| TpMuguga_04g00868 | TpMuguga_04g00867 | 169 | 0 | FALSE |
| TpMuguga_04g00872 | TpMuguga_04g00871 | 18 | 0 | TRUE |
| TpMuguga_04g00871 | TpMuguga_04g00872 | 18 | 0 | TRUE |
| TpMuguga_04g02320 | TpMuguga_04g02435 | 16 | 11 | TRUE |
| TpMuguga_04g02435 | TpMuguga_04g02320 | 16 | 11 | TRUE |
| TpMuguga_04g00880 | TpMuguga_04g00879 | 83 | 0 | FALSE |
| TpMuguga_04g00879 | TpMuguga_04g00880 | 83 | 0 | FALSE |
| TpMuguga_04g02420 | TpMuguga_04g00901 | 51 | 0 | FALSE |
| TpMuguga_04g00901 | TpMuguga_04g02420 | 51 | 0 | FALSE |

| | | | | |
|---|---|---|---|---|
| TpMuguga_03g00012 | TpMuguga_03g00011 | 134 | 134 | FALSE |
| TpMuguga_03g00011 | TpMuguga_03g00012 | 134 | 134 | FALSE |
| TpMuguga_03g00015 | TpMuguga_03g02460 | 53 | 0 | FALSE |
| TpMuguga_03g02460 | TpMuguga_03g00015 | 53 | 0 | FALSE |
| TpMuguga_03g02460 | TpMuguga_03g02465 | 1 | 1 | TRUE |
| TpMuguga_03g02465 | TpMuguga_03g02460 | 1 | 1 | TRUE |
| TpMuguga_03g00020 | TpMuguga_03g00019 | 27 | 0 | FALSE |
| TpMuguga_03g00019 | TpMuguga_03g00020 | 27 | 0 | FALSE |
| TpMuguga_03g00023 | TpMuguga_03g00024 | 185 | 0 | FALSE |
| TpMuguga_03g00024 | TpMuguga_03g00023 | 185 | 0 | FALSE |
| TpMuguga_03g02215 | TpMuguga_03g00065 | 29 | 0 | FALSE |
| TpMuguga_03g02285 | TpMuguga_03g00065 | 294 | 0 | FALSE |
| TpMuguga_03g00065 | TpMuguga_03g02215 | 29 | 0 | FALSE |
| TpMuguga_03g00065 | TpMuguga_03g02285 | 294 | 0 | FALSE |
| TpMuguga_03g00072 | TpMuguga_03g00071 | 14 | 0 | FALSE |
| TpMuguga_03g00071 | TpMuguga_03g00072 | 14 | 0 | FALSE |
| TpMuguga_03g00076 | TpMuguga_03g00077 | 24 | 0 | TRUE |
| TpMuguga_03g00077 | TpMuguga_03g00076 | 24 | 0 | TRUE |
| TpMuguga_03g00083 | TpMuguga_03g00082 | 16 | 0 | TRUE |
| TpMuguga_03g00083 | TpMuguga_03g00084 | 7 | 0 | FALSE |
| TpMuguga_03g00082 | TpMuguga_03g00083 | 16 | 0 | TRUE |
| TpMuguga_03g00084 | TpMuguga_03g00083 | 7 | 0 | FALSE |
| TpMuguga_03g00090 | TpMuguga_03g00092 | 43 | 0 | FALSE |
| TpMuguga_03g00092 | TpMuguga_03g00090 | 43 | 0 | FALSE |
| TpMuguga_03g02315 | TpMuguga_03g00098 | 24 | 0 | TRUE |
| TpMuguga_03g00098 | TpMuguga_03g02315 | 24 | 0 | TRUE |
| TpMuguga_03g00111 | TpMuguga_03g00110 | 24 | 0 | TRUE |
| TpMuguga_03g00110 | TpMuguga_03g00111 | 24 | 0 | TRUE |
| TpMuguga_03g00110 | TpMuguga_03g00109 | 91 | 0 | FALSE |
| TpMuguga_03g00109 | TpMuguga_03g00110 | 91 | 0 | FALSE |
| TpMuguga_03g02495 | TpMuguga_03g00115 | 170 | 0 | FALSE |
| TpMuguga_03g00115 | TpMuguga_03g02495 | 170 | 0 | FALSE |
| TpMuguga_03g00118 | TpMuguga_03g02565 | 7 | 0 | FALSE |
| TpMuguga_03g02565 | TpMuguga_03g00118 | 7 | 0 | FALSE |
| TpMuguga_03g00125 | TpMuguga_03g00126 | 98 | 0 | FALSE |
| TpMuguga_03g00126 | TpMuguga_03g00125 | 98 | 0 | FALSE |
| TpMuguga_03g00127 | TpMuguga_03g00128 | 50 | 21 | FALSE |
| TpMuguga_03g00128 | TpMuguga_03g00127 | 50 | 21 | FALSE |
| TpMuguga_03g00155 | TpMuguga_03g00154 | 210 | 10 | FALSE |
| TpMuguga_03g00155 | TpMuguga_03g02685 | 79 | 0 | FALSE |
| TpMuguga_03g00154 | TpMuguga_03g00155 | 210 | 10 | FALSE |
| TpMuguga_03g02685 | TpMuguga_03g00155 | 79 | 0 | FALSE |
| TpMuguga_03g00161 | TpMuguga_03g00162 | 160 | 0 | FALSE |
| TpMuguga_03g00162 | TpMuguga_03g00161 | 160 | 0 | FALSE |
| TpMuguga_03g00162 | TpMuguga_03g00163 | 249 | 0 | FALSE |
| TpMuguga_03g00163 | TpMuguga_03g00162 | 249 | 0 | FALSE |
| TpMuguga_03g00179 | TpMuguga_03g02605 | 1 | 0 | TRUE |
| TpMuguga_03g02605 | TpMuguga_03g00179 | 1 | 0 | TRUE |
| TpMuguga_03g02545 | TpMuguga_03g00200 | 172 | 0 | FALSE |
| TpMuguga_03g00200 | TpMuguga_03g02545 | 172 | 0 | FALSE |
| TpMuguga_03g00208 | TpMuguga_03g00207 | 7 | 0 | TRUE |
| TpMuguga_03g00207 | TpMuguga_03g00208 | 7 | 0 | TRUE |
| TpMuguga_03g00214 | TpMuguga_03g00213 | 20 | 0 | FALSE |
| TpMuguga_03g00213 | TpMuguga_03g00214 | 20 | 0 | FALSE |
| TpMuguga_03g00217 | TpMuguga_03g00216 | 25 | 0 | FALSE |
| TpMuguga_03g00216 | TpMuguga_03g00217 | 25 | 0 | FALSE |
| TpMuguga_03g00225 | TpMuguga_03g00226 | 330 | 0 | FALSE |
| TpMuguga_03g00226 | TpMuguga_03g00225 | 330 | 0 | FALSE |
| TpMuguga_03g02300 | TpMuguga_03g00237 | 7 | 0 | TRUE |
| TpMuguga_03g00237 | TpMuguga_03g02300 | 7 | 0 | TRUE |
| TpMuguga_03g00246 | TpMuguga_03g00245 | 410 | 0 | FALSE |
| TpMuguga_03g00245 | TpMuguga_03g00246 | 410 | 0 | FALSE |
| TpMuguga_03g00250 | TpMuguga_03g00249 | 21 | 0 | FALSE |
| TpMuguga_03g00249 | TpMuguga_03g00250 | 21 | 0 | FALSE |
| TpMuguga_03g00254 | TpMuguga_03g02370 | 5 | 0 | FALSE |
| TpMuguga_03g00257 | TpMuguga_03g02370 | 24 | 0 | TRUE |
| TpMuguga_03g02370 | TpMuguga_03g00254 | 5 | 0 | FALSE |
| TpMuguga_03g02370 | TpMuguga_03g00257 | 24 | 0 | TRUE |
| TpMuguga_03g00270 | TpMuguga_03g02310 | 4 | 4 | TRUE |
| TpMuguga_03g02310 | TpMuguga_03g00270 | 4 | 4 | TRUE |
| TpMuguga_03g00278 | TpMuguga_03g00279 | 57 | 0 | FALSE |

| | | | | |
|---|---|---|---|---|
| TpMuguga_03g00279 | TpMuguga_03g00278 | 57 | 0 | FALSE |
| TpMuguga_03g00290 | TpMuguga_03g00291 | 180 | 0 | TRUE |
| TpMuguga_03g00291 | TpMuguga_03g00290 | 180 | 0 | TRUE |
| TpMuguga_03g02555 | TpMuguga_03g00308 | 124 | 0 | FALSE |
| TpMuguga_03g00308 | TpMuguga_03g02555 | 124 | 0 | FALSE |
| TpMuguga_03g02190 | TpMuguga_03g00314 | 10 | 0 | TRUE |
| TpMuguga_03g00314 | TpMuguga_03g02190 | 10 | 0 | TRUE |
| TpMuguga_03g02265 | TpMuguga_03g02270 | 19 | 0 | TRUE |
| TpMuguga_03g02270 | TpMuguga_03g02265 | 19 | 0 | TRUE |
| TpMuguga_03g02245 | TpMuguga_03g00320 | 22 | 0 | TRUE |
| TpMuguga_03g00320 | TpMuguga_03g02245 | 22 | 0 | TRUE |
| TpMuguga_03g00334 | TpMuguga_03g02680 | 44 | 0 | FALSE |
| TpMuguga_03g02680 | TpMuguga_03g00334 | 44 | 0 | FALSE |
| TpMuguga_03g00362 | TpMuguga_03g00361 | 3 | 0 | FALSE |
| TpMuguga_03g00361 | TpMuguga_03g00362 | 3 | 0 | FALSE |
| TpMuguga_03g00384 | TpMuguga_03g00382 | 4 | 0 | FALSE |
| TpMuguga_03g00382 | TpMuguga_03g00384 | 4 | 0 | FALSE |
| TpMuguga_03g00389 | TpMuguga_03g00390 | 8 | 0 | FALSE |
| TpMuguga_03g00390 | TpMuguga_03g00389 | 8 | 0 | FALSE |
| TpMuguga_03g00413 | TpMuguga_03g00412 | 304 | 4 | FALSE |
| TpMuguga_03g00412 | TpMuguga_03g00413 | 304 | 4 | FALSE |
| TpMuguga_03g00420 | TpMuguga_03g00418 | 370 | 0 | FALSE |
| TpMuguga_03g00418 | TpMuguga_03g00420 | 370 | 0 | FALSE |
| TpMuguga_03g00444 | TpMuguga_03g00445 | 23 | 0 | FALSE |
| TpMuguga_03g00445 | TpMuguga_03g00444 | 23 | 0 | FALSE |
| TpMuguga_03g00451 | TpMuguga_03g00452 | 246 | 0 | TRUE |
| TpMuguga_03g00452 | TpMuguga_03g00451 | 246 | 0 | TRUE |
| TpMuguga_03g00456 | TpMuguga_03g00457 | 38 | 0 | TRUE |
| TpMuguga_03g00457 | TpMuguga_03g00456 | 38 | 0 | TRUE |
| TpMuguga_03g00472 | TpMuguga_03g00473 | 142 | 0 | FALSE |
| TpMuguga_03g00473 | TpMuguga_03g00472 | 142 | 0 | FALSE |
| TpMuguga_03g00484 | TpMuguga_03g00483 | 247 | 0 | FALSE |
| TpMuguga_03g00483 | TpMuguga_03g00484 | 247 | 0 | FALSE |
| TpMuguga_03g00487 | TpMuguga_03g00488 | 228 | 0 | FALSE |
| TpMuguga_03g00488 | TpMuguga_03g00487 | 228 | 0 | FALSE |
| TpMuguga_03g00488 | TpMuguga_03g00489 | 19 | 0 | TRUE |
| TpMuguga_03g00489 | TpMuguga_03g00488 | 19 | 0 | TRUE |
| TpMuguga_03g00489 | TpMuguga_03g00493 | 106 | 0 | FALSE |
| TpMuguga_03g00493 | TpMuguga_03g00489 | 106 | 0 | FALSE |
| TpMuguga_03g00496 | TpMuguga_03g00495 | 1184 | 0 | FALSE |
| TpMuguga_03g00495 | TpMuguga_03g00496 | 1184 | 0 | FALSE |
| TpMuguga_03g00504 | TpMuguga_03g00503 | 81 | 0 | FALSE |
| TpMuguga_03g00503 | TpMuguga_03g00504 | 81 | 0 | FALSE |
| TpMuguga_03g00508 | TpMuguga_03g00507 | 129 | 0 | FALSE |
| TpMuguga_03g00507 | TpMuguga_03g00508 | 129 | 0 | FALSE |
| TpMuguga_03g00515 | TpMuguga_03g00514 | 190 | 0 | TRUE |
| TpMuguga_03g00514 | TpMuguga_03g00515 | 190 | 0 | TRUE |
| TpMuguga_03g02480 | TpMuguga_03g00519 | 24 | 0 | TRUE |
| TpMuguga_03g00519 | TpMuguga_03g02480 | 24 | 0 | TRUE |
| TpMuguga_03g00523 | TpMuguga_03g02210 | 154 | 0 | FALSE |
| TpMuguga_03g02210 | TpMuguga_03g00523 | 154 | 0 | FALSE |
| TpMuguga_03g02210 | TpMuguga_03g00522 | 276 | 0 | FALSE |
| TpMuguga_03g00522 | TpMuguga_03g02210 | 276 | 0 | FALSE |
| TpMuguga_03g00525 | TpMuguga_03g00524 | 58 | 0 | FALSE |
| TpMuguga_03g00525 | TpMuguga_03g02585 | 93 | 0 | FALSE |
| TpMuguga_03g00524 | TpMuguga_03g00525 | 58 | 0 | FALSE |
| TpMuguga_03g02585 | TpMuguga_03g00525 | 93 | 0 | FALSE |
| TpMuguga_03g00531 | TpMuguga_03g00532 | 164 | 0 | FALSE |
| TpMuguga_03g00532 | TpMuguga_03g00531 | 164 | 0 | FALSE |
| TpMuguga_03g00552 | TpMuguga_03g00551 | 354 | 0 | FALSE |
| TpMuguga_03g00551 | TpMuguga_03g00552 | 354 | 0 | FALSE |
| TpMuguga_03g00554 | TpMuguga_03g00553 | 402 | 0 | FALSE |
| TpMuguga_03g00554 | TpMuguga_03g02220 | 14 | 0 | TRUE |
| TpMuguga_03g00553 | TpMuguga_03g00554 | 402 | 0 | FALSE |
| TpMuguga_03g02220 | TpMuguga_03g00554 | 14 | 0 | TRUE |
| TpMuguga_03g02185 | TpMuguga_03g00565 | 915 | 221 | FALSE |
| TpMuguga_03g00565 | TpMuguga_03g02185 | 915 | 221 | FALSE |
| TpMuguga_03g00567 | TpMuguga_03g00566 | 493 | 0 | FALSE |
| TpMuguga_03g00567 | TpMuguga_03g02660 | 1 | 1 | TRUE |
| TpMuguga_03g00566 | TpMuguga_03g00567 | 493 | 0 | FALSE |
| TpMuguga_03g02660 | TpMuguga_03g00567 | 1 | 1 | TRUE |

| | | | | |
|---|---|---|---|---|
| TpMuguga_03g00570 | TpMuguga_03g00571 | 96 | 0 | FALSE |
| TpMuguga_03g00571 | TpMuguga_03g00570 | 96 | 0 | FALSE |
| TpMuguga_03g00577 | TpMuguga_03g00576 | 84 | 0 | FALSE |
| TpMuguga_03g00577 | TpMuguga_03g00578 | 578 | 0 | FALSE |
| TpMuguga_03g00576 | TpMuguga_03g00577 | 84 | 0 | FALSE |
| TpMuguga_03g00578 | TpMuguga_03g00577 | 578 | 0 | FALSE |
| TpMuguga_03g02630 | TpMuguga_03g00607 | 243 | 4 | FALSE |
| TpMuguga_03g00607 | TpMuguga_03g02630 | 243 | 4 | FALSE |
| TpMuguga_03g00627 | TpMuguga_03g02100 | 222 | 0 | FALSE |
| TpMuguga_03g02100 | TpMuguga_03g00627 | 222 | 0 | FALSE |
| TpMuguga_03g02115 | TpMuguga_03g00629 | 80 | 11 | FALSE |
| TpMuguga_03g00629 | TpMuguga_03g02115 | 80 | 11 | FALSE |
| TpMuguga_03g00641 | TpMuguga_03g00642 | 39 | 0 | FALSE |
| TpMuguga_03g00642 | TpMuguga_03g00641 | 39 | 0 | FALSE |
| TpMuguga_03g00644 | TpMuguga_03g00643 | 344 | 0 | FALSE |
| TpMuguga_03g00643 | TpMuguga_03g00644 | 344 | 0 | FALSE |
| TpMuguga_03g00652 | TpMuguga_03g00653 | 682 | 0 | FALSE |
| TpMuguga_03g00653 | TpMuguga_03g00652 | 682 | 0 | FALSE |
| TpMuguga_03g00678 | TpMuguga_03g00679 | 18 | 0 | TRUE |
| TpMuguga_03g00679 | TpMuguga_03g00678 | 18 | 0 | TRUE |
| TpMuguga_03g00684 | TpMuguga_03g02105 | 23 | 0 | FALSE |
| TpMuguga_03g02105 | TpMuguga_03g00684 | 23 | 0 | FALSE |
| TpMuguga_03g00691 | TpMuguga_03g00692 | 14 | 0 | TRUE |
| TpMuguga_03g00692 | TpMuguga_03g00691 | 14 | 0 | TRUE |
| TpMuguga_03g02045 | TpMuguga_03g00701 | 247 | 0 | FALSE |
| TpMuguga_03g00701 | TpMuguga_03g02045 | 247 | 0 | FALSE |
| TpMuguga_03g00712 | TpMuguga_03g00711 | 316 | 0 | FALSE |
| TpMuguga_03g00711 | TpMuguga_03g00712 | 316 | 0 | FALSE |
| TpMuguga_03g00714 | TpMuguga_03g00715 | 16 | 0 | TRUE |
| TpMuguga_03g00715 | TpMuguga_03g00714 | 16 | 0 | TRUE |
| TpMuguga_03g00728 | TpMuguga_03g00727 | 23 | 0 | TRUE |
| TpMuguga_03g00727 | TpMuguga_03g00728 | 23 | 0 | TRUE |
| TpMuguga_03g00733 | TpMuguga_03g00732 | 235 | 4 | FALSE |
| TpMuguga_03g00733 | TpMuguga_03g00734 | 7 | 0 | FALSE |
| TpMuguga_03g00732 | TpMuguga_03g00733 | 235 | 4 | FALSE |
| TpMuguga_03g00736 | TpMuguga_03g00735 | 19 | 0 | TRUE |
| TpMuguga_03g00735 | TpMuguga_03g00736 | 19 | 0 | TRUE |
| TpMuguga_03g00734 | TpMuguga_03g00733 | 7 | 0 | FALSE |
| TpMuguga_03g00756 | TpMuguga_03g00757 | 15 | 0 | TRUE |
| TpMuguga_03g00757 | TpMuguga_03g00756 | 15 | 0 | TRUE |
| TpMuguga_03g00792 | TpMuguga_03g00793 | 204 | 0 | FALSE |
| TpMuguga_03g00794 | TpMuguga_03g00795 | 99 | 0 | FALSE |
| TpMuguga_03g00793 | TpMuguga_03g00792 | 204 | 0 | FALSE |
| TpMuguga_03g00795 | TpMuguga_03g00794 | 99 | 0 | FALSE |
| TpMuguga_03g00798 | TpMuguga_03g02160 | 1125 | 0 | FALSE |
| TpMuguga_03g02160 | TpMuguga_03g00798 | 1125 | 0 | FALSE |
| TpMuguga_03g00801 | TpMuguga_03g00802 | 33 | 0 | TRUE |
| TpMuguga_03g02025 | TpMuguga_03g00802 | 140 | 0 | FALSE |
| TpMuguga_03g02025 | TpMuguga_03g00803 | 867 | 371 | FALSE |
| TpMuguga_03g00802 | TpMuguga_03g00801 | 33 | 0 | TRUE |
| TpMuguga_03g00802 | TpMuguga_03g02025 | 140 | 0 | FALSE |
| TpMuguga_03g00803 | TpMuguga_03g02025 | 867 | 371 | FALSE |
| TpMuguga_03g00809 | TpMuguga_03g00808 | 24 | 0 | TRUE |
| TpMuguga_03g00808 | TpMuguga_03g00809 | 24 | 0 | TRUE |
| TpMuguga_03g00811 | TpMuguga_03g00812 | 593 | 0 | FALSE |
| TpMuguga_03g00812 | TpMuguga_03g00811 | 593 | 0 | FALSE |
| TpMuguga_03g00817 | TpMuguga_03g00816 | 30 | 0 | TRUE |
| TpMuguga_03g00817 | TpMuguga_03g00818 | 36 | 0 | FALSE |
| TpMuguga_03g00816 | TpMuguga_03g00817 | 30 | 0 | TRUE |
| TpMuguga_03g00818 | TpMuguga_03g00817 | 36 | 0 | FALSE |
| TpMuguga_03g00833 | TpMuguga_03g00832 | 218 | 4 | FALSE |
| TpMuguga_03g00832 | TpMuguga_03g00833 | 218 | 4 | FALSE |
| TpMuguga_03g02000 | TpMuguga_03g00841 | 40 | 0 | FALSE |
| TpMuguga_03g02000 | TpMuguga_03g00842 | 93 | 9 | FALSE |
| TpMuguga_03g00841 | TpMuguga_03g02000 | 40 | 0 | FALSE |
| TpMuguga_03g00842 | TpMuguga_03g02000 | 93 | 9 | FALSE |
| TpMuguga_03g00846 | TpMuguga_03g02135 | 24 | 0 | TRUE |
| TpMuguga_03g02135 | TpMuguga_03g00846 | 24 | 0 | TRUE |
| TpMuguga_03g00854 | TpMuguga_03g00853 | 5 | 0 | FALSE |
| TpMuguga_03g00853 | TpMuguga_03g00854 | 5 | 0 | FALSE |
| TpMuguga_03g00940 | TpMuguga_03g00857 | 60 | 0 | TRUE |

| | | | | |
|---|---|---|---|---|
| TpMuguga_03g00858 | TpMuguga_03g00857 | 466 | 0 | FALSE |
| TpMuguga_03g00857 | TpMuguga_03g00940 | 60 | 0 | TRUE |
| TpMuguga_03g00857 | TpMuguga_03g00858 | 466 | 0 | FALSE |
| TpMuguga_05g00004 | TpMuguga_05g00003 | 17 | 17 | TRUE |
| TpMuguga_05g00004 | TpMuguga_05g00005 | 8 | 8 | TRUE |
| TpMuguga_05g00003 | TpMuguga_05g00004 | 17 | 17 | TRUE |
| TpMuguga_05g00005 | TpMuguga_05g00004 | 8 | 8 | TRUE |
| TpMuguga_05g00006 | TpMuguga_05g00007 | 11 | 11 | TRUE |
| TpMuguga_05g00007 | TpMuguga_05g00006 | 11 | 11 | TRUE |
| TpMuguga_05g00007 | TpMuguga_05g00008 | 8 | 8 | TRUE |
| TpMuguga_05g00008 | TpMuguga_05g00007 | 8 | 8 | TRUE |
| TpMuguga_05g00011 | TpMuguga_05g00010 | 1 | 1 | TRUE |
| TpMuguga_05g00011 | TpMuguga_05g00012 | 53 | 53 | TRUE |
| TpMuguga_05g00010 | TpMuguga_05g00011 | 1 | 1 | TRUE |
| TpMuguga_05g00013 | TpMuguga_05g00012 | 1 | 1 | TRUE |
| TpMuguga_05g00012 | TpMuguga_05g00011 | 53 | 53 | TRUE |
| TpMuguga_05g00012 | TpMuguga_05g00013 | 1 | 1 | TRUE |
| TpMuguga_05g00015 | TpMuguga_05g00014 | 35 | 35 | TRUE |
| TpMuguga_05g00015 | TpMuguga_05g00016 | 4 | 4 | TRUE |
| TpMuguga_05g00014 | TpMuguga_05g00015 | 35 | 35 | TRUE |
| TpMuguga_05g00016 | TpMuguga_05g00015 | 4 | 4 | TRUE |
| TpMuguga_05g00018 | TpMuguga_05g00017 | 35 | 35 | TRUE |
| TpMuguga_05g00017 | TpMuguga_05g00018 | 35 | 35 | TRUE |
| TpMuguga_05g00024 | TpMuguga_05g00023 | 16 | 16 | TRUE |
| TpMuguga_05g00023 | TpMuguga_05g00024 | 16 | 16 | TRUE |
| TpMuguga_05g00026 | TpMuguga_05g00025 | 8 | 8 | TRUE |
| TpMuguga_05g00025 | TpMuguga_05g00026 | 8 | 8 | TRUE |
