## Additional Data 3 for "Re-annotation of the *Theileria parva* genome refines 53% of the proteome and uncovers essential components of N-glycosylation, a conserved pathway in many organisms"

**Additional file 3: Ratio of intron_coverage by average_CDS_coverage, for introns with read_coverage>0**

| Gene_LocusTag | Intron_num | RatioPerIntronExonCov |
|---|---|---|
| TpMuguga_01g00007 | 1 | 0.027586207 |
| TpMuguga_01g00012 | 1 | 0 |
| TpMuguga_01g00012 | 2 | 0 |
| TpMuguga_01g00012 | 3 | 0 |
| TpMuguga_01g00013 | 1 | 0 |
| TpMuguga_01g00013 | 2 | 0 |
| TpMuguga_01g00013 | 3 | 0 |
| TpMuguga_01g00013 | 4 | 0 |
| TpMuguga_01g00013 | 5 | 0 |
| TpMuguga_01g00013 | 6 | 0 |
| TpMuguga_01g00013 | 7 | 0 |
| TpMuguga_01g00014 | 1 | 0 |
| TpMuguga_01g00014 | 2 | 0 |
| TpMuguga_01g00015 | 1 | 0 |
| TpMuguga_01g00015 | 2 | 5.333333333 |
| TpMuguga_01g00015 | 3 | 0 |
| TpMuguga_01g00015 | 4 | 0 |
| TpMuguga_01g00015 | 5 | 0.666666667 |
| TpMuguga_01g00015 | 6 | 0 |
| TpMuguga_01g00015 | 7 | 0 |
| TpMuguga_01g00016 | 1 | 3 |
| TpMuguga_01g00016 | 2 | 0 |
| TpMuguga_01g00017 | 1 | 6.642857143 |
| TpMuguga_01g00017 | 2 | 0.428571429 |
| TpMuguga_01g00017 | 3 | 0 |
| TpMuguga_01g00017 | 4 | 0 |
| TpMuguga_01g00017 | 5 | 0 |
| TpMuguga_01g00017 | 6 | 0 |
| TpMuguga_01g00017 | 7 | 0 |
| TpMuguga_01g00017 | 8 | 0 |
| TpMuguga_01g00017 | 9 | 0.171428571 |
| TpMuguga_01g00017 | 10 | 0 |
| TpMuguga_01g00017 | 11 | 0 |
| TpMuguga_01g00018 | 1 | 2.666666667 |
| TpMuguga_01g00019 | 1 | 0 |
| TpMuguga_01g00020 | 1 | 0 |

| | | |
|---|---|---|
| TpMuguga_01g00020 | 2 | 2.25 |
| TpMuguga_01g02140 | 1 | 0 |
| TpMuguga_01g02140 | 2 | 0 |
| TpMuguga_01g02140 | 3 | 0 |
| TpMuguga_01g02140 | 4 | 0 |
| TpMuguga_01g02140 | 5 | 0 |
| TpMuguga_01g00023 | 1 | 0.074074074 |
| TpMuguga_01g00023 | 2 | 3.333333333 |
| TpMuguga_01g00024 | 1 | 0 |
| TpMuguga_01g02185 | 1 | 3.076923077 |
| TpMuguga_01g02180 | 1 | 0 |
| TpMuguga_01g02190 | 1 | 0 |
| TpMuguga_01g02190 | 2 | 0 |
| TpMuguga_01g00028 | 1 | 4.780487805 |
| TpMuguga_01g00028 | 2 | 1.62195122 |
| TpMuguga_01g00028 | 3 | 0.597560976 |
| TpMuguga_01g00028 | 4 | 0.426829268 |
| TpMuguga_01g00028 | 5 | 1.024390244 |
| TpMuguga_01g00028 | 6 | 1.365853659 |
| TpMuguga_01g00030 | 1 | 1.8 |
| TpMuguga_01g00030 | 2 | 0 |
| TpMuguga_01g00030 | 3 | 2.04 |
| TpMuguga_01g00030 | 4 | 0.96 |
| TpMuguga_01g00030 | 5 | 1.08 |
| TpMuguga_01g00031 | 1 | 0.136546185 |
| TpMuguga_01g02620 | 1 | 0 |
| TpMuguga_01g02620 | 2 | 0.581395349 |
| TpMuguga_01g02620 | 3 | 0.465116279 |
| TpMuguga_01g02620 | 4 | 0 |
| TpMuguga_01g00035 | 1 | 0 |
| TpMuguga_01g00035 | 2 | 0 |
| TpMuguga_01g00036 | 1 | 9.308411215 |
| TpMuguga_01g00036 | 2 | 0 |
| TpMuguga_01g00036 | 3 | 0 |
| TpMuguga_01g00036 | 4 | 0 |
| TpMuguga_01g00036 | 5 | 0 |
| TpMuguga_01g00036 | 6 | 0 |
| TpMuguga_01g00036 | 7 | 0 |
| TpMuguga_01g00036 | 8 | 0 |
| TpMuguga_01g00036 | 9 | 0 |
| TpMuguga_01g00036 | 10 | 0 |
| TpMuguga_01g00036 | 11 | 0 |

| | | |
|---|---|---|
| TpMuguga_01g00037 | 1 | 1.777777778 |
| TpMuguga_01g00037 | 2 | 0 |
| TpMuguga_01g00037 | 3 | 0 |
| TpMuguga_01g00040 | 1 | 0 |
| TpMuguga_01g00040 | 2 | 0 |
| TpMuguga_01g00039 | 1 | 0 |
| TpMuguga_01g00039 | 2 | 0 |
| TpMuguga_01g00039 | 3 | 0 |
| TpMuguga_01g00042 | 1 | 0.008069939 |
| TpMuguga_01g00042 | 2 | 0.010087424 |
| TpMuguga_01g00041 | 1 | 0.318181818 |
| TpMuguga_01g00041 | 2 | 0 |
| TpMuguga_01g00041 | 3 | 1.590909091 |
| TpMuguga_01g00041 | 4 | 0 |
| TpMuguga_01g00041 | 5 | 1.272727273 |
| TpMuguga_01g00041 | 6 | 0 |
| TpMuguga_01g00044 | 1 | 0 |
| TpMuguga_01g00044 | 2 | 0 |
| TpMuguga_01g00043 | 1 | 0 |
| TpMuguga_01g00046 | 1 | 10 |
| TpMuguga_01g00048 | 1 | 0 |
| TpMuguga_01g00048 | 2 | 0 |
| TpMuguga_01g00048 | 3 | 0 |
| TpMuguga_01g00048 | 4 | 0 |
| TpMuguga_01g00048 | 5 | 0 |
| TpMuguga_01g00048 | 6 | 0 |
| TpMuguga_01g00048 | 7 | 0 |
| TpMuguga_01g00048 | 8 | 27.5 |
| TpMuguga_01g00048 | 9 | 0 |
| TpMuguga_01g00048 | 10 | 0 |
| TpMuguga_01g00047 | 1 | 0 |
| TpMuguga_01g00047 | 2 | 0 |
| TpMuguga_01g00047 | 3 | 0 |
| TpMuguga_01g00047 | 4 | 0 |
| TpMuguga_01g00047 | 5 | 0 |
| TpMuguga_01g00047 | 6 | 0 |
| TpMuguga_01g00047 | 7 | 0 |
| TpMuguga_01g00047 | 8 | 0 |
| TpMuguga_01g00047 | 9 | 0 |
| TpMuguga_01g02170 | 1 | 1.224043716 |
| TpMuguga_01g02170 | 2 | 0 |
| TpMuguga_01g02170 | 3 | 0 |

| | | |
|---|---|---|
| TpMuguga_01g02170 | 4 | 1.836065574 |
| TpMuguga_01g02170 | 5 | 0 |
| TpMuguga_01g02170 | 6 | 0 |
| TpMuguga_01g02170 | 7 | 0 |
| TpMuguga_01g02170 | 8 | 0 |
| TpMuguga_01g02170 | 9 | 0.918032787 |
| TpMuguga_01g02170 | 10 | 0.306010929 |
| TpMuguga_01g02170 | 11 | 0.459016393 |
| TpMuguga_01g02170 | 12 | 0.153005464 |
| TpMuguga_01g02170 | 13 | 0.153005464 |
| TpMuguga_01g02605 | 1 | 0 |
| TpMuguga_01g02605 | 2 | 0 |
| TpMuguga_01g02605 | 3 | 0 |
| TpMuguga_01g02605 | 4 | 0 |
| TpMuguga_01g02605 | 5 | 0 |
| TpMuguga_01g00055 | 1 | 0 |
| TpMuguga_01g00055 | 2 | 0 |
| TpMuguga_01g00055 | 3 | 0 |
| TpMuguga_01g00055 | 4 | 0 |
| TpMuguga_01g00055 | 5 | 0 |
| TpMuguga_01g00055 | 6 | 0 |
| TpMuguga_01g00055 | 7 | 0 |
| TpMuguga_01g00055 | 8 | 0 |
| TpMuguga_01g00055 | 9 | 0 |
| TpMuguga_01g00055 | 10 | 0 |
| TpMuguga_01g00054 | 1 | 4.666666667 |
| TpMuguga_01g00054 | 2 | 0 |
| TpMuguga_01g00054 | 3 | 37.33333333 |
| TpMuguga_01g00054 | 4 | 0 |
| TpMuguga_01g00054 | 5 | 0 |
| TpMuguga_01g00054 | 6 | 0 |
| TpMuguga_01g00054 | 7 | 0 |
| TpMuguga_01g00054 | 8 | 0 |
| TpMuguga_01g00054 | 9 | 0 |
| TpMuguga_01g00054 | 10 | 0 |
| TpMuguga_01g00054 | 11 | 0 |
| TpMuguga_01g00054 | 12 | 0 |
| TpMuguga_01g00054 | 13 | 0 |
| TpMuguga_01g00057 | 1 | 12 |
| TpMuguga_01g00057 | 2 | 4 |
| TpMuguga_01g00057 | 3 | 0 |
| TpMuguga_01g00060 | 1 | 0.916666667 |

| | | |
|---|---|---|
| TpMuguga_01g00061 | 1 | 0 |
| TpMuguga_01g00061 | 2 | 1.217391304 |
| TpMuguga_01g00061 | 3 | 0 |
| TpMuguga_01g00061 | 4 | 0 |
| TpMuguga_01g00061 | 5 | 0 |
| TpMuguga_01g00061 | 6 | 0 |
| TpMuguga_01g00062 | 1 | 2 |
| TpMuguga_01g00064 | 1 | 0 |
| TpMuguga_01g00064 | 2 | 0 |
| TpMuguga_01g00064 | 3 | 6 |
| TpMuguga_01g00064 | 4 | 0 |
| TpMuguga_01g00064 | 5 | 0 |
| TpMuguga_01g00063 | 1 | 0 |
| TpMuguga_01g00063 | 2 | 0 |
| TpMuguga_01g00063 | 3 | 0 |
| TpMuguga_01g00063 | 4 | 0 |
| TpMuguga_01g00063 | 5 | 0 |
| TpMuguga_01g00063 | 6 | 0 |
| TpMuguga_01g00063 | 7 | 0 |
| TpMuguga_01g00063 | 8 | 0 |
| TpMuguga_01g00065 | 1 | 0 |
| TpMuguga_01g00065 | 2 | 0 |
| TpMuguga_01g00065 | 3 | 0 |
| TpMuguga_01g00065 | 4 | 0 |
| TpMuguga_01g00065 | 5 | 0 |
| TpMuguga_01g00065 | 6 | 0 |
| TpMuguga_01g00065 | 7 | 0 |
| TpMuguga_01g00065 | 8 | 13 |
| TpMuguga_01g00065 | 9 | 0 |
| TpMuguga_01g00065 | 10 | 0 |
| TpMuguga_01g00065 | 11 | 0 |
| TpMuguga_01g00065 | 12 | 0 |
| TpMuguga_01g00066 | 1 | 0 |
| TpMuguga_01g02310 | 1 | 0 |
| TpMuguga_01g02310 | 2 | 0 |
| TpMuguga_01g02310 | 3 | 0 |
| TpMuguga_01g02310 | 4 | 0 |
| TpMuguga_01g02310 | 5 | 0 |
| TpMuguga_01g02310 | 6 | 0 |
| TpMuguga_01g02310 | 7 | 0 |
| TpMuguga_01g02310 | 8 | 0 |
| TpMuguga_01g02310 | 9 | 0 |

| | | |
|---|---|---|
| TpMuguga_01g02310 | 10 | 12 |
| TpMuguga_01g02310 | 11 | 0 |
| TpMuguga_01g00071 | 1 | 1.006681514 |
| TpMuguga_01g00071 | 2 | 0 |
| TpMuguga_01g00071 | 3 | 0 |
| TpMuguga_01g00072 | 1 | 0 |
| TpMuguga_01g00072 | 2 | 0 |
| TpMuguga_01g00073 | 1 | 0 |
| TpMuguga_01g00073 | 2 | 4.363636364 |
| TpMuguga_01g00075 | 1 | 0.025210084 |
| TpMuguga_01g00075 | 2 | 0.100840336 |
| TpMuguga_01g00076 | 1 | 0 |
| TpMuguga_01g00076 | 2 | 0 |
| TpMuguga_01g00076 | 3 | 0 |
| TpMuguga_01g00076 | 4 | 0 |
| TpMuguga_01g00076 | 5 | 0 |
| TpMuguga_01g00076 | 6 | 0 |
| TpMuguga_01g00077 | 1 | 0.004395604 |
| TpMuguga_01g00077 | 2 | 0 |
| TpMuguga_01g00077 | 3 | 0.145054945 |
| TpMuguga_01g00078 | 1 | 0.316455696 |
| TpMuguga_01g00078 | 2 | 0 |
| TpMuguga_01g00078 | 3 | 0 |
| TpMuguga_01g00078 | 4 | 0 |
| TpMuguga_01g00079 | 1 | 2.028571429 |
| TpMuguga_01g00080 | 1 | 1 |
| TpMuguga_01g00080 | 2 | 9.5 |
| TpMuguga_01g00081 | 1 | 0 |
| TpMuguga_01g00082 | 1 | 1.555555556 |
| TpMuguga_01g00082 | 2 | 0 |
| TpMuguga_01g00082 | 3 | 0.444444444 |
| TpMuguga_01g00083 | 1 | 0.796278159 |
| TpMuguga_01g00083 | 2 | 0.96963761 |
| TpMuguga_01g00084 | 1 | 0 |
| TpMuguga_01g00084 | 2 | 0 |
| TpMuguga_01g00084 | 3 | 0 |
| TpMuguga_01g00086 | 1 | 0.883227176 |
| TpMuguga_01g00086 | 2 | 0.067940552 |
| TpMuguga_01g00086 | 3 | 0.042462845 |
| TpMuguga_01g00085 | 1 | 0 |
| TpMuguga_01g00085 | 2 | 0 |
| TpMuguga_01g00085 | 3 | 0.166666667 |

| | | |
|---|---|---|
| TpMuguga_01g00085 | 4 | 0.5 |
| TpMuguga_01g00085 | 5 | 0.833333333 |
| TpMuguga_01g00087 | 1 | 0 |
| TpMuguga_01g00087 | 2 | 0 |
| TpMuguga_01g00087 | 3 | 0 |
| TpMuguga_01g00087 | 4 | 0 |
| TpMuguga_01g02570 | 1 | 0.098360656 |
| TpMuguga_01g00091 | 1 | 0 |
| TpMuguga_01g00091 | 2 | 0 |
| TpMuguga_01g00091 | 3 | 0 |
| TpMuguga_01g00091 | 4 | 0 |
| TpMuguga_01g00091 | 5 | 0 |
| TpMuguga_01g00091 | 6 | 0 |
| TpMuguga_01g00091 | 7 | 0 |
| TpMuguga_01g00091 | 8 | 0 |
| TpMuguga_01g00091 | 9 | 0 |
| TpMuguga_01g00092 | 1 | 0 |
| TpMuguga_01g00092 | 2 | 0 |
| TpMuguga_01g00092 | 3 | 0 |
| TpMuguga_01g00092 | 4 | 0 |
| TpMuguga_01g00092 | 5 | 0 |
| TpMuguga_01g00096 | 1 | 0.694444444 |
| TpMuguga_01g00096 | 2 | 0 |
| TpMuguga_01g00096 | 3 | 0 |
| TpMuguga_01g00096 | 4 | 0 |
| TpMuguga_01g00097 | 1 | 0 |
| TpMuguga_01g00097 | 2 | 7 |
| TpMuguga_01g00097 | 3 | 0 |
| TpMuguga_01g00097 | 4 | 0 |
| TpMuguga_01g00097 | 5 | 0 |
| TpMuguga_01g00097 | 6 | 1 |
| TpMuguga_01g00097 | 7 | 0 |
| TpMuguga_01g00099 | 1 | 0.523809524 |
| TpMuguga_01g00099 | 2 | 0 |
| TpMuguga_01g00099 | 3 | 0 |
| TpMuguga_01g00099 | 4 | 3.142857143 |
| TpMuguga_01g00099 | 5 | 0 |
| TpMuguga_01g00099 | 6 | 0 |
| TpMuguga_01g00099 | 7 | 0 |
| TpMuguga_01g00099 | 8 | 0 |
| TpMuguga_01g00099 | 9 | 0 |
| TpMuguga_01g00099 | 10 | 0 |

| | | |
|---|---|---|
| TpMuguga_01g00102 | 1 | 0 |
| TpMuguga_01g00102 | 2 | 0 |
| TpMuguga_01g00102 | 3 | 0 |
| TpMuguga_01g00103 | 1 | 0 |
| TpMuguga_01g00103 | 2 | 0 |
| TpMuguga_01g00103 | 3 | 1.129032258 |
| TpMuguga_01g00103 | 4 | 0.677419355 |
| TpMuguga_01g00103 | 5 | 3.612903226 |
| TpMuguga_01g00103 | 6 | 0 |
| TpMuguga_01g00104 | 1 | 0 |
| TpMuguga_01g00104 | 2 | 0 |
| TpMuguga_01g00104 | 3 | 0 |
| TpMuguga_01g00104 | 4 | 0 |
| TpMuguga_01g00105 | 1 | 0 |
| TpMuguga_01g00105 | 2 | 0 |
| TpMuguga_01g00105 | 3 | 1.714285714 |
| TpMuguga_01g00105 | 4 | 5.142857143 |
| TpMuguga_01g00105 | 5 | 3.428571429 |
| TpMuguga_01g00106 | 1 | 0 |
| TpMuguga_01g00106 | 2 | 0 |
| TpMuguga_01g00107 | 1 | 0 |
| TpMuguga_01g00107 | 2 | 0 |
| TpMuguga_01g00107 | 3 | 0 |
| TpMuguga_01g00107 | 4 | 0 |
| TpMuguga_01g00107 | 5 | 2.255555556 |
| TpMuguga_01g00107 | 6 | 10.96666667 |
| TpMuguga_01g02240 | 1 | 0 |
| TpMuguga_01g02240 | 2 | 0 |
| TpMuguga_01g02240 | 3 | 0 |
| TpMuguga_01g02240 | 4 | 0.55 |
| TpMuguga_01g02240 | 5 | 0 |
| TpMuguga_01g02240 | 6 | 0 |
| TpMuguga_01g02240 | 7 | 0 |
| TpMuguga_01g02240 | 8 | 0 |
| TpMuguga_01g02240 | 9 | 0 |
| TpMuguga_01g02240 | 10 | 0 |
| TpMuguga_01g00109 | 1 | 0 |
| TpMuguga_01g00109 | 2 | 8.75 |
| TpMuguga_01g00109 | 3 | 0 |
| TpMuguga_01g00109 | 4 | 0 |
| TpMuguga_01g00109 | 5 | 0 |
| TpMuguga_01g00109 | 6 | 0 |

| | | |
|---|---|---|
| TpMuguga_01g00108 | 1 | 0 |
| TpMuguga_01g00108 | 2 | 0 |
| TpMuguga_01g00108 | 3 | 0 |
| TpMuguga_01g00110 | 1 | 0 |
| TpMuguga_01g00113 | 1 | 0 |
| TpMuguga_01g00113 | 2 | 0 |
| TpMuguga_01g00113 | 3 | 5.133333333 |
| TpMuguga_01g00113 | 4 | 8.4 |
| TpMuguga_01g00113 | 5 | 0 |
| TpMuguga_01g00113 | 6 | 0 |
| TpMuguga_01g00112 | 1 | 0.026548673 |
| TpMuguga_01g00112 | 2 | 0 |
| TpMuguga_01g02110 | 1 | 0.078431373 |
| TpMuguga_01g02110 | 2 | 0 |
| TpMuguga_01g02110 | 3 | 0.339869281 |
| TpMuguga_01g00115 | 1 | 5 |
| TpMuguga_01g00115 | 2 | 0 |
| TpMuguga_01g00115 | 3 | 0 |
| TpMuguga_01g00115 | 4 | 0 |
| TpMuguga_01g00115 | 5 | 0 |
| TpMuguga_01g00115 | 6 | 0 |
| TpMuguga_01g00115 | 7 | 0 |
| TpMuguga_01g00115 | 8 | 0 |
| TpMuguga_01g00118 | 1 | 6.857142857 |
| TpMuguga_01g00118 | 2 | 1.714285714 |
| TpMuguga_01g00118 | 3 | 0 |
| TpMuguga_01g00118 | 4 | 0 |
| TpMuguga_01g00118 | 5 | 0 |
| TpMuguga_01g00118 | 6 | 0 |
| TpMuguga_01g00118 | 7 | 0 |
| TpMuguga_01g00118 | 8 | 0 |
| TpMuguga_01g00118 | 9 | 0 |
| TpMuguga_01g00118 | 10 | 0 |
| TpMuguga_01g00118 | 11 | 0 |
| TpMuguga_01g02070 | 1 | 0 |
| TpMuguga_01g02070 | 2 | 0 |
| TpMuguga_01g02070 | 3 | 0 |
| TpMuguga_01g02070 | 4 | 0 |
| TpMuguga_01g02070 | 5 | 0 |
| TpMuguga_01g02070 | 6 | 0 |
| TpMuguga_01g02070 | 7 | 0 |
| TpMuguga_01g02070 | 8 | 0 |

| | | |
|---|---|---|
| TpMuguga_01g02070 | 9 | 0 |
| TpMuguga_01g02070 | 10 | 0 |
| TpMuguga_01g02070 | 11 | 0 |
| TpMuguga_01g02070 | 12 | 0 |
| TpMuguga_01g02070 | 13 | 0 |
| TpMuguga_01g02070 | 14 | 0 |
| TpMuguga_01g02070 | 15 | 0 |
| TpMuguga_01g02070 | 16 | 0 |
| TpMuguga_01g02070 | 17 | 0 |
| TpMuguga_01g02070 | 18 | 0 |
| TpMuguga_01g02070 | 19 | 0 |
| TpMuguga_01g02070 | 20 | 0 |
| TpMuguga_01g02070 | 21 | 0 |
| TpMuguga_01g02070 | 22 | 0 |
| TpMuguga_01g02070 | 23 | 0 |
| TpMuguga_01g00119 | 1 | 7.5 |
| TpMuguga_01g00119 | 2 | 0 |
| TpMuguga_01g00119 | 3 | 0 |
| TpMuguga_01g00119 | 4 | 0 |
| TpMuguga_01g00119 | 5 | 0 |
| TpMuguga_01g00119 | 6 | 0 |
| TpMuguga_01g00119 | 7 | 0 |
| TpMuguga_01g00119 | 8 | 0 |
| TpMuguga_01g00119 | 9 | 5 |
| TpMuguga_01g00124 | 1 | 0 |
| TpMuguga_01g00124 | 2 | 0 |
| TpMuguga_01g00124 | 3 | 0 |
| TpMuguga_01g00124 | 4 | 0 |
| TpMuguga_01g00125 | 1 | 0.534246575 |
| TpMuguga_01g00125 | 2 | 7.315068493 |
| TpMuguga_01g00125 | 3 | 5.506849315 |
| TpMuguga_01g00125 | 4 | 3 |
| TpMuguga_01g00125 | 5 | 0 |
| TpMuguga_01g00125 | 6 | 0 |
| TpMuguga_01g00125 | 7 | 0 |
| TpMuguga_01g00125 | 8 | 0 |
| TpMuguga_01g00126 | 1 | 0 |
| TpMuguga_01g00126 | 2 | 0 |
| TpMuguga_01g00126 | 3 | 0 |
| TpMuguga_01g00126 | 4 | 0 |
| TpMuguga_01g00126 | 5 | 0 |
| TpMuguga_01g00126 | 6 | 0 |

| | | |
|---|---|---|
| TpMuguga_01g00126 | 7 | 0 |
| TpMuguga_01g00126 | 8 | 0 |
| TpMuguga_01g00126 | 9 | 0 |
| TpMuguga_01g00127 | 1 | 0 |
| TpMuguga_01g00127 | 2 | 0 |
| TpMuguga_01g00128 | 1 | 0.021459227 |
| TpMuguga_01g00128 | 2 | 0 |
| TpMuguga_01g00128 | 3 | 0 |
| TpMuguga_01g00128 | 4 | 0 |
| TpMuguga_01g00129 | 1 | 0 |
| TpMuguga_01g00129 | 2 | 0.205882353 |
| TpMuguga_01g00129 | 3 | 0 |
| TpMuguga_01g00129 | 4 | 0 |
| TpMuguga_01g00129 | 5 | 0 |
| TpMuguga_01g00129 | 6 | 0 |
| TpMuguga_01g00130 | 1 | 0.158311346 |
| TpMuguga_01g00130 | 2 | 0 |
| TpMuguga_01g00130 | 3 | 0 |
| TpMuguga_01g00130 | 4 | 0.158311346 |
| TpMuguga_01g00130 | 5 | 0 |
| TpMuguga_01g00130 | 6 | 0 |
| TpMuguga_01g00130 | 7 | 0.094986807 |
| TpMuguga_01g00130 | 8 | 0 |
| TpMuguga_01g00130 | 9 | 0 |
| TpMuguga_01g00130 | 10 | 0 |
| TpMuguga_01g00130 | 11 | 0 |
| TpMuguga_01g00132 | 1 | 0 |
| TpMuguga_01g00132 | 2 | 0 |
| TpMuguga_01g00132 | 3 | 0 |
| TpMuguga_01g00132 | 4 | 0 |
| TpMuguga_01g00132 | 5 | 0 |
| TpMuguga_01g00138 | 1 | 24 |
| TpMuguga_01g00137 | 1 | 0.134328358 |
| TpMuguga_01g00137 | 2 | 0.223880597 |
| TpMuguga_01g00136 | 1 | 3.408 |
| TpMuguga_01g00136 | 2 | 0.288 |
| TpMuguga_01g00136 | 3 | 0.24 |
| TpMuguga_01g00136 | 4 | 1.152 |
| TpMuguga_01g00136 | 5 | 0.528 |
| TpMuguga_01g00139 | 1 | 0 |
| TpMuguga_01g00139 | 2 | 0.168224299 |
| TpMuguga_01g00139 | 3 | 0.224299065 |

| | | |
|---|---|---|
| TpMuguga_01g00139 | 4 | 0.560747664 |
| TpMuguga_01g00139 | 5 | 6.336448598 |
| TpMuguga_01g00140 | 1 | 0.562352941 |
| TpMuguga_01g00140 | 2 | 0.788235294 |
| TpMuguga_01g00140 | 3 | 0.632941176 |
| TpMuguga_01g02465 | 1 | 0 |
| TpMuguga_01g02465 | 2 | 6.035714286 |
| TpMuguga_01g02465 | 3 | 0.464285714 |
| TpMuguga_01g02465 | 4 | 0 |
| TpMuguga_01g02465 | 5 | 0 |
| TpMuguga_01g02465 | 6 | 0 |
| TpMuguga_01g02465 | 7 | 0.232142857 |
| TpMuguga_01g02465 | 8 | 0 |
| TpMuguga_01g02465 | 9 | 0 |
| TpMuguga_01g02465 | 10 | 0 |
| TpMuguga_01g02465 | 11 | 0 |
| TpMuguga_01g02465 | 12 | 0 |
| TpMuguga_01g00145 | 1 | 0 |
| TpMuguga_01g00145 | 2 | 0 |
| TpMuguga_01g00145 | 3 | 0 |
| TpMuguga_01g00145 | 4 | 0 |
| TpMuguga_01g02200 | 1 | 0 |
| TpMuguga_01g02200 | 2 | 0 |
| TpMuguga_01g02200 | 3 | 0 |
| TpMuguga_01g02200 | 4 | 0 |
| TpMuguga_01g02200 | 5 | 0 |
| TpMuguga_01g02200 | 6 | 0 |
| TpMuguga_01g02200 | 7 | 0 |
| TpMuguga_01g02200 | 8 | 0 |
| TpMuguga_01g02200 | 9 | 0 |
| TpMuguga_01g00148 | 1 | 0 |
| TpMuguga_01g00148 | 2 | 0 |
| TpMuguga_01g00148 | 3 | 0 |
| TpMuguga_01g00148 | 4 | 0 |
| TpMuguga_01g00148 | 5 | 0 |
| TpMuguga_01g00148 | 6 | 0 |
| TpMuguga_01g00148 | 7 | 0 |
| TpMuguga_01g00148 | 8 | 0 |
| TpMuguga_01g00148 | 9 | 0 |
| TpMuguga_01g00149 | 1 | 0 |
| TpMuguga_01g00149 | 2 | 0 |
| TpMuguga_01g00149 | 3 | 0 |

| | | |
|---|---|---|
| TpMuguga_01g00149 | 4 | 0 |
| TpMuguga_01g00149 | 5 | 0 |
| TpMuguga_01g00150 | 1 | 0 |
| TpMuguga_01g00150 | 2 | 0.156028369 |
| TpMuguga_01g00150 | 3 | 0 |
| TpMuguga_01g00150 | 4 | 0.286052009 |
| TpMuguga_01g00150 | 5 | 0.182033097 |
| TpMuguga_01g00150 | 6 | 1.144208038 |
| TpMuguga_01g00150 | 7 | 1.014184397 |
| TpMuguga_01g00150 | 8 | 1.144208038 |
| TpMuguga_01g00150 | 9 | 2.808510638 |
| TpMuguga_01g00150 | 10 | 4.108747045 |
| TpMuguga_01g00151 | 1 | 0 |
| TpMuguga_01g00151 | 2 | 0 |
| TpMuguga_01g00151 | 3 | 0 |
| TpMuguga_01g00151 | 4 | 0 |
| TpMuguga_01g00151 | 5 | 0 |
| TpMuguga_01g00151 | 6 | 0 |
| TpMuguga_01g00151 | 7 | 0 |
| TpMuguga_01g00151 | 8 | 0 |
| TpMuguga_01g00151 | 9 | 0 |
| TpMuguga_01g00151 | 10 | 0 |
| TpMuguga_01g02625 | 1 | 0.178571429 |
| TpMuguga_01g02625 | 2 | 0.178571429 |
| TpMuguga_01g02625 | 3 | 0 |
| TpMuguga_01g02625 | 4 | 0.178571429 |
| TpMuguga_01g00153 | 1 | 0 |
| TpMuguga_01g00155 | 1 | 0.104672897 |
| TpMuguga_01g00155 | 2 | 0.052336449 |
| TpMuguga_01g00155 | 3 | 0 |
| TpMuguga_01g00155 | 4 | 0 |
| TpMuguga_01g00155 | 5 | 0.798130841 |
| TpMuguga_01g00155 | 6 | 4.998130841 |
| TpMuguga_01g00156 | 1 | 0.185840708 |
| TpMuguga_01g00156 | 2 | 0 |
| TpMuguga_01g00157 | 1 | 0.984615385 |
| TpMuguga_01g00158 | 1 | 0 |
| TpMuguga_01g00158 | 2 | 0 |
| TpMuguga_01g00158 | 3 | 0 |
| TpMuguga_01g00158 | 4 | 0 |
| TpMuguga_01g00158 | 5 | 0 |
| TpMuguga_01g00159 | 1 | 0 |

| | | |
|---|---|---|
| TpMuguga_01g00159 | 2 | 0 |
| TpMuguga_01g00159 | 3 | 0 |
| TpMuguga_01g00159 | 4 | 0.168067227 |
| TpMuguga_01g00161 | 1 | 1.097560976 |
| TpMuguga_01g00161 | 2 | 0.43902439 |
| TpMuguga_01g00162 | 1 | 0 |
| TpMuguga_01g00162 | 2 | 0 |
| TpMuguga_01g00162 | 3 | 0 |
| TpMuguga_01g00162 | 4 | 7.954545455 |
| TpMuguga_01g00162 | 5 | 9.227272727 |
| TpMuguga_01g00162 | 6 | 3.181818182 |
| TpMuguga_01g00163 | 1 | 3.321428571 |
| TpMuguga_01g00163 | 2 | 0.107142857 |
| TpMuguga_01g00164 | 1 | 0 |
| TpMuguga_01g00164 | 2 | 0 |
| TpMuguga_01g00164 | 3 | 0 |
| TpMuguga_01g00164 | 4 | 0 |
| TpMuguga_01g00164 | 5 | 0 |
| TpMuguga_01g00164 | 6 | 1.911764706 |
| TpMuguga_01g00164 | 7 | 0 |
| TpMuguga_01g00164 | 8 | 2.5 |
| TpMuguga_01g00164 | 9 | 2.205882353 |
| TpMuguga_01g00165 | 1 | 1.119047619 |
| TpMuguga_01g00166 | 1 | 0 |
| TpMuguga_01g00166 | 2 | 0 |
| TpMuguga_01g00166 | 3 | 0 |
| TpMuguga_01g00166 | 4 | 0 |
| TpMuguga_01g00166 | 5 | 0 |
| TpMuguga_01g00166 | 6 | 0 |
| TpMuguga_01g00166 | 7 | 0 |
| TpMuguga_01g00166 | 8 | 0 |
| TpMuguga_01g00166 | 9 | 0 |
| TpMuguga_01g00167 | 1 | 0 |
| TpMuguga_01g00167 | 2 | 0 |
| TpMuguga_01g00168 | 1 | 0.166666667 |
| TpMuguga_01g02120 | 1 | 0 |
| TpMuguga_01g02120 | 2 | 0 |
| TpMuguga_01g02120 | 3 | 0 |
| TpMuguga_01g02120 | 4 | 0 |
| TpMuguga_01g02120 | 5 | 0 |
| TpMuguga_01g02120 | 6 | 0.863636364 |
| TpMuguga_01g02120 | 7 | 0 |

| | | |
|---|---|---|
| TpMuguga_01g00171 | 1 | 0 |
| TpMuguga_01g00173 | 1 | 0 |
| TpMuguga_01g00173 | 2 | 0 |
| TpMuguga_01g00173 | 3 | 0 |
| TpMuguga_01g00172 | 1 | 0 |
| TpMuguga_01g00172 | 2 | 0 |
| TpMuguga_01g00172 | 3 | 0 |
| TpMuguga_01g00172 | 4 | 0 |
| TpMuguga_01g00175 | 1 | 9.942857143 |
| TpMuguga_01g00175 | 2 | 0 |
| TpMuguga_01g00175 | 3 | 1.885714286 |
| TpMuguga_01g00175 | 4 | 0 |
| TpMuguga_01g00175 | 5 | 0 |
| TpMuguga_01g00174 | 1 | 1.431818182 |
| TpMuguga_01g00174 | 2 | 0.068181818 |
| TpMuguga_01g00176 | 1 | 0 |
| TpMuguga_01g00176 | 2 | 3.108108108 |
| TpMuguga_01g00176 | 3 | 0 |
| TpMuguga_01g00176 | 4 | 0.405405405 |
| TpMuguga_01g00179 | 1 | 0.666666667 |
| TpMuguga_01g00179 | 2 | 0 |
| TpMuguga_01g00179 | 3 | 0 |
| TpMuguga_01g00181 | 1 | 1.5 |
| TpMuguga_01g00181 | 2 | 4.5 |
| TpMuguga_01g00182 | 1 | 0.0625 |
| TpMuguga_01g00182 | 2 | 0.1875 |
| TpMuguga_01g00182 | 3 | 0 |
| TpMuguga_01g00182 | 4 | 0 |
| TpMuguga_01g00182 | 5 | 0 |
| TpMuguga_01g00182 | 6 | 0 |
| TpMuguga_01g00182 | 7 | 0.0625 |
| TpMuguga_01g00182 | 8 | 0 |
| TpMuguga_01g00184 | 1 | 0 |
| TpMuguga_01g00184 | 2 | 0 |
| TpMuguga_01g00185 | 1 | 0 |
| TpMuguga_01g00185 | 2 | 0.424242424 |
| TpMuguga_01g00185 | 3 | 2.757575758 |
| TpMuguga_01g00185 | 4 | 0 |
| TpMuguga_01g00185 | 5 | 0 |
| TpMuguga_01g00185 | 6 | 0 |
| TpMuguga_01g00186 | 1 | 0 |
| TpMuguga_01g00186 | 2 | 0 |

| | | |
|---|---|---|
| TpMuguga_01g00186 | 3 | 0 |
| TpMuguga_01g00186 | 4 | 0 |
| TpMuguga_01g00186 | 5 | 0 |
| TpMuguga_01g00186 | 6 | 0 |
| TpMuguga_01g00186 | 7 | 0 |
| TpMuguga_01g00186 | 8 | 0 |
| TpMuguga_01g00186 | 9 | 0 |
| TpMuguga_01g00186 | 10 | 0 |
| TpMuguga_01g00186 | 11 | 4 |
| TpMuguga_01g00187 | 1 | 0 |
| TpMuguga_01g00188 | 1 | 0 |
| TpMuguga_01g00190 | 1 | 0.333333333 |
| TpMuguga_01g00190 | 2 | 0 |
| TpMuguga_01g00189 | 1 | 0 |
| TpMuguga_01g00189 | 2 | 1.107142857 |
| TpMuguga_01g00191 | 1 | 0 |
| TpMuguga_01g00191 | 2 | 0 |
| TpMuguga_01g00191 | 3 | 0 |
| TpMuguga_01g00191 | 4 | 0 |
| TpMuguga_01g00192 | 1 | 0 |
| TpMuguga_01g00193 | 1 | 0 |
| TpMuguga_01g00193 | 2 | 0 |
| TpMuguga_01g00193 | 3 | 0 |
| TpMuguga_01g00193 | 4 | 0 |
| TpMuguga_01g00193 | 5 | 0 |
| TpMuguga_01g00193 | 6 | 0 |
| TpMuguga_01g00193 | 7 | 0 |
| TpMuguga_01g00193 | 8 | 0 |
| TpMuguga_01g00195 | 1 | 0 |
| TpMuguga_01g00195 | 2 | 0 |
| TpMuguga_01g00195 | 3 | 0 |
| TpMuguga_01g02045 | 1 | 0 |
| TpMuguga_01g02045 | 2 | 0 |
| TpMuguga_01g02045 | 3 | 0 |
| TpMuguga_01g02045 | 4 | 0 |
| TpMuguga_01g00196 | 1 | 0 |
| TpMuguga_01g00196 | 2 | 0 |
| TpMuguga_01g00197 | 1 | 0 |
| TpMuguga_01g00197 | 2 | 0 |
| TpMuguga_01g00197 | 3 | 0 |
| TpMuguga_01g00197 | 4 | 0 |
| TpMuguga_01g00197 | 5 | 0 |

| | | |
|---|---|---|
| TpMuguga_01g00197 | 6 | 0 |
| TpMuguga_01g00197 | 7 | 0 |
| TpMuguga_01g00197 | 8 | 0 |
| TpMuguga_01g00198 | 1 | 0 |
| TpMuguga_01g00199 | 1 | 0 |
| TpMuguga_01g00199 | 2 | 0 |
| TpMuguga_01g00200 | 1 | 0 |
| TpMuguga_01g00201 | 1 | 0 |
| TpMuguga_01g00201 | 2 | 0 |
| TpMuguga_01g00201 | 3 | 0 |
| TpMuguga_01g00201 | 4 | 0 |
| TpMuguga_01g00201 | 5 | 2.4 |
| TpMuguga_01g00202 | 1 | 0 |
| TpMuguga_01g00202 | 2 | 0 |
| TpMuguga_01g00202 | 3 | 0 |
| TpMuguga_01g00202 | 4 | 0 |
| TpMuguga_01g00202 | 5 | 0 |
| TpMuguga_01g00202 | 6 | 0 |
| TpMuguga_01g00202 | 7 | 0 |
| TpMuguga_01g00202 | 8 | 2 |
| TpMuguga_01g00202 | 9 | 0 |
| TpMuguga_01g00202 | 10 | 0 |
| TpMuguga_01g00202 | 11 | 0 |
| TpMuguga_01g00202 | 12 | 24 |
| TpMuguga_01g00202 | 13 | 38 |
| TpMuguga_01g02055 | 1 | 0 |
| TpMuguga_01g02055 | 2 | 0 |
| TpMuguga_01g02055 | 3 | 0 |
| TpMuguga_01g02055 | 4 | 0 |
| TpMuguga_01g02055 | 5 | 0 |
| TpMuguga_01g02055 | 6 | 0 |
| TpMuguga_01g02055 | 7 | 0 |
| TpMuguga_01g02055 | 8 | 0 |
| TpMuguga_01g02055 | 9 | 0 |
| TpMuguga_01g02055 | 10 | 0 |
| TpMuguga_01g02055 | 11 | 0 |
| TpMuguga_01g02055 | 12 | 0 |
| TpMuguga_01g02055 | 13 | 0 |
| TpMuguga_01g02055 | 14 | 0 |
| TpMuguga_01g02055 | 15 | 7.98185941 |
| TpMuguga_01g00206 | 1 | 0 |
| TpMuguga_01g00206 | 2 | 0 |

| | | |
|---|---|---|
| TpMuguga_01g00206 | 3 | 0 |
| TpMuguga_01g00206 | 4 | 0 |
| TpMuguga_01g00207 | 1 | 0 |
| TpMuguga_01g00208 | 1 | 0 |
| TpMuguga_01g00208 | 2 | 0 |
| TpMuguga_01g00208 | 3 | 0 |
| TpMuguga_01g00208 | 4 | 0 |
| TpMuguga_01g00208 | 5 | 0 |
| TpMuguga_01g00209 | 1 | 0 |
| TpMuguga_01g00209 | 2 | 0 |
| TpMuguga_01g00210 | 1 | 0 |
| TpMuguga_01g00210 | 2 | 0 |
| TpMuguga_01g00210 | 3 | 0 |
| TpMuguga_01g00210 | 4 | 0 |
| TpMuguga_01g00210 | 5 | 0 |
| TpMuguga_01g02015 | 1 | 0 |
| TpMuguga_01g02015 | 2 | 0 |
| TpMuguga_01g02015 | 3 | 0 |
| TpMuguga_01g02015 | 4 | 0 |
| TpMuguga_01g02015 | 5 | 0 |
| TpMuguga_01g02015 | 6 | 0 |
| TpMuguga_01g02015 | 7 | 0 |
| TpMuguga_01g02355 | 1 | 0 |
| TpMuguga_01g02355 | 2 | 0 |
| TpMuguga_01g00213 | 1 | 0 |
| TpMuguga_01g00214 | 1 | 0.571428571 |
| TpMuguga_01g02585 | 1 | 0 |
| TpMuguga_01g02585 | 2 | 0 |
| TpMuguga_01g02585 | 3 | 0 |
| TpMuguga_01g02585 | 4 | 0 |
| TpMuguga_01g02585 | 5 | 0 |
| TpMuguga_01g02585 | 6 | 0 |
| TpMuguga_01g02585 | 7 | 0 |
| TpMuguga_01g02585 | 8 | 0 |
| TpMuguga_01g02780 | 1 | 0 |
| TpMuguga_01g02780 | 2 | 0 |
| TpMuguga_01g02780 | 3 | 0 |
| TpMuguga_01g02780 | 4 | 0 |
| TpMuguga_01g02780 | 5 | 0 |
| TpMuguga_01g00219 | 1 | 0 |
| TpMuguga_01g00219 | 2 | 0 |
| TpMuguga_01g00219 | 3 | 0 |

| | | |
|---|---|---|
| TpMuguga_01g00219 | 4 | 0 |
| TpMuguga_01g00219 | 5 | 0 |
| TpMuguga_01g00219 | 6 | 10 |
| TpMuguga_01g00219 | 7 | 0 |
| TpMuguga_01g00219 | 8 | 0 |
| TpMuguga_01g00219 | 9 | 0 |
| TpMuguga_01g00220 | 1 | 0 |
| TpMuguga_01g00222 | 1 | 0 |
| TpMuguga_01g00222 | 2 | 0 |
| TpMuguga_01g00222 | 3 | 0 |
| TpMuguga_01g00222 | 4 | 0 |
| TpMuguga_01g00222 | 5 | 0 |
| TpMuguga_01g00222 | 6 | 0 |
| TpMuguga_01g00222 | 7 | 0 |
| TpMuguga_01g00222 | 8 | 0 |
| TpMuguga_01g00222 | 9 | 0 |
| TpMuguga_01g00222 | 10 | 0 |
| TpMuguga_01g00222 | 11 | 0 |
| TpMuguga_01g00222 | 12 | 0 |
| TpMuguga_01g00224 | 1 | 0.324324324 |
| TpMuguga_01g00224 | 2 | 1.72972973 |
| TpMuguga_01g00224 | 3 | 0.648648649 |
| TpMuguga_01g00223 | 1 | 0 |
| TpMuguga_01g00223 | 2 | 0 |
| TpMuguga_01g00223 | 3 | 0 |
| TpMuguga_01g00223 | 4 | 0 |
| TpMuguga_01g00223 | 5 | 0 |
| TpMuguga_01g00223 | 6 | 0 |
| TpMuguga_01g00223 | 7 | 0 |
| TpMuguga_01g00223 | 8 | 0 |
| TpMuguga_01g00223 | 9 | 0 |
| TpMuguga_01g00223 | 10 | 0 |
| TpMuguga_01g00223 | 11 | 0 |
| TpMuguga_01g00223 | 12 | 0 |
| TpMuguga_01g00223 | 13 | 0 |
| TpMuguga_01g00223 | 14 | 0 |
| TpMuguga_01g00225 | 1 | 0.326923077 |
| TpMuguga_01g00226 | 1 | 4 |
| TpMuguga_01g00227 | 1 | 0 |
| TpMuguga_01g00227 | 2 | 0 |
| TpMuguga_01g00227 | 3 | 0 |
| TpMuguga_01g00227 | 4 | 4.666666667 |

| | | |
|---|---|---|
| TpMuguga_01g00227 | 5 | 0 |
| TpMuguga_01g02025 | 1 | 0 |
| TpMuguga_01g02025 | 2 | 0 |
| TpMuguga_01g02025 | 3 | 5.25 |
| TpMuguga_01g02025 | 4 | 0 |
| TpMuguga_01g02025 | 5 | 0 |
| TpMuguga_01g02025 | 6 | 0 |
| TpMuguga_01g00230 | 1 | 0 |
| TpMuguga_01g00230 | 2 | 0 |
| TpMuguga_01g00231 | 1 | 5.294117647 |
| TpMuguga_01g00231 | 2 | 3.176470588 |
| TpMuguga_01g00231 | 3 | 0 |
| TpMuguga_01g00231 | 4 | 3.176470588 |
| TpMuguga_01g00231 | 5 | 0 |
| TpMuguga_01g00231 | 6 | 0 |
| TpMuguga_01g00231 | 7 | 6.882352941 |
| TpMuguga_01g00231 | 8 | 8.470588235 |
| TpMuguga_01g00232 | 1 | 0.829268293 |
| TpMuguga_01g00233 | 1 | 0 |
| TpMuguga_01g00233 | 2 | 0 |
| TpMuguga_01g00235 | 1 | 0.888888889 |
| TpMuguga_01g00237 | 1 | 0 |
| TpMuguga_01g00237 | 2 | 0 |
| TpMuguga_01g00237 | 3 | 1.125 |
| TpMuguga_01g00237 | 4 | 0.375 |
| TpMuguga_01g00237 | 5 | 7.875 |
| TpMuguga_01g00238 | 1 | 0 |
| TpMuguga_01g00238 | 2 | 0 |
| TpMuguga_01g00238 | 3 | 0.095238095 |
| TpMuguga_01g00240 | 1 | 0 |
| TpMuguga_01g00240 | 2 | 0 |
| TpMuguga_01g00240 | 3 | 0 |
| TpMuguga_01g00240 | 4 | 0 |
| TpMuguga_01g00241 | 1 | 0 |
| TpMuguga_01g00241 | 2 | 0 |
| TpMuguga_01g00241 | 3 | 0 |
| TpMuguga_01g00241 | 4 | 0 |
| TpMuguga_01g00241 | 5 | 0 |
| TpMuguga_01g00241 | 6 | 0 |
| TpMuguga_01g00241 | 7 | 0 |
| TpMuguga_01g00242 | 1 | 0 |
| TpMuguga_01g00242 | 2 | 0 |

| | | |
|---|---|---|
| TpMuguga_01g00242 | 3 | 0 |
| TpMuguga_01g00242 | 4 | 0 |
| TpMuguga_01g00242 | 5 | 0 |
| TpMuguga_01g00242 | 6 | 0 |
| TpMuguga_01g00242 | 7 | 0 |
| TpMuguga_01g00242 | 8 | 0 |
| TpMuguga_01g00243 | 1 | 0 |
| TpMuguga_01g00243 | 2 | 0 |
| TpMuguga_01g00244 | 1 | 0.49122807 |
| TpMuguga_01g00244 | 2 | 0 |
| TpMuguga_01g00244 | 3 | 0.140350877 |
| TpMuguga_01g00245 | 1 | 0 |
| TpMuguga_01g00246 | 1 | 0 |
| TpMuguga_01g00246 | 2 | 0 |
| TpMuguga_01g00247 | 1 | 0 |
| TpMuguga_01g00247 | 2 | 0 |
| TpMuguga_01g00247 | 3 | 0 |
| TpMuguga_01g00247 | 4 | 0 |
| TpMuguga_01g00247 | 5 | 0 |
| TpMuguga_01g00247 | 6 | 0 |
| TpMuguga_01g00247 | 7 | 0 |
| TpMuguga_01g00247 | 8 | 0 |
| TpMuguga_01g00247 | 9 | 0 |
| TpMuguga_01g00247 | 10 | 0 |
| TpMuguga_01g00247 | 11 | 0 |
| TpMuguga_01g00248 | 1 | 7.5 |
| TpMuguga_01g00248 | 2 | 0 |
| TpMuguga_01g00248 | 3 | 6 |
| TpMuguga_01g00248 | 4 | 0 |
| TpMuguga_01g00248 | 5 | 0 |
| TpMuguga_01g00249 | 1 | 0 |
| TpMuguga_01g00249 | 2 | 0 |
| TpMuguga_01g00249 | 3 | 0.224137931 |
| TpMuguga_01g00249 | 4 | 0.327586207 |
| TpMuguga_01g00250 | 1 | 0 |
| TpMuguga_01g00250 | 2 | 0.161616162 |
| TpMuguga_01g00250 | 3 | 0 |
| TpMuguga_01g00250 | 4 | 0 |
| TpMuguga_01g00250 | 5 | 0 |
| TpMuguga_01g00250 | 6 | 0 |
| TpMuguga_01g00250 | 7 | 0 |
| TpMuguga_01g00251 | 1 | 0 |

| | | |
|---|---|---|
| TpMuguga_01g00251 | 2 | 0 |
| TpMuguga_01g00251 | 3 | 0 |
| TpMuguga_01g00251 | 4 | 0 |
| TpMuguga_01g00251 | 5 | 4.5 |
| TpMuguga_01g00252 | 1 | 0 |
| TpMuguga_01g00252 | 2 | 1.142857143 |
| TpMuguga_01g00252 | 3 | 0 |
| TpMuguga_01g00252 | 4 | 0.761904762 |
| TpMuguga_01g00252 | 5 | 0 |
| TpMuguga_01g00252 | 6 | 0 |
| TpMuguga_01g00252 | 7 | 0.952380952 |
| TpMuguga_01g00253 | 1 | 6.042654028 |
| TpMuguga_01g00253 | 2 | 0.639810427 |
| TpMuguga_01g00253 | 3 | 0.971563981 |
| TpMuguga_01g00253 | 4 | 0.710900474 |
| TpMuguga_01g00254 | 1 | 1.38028169 |
| TpMuguga_01g00254 | 2 | 0 |
| TpMuguga_01g00254 | 3 | 0.098591549 |
| TpMuguga_01g00254 | 4 | 1.41314554 |
| TpMuguga_01g00254 | 5 | 1.018779343 |
| TpMuguga_01g00254 | 6 | 0.887323944 |
| TpMuguga_01g00255 | 1 | 0 |
| TpMuguga_01g00255 | 2 | 0 |
| TpMuguga_01g00255 | 3 | 0 |
| TpMuguga_01g00255 | 4 | 0 |
| TpMuguga_01g00257 | 1 | 0 |
| TpMuguga_01g02275 | 1 | 0 |
| TpMuguga_01g02275 | 2 | 0 |
| TpMuguga_01g00262 | 1 | 0 |
| TpMuguga_01g00262 | 2 | 0 |
| TpMuguga_01g00262 | 3 | 0 |
| TpMuguga_01g00262 | 4 | 0 |
| TpMuguga_01g00262 | 5 | 0 |
| TpMuguga_01g00262 | 6 | 0 |
| TpMuguga_01g00262 | 7 | 0 |
| TpMuguga_01g00264 | 1 | 0.026086957 |
| TpMuguga_01g00264 | 2 | 0.07826087 |
| TpMuguga_01g00265 | 1 | 1.333333333 |
| TpMuguga_01g00265 | 2 | 0 |
| TpMuguga_01g00266 | 1 | 0 |
| TpMuguga_01g00266 | 2 | 0 |
| TpMuguga_01g00267 | 1 | 0 |

| | | |
|---|---|---|
| TpMuguga_01g00267 | 2 | 0 |
| TpMuguga_01g00267 | 3 | 0 |
| TpMuguga_01g00267 | 4 | 0 |
| TpMuguga_01g00268 | 1 | 0 |
| TpMuguga_01g00269 | 1 | 0 |
| TpMuguga_01g00269 | 2 | 0 |
| TpMuguga_01g00269 | 3 | 0 |
| TpMuguga_01g00269 | 4 | 3.666666667 |
| TpMuguga_01g00269 | 5 | 0 |
| TpMuguga_01g00269 | 6 | 0 |
| TpMuguga_01g00269 | 7 | 0 |
| TpMuguga_01g00269 | 8 | 7.333333333 |
| TpMuguga_01g00269 | 9 | 0 |
| TpMuguga_01g00269 | 10 | 0 |
| TpMuguga_01g00270 | 1 | 0 |
| TpMuguga_01g00270 | 2 | 0 |
| TpMuguga_01g00271 | 1 | 0 |
| TpMuguga_01g00271 | 2 | 0 |
| TpMuguga_01g02345 | 1 | 0 |
| TpMuguga_01g02345 | 2 | 0 |
| TpMuguga_01g02345 | 3 | 0 |
| TpMuguga_01g02345 | 4 | 0 |
| TpMuguga_01g02345 | 5 | 0 |
| TpMuguga_01g02345 | 6 | 0 |
| TpMuguga_01g02345 | 7 | 0 |
| TpMuguga_01g00272 | 1 | 0 |
| TpMuguga_01g00272 | 2 | 0 |
| TpMuguga_01g00272 | 3 | 0 |
| TpMuguga_01g00272 | 4 | 0.330188679 |
| TpMuguga_01g00277 | 1 | 0 |
| TpMuguga_01g00277 | 2 | 0 |
| TpMuguga_01g00277 | 3 | 0.111111111 |
| TpMuguga_01g00277 | 4 | 0 |
| TpMuguga_01g00276 | 1 | 0 |
| TpMuguga_01g00276 | 2 | 0.636363636 |
| TpMuguga_01g00278 | 1 | 0 |
| TpMuguga_01g00278 | 2 | 0 |
| TpMuguga_01g00278 | 3 | 0 |
| TpMuguga_01g00278 | 4 | 0 |
| TpMuguga_01g00278 | 5 | 0 |
| TpMuguga_01g00278 | 6 | 0 |
| TpMuguga_01g00278 | 7 | 0 |

| | | |
|---|---|---|
| TpMuguga_01g00278 | 8 | 0 |
| TpMuguga_01g00278 | 9 | 0 |
| TpMuguga_01g00278 | 10 | 0 |
| TpMuguga_01g00278 | 11 | 0 |
| TpMuguga_01g00278 | 12 | 0 |
| TpMuguga_01g00278 | 13 | 0 |
| TpMuguga_01g00279 | 1 | 11.2 |
| TpMuguga_01g00279 | 2 | 0 |
| TpMuguga_01g00279 | 3 | 1.6 |
| TpMuguga_01g00280 | 1 | 0 |
| TpMuguga_01g00281 | 1 | 0 |
| TpMuguga_01g00281 | 2 | 0 |
| TpMuguga_01g00281 | 3 | 0 |
| TpMuguga_01g00281 | 4 | 0 |
| TpMuguga_01g00281 | 5 | 0 |
| TpMuguga_01g00281 | 6 | 0 |
| TpMuguga_01g00282 | 1 | 10 |
| TpMuguga_01g00282 | 2 | 0 |
| TpMuguga_01g00282 | 3 | 0 |
| TpMuguga_01g00282 | 4 | 0 |
| TpMuguga_01g00283 | 1 | 0.015321757 |
| TpMuguga_01g00283 | 2 | 0.306435138 |
| TpMuguga_01g00283 | 3 | 1.488764045 |
| TpMuguga_01g00283 | 4 | 0.459652707 |
| TpMuguga_01g00284 | 1 | 0 |
| TpMuguga_01g00284 | 2 | 0 |
| TpMuguga_01g00284 | 3 | 0 |
| TpMuguga_01g00284 | 4 | 0 |
| TpMuguga_01g00284 | 5 | 0 |
| TpMuguga_01g00284 | 6 | 0 |
| TpMuguga_01g00284 | 7 | 0 |
| TpMuguga_01g00284 | 8 | 0 |
| TpMuguga_01g00284 | 9 | 0 |
| TpMuguga_01g00285 | 1 | 1.475409836 |
| TpMuguga_01g00285 | 2 | 0 |
| TpMuguga_01g00285 | 3 | 0 |
| TpMuguga_01g00285 | 4 | 0.491803279 |
| TpMuguga_01g00285 | 5 | 0.786885246 |
| TpMuguga_01g00286 | 1 | 0 |
| TpMuguga_01g00286 | 2 | 0 |
| TpMuguga_01g00286 | 3 | 0 |
| TpMuguga_01g00286 | 4 | 0 |

| | | |
|---|---|---|
| TpMuguga_01g00286 | 5 | 0 |
| TpMuguga_01g00286 | 6 | 0 |
| TpMuguga_01g00287 | 1 | 2.4 |
| TpMuguga_01g00287 | 2 | 1.4 |
| TpMuguga_01g00287 | 3 | 0.8 |
| TpMuguga_01g00287 | 4 | 0.2 |
| TpMuguga_01g00287 | 5 | 0 |
| TpMuguga_01g00290 | 1 | 0 |
| TpMuguga_01g00290 | 2 | 0 |
| TpMuguga_01g00290 | 3 | 0 |
| TpMuguga_01g00290 | 4 | 0 |
| TpMuguga_01g00290 | 5 | 0 |
| TpMuguga_01g00290 | 6 | 0 |
| TpMuguga_01g00290 | 7 | 0 |
| TpMuguga_01g00290 | 8 | 0 |
| TpMuguga_01g00289 | 1 | 0 |
| TpMuguga_01g00289 | 2 | 0 |
| TpMuguga_01g00289 | 3 | 0 |
| TpMuguga_01g00289 | 4 | 0 |
| TpMuguga_01g00289 | 5 | 0 |
| TpMuguga_01g00289 | 6 | 0.972881356 |
| TpMuguga_01g00292 | 1 | 0.073255814 |
| TpMuguga_01g00292 | 2 | 0.027131783 |
| TpMuguga_01g00292 | 3 | 0.002713178 |
| TpMuguga_01g00292 | 4 | 0.002713178 |
| TpMuguga_01g00292 | 5 | 0 |
| TpMuguga_01g00292 | 6 | 0 |
| TpMuguga_01g00291 | 1 | 0 |
| TpMuguga_01g00291 | 2 | 0.913461538 |
| TpMuguga_01g00291 | 3 | 1.346153846 |
| TpMuguga_01g00291 | 4 | 1.634615385 |
| TpMuguga_01g02155 | 1 | 0 |
| TpMuguga_01g02685 | 1 | 0.452674897 |
| TpMuguga_01g02685 | 2 | 0 |
| TpMuguga_01g02685 | 3 | 0 |
| TpMuguga_01g02685 | 4 | 0 |
| TpMuguga_01g02685 | 5 | 0 |
| TpMuguga_01g02685 | 6 | 0.271604938 |
| TpMuguga_01g02685 | 7 | 0.090534979 |
| TpMuguga_01g02685 | 8 | 0 |
| TpMuguga_01g02685 | 9 | 0.271604938 |
| TpMuguga_01g02685 | 10 | 2.263374486 |

| | | |
|---|---|---|
| TpMuguga_01g00297 | 1 | 0.146341463 |
| TpMuguga_01g00297 | 2 | 0.054878049 |
| TpMuguga_01g02875 | 1 | 0 |
| TpMuguga_01g02875 | 2 | 0 |
| TpMuguga_01g02875 | 3 | 0 |
| TpMuguga_01g02875 | 4 | 0 |
| TpMuguga_01g02875 | 5 | 0 |
| TpMuguga_01g02875 | 6 | 0 |
| TpMuguga_01g02875 | 7 | 0 |
| TpMuguga_01g02875 | 8 | 0 |
| TpMuguga_01g00301 | 1 | 0.666666667 |
| TpMuguga_01g00301 | 2 | 0 |
| TpMuguga_01g00301 | 3 | 0 |
| TpMuguga_01g00301 | 4 | 2 |
| TpMuguga_01g00301 | 5 | 5.333333333 |
| TpMuguga_01g00302 | 1 | 0 |
| TpMuguga_01g00302 | 2 | 0 |
| TpMuguga_01g00302 | 3 | 5.925925926 |
| TpMuguga_01g00302 | 4 | 0.592592593 |
| TpMuguga_01g00302 | 5 | 0 |
| TpMuguga_01g00302 | 6 | 6.518518519 |
| TpMuguga_01g00302 | 7 | 0 |
| TpMuguga_01g00304 | 1 | 0 |
| TpMuguga_01g00304 | 2 | 0 |
| TpMuguga_01g00304 | 3 | 0 |
| TpMuguga_01g00303 | 1 | 0 |
| TpMuguga_01g00303 | 2 | 0 |
| TpMuguga_01g00303 | 3 | 0 |
| TpMuguga_01g00303 | 4 | 0 |
| TpMuguga_01g00303 | 5 | 0 |
| TpMuguga_01g00303 | 6 | 1.230769231 |
| TpMuguga_01g00303 | 7 | 0 |
| TpMuguga_01g02445 | 1 | 0 |
| TpMuguga_01g02445 | 2 | 0 |
| TpMuguga_01g00305 | 1 | 0 |
| TpMuguga_01g00305 | 2 | 0.166666667 |
| TpMuguga_01g00305 | 3 | 0 |
| TpMuguga_01g00305 | 4 | 0 |
| TpMuguga_01g00308 | 1 | 0.264150943 |
| TpMuguga_01g00308 | 2 | 0 |
| TpMuguga_01g00308 | 3 | 0 |
| TpMuguga_01g00308 | 4 | 0 |

| | | |
|---|---|---|
| TpMuguga_01g00308 | 5 | 0 |
| TpMuguga_01g00308 | 6 | 1.188679245 |
| TpMuguga_01g02410 | 1 | 0 |
| TpMuguga_01g02410 | 2 | 0 |
| TpMuguga_01g02410 | 3 | 0 |
| TpMuguga_01g02410 | 4 | 0 |
| TpMuguga_01g02410 | 5 | 0 |
| TpMuguga_01g00313 | 1 | 1.666666667 |
| TpMuguga_01g00313 | 2 | 0 |
| TpMuguga_01g00313 | 3 | 3.333333333 |
| TpMuguga_01g00313 | 4 | 0 |
| TpMuguga_01g00313 | 5 | 1.666666667 |
| TpMuguga_01g00313 | 6 | 0 |
| TpMuguga_01g00313 | 7 | 0 |
| TpMuguga_01g00313 | 8 | 8.333333333 |
| TpMuguga_01g00313 | 9 | 6.666666667 |
| TpMuguga_01g00315 | 1 | 0.143153049 |
| TpMuguga_01g00314 | 1 | 0 |
| TpMuguga_01g00314 | 2 | 0 |
| TpMuguga_01g00314 | 3 | 0 |
| TpMuguga_01g00314 | 4 | 0 |
| TpMuguga_01g00314 | 5 | 0 |
| TpMuguga_01g00314 | 6 | 0.103896104 |
| TpMuguga_01g00314 | 7 | 11.01298701 |
| TpMuguga_01g00316 | 1 | 0 |
| TpMuguga_01g00316 | 2 | 1.961538462 |
| TpMuguga_01g00316 | 3 | 0 |
| TpMuguga_01g00316 | 4 | 2.615384615 |
| TpMuguga_01g00316 | 5 | 0 |
| TpMuguga_01g00316 | 6 | 0 |
| TpMuguga_01g00316 | 7 | 0 |
| TpMuguga_01g00316 | 8 | 0 |
| TpMuguga_01g00316 | 9 | 0 |
| TpMuguga_01g00316 | 10 | 0 |
| TpMuguga_01g00316 | 11 | 0 |
| TpMuguga_01g00316 | 12 | 0 |
| TpMuguga_01g00316 | 13 | 0 |
| TpMuguga_01g00316 | 14 | 0 |
| TpMuguga_01g00316 | 15 | 0 |
| TpMuguga_01g00316 | 16 | 0 |
| TpMuguga_01g02145 | 1 | 6.08 |
| TpMuguga_01g02145 | 2 | 0 |

| | | |
|---|---|---|
| TpMuguga_01g02145 | 3 | 1.92 |
| TpMuguga_01g02145 | 4 | 0 |
| TpMuguga_01g02145 | 5 | 0 |
| TpMuguga_01g02145 | 6 | 0 |
| TpMuguga_01g02145 | 7 | 0 |
| TpMuguga_01g00320 | 1 | 0 |
| TpMuguga_01g00320 | 2 | 0.132231405 |
| TpMuguga_01g00320 | 3 | 0.867768595 |
| TpMuguga_01g00320 | 4 | 1.900826446 |
| TpMuguga_01g00322 | 1 | 1.86407767 |
| TpMuguga_01g00321 | 1 | 0.272727273 |
| TpMuguga_01g00321 | 2 | 0 |
| TpMuguga_01g00323 | 1 | 0 |
| TpMuguga_01g00324 | 1 | 0.595813205 |
| TpMuguga_01g00324 | 2 | 0.990338164 |
| TpMuguga_01g00324 | 3 | 0.748792271 |
| TpMuguga_01g00324 | 4 | 0.845410628 |
| TpMuguga_01g00326 | 1 | 0.818181818 |
| TpMuguga_01g00326 | 2 | 0.636363636 |
| TpMuguga_01g00327 | 1 | 0 |
| TpMuguga_01g00327 | 2 | 0 |
| TpMuguga_01g00327 | 3 | 0 |
| TpMuguga_01g00327 | 4 | 2.003831418 |
| TpMuguga_01g00329 | 1 | 2.683585313 |
| TpMuguga_01g00329 | 2 | 1.754859611 |
| TpMuguga_01g00329 | 3 | 1.668466523 |
| TpMuguga_01g00329 | 4 | 0.998920086 |
| TpMuguga_01g00330 | 1 | 0 |
| TpMuguga_01g00330 | 2 | 1.596774194 |
| TpMuguga_01g00330 | 3 | 9.798387097 |
| TpMuguga_01g00330 | 4 | 14.80645161 |
| TpMuguga_01g00330 | 5 | 0.14516129 |
| TpMuguga_01g00330 | 6 | 0.217741935 |
| TpMuguga_01g00330 | 7 | 0 |
| TpMuguga_01g00330 | 8 | 0.580645161 |
| TpMuguga_01g00331 | 1 | 0.279069767 |
| TpMuguga_01g00331 | 2 | 0.948837209 |
| TpMuguga_01g00332 | 1 | 0 |
| TpMuguga_01g00332 | 2 | 0 |
| TpMuguga_01g00332 | 3 | 0 |
| TpMuguga_01g00332 | 4 | 0 |
| TpMuguga_01g00332 | 5 | 0 |

| | | |
|---|---|---|
| TpMuguga_01g00333 | 1 | 0 |
| TpMuguga_01g00333 | 2 | 0 |
| TpMuguga_01g00335 | 1 | 0 |
| TpMuguga_01g00335 | 2 | 0 |
| TpMuguga_01g00335 | 3 | 0 |
| TpMuguga_01g00335 | 4 | 0 |
| TpMuguga_01g00335 | 5 | 0 |
| TpMuguga_01g00335 | 6 | 0 |
| TpMuguga_01g00335 | 7 | 0 |
| TpMuguga_01g00336 | 1 | 0.147058824 |
| TpMuguga_01g00337 | 1 | 0.282982114 |
| TpMuguga_01g00339 | 1 | 3.012383901 |
| TpMuguga_01g00339 | 2 | 4.464396285 |
| TpMuguga_01g00339 | 3 | 0.606811146 |
| TpMuguga_01g00339 | 4 | 0.06501548 |
| TpMuguga_01g00339 | 5 | 0 |
| TpMuguga_01g00339 | 6 | 0.498452012 |
| TpMuguga_01g00340 | 1 | 0 |
| TpMuguga_01g00341 | 1 | 0.736196319 |
| TpMuguga_01g00341 | 2 | 0 |
| TpMuguga_01g00341 | 3 | 0 |
| TpMuguga_01g00341 | 4 | 0 |
| TpMuguga_01g00341 | 5 | 0 |
| TpMuguga_01g00341 | 6 | 0 |
| TpMuguga_01g00341 | 7 | 0 |
| TpMuguga_01g00343 | 1 | 0 |
| TpMuguga_01g00343 | 2 | 0 |
| TpMuguga_01g00343 | 3 | 0 |
| TpMuguga_01g00343 | 4 | 0 |
| TpMuguga_01g00343 | 5 | 0 |
| TpMuguga_01g00343 | 6 | 2.285714286 |
| TpMuguga_01g00343 | 7 | 6.857142857 |
| TpMuguga_01g00344 | 1 | 0 |
| TpMuguga_01g00344 | 2 | 0 |
| TpMuguga_01g00344 | 3 | 0 |
| TpMuguga_01g00344 | 4 | 0 |
| TpMuguga_01g00344 | 5 | 0 |
| TpMuguga_01g00344 | 6 | 0 |
| TpMuguga_01g00344 | 7 | 0 |
| TpMuguga_01g00346 | 1 | 1.395 |
| TpMuguga_01g00346 | 2 | 1.5 |
| TpMuguga_01g00347 | 1 | 0 |

| | | |
|---|---|---|
| TpMuguga_01g00347 | 2 | 0 |
| TpMuguga_01g00347 | 3 | 1.666666667 |
| TpMuguga_01g00347 | 4 | 4.444444444 |
| TpMuguga_01g00350 | 1 | 0.228571429 |
| TpMuguga_01g00350 | 2 | 0.457142857 |
| TpMuguga_01g00350 | 3 | 0 |
| TpMuguga_01g00350 | 4 | 0 |
| TpMuguga_01g00350 | 5 | 0 |
| TpMuguga_01g00350 | 6 | 0.685714286 |
| TpMuguga_01g00350 | 7 | 0 |
| TpMuguga_01g00349 | 1 | 0.909090909 |
| TpMuguga_01g00349 | 2 | 1.272727273 |
| TpMuguga_01g00349 | 3 | 0.545454545 |
| TpMuguga_01g00352 | 1 | 2.2 |
| TpMuguga_01g00352 | 2 | 0 |
| TpMuguga_01g02505 | 1 | 1.129032258 |
| TpMuguga_01g02505 | 2 | 0.64516129 |
| TpMuguga_01g02505 | 3 | 0 |
| TpMuguga_01g02505 | 4 | 0 |
| TpMuguga_01g02505 | 5 | 0 |
| TpMuguga_01g02505 | 6 | 0 |
| TpMuguga_01g02505 | 7 | 0 |
| TpMuguga_01g02505 | 8 | 0 |
| TpMuguga_01g02505 | 9 | 0 |
| TpMuguga_01g00356 | 1 | 0 |
| TpMuguga_01g00356 | 2 | 0 |
| TpMuguga_01g00356 | 3 | 0 |
| TpMuguga_01g00356 | 4 | 0 |
| TpMuguga_01g00357 | 1 | 25.5 |
| TpMuguga_01g00357 | 2 | 0 |
| TpMuguga_01g00358 | 1 | 0 |
| TpMuguga_01g00358 | 2 | 0 |
| TpMuguga_01g00358 | 3 | 0 |
| TpMuguga_01g02005 | 1 | 0 |
| TpMuguga_01g02005 | 2 | 0 |
| TpMuguga_01g02005 | 3 | 0 |
| TpMuguga_01g02005 | 4 | 0 |
| TpMuguga_01g02005 | 5 | 0 |
| TpMuguga_01g02005 | 6 | 0 |
| TpMuguga_01g02005 | 7 | 0 |
| TpMuguga_01g00362 | 1 | 0 |
| TpMuguga_01g00362 | 2 | 0 |

| | | |
|---|---|---|
| TpMuguga_01g00362 | 3 | 0 |
| TpMuguga_01g00362 | 4 | 0 |
| TpMuguga_01g00362 | 5 | 0 |
| TpMuguga_01g00362 | 6 | 0 |
| TpMuguga_01g00362 | 7 | 0 |
| TpMuguga_01g00362 | 8 | 0 |
| TpMuguga_01g00362 | 9 | 0 |
| TpMuguga_01g00364 | 1 | 0 |
| TpMuguga_01g00364 | 2 | 0 |
| TpMuguga_01g00364 | 3 | 5 |
| TpMuguga_01g00364 | 4 | 0 |
| TpMuguga_01g00365 | 1 | 2.25 |
| TpMuguga_01g00366 | 1 | 0 |
| TpMuguga_01g00366 | 2 | 0 |
| TpMuguga_01g00366 | 3 | 13 |
| TpMuguga_01g00366 | 4 | 0 |
| TpMuguga_01g00366 | 5 | 0 |
| TpMuguga_01g00366 | 6 | 0 |
| TpMuguga_01g00366 | 7 | 0 |
| TpMuguga_01g00366 | 8 | 0 |
| TpMuguga_01g00366 | 9 | 0 |
| TpMuguga_01g00366 | 10 | 0 |
| TpMuguga_01g00366 | 11 | 0 |
| TpMuguga_01g00366 | 12 | 0 |
| TpMuguga_01g00368 | 1 | 0 |
| TpMuguga_01g00368 | 2 | 0 |
| TpMuguga_01g00368 | 3 | 0 |
| TpMuguga_01g00368 | 4 | 0 |
| TpMuguga_01g00368 | 5 | 0 |
| TpMuguga_01g00368 | 6 | 0 |
| TpMuguga_01g00368 | 7 | 0 |
| TpMuguga_01g00368 | 8 | 0 |
| TpMuguga_01g00369 | 1 | 6 |
| TpMuguga_01g00369 | 2 | 0 |
| TpMuguga_01g00369 | 3 | 0 |
| TpMuguga_01g00369 | 4 | 1.8 |
| TpMuguga_01g00369 | 5 | 0 |
| TpMuguga_01g00370 | 1 | 0 |
| TpMuguga_01g00370 | 2 | 0 |
| TpMuguga_01g00370 | 3 | 0 |
| TpMuguga_01g00370 | 4 | 0 |
| TpMuguga_01g00370 | 5 | 0 |

| | | |
|---|---|---|
| TpMuguga_01g00370 | 6 | 0 |
| TpMuguga_01g00370 | 7 | 0 |
| TpMuguga_01g00370 | 8 | 0 |
| TpMuguga_01g00370 | 9 | 0 |
| TpMuguga_01g00370 | 10 | 0 |
| TpMuguga_01g02775 | 1 | 0 |
| TpMuguga_01g02775 | 2 | 0 |
| TpMuguga_01g02775 | 3 | 0 |
| TpMuguga_01g02775 | 4 | 0 |
| TpMuguga_01g02775 | 5 | 0 |
| TpMuguga_01g02775 | 6 | 0 |
| TpMuguga_01g02775 | 7 | 0 |
| TpMuguga_01g00373 | 1 | 0 |
| TpMuguga_01g00374 | 1 | 0 |
| TpMuguga_01g00374 | 2 | 0 |
| TpMuguga_01g00374 | 3 | 0 |
| TpMuguga_01g00374 | 4 | 0 |
| TpMuguga_01g00374 | 5 | 0 |
| TpMuguga_01g00374 | 6 | 0 |
| TpMuguga_01g00374 | 7 | 0 |
| TpMuguga_01g00374 | 8 | 0 |
| TpMuguga_01g00374 | 9 | 0 |
| TpMuguga_01g00374 | 10 | 0 |
| TpMuguga_01g00375 | 1 | 0 |
| TpMuguga_01g00375 | 2 | 0 |
| TpMuguga_01g00375 | 3 | 0 |
| TpMuguga_01g00375 | 4 | 0 |
| TpMuguga_01g00375 | 5 | 0 |
| TpMuguga_01g00375 | 6 | 0.636363636 |
| TpMuguga_01g00377 | 1 | 0.388601036 |
| TpMuguga_01g00377 | 2 | 0 |
| TpMuguga_01g00377 | 3 | 0 |
| TpMuguga_01g00377 | 4 | 0.093264249 |
| TpMuguga_01g00377 | 5 | 0.21761658 |
| TpMuguga_01g00377 | 6 | 0.730569948 |
| TpMuguga_01g00377 | 7 | 2.067357513 |
| TpMuguga_01g00377 | 8 | 4.321243523 |
| TpMuguga_01g00376 | 1 | 0 |
| TpMuguga_01g00376 | 2 | 0 |
| TpMuguga_01g00376 | 3 | 0 |
| TpMuguga_01g00380 | 1 | 0 |
| TpMuguga_01g00381 | 1 | 0 |

| | | |
|---|---|---|
| TpMuguga_01g00381 | 2 | 0 |
| TpMuguga_01g00382 | 1 | 0 |
| TpMuguga_01g00382 | 2 | 0 |
| TpMuguga_01g02460 | 1 | 0 |
| TpMuguga_01g00385 | 1 | 0 |
| TpMuguga_01g00385 | 2 | 0 |
| TpMuguga_01g00385 | 3 | 0.266666667 |
| TpMuguga_01g00386 | 1 | 0 |
| TpMuguga_01g00386 | 2 | 0.108910891 |
| TpMuguga_01g00386 | 3 | 0 |
| TpMuguga_01g00386 | 4 | 0.290429043 |
| TpMuguga_01g00386 | 5 | 0 |
| TpMuguga_01g00386 | 6 | 0 |
| TpMuguga_01g00386 | 7 | 0 |
| TpMuguga_01g00386 | 8 | 0 |
| TpMuguga_01g00386 | 9 | 0 |
| TpMuguga_01g00386 | 10 | 0 |
| TpMuguga_01g02475 | 1 | 0 |
| TpMuguga_01g02475 | 2 | 0 |
| TpMuguga_01g02475 | 3 | 0 |
| TpMuguga_01g02475 | 4 | 0 |
| TpMuguga_01g02475 | 5 | 0 |
| TpMuguga_01g02475 | 6 | 0 |
| TpMuguga_01g02475 | 7 | 0 |
| TpMuguga_01g02475 | 8 | 0 |
| TpMuguga_01g02475 | 9 | 0 |
| TpMuguga_01g02475 | 10 | 0 |
| TpMuguga_01g02475 | 11 | 0 |
| TpMuguga_01g00389 | 1 | 0 |
| TpMuguga_01g00389 | 2 | 0 |
| TpMuguga_01g00390 | 1 | 0 |
| TpMuguga_01g00390 | 2 | 0 |
| TpMuguga_01g00390 | 3 | 0 |
| TpMuguga_01g00390 | 4 | 1.428571429 |
| TpMuguga_01g00391 | 1 | 0.130434783 |
| TpMuguga_01g00391 | 2 | 0.244565217 |
| TpMuguga_01g00394 | 1 | 0 |
| TpMuguga_01g00394 | 2 | 0 |
| TpMuguga_01g00394 | 3 | 0 |
| TpMuguga_01g00394 | 4 | 0 |
| TpMuguga_01g00394 | 5 | 0 |
| TpMuguga_01g00395 | 1 | 0.930232558 |

| | | |
|---|---|---|
| TpMuguga_01g00395 | 2 | 0.930232558 |
| TpMuguga_01g00395 | 3 | 0 |
| TpMuguga_01g00397 | 1 | 0.056338028 |
| TpMuguga_01g00398 | 1 | 1 |
| TpMuguga_01g00398 | 2 | 14 |
| TpMuguga_01g00398 | 3 | 0 |
| TpMuguga_01g00399 | 1 | 10.625 |
| TpMuguga_01g00399 | 2 | 1.875 |
| TpMuguga_01g00399 | 3 | 0 |
| TpMuguga_01g00399 | 4 | 3.75 |
| TpMuguga_01g00400 | 1 | 0 |
| TpMuguga_01g00400 | 2 | 0 |
| TpMuguga_01g00400 | 3 | 0 |
| TpMuguga_01g00400 | 4 | 0 |
| TpMuguga_01g00400 | 5 | 0 |
| TpMuguga_01g00400 | 6 | 0 |
| TpMuguga_01g00401 | 1 | 0 |
| TpMuguga_01g00402 | 1 | 0 |
| TpMuguga_01g00403 | 1 | 3.2 |
| TpMuguga_01g00403 | 2 | 0 |
| TpMuguga_01g00403 | 3 | 0 |
| TpMuguga_01g00405 | 1 | 0 |
| TpMuguga_01g00405 | 2 | 0 |
| TpMuguga_01g00405 | 3 | 0 |
| TpMuguga_01g02525 | 1 | 0 |
| TpMuguga_01g02525 | 2 | 0 |
| TpMuguga_01g02525 | 3 | 0 |
| TpMuguga_01g02525 | 4 | 0 |
| TpMuguga_01g02525 | 5 | 0 |
| TpMuguga_01g00409 | 1 | 0 |
| TpMuguga_01g00410 | 1 | 0 |
| TpMuguga_01g00410 | 2 | 0 |
| TpMuguga_01g02735 | 1 | 0 |
| TpMuguga_01g02735 | 2 | 0 |
| TpMuguga_01g02735 | 3 | 0 |
| TpMuguga_01g02735 | 4 | 0 |
| TpMuguga_01g02735 | 5 | 0 |
| TpMuguga_01g02735 | 6 | 0 |
| TpMuguga_01g02735 | 7 | 0 |
| TpMuguga_01g02735 | 8 | 0 |
| TpMuguga_01g02735 | 9 | 0 |
| TpMuguga_01g02735 | 10 | 0 |

| | | |
|---|---|---|
| TpMuguga_01g00413 | 1 | 0.727272727 |
| TpMuguga_01g00414 | 1 | 0 |
| TpMuguga_01g00414 | 2 | 2 |
| TpMuguga_01g00414 | 3 | 0 |
| TpMuguga_01g00414 | 4 | 0 |
| TpMuguga_01g00414 | 5 | 0 |
| TpMuguga_01g02865 | 1 | 0 |
| TpMuguga_01g02865 | 2 | 0 |
| TpMuguga_01g02865 | 3 | 0 |
| TpMuguga_01g02865 | 4 | 0 |
| TpMuguga_01g02865 | 5 | 0 |
| TpMuguga_01g02865 | 6 | 0 |
| TpMuguga_01g02865 | 7 | 0 |
| TpMuguga_01g02865 | 8 | 0 |
| TpMuguga_01g02865 | 9 | 0 |
| TpMuguga_01g02865 | 10 | 0 |
| TpMuguga_01g02040 | 1 | 0 |
| TpMuguga_01g02325 | 1 | 0.631578947 |
| TpMuguga_01g02325 | 2 | 0 |
| TpMuguga_01g02830 | 1 | 3.745033113 |
| TpMuguga_01g02830 | 2 | 0.238410596 |
| TpMuguga_01g00422 | 1 | 0 |
| TpMuguga_01g00422 | 2 | 0 |
| TpMuguga_01g00422 | 3 | 0 |
| TpMuguga_01g00422 | 4 | 0 |
| TpMuguga_01g00422 | 5 | 0 |
| TpMuguga_01g00422 | 6 | 2.823529412 |
| TpMuguga_01g00422 | 7 | 12.23529412 |
| TpMuguga_01g00421 | 1 | 0.083018868 |
| TpMuguga_01g02800 | 1 | 0 |
| TpMuguga_01g02800 | 2 | 2.230263158 |
| TpMuguga_01g00426 | 1 | 0 |
| TpMuguga_01g00425 | 1 | 0.363636364 |
| TpMuguga_01g00425 | 2 | 0.727272727 |
| TpMuguga_01g00425 | 3 | 0.727272727 |
| TpMuguga_01g02125 | 1 | 0 |
| TpMuguga_01g02125 | 2 | 0 |
| TpMuguga_01g00430 | 1 | 0 |
| TpMuguga_01g00430 | 2 | 0 |
| TpMuguga_01g00430 | 3 | 0 |
| TpMuguga_01g00432 | 1 | 6 |
| TpMuguga_01g00432 | 2 | 0 |

| | | |
|---|---|---|
| TpMuguga_01g00432 | 3 | 0 |
| TpMuguga_01g00432 | 4 | 48 |
| TpMuguga_01g00432 | 5 | 30 |
| TpMuguga_01g00433 | 1 | 12 |
| TpMuguga_01g00434 | 1 | 0 |
| TpMuguga_01g00434 | 2 | 0 |
| TpMuguga_01g00434 | 3 | 0 |
| TpMuguga_01g00435 | 1 | 2.666666667 |
| TpMuguga_01g00436 | 1 | 0 |
| TpMuguga_01g00436 | 2 | 0 |
| TpMuguga_01g00437 | 1 | 0.357142857 |
| TpMuguga_01g02100 | 1 | 0 |
| TpMuguga_01g02100 | 2 | 0 |
| TpMuguga_01g02100 | 3 | 0 |
| TpMuguga_01g02100 | 4 | 0.125 |
| TpMuguga_01g02100 | 5 | 0 |
| TpMuguga_01g02100 | 6 | 0.875 |
| TpMuguga_01g00440 | 1 | 0 |
| TpMuguga_01g00440 | 2 | 0 |
| TpMuguga_01g00440 | 3 | 0 |
| TpMuguga_01g00440 | 4 | 0 |
| TpMuguga_01g00440 | 5 | 0 |
| TpMuguga_01g00440 | 6 | 0 |
| TpMuguga_01g00440 | 7 | 0.326018809 |
| TpMuguga_01g00439 | 1 | 0 |
| TpMuguga_01g00439 | 2 | 0 |
| TpMuguga_01g00439 | 3 | 0 |
| TpMuguga_01g00439 | 4 | 0 |
| TpMuguga_01g00439 | 5 | 0 |
| TpMuguga_01g00439 | 6 | 0 |
| TpMuguga_01g02470 | 1 | 0 |
| TpMuguga_01g00443 | 1 | 0.143678161 |
| TpMuguga_01g00443 | 2 | 0 |
| TpMuguga_01g00443 | 3 | 1.149425287 |
| TpMuguga_01g00443 | 4 | 0.603448276 |
| TpMuguga_01g00445 | 1 | 0 |
| TpMuguga_01g00445 | 2 | 0.170886076 |
| TpMuguga_01g00445 | 3 | 0.028481013 |
| TpMuguga_01g00445 | 4 | 0 |
| TpMuguga_01g00445 | 5 | 0 |
| TpMuguga_01g00445 | 6 | 0 |
| TpMuguga_01g00445 | 7 | 0 |

| | | |
|---|---|---|
| TpMuguga_01g00445 | 8 | 0.626582278 |
| TpMuguga_01g00444 | 1 | 0 |
| TpMuguga_01g00444 | 2 | 6.5 |
| TpMuguga_01g00444 | 3 | 1.181818182 |
| TpMuguga_01g00444 | 4 | 4.136363636 |
| TpMuguga_01g00444 | 5 | 7.681818182 |
| TpMuguga_01g00444 | 6 | 3.545454545 |
| TpMuguga_01g00444 | 7 | 0.590909091 |
| TpMuguga_01g00444 | 8 | 1.772727273 |
| TpMuguga_01g00444 | 9 | 2.954545455 |
| TpMuguga_01g00444 | 10 | 2.363636364 |
| TpMuguga_01g00444 | 11 | 3.545454545 |
| TpMuguga_01g00444 | 12 | 0.590909091 |
| TpMuguga_01g01232 | 1 | 0.078431373 |
| TpMuguga_01g01232 | 2 | 0.215686275 |
| TpMuguga_01g01233 | 1 | 0.016064257 |
| TpMuguga_01g00456 | 1 | 0 |
| TpMuguga_01g00456 | 2 | 0 |
| TpMuguga_01g00456 | 3 | 0 |
| TpMuguga_01g00456 | 4 | 0 |
| TpMuguga_01g00456 | 5 | 0 |
| TpMuguga_01g00456 | 6 | 0 |
| TpMuguga_01g00456 | 7 | 0 |
| TpMuguga_01g00456 | 8 | 0 |
| TpMuguga_01g02000 | 1 | 3.375 |
| TpMuguga_01g02000 | 2 | 0 |
| TpMuguga_01g02000 | 3 | 0 |
| TpMuguga_01g02000 | 4 | 0 |
| TpMuguga_01g02000 | 5 | 0 |
| TpMuguga_01g00459 | 1 | 9.5 |
| TpMuguga_01g00459 | 2 | 0 |
| TpMuguga_01g00459 | 3 | 0 |
| TpMuguga_01g00459 | 4 | 0 |
| TpMuguga_01g00461 | 1 | 0 |
| TpMuguga_01g00466 | 1 | 0.866666667 |
| TpMuguga_01g00468 | 1 | 0 |
| TpMuguga_01g00468 | 2 | 9 |
| TpMuguga_01g00468 | 3 | 0 |
| TpMuguga_01g00468 | 4 | 0 |
| TpMuguga_01g00468 | 5 | 0 |
| TpMuguga_01g00468 | 6 | 0 |
| TpMuguga_01g00468 | 7 | 0 |

| | | |
|---|---|---|
| TpMuguga_01g00468 | 8 | 0 |
| TpMuguga_01g02430 | 1 | 0 |
| TpMuguga_01g02430 | 2 | 0 |
| TpMuguga_01g02490 | 1 | 0.271186441 |
| TpMuguga_01g02490 | 2 | 1.288135593 |
| TpMuguga_01g02490 | 3 | 1.186440678 |
| TpMuguga_01g00471 | 1 | 0 |
| TpMuguga_01g00471 | 2 | 0 |
| TpMuguga_01g00471 | 3 | 0 |
| TpMuguga_01g00471 | 4 | 5 |
| TpMuguga_01g00472 | 1 | 0 |
| TpMuguga_01g00472 | 2 | 0 |
| TpMuguga_01g00472 | 3 | 0 |
| TpMuguga_01g00473 | 1 | 0 |
| TpMuguga_01g00473 | 2 | 0 |
| TpMuguga_01g00473 | 3 | 0 |
| TpMuguga_01g00473 | 4 | 0 |
| TpMuguga_01g00474 | 1 | 4 |
| TpMuguga_01g00474 | 2 | 0 |
| TpMuguga_01g00474 | 3 | 0 |
| TpMuguga_01g00475 | 1 | 0 |
| TpMuguga_01g00475 | 2 | 0 |
| TpMuguga_01g00475 | 3 | 0 |
| TpMuguga_01g00475 | 4 | 0 |
| TpMuguga_01g00475 | 5 | 0 |
| TpMuguga_01g00476 | 1 | 0.5 |
| TpMuguga_01g00476 | 2 | 0 |
| TpMuguga_01g00477 | 1 | 1.878787879 |
| TpMuguga_01g00480 | 1 | 0 |
| TpMuguga_01g00480 | 2 | 0 |
| TpMuguga_01g00480 | 3 | 0 |
| TpMuguga_01g00481 | 1 | 1.588235294 |
| TpMuguga_01g00484 | 1 | 0 |
| TpMuguga_01g00484 | 2 | 2.5 |
| TpMuguga_01g00484 | 3 | 0 |
| TpMuguga_01g00484 | 4 | 0 |
| TpMuguga_01g02390 | 1 | 0.833333333 |
| TpMuguga_01g02390 | 2 | 0 |
| TpMuguga_01g02390 | 3 | 0 |
| TpMuguga_01g02390 | 4 | 0 |
| TpMuguga_01g00488 | 1 | 0 |
| TpMuguga_01g00488 | 2 | 0 |

| | | |
|---|---|---|
| TpMuguga_01g00488 | 3 | 0 |
| TpMuguga_01g00489 | 1 | 0.277777778 |
| TpMuguga_01g00490 | 1 | 0 |
| TpMuguga_01g00490 | 2 | 0 |
| TpMuguga_01g00490 | 3 | 0 |
| TpMuguga_01g02135 | 1 | 0 |
| TpMuguga_01g00495 | 1 | 0 |
| TpMuguga_01g00495 | 2 | 0.133333333 |
| TpMuguga_01g00495 | 3 | 0.133333333 |
| TpMuguga_01g00496 | 1 | 0 |
| TpMuguga_01g00496 | 2 | 0 |
| TpMuguga_01g00496 | 3 | 1 |
| TpMuguga_01g00496 | 4 | 13 |
| TpMuguga_01g00496 | 5 | 4.5 |
| TpMuguga_01g00496 | 6 | 1.5 |
| TpMuguga_01g00496 | 7 | 2 |
| TpMuguga_01g00496 | 8 | 2.5 |
| TpMuguga_01g00496 | 9 | 0 |
| TpMuguga_01g00496 | 10 | 0 |
| TpMuguga_01g00496 | 11 | 0 |
| TpMuguga_01g00498 | 1 | 0 |
| TpMuguga_01g00499 | 1 | 0 |
| TpMuguga_01g00499 | 2 | 0.631578947 |
| TpMuguga_01g00499 | 3 | 0 |
| TpMuguga_01g00499 | 4 | 0 |
| TpMuguga_01g00499 | 5 | 0 |
| TpMuguga_01g00499 | 6 | 0 |
| TpMuguga_01g00499 | 7 | 0 |
| TpMuguga_01g00499 | 8 | 0 |
| TpMuguga_01g00499 | 9 | 0 |
| TpMuguga_01g00499 | 10 | 9.473684211 |
| TpMuguga_01g00499 | 11 | 0.631578947 |
| TpMuguga_01g00500 | 1 | 1.567164179 |
| TpMuguga_01g00500 | 2 | 0.194029851 |
| TpMuguga_01g00501 | 1 | 0 |
| TpMuguga_01g00501 | 2 | 0 |
| TpMuguga_01g00502 | 1 | 0 |
| TpMuguga_01g00503 | 1 | 0 |
| TpMuguga_01g00503 | 2 | 0 |
| TpMuguga_01g00503 | 3 | 0 |
| TpMuguga_01g00503 | 4 | 0 |
| TpMuguga_01g00503 | 5 | 0 |

| | | |
|---|---|---|
| TpMuguga_01g00504 | 1 | 0 |
| TpMuguga_01g00504 | 2 | 1.03125 |
| TpMuguga_01g00504 | 3 | 0 |
| TpMuguga_01g00504 | 4 | 0 |
| TpMuguga_01g00504 | 5 | 0 |
| TpMuguga_01g00504 | 6 | 1.71875 |
| TpMuguga_01g00504 | 7 | 0 |
| TpMuguga_01g00504 | 8 | 0 |
| TpMuguga_01g00504 | 9 | 0.6875 |
| TpMuguga_01g00504 | 10 | 1.03125 |
| TpMuguga_01g00506 | 1 | 1.405405405 |
| TpMuguga_01g00506 | 2 | 0 |
| TpMuguga_01g00506 | 3 | 0 |
| TpMuguga_01g00507 | 1 | 0 |
| TpMuguga_01g00507 | 2 | 0 |
| TpMuguga_01g00507 | 3 | 0 |
| TpMuguga_01g00508 | 1 | 0 |
| TpMuguga_01g00509 | 1 | 0 |
| TpMuguga_01g00509 | 2 | 0 |
| TpMuguga_01g00509 | 3 | 0 |
| TpMuguga_01g00509 | 4 | 0.111111111 |
| TpMuguga_01g00509 | 5 | 0 |
| TpMuguga_01g00509 | 6 | 0.5 |
| TpMuguga_01g02850 | 1 | 0 |
| TpMuguga_01g02755 | 1 | 0.164705882 |
| TpMuguga_01g02755 | 2 | 0 |
| TpMuguga_01g02755 | 3 | 0 |
| TpMuguga_01g02755 | 4 | 0 |
| TpMuguga_01g02755 | 5 | 0.263529412 |
| TpMuguga_01g02755 | 6 | 4.595294118 |
| TpMuguga_01g00512 | 1 | 1.911111111 |
| TpMuguga_01g00512 | 2 | 0 |
| TpMuguga_01g00511 | 1 | 0 |
| TpMuguga_01g00511 | 2 | 0 |
| TpMuguga_01g00513 | 1 | 0 |
| TpMuguga_01g00513 | 2 | 7.8 |
| TpMuguga_01g00514 | 1 | 3.571428571 |
| TpMuguga_01g00514 | 2 | 1.428571429 |
| TpMuguga_01g00514 | 3 | 0.476190476 |
| TpMuguga_01g00514 | 4 | 0 |
| TpMuguga_01g00515 | 1 | 0 |
| TpMuguga_01g00515 | 2 | 0 |

| | | |
|---|---|---|
| TpMuguga_01g00515 | 3 | 0 |
| TpMuguga_01g00515 | 4 | 0 |
| TpMuguga_01g00515 | 5 | 0 |
| TpMuguga_01g00515 | 6 | 9.916666667 |
| TpMuguga_01g00515 | 7 | 0 |
| TpMuguga_01g00515 | 8 | 0 |
| TpMuguga_01g00515 | 9 | 0 |
| TpMuguga_01g00515 | 10 | 0 |
| TpMuguga_01g00515 | 11 | 2.333333333 |
| TpMuguga_01g00515 | 12 | 8.75 |
| TpMuguga_01g00515 | 13 | 4.666666667 |
| TpMuguga_01g00516 | 1 | 0 |
| TpMuguga_01g00517 | 1 | 0 |
| TpMuguga_01g00517 | 2 | 0 |
| TpMuguga_01g00517 | 3 | 20 |
| TpMuguga_01g00518 | 1 | 0 |
| TpMuguga_01g00518 | 2 | 0 |
| TpMuguga_01g00518 | 3 | 0 |
| TpMuguga_01g00518 | 4 | 0 |
| TpMuguga_01g00518 | 5 | 0 |
| TpMuguga_01g00518 | 6 | 0 |
| TpMuguga_01g00518 | 7 | 0 |
| TpMuguga_01g00518 | 8 | 0 |
| TpMuguga_01g00518 | 9 | 0 |
| TpMuguga_01g02640 | 1 | 0 |
| TpMuguga_01g02640 | 2 | 0 |
| TpMuguga_01g02640 | 3 | 0 |
| TpMuguga_01g00520 | 1 | 0 |
| TpMuguga_01g00520 | 2 | 0 |
| TpMuguga_01g00520 | 3 | 0 |
| TpMuguga_01g00520 | 4 | 7.14084507 |
| TpMuguga_01g00520 | 5 | 2.563380282 |
| TpMuguga_01g00520 | 6 | 0.183098592 |
| TpMuguga_01g00520 | 7 | 0 |
| TpMuguga_01g00520 | 8 | 0 |
| TpMuguga_01g00520 | 9 | 0 |
| TpMuguga_01g00520 | 10 | 0 |
| TpMuguga_01g00520 | 11 | 0.457746479 |
| TpMuguga_01g00520 | 12 | 1.830985915 |
| TpMuguga_01g00522 | 1 | 0 |
| TpMuguga_01g00522 | 2 | 0 |
| TpMuguga_01g00523 | 1 | 0.726804124 |

| | | |
|---|---|---|
| TpMuguga_01g00523 | 2 | 0.937113402 |
| TpMuguga_01g00524 | 1 | 0 |
| TpMuguga_01g00524 | 2 | 3.75 |
| TpMuguga_01g00525 | 1 | 0 |
| TpMuguga_01g02250 | 1 | 0 |
| TpMuguga_01g00526 | 1 | 0 |
| TpMuguga_01g00527 | 1 | 0 |
| TpMuguga_01g00527 | 2 | 0 |
| TpMuguga_01g00527 | 3 | 1.217391304 |
| TpMuguga_01g00528 | 1 | 0 |
| TpMuguga_01g00528 | 2 | 0 |
| TpMuguga_01g00528 | 3 | 0 |
| TpMuguga_01g00529 | 1 | 0.765987025 |
| TpMuguga_01g00529 | 2 | 0 |
| TpMuguga_01g00530 | 1 | 0.314606742 |
| TpMuguga_01g00532 | 1 | 0 |
| TpMuguga_01g00532 | 2 | 0.7 |
| TpMuguga_01g00534 | 1 | 0 |
| TpMuguga_01g00534 | 2 | 0 |
| TpMuguga_01g00535 | 1 | 0.620192308 |
| TpMuguga_01g00535 | 2 | 0.028846154 |
| TpMuguga_01g00536 | 1 | 9 |
| TpMuguga_01g00536 | 2 | 0 |
| TpMuguga_01g00539 | 1 | 0 |
| TpMuguga_01g00542 | 1 | 0 |
| TpMuguga_01g00542 | 2 | 0 |
| TpMuguga_01g00543 | 1 | 0.816326531 |
| TpMuguga_01g00543 | 2 | 0.816326531 |
| TpMuguga_01g00543 | 3 | 1.93877551 |
| TpMuguga_01g00543 | 4 | 1.632653061 |
| TpMuguga_01g00544 | 1 | 0 |
| TpMuguga_01g00544 | 2 | 0 |
| TpMuguga_01g00544 | 3 | 2.638888889 |
| TpMuguga_01g00544 | 4 | 0.694444444 |
| TpMuguga_01g00544 | 5 | 0.694444444 |
| TpMuguga_01g00544 | 6 | 2.777777778 |
| TpMuguga_01g00544 | 7 | 2.638888889 |
| TpMuguga_01g00544 | 8 | 3.055555556 |
| TpMuguga_01g00544 | 9 | 0 |
| TpMuguga_01g00544 | 10 | 0 |
| TpMuguga_01g00544 | 11 | 0 |
| TpMuguga_01g00544 | 12 | 0 |

| | | |
|---|---|---|
| TpMuguga_01g00544 | 13 | 1.388888889 |
| TpMuguga_01g00544 | 14 | 0.833333333 |
| TpMuguga_01g00545 | 1 | 0 |
| TpMuguga_01g00545 | 2 | 0 |
| TpMuguga_01g00546 | 1 | 0.000486381 |
| TpMuguga_01g01229 | 1 | 1.6 |
| TpMuguga_01g02515 | 1 | 0 |
| TpMuguga_01g00552 | 1 | 0 |
| TpMuguga_01g00552 | 2 | 0 |
| TpMuguga_01g00553 | 1 | 0.666666667 |
| TpMuguga_01g00554 | 1 | 0.611111111 |
| TpMuguga_01g00554 | 2 | 0.740740741 |
| TpMuguga_01g00557 | 1 | 0.018115942 |
| TpMuguga_01g00557 | 2 | 0 |
| TpMuguga_01g00557 | 3 | 0 |
| TpMuguga_01g00557 | 4 | 0 |
| TpMuguga_01g00556 | 1 | 0 |
| TpMuguga_01g00556 | 2 | 0.014040014 |
| TpMuguga_01g00556 | 3 | 0 |
| TpMuguga_01g00556 | 4 | 0 |
| TpMuguga_01g00555 | 1 | 0.214187328 |
| TpMuguga_01g00559 | 1 | 1.714285714 |
| TpMuguga_01g00559 | 2 | 0 |
| TpMuguga_01g00560 | 1 | 0.714285714 |
| TpMuguga_01g00562 | 1 | 0.829268293 |
| TpMuguga_01g00563 | 1 | 1.846153846 |
| TpMuguga_01g00565 | 1 | 6.84 |
| TpMuguga_01g00565 | 2 | 0 |
| TpMuguga_01g00565 | 3 | 0.72 |
| TpMuguga_01g00565 | 4 | 1.08 |
| TpMuguga_01g00565 | 5 | 0 |
| TpMuguga_01g00565 | 6 | 0.18 |
| TpMuguga_01g00565 | 7 | 0 |
| TpMuguga_01g00565 | 8 | 0 |
| TpMuguga_01g00566 | 1 | 1.330024814 |
| TpMuguga_01g00566 | 2 | 1.46898263 |
| TpMuguga_01g00566 | 3 | 1.836228288 |
| TpMuguga_01g00568 | 1 | 0 |
| TpMuguga_01g00568 | 2 | 0.056939502 |
| TpMuguga_01g00568 | 3 | 0 |
| TpMuguga_01g00567 | 1 | 0 |
| TpMuguga_01g00567 | 2 | 0 |

| | | |
|---|---|---|
| TpMuguga_01g00567 | 3 | 0.274566474 |
| TpMuguga_01g00567 | 4 | 4.26300578 |
| TpMuguga_01g00569 | 1 | 0 |
| TpMuguga_01g00569 | 2 | 0 |
| TpMuguga_01g00569 | 3 | 0 |
| TpMuguga_01g00569 | 4 | 1.434782609 |
| TpMuguga_01g00569 | 5 | 0 |
| TpMuguga_01g00569 | 6 | 0 |
| TpMuguga_01g00569 | 7 | 0 |
| TpMuguga_01g00569 | 8 | 0 |
| TpMuguga_01g00569 | 9 | 0.956521739 |
| TpMuguga_01g00569 | 10 | 6.217391304 |
| TpMuguga_01g00570 | 1 | 0 |
| TpMuguga_01g00570 | 2 | 0 |
| TpMuguga_01g02555 | 1 | 0 |
| TpMuguga_01g00573 | 1 | 0.06557377 |
| TpMuguga_01g02580 | 1 | 1.270588235 |
| TpMuguga_01g00574 | 1 | 0.114359609 |
| TpMuguga_01g00574 | 2 | 0.019903295 |
| TpMuguga_01g00575 | 1 | 0 |
| TpMuguga_01g02270 | 1 | 0 |
| TpMuguga_01g02010 | 1 | 0 |
| TpMuguga_01g02010 | 2 | 0.535135135 |
| TpMuguga_01g02820 | 1 | 0 |
| TpMuguga_01g01230 | 1 | 0.465116279 |
| TpMuguga_01g01230 | 2 | 0 |
| TpMuguga_01g01230 | 3 | 0 |
| TpMuguga_01g01230 | 4 | 0 |
| TpMuguga_01g01230 | 5 | 0 |
| TpMuguga_01g01230 | 6 | 10.69767442 |
| TpMuguga_01g01230 | 7 | 21.62790698 |
| TpMuguga_01g01230 | 8 | 0 |
| TpMuguga_01g01230 | 9 | 3.023255814 |
| TpMuguga_01g02360 | 1 | 0 |
| TpMuguga_01g02360 | 2 | 0 |
| TpMuguga_01g02360 | 3 | 0 |
| TpMuguga_01g01231 | 1 | 0 |
| TpMuguga_01g01231 | 2 | 0 |
| TpMuguga_01g01231 | 3 | 1 |
| TpMuguga_01g00578 | 1 | 0 |
| TpMuguga_01g00578 | 2 | 0 |
| TpMuguga_01g00580 | 1 | 0 |

| | | |
|---|---|---|
| TpMuguga_01g02210 | 1 | 1.166666667 |
| TpMuguga_01g00581 | 1 | 0 |
| TpMuguga_01g00581 | 2 | 0 |
| TpMuguga_01g00582 | 1 | 0 |
| TpMuguga_01g00582 | 2 | 0 |
| TpMuguga_01g00582 | 3 | 0 |
| TpMuguga_01g00582 | 4 | 0 |
| TpMuguga_01g00582 | 5 | 0.893617021 |
| TpMuguga_01g00582 | 6 | 0 |
| TpMuguga_01g00582 | 7 | 0 |
| TpMuguga_01g00582 | 8 | 0.893617021 |
| TpMuguga_01g00582 | 9 | 3.574468085 |
| TpMuguga_01g00582 | 10 | 0 |
| TpMuguga_01g00582 | 11 | 0 |
| TpMuguga_01g00582 | 12 | 0 |
| TpMuguga_01g00582 | 13 | 3.872340426 |
| TpMuguga_01g00584 | 1 | 0 |
| TpMuguga_01g00584 | 2 | 0 |
| TpMuguga_01g00584 | 3 | 0 |
| TpMuguga_01g00584 | 4 | 0 |
| TpMuguga_01g00585 | 1 | 0.714285714 |
| TpMuguga_01g00585 | 2 | 0 |
| TpMuguga_01g00585 | 3 | 0 |
| TpMuguga_01g00585 | 4 | 0 |
| TpMuguga_01g00587 | 1 | 0 |
| TpMuguga_01g00587 | 2 | 0 |
| TpMuguga_01g00587 | 3 | 0 |
| TpMuguga_01g00588 | 1 | 0 |
| TpMuguga_01g00588 | 2 | 0 |
| TpMuguga_01g00588 | 3 | 0 |
| TpMuguga_01g00589 | 1 | 0 |
| TpMuguga_01g00589 | 2 | 0 |
| TpMuguga_01g00589 | 3 | 0 |
| TpMuguga_01g02835 | 1 | 0 |
| TpMuguga_01g02835 | 2 | 0 |
| TpMuguga_01g02835 | 3 | 0 |
| TpMuguga_01g02835 | 4 | 0 |
| TpMuguga_01g02835 | 5 | 0 |
| TpMuguga_01g02835 | 6 | 0 |
| TpMuguga_01g00594 | 1 | 0.218211649 |
| TpMuguga_01g00594 | 2 | 0 |
| TpMuguga_01g00594 | 3 | 0.723543888 |

| | | |
|---|---|---|
| TpMuguga_01g00594 | 4 | 0.844134537 |
| TpMuguga_01g00594 | 5 | 0.114848236 |
| TpMuguga_01g00594 | 6 | 0.011484824 |
| TpMuguga_01g00595 | 1 | 0 |
| TpMuguga_01g00595 | 2 | 0 |
| TpMuguga_01g00595 | 3 | 15 |
| TpMuguga_01g00595 | 4 | 0 |
| TpMuguga_01g00596 | 1 | 0 |
| TpMuguga_01g00596 | 2 | 0 |
| TpMuguga_01g00596 | 3 | 0 |
| TpMuguga_01g00596 | 4 | 0 |
| TpMuguga_01g00596 | 5 | 0 |
| TpMuguga_01g00596 | 6 | 3.538461538 |
| TpMuguga_01g00597 | 1 | 0 |
| TpMuguga_01g00597 | 2 | 0 |
| TpMuguga_01g00598 | 1 | 0 |
| TpMuguga_01g00598 | 2 | 0 |
| TpMuguga_01g00598 | 3 | 0.569105691 |
| TpMuguga_01g00598 | 4 | 2.674796748 |
| TpMuguga_01g00598 | 5 | 0.227642276 |
| TpMuguga_01g00598 | 6 | 0 |
| TpMuguga_01g02510 | 1 | 0 |
| TpMuguga_01g02510 | 2 | 0 |
| TpMuguga_01g02510 | 3 | 0 |
| TpMuguga_01g02510 | 4 | 0 |
| TpMuguga_01g02510 | 5 | 0 |
| TpMuguga_01g02510 | 6 | 0 |
| TpMuguga_01g02510 | 7 | 0 |
| TpMuguga_01g02510 | 8 | 0 |
| TpMuguga_01g00601 | 1 | 0.972762646 |
| TpMuguga_01g00601 | 2 | 1.089494163 |
| TpMuguga_01g00601 | 3 | 1.381322957 |
| TpMuguga_01g00601 | 4 | 1.498054475 |
| TpMuguga_01g00601 | 5 | 1.070038911 |
| TpMuguga_01g00601 | 6 | 2.470817121 |
| TpMuguga_01g00601 | 7 | 2.996108949 |
| TpMuguga_01g00601 | 8 | 1.050583658 |
| TpMuguga_01g00601 | 9 | 4.6692607 |
| TpMuguga_01g00602 | 1 | 0.058287796 |
| TpMuguga_01g00606 | 1 | 1.125 |
| TpMuguga_01g00622 | 1 | 0 |
| TpMuguga_01g00622 | 2 | 0 |

| | | |
|---|---|---|
| TpMuguga_01g00622 | 3 | 0.2 |
| TpMuguga_01g00622 | 4 | 6.6 |
| TpMuguga_01g00624 | 1 | 1.163636364 |
| TpMuguga_01g00624 | 2 | 0.290909091 |
| TpMuguga_01g00624 | 3 | 0.072727273 |
| TpMuguga_01g00625 | 1 | 0 |
| TpMuguga_01g00625 | 2 | 0 |
| TpMuguga_01g00625 | 3 | 0 |
| TpMuguga_01g00625 | 4 | 0 |
| TpMuguga_01g00626 | 1 | 0 |
| TpMuguga_01g00626 | 2 | 0 |
| TpMuguga_01g00626 | 3 | 0 |
| TpMuguga_01g00626 | 4 | 0.054995417 |
| TpMuguga_01g02825 | 1 | 0.002290951 |
| TpMuguga_01g02825 | 2 | 0.041237113 |
| TpMuguga_01g02825 | 3 | 0.006872852 |
| TpMuguga_01g00630 | 1 | 0 |
| TpMuguga_01g00630 | 2 | 0 |
| TpMuguga_01g00630 | 3 | 0 |
| TpMuguga_01g00632 | 1 | 0.9 |
| TpMuguga_01g00632 | 2 | 0 |
| TpMuguga_01g00633 | 1 | 0 |
| TpMuguga_01g00633 | 2 | 0 |
| TpMuguga_01g00633 | 3 | 0 |
| TpMuguga_01g00633 | 4 | 0 |
| TpMuguga_01g00634 | 1 | 0.249201278 |
| TpMuguga_01g00634 | 2 | 0.095846645 |
| TpMuguga_01g00634 | 3 | 0.325878594 |
| TpMuguga_01g00634 | 4 | 0.095846645 |
| TpMuguga_01g00634 | 5 | 0.076677316 |
| TpMuguga_01g00635 | 1 | 1.296296296 |
| TpMuguga_01g00635 | 2 | 0.24691358 |
| TpMuguga_01g00635 | 3 | 0.679012346 |
| TpMuguga_01g00635 | 4 | 0.185185185 |
| TpMuguga_01g00635 | 5 | 0.555555556 |
| TpMuguga_01g00635 | 6 | 0.679012346 |
| TpMuguga_01g00635 | 7 | 0 |
| TpMuguga_01g00635 | 8 | 0 |
| TpMuguga_01g00635 | 9 | 1.481481481 |
| TpMuguga_01g02540 | 1 | 0 |
| TpMuguga_01g02220 | 1 | 0 |
| TpMuguga_01g02220 | 2 | 0 |

| | | |
|---|---|---|
| TpMuguga_01g02220 | 3 | 0 |
| TpMuguga_01g02225 | 1 | 0 |
| TpMuguga_01g02225 | 2 | 0 |
| TpMuguga_01g02225 | 3 | 0 |
| TpMuguga_01g02225 | 4 | 0 |
| TpMuguga_01g02225 | 5 | 0 |
| TpMuguga_01g00639 | 1 | 0 |
| TpMuguga_01g00639 | 2 | 9.333333333 |
| TpMuguga_01g00639 | 3 | 0 |
| TpMuguga_01g00639 | 4 | 0 |
| TpMuguga_01g00639 | 5 | 0 |
| TpMuguga_01g00639 | 6 | 2.333333333 |
| TpMuguga_01g00641 | 1 | 2.5 |
| TpMuguga_01g00641 | 2 | 0 |
| TpMuguga_01g00641 | 3 | 0 |
| TpMuguga_01g00641 | 4 | 0 |
| TpMuguga_01g00642 | 1 | 0 |
| TpMuguga_01g00642 | 2 | 3.818181818 |
| TpMuguga_01g00642 | 3 | 0 |
| TpMuguga_01g00642 | 4 | 1.636363636 |
| TpMuguga_01g00642 | 5 | 0.545454545 |
| TpMuguga_01g02450 | 1 | 0 |
| TpMuguga_01g02450 | 2 | 0 |
| TpMuguga_01g02450 | 3 | 0 |
| TpMuguga_01g02450 | 4 | 0 |
| TpMuguga_01g02450 | 5 | 0 |
| TpMuguga_01g02450 | 6 | 0 |
| TpMuguga_01g02450 | 7 | 0 |
| TpMuguga_01g02450 | 8 | 0 |
| TpMuguga_01g02450 | 9 | 0 |
| TpMuguga_01g02450 | 10 | 0 |
| TpMuguga_01g02450 | 11 | 0 |
| TpMuguga_01g00648 | 1 | 1.826086957 |
| TpMuguga_01g00648 | 2 | 0 |
| TpMuguga_01g00649 | 1 | 0 |
| TpMuguga_01g00649 | 2 | 0 |
| TpMuguga_01g00649 | 3 | 0 |
| TpMuguga_01g02350 | 1 | 0 |
| TpMuguga_01g02350 | 2 | 0 |
| TpMuguga_01g02350 | 3 | 3.571428571 |
| TpMuguga_01g02350 | 4 | 0 |
| TpMuguga_01g02650 | 1 | 0.363636364 |

| | | |
|---|---|---|
| TpMuguga_01g00652 | 1 | 0 |
| TpMuguga_01g00652 | 2 | 0 |
| TpMuguga_01g00652 | 3 | 5.9375 |
| TpMuguga_01g00652 | 4 | 9.375 |
| TpMuguga_01g00653 | 1 | 0 |
| TpMuguga_01g00653 | 2 | 0 |
| TpMuguga_01g00654 | 1 | 0 |
| TpMuguga_01g00655 | 1 | 0 |
| TpMuguga_01g00655 | 2 | 0 |
| TpMuguga_01g00655 | 3 | 0 |
| TpMuguga_01g00655 | 4 | 0 |
| TpMuguga_01g00655 | 5 | 0 |
| TpMuguga_01g00655 | 6 | 0 |
| TpMuguga_01g00655 | 7 | 0 |
| TpMuguga_01g00655 | 8 | 0 |
| TpMuguga_01g00655 | 9 | 0 |
| TpMuguga_01g00657 | 1 | 0 |
| TpMuguga_01g00657 | 2 | 0.818181818 |
| TpMuguga_01g02535 | 1 | 1.125 |
| TpMuguga_01g02535 | 2 | 0 |
| TpMuguga_01g00658 | 1 | 0 |
| TpMuguga_01g00661 | 1 | 0 |
| TpMuguga_01g00661 | 2 | 0 |
| TpMuguga_01g00661 | 3 | 0 |
| TpMuguga_01g00661 | 4 | 0 |
| TpMuguga_01g00660 | 1 | 0 |
| TpMuguga_01g00663 | 1 | 0 |
| TpMuguga_01g00663 | 2 | 0 |
| TpMuguga_01g00663 | 3 | 0 |
| TpMuguga_01g00663 | 4 | 12.5 |
| TpMuguga_01g00663 | 5 | 7 |
| TpMuguga_01g00664 | 1 | 0 |
| TpMuguga_01g00664 | 2 | 1.044776119 |
| TpMuguga_01g00664 | 3 | 2.910447761 |
| TpMuguga_01g00664 | 4 | 1.902985075 |
| TpMuguga_01g00665 | 1 | 0 |
| TpMuguga_01g00665 | 2 | 0 |
| TpMuguga_01g00665 | 3 | 0 |
| TpMuguga_01g00665 | 4 | 0 |
| TpMuguga_01g00665 | 5 | 0 |
| TpMuguga_01g00666 | 1 | 0 |
| TpMuguga_01g00666 | 2 | 0 |

| | | |
|---|---|---|
| TpMuguga_01g00666 | 3 | 0 |
| TpMuguga_01g00666 | 4 | 0 |
| TpMuguga_01g00666 | 5 | 0 |
| TpMuguga_01g00666 | 6 | 0 |
| TpMuguga_01g00666 | 7 | 0 |
| TpMuguga_01g00666 | 8 | 0 |
| TpMuguga_01g00666 | 9 | 0 |
| TpMuguga_01g00666 | 10 | 0 |
| TpMuguga_01g00666 | 11 | 2.8 |
| TpMuguga_01g00666 | 12 | 0 |
| TpMuguga_01g00666 | 13 | 0 |
| TpMuguga_01g00667 | 1 | 3 |
| TpMuguga_01g00667 | 2 | 0 |
| TpMuguga_01g00668 | 1 | 0 |
| TpMuguga_01g00668 | 2 | 0 |
| TpMuguga_01g00668 | 3 | 2.5 |
| TpMuguga_01g00668 | 4 | 2.5 |
| TpMuguga_01g00669 | 1 | 0 |
| TpMuguga_01g00671 | 1 | 0.828947368 |
| TpMuguga_01g00671 | 2 | 0.184210526 |
| TpMuguga_01g00671 | 3 | 0 |
| TpMuguga_01g00671 | 4 | 0 |
| TpMuguga_01g00671 | 5 | 0.092105263 |
| TpMuguga_01g00671 | 6 | 0.092105263 |
| TpMuguga_01g00672 | 1 | 1 |
| TpMuguga_01g00672 | 2 | 6.333333333 |
| TpMuguga_01g00673 | 1 | 0.5 |
| TpMuguga_01g02495 | 1 | 0 |
| TpMuguga_01g02495 | 2 | 3.75 |
| TpMuguga_01g02495 | 3 | 0 |
| TpMuguga_01g02495 | 4 | 5 |
| TpMuguga_01g02050 | 1 | 0.147058824 |
| TpMuguga_01g02050 | 2 | 0.147058824 |
| TpMuguga_01g02050 | 3 | 2.941176471 |
| TpMuguga_01g02050 | 4 | 0.441176471 |
| TpMuguga_01g00675 | 1 | 0.24 |
| TpMuguga_01g00675 | 2 | 0 |
| TpMuguga_01g00675 | 3 | 0.04 |
| TpMuguga_01g00676 | 1 | 0 |
| TpMuguga_01g00676 | 2 | 0 |
| TpMuguga_01g00676 | 3 | 0 |
| TpMuguga_01g00676 | 4 | 0 |

| | | |
|---|---|---|
| TpMuguga_01g00676 | 5 | 0 |
| TpMuguga_01g00678 | 1 | 0 |
| TpMuguga_01g00679 | 1 | 0 |
| TpMuguga_01g00679 | 2 | 0 |
| TpMuguga_01g00681 | 1 | 0 |
| TpMuguga_01g00681 | 2 | 0 |
| TpMuguga_01g00681 | 3 | 0 |
| TpMuguga_01g00681 | 4 | 0 |
| TpMuguga_01g00681 | 5 | 0 |
| TpMuguga_01g02130 | 1 | 0 |
| TpMuguga_01g02130 | 2 | 0 |
| TpMuguga_01g02130 | 3 | 0 |
| TpMuguga_01g02130 | 4 | 0 |
| TpMuguga_01g02130 | 5 | 0 |
| TpMuguga_01g02150 | 1 | 0 |
| TpMuguga_01g02150 | 2 | 0 |
| TpMuguga_01g02150 | 3 | 0 |
| TpMuguga_01g02150 | 4 | 0 |
| TpMuguga_01g02150 | 5 | 23.14285714 |
| TpMuguga_01g02150 | 6 | 15.42857143 |
| TpMuguga_01g02150 | 7 | 0 |
| TpMuguga_01g02150 | 8 | 0 |
| TpMuguga_01g02160 | 1 | 0.782128514 |
| TpMuguga_01g02160 | 2 | 0.09939759 |
| TpMuguga_01g02420 | 1 | 8.68 |
| TpMuguga_01g02420 | 2 | 4.853333333 |
| TpMuguga_01g02420 | 3 | 0 |
| TpMuguga_01g02420 | 4 | 0 |
| TpMuguga_01g02420 | 5 | 0.186666667 |
| TpMuguga_01g02420 | 6 | 0.28 |
| TpMuguga_01g00687 | 1 | 6.470588235 |
| TpMuguga_01g00687 | 2 | 6.470588235 |
| TpMuguga_01g00687 | 3 | 9.705882353 |
| TpMuguga_01g00687 | 4 | 0 |
| TpMuguga_01g00687 | 5 | 0 |
| TpMuguga_01g00687 | 6 | 0 |
| TpMuguga_01g00687 | 7 | 0 |
| TpMuguga_01g00687 | 8 | 0 |
| TpMuguga_01g00687 | 9 | 0 |
| TpMuguga_01g00687 | 10 | 0 |
| TpMuguga_01g00688 | 1 | 0 |
| TpMuguga_01g00688 | 2 | 0 |

| | | |
|---|---|---|
| TpMuguga_01g00688 | 3 | 0 |
| TpMuguga_01g00688 | 4 | 0 |
| TpMuguga_01g00688 | 5 | 0 |
| TpMuguga_01g00689 | 1 | 0 |
| TpMuguga_01g00689 | 2 | 0 |
| TpMuguga_01g00689 | 3 | 0 |
| TpMuguga_01g00689 | 4 | 0 |
| TpMuguga_01g00689 | 5 | 0 |
| TpMuguga_01g00691 | 1 | 4 |
| TpMuguga_01g00691 | 2 | 4.571428571 |
| TpMuguga_01g00691 | 3 | 2.285714286 |
| TpMuguga_01g00692 | 1 | 3.545454545 |
| TpMuguga_01g00692 | 2 | 0.409090909 |
| TpMuguga_01g00692 | 3 | 0.136363636 |
| TpMuguga_01g00692 | 4 | 0 |
| TpMuguga_01g00692 | 5 | 0 |
| TpMuguga_01g00693 | 1 | 0 |
| TpMuguga_01g00695 | 1 | 0 |
| TpMuguga_01g00695 | 2 | 0 |
| TpMuguga_01g00695 | 3 | 0.857142857 |
| TpMuguga_01g00695 | 4 | 0 |
| TpMuguga_01g00695 | 5 | 0 |
| TpMuguga_01g00694 | 1 | 0.776699029 |
| TpMuguga_01g00694 | 2 | 0.291262136 |
| TpMuguga_01g00694 | 3 | 0.145631068 |
| TpMuguga_01g00694 | 4 | 0 |
| TpMuguga_01g00696 | 1 | 1.442307692 |
| TpMuguga_01g00696 | 2 | 0.115384615 |
| TpMuguga_01g00697 | 1 | 0 |
| TpMuguga_01g00697 | 2 | 0 |
| TpMuguga_01g00698 | 1 | 0 |
| TpMuguga_01g00698 | 2 | 40 |
| TpMuguga_01g00698 | 3 | 0 |
| TpMuguga_01g00698 | 4 | 0 |
| TpMuguga_01g00698 | 5 | 0 |
| TpMuguga_01g00698 | 6 | 0 |
| TpMuguga_01g00698 | 7 | 0 |
| TpMuguga_01g00698 | 8 | 20 |
| TpMuguga_01g00698 | 9 | 0 |
| TpMuguga_01g00699 | 1 | 0 |
| TpMuguga_01g00699 | 2 | 0 |
| TpMuguga_01g00699 | 3 | 0 |

| | | |
|---|---|---|
| TpMuguga_01g00699 | 4 | 0 |
| TpMuguga_01g00705 | 1 | 1.514563107 |
| TpMuguga_01g00705 | 2 | 0.233009709 |
| TpMuguga_01g00705 | 3 | 0.233009709 |
| TpMuguga_01g00705 | 4 | 1.45631068 |
| TpMuguga_01g00705 | 5 | 0.873786408 |
| TpMuguga_01g00706 | 1 | 0 |
| TpMuguga_01g00706 | 2 | 0 |
| TpMuguga_01g00706 | 3 | 0 |
| TpMuguga_01g00706 | 4 | 0 |
| TpMuguga_01g00706 | 5 | 0 |
| TpMuguga_01g00707 | 1 | 0 |
| TpMuguga_01g00707 | 2 | 0 |
| TpMuguga_01g00707 | 3 | 0 |
| TpMuguga_01g00707 | 4 | 0 |
| TpMuguga_01g00707 | 5 | 0 |
| TpMuguga_01g00707 | 6 | 0 |
| TpMuguga_01g00707 | 7 | 0 |
| TpMuguga_01g00707 | 8 | 6 |
| TpMuguga_01g00707 | 9 | 0 |
| TpMuguga_01g00707 | 10 | 0 |
| TpMuguga_01g00707 | 11 | 0 |
| TpMuguga_01g00708 | 1 | 0 |
| TpMuguga_01g00709 | 1 | 0 |
| TpMuguga_01g00710 | 1 | 1.368421053 |
| TpMuguga_01g00710 | 2 | 0.736842105 |
| TpMuguga_01g00710 | 3 | 0 |
| TpMuguga_01g00710 | 4 | 0 |
| TpMuguga_01g00710 | 5 | 0 |
| TpMuguga_01g00711 | 1 | 0 |
| TpMuguga_01g00711 | 2 | 0 |
| TpMuguga_01g00711 | 3 | 0 |
| TpMuguga_01g00711 | 4 | 0 |
| TpMuguga_01g00711 | 5 | 0 |
| TpMuguga_01g00711 | 6 | 0 |
| TpMuguga_01g00711 | 7 | 0 |
| TpMuguga_01g02690 | 1 | 0 |
| TpMuguga_01g02690 | 2 | 0 |
| TpMuguga_01g02690 | 3 | 0 |
| TpMuguga_01g02690 | 4 | 7.5 |
| TpMuguga_01g02690 | 5 | 0 |
| TpMuguga_01g02690 | 6 | 0 |

| | | |
|---|---|---|
| TpMuguga_01g02690 | 7 | 0 |
| TpMuguga_01g02690 | 8 | 15 |
| TpMuguga_01g02690 | 9 | 0 |
| TpMuguga_01g02690 | 10 | 0 |
| TpMuguga_01g02690 | 11 | 0 |
| TpMuguga_01g02690 | 12 | 17.5 |
| TpMuguga_01g02690 | 13 | 15 |
| TpMuguga_01g02690 | 14 | 12.5 |
| TpMuguga_01g00715 | 1 | 0 |
| TpMuguga_01g00716 | 1 | 8 |
| TpMuguga_01g00719 | 1 | 0.144404332 |
| TpMuguga_01g00718 | 1 | 0 |
| TpMuguga_01g00718 | 2 | 0.20979021 |
| TpMuguga_01g00718 | 3 | 0 |
| TpMuguga_01g00718 | 4 | 0.839160839 |
| TpMuguga_01g00722 | 1 | 7.764705882 |
| TpMuguga_01g00722 | 2 | 0 |
| TpMuguga_01g00722 | 3 | 0 |
| TpMuguga_01g00722 | 4 | 0 |
| TpMuguga_01g00722 | 5 | 0 |
| TpMuguga_01g00722 | 6 | 0 |
| TpMuguga_01g00722 | 7 | 0 |
| TpMuguga_01g00722 | 8 | 0 |
| TpMuguga_01g00722 | 9 | 0 |
| TpMuguga_01g00722 | 10 | 0 |
| TpMuguga_01g00722 | 11 | 0 |
| TpMuguga_01g02520 | 1 | 0.571428571 |
| TpMuguga_01g02520 | 2 | 1.257142857 |
| TpMuguga_01g02520 | 3 | 0 |
| TpMuguga_01g02725 | 1 | 0 |
| TpMuguga_01g02725 | 2 | 0 |
| TpMuguga_01g02710 | 1 | 0.185185185 |
| TpMuguga_01g00724 | 1 | 1.228813559 |
| TpMuguga_01g00724 | 2 | 0.296610169 |
| TpMuguga_01g00724 | 3 | 0 |
| TpMuguga_01g00724 | 4 | 0 |
| TpMuguga_01g00725 | 1 | 0.113207547 |
| TpMuguga_01g00725 | 2 | 0 |
| TpMuguga_01g00725 | 3 | 0 |
| TpMuguga_01g00725 | 4 | 0 |
| TpMuguga_01g00725 | 5 | 8.037735849 |
| TpMuguga_01g00727 | 1 | 0 |

| | | |
|---|---|---|
| TpMuguga_01g00727 | 2 | 0 |
| TpMuguga_01g00727 | 3 | 0 |
| TpMuguga_01g00727 | 4 | 0 |
| TpMuguga_01g00727 | 5 | 0 |
| TpMuguga_01g00727 | 6 | 1.266666667 |
| TpMuguga_01g00727 | 7 | 0 |
| TpMuguga_01g00727 | 8 | 11.03333333 |
| TpMuguga_01g00727 | 9 | 12.93333333 |
| TpMuguga_01g00728 | 1 | 2.460431655 |
| TpMuguga_01g00728 | 2 | 1.294964029 |
| TpMuguga_01g00728 | 3 | 0 |
| TpMuguga_01g00728 | 4 | 0.863309353 |
| TpMuguga_01g00728 | 5 | 4.316546763 |
| TpMuguga_01g02670 | 1 | 0 |
| TpMuguga_01g02670 | 2 | 0 |
| TpMuguga_01g02670 | 3 | 0 |
| TpMuguga_01g02205 | 1 | 0 |
| TpMuguga_01g02205 | 2 | 0 |
| TpMuguga_01g02205 | 3 | 0 |
| TpMuguga_01g02205 | 4 | 0 |
| TpMuguga_01g02205 | 5 | 0 |
| TpMuguga_01g02205 | 6 | 0 |
| TpMuguga_01g00730 | 1 | 0.45 |
| TpMuguga_01g00730 | 2 | 0.1 |
| TpMuguga_01g00730 | 3 | 0.3 |
| TpMuguga_01g00730 | 4 | 0.65 |
| TpMuguga_01g00730 | 5 | 0 |
| TpMuguga_01g00732 | 1 | 0 |
| TpMuguga_01g00732 | 2 | 0 |
| TpMuguga_01g00732 | 3 | 0 |
| TpMuguga_01g00732 | 4 | 0 |
| TpMuguga_01g00732 | 5 | 0 |
| TpMuguga_01g00732 | 6 | 0 |
| TpMuguga_01g00732 | 7 | 0 |
| TpMuguga_01g00733 | 1 | 0 |
| TpMuguga_01g00733 | 2 | 9 |
| TpMuguga_01g00734 | 1 | 0 |
| TpMuguga_01g00734 | 2 | 0 |
| TpMuguga_01g00734 | 3 | 0 |
| TpMuguga_01g00734 | 4 | 0 |
| TpMuguga_01g00734 | 5 | 0 |
| TpMuguga_01g00735 | 1 | 0 |

| | | |
|---|---|---|
| TpMuguga_01g00735 | 2 | 0 |
| TpMuguga_01g00735 | 3 | 0 |
| TpMuguga_01g02305 | 1 | 0.054200542 |
| TpMuguga_01g02305 | 2 | 0 |
| TpMuguga_01g02305 | 3 | 0 |
| TpMuguga_01g02305 | 4 | 0.021680217 |
| TpMuguga_01g02305 | 5 | 0.032520325 |
| TpMuguga_01g02305 | 6 | 0.411924119 |
| TpMuguga_01g02305 | 7 | 0.281842818 |
| TpMuguga_01g00736 | 1 | 0 |
| TpMuguga_01g00736 | 2 | 0 |
| TpMuguga_01g00736 | 3 | 0 |
| TpMuguga_01g00736 | 4 | 1.818181818 |
| TpMuguga_01g02175 | 1 | 0 |
| TpMuguga_01g02175 | 2 | 0 |
| TpMuguga_01g02175 | 3 | 0 |
| TpMuguga_01g02175 | 4 | 0 |
| TpMuguga_01g02175 | 5 | 0 |
| TpMuguga_01g02175 | 6 | 0 |
| TpMuguga_01g02175 | 7 | 0 |
| TpMuguga_01g02175 | 8 | 20.25 |
| TpMuguga_01g00742 | 1 | 0 |
| TpMuguga_01g00742 | 2 | 0 |
| TpMuguga_01g00742 | 3 | 0 |
| TpMuguga_01g00743 | 1 | 0 |
| TpMuguga_01g00743 | 2 | 0 |
| TpMuguga_01g00743 | 3 | 0 |
| TpMuguga_01g00743 | 4 | 0 |
| TpMuguga_01g00743 | 5 | 0 |
| TpMuguga_01g00743 | 6 | 7 |
| TpMuguga_01g00744 | 1 | 0 |
| TpMuguga_01g00744 | 2 | 0 |
| TpMuguga_01g00744 | 3 | 0 |
| TpMuguga_01g00744 | 4 | 0 |
| TpMuguga_01g00744 | 5 | 0 |
| TpMuguga_01g00744 | 6 | 0.250596659 |
| TpMuguga_01g00745 | 1 | 1.228668942 |
| TpMuguga_01g00745 | 2 | 0.030716724 |
| TpMuguga_01g00745 | 3 | 0.030716724 |
| TpMuguga_01g00745 | 4 | 0.092150171 |
| TpMuguga_01g00745 | 5 | 0 |
| TpMuguga_01g00745 | 6 | 0 |

| | | |
|---|---|---|
| TpMuguga_01g00745 | 7 | 0.215017065 |
| TpMuguga_01g00745 | 8 | 0 |
| TpMuguga_01g00746 | 1 | 0 |
| TpMuguga_01g00746 | 2 | 0 |
| TpMuguga_01g00746 | 3 | 0 |
| TpMuguga_01g00746 | 4 | 7.2 |
| TpMuguga_01g00746 | 5 | 0 |
| TpMuguga_01g00746 | 6 | 0 |
| TpMuguga_01g00746 | 7 | 7.2 |
| TpMuguga_01g00748 | 1 | 4.4 |
| TpMuguga_01g02600 | 1 | 1 |
| TpMuguga_01g02600 | 2 | 0 |
| TpMuguga_01g02600 | 3 | 6 |
| TpMuguga_01g02600 | 4 | 0 |
| TpMuguga_01g02600 | 5 | 0 |
| TpMuguga_01g02600 | 6 | 10 |
| TpMuguga_01g02600 | 7 | 4 |
| TpMuguga_01g02500 | 1 | 0.617647059 |
| TpMuguga_01g02500 | 2 | 0.411764706 |
| TpMuguga_01g02500 | 3 | 0.617647059 |
| TpMuguga_01g02500 | 4 | 0 |
| TpMuguga_01g02500 | 5 | 0 |
| TpMuguga_01g02500 | 6 | 0 |
| TpMuguga_01g00754 | 1 | 0 |
| TpMuguga_01g00754 | 2 | 9 |
| TpMuguga_01g00753 | 1 | 0 |
| TpMuguga_01g00753 | 2 | 0 |
| TpMuguga_01g00753 | 3 | 0 |
| TpMuguga_01g00753 | 4 | 0 |
| TpMuguga_01g00753 | 5 | 0 |
| TpMuguga_01g00753 | 6 | 0 |
| TpMuguga_01g00753 | 7 | 0 |
| TpMuguga_01g00753 | 8 | 0 |
| TpMuguga_01g00753 | 9 | 1.538461538 |
| TpMuguga_01g00755 | 1 | 0.916666667 |
| TpMuguga_01g00755 | 2 | 0 |
| TpMuguga_01g00755 | 3 | 0 |
| TpMuguga_01g00756 | 1 | 0 |
| TpMuguga_01g00756 | 2 | 0 |
| TpMuguga_01g00756 | 3 | 0 |
| TpMuguga_01g00756 | 4 | 0 |
| TpMuguga_01g00756 | 5 | 0 |

| | | |
|---|---|---|
| TpMuguga_01g00756 | 6 | 0 |
| TpMuguga_01g00756 | 7 | 0 |
| TpMuguga_01g00756 | 8 | 0 |
| TpMuguga_01g00757 | 1 | 0 |
| TpMuguga_01g00757 | 2 | 1.75862069 |
| TpMuguga_01g00758 | 1 | 0 |
| TpMuguga_01g00758 | 2 | 0 |
| TpMuguga_01g00758 | 3 | 0 |
| TpMuguga_01g00758 | 4 | 0 |
| TpMuguga_01g00758 | 5 | 0 |
| TpMuguga_01g00758 | 6 | 0 |
| TpMuguga_01g00758 | 7 | 0 |
| TpMuguga_01g00758 | 8 | 4 |
| TpMuguga_01g00758 | 9 | 0 |
| TpMuguga_01g00758 | 10 | 0 |
| TpMuguga_01g00758 | 11 | 0 |
| TpMuguga_01g00759 | 1 | 6.125 |
| TpMuguga_01g00759 | 2 | 0 |
| TpMuguga_01g00759 | 3 | 0 |
| TpMuguga_01g00759 | 4 | 0 |
| TpMuguga_01g00759 | 5 | 0 |
| TpMuguga_01g00759 | 6 | 0 |
| TpMuguga_01g00759 | 7 | 0 |
| TpMuguga_01g00759 | 8 | 0 |
| TpMuguga_01g00759 | 9 | 0 |
| TpMuguga_01g00759 | 10 | 3.5 |
| TpMuguga_01g00759 | 11 | 0 |
| TpMuguga_01g00759 | 12 | 0 |
| TpMuguga_01g00759 | 13 | 7.875 |
| TpMuguga_01g00760 | 1 | 0 |
| TpMuguga_01g00760 | 2 | 0 |
| TpMuguga_01g00760 | 3 | 0 |
| TpMuguga_01g00760 | 4 | 0 |
| TpMuguga_01g00760 | 5 | 0 |
| TpMuguga_01g00760 | 6 | 0 |
| TpMuguga_01g00761 | 1 | 0 |
| TpMuguga_01g00762 | 1 | 0 |
| TpMuguga_01g00762 | 2 | 0 |
| TpMuguga_01g00762 | 3 | 0 |
| TpMuguga_01g00763 | 1 | 0 |
| TpMuguga_01g02080 | 1 | 0 |
| TpMuguga_01g02080 | 2 | 0 |

| | | |
|---|---|---|
| TpMuguga_01g02080 | 3 | 0 |
| TpMuguga_01g02080 | 4 | 0 |
| TpMuguga_01g02080 | 5 | 0 |
| TpMuguga_01g00765 | 1 | 0 |
| TpMuguga_01g00765 | 2 | 0 |
| TpMuguga_01g00765 | 3 | 0 |
| TpMuguga_01g00768 | 1 | 0 |
| TpMuguga_01g00768 | 2 | 0 |
| TpMuguga_01g00768 | 3 | 0 |
| TpMuguga_01g00768 | 4 | 0 |
| TpMuguga_01g00769 | 1 | 0 |
| TpMuguga_01g00769 | 2 | 0 |
| TpMuguga_01g02480 | 1 | 0 |
| TpMuguga_01g02480 | 2 | 0 |
| TpMuguga_01g02480 | 3 | 0 |
| TpMuguga_01g02480 | 4 | 0 |
| TpMuguga_01g02480 | 5 | 0 |
| TpMuguga_01g02480 | 6 | 0 |
| TpMuguga_01g02480 | 7 | 0 |
| TpMuguga_01g02480 | 8 | 0 |
| TpMuguga_01g02480 | 9 | 0 |
| TpMuguga_01g02480 | 10 | 0 |
| TpMuguga_01g02480 | 11 | 0 |
| TpMuguga_01g02480 | 12 | 0 |
| TpMuguga_01g02480 | 13 | 0 |
| TpMuguga_01g02480 | 14 | 0 |
| TpMuguga_01g02480 | 15 | 0 |
| TpMuguga_01g02480 | 16 | 0 |
| TpMuguga_01g02480 | 17 | 0 |
| TpMuguga_01g02480 | 18 | 0 |
| TpMuguga_01g00775 | 1 | 0 |
| TpMuguga_01g00777 | 1 | 0 |
| TpMuguga_01g00777 | 2 | 0 |
| TpMuguga_01g02795 | 1 | 4.083333333 |
| TpMuguga_01g02795 | 2 | 0 |
| TpMuguga_01g02795 | 3 | 0 |
| TpMuguga_01g02795 | 4 | 0 |
| TpMuguga_01g02795 | 5 | 0 |
| TpMuguga_01g02795 | 6 | 0 |
| TpMuguga_01g00778 | 1 | 0 |
| TpMuguga_01g00779 | 1 | 0 |
| TpMuguga_01g00779 | 2 | 0 |

| | | |
|---|---|---|
| TpMuguga_01g00779 | 3 | 0 |
| TpMuguga_01g00779 | 4 | 0 |
| TpMuguga_01g00779 | 5 | 0 |
| TpMuguga_01g00779 | 6 | 0 |
| TpMuguga_01g00779 | 7 | 1.567839196 |
| TpMuguga_01g02235 | 1 | 0.580645161 |
| TpMuguga_01g00781 | 1 | 0 |
| TpMuguga_01g00781 | 2 | 0.196721311 |
| TpMuguga_01g00781 | 3 | 2.491803279 |
| TpMuguga_01g00782 | 1 | 0 |
| TpMuguga_01g00782 | 2 | 0.136986301 |
| TpMuguga_01g00782 | 3 | 1.095890411 |
| TpMuguga_01g00782 | 4 | 0.684931507 |
| TpMuguga_01g00782 | 5 | 0.136986301 |
| TpMuguga_01g00782 | 6 | 0.273972603 |
| TpMuguga_01g00782 | 7 | 0.684931507 |
| TpMuguga_01g00782 | 8 | 0.273972603 |
| TpMuguga_01g00782 | 9 | 1.506849315 |
| TpMuguga_01g00783 | 1 | 0 |
| TpMuguga_01g00783 | 2 | 0 |
| TpMuguga_01g00784 | 1 | 0 |
| TpMuguga_01g02370 | 1 | 1.684210526 |
| TpMuguga_01g02370 | 2 | 0.421052632 |
| TpMuguga_01g02370 | 3 | 2.315789474 |
| TpMuguga_01g02370 | 4 | 0.842105263 |
| TpMuguga_01g02370 | 5 | 0 |
| TpMuguga_01g02370 | 6 | 0 |
| TpMuguga_01g02370 | 7 | 0 |
| TpMuguga_01g00785 | 1 | 0 |
| TpMuguga_01g00785 | 2 | 0 |
| TpMuguga_01g00785 | 3 | 0 |
| TpMuguga_01g00785 | 4 | 0 |
| TpMuguga_01g00785 | 5 | 0 |
| TpMuguga_01g00785 | 6 | 0 |
| TpMuguga_01g00785 | 7 | 0 |
| TpMuguga_01g00786 | 1 | 0.483870968 |
| TpMuguga_01g00786 | 2 | 0.22983871 |
| TpMuguga_01g00787 | 1 | 0 |
| TpMuguga_01g00787 | 2 | 0 |
| TpMuguga_01g00787 | 3 | 0 |
| TpMuguga_01g00787 | 4 | 0 |
| TpMuguga_01g00787 | 5 | 0 |

| | | |
|---|---|---|
| TpMuguga_01g00787 | 6 | 10.66666667 |
| TpMuguga_01g00787 | 7 | 0 |
| TpMuguga_01g00788 | 1 | 0.658536585 |
| TpMuguga_01g00788 | 2 | 0 |
| TpMuguga_01g00791 | 1 | 1.137931034 |
| TpMuguga_01g00791 | 2 | 0.517241379 |
| TpMuguga_01g00792 | 1 | 0 |
| TpMuguga_01g00792 | 2 | 0 |
| TpMuguga_01g00792 | 3 | 0 |
| TpMuguga_01g00792 | 4 | 0 |
| TpMuguga_01g00792 | 5 | 0 |
| TpMuguga_01g00792 | 6 | 0 |
| TpMuguga_01g00792 | 7 | 0 |
| TpMuguga_01g00792 | 8 | 0 |
| TpMuguga_01g00792 | 9 | 0 |
| TpMuguga_01g00792 | 10 | 0 |
| TpMuguga_01g00793 | 1 | 3.84279476 |
| TpMuguga_01g00793 | 2 | 3.218340611 |
| TpMuguga_01g00794 | 1 | 0 |
| TpMuguga_01g00794 | 2 | 0 |
| TpMuguga_01g00794 | 3 | 0 |
| TpMuguga_01g00794 | 4 | 0 |
| TpMuguga_01g00794 | 5 | 0 |
| TpMuguga_01g00794 | 6 | 0 |
| TpMuguga_01g00796 | 1 | 0 |
| TpMuguga_01g00795 | 1 | 0.387096774 |
| TpMuguga_01g00795 | 2 | 0 |
| TpMuguga_01g00797 | 1 | 0 |
| TpMuguga_01g00797 | 2 | 2.709677419 |
| TpMuguga_01g00797 | 3 | 0 |
| TpMuguga_01g00797 | 4 | 0 |
| TpMuguga_01g00797 | 5 | 0 |
| TpMuguga_01g00797 | 6 | 0 |
| TpMuguga_01g00797 | 7 | 0 |
| TpMuguga_01g00797 | 8 | 0 |
| TpMuguga_01g00797 | 9 | 0 |
| TpMuguga_01g00797 | 10 | 0 |
| TpMuguga_01g00797 | 11 | 0 |
| TpMuguga_01g00797 | 12 | 0 |
| TpMuguga_01g00797 | 13 | 0 |
| TpMuguga_01g02565 | 1 | 0 |
| TpMuguga_01g02565 | 2 | 0 |

| | | |
|---|---|---|
| TpMuguga_01g02565 | 3 | 0.24691358 |
| TpMuguga_01g02565 | 4 | 0 |
| TpMuguga_01g00798 | 1 | 0 |
| TpMuguga_01g00798 | 2 | 0 |
| TpMuguga_01g00798 | 3 | 0 |
| TpMuguga_01g00798 | 4 | 0 |
| TpMuguga_01g00798 | 5 | 0 |
| TpMuguga_01g00798 | 6 | 0 |
| TpMuguga_01g00798 | 7 | 0 |
| TpMuguga_01g00798 | 8 | 0 |
| TpMuguga_01g00798 | 9 | 0 |
| TpMuguga_01g00798 | 10 | 77 |
| TpMuguga_01g02340 | 1 | 0 |
| TpMuguga_01g02340 | 2 | 0 |
| TpMuguga_01g02340 | 3 | 0 |
| TpMuguga_01g02340 | 4 | 0 |
| TpMuguga_01g02340 | 5 | 0 |
| TpMuguga_01g00800 | 1 | 0 |
| TpMuguga_01g00800 | 2 | 0 |
| TpMuguga_01g02720 | 1 | 0 |
| TpMuguga_01g02720 | 2 | 0 |
| TpMuguga_01g02720 | 3 | 3 |
| TpMuguga_01g02720 | 4 | 0 |
| TpMuguga_01g02720 | 5 | 0 |
| TpMuguga_01g02720 | 6 | 0 |
| TpMuguga_01g02720 | 7 | 0 |
| TpMuguga_01g02720 | 8 | 4 |
| TpMuguga_01g02720 | 9 | 0 |
| TpMuguga_01g00804 | 1 | 0 |
| TpMuguga_01g00804 | 2 | 0 |
| TpMuguga_01g00804 | 3 | 0 |
| TpMuguga_01g00804 | 4 | 0 |
| TpMuguga_01g00804 | 5 | 5.1 |
| TpMuguga_01g00805 | 1 | 0 |
| TpMuguga_01g00805 | 2 | 0 |
| TpMuguga_01g00805 | 3 | 0 |
| TpMuguga_01g00805 | 4 | 0 |
| TpMuguga_01g00805 | 5 | 0 |
| TpMuguga_01g00805 | 6 | 0 |
| TpMuguga_01g00805 | 7 | 0 |
| TpMuguga_01g00805 | 8 | 0 |
| TpMuguga_01g00805 | 9 | 0 |

| | | |
|---|---|---|
| TpMuguga_01g00805 | 10 | 0 |
| TpMuguga_01g00805 | 11 | 0 |
| TpMuguga_01g00805 | 12 | 0 |
| TpMuguga_01g00805 | 13 | 0 |
| TpMuguga_01g00805 | 14 | 0 |
| TpMuguga_01g00805 | 15 | 0 |
| TpMuguga_01g00805 | 16 | 0 |
| TpMuguga_01g00805 | 17 | 0 |
| TpMuguga_01g00805 | 18 | 0 |
| TpMuguga_01g00805 | 19 | 0 |
| TpMuguga_01g00805 | 20 | 0.535149385 |
| TpMuguga_01g00806 | 1 | 1.559701493 |
| TpMuguga_01g00809 | 1 | 0 |
| TpMuguga_01g00809 | 2 | 0 |
| TpMuguga_01g00809 | 3 | 0 |
| TpMuguga_01g00809 | 4 | 0 |
| TpMuguga_01g00809 | 5 | 0 |
| TpMuguga_01g00809 | 6 | 0 |
| TpMuguga_01g00810 | 1 | 0 |
| TpMuguga_01g00810 | 2 | 0 |
| TpMuguga_01g00810 | 3 | 0 |
| TpMuguga_01g00811 | 1 | 0 |
| TpMuguga_01g00811 | 2 | 18 |
| TpMuguga_01g00811 | 3 | 0 |
| TpMuguga_01g00811 | 4 | 12 |
| TpMuguga_01g00811 | 5 | 9 |
| TpMuguga_01g00812 | 1 | 0 |
| TpMuguga_01g00812 | 2 | 0 |
| TpMuguga_01g00812 | 3 | 0 |
| TpMuguga_01g02255 | 1 | 0 |
| TpMuguga_01g02255 | 2 | 0 |
| TpMuguga_01g00815 | 1 | 0 |
| TpMuguga_01g00815 | 2 | 0 |
| TpMuguga_01g00815 | 3 | 0 |
| TpMuguga_01g00815 | 4 | 0 |
| TpMuguga_01g00815 | 5 | 0 |
| TpMuguga_01g00815 | 6 | 0 |
| TpMuguga_01g00815 | 7 | 0 |
| TpMuguga_01g00815 | 8 | 0 |
| TpMuguga_01g00815 | 9 | 0 |
| TpMuguga_01g00815 | 10 | 0 |
| TpMuguga_01g00815 | 11 | 0 |

| | | |
|---|---|---|
| TpMuguga_01g00815 | 12 | 0 |
| TpMuguga_01g00816 | 1 | 2.181818182 |
| TpMuguga_01g00816 | 2 | 1.454545455 |
| TpMuguga_01g00816 | 3 | 0.727272727 |
| TpMuguga_01g00816 | 4 | 0.484848485 |
| TpMuguga_01g00816 | 5 | 4.121212121 |
| TpMuguga_01g00816 | 6 | 0 |
| TpMuguga_01g00816 | 7 | 0 |
| TpMuguga_01g00818 | 1 | 3.096774194 |
| TpMuguga_01g00818 | 2 | 2.322580645 |
| TpMuguga_01g00818 | 3 | 0 |
| TpMuguga_01g00818 | 4 | 0 |
| TpMuguga_01g00818 | 5 | 0 |
| TpMuguga_01g00818 | 6 | 0 |
| TpMuguga_01g00818 | 7 | 0 |
| TpMuguga_01g00818 | 8 | 0 |
| TpMuguga_01g00818 | 9 | 0 |
| TpMuguga_01g00818 | 10 | 0 |
| TpMuguga_01g00818 | 11 | 0 |
| TpMuguga_01g00819 | 1 | 0 |
| TpMuguga_01g00819 | 2 | 0 |
| TpMuguga_01g00819 | 3 | 0 |
| TpMuguga_01g00819 | 4 | 0 |
| TpMuguga_01g00820 | 1 | 0 |
| TpMuguga_01g00820 | 2 | 0 |
| TpMuguga_01g00820 | 3 | 0 |
| TpMuguga_01g00821 | 1 | 0 |
| TpMuguga_01g00822 | 1 | 0 |
| TpMuguga_01g00822 | 2 | 0 |
| TpMuguga_01g00822 | 3 | 0 |
| TpMuguga_01g00822 | 4 | 0 |
| TpMuguga_01g00822 | 5 | 0 |
| TpMuguga_01g00822 | 6 | 0 |
| TpMuguga_01g00822 | 7 | 0 |
| TpMuguga_01g02715 | 1 | 0 |
| TpMuguga_01g02715 | 2 | 0 |
| TpMuguga_01g02715 | 3 | 0 |
| TpMuguga_01g02715 | 4 | 0 |
| TpMuguga_01g02715 | 5 | 0 |
| TpMuguga_01g02715 | 6 | 0 |
| TpMuguga_01g02715 | 7 | 0 |
| TpMuguga_01g02715 | 8 | 0 |

| | | |
|---|---|---|
| TpMuguga_01g02715 | 9 | 0 |
| TpMuguga_01g02715 | 10 | 0 |
| TpMuguga_01g00826 | 1 | 0.747126437 |
| TpMuguga_01g00826 | 2 | 0 |
| TpMuguga_01g00826 | 3 | 0.298850575 |
| TpMuguga_01g00826 | 4 | 1.195402299 |
| TpMuguga_01g00826 | 5 | 0 |
| TpMuguga_01g00826 | 6 | 0.896551724 |
| TpMuguga_01g00826 | 7 | 0 |
| TpMuguga_01g00826 | 8 | 1.195402299 |
| TpMuguga_01g00826 | 9 | 0.448275862 |
| TpMuguga_01g00826 | 10 | 3.586206897 |
| TpMuguga_01g00826 | 11 | 3.586206897 |
| TpMuguga_01g00826 | 12 | 12.70114943 |
| TpMuguga_01g02635 | 1 | 0 |
| TpMuguga_01g02635 | 2 | 0 |
| TpMuguga_01g02635 | 3 | 0.583333333 |
| TpMuguga_01g02635 | 4 | 0.25 |
| TpMuguga_01g02635 | 5 | 1.416666667 |
| TpMuguga_01g00827 | 1 | 6.769230769 |
| TpMuguga_01g00827 | 2 | 4 |
| TpMuguga_01g00827 | 3 | 0 |
| TpMuguga_01g00828 | 1 | 0 |
| TpMuguga_01g00828 | 2 | 0 |
| TpMuguga_01g00828 | 3 | 0 |
| TpMuguga_01g00828 | 4 | 0 |
| TpMuguga_01g00828 | 5 | 0 |
| TpMuguga_01g00829 | 1 | 0 |
| TpMuguga_01g00829 | 2 | 0 |
| TpMuguga_01g00829 | 3 | 0 |
| TpMuguga_01g00829 | 4 | 0 |
| TpMuguga_01g00829 | 5 | 0 |
| TpMuguga_01g00829 | 6 | 0 |
| TpMuguga_01g02770 | 1 | 0 |
| TpMuguga_01g02770 | 2 | 0 |
| TpMuguga_01g02770 | 3 | 0 |
| TpMuguga_01g02770 | 4 | 0 |
| TpMuguga_01g02770 | 5 | 0 |
| TpMuguga_01g02770 | 6 | 0 |
| TpMuguga_01g02770 | 7 | 0 |
| TpMuguga_01g02770 | 8 | 0 |
| TpMuguga_01g02770 | 9 | 0.221276596 |

| | | |
|---|---|---|
| TpMuguga_01g02770 | 10 | 0 |
| TpMuguga_01g02770 | 11 | 0 |
| TpMuguga_01g02770 | 12 | 1.604255319 |
| TpMuguga_01g00833 | 1 | 0.177777778 |
| TpMuguga_01g00835 | 1 | 0 |
| TpMuguga_01g00835 | 2 | 0 |
| TpMuguga_01g00835 | 3 | 0 |
| TpMuguga_01g00836 | 1 | 0 |
| TpMuguga_01g00836 | 2 | 0 |
| TpMuguga_01g00836 | 3 | 0 |
| TpMuguga_01g00836 | 4 | 0 |
| TpMuguga_01g00836 | 5 | 0 |
| TpMuguga_01g00836 | 6 | 0 |
| TpMuguga_01g00838 | 1 | 0 |
| TpMuguga_01g00838 | 2 | 0 |
| TpMuguga_01g00839 | 1 | 0 |
| TpMuguga_01g00840 | 1 | 0 |
| TpMuguga_01g00840 | 2 | 0 |
| TpMuguga_01g00840 | 3 | 0 |
| TpMuguga_01g00840 | 4 | 0 |
| TpMuguga_01g00840 | 5 | 0 |
| TpMuguga_01g00840 | 6 | 0 |
| TpMuguga_01g00840 | 7 | 0 |
| TpMuguga_01g00840 | 8 | 0 |
| TpMuguga_01g00840 | 9 | 0 |
| TpMuguga_01g00840 | 10 | 0 |
| TpMuguga_01g00840 | 11 | 0 |
| TpMuguga_01g00840 | 12 | 0 |
| TpMuguga_01g00840 | 13 | 0 |
| TpMuguga_01g00840 | 14 | 0 |
| TpMuguga_01g00842 | 1 | 0 |
| TpMuguga_01g00842 | 2 | 0 |
| TpMuguga_01g00842 | 3 | 0 |
| TpMuguga_01g00842 | 4 | 0 |
| TpMuguga_01g00842 | 5 | 0 |
| TpMuguga_01g00842 | 6 | 0 |
| TpMuguga_01g00842 | 7 | 0 |
| TpMuguga_01g00842 | 8 | 3.333333333 |
| TpMuguga_01g00842 | 9 | 3.333333333 |
| TpMuguga_01g00842 | 10 | 0 |
| TpMuguga_01g00842 | 11 | 7.333333333 |
| TpMuguga_01g00842 | 12 | 1.333333333 |

| | | |
|---|---|---|
| TpMuguga_01g00842 | 13 | 1.333333333 |
| TpMuguga_01g00842 | 14 | 2 |
| TpMuguga_01g00842 | 15 | 0 |
| TpMuguga_01g00841 | 1 | 0 |
| TpMuguga_01g00841 | 2 | 0 |
| TpMuguga_01g00841 | 3 | 0 |
| TpMuguga_01g02550 | 1 | 9 |
| TpMuguga_01g02550 | 2 | 6.5 |
| TpMuguga_01g02665 | 1 | 0 |
| TpMuguga_01g02665 | 2 | 0 |
| TpMuguga_01g02665 | 3 | 0 |
| TpMuguga_01g02665 | 4 | 0 |
| TpMuguga_01g02665 | 5 | 0 |
| TpMuguga_01g00845 | 1 | 0 |
| TpMuguga_01g00845 | 2 | 0 |
| TpMuguga_01g00845 | 3 | 0 |
| TpMuguga_01g00845 | 4 | 0 |
| TpMuguga_01g00845 | 5 | 0 |
| TpMuguga_01g00846 | 1 | 0 |
| TpMuguga_01g00846 | 2 | 0 |
| TpMuguga_01g00846 | 3 | 0 |
| TpMuguga_01g00846 | 4 | 0 |
| TpMuguga_01g00846 | 5 | 0 |
| TpMuguga_01g02695 | 1 | 2.857142857 |
| TpMuguga_01g02695 | 2 | 7.142857143 |
| TpMuguga_01g02695 | 3 | 1.428571429 |
| TpMuguga_01g02695 | 4 | 0 |
| TpMuguga_01g02695 | 5 | 0 |
| TpMuguga_01g02695 | 6 | 0 |
| TpMuguga_01g02695 | 7 | 0 |
| TpMuguga_01g02695 | 8 | 0 |
| TpMuguga_01g02695 | 9 | 0 |
| TpMuguga_01g02860 | 1 | 0 |
| TpMuguga_01g00850 | 1 | 0 |
| TpMuguga_01g00850 | 2 | 0 |
| TpMuguga_01g00850 | 3 | 0 |
| TpMuguga_01g00850 | 4 | 0 |
| TpMuguga_01g00850 | 5 | 0 |
| TpMuguga_01g00850 | 6 | 0 |
| TpMuguga_01g00851 | 1 | 0 |
| TpMuguga_01g00852 | 1 | 0 |
| TpMuguga_01g00852 | 2 | 0 |

| | | |
|---|---|---|
| TpMuguga_01g00852 | 3 | 0 |
| TpMuguga_01g00852 | 4 | 0 |
| TpMuguga_01g00852 | 5 | 0 |
| TpMuguga_01g00852 | 6 | 0 |
| TpMuguga_01g00852 | 7 | 0 |
| TpMuguga_01g00852 | 8 | 0 |
| TpMuguga_01g00853 | 1 | 0 |
| TpMuguga_01g00853 | 2 | 0 |
| TpMuguga_01g00853 | 3 | 0 |
| TpMuguga_01g00853 | 4 | 0 |
| TpMuguga_01g00853 | 5 | 0.35 |
| TpMuguga_01g00853 | 6 | 1.4 |
| TpMuguga_01g00855 | 1 | 0.672413793 |
| TpMuguga_01g00855 | 2 | 0.206896552 |
| TpMuguga_01g00855 | 3 | 0.336206897 |
| TpMuguga_01g00855 | 4 | 0 |
| TpMuguga_01g00855 | 5 | 0 |
| TpMuguga_01g00854 | 1 | 0 |
| TpMuguga_01g00854 | 2 | 0 |
| TpMuguga_01g00854 | 3 | 0 |
| TpMuguga_01g00854 | 4 | 0 |
| TpMuguga_01g00854 | 5 | 0 |
| TpMuguga_01g00854 | 6 | 0 |
| TpMuguga_01g00854 | 7 | 0 |
| TpMuguga_01g00856 | 1 | 1.2 |
| TpMuguga_01g00857 | 1 | 56.88888889 |
| TpMuguga_01g00857 | 2 | 16.88888889 |
| TpMuguga_01g00857 | 3 | 14.22222222 |
| TpMuguga_01g00857 | 4 | 0 |
| TpMuguga_01g00857 | 5 | 0 |
| TpMuguga_01g00857 | 6 | 0 |
| TpMuguga_01g00857 | 7 | 0 |
| TpMuguga_01g00858 | 1 | 0.339285714 |
| TpMuguga_01g00859 | 1 | 7.076923077 |
| TpMuguga_01g00859 | 2 | 1.153846154 |
| TpMuguga_01g00859 | 3 | 0 |
| TpMuguga_01g00859 | 4 | 0 |
| TpMuguga_01g00860 | 1 | 0 |
| TpMuguga_01g00860 | 2 | 0 |
| TpMuguga_01g00861 | 1 | 0 |
| TpMuguga_01g00862 | 1 | 0 |
| TpMuguga_01g00864 | 1 | 1.714285714 |

| | | |
|---|---|---|
| TpMuguga_01g00864 | 2 | 0 |
| TpMuguga_01g00864 | 3 | 0 |
| TpMuguga_01g00864 | 4 | 0 |
| TpMuguga_01g00864 | 5 | 0 |
| TpMuguga_01g00864 | 6 | 3.428571429 |
| TpMuguga_01g00864 | 7 | 0.857142857 |
| TpMuguga_01g00864 | 8 | 12 |
| TpMuguga_01g00864 | 9 | 0 |
| TpMuguga_01g00864 | 10 | 0 |
| TpMuguga_01g00864 | 11 | 1.714285714 |
| TpMuguga_01g00863 | 1 | 0 |
| TpMuguga_01g00863 | 2 | 0 |
| TpMuguga_01g00863 | 3 | 0 |
| TpMuguga_01g00863 | 4 | 0 |
| TpMuguga_01g00863 | 5 | 0 |
| TpMuguga_01g00863 | 6 | 0 |
| TpMuguga_01g00863 | 7 | 0 |
| TpMuguga_01g00865 | 1 | 0 |
| TpMuguga_01g00865 | 2 | 0 |
| TpMuguga_01g00865 | 3 | 0 |
| TpMuguga_01g00868 | 1 | 0 |
| TpMuguga_01g00869 | 1 | 0 |
| TpMuguga_01g00869 | 2 | 0 |
| TpMuguga_01g00869 | 3 | 0 |
| TpMuguga_01g00869 | 4 | 0 |
| TpMuguga_01g00869 | 5 | 4.55 |
| TpMuguga_01g00869 | 6 | 0 |
| TpMuguga_01g00870 | 1 | 0 |
| TpMuguga_01g00870 | 2 | 0 |
| TpMuguga_01g00870 | 3 | 0 |
| TpMuguga_01g00870 | 4 | 0 |
| TpMuguga_01g00870 | 5 | 0 |
| TpMuguga_01g00870 | 6 | 9 |
| TpMuguga_01g00870 | 7 | 0 |
| TpMuguga_01g00870 | 8 | 0 |
| TpMuguga_01g00871 | 1 | 4.235294118 |
| TpMuguga_01g00871 | 2 | 2.470588235 |
| TpMuguga_01g00871 | 3 | 0.352941176 |
| TpMuguga_01g00871 | 4 | 0 |
| TpMuguga_01g00871 | 5 | 2.470588235 |
| TpMuguga_01g00873 | 1 | 0.127659574 |
| TpMuguga_01g00873 | 2 | 0 |

| | | |
|---|---|---|
| TpMuguga_01g00873 | 3 | 0.127659574 |
| TpMuguga_01g00873 | 4 | 1.787234043 |
| TpMuguga_01g00873 | 5 | 4.468085106 |
| TpMuguga_01g00874 | 1 | 0 |
| TpMuguga_01g00874 | 2 | 0 |
| TpMuguga_01g00874 | 3 | 0 |
| TpMuguga_01g00874 | 4 | 0 |
| TpMuguga_01g00874 | 5 | 0 |
| TpMuguga_01g00874 | 6 | 0 |
| TpMuguga_01g00875 | 1 | 0 |
| TpMuguga_01g00875 | 2 | 0 |
| TpMuguga_01g00875 | 3 | 0 |
| TpMuguga_01g00875 | 4 | 0 |
| TpMuguga_01g00875 | 5 | 0 |
| TpMuguga_01g00875 | 6 | 0 |
| TpMuguga_01g00875 | 7 | 0 |
| TpMuguga_01g00877 | 1 | 0 |
| TpMuguga_01g00877 | 2 | 0 |
| TpMuguga_01g00877 | 3 | 0 |
| TpMuguga_01g00877 | 4 | 0 |
| TpMuguga_01g00877 | 5 | 0 |
| TpMuguga_01g00877 | 6 | 0 |
| TpMuguga_01g00876 | 1 | 0 |
| TpMuguga_01g00876 | 2 | 0 |
| TpMuguga_01g00876 | 3 | 0 |
| TpMuguga_01g00878 | 1 | 0.157894737 |
| TpMuguga_01g00878 | 2 | 0 |
| TpMuguga_01g00878 | 3 | 0.710526316 |
| TpMuguga_01g00878 | 4 | 0.078947368 |
| TpMuguga_01g00878 | 5 | 3.157894737 |
| TpMuguga_01g02560 | 1 | 0 |
| TpMuguga_01g02560 | 2 | 0.769230769 |
| TpMuguga_01g02560 | 3 | 0 |
| TpMuguga_01g02560 | 4 | 2.115384615 |
| TpMuguga_01g00881 | 1 | 1.2 |
| TpMuguga_01g00881 | 2 | 0 |
| TpMuguga_01g00881 | 3 | 0 |
| TpMuguga_01g00881 | 4 | 3.2 |
| TpMuguga_01g00881 | 5 | 8.8 |
| TpMuguga_01g00883 | 1 | 0 |
| TpMuguga_01g00885 | 1 | 0 |
| TpMuguga_01g00885 | 2 | 0 |

| | | |
|---|---|---|
| TpMuguga_01g00886 | 1 | 1 |
| TpMuguga_01g00888 | 1 | 0 |
| TpMuguga_01g00889 | 1 | 0 |
| TpMuguga_01g00891 | 1 | 0 |
| TpMuguga_01g00891 | 2 | 0 |
| TpMuguga_01g00891 | 3 | 0 |
| TpMuguga_01g00891 | 4 | 0 |
| TpMuguga_01g00891 | 5 | 0 |
| TpMuguga_01g00893 | 1 | 5.333333333 |
| TpMuguga_01g00893 | 2 | 2.666666667 |
| TpMuguga_01g00893 | 3 | 14.66666667 |
| TpMuguga_01g00892 | 1 | 0 |
| TpMuguga_01g00892 | 2 | 0 |
| TpMuguga_01g00892 | 3 | 0 |
| TpMuguga_01g00892 | 4 | 0 |
| TpMuguga_01g00892 | 5 | 0 |
| TpMuguga_01g00892 | 6 | 1 |
| TpMuguga_01g00892 | 7 | 0 |
| TpMuguga_01g00892 | 8 | 0 |
| TpMuguga_01g00892 | 9 | 0 |
| TpMuguga_01g00895 | 1 | 0 |
| TpMuguga_01g00895 | 2 | 0 |
| TpMuguga_01g00895 | 3 | 1.042105263 |
| TpMuguga_01g00895 | 4 | 0.757894737 |
| TpMuguga_01g00895 | 5 | 0 |
| TpMuguga_01g00895 | 6 | 0.094736842 |
| TpMuguga_01g00895 | 7 | 0 |
| TpMuguga_01g00895 | 8 | 0.947368421 |
| TpMuguga_01g00896 | 1 | 0.929577465 |
| TpMuguga_01g00896 | 2 | 3.507042254 |
| TpMuguga_01g00898 | 1 | 0 |
| TpMuguga_01g00898 | 2 | 0 |
| TpMuguga_01g00898 | 3 | 0 |
| TpMuguga_01g00899 | 1 | 6.348837209 |
| TpMuguga_01g00899 | 2 | 5.372093023 |
| TpMuguga_01g00899 | 3 | 0 |
| TpMuguga_01g00899 | 4 | 0 |
| TpMuguga_01g00899 | 5 | 0 |
| TpMuguga_01g00899 | 6 | 0 |
| TpMuguga_01g00899 | 7 | 0 |
| TpMuguga_01g00899 | 8 | 0 |
| TpMuguga_01g00899 | 9 | 0 |

| | | |
|---|---|---|
| TpMuguga_01g00899 | 10 | 0.976744186 |
| TpMuguga_01g00899 | 11 | 0 |
| TpMuguga_01g00899 | 12 | 0 |
| TpMuguga_01g00899 | 13 | 0.976744186 |
| TpMuguga_01g00899 | 14 | 0 |
| TpMuguga_01g00899 | 15 | 0 |
| TpMuguga_01g00899 | 16 | 0 |
| TpMuguga_01g00899 | 17 | 0 |
| TpMuguga_01g00899 | 18 | 0 |
| TpMuguga_01g00899 | 19 | 0 |
| TpMuguga_01g00899 | 20 | 0 |
| TpMuguga_01g00900 | 1 | 0 |
| TpMuguga_01g00900 | 2 | 0 |
| TpMuguga_01g00901 | 1 | 0.333333333 |
| TpMuguga_01g00901 | 2 | 0 |
| TpMuguga_01g00901 | 3 | 0 |
| TpMuguga_01g00902 | 1 | 0.75 |
| TpMuguga_01g00902 | 2 | 1.875 |
| TpMuguga_01g00902 | 3 | 0 |
| TpMuguga_01g00902 | 4 | 0 |
| TpMuguga_01g00902 | 5 | 0 |
| TpMuguga_01g00903 | 1 | 0 |
| TpMuguga_01g00905 | 1 | 1.153846154 |
| TpMuguga_01g00905 | 2 | 0.769230769 |
| TpMuguga_01g00905 | 3 | 0.641025641 |
| TpMuguga_01g00905 | 4 | 0 |
| TpMuguga_01g00908 | 1 | 0 |
| TpMuguga_01g00908 | 2 | 0 |
| TpMuguga_01g00907 | 1 | 2.666666667 |
| TpMuguga_01g00907 | 2 | 3 |
| TpMuguga_01g00909 | 1 | 0.789473684 |
| TpMuguga_01g00909 | 2 | 0.394736842 |
| TpMuguga_01g00910 | 1 | 0 |
| TpMuguga_01g00910 | 2 | 0 |
| TpMuguga_01g00910 | 3 | 2.647058824 |
| TpMuguga_01g00910 | 4 | 3.044117647 |
| TpMuguga_01g00910 | 5 | 0.397058824 |
| TpMuguga_01g00910 | 6 | 4.897058824 |
| TpMuguga_01g00910 | 7 | 2.514705882 |
| TpMuguga_01g00910 | 8 | 1.323529412 |
| TpMuguga_01g00912 | 1 | 0 |
| TpMuguga_01g00913 | 1 | 0.124293785 |

| | | |
|---|---|---|
| TpMuguga_01g00914 | 1 | 0.505263158 |
| TpMuguga_01g00915 | 1 | 0 |
| TpMuguga_01g00915 | 2 | 0 |
| TpMuguga_01g00915 | 3 | 0 |
| TpMuguga_01g00915 | 4 | 0 |
| TpMuguga_01g00915 | 5 | 0.070175439 |
| TpMuguga_01g00915 | 6 | 0.842105263 |
| TpMuguga_01g00915 | 7 | 0 |
| TpMuguga_01g00918 | 1 | 0 |
| TpMuguga_01g00919 | 1 | 0 |
| TpMuguga_01g00919 | 2 | 0 |
| TpMuguga_01g00919 | 3 | 0 |
| TpMuguga_01g00919 | 4 | 0 |
| TpMuguga_01g00920 | 1 | 0 |
| TpMuguga_01g00920 | 2 | 0 |
| TpMuguga_01g00920 | 3 | 6.363636364 |
| TpMuguga_01g00920 | 4 | 3.636363636 |
| TpMuguga_01g00921 | 1 | 0 |
| TpMuguga_01g00921 | 2 | 0 |
| TpMuguga_01g00921 | 3 | 15.42857143 |
| TpMuguga_01g00921 | 4 | 0 |
| TpMuguga_01g00921 | 5 | 0 |
| TpMuguga_01g00921 | 6 | 0 |
| TpMuguga_01g00921 | 7 | 0 |
| TpMuguga_01g00921 | 8 | 0 |
| TpMuguga_01g00921 | 9 | 0 |
| TpMuguga_01g00921 | 10 | 0 |
| TpMuguga_01g00921 | 11 | 0 |
| TpMuguga_01g00921 | 12 | 0 |
| TpMuguga_01g00921 | 13 | 0 |
| TpMuguga_01g00921 | 14 | 0 |
| TpMuguga_01g00921 | 15 | 0 |
| TpMuguga_01g00921 | 16 | 0 |
| TpMuguga_01g00921 | 17 | 0 |
| TpMuguga_01g00923 | 1 | 0 |
| TpMuguga_01g00923 | 2 | 0 |
| TpMuguga_01g00923 | 3 | 0 |
| TpMuguga_01g00923 | 4 | 0 |
| TpMuguga_01g00923 | 5 | 0 |
| TpMuguga_01g00923 | 6 | 0 |
| TpMuguga_01g00923 | 7 | 0 |
| TpMuguga_01g00923 | 8 | 0 |

| | | |
|---|---|---|
| TpMuguga_01g00923 | 9 | 0 |
| TpMuguga_01g00923 | 10 | 0 |
| TpMuguga_01g00923 | 11 | 0 |
| TpMuguga_01g00924 | 1 | 0 |
| TpMuguga_01g00924 | 2 | 0 |
| TpMuguga_01g00925 | 1 | 9 |
| TpMuguga_01g00925 | 2 | 2.25 |
| TpMuguga_01g00925 | 3 | 1.125 |
| TpMuguga_01g00925 | 4 | 0 |
| TpMuguga_01g00925 | 5 | 0 |
| TpMuguga_01g00925 | 6 | 0 |
| TpMuguga_01g00925 | 7 | 0 |
| TpMuguga_01g00925 | 8 | 0 |
| TpMuguga_01g00926 | 1 | 0 |
| TpMuguga_01g00927 | 1 | 0 |
| TpMuguga_01g00927 | 2 | 0 |
| TpMuguga_01g00927 | 3 | 0 |
| TpMuguga_01g00927 | 4 | 0 |
| TpMuguga_01g00927 | 5 | 6 |
| TpMuguga_01g00928 | 1 | 1.353220339 |
| TpMuguga_01g02265 | 1 | 0 |
| TpMuguga_01g02265 | 2 | 0 |
| TpMuguga_01g02265 | 3 | 0 |
| TpMuguga_01g02265 | 4 | 0 |
| TpMuguga_01g02265 | 5 | 0 |
| TpMuguga_01g02740 | 1 | 0 |
| TpMuguga_01g02740 | 2 | 0 |
| TpMuguga_01g02740 | 3 | 0 |
| TpMuguga_01g02745 | 1 | 4.4 |
| TpMuguga_01g02745 | 2 | 35.2 |
| TpMuguga_01g02745 | 3 | 0.4 |
| TpMuguga_01g02745 | 4 | 0 |
| TpMuguga_01g02745 | 5 | 6.8 |
| TpMuguga_01g02745 | 6 | 0 |
| TpMuguga_01g02745 | 7 | 0 |
| TpMuguga_01g02745 | 8 | 0 |
| TpMuguga_01g02745 | 9 | 0 |
| TpMuguga_01g02575 | 1 | 0 |
| TpMuguga_01g02575 | 2 | 0 |
| TpMuguga_01g02575 | 3 | 0 |
| TpMuguga_01g02575 | 4 | 0 |
| TpMuguga_01g02575 | 5 | 0 |

| | | |
|---|---|---|
| TpMuguga_01g02575 | 6 | 33 |
| TpMuguga_01g02575 | 7 | 3 |
| TpMuguga_01g02575 | 8 | 6 |
| TpMuguga_01g00937 | 1 | 1.125 |
| TpMuguga_01g00937 | 2 | 0 |
| TpMuguga_01g00937 | 3 | 0 |
| TpMuguga_01g00937 | 4 | 0 |
| TpMuguga_01g00937 | 5 | 0 |
| TpMuguga_01g00938 | 1 | 0 |
| TpMuguga_01g00938 | 2 | 0 |
| TpMuguga_01g00938 | 3 | 1.12 |
| TpMuguga_01g00940 | 1 | 2 |
| TpMuguga_01g00942 | 1 | 2.333333333 |
| TpMuguga_01g00942 | 2 | 0 |
| TpMuguga_01g00942 | 3 | 0 |
| TpMuguga_01g00942 | 4 | 0.636363636 |
| TpMuguga_01g00942 | 5 | 0.212121212 |
| TpMuguga_01g00942 | 6 | 0 |
| TpMuguga_01g00941 | 1 | 0.430769231 |
| TpMuguga_01g00941 | 2 | 0 |
| TpMuguga_01g00941 | 3 | 0 |
| TpMuguga_01g00941 | 4 | 0 |
| TpMuguga_01g00941 | 5 | 0 |
| TpMuguga_01g00941 | 6 | 0 |
| TpMuguga_01g00943 | 1 | 3.139534884 |
| TpMuguga_01g00943 | 2 | 5.023255814 |
| TpMuguga_01g00943 | 3 | 0.209302326 |
| TpMuguga_01g00943 | 4 | 1.046511628 |
| TpMuguga_01g00943 | 5 | 1.046511628 |
| TpMuguga_01g00943 | 6 | 1.255813953 |
| TpMuguga_01g00943 | 7 | 1.674418605 |
| TpMuguga_01g00943 | 8 | 0 |
| TpMuguga_01g00944 | 1 | 0 |
| TpMuguga_01g00946 | 1 | 4.9 |
| TpMuguga_01g00946 | 2 | 2.1 |
| TpMuguga_01g00946 | 3 | 0 |
| TpMuguga_01g00946 | 4 | 0.35 |
| TpMuguga_01g00946 | 5 | 0 |
| TpMuguga_01g00946 | 6 | 0 |
| TpMuguga_01g00947 | 1 | 0 |
| TpMuguga_01g00948 | 1 | 0 |
| TpMuguga_01g00951 | 1 | 0.565445026 |

| | | |
|---|---|---|
| TpMuguga_01g00951 | 2 | 0 |
| TpMuguga_01g00950 | 1 | 0 |
| TpMuguga_01g00950 | 2 | 0 |
| TpMuguga_01g00950 | 3 | 0 |
| TpMuguga_01g00950 | 4 | 0 |
| TpMuguga_01g00950 | 5 | 0 |
| TpMuguga_01g00950 | 6 | 0.389275074 |
| TpMuguga_01g00955 | 1 | 45 |
| TpMuguga_01g00955 | 2 | 0 |
| TpMuguga_01g00955 | 3 | 0 |
| TpMuguga_01g00955 | 4 | 0 |
| TpMuguga_01g00955 | 5 | 0 |
| TpMuguga_01g00955 | 6 | 0 |
| TpMuguga_01g00955 | 7 | 0 |
| TpMuguga_01g00955 | 8 | 0 |
| TpMuguga_01g02280 | 1 | 0 |
| TpMuguga_01g02280 | 2 | 0 |
| TpMuguga_01g02280 | 3 | 0 |
| TpMuguga_01g02280 | 4 | 0 |
| TpMuguga_01g02280 | 5 | 0 |
| TpMuguga_01g02280 | 6 | 0 |
| TpMuguga_01g02280 | 7 | 0 |
| TpMuguga_01g02280 | 8 | 0 |
| TpMuguga_01g02280 | 9 | 0 |
| TpMuguga_01g02280 | 10 | 0 |
| TpMuguga_01g02280 | 11 | 0 |
| TpMuguga_01g00952 | 1 | 0 |
| TpMuguga_01g00952 | 2 | 0.954545455 |
| TpMuguga_01g00956 | 1 | 0.9 |
| TpMuguga_01g00956 | 2 | 0.6 |
| TpMuguga_01g00956 | 3 | 0 |
| TpMuguga_01g00956 | 4 | 0 |
| TpMuguga_01g00956 | 5 | 2.4 |
| TpMuguga_01g02335 | 1 | 0 |
| TpMuguga_01g02335 | 2 | 0 |
| TpMuguga_01g02335 | 3 | 0 |
| TpMuguga_01g02335 | 4 | 0 |
| TpMuguga_01g02335 | 5 | 0 |
| TpMuguga_01g02335 | 6 | 0 |
| TpMuguga_01g02335 | 7 | 0 |
| TpMuguga_01g02335 | 8 | 0 |
| TpMuguga_01g02335 | 9 | 0 |

| | | |
|---|---|---|
| TpMuguga_01g02335 | 10 | 0 |
| TpMuguga_01g02335 | 11 | 0 |
| TpMuguga_01g02335 | 12 | 35 |
| TpMuguga_01g02335 | 13 | 70 |
| TpMuguga_01g02400 | 1 | 0 |
| TpMuguga_01g02400 | 2 | 0 |
| TpMuguga_01g02400 | 3 | 0 |
| TpMuguga_01g02400 | 4 | 0 |
| TpMuguga_01g02400 | 5 | 0 |
| TpMuguga_01g02400 | 6 | 0 |
| TpMuguga_01g02400 | 7 | 0 |
| TpMuguga_01g00959 | 1 | 0 |
| TpMuguga_01g00961 | 1 | 0.153846154 |
| TpMuguga_01g00962 | 1 | 0 |
| TpMuguga_01g00962 | 2 | 0 |
| TpMuguga_01g00963 | 1 | 0.178571429 |
| TpMuguga_01g02790 | 1 | 0 |
| TpMuguga_01g02790 | 2 | 0 |
| TpMuguga_01g00964 | 1 | 0 |
| TpMuguga_01g00965 | 1 | 0 |
| TpMuguga_01g00967 | 1 | 3.9 |
| TpMuguga_01g00967 | 2 | 3.9 |
| TpMuguga_01g00968 | 1 | 0 |
| TpMuguga_01g00968 | 2 | 0 |
| TpMuguga_01g00968 | 3 | 0 |
| TpMuguga_01g00968 | 4 | 0 |
| TpMuguga_01g00968 | 5 | 0 |
| TpMuguga_01g00969 | 1 | 0 |
| TpMuguga_01g00969 | 2 | 0 |
| TpMuguga_01g00969 | 3 | 0 |
| TpMuguga_01g00969 | 4 | 0.007216495 |
| TpMuguga_01g00969 | 5 | 1.003092784 |
| TpMuguga_01g00969 | 6 | 2.778350515 |
| TpMuguga_01g00970 | 1 | 1.723515109 |
| TpMuguga_01g00970 | 2 | 0.10316082 |
| TpMuguga_01g00971 | 1 | 2 |
| TpMuguga_01g00971 | 2 | 2 |
| TpMuguga_01g00971 | 3 | 0 |
| TpMuguga_01g00971 | 4 | 0 |
| TpMuguga_01g00973 | 1 | 0.166666667 |
| TpMuguga_01g00973 | 2 | 0 |
| TpMuguga_01g00973 | 3 | 0 |

| | | |
|---|---|---|
| TpMuguga_01g00973 | 4 | 0 |
| TpMuguga_01g00973 | 5 | 0 |
| TpMuguga_01g02615 | 1 | 0 |
| TpMuguga_01g00974 | 1 | 0.875 |
| TpMuguga_01g00974 | 2 | 6.125 |
| TpMuguga_01g00974 | 3 | 0 |
| TpMuguga_01g00974 | 4 | 0 |
| TpMuguga_01g00974 | 5 | 1.75 |
| TpMuguga_01g00974 | 6 | 0 |
| TpMuguga_01g00976 | 1 | 2.53164557 |
| TpMuguga_01g00976 | 2 | 0.759493671 |
| TpMuguga_01g00976 | 3 | 0 |
| TpMuguga_01g00976 | 4 | 0 |
| TpMuguga_01g00976 | 5 | 0 |
| TpMuguga_01g00976 | 6 | 0 |
| TpMuguga_01g00976 | 7 | 0 |
| TpMuguga_01g00976 | 8 | 0 |
| TpMuguga_01g00976 | 9 | 0 |
| TpMuguga_01g00977 | 1 | 0 |
| TpMuguga_01g00977 | 2 | 0 |
| TpMuguga_01g00977 | 3 | 0 |
| TpMuguga_01g00979 | 1 | 0.035389282 |
| TpMuguga_01g00979 | 2 | 0.007077856 |
| TpMuguga_01g00979 | 3 | 0 |
| TpMuguga_01g00979 | 4 | 0 |
| TpMuguga_01g00979 | 5 | 0.031850354 |
| TpMuguga_01g00979 | 6 | 0 |
| TpMuguga_01g00978 | 1 | 0 |
| TpMuguga_01g00978 | 2 | 0 |
| TpMuguga_01g00978 | 3 | 0 |
| TpMuguga_01g00978 | 4 | 0 |
| TpMuguga_01g00978 | 5 | 0 |
| TpMuguga_01g00978 | 6 | 0 |
| TpMuguga_01g00978 | 7 | 0 |
| TpMuguga_01g00978 | 8 | 0 |
| TpMuguga_01g00978 | 9 | 0 |
| TpMuguga_01g00978 | 10 | 0 |
| TpMuguga_01g00981 | 1 | 0.181818182 |
| TpMuguga_01g00981 | 2 | 0.545454545 |
| TpMuguga_01g00981 | 3 | 0 |
| TpMuguga_01g00980 | 1 | 0.034972678 |
| TpMuguga_01g00980 | 2 | 0 |

| | | |
|---|---|---|
| TpMuguga_01g00983 | 1 | 0.366935484 |
| TpMuguga_01g00983 | 2 | 0.959677419 |
| TpMuguga_01g00983 | 3 | 0.762096774 |
| TpMuguga_01g00983 | 4 | 1.580645161 |
| TpMuguga_01g00983 | 5 | 0 |
| TpMuguga_01g00983 | 6 | 0 |
| TpMuguga_01g02675 | 1 | 0 |
| TpMuguga_01g02675 | 2 | 0 |
| TpMuguga_01g02675 | 3 | 0 |
| TpMuguga_01g02675 | 4 | 0 |
| TpMuguga_01g02675 | 5 | 0 |
| TpMuguga_01g02675 | 6 | 10 |
| TpMuguga_01g02675 | 7 | 0 |
| TpMuguga_01g02675 | 8 | 0 |
| TpMuguga_01g02675 | 9 | 0 |
| TpMuguga_01g02680 | 1 | 6 |
| TpMuguga_01g02680 | 2 | 0 |
| TpMuguga_01g02680 | 3 | 0 |
| TpMuguga_01g02680 | 4 | 0 |
| TpMuguga_01g00984 | 1 | 3.183673469 |
| TpMuguga_01g00984 | 2 | 0.816326531 |
| TpMuguga_01g00984 | 3 | 0.816326531 |
| TpMuguga_01g02425 | 1 | 0 |
| TpMuguga_01g00990 | 1 | 0.925996205 |
| TpMuguga_01g00990 | 2 | 0.280834915 |
| TpMuguga_01g00990 | 3 | 1.404174573 |
| TpMuguga_01g00991 | 1 | 0 |
| TpMuguga_01g00991 | 2 | 0 |
| TpMuguga_01g00991 | 3 | 0 |
| TpMuguga_01g00991 | 4 | 0 |
| TpMuguga_01g00992 | 1 | 0 |
| TpMuguga_01g00992 | 2 | 0 |
| TpMuguga_01g00994 | 1 | 0.727272727 |
| TpMuguga_01g00996 | 1 | 0.354223433 |
| TpMuguga_01g00996 | 2 | 0.081743869 |
| TpMuguga_01g00996 | 3 | 1.371480472 |
| TpMuguga_01g00996 | 4 | 3.065395095 |
| TpMuguga_01g00997 | 1 | 0.128205128 |
| TpMuguga_01g00997 | 2 | 0 |
| TpMuguga_01g00997 | 3 | 0.769230769 |
| TpMuguga_01g00997 | 4 | 0 |
| TpMuguga_01g00999 | 1 | 0 |

| | | |
|---|---|---|
| TpMuguga_01g00999 | 2 | 0 |
| TpMuguga_01g00999 | 3 | 0 |
| TpMuguga_01g00998 | 1 | 0 |
| TpMuguga_01g00998 | 2 | 0 |
| TpMuguga_01g00998 | 3 | 0 |
| TpMuguga_01g00998 | 4 | 0 |
| TpMuguga_01g00998 | 5 | 0 |
| TpMuguga_01g01000 | 1 | 0 |
| TpMuguga_01g01000 | 2 | 0 |
| TpMuguga_01g01000 | 3 | 0 |
| TpMuguga_01g01000 | 4 | 0 |
| TpMuguga_01g01002 | 1 | 0 |
| TpMuguga_01g01002 | 2 | 0 |
| TpMuguga_01g01003 | 1 | 0 |
| TpMuguga_01g01003 | 2 | 0 |
| TpMuguga_01g01003 | 3 | 0 |
| TpMuguga_01g01003 | 4 | 0 |
| TpMuguga_01g01003 | 5 | 0 |
| TpMuguga_01g01004 | 1 | 0 |
| TpMuguga_01g01004 | 2 | 2.75862069 |
| TpMuguga_01g01004 | 3 | 0 |
| TpMuguga_01g01004 | 4 | 0 |
| TpMuguga_01g02810 | 1 | 0 |
| TpMuguga_01g02810 | 2 | 0 |
| TpMuguga_01g02810 | 3 | 0 |
| TpMuguga_01g02810 | 4 | 0 |
| TpMuguga_01g02810 | 5 | 0 |
| TpMuguga_01g02810 | 6 | 0 |
| TpMuguga_01g02810 | 7 | 0 |
| TpMuguga_01g02810 | 8 | 0 |
| TpMuguga_01g01009 | 1 | 0.103448276 |
| TpMuguga_01g01009 | 2 | 0.806896552 |
| TpMuguga_01g01010 | 1 | 0 |
| TpMuguga_01g01010 | 2 | 0 |
| TpMuguga_01g01010 | 3 | 0 |
| TpMuguga_01g01010 | 4 | 0 |
| TpMuguga_01g01010 | 5 | 0 |
| TpMuguga_01g01013 | 1 | 0 |
| TpMuguga_01g01013 | 2 | 0 |
| TpMuguga_01g01012 | 1 | 0.666666667 |
| TpMuguga_01g01014 | 1 | 2.923076923 |
| TpMuguga_01g01014 | 2 | 0 |

| | | |
|---|---|---|
| TpMuguga_01g01014 | 3 | 0 |
| TpMuguga_01g01015 | 1 | 0 |
| TpMuguga_01g01015 | 2 | 0 |
| TpMuguga_01g01015 | 3 | 0 |
| TpMuguga_01g01016 | 1 | 0 |
| TpMuguga_01g01016 | 2 | 5.6 |
| TpMuguga_01g01016 | 3 | 0 |
| TpMuguga_01g01016 | 4 | 0 |
| TpMuguga_01g01016 | 5 | 0 |
| TpMuguga_01g01016 | 6 | 5.6 |
| TpMuguga_01g01017 | 1 | 0 |
| TpMuguga_01g01017 | 2 | 0 |
| TpMuguga_01g01017 | 3 | 0 |
| TpMuguga_01g01017 | 4 | 0.7 |
| TpMuguga_01g01017 | 5 | 0 |
| TpMuguga_01g01017 | 6 | 0 |
| TpMuguga_01g01018 | 1 | 1.066666667 |
| TpMuguga_01g01019 | 1 | 0 |
| TpMuguga_01g01019 | 2 | 0 |
| TpMuguga_01g01020 | 1 | 3.181818182 |
| TpMuguga_01g01020 | 2 | 2.272727273 |
| TpMuguga_01g01020 | 3 | 0.454545455 |
| TpMuguga_01g01020 | 4 | 0 |
| TpMuguga_01g01021 | 1 | 1.508474576 |
| TpMuguga_01g01021 | 2 | 0.118644068 |
| TpMuguga_01g01021 | 3 | 0 |
| TpMuguga_01g01021 | 4 | 0.101694915 |
| TpMuguga_01g01021 | 5 | 0 |
| TpMuguga_01g01022 | 1 | 0 |
| TpMuguga_01g01022 | 2 | 0 |
| TpMuguga_01g01024 | 1 | 0 |
| TpMuguga_01g01024 | 2 | 0 |
| TpMuguga_01g01024 | 3 | 0 |
| TpMuguga_01g01024 | 4 | 0 |
| TpMuguga_01g01024 | 5 | 0 |
| TpMuguga_01g01024 | 6 | 0 |
| TpMuguga_01g01024 | 7 | 1.635514019 |
| TpMuguga_01g01023 | 1 | 0 |
| TpMuguga_01g01023 | 2 | 0 |
| TpMuguga_01g01023 | 3 | 0 |
| TpMuguga_01g01023 | 4 | 0 |
| TpMuguga_01g01025 | 1 | 0.00152439 |

| | | |
|---|---|---|
| TpMuguga_01g01027 | 1 | 0.375 |
| TpMuguga_01g01027 | 2 | 0.375 |
| TpMuguga_01g01026 | 1 | 0 |
| TpMuguga_01g02245 | 1 | 0 |
| TpMuguga_01g02245 | 2 | 0 |
| TpMuguga_01g02245 | 3 | 0 |
| TpMuguga_01g02245 | 4 | 13.33333333 |
| TpMuguga_01g02435 | 1 | 0.972222222 |
| TpMuguga_01g02435 | 2 | 0 |
| TpMuguga_01g02435 | 3 | 0 |
| TpMuguga_01g02435 | 4 | 0.277777778 |
| TpMuguga_01g01032 | 1 | 0 |
| TpMuguga_01g01032 | 2 | 0 |
| TpMuguga_01g01034 | 1 | 0 |
| TpMuguga_01g01033 | 1 | 0 |
| TpMuguga_01g01033 | 2 | 23.71428571 |
| TpMuguga_01g01033 | 3 | 0 |
| TpMuguga_01g01035 | 1 | 0 |
| TpMuguga_01g01036 | 1 | 0.089285714 |
| TpMuguga_01g02300 | 1 | 4.814814815 |
| TpMuguga_01g01037 | 1 | 0 |
| TpMuguga_01g01037 | 2 | 0 |
| TpMuguga_01g01037 | 3 | 0 |
| TpMuguga_01g01037 | 4 | 0 |
| TpMuguga_01g01037 | 5 | 0 |
| TpMuguga_01g01037 | 6 | 0 |
| TpMuguga_01g01040 | 1 | 0 |
| TpMuguga_01g01040 | 2 | 0 |
| TpMuguga_01g01041 | 1 | 0 |
| TpMuguga_01g01041 | 2 | 0 |
| TpMuguga_01g01041 | 3 | 0 |
| TpMuguga_01g01041 | 4 | 0 |
| TpMuguga_01g01041 | 5 | 0 |
| TpMuguga_01g01041 | 6 | 0 |
| TpMuguga_01g01042 | 1 | 0 |
| TpMuguga_01g01042 | 2 | 0.105726872 |
| TpMuguga_01g01043 | 1 | 2.121212121 |
| TpMuguga_01g01043 | 2 | 0 |
| TpMuguga_01g01043 | 3 | 2.121212121 |
| TpMuguga_01g01043 | 4 | 2.121212121 |
| TpMuguga_01g01044 | 1 | 0 |
| TpMuguga_01g01044 | 2 | 0 |

| | | |
|---|---|---|
| TpMuguga_01g01044 | 3 | 0 |
| TpMuguga_01g01044 | 4 | 0 |
| TpMuguga_01g01044 | 5 | 0 |
| TpMuguga_01g01044 | 6 | 0 |
| TpMuguga_01g01044 | 7 | 0 |
| TpMuguga_01g01045 | 1 | 0 |
| TpMuguga_01g01045 | 2 | 0 |
| TpMuguga_01g01045 | 3 | 0 |
| TpMuguga_01g01045 | 4 | 0 |
| TpMuguga_01g01045 | 5 | 0 |
| TpMuguga_01g01045 | 6 | 0 |
| TpMuguga_01g01045 | 7 | 0 |
| TpMuguga_01g01045 | 8 | 0 |
| TpMuguga_01g01045 | 9 | 0 |
| TpMuguga_01g01045 | 10 | 0 |
| TpMuguga_01g01046 | 1 | 0 |
| TpMuguga_01g01046 | 2 | 3.5 |
| TpMuguga_01g01046 | 3 | 3.5 |
| TpMuguga_01g01046 | 4 | 0 |
| TpMuguga_01g01046 | 5 | 3.5 |
| TpMuguga_01g01046 | 6 | 0 |
| TpMuguga_01g01047 | 1 | 0 |
| TpMuguga_01g02595 | 1 | 0 |
| TpMuguga_01g02595 | 2 | 0 |
| TpMuguga_01g02595 | 3 | 1.25 |
| TpMuguga_01g02595 | 4 | 1.5 |
| TpMuguga_01g02595 | 5 | 0 |
| TpMuguga_01g02595 | 6 | 0 |
| TpMuguga_01g01050 | 1 | 0 |
| TpMuguga_01g01050 | 2 | 0.131578947 |
| TpMuguga_01g01050 | 3 | 0.394736842 |
| TpMuguga_01g01050 | 4 | 3.947368421 |
| TpMuguga_01g01050 | 5 | 1.052631579 |
| TpMuguga_01g01050 | 6 | 0 |
| TpMuguga_01g01050 | 7 | 0 |
| TpMuguga_01g01050 | 8 | 0 |
| TpMuguga_01g01050 | 9 | 0 |
| TpMuguga_01g01051 | 1 | 0.574850299 |
| TpMuguga_01g01052 | 1 | 0 |
| TpMuguga_01g01052 | 2 | 0 |
| TpMuguga_01g01052 | 3 | 0 |
| TpMuguga_01g01052 | 4 | 0 |

| | | |
|---|---|---|
| TpMuguga_01g01054 | 1 | 0 |
| TpMuguga_01g01054 | 2 | 0 |
| TpMuguga_01g01054 | 3 | 0.062084257 |
| TpMuguga_01g01053 | 1 | 0 |
| TpMuguga_01g01053 | 2 | 2.285714286 |
| TpMuguga_01g01053 | 3 | 0 |
| TpMuguga_01g01053 | 4 | 0 |
| TpMuguga_01g01053 | 5 | 0 |
| TpMuguga_01g01053 | 6 | 0 |
| TpMuguga_01g01053 | 7 | 0 |
| TpMuguga_01g01055 | 1 | 0 |
| TpMuguga_01g01055 | 2 | 0 |
| TpMuguga_01g01055 | 3 | 0 |
| TpMuguga_01g01055 | 4 | 0 |
| TpMuguga_01g01055 | 5 | 0 |
| TpMuguga_01g01055 | 6 | 1.28 |
| TpMuguga_01g01055 | 7 | 0 |
| TpMuguga_01g01057 | 1 | 0 |
| TpMuguga_01g01058 | 1 | 0.5 |
| TpMuguga_01g01058 | 2 | 0 |
| TpMuguga_01g01059 | 1 | 16 |
| TpMuguga_01g01059 | 2 | 0 |
| TpMuguga_01g01059 | 3 | 0 |
| TpMuguga_01g01059 | 4 | 0 |
| TpMuguga_01g01059 | 5 | 0 |
| TpMuguga_01g01059 | 6 | 0 |
| TpMuguga_01g01059 | 7 | 0 |
| TpMuguga_01g01059 | 8 | 0 |
| TpMuguga_01g01059 | 9 | 0 |
| TpMuguga_01g01059 | 10 | 0 |
| TpMuguga_01g01059 | 11 | 8 |
| TpMuguga_01g01060 | 1 | 0 |
| TpMuguga_01g01060 | 2 | 0 |
| TpMuguga_01g01060 | 3 | 0 |
| TpMuguga_01g01060 | 4 | 0 |
| TpMuguga_01g01060 | 5 | 0 |
| TpMuguga_01g01060 | 6 | 0 |
| TpMuguga_01g01060 | 7 | 0 |
| TpMuguga_01g01060 | 8 | 0 |
| TpMuguga_01g01061 | 1 | 0 |
| TpMuguga_01g01061 | 2 | 0 |
| TpMuguga_01g01061 | 3 | 9.333333333 |

| | | |
|---|---|---|
| TpMuguga_01g01061 | 4 | 3.5 |
| TpMuguga_01g01061 | 5 | 5.833333333 |
| TpMuguga_01g01061 | 6 | 0 |
| TpMuguga_01g01062 | 1 | 0 |
| TpMuguga_01g01062 | 2 | 0 |
| TpMuguga_01g01062 | 3 | 0 |
| TpMuguga_01g01062 | 4 | 0 |
| TpMuguga_01g01062 | 5 | 0 |
| TpMuguga_01g01062 | 6 | 0 |
| TpMuguga_01g01062 | 7 | 0 |
| TpMuguga_01g02105 | 1 | 6 |
| TpMuguga_01g02105 | 2 | 1 |
| TpMuguga_01g02105 | 3 | 3 |
| TpMuguga_01g02105 | 4 | 12 |
| TpMuguga_01g02105 | 5 | 0 |
| TpMuguga_01g02105 | 6 | 2 |
| TpMuguga_01g02105 | 7 | 8 |
| TpMuguga_01g02610 | 1 | 0 |
| TpMuguga_01g02610 | 2 | 0 |
| TpMuguga_01g02455 | 1 | 8 |
| TpMuguga_01g02455 | 2 | 5.714285714 |
| TpMuguga_01g02455 | 3 | 2.142857143 |
| TpMuguga_01g02455 | 4 | 0 |
| TpMuguga_01g02455 | 5 | 0 |
| TpMuguga_01g02455 | 6 | 0 |
| TpMuguga_01g02455 | 7 | 0 |
| TpMuguga_01g02455 | 8 | 0 |
| TpMuguga_01g01065 | 1 | 27 |
| TpMuguga_01g01065 | 2 | 0 |
| TpMuguga_01g01065 | 3 | 0 |
| TpMuguga_01g01065 | 4 | 0 |
| TpMuguga_01g01065 | 5 | 0 |
| TpMuguga_01g01065 | 6 | 1 |
| TpMuguga_01g01064 | 1 | 0.955974843 |
| TpMuguga_01g01064 | 2 | 1.559748428 |
| TpMuguga_01g01064 | 3 | 1.081761006 |
| TpMuguga_01g01066 | 1 | 0 |
| TpMuguga_01g01066 | 2 | 0 |
| TpMuguga_01g01066 | 3 | 0.615384615 |
| TpMuguga_01g01066 | 4 | 0 |
| TpMuguga_01g01066 | 5 | 0 |
| TpMuguga_01g01066 | 6 | 1.230769231 |

| | | |
|---|---|---|
| TpMuguga_01g01066 | 7 | 11.69230769 |
| TpMuguga_01g01067 | 1 | 1.891891892 |
| TpMuguga_01g01067 | 2 | 2.432432432 |
| TpMuguga_01g01067 | 3 | 2.972972973 |
| TpMuguga_01g01067 | 4 | 1.689189189 |
| TpMuguga_01g02815 | 1 | 0 |
| TpMuguga_01g02815 | 2 | 0 |
| TpMuguga_01g02815 | 3 | 0 |
| TpMuguga_01g01068 | 1 | 0 |
| TpMuguga_01g01068 | 2 | 0 |
| TpMuguga_01g01068 | 3 | 0 |
| TpMuguga_01g01071 | 1 | 0 |
| TpMuguga_01g01071 | 2 | 2.068965517 |
| TpMuguga_01g01071 | 3 | 1.034482759 |
| TpMuguga_01g01071 | 4 | 0 |
| TpMuguga_01g01071 | 5 | 0 |
| TpMuguga_01g01070 | 1 | 0.352941176 |
| TpMuguga_01g01073 | 1 | 0 |
| TpMuguga_01g01073 | 2 | 0 |
| TpMuguga_01g01073 | 3 | 0 |
| TpMuguga_01g01073 | 4 | 0 |
| TpMuguga_01g01072 | 1 | 0 |
| TpMuguga_01g01072 | 2 | 0 |
| TpMuguga_01g01072 | 3 | 0 |
| TpMuguga_01g01072 | 4 | 0 |
| TpMuguga_01g01072 | 5 | 0 |
| TpMuguga_01g01072 | 6 | 0 |
| TpMuguga_01g01072 | 7 | 0 |
| TpMuguga_01g01072 | 8 | 0 |
| TpMuguga_01g01074 | 1 | 0 |
| TpMuguga_01g01074 | 2 | 0 |
| TpMuguga_01g01075 | 1 | 0.12 |
| TpMuguga_01g01076 | 1 | 0.970711297 |
| TpMuguga_01g01077 | 1 | 0.929824561 |
| TpMuguga_01g01078 | 1 | 0.021276596 |
| TpMuguga_01g01079 | 1 | 0 |
| TpMuguga_01g01080 | 1 | 0 |
| TpMuguga_01g01080 | 2 | 0 |
| TpMuguga_01g01080 | 3 | 0 |
| TpMuguga_01g01080 | 4 | 0 |
| TpMuguga_01g01082 | 1 | 0 |
| TpMuguga_01g01081 | 1 | 1.6 |

| | | |
|---|---|---|
| TpMuguga_01g01083 | 1 | 0 |
| TpMuguga_01g01085 | 1 | 0.375 |
| TpMuguga_01g01085 | 2 | 0 |
| TpMuguga_01g01085 | 3 | 0.1875 |
| TpMuguga_01g01085 | 4 | 0.375 |
| TpMuguga_01g01085 | 5 | 2.0625 |
| TpMuguga_01g01086 | 1 | 0.347970174 |
| TpMuguga_01g02035 | 1 | 0 |
| TpMuguga_01g01087 | 1 | 1.813953488 |
| TpMuguga_01g01087 | 2 | 0.76744186 |
| TpMuguga_01g01088 | 1 | 0 |
| TpMuguga_01g01088 | 2 | 0 |
| TpMuguga_01g01088 | 3 | 0 |
| TpMuguga_01g01088 | 4 | 0 |
| TpMuguga_01g01088 | 5 | 0 |
| TpMuguga_01g01088 | 6 | 0 |
| TpMuguga_01g01088 | 7 | 1.565217391 |
| TpMuguga_01g01088 | 8 | 0 |
| TpMuguga_01g01088 | 9 | 0 |
| TpMuguga_01g01088 | 10 | 0 |
| TpMuguga_01g01088 | 11 | 10.95652174 |
| TpMuguga_01g01089 | 1 | 0.170212766 |
| TpMuguga_01g01090 | 1 | 1.130434783 |
| TpMuguga_01g01090 | 2 | 0.565217391 |
| TpMuguga_01g01092 | 1 | 0 |
| TpMuguga_01g01092 | 2 | 0 |
| TpMuguga_01g01092 | 3 | 0 |
| TpMuguga_01g01092 | 4 | 0 |
| TpMuguga_01g01092 | 5 | 0 |
| TpMuguga_01g01092 | 6 | 0 |
| TpMuguga_01g02260 | 1 | 0 |
| TpMuguga_01g02260 | 2 | 0 |
| TpMuguga_01g02260 | 3 | 0 |
| TpMuguga_01g02260 | 4 | 0 |
| TpMuguga_01g02260 | 5 | 0 |
| TpMuguga_01g02260 | 6 | 0 |
| TpMuguga_01g02290 | 1 | 0 |
| TpMuguga_01g02290 | 2 | 0 |
| TpMuguga_01g02290 | 3 | 0 |
| TpMuguga_01g02290 | 4 | 0 |
| TpMuguga_01g02290 | 5 | 6.545454545 |
| TpMuguga_01g02290 | 6 | 0 |

| | | |
|---|---|---|
| TpMuguga_01g02290 | 7 | 0 |
| TpMuguga_01g02290 | 8 | 3.272727273 |
| TpMuguga_01g01095 | 1 | 0 |
| TpMuguga_01g01095 | 2 | 0 |
| TpMuguga_01g01095 | 3 | 0 |
| TpMuguga_01g01095 | 4 | 5 |
| TpMuguga_01g01096 | 1 | 1.818181818 |
| TpMuguga_01g01097 | 1 | 0 |
| TpMuguga_01g01098 | 1 | 0 |
| TpMuguga_01g01098 | 2 | 0 |
| TpMuguga_01g01098 | 3 | 0 |
| TpMuguga_01g01098 | 4 | 0 |
| TpMuguga_01g01098 | 5 | 0 |
| TpMuguga_01g01098 | 6 | 0 |
| TpMuguga_01g01099 | 1 | 0 |
| TpMuguga_01g01099 | 2 | 0 |
| TpMuguga_01g01099 | 3 | 0 |
| TpMuguga_01g01099 | 4 | 0 |
| TpMuguga_01g01099 | 5 | 0 |
| TpMuguga_01g01099 | 6 | 0 |
| TpMuguga_01g01099 | 7 | 0 |
| TpMuguga_01g01100 | 1 | 1.046153846 |
| TpMuguga_01g01100 | 2 | 0.584615385 |
| TpMuguga_01g01100 | 3 | 1.015384615 |
| TpMuguga_01g01101 | 1 | 0 |
| TpMuguga_01g01101 | 2 | 0 |
| TpMuguga_01g01101 | 3 | 0 |
| TpMuguga_01g01101 | 4 | 3.548387097 |
| TpMuguga_01g01101 | 5 | 1.064516129 |
| TpMuguga_01g01101 | 6 | 6.387096774 |
| TpMuguga_01g01101 | 7 | 12.41935484 |
| TpMuguga_01g01101 | 8 | 3.903225806 |
| TpMuguga_01g01101 | 9 | 0 |
| TpMuguga_01g01101 | 10 | 0 |
| TpMuguga_01g01102 | 1 | 3.125 |
| TpMuguga_01g01102 | 2 | 0 |
| TpMuguga_01g01102 | 3 | 0 |
| TpMuguga_01g01102 | 4 | 0.625 |
| TpMuguga_01g01103 | 1 | 0 |
| TpMuguga_01g01106 | 1 | 0 |
| TpMuguga_01g01107 | 1 | 0 |
| TpMuguga_01g01107 | 2 | 0 |

| | | |
|---|---|---|
| TpMuguga_01g01107 | 3 | 0 |
| TpMuguga_01g01107 | 4 | 0 |
| TpMuguga_01g01107 | 5 | 0 |
| TpMuguga_01g01107 | 6 | 0 |
| TpMuguga_01g01109 | 1 | 0 |
| TpMuguga_01g01109 | 2 | 0.06122449 |
| TpMuguga_01g01110 | 1 | 0 |
| TpMuguga_01g01110 | 2 | 0 |
| TpMuguga_01g01110 | 3 | 0 |
| TpMuguga_01g01112 | 1 | 0.051001821 |
| TpMuguga_01g01112 | 2 | 0 |
| TpMuguga_01g01112 | 3 | 0.422586521 |
| TpMuguga_01g01113 | 1 | 0 |
| TpMuguga_01g01114 | 1 | 0 |
| TpMuguga_01g01114 | 2 | 0 |
| TpMuguga_01g01114 | 3 | 0 |
| TpMuguga_01g01115 | 1 | 0 |
| TpMuguga_01g01116 | 1 | 0 |
| TpMuguga_01g01116 | 2 | 0 |
| TpMuguga_01g01116 | 3 | 0 |
| TpMuguga_01g02805 | 1 | 0 |
| TpMuguga_01g02805 | 2 | 0 |
| TpMuguga_01g02805 | 3 | 0 |
| TpMuguga_01g02805 | 4 | 0 |
| TpMuguga_01g02805 | 5 | 0 |
| TpMuguga_01g01117 | 1 | 0.286624204 |
| TpMuguga_01g01117 | 2 | 0.171974522 |
| TpMuguga_01g02215 | 1 | 0 |
| TpMuguga_01g02215 | 2 | 0 |
| TpMuguga_01g02870 | 1 | 2.25 |
| TpMuguga_01g02870 | 2 | 2.25 |
| TpMuguga_01g02870 | 3 | 13.5 |
| TpMuguga_01g02870 | 4 | 9 |
| TpMuguga_01g02870 | 5 | 3 |
| TpMuguga_01g01120 | 1 | 2.037735849 |
| TpMuguga_01g01120 | 2 | 0.075471698 |
| TpMuguga_01g01120 | 3 | 0 |
| TpMuguga_01g01122 | 1 | 0 |
| TpMuguga_01g01122 | 2 | 0 |
| TpMuguga_01g01122 | 3 | 0 |
| TpMuguga_01g02750 | 1 | 11.2 |
| TpMuguga_01g02750 | 2 | 0 |

| | | |
|---|---|---|
| TpMuguga_01g02750 | 3 | 0 |
| TpMuguga_01g02750 | 4 | 0 |
| TpMuguga_01g02750 | 5 | 0 |
| TpMuguga_01g02750 | 6 | 0 |
| TpMuguga_01g01125 | 1 | 0 |
| TpMuguga_01g01125 | 2 | 0 |
| TpMuguga_01g01125 | 3 | 0 |
| TpMuguga_01g01125 | 4 | 0 |
| TpMuguga_01g01125 | 5 | 0 |
| TpMuguga_01g01125 | 6 | 0 |
| TpMuguga_01g01125 | 7 | 0 |
| TpMuguga_01g01125 | 8 | 0 |
| TpMuguga_01g02365 | 1 | 0 |
| TpMuguga_01g02365 | 2 | 0 |
| TpMuguga_01g02365 | 3 | 0 |
| TpMuguga_01g02365 | 4 | 0 |
| TpMuguga_01g02365 | 5 | 0 |
| TpMuguga_01g02365 | 6 | 7 |
| TpMuguga_01g02660 | 1 | 0 |
| TpMuguga_01g02660 | 2 | 0 |
| TpMuguga_01g02660 | 3 | 0 |
| TpMuguga_01g02655 | 1 | 0 |
| TpMuguga_01g02655 | 2 | 0 |
| TpMuguga_01g02655 | 3 | 0.285714286 |
| TpMuguga_01g01129 | 1 | 4.976470588 |
| TpMuguga_01g01129 | 2 | 1.023529412 |
| TpMuguga_01g01131 | 1 | 0 |
| TpMuguga_01g01131 | 2 | 0 |
| TpMuguga_01g01131 | 3 | 0 |
| TpMuguga_01g01132 | 1 | 0 |
| TpMuguga_01g01132 | 2 | 0 |
| TpMuguga_01g01132 | 3 | 0 |
| TpMuguga_01g01132 | 4 | 0 |
| TpMuguga_01g01132 | 5 | 0 |
| TpMuguga_01g01133 | 1 | 0 |
| TpMuguga_01g01133 | 2 | 0 |
| TpMuguga_01g01133 | 3 | 0 |
| TpMuguga_01g01133 | 4 | 0 |
| TpMuguga_01g01135 | 1 | 0.631578947 |
| TpMuguga_01g01135 | 2 | 0 |
| TpMuguga_01g01135 | 3 | 1.894736842 |
| TpMuguga_01g01138 | 1 | 0 |

| | | |
|---|---|---|
| TpMuguga_01g01138 | 2 | 0 |
| TpMuguga_01g01138 | 3 | 0 |
| TpMuguga_01g01138 | 4 | 0 |
| TpMuguga_01g01138 | 5 | 0 |
| TpMuguga_01g01138 | 6 | 0 |
| TpMuguga_01g01138 | 7 | 0 |
| TpMuguga_01g01138 | 8 | 0 |
| TpMuguga_01g01137 | 1 | 1 |
| TpMuguga_01g01137 | 2 | 7 |
| TpMuguga_01g01137 | 3 | 12 |
| TpMuguga_01g01137 | 4 | 6 |
| TpMuguga_01g01137 | 5 | 0 |
| TpMuguga_01g01137 | 6 | 13 |
| TpMuguga_01g01139 | 1 | 0 |
| TpMuguga_01g01139 | 2 | 0 |
| TpMuguga_01g01139 | 3 | 4 |
| TpMuguga_01g01140 | 1 | 0 |
| TpMuguga_01g01140 | 2 | 0 |
| TpMuguga_01g01142 | 1 | 0 |
| TpMuguga_01g01142 | 2 | 0 |
| TpMuguga_01g01142 | 3 | 0 |
| TpMuguga_01g01142 | 4 | 1.666666667 |
| TpMuguga_01g01143 | 1 | 0 |
| TpMuguga_01g01145 | 1 | 0 |
| TpMuguga_01g02700 | 1 | 0 |
| TpMuguga_01g02700 | 2 | 0 |
| TpMuguga_01g02700 | 3 | 0 |
| TpMuguga_01g02700 | 4 | 0 |
| TpMuguga_01g02700 | 5 | 0 |
| TpMuguga_01g02700 | 6 | 0 |
| TpMuguga_01g02700 | 7 | 0 |
| TpMuguga_01g02700 | 8 | 0 |
| TpMuguga_01g02700 | 9 | 0 |
| TpMuguga_01g02700 | 10 | 0 |
| TpMuguga_01g02700 | 11 | 0 |
| TpMuguga_01g02705 | 1 | 0 |
| TpMuguga_01g02705 | 2 | 0 |
| TpMuguga_01g01148 | 1 | 0 |
| TpMuguga_01g01148 | 2 | 0 |
| TpMuguga_01g01148 | 3 | 0 |
| TpMuguga_01g01148 | 4 | 1.777777778 |
| TpMuguga_01g01148 | 5 | 0.666666667 |

| | | |
|---|---|---|
| TpMuguga_01g01148 | 6 | 0.777777778 |
| TpMuguga_01g01148 | 7 | 1.888888889 |
| TpMuguga_01g01149 | 1 | 0 |
| TpMuguga_01g01149 | 2 | 0 |
| TpMuguga_01g01149 | 3 | 0.25 |
| TpMuguga_01g01149 | 4 | 0 |
| TpMuguga_01g01149 | 5 | 0 |
| TpMuguga_01g01149 | 6 | 0.75 |
| TpMuguga_01g01151 | 1 | 30 |
| TpMuguga_01g01151 | 2 | 0 |
| TpMuguga_01g01151 | 3 | 0 |
| TpMuguga_01g01151 | 4 | 0 |
| TpMuguga_01g01151 | 5 | 0 |
| TpMuguga_01g01151 | 6 | 0 |
| TpMuguga_01g01151 | 7 | 45 |
| TpMuguga_01g01151 | 8 | 30 |
| TpMuguga_01g01151 | 9 | 0 |
| TpMuguga_01g01150 | 1 | 0 |
| TpMuguga_01g01150 | 2 | 0 |
| TpMuguga_01g01150 | 3 | 0 |
| TpMuguga_01g01150 | 4 | 0 |
| TpMuguga_01g01150 | 5 | 0 |
| TpMuguga_01g01150 | 6 | 0.5 |
| TpMuguga_01g01150 | 7 | 0 |
| TpMuguga_01g01150 | 8 | 0.5 |
| TpMuguga_01g01153 | 1 | 0 |
| TpMuguga_01g01153 | 2 | 0 |
| TpMuguga_01g01153 | 3 | 0.032171582 |
| TpMuguga_01g01153 | 4 | 0.498659517 |
| TpMuguga_01g01153 | 5 | 0 |
| TpMuguga_01g01152 | 1 | 0.494505495 |
| TpMuguga_01g01152 | 2 | 0 |
| TpMuguga_01g01152 | 3 | 0.494505495 |
| TpMuguga_01g01152 | 4 | 1.978021978 |
| TpMuguga_01g01156 | 1 | 0 |
| TpMuguga_01g01156 | 2 | 2.5 |
| TpMuguga_01g01156 | 3 | 0.416666667 |
| TpMuguga_01g01156 | 4 | 2.5 |
| TpMuguga_01g02415 | 1 | 2.675675676 |
| TpMuguga_01g02415 | 2 | 3.567567568 |
| TpMuguga_01g02415 | 3 | 7.432432432 |
| TpMuguga_01g02415 | 4 | 1.189189189 |

| | | |
|---|---|---|
| TpMuguga_01g02415 | 5 | 0 |
| TpMuguga_01g02415 | 6 | 0 |
| TpMuguga_01g02415 | 7 | 0 |
| TpMuguga_01g02415 | 8 | 0 |
| TpMuguga_01g02415 | 9 | 0 |
| TpMuguga_01g02415 | 10 | 0 |
| TpMuguga_01g01157 | 1 | 0 |
| TpMuguga_01g01157 | 2 | 0 |
| TpMuguga_01g01157 | 3 | 7.857142857 |
| TpMuguga_01g01157 | 4 | 11.42857143 |
| TpMuguga_01g01157 | 5 | 0 |
| TpMuguga_01g01157 | 6 | 0 |
| TpMuguga_01g01157 | 7 | 0 |
| TpMuguga_01g01157 | 8 | 0 |
| TpMuguga_01g01157 | 9 | 0 |
| TpMuguga_01g01158 | 1 | 0 |
| TpMuguga_01g01158 | 2 | 0 |
| TpMuguga_01g01158 | 3 | 10.90909091 |
| TpMuguga_01g01158 | 4 | 0 |
| TpMuguga_01g01158 | 5 | 0 |
| TpMuguga_01g01158 | 6 | 1.636363636 |
| TpMuguga_01g01158 | 7 | 0 |
| TpMuguga_01g01158 | 8 | 0 |
| TpMuguga_01g01158 | 9 | 0 |
| TpMuguga_01g01158 | 10 | 0 |
| TpMuguga_01g01158 | 11 | 7.090909091 |
| TpMuguga_01g01159 | 1 | 0 |
| TpMuguga_01g01159 | 2 | 0 |
| TpMuguga_01g01160 | 1 | 0 |
| TpMuguga_01g01161 | 1 | 0 |
| TpMuguga_01g01161 | 2 | 0 |
| TpMuguga_01g01162 | 1 | 0 |
| TpMuguga_01g01162 | 2 | 0 |
| TpMuguga_01g01162 | 3 | 0 |
| TpMuguga_01g01162 | 4 | 0 |
| TpMuguga_01g01162 | 5 | 0 |
| TpMuguga_01g01162 | 6 | 0 |
| TpMuguga_01g01163 | 1 | 5.177419355 |
| TpMuguga_01g01163 | 2 | 0.870967742 |
| TpMuguga_01g01163 | 3 | 1.209677419 |
| TpMuguga_01g01163 | 4 | 0.338709677 |
| TpMuguga_01g01163 | 5 | 0 |

| | | |
|---|---|---|
| TpMuguga_01g01165 | 1 | 0 |
| TpMuguga_01g01165 | 2 | 0 |
| TpMuguga_01g01165 | 3 | 0 |
| TpMuguga_01g01165 | 4 | 0 |
| TpMuguga_01g01165 | 5 | 0 |
| TpMuguga_01g01165 | 6 | 0 |
| TpMuguga_01g01165 | 7 | 0 |
| TpMuguga_01g01166 | 1 | 0 |
| TpMuguga_01g01166 | 2 | 0 |
| TpMuguga_01g01166 | 3 | 0 |
| TpMuguga_01g01167 | 1 | 0 |
| TpMuguga_01g01167 | 2 | 0 |
| TpMuguga_01g01167 | 3 | 0 |
| TpMuguga_01g01167 | 4 | 0 |
| TpMuguga_01g01168 | 1 | 1.825396825 |
| TpMuguga_01g01168 | 2 | 0 |
| TpMuguga_01g01168 | 3 | 0 |
| TpMuguga_01g01168 | 4 | 0.396825397 |
| TpMuguga_01g02785 | 1 | 0 |
| TpMuguga_01g02785 | 2 | 0 |
| TpMuguga_01g02785 | 3 | 0 |
| TpMuguga_01g02785 | 4 | 0 |
| TpMuguga_01g02785 | 5 | 0 |
| TpMuguga_01g02785 | 6 | 0 |
| TpMuguga_01g01174 | 1 | 0.586206897 |
| TpMuguga_01g01173 | 1 | 0.705882353 |
| TpMuguga_01g01173 | 2 | 0 |
| TpMuguga_01g01173 | 3 | 0 |
| TpMuguga_01g01175 | 1 | 0 |
| TpMuguga_01g01175 | 2 | 0 |
| TpMuguga_01g02085 | 1 | 0 |
| TpMuguga_01g02085 | 2 | 0 |
| TpMuguga_01g02085 | 3 | 0 |
| TpMuguga_01g02090 | 1 | 0 |
| TpMuguga_01g02090 | 2 | 0.090909091 |
| TpMuguga_01g02090 | 3 | 0 |
| TpMuguga_01g02090 | 4 | 0.181818182 |
| TpMuguga_01g01180 | 1 | 0 |
| TpMuguga_01g01180 | 2 | 0 |
| TpMuguga_01g01180 | 3 | 0.025380711 |
| TpMuguga_01g01180 | 4 | 0 |
| TpMuguga_01g01181 | 1 | 0.774193548 |

| | | |
|---|---|---|
| TpMuguga_01g01182 | 1 | 0.221052632 |
| TpMuguga_01g01182 | 2 | 2.021052632 |
| TpMuguga_01g01183 | 1 | 0 |
| TpMuguga_01g01183 | 2 | 0 |
| TpMuguga_01g01183 | 3 | 0 |
| TpMuguga_01g01183 | 4 | 0 |
| TpMuguga_01g01183 | 5 | 0 |
| TpMuguga_01g01183 | 6 | 0 |
| TpMuguga_01g01183 | 7 | 0 |
| TpMuguga_01g01183 | 8 | 1.125 |
| TpMuguga_01g01184 | 1 | 0 |
| TpMuguga_01g02645 | 1 | 0 |
| TpMuguga_01g02645 | 2 | 0 |
| TpMuguga_01g01188 | 1 | 0 |
| TpMuguga_01g01188 | 2 | 0 |
| TpMuguga_01g01188 | 3 | 0 |
| TpMuguga_01g01188 | 4 | 0 |
| TpMuguga_01g01188 | 5 | 0 |
| TpMuguga_01g01188 | 6 | 0 |
| TpMuguga_01g01189 | 1 | 0 |
| TpMuguga_01g01189 | 2 | 3.741935484 |
| TpMuguga_01g01189 | 3 | 2.322580645 |
| TpMuguga_01g01190 | 1 | 0 |
| TpMuguga_01g01190 | 2 | 0 |
| TpMuguga_01g01191 | 1 | 0.881355932 |
| TpMuguga_01g01191 | 2 | 1.355932203 |
| TpMuguga_01g01191 | 3 | 2.169491525 |
| TpMuguga_01g01191 | 4 | 5.084745763 |
| TpMuguga_01g01191 | 5 | 1.949152542 |
| TpMuguga_01g01191 | 6 | 1.677966102 |
| TpMuguga_01g01191 | 7 | 1.288135593 |
| TpMuguga_01g01191 | 8 | 0 |
| TpMuguga_01g01191 | 9 | 0 |
| TpMuguga_01g01191 | 10 | 0.542372881 |
| TpMuguga_01g01192 | 1 | 2.471910112 |
| TpMuguga_01g01192 | 2 | 0.449438202 |
| TpMuguga_01g01192 | 3 | 0 |
| TpMuguga_01g01192 | 4 | 0 |
| TpMuguga_01g01192 | 5 | 0 |
| TpMuguga_01g01192 | 6 | 0 |
| TpMuguga_01g01192 | 7 | 0 |
| TpMuguga_01g01192 | 8 | 0 |

| | | |
|---|---|---|
| TpMuguga_01g01192 | 9 | 0 |
| TpMuguga_01g01195 | 1 | 0.074725275 |
| TpMuguga_01g01195 | 2 | 0.052747253 |
| TpMuguga_01g01195 | 3 | 0.083516484 |
| TpMuguga_01g01194 | 1 | 0 |
| TpMuguga_01g01194 | 2 | 0.015317287 |
| TpMuguga_01g01194 | 3 | 0.038293217 |
| TpMuguga_01g01194 | 4 | 0 |
| TpMuguga_01g01194 | 5 | 0 |
| TpMuguga_01g01194 | 6 | 0 |
| TpMuguga_01g01193 | 1 | 0 |
| TpMuguga_01g01193 | 2 | 0 |
| TpMuguga_01g01193 | 3 | 0 |
| TpMuguga_01g01193 | 4 | 0 |
| TpMuguga_01g01193 | 5 | 0 |
| TpMuguga_01g01196 | 1 | 0.9 |
| TpMuguga_01g01196 | 2 | 1.9 |
| TpMuguga_01g01196 | 3 | 0.166666667 |
| TpMuguga_01g01196 | 4 | 0.066666667 |
| TpMuguga_01g01196 | 5 | 0.033333333 |
| TpMuguga_01g01199 | 1 | 0.865384615 |
| TpMuguga_01g01199 | 2 | 0 |
| TpMuguga_01g01200 | 1 | 0.6 |
| TpMuguga_01g01200 | 2 | 1.1 |
| TpMuguga_01g01200 | 3 | 0 |
| TpMuguga_01g01200 | 4 | 0 |
| TpMuguga_01g01200 | 5 | 0 |
| TpMuguga_01g01200 | 6 | 0 |
| TpMuguga_01g01200 | 7 | 0.6 |
| TpMuguga_01g01200 | 8 | 6.7 |
| TpMuguga_01g01201 | 1 | 3 |
| TpMuguga_01g01201 | 2 | 3 |
| TpMuguga_01g01203 | 1 | 0.00907441 |
| TpMuguga_01g01204 | 1 | 0 |
| TpMuguga_01g01204 | 2 | 0.000746083 |
| TpMuguga_01g02320 | 1 | 0.637119114 |
| TpMuguga_01g02320 | 2 | 0.650969529 |
| TpMuguga_01g02320 | 3 | 2.645429363 |
| TpMuguga_01g02320 | 4 | 0.927977839 |
| TpMuguga_01g01207 | 1 | 0.266666667 |
| TpMuguga_01g01208 | 1 | 0 |
| TpMuguga_01g01208 | 2 | 0 |

| | | |
|---|---|---|
| TpMuguga_01g01210 | 1 | 1.532751092 |
| TpMuguga_01g01210 | 2 | 0.458515284 |
| TpMuguga_01g01210 | 3 | 0.104803493 |
| TpMuguga_01g01210 | 4 | 0 |
| TpMuguga_01g01210 | 5 | 0.104803493 |
| TpMuguga_01g01209 | 1 | 0 |
| TpMuguga_01g01209 | 2 | 0.005724098 |
| TpMuguga_01g01209 | 3 | 0.008013738 |
| TpMuguga_01g01211 | 1 | 0 |
| TpMuguga_01g01211 | 2 | 1.68 |
| TpMuguga_01g01211 | 3 | 0 |
| TpMuguga_01g01211 | 4 | 0 |
| TpMuguga_01g01211 | 5 | 0 |
| TpMuguga_01g01211 | 6 | 0 |
| TpMuguga_01g01213 | 1 | 0 |
| TpMuguga_01g01213 | 2 | 0 |
| TpMuguga_01g01213 | 3 | 0 |
| TpMuguga_01g01213 | 4 | 0 |
| TpMuguga_01g01213 | 5 | 0.567567568 |
| TpMuguga_01g01213 | 6 | 4.540540541 |
| TpMuguga_01g01214 | 1 | 0.513157895 |
| TpMuguga_01g01214 | 2 | 1.894736842 |
| TpMuguga_01g01215 | 1 | 0 |
| TpMuguga_01g01215 | 2 | 0.774193548 |
| TpMuguga_01g01215 | 3 | 0.516129032 |
| TpMuguga_01g01216 | 1 | 0 |
| TpMuguga_01g01216 | 2 | 0 |
| TpMuguga_01g01216 | 3 | 0 |
| TpMuguga_01g01216 | 4 | 0 |
| TpMuguga_01g01216 | 5 | 0 |
| TpMuguga_01g01225 | 1 | 0.36 |
| TpMuguga_02g02590 | 1 | 0.741847826 |
| TpMuguga_02g02055 | 1 | 0.032967033 |
| TpMuguga_02g00018 | 1 | 0.010484928 |
| TpMuguga_02g00021 | 1 | 1.625570776 |
| TpMuguga_02g00021 | 2 | 1.059360731 |
| TpMuguga_02g00021 | 3 | 0.584474886 |
| TpMuguga_02g00022 | 1 | 0 |
| TpMuguga_02g00022 | 2 | 1.125 |
| TpMuguga_02g00022 | 3 | 4.5 |
| TpMuguga_02g00022 | 4 | 0 |
| TpMuguga_02g00022 | 5 | 0 |

| | | |
|---|---|---|
| TpMuguga_02g02250 | 1 | 0 |
| TpMuguga_02g02250 | 2 | 0 |
| TpMuguga_02g02250 | 3 | 1.125 |
| TpMuguga_02g02250 | 4 | 3 |
| TpMuguga_02g02250 | 5 | 0 |
| TpMuguga_02g02250 | 6 | 0.5625 |
| TpMuguga_02g02250 | 7 | 0 |
| TpMuguga_02g02250 | 8 | 1.125 |
| TpMuguga_02g00025 | 1 | 0 |
| TpMuguga_02g00025 | 2 | 0 |
| TpMuguga_02g00026 | 1 | 0 |
| TpMuguga_02g00026 | 2 | 1.2 |
| TpMuguga_02g02355 | 1 | 0 |
| TpMuguga_02g02595 | 1 | 0 |
| TpMuguga_02g02595 | 2 | 0 |
| TpMuguga_02g02595 | 3 | 2 |
| TpMuguga_02g02595 | 4 | 0.285714286 |
| TpMuguga_02g02595 | 5 | 0.285714286 |
| TpMuguga_02g02595 | 6 | 5.428571429 |
| TpMuguga_02g02595 | 7 | 2 |
| TpMuguga_02g00031 | 1 | 0 |
| TpMuguga_02g00033 | 1 | 0 |
| TpMuguga_02g00033 | 2 | 0.409090909 |
| TpMuguga_02g00033 | 3 | 0.409090909 |
| TpMuguga_02g00033 | 4 | 0 |
| TpMuguga_02g00033 | 5 | 0 |
| TpMuguga_02g00033 | 6 | 0 |
| TpMuguga_02g00033 | 7 | 0 |
| TpMuguga_02g00033 | 8 | 0 |
| TpMuguga_02g00035 | 1 | 2.307692308 |
| TpMuguga_02g00034 | 1 | 0 |
| TpMuguga_02g00034 | 2 | 1.536585366 |
| TpMuguga_02g00036 | 1 | 0 |
| TpMuguga_02g00036 | 2 | 0 |
| TpMuguga_02g00036 | 3 | 0 |
| TpMuguga_02g00036 | 4 | 0 |
| TpMuguga_02g00039 | 1 | 0.296296296 |
| TpMuguga_02g00040 | 1 | 0 |
| TpMuguga_02g00040 | 2 | 0 |
| TpMuguga_02g00040 | 3 | 0.454545455 |
| TpMuguga_02g00040 | 4 | 0.568181818 |
| TpMuguga_02g02485 | 1 | 0 |

| | | |
|---|---|---|
| TpMuguga_02g02485 | 2 | 2.029411765 |
| TpMuguga_02g02060 | 1 | 0 |
| TpMuguga_02g00044 | 1 | 0.6 |
| TpMuguga_02g00044 | 2 | 0 |
| TpMuguga_02g00045 | 1 | 0.531428571 |
| TpMuguga_02g00045 | 2 | 2.588571429 |
| TpMuguga_02g00046 | 1 | 0.588235294 |
| TpMuguga_02g00046 | 2 | 1.294117647 |
| TpMuguga_02g00046 | 3 | 0.647058824 |
| TpMuguga_02g00047 | 1 | 0 |
| TpMuguga_02g00048 | 1 | 0 |
| TpMuguga_02g00048 | 2 | 0 |
| TpMuguga_02g00050 | 1 | 2.875 |
| TpMuguga_02g00050 | 2 | 0.875 |
| TpMuguga_02g00050 | 3 | 0.75 |
| TpMuguga_02g02265 | 1 | 0 |
| TpMuguga_02g02265 | 2 | 2.902439024 |
| TpMuguga_02g02265 | 3 | 2.219512195 |
| TpMuguga_02g02265 | 4 | 1.024390244 |
| TpMuguga_02g02265 | 5 | 0 |
| TpMuguga_02g02265 | 6 | 0 |
| TpMuguga_02g00054 | 1 | 0 |
| TpMuguga_02g00055 | 1 | 0 |
| TpMuguga_02g00056 | 1 | 0.16875 |
| TpMuguga_02g00056 | 2 | 0.75 |
| TpMuguga_02g00057 | 1 | 0 |
| TpMuguga_02g00057 | 2 | 0 |
| TpMuguga_02g00057 | 3 | 0 |
| TpMuguga_02g00057 | 4 | 0 |
| TpMuguga_02g00057 | 5 | 0 |
| TpMuguga_02g00057 | 6 | 0 |
| TpMuguga_02g00057 | 7 | 0 |
| TpMuguga_02g00057 | 8 | 0 |
| TpMuguga_02g00057 | 9 | 0 |
| TpMuguga_02g00057 | 10 | 0 |
| TpMuguga_02g00057 | 11 | 0 |
| TpMuguga_02g00057 | 12 | 0 |
| TpMuguga_02g00059 | 1 | 0 |
| TpMuguga_02g00059 | 2 | 0 |
| TpMuguga_02g00059 | 3 | 0.303030303 |
| TpMuguga_02g00059 | 4 | 1.515151515 |
| TpMuguga_02g00060 | 1 | 0 |

| | | |
|---|---|---|
| TpMuguga_02g00060 | 2 | 0 |
| TpMuguga_02g00060 | 3 | 0 |
| TpMuguga_02g00060 | 4 | 0 |
| TpMuguga_02g00060 | 5 | 0 |
| TpMuguga_02g00061 | 1 | 1.428571429 |
| TpMuguga_02g00062 | 1 | 0 |
| TpMuguga_02g00063 | 1 | 0 |
| TpMuguga_02g00063 | 2 | 0 |
| TpMuguga_02g00066 | 1 | 0.935483871 |
| TpMuguga_02g00067 | 1 | 4 |
| TpMuguga_02g00067 | 2 | 1.25 |
| TpMuguga_02g00067 | 3 | 1.75 |
| TpMuguga_02g02645 | 1 | 0 |
| TpMuguga_02g00071 | 1 | 0 |
| TpMuguga_02g02165 | 1 | 0 |
| TpMuguga_02g02165 | 2 | 0 |
| TpMuguga_02g02165 | 3 | 0 |
| TpMuguga_02g02165 | 4 | 0 |
| TpMuguga_02g02165 | 5 | 0 |
| TpMuguga_02g02165 | 6 | 0 |
| TpMuguga_02g02165 | 7 | 0 |
| TpMuguga_02g00074 | 1 | 0 |
| TpMuguga_02g00074 | 2 | 0 |
| TpMuguga_02g00074 | 3 | 0.187319885 |
| TpMuguga_02g00074 | 4 | 0.90778098 |
| TpMuguga_02g02465 | 1 | 0 |
| TpMuguga_02g02465 | 2 | 0 |
| TpMuguga_02g02465 | 3 | 0 |
| TpMuguga_02g02465 | 4 | 2.692307692 |
| TpMuguga_02g00075 | 1 | 0 |
| TpMuguga_02g00075 | 2 | 0 |
| TpMuguga_02g00075 | 3 | 0 |
| TpMuguga_02g00075 | 4 | 0 |
| TpMuguga_02g00075 | 5 | 0 |
| TpMuguga_02g00075 | 6 | 0 |
| TpMuguga_02g00076 | 1 | 0 |
| TpMuguga_02g00076 | 2 | 0 |
| TpMuguga_02g00076 | 3 | 0 |
| TpMuguga_02g00076 | 4 | 6 |
| TpMuguga_02g00076 | 5 | 0 |
| TpMuguga_02g00078 | 1 | 0 |
| TpMuguga_02g00078 | 2 | 0 |

| | | |
|---|---|---|
| TpMuguga_02g00077 | 1 | 0 |
| TpMuguga_02g02495 | 1 | 0 |
| TpMuguga_02g00079 | 1 | 0 |
| TpMuguga_02g00079 | 2 | 0 |
| TpMuguga_02g00079 | 3 | 0 |
| TpMuguga_02g00080 | 1 | 1.558441558 |
| TpMuguga_02g00080 | 2 | 0.974025974 |
| TpMuguga_02g00081 | 1 | 2 |
| TpMuguga_02g00081 | 2 | 4 |
| TpMuguga_02g00081 | 3 | 0.5 |
| TpMuguga_02g00082 | 1 | 0 |
| TpMuguga_02g00082 | 2 | 0 |
| TpMuguga_02g00083 | 1 | 0 |
| TpMuguga_02g00084 | 1 | 0 |
| TpMuguga_02g00084 | 2 | 0 |
| TpMuguga_02g00084 | 3 | 0 |
| TpMuguga_02g00084 | 4 | 0 |
| TpMuguga_02g02180 | 1 | 0 |
| TpMuguga_02g02180 | 2 | 0 |
| TpMuguga_02g02180 | 3 | 0 |
| TpMuguga_02g00088 | 1 | 0 |
| TpMuguga_02g00088 | 2 | 0 |
| TpMuguga_02g00088 | 3 | 0 |
| TpMuguga_02g00088 | 4 | 21 |
| TpMuguga_02g00088 | 5 | 0 |
| TpMuguga_02g00089 | 1 | 0 |
| TpMuguga_02g02520 | 1 | 0 |
| TpMuguga_02g02520 | 2 | 0 |
| TpMuguga_02g00093 | 1 | 1.862068966 |
| TpMuguga_02g00093 | 2 | 0 |
| TpMuguga_02g00094 | 1 | 7.5 |
| TpMuguga_02g00094 | 2 | 0 |
| TpMuguga_02g00094 | 3 | 0 |
| TpMuguga_02g00094 | 4 | 8.333333333 |
| TpMuguga_02g00095 | 1 | 17 |
| TpMuguga_02g00095 | 2 | 0 |
| TpMuguga_02g00095 | 3 | 0 |
| TpMuguga_02g00095 | 4 | 0 |
| TpMuguga_02g00095 | 5 | 0 |
| TpMuguga_02g00095 | 6 | 0 |
| TpMuguga_02g00095 | 7 | 0 |
| TpMuguga_02g00095 | 8 | 0 |

| | | |
|---|---|---|
| TpMuguga_02g00095 | 9 | 0 |
| TpMuguga_02g00095 | 10 | 0 |
| TpMuguga_02g00095 | 11 | 0 |
| TpMuguga_02g00095 | 12 | 0 |
| TpMuguga_02g00095 | 13 | 0 |
| TpMuguga_02g00095 | 14 | 0 |
| TpMuguga_02g00095 | 15 | 0 |
| TpMuguga_02g00095 | 16 | 0 |
| TpMuguga_02g00096 | 1 | 0 |
| TpMuguga_02g00096 | 2 | 0 |
| TpMuguga_02g00096 | 3 | 0 |
| TpMuguga_02g00096 | 4 | 0 |
| TpMuguga_02g00096 | 5 | 0 |
| TpMuguga_02g00097 | 1 | 0.050307434 |
| TpMuguga_02g00097 | 2 | 0.006707658 |
| TpMuguga_02g00098 | 1 | 0 |
| TpMuguga_02g02195 | 1 | 0 |
| TpMuguga_02g00099 | 1 | 0 |
| TpMuguga_02g00100 | 1 | 0 |
| TpMuguga_02g00100 | 2 | 0 |
| TpMuguga_02g00100 | 3 | 0 |
| TpMuguga_02g00102 | 1 | 0.261874198 |
| TpMuguga_02g00102 | 2 | 0.189987163 |
| TpMuguga_02g00102 | 3 | 0.410783055 |
| TpMuguga_02g02575 | 1 | 0.32 |
| TpMuguga_02g02575 | 2 | 0 |
| TpMuguga_02g02575 | 3 | 0 |
| TpMuguga_02g02575 | 4 | 0 |
| TpMuguga_02g02575 | 5 | 1.6 |
| TpMuguga_02g02575 | 6 | 0 |
| TpMuguga_02g02575 | 7 | 0 |
| TpMuguga_02g00106 | 1 | 4 |
| TpMuguga_02g00106 | 2 | 12 |
| TpMuguga_02g00106 | 3 | 0 |
| TpMuguga_02g00107 | 1 | 2.631578947 |
| TpMuguga_02g00107 | 2 | 0 |
| TpMuguga_02g00107 | 3 | 0 |
| TpMuguga_02g00107 | 4 | 0 |
| TpMuguga_02g02005 | 1 | 0 |
| TpMuguga_02g02005 | 2 | 0 |
| TpMuguga_02g02005 | 3 | 0 |
| TpMuguga_02g02005 | 4 | 0 |

| | | |
|---|---|---|
| TpMuguga_02g00110 | 1 | 0.6 |
| TpMuguga_02g00110 | 2 | 2.4 |
| TpMuguga_02g00110 | 3 | 0 |
| TpMuguga_02g00110 | 4 | 0 |
| TpMuguga_02g00111 | 1 | 0 |
| TpMuguga_02g00111 | 2 | 0 |
| TpMuguga_02g00111 | 3 | 0 |
| TpMuguga_02g00113 | 1 | 0 |
| TpMuguga_02g00113 | 2 | 0 |
| TpMuguga_02g00112 | 1 | 0 |
| TpMuguga_02g00112 | 2 | 0 |
| TpMuguga_02g00112 | 3 | 0.082644628 |
| TpMuguga_02g00112 | 4 | 0.082644628 |
| TpMuguga_02g00112 | 5 | 0.115702479 |
| TpMuguga_02g00112 | 6 | 0 |
| TpMuguga_02g00112 | 7 | 0.26446281 |
| TpMuguga_02g00114 | 1 | 0 |
| TpMuguga_02g00114 | 2 | 0 |
| TpMuguga_02g00114 | 3 | 3.6 |
| TpMuguga_02g00116 | 1 | 1.153846154 |
| TpMuguga_02g00116 | 2 | 0 |
| TpMuguga_02g00118 | 1 | 0.756176155 |
| TpMuguga_02g00118 | 2 | 0.833512352 |
| TpMuguga_02g00118 | 3 | 1.86895811 |
| TpMuguga_02g00117 | 1 | 0 |
| TpMuguga_02g00117 | 2 | 0 |
| TpMuguga_02g00117 | 3 | 6.035087719 |
| TpMuguga_02g00119 | 1 | 0 |
| TpMuguga_02g00121 | 1 | 2.747252747 |
| TpMuguga_02g00121 | 2 | 0 |
| TpMuguga_02g00121 | 3 | 0.274725275 |
| TpMuguga_02g00121 | 4 | 0 |
| TpMuguga_02g00123 | 1 | 0 |
| TpMuguga_02g00123 | 2 | 0 |
| TpMuguga_02g00123 | 3 | 0 |
| TpMuguga_02g00123 | 4 | 0 |
| TpMuguga_02g00124 | 1 | 0 |
| TpMuguga_02g00124 | 2 | 0 |
| TpMuguga_02g00124 | 3 | 0 |
| TpMuguga_02g00124 | 4 | 0 |
| TpMuguga_02g00124 | 5 | 0 |
| TpMuguga_02g00124 | 6 | 0 |

| | | |
|---|---|---|
| TpMuguga_02g00124 | 7 | 0 |
| TpMuguga_02g00124 | 8 | 0 |
| TpMuguga_02g00124 | 9 | 0 |
| TpMuguga_02g00124 | 10 | 0 |
| TpMuguga_02g00124 | 11 | 0 |
| TpMuguga_02g00124 | 12 | 0 |
| TpMuguga_02g00124 | 13 | 0 |
| TpMuguga_02g00126 | 1 | 1.111111111 |
| TpMuguga_02g00126 | 2 | 0.317460317 |
| TpMuguga_02g00126 | 3 | 0 |
| TpMuguga_02g00126 | 4 | 0 |
| TpMuguga_02g00127 | 1 | 0 |
| TpMuguga_02g00127 | 2 | 0 |
| TpMuguga_02g00127 | 3 | 0 |
| TpMuguga_02g00127 | 4 | 0 |
| TpMuguga_02g00127 | 5 | 0 |
| TpMuguga_02g00127 | 6 | 1.074626866 |
| TpMuguga_02g00127 | 7 | 2.686567164 |
| TpMuguga_02g00127 | 8 | 0.179104478 |
| TpMuguga_02g00127 | 9 | 0.537313433 |
| TpMuguga_02g00127 | 10 | 1.074626866 |
| TpMuguga_02g00127 | 11 | 2.328358209 |
| TpMuguga_02g00128 | 1 | 0 |
| TpMuguga_02g00128 | 2 | 0 |
| TpMuguga_02g00128 | 3 | 0 |
| TpMuguga_02g00129 | 1 | 0 |
| TpMuguga_02g00129 | 2 | 0 |
| TpMuguga_02g00129 | 3 | 0 |
| TpMuguga_02g00129 | 4 | 0 |
| TpMuguga_02g00129 | 5 | 0 |
| TpMuguga_02g00129 | 6 | 0 |
| TpMuguga_02g00129 | 7 | 0 |
| TpMuguga_02g00131 | 1 | 1.25 |
| TpMuguga_02g00131 | 2 | 7.5 |
| TpMuguga_02g00131 | 3 | 18.75 |
| TpMuguga_02g00131 | 4 | 16.25 |
| TpMuguga_02g00131 | 5 | 12.5 |
| TpMuguga_02g00131 | 6 | 0 |
| TpMuguga_02g00131 | 7 | 0 |
| TpMuguga_02g00131 | 8 | 0 |
| TpMuguga_02g00131 | 9 | 0 |
| TpMuguga_02g00131 | 10 | 0 |

| | | |
|---|---|---|
| TpMuguga_02g00131 | 11 | 0 |
| TpMuguga_02g00131 | 12 | 0 |
| TpMuguga_02g00131 | 13 | 8.75 |
| TpMuguga_02g00131 | 14 | 0 |
| TpMuguga_02g00132 | 1 | 0.056461731 |
| TpMuguga_02g00132 | 2 | 0 |
| TpMuguga_02g00132 | 3 | 0 |
| TpMuguga_02g00132 | 4 | 0 |
| TpMuguga_02g00132 | 5 | 0 |
| TpMuguga_02g00132 | 6 | 0 |
| TpMuguga_02g00132 | 7 | 0.188205772 |
| TpMuguga_02g00132 | 8 | 0 |
| TpMuguga_02g00132 | 9 | 0 |
| TpMuguga_02g00132 | 10 | 0 |
| TpMuguga_02g00132 | 11 | 0.018820577 |
| TpMuguga_02g00132 | 12 | 0 |
| TpMuguga_02g00132 | 13 | 0.018820577 |
| TpMuguga_02g00132 | 14 | 0.225846926 |
| TpMuguga_02g02405 | 1 | 0.04432133 |
| TpMuguga_02g02405 | 2 | 0 |
| TpMuguga_02g02405 | 3 | 0 |
| TpMuguga_02g02410 | 1 | 0 |
| TpMuguga_02g02410 | 2 | 0 |
| TpMuguga_02g00135 | 1 | 0 |
| TpMuguga_02g00136 | 1 | 0.714285714 |
| TpMuguga_02g00136 | 2 | 0 |
| TpMuguga_02g00136 | 3 | 0 |
| TpMuguga_02g00136 | 4 | 0 |
| TpMuguga_02g00137 | 1 | 0 |
| TpMuguga_02g00137 | 2 | 0 |
| TpMuguga_02g00137 | 3 | 0 |
| TpMuguga_02g00137 | 4 | 0 |
| TpMuguga_02g00139 | 1 | 0 |
| TpMuguga_02g00139 | 2 | 0 |
| TpMuguga_02g00143 | 1 | 0 |
| TpMuguga_02g00143 | 2 | 0.41509434 |
| TpMuguga_02g00143 | 3 | 0 |
| TpMuguga_02g00143 | 4 | 0 |
| TpMuguga_02g00143 | 5 | 0 |
| TpMuguga_02g00143 | 6 | 2.905660377 |
| TpMuguga_02g00143 | 7 | 8.924528302 |
| TpMuguga_02g00143 | 8 | 6.641509434 |

| | | |
|---|---|---|
| TpMuguga_02g00143 | 9 | 9.132075472 |
| TpMuguga_02g00143 | 10 | 8.509433962 |
| TpMuguga_02g00142 | 1 | 0 |
| TpMuguga_02g00142 | 2 | 0 |
| TpMuguga_02g00142 | 3 | 0 |
| TpMuguga_02g00142 | 4 | 0 |
| TpMuguga_02g00144 | 1 | 2 |
| TpMuguga_02g00144 | 2 | 0 |
| TpMuguga_02g00144 | 3 | 0 |
| TpMuguga_02g00144 | 4 | 0 |
| TpMuguga_02g00144 | 5 | 0 |
| TpMuguga_02g00144 | 6 | 0 |
| TpMuguga_02g00144 | 7 | 12 |
| TpMuguga_02g00146 | 1 | 0 |
| TpMuguga_02g00146 | 2 | 0 |
| TpMuguga_02g00145 | 1 | 0.064935065 |
| TpMuguga_02g00145 | 2 | 0 |
| TpMuguga_02g00145 | 3 | 0 |
| TpMuguga_02g00145 | 4 | 0.111317254 |
| TpMuguga_02g00149 | 1 | 0 |
| TpMuguga_02g02435 | 1 | 0 |
| TpMuguga_02g02435 | 2 | 0 |
| TpMuguga_02g02435 | 3 | 0 |
| TpMuguga_02g02435 | 4 | 0 |
| TpMuguga_02g00152 | 1 | 0 |
| TpMuguga_02g00153 | 1 | 0 |
| TpMuguga_02g00154 | 1 | 0 |
| TpMuguga_02g00154 | 2 | 0 |
| TpMuguga_02g00154 | 3 | 0 |
| TpMuguga_02g00154 | 4 | 0 |
| TpMuguga_02g00154 | 5 | 0 |
| TpMuguga_02g00154 | 6 | 0 |
| TpMuguga_02g00154 | 7 | 0 |
| TpMuguga_02g00154 | 8 | 0 |
| TpMuguga_02g00154 | 9 | 0 |
| TpMuguga_02g00154 | 10 | 0 |
| TpMuguga_02g00154 | 11 | 0 |
| TpMuguga_02g00154 | 12 | 0 |
| TpMuguga_02g00154 | 13 | 0 |
| TpMuguga_02g00154 | 14 | 0 |
| TpMuguga_02g00154 | 15 | 0 |
| TpMuguga_02g00154 | 16 | 0 |

| | | |
|---|---|---|
| TpMuguga_02g00154 | 17 | 0 |
| TpMuguga_02g00154 | 18 | 0 |
| TpMuguga_02g00154 | 19 | 0 |
| TpMuguga_02g00154 | 20 | 0 |
| TpMuguga_02g00154 | 21 | 0 |
| TpMuguga_02g00154 | 22 | 0 |
| TpMuguga_02g00154 | 23 | 0 |
| TpMuguga_02g00154 | 24 | 0 |
| TpMuguga_02g00154 | 25 | 0 |
| TpMuguga_02g00154 | 26 | 0 |
| TpMuguga_02g00154 | 27 | 0 |
| TpMuguga_02g00154 | 28 | 0 |
| TpMuguga_02g00154 | 29 | 0 |
| TpMuguga_02g00154 | 30 | 0 |
| TpMuguga_02g00154 | 31 | 0 |
| TpMuguga_02g00154 | 32 | 0 |
| TpMuguga_02g00160 | 1 | 0 |
| TpMuguga_02g00160 | 2 | 0 |
| TpMuguga_02g00161 | 1 | 14 |
| TpMuguga_02g00161 | 2 | 0 |
| TpMuguga_02g00161 | 3 | 0 |
| TpMuguga_02g00161 | 4 | 7 |
| TpMuguga_02g00161 | 5 | 0 |
| TpMuguga_02g00161 | 6 | 10.5 |
| TpMuguga_02g02255 | 1 | 4.5 |
| TpMuguga_02g02255 | 2 | 0.176470588 |
| TpMuguga_02g00162 | 1 | 0 |
| TpMuguga_02g00162 | 2 | 0 |
| TpMuguga_02g00162 | 3 | 0 |
| TpMuguga_02g00162 | 4 | 0 |
| TpMuguga_02g00162 | 5 | 0.003668042 |
| TpMuguga_02g00162 | 6 | 0 |
| TpMuguga_02g00162 | 7 | 0 |
| TpMuguga_02g00167 | 1 | 0 |
| TpMuguga_02g00166 | 1 | 0 |
| TpMuguga_02g00166 | 2 | 0 |
| TpMuguga_02g00166 | 3 | 0 |
| TpMuguga_02g00166 | 4 | 0 |
| TpMuguga_02g00166 | 5 | 0 |
| TpMuguga_02g00166 | 6 | 0 |
| TpMuguga_02g00166 | 7 | 0 |
| TpMuguga_02g00166 | 8 | 0 |

| | | |
|---|---|---|
| TpMuguga_02g00166 | 9 | 0 |
| TpMuguga_02g00169 | 1 | 0 |
| TpMuguga_02g00170 | 1 | 0 |
| TpMuguga_02g00172 | 1 | 0 |
| TpMuguga_02g00172 | 2 | 0 |
| TpMuguga_02g00172 | 3 | 0 |
| TpMuguga_02g00172 | 4 | 0 |
| TpMuguga_02g00172 | 5 | 0 |
| TpMuguga_02g00172 | 6 | 0 |
| TpMuguga_02g00172 | 7 | 0 |
| TpMuguga_02g00172 | 8 | 0 |
| TpMuguga_02g00172 | 9 | 0 |
| TpMuguga_02g00172 | 10 | 0 |
| TpMuguga_02g00172 | 11 | 14 |
| TpMuguga_02g00172 | 12 | 98 |
| TpMuguga_02g00172 | 13 | 84 |
| TpMuguga_02g00171 | 1 | 0 |
| TpMuguga_02g00171 | 2 | 0 |
| TpMuguga_02g00171 | 3 | 0 |
| TpMuguga_02g00171 | 4 | 0 |
| TpMuguga_02g00171 | 5 | 0 |
| TpMuguga_02g00171 | 6 | 0 |
| TpMuguga_02g00171 | 7 | 0.423076923 |
| TpMuguga_02g00171 | 8 | 0 |
| TpMuguga_02g00171 | 9 | 1.269230769 |
| TpMuguga_02g00171 | 10 | 0 |
| TpMuguga_02g00174 | 1 | 0.888888889 |
| TpMuguga_02g00175 | 1 | 0 |
| TpMuguga_02g02500 | 1 | 0 |
| TpMuguga_02g00178 | 1 | 0 |
| TpMuguga_02g00178 | 2 | 0 |
| TpMuguga_02g00180 | 1 | 0.827586207 |
| TpMuguga_02g00180 | 2 | 3.310344828 |
| TpMuguga_02g00179 | 1 | 0 |
| TpMuguga_02g00181 | 1 | 0.194059406 |
| TpMuguga_02g00181 | 2 | 0 |
| TpMuguga_02g00181 | 3 | 0 |
| TpMuguga_02g00181 | 4 | 0 |
| TpMuguga_02g00181 | 5 | 0.60990099 |
| TpMuguga_02g00181 | 6 | 0.194059406 |
| TpMuguga_02g00181 | 7 | 0.388118812 |
| TpMuguga_02g00181 | 8 | 0.110891089 |

| | | |
|---|---|---|
| TpMuguga_02g00181 | 9 | 0.277227723 |
| TpMuguga_02g00181 | 10 | 0 |
| TpMuguga_02g00181 | 11 | 0 |
| TpMuguga_02g00181 | 12 | 0 |
| TpMuguga_02g00181 | 13 | 0 |
| TpMuguga_02g00182 | 1 | 0 |
| TpMuguga_02g00182 | 2 | 0 |
| TpMuguga_02g00183 | 1 | 0 |
| TpMuguga_02g00183 | 2 | 0 |
| TpMuguga_02g00183 | 3 | 0 |
| TpMuguga_02g00183 | 4 | 0 |
| TpMuguga_02g00183 | 5 | 0 |
| TpMuguga_02g00183 | 6 | 0 |
| TpMuguga_02g00183 | 7 | 0 |
| TpMuguga_02g00183 | 8 | 0 |
| TpMuguga_02g00183 | 9 | 0.344827586 |
| TpMuguga_02g00184 | 1 | 0 |
| TpMuguga_02g00184 | 2 | 60 |
| TpMuguga_02g00184 | 3 | 0 |
| TpMuguga_02g00184 | 4 | 0 |
| TpMuguga_02g00184 | 5 | 40 |
| TpMuguga_02g00184 | 6 | 10 |
| TpMuguga_02g00184 | 7 | 0 |
| TpMuguga_02g00184 | 8 | 0 |
| TpMuguga_02g00184 | 9 | 0 |
| TpMuguga_02g02675 | 1 | 0 |
| TpMuguga_02g02675 | 2 | 0 |
| TpMuguga_02g02675 | 3 | 0 |
| TpMuguga_02g02675 | 4 | 0 |
| TpMuguga_02g02675 | 5 | 0 |
| TpMuguga_02g02675 | 6 | 0 |
| TpMuguga_02g02675 | 7 | 0 |
| TpMuguga_02g02530 | 1 | 0 |
| TpMuguga_02g02530 | 2 | 1.125 |
| TpMuguga_02g02530 | 3 | 0 |
| TpMuguga_02g02530 | 4 | 0 |
| TpMuguga_02g02530 | 5 | 0 |
| TpMuguga_02g00188 | 1 | 0.344262295 |
| TpMuguga_02g00188 | 2 | 0 |
| TpMuguga_02g00188 | 3 | 0 |
| TpMuguga_02g00188 | 4 | 0 |
| TpMuguga_02g00188 | 5 | 0 |

| | | |
|---|---|---|
| TpMuguga_02g00188 | 6 | 5.795081967 |
| TpMuguga_02g00189 | 1 | 1.595505618 |
| TpMuguga_02g00190 | 1 | 1.285714286 |
| TpMuguga_02g00190 | 2 | 0 |
| TpMuguga_02g00190 | 3 | 0.857142857 |
| TpMuguga_02g00190 | 4 | 0 |
| TpMuguga_02g00190 | 5 | 0 |
| TpMuguga_02g00191 | 1 | 0 |
| TpMuguga_02g00191 | 2 | 0 |
| TpMuguga_02g00191 | 3 | 0 |
| TpMuguga_02g02585 | 1 | 13.33333333 |
| TpMuguga_02g02585 | 2 | 0 |
| TpMuguga_02g02585 | 3 | 0 |
| TpMuguga_02g00194 | 1 | 0 |
| TpMuguga_02g00194 | 2 | 0 |
| TpMuguga_02g00194 | 3 | 0 |
| TpMuguga_02g00194 | 4 | 0 |
| TpMuguga_02g00194 | 5 | 0 |
| TpMuguga_02g00196 | 1 | 0 |
| TpMuguga_02g00196 | 2 | 0 |
| TpMuguga_02g00196 | 3 | 24 |
| TpMuguga_02g00966 | 1 | 0 |
| TpMuguga_02g00966 | 2 | 5 |
| TpMuguga_02g00966 | 3 | 0 |
| TpMuguga_02g00966 | 4 | 40 |
| TpMuguga_02g00198 | 1 | 0 |
| TpMuguga_02g00198 | 2 | 0 |
| TpMuguga_02g00198 | 3 | 0 |
| TpMuguga_02g00198 | 4 | 0 |
| TpMuguga_02g00198 | 5 | 0 |
| TpMuguga_02g00198 | 6 | 0 |
| TpMuguga_02g00199 | 1 | 0 |
| TpMuguga_02g00199 | 2 | 0 |
| TpMuguga_02g00199 | 3 | 0 |
| TpMuguga_02g00200 | 1 | 0 |
| TpMuguga_02g00200 | 2 | 0 |
| TpMuguga_02g00201 | 1 | 0 |
| TpMuguga_02g00201 | 2 | 0 |
| TpMuguga_02g00201 | 3 | 0 |
| TpMuguga_02g00201 | 4 | 0 |
| TpMuguga_02g00201 | 5 | 0 |
| TpMuguga_02g02125 | 1 | 0 |

| | | |
|---|---|---|
| TpMuguga_02g02125 | 2 | 0 |
| TpMuguga_02g02125 | 3 | 0 |
| TpMuguga_02g02125 | 4 | 0 |
| TpMuguga_02g02125 | 5 | 0.055147059 |
| TpMuguga_02g02345 | 1 | 1.63507109 |
| TpMuguga_02g02345 | 2 | 11.31753555 |
| TpMuguga_02g02345 | 3 | 2.687203791 |
| TpMuguga_02g02345 | 4 | 0.099526066 |
| TpMuguga_02g02345 | 5 | 0.312796209 |
| TpMuguga_02g00206 | 1 | 0.005007153 |
| TpMuguga_02g00205 | 1 | 0 |
| TpMuguga_02g00205 | 2 | 0.242424242 |
| TpMuguga_02g00207 | 1 | 2.117647059 |
| TpMuguga_02g00207 | 2 | 0 |
| TpMuguga_02g00207 | 3 | 0 |
| TpMuguga_02g00207 | 4 | 0 |
| TpMuguga_02g00207 | 5 | 4.235294118 |
| TpMuguga_02g00207 | 6 | 5.823529412 |
| TpMuguga_02g00207 | 7 | 0.529411765 |
| TpMuguga_02g00207 | 8 | 8.470588235 |
| TpMuguga_02g02225 | 1 | 4.666666667 |
| TpMuguga_02g02225 | 2 | 12.66666667 |
| TpMuguga_02g02225 | 3 | 4.666666667 |
| TpMuguga_02g00209 | 1 | 0.171428571 |
| TpMuguga_02g00209 | 2 | 0.942857143 |
| TpMuguga_02g00209 | 3 | 0.771428571 |
| TpMuguga_02g00209 | 4 | 0.171428571 |
| TpMuguga_02g00209 | 5 | 0.857142857 |
| TpMuguga_02g00217 | 1 | 0.732394366 |
| TpMuguga_02g00218 | 1 | 0.068965517 |
| TpMuguga_02g00218 | 2 | 0.068965517 |
| TpMuguga_02g00218 | 3 | 0.689655172 |
| TpMuguga_02g00219 | 1 | 0.279069767 |
| TpMuguga_02g00220 | 1 | 4 |
| TpMuguga_02g00223 | 1 | 1.125 |
| TpMuguga_02g00223 | 2 | 1.125 |
| TpMuguga_02g00223 | 3 | 4.78125 |
| TpMuguga_02g00223 | 4 | 0 |
| TpMuguga_02g00223 | 5 | 5.625 |
| TpMuguga_02g00223 | 6 | 0.5625 |
| TpMuguga_02g00223 | 7 | 3.9375 |
| TpMuguga_02g00223 | 8 | 6.75 |

| | | |
|---|---|---|
| TpMuguga_02g00222 | 1 | 0 |
| TpMuguga_02g00222 | 2 | 0 |
| TpMuguga_02g00225 | 1 | 0 |
| TpMuguga_02g00225 | 2 | 0 |
| TpMuguga_02g00225 | 3 | 0 |
| TpMuguga_02g00225 | 4 | 0.3125 |
| TpMuguga_02g02000 | 1 | 0 |
| TpMuguga_02g02000 | 2 | 0 |
| TpMuguga_02g02000 | 3 | 0 |
| TpMuguga_02g02000 | 4 | 0 |
| TpMuguga_02g02000 | 5 | 0 |
| TpMuguga_02g02000 | 6 | 3.428571429 |
| TpMuguga_02g02000 | 7 | 0 |
| TpMuguga_02g00229 | 1 | 0 |
| TpMuguga_02g00230 | 1 | 0 |
| TpMuguga_02g00230 | 2 | 0 |
| TpMuguga_02g00230 | 3 | 0 |
| TpMuguga_02g00230 | 4 | 0 |
| TpMuguga_02g02515 | 1 | 0 |
| TpMuguga_02g02515 | 2 | 0 |
| TpMuguga_02g02515 | 3 | 0 |
| TpMuguga_02g02515 | 4 | 0 |
| TpMuguga_02g00231 | 1 | 0 |
| TpMuguga_02g00231 | 2 | 0 |
| TpMuguga_02g00231 | 3 | 0 |
| TpMuguga_02g00231 | 4 | 0 |
| TpMuguga_02g00231 | 5 | 2.142857143 |
| TpMuguga_02g00233 | 1 | 0 |
| TpMuguga_02g00233 | 2 | 0 |
| TpMuguga_02g00233 | 3 | 0 |
| TpMuguga_02g00233 | 4 | 0 |
| TpMuguga_02g00233 | 5 | 0 |
| TpMuguga_02g02300 | 1 | 0.239316239 |
| TpMuguga_02g00235 | 1 | 0 |
| TpMuguga_02g00236 | 1 | 0 |
| TpMuguga_02g00236 | 2 | 0 |
| TpMuguga_02g00236 | 3 | 0 |
| TpMuguga_02g00236 | 4 | 0 |
| TpMuguga_02g00236 | 5 | 0 |
| TpMuguga_02g00236 | 6 | 2.8 |
| TpMuguga_02g00237 | 1 | 0.185328185 |
| TpMuguga_02g00237 | 2 | 0 |

| | | |
|---|---|---|
| TpMuguga_02g00237 | 3 | 0 |
| TpMuguga_02g00238 | 1 | 0 |
| TpMuguga_02g00238 | 2 | 0 |
| TpMuguga_02g00240 | 1 | 24 |
| TpMuguga_02g00240 | 2 | 18 |
| TpMuguga_02g00240 | 3 | 0 |
| TpMuguga_02g00240 | 4 | 0 |
| TpMuguga_02g00240 | 5 | 0 |
| TpMuguga_02g02665 | 1 | 0 |
| TpMuguga_02g02665 | 2 | 0.142857143 |
| TpMuguga_02g02665 | 3 | 0.571428571 |
| TpMuguga_02g00241 | 1 | 0 |
| TpMuguga_02g00241 | 2 | 0 |
| TpMuguga_02g00241 | 3 | 0 |
| TpMuguga_02g00242 | 1 | 0 |
| TpMuguga_02g00242 | 2 | 0 |
| TpMuguga_02g00242 | 3 | 0 |
| TpMuguga_02g00242 | 4 | 0 |
| TpMuguga_02g00242 | 5 | 3 |
| TpMuguga_02g00242 | 6 | 0 |
| TpMuguga_02g00242 | 7 | 0 |
| TpMuguga_02g00242 | 8 | 0 |
| TpMuguga_02g00243 | 1 | 0 |
| TpMuguga_02g00243 | 2 | 0 |
| TpMuguga_02g00243 | 3 | 0 |
| TpMuguga_02g00243 | 4 | 0 |
| TpMuguga_02g00244 | 1 | 0 |
| TpMuguga_02g00244 | 2 | 0.153846154 |
| TpMuguga_02g00244 | 3 | 0.538461538 |
| TpMuguga_02g00245 | 1 | 0 |
| TpMuguga_02g00245 | 2 | 0 |
| TpMuguga_02g00247 | 1 | 0 |
| TpMuguga_02g00247 | 2 | 0 |
| TpMuguga_02g00247 | 3 | 0 |
| TpMuguga_02g00247 | 4 | 0 |
| TpMuguga_02g00247 | 5 | 0 |
| TpMuguga_02g00248 | 1 | 0.818181818 |
| TpMuguga_02g00248 | 2 | 0 |
| TpMuguga_02g00248 | 3 | 0 |
| TpMuguga_02g00248 | 4 | 0 |
| TpMuguga_02g00248 | 5 | 0 |
| TpMuguga_02g00250 | 1 | 0.4 |

| | | |
|---|---|---|
| TpMuguga_02g00251 | 1 | 0 |
| TpMuguga_02g00251 | 2 | 0 |
| TpMuguga_02g00251 | 3 | 0 |
| TpMuguga_02g02400 | 1 | 0 |
| TpMuguga_02g02400 | 2 | 0 |
| TpMuguga_02g02400 | 3 | 0 |
| TpMuguga_02g02400 | 4 | 0 |
| TpMuguga_02g00252 | 1 | 0 |
| TpMuguga_02g00252 | 2 | 0.333333333 |
| TpMuguga_02g00252 | 3 | 0 |
| TpMuguga_02g00254 | 1 | 1.875 |
| TpMuguga_02g00254 | 2 | 0.25 |
| TpMuguga_02g00253 | 1 | 0.310679612 |
| TpMuguga_02g00253 | 2 | 0 |
| TpMuguga_02g00253 | 3 | 1.126213592 |
| TpMuguga_02g00255 | 1 | 0 |
| TpMuguga_02g00255 | 2 | 0 |
| TpMuguga_02g00256 | 1 | 0 |
| TpMuguga_02g00256 | 2 | 0 |
| TpMuguga_02g00256 | 3 | 0 |
| TpMuguga_02g00256 | 4 | 0 |
| TpMuguga_02g00256 | 5 | 0.416918429 |
| TpMuguga_02g02035 | 1 | 0 |
| TpMuguga_02g02035 | 2 | 0 |
| TpMuguga_02g02035 | 3 | 0 |
| TpMuguga_02g02035 | 4 | 0 |
| TpMuguga_02g02035 | 5 | 0 |
| TpMuguga_02g02035 | 6 | 0 |
| TpMuguga_02g02035 | 7 | 0 |
| TpMuguga_02g02035 | 8 | 0 |
| TpMuguga_02g02035 | 9 | 0 |
| TpMuguga_02g02035 | 10 | 0 |
| TpMuguga_02g02035 | 11 | 0 |
| TpMuguga_02g02035 | 12 | 0 |
| TpMuguga_02g02035 | 13 | 0 |
| TpMuguga_02g02035 | 14 | 0 |
| TpMuguga_02g02035 | 15 | 0 |
| TpMuguga_02g02035 | 16 | 0 |
| TpMuguga_02g02035 | 17 | 0 |
| TpMuguga_02g00262 | 1 | 1 |
| TpMuguga_02g00263 | 1 | 0 |
| TpMuguga_02g00263 | 2 | 0 |

| | | |
|---|---|---|
| TpMuguga_02g00265 | 1 | 0 |
| TpMuguga_02g00265 | 2 | 3 |
| TpMuguga_02g02080 | 1 | 0 |
| TpMuguga_02g02080 | 2 | 0 |
| TpMuguga_02g02080 | 3 | 0 |
| TpMuguga_02g02080 | 4 | 0 |
| TpMuguga_02g00269 | 1 | 3.6 |
| TpMuguga_02g00269 | 2 | 0 |
| TpMuguga_02g00269 | 3 | 2.8 |
| TpMuguga_02g00269 | 4 | 0 |
| TpMuguga_02g00269 | 5 | 0 |
| TpMuguga_02g02200 | 1 | 0 |
| TpMuguga_02g02200 | 2 | 0 |
| TpMuguga_02g02200 | 3 | 0 |
| TpMuguga_02g02200 | 4 | 9 |
| TpMuguga_02g02200 | 5 | 2 |
| TpMuguga_02g02200 | 6 | 0 |
| TpMuguga_02g02200 | 7 | 4 |
| TpMuguga_02g02200 | 8 | 1 |
| TpMuguga_02g02200 | 9 | 0 |
| TpMuguga_02g02200 | 10 | 0 |
| TpMuguga_02g00272 | 1 | 0 |
| TpMuguga_02g00272 | 2 | 0 |
| TpMuguga_02g02395 | 1 | 0 |
| TpMuguga_02g02395 | 2 | 5 |
| TpMuguga_02g02395 | 3 | 5 |
| TpMuguga_02g02395 | 4 | 15 |
| TpMuguga_02g02065 | 1 | 2.842105263 |
| TpMuguga_02g02295 | 1 | 3.658536585 |
| TpMuguga_02g02295 | 2 | 5.853658537 |
| TpMuguga_02g02295 | 3 | 2.926829268 |
| TpMuguga_02g02295 | 4 | 2.317073171 |
| TpMuguga_02g02295 | 5 | 2.682926829 |
| TpMuguga_02g02295 | 6 | 5.243902439 |
| TpMuguga_02g02295 | 7 | 0.243902439 |
| TpMuguga_02g02295 | 8 | 1.219512195 |
| TpMuguga_02g02295 | 9 | 3.536585366 |
| TpMuguga_02g02285 | 1 | 2.142857143 |
| TpMuguga_02g02285 | 2 | 7.857142857 |
| TpMuguga_02g02285 | 3 | 1.428571429 |
| TpMuguga_02g02285 | 4 | 0 |
| TpMuguga_02g02535 | 1 | 1.875 |

| | | |
|---|---|---|
| TpMuguga_02g02535 | 2 | 0.833333333 |
| TpMuguga_02g02535 | 3 | 0 |
| TpMuguga_02g02535 | 4 | 0 |
| TpMuguga_02g00280 | 1 | 6.230769231 |
| TpMuguga_02g00280 | 2 | 1.153846154 |
| TpMuguga_02g00280 | 3 | 0 |
| TpMuguga_02g00280 | 4 | 0 |
| TpMuguga_02g00280 | 5 | 0 |
| TpMuguga_02g00281 | 1 | 0 |
| TpMuguga_02g00281 | 2 | 0 |
| TpMuguga_02g00281 | 3 | 0 |
| TpMuguga_02g00281 | 4 | 0 |
| TpMuguga_02g00281 | 5 | 0 |
| TpMuguga_02g00282 | 1 | 0 |
| TpMuguga_02g00282 | 2 | 0 |
| TpMuguga_02g00282 | 3 | 0.028169014 |
| TpMuguga_02g00282 | 4 | 0.070422535 |
| TpMuguga_02g00282 | 5 | 0 |
| TpMuguga_02g02610 | 1 | 0 |
| TpMuguga_02g02610 | 2 | 0 |
| TpMuguga_02g00283 | 1 | 0 |
| TpMuguga_02g00283 | 2 | 0.071090047 |
| TpMuguga_02g02445 | 1 | 0.596385542 |
| TpMuguga_02g02445 | 2 | 1.084337349 |
| TpMuguga_02g00285 | 1 | 0 |
| TpMuguga_02g00285 | 2 | 0.260869565 |
| TpMuguga_02g00285 | 3 | 2.217391304 |
| TpMuguga_02g00285 | 4 | 0 |
| TpMuguga_02g00285 | 5 | 0 |
| TpMuguga_02g00287 | 1 | 2.113821138 |
| TpMuguga_02g00287 | 2 | 0 |
| TpMuguga_02g00287 | 3 | 0 |
| TpMuguga_02g00287 | 4 | 0.243902439 |
| TpMuguga_02g00288 | 1 | 0 |
| TpMuguga_02g00288 | 2 | 0 |
| TpMuguga_02g00289 | 1 | 0 |
| TpMuguga_02g00289 | 2 | 0 |
| TpMuguga_02g00289 | 3 | 0 |
| TpMuguga_02g00289 | 4 | 0 |
| TpMuguga_02g00289 | 5 | 0 |
| TpMuguga_02g00289 | 6 | 0 |
| TpMuguga_02g00289 | 7 | 0 |

| | | |
|---|---|---|
| TpMuguga_02g00290 | 1 | 0 |
| TpMuguga_02g00290 | 2 | 0 |
| TpMuguga_02g00291 | 1 | 0 |
| TpMuguga_02g00291 | 2 | 0 |
| TpMuguga_02g00291 | 3 | 0 |
| TpMuguga_02g00291 | 4 | 0 |
| TpMuguga_02g00292 | 1 | 0 |
| TpMuguga_02g00292 | 2 | 0 |
| TpMuguga_02g00292 | 3 | 0 |
| TpMuguga_02g00292 | 4 | 0 |
| TpMuguga_02g00292 | 5 | 0 |
| TpMuguga_02g00292 | 6 | 0 |
| TpMuguga_02g00292 | 7 | 0 |
| TpMuguga_02g00292 | 8 | 0 |
| TpMuguga_02g00294 | 1 | 0.296296296 |
| TpMuguga_02g00294 | 2 | 0.444444444 |
| TpMuguga_02g00294 | 3 | 5.481481481 |
| TpMuguga_02g02130 | 1 | 0 |
| TpMuguga_02g02130 | 2 | 0 |
| TpMuguga_02g02130 | 3 | 0 |
| TpMuguga_02g02130 | 4 | 0 |
| TpMuguga_02g02130 | 5 | 0 |
| TpMuguga_02g02130 | 6 | 0 |
| TpMuguga_02g02130 | 7 | 0.028933092 |
| TpMuguga_02g02130 | 8 | 0 |
| TpMuguga_02g02130 | 9 | 0 |
| TpMuguga_02g02130 | 10 | 0 |
| TpMuguga_02g02130 | 11 | 0 |
| TpMuguga_02g00297 | 1 | 0.52173913 |
| TpMuguga_02g00297 | 2 | 0 |
| TpMuguga_02g00298 | 1 | 1.252941176 |
| TpMuguga_02g02580 | 1 | 0 |
| TpMuguga_02g02580 | 2 | 0 |
| TpMuguga_02g02580 | 3 | 0 |
| TpMuguga_02g02580 | 4 | 0 |
| TpMuguga_02g02580 | 5 | 0 |
| TpMuguga_02g02580 | 6 | 0 |
| TpMuguga_02g02580 | 7 | 0 |
| TpMuguga_02g02580 | 8 | 0 |
| TpMuguga_02g02580 | 9 | 0.002658378 |
| TpMuguga_02g02580 | 10 | 0 |
| TpMuguga_02g02580 | 11 | 0 |

| | | |
|---|---|---|
| TpMuguga_02g02580 | 12 | 0 |
| TpMuguga_02g00300 | 1 | 0 |
| TpMuguga_02g00302 | 1 | 0 |
| TpMuguga_02g00302 | 2 | 0.425531915 |
| TpMuguga_02g00302 | 3 | 0 |
| TpMuguga_02g00302 | 4 | 0 |
| TpMuguga_02g00301 | 1 | 3.138888889 |
| TpMuguga_02g00301 | 2 | 0.472222222 |
| TpMuguga_02g00304 | 1 | 0 |
| TpMuguga_02g00304 | 2 | 0 |
| TpMuguga_02g00305 | 1 | 3.4 |
| TpMuguga_02g00307 | 1 | 0 |
| TpMuguga_02g00308 | 1 | 0 |
| TpMuguga_02g00308 | 2 | 0 |
| TpMuguga_02g00308 | 3 | 0 |
| TpMuguga_02g00308 | 4 | 0.857142857 |
| TpMuguga_02g00310 | 1 | 0 |
| TpMuguga_02g00310 | 2 | 0 |
| TpMuguga_02g00310 | 3 | 0 |
| TpMuguga_02g00310 | 4 | 0 |
| TpMuguga_02g00310 | 5 | 0 |
| TpMuguga_02g00310 | 6 | 0 |
| TpMuguga_02g00310 | 7 | 0 |
| TpMuguga_02g00311 | 1 | 0 |
| TpMuguga_02g00311 | 2 | 0 |
| TpMuguga_02g00311 | 3 | 0 |
| TpMuguga_02g00311 | 4 | 0 |
| TpMuguga_02g00311 | 5 | 0 |
| TpMuguga_02g00312 | 1 | 0 |
| TpMuguga_02g00312 | 2 | 0 |
| TpMuguga_02g00312 | 3 | 0 |
| TpMuguga_02g00312 | 4 | 0 |
| TpMuguga_02g00312 | 5 | 0 |
| TpMuguga_02g00312 | 6 | 0 |
| TpMuguga_02g00312 | 7 | 0 |
| TpMuguga_02g00312 | 8 | 0 |
| TpMuguga_02g00312 | 9 | 0 |
| TpMuguga_02g00315 | 1 | 0 |
| TpMuguga_02g00315 | 2 | 0 |
| TpMuguga_02g00315 | 3 | 0 |
| TpMuguga_02g00315 | 4 | 0 |
| TpMuguga_02g00315 | 5 | 0 |

| | | |
|---|---|---|
| TpMuguga_02g00315 | 6 | 0 |
| TpMuguga_02g00315 | 7 | 0 |
| TpMuguga_02g00315 | 8 | 6.387096774 |
| TpMuguga_02g02670 | 1 | 0 |
| TpMuguga_02g02670 | 2 | 0 |
| TpMuguga_02g02670 | 3 | 24 |
| TpMuguga_02g00316 | 1 | 0.428571429 |
| TpMuguga_02g00316 | 2 | 1.714285714 |
| TpMuguga_02g00316 | 3 | 0.285714286 |
| TpMuguga_02g00316 | 4 | 0.428571429 |
| TpMuguga_02g00316 | 5 | 0 |
| TpMuguga_02g00317 | 1 | 0 |
| TpMuguga_02g00319 | 1 | 0 |
| TpMuguga_02g00319 | 2 | 0 |
| TpMuguga_02g00319 | 3 | 0 |
| TpMuguga_02g00319 | 4 | 0.933333333 |
| TpMuguga_02g00319 | 5 | 1.4 |
| TpMuguga_02g00319 | 6 | 5.6 |
| TpMuguga_02g00320 | 1 | 0 |
| TpMuguga_02g00320 | 2 | 0 |
| TpMuguga_02g00320 | 3 | 0 |
| TpMuguga_02g00321 | 1 | 1.363636364 |
| TpMuguga_02g02375 | 1 | 0 |
| TpMuguga_02g02375 | 2 | 0 |
| TpMuguga_02g00322 | 1 | 0.068027211 |
| TpMuguga_02g00322 | 2 | 0.05952381 |
| TpMuguga_02g00322 | 3 | 0.068027211 |
| TpMuguga_02g00322 | 4 | 0.204081633 |
| TpMuguga_02g00323 | 1 | 3 |
| TpMuguga_02g00323 | 2 | 4 |
| TpMuguga_02g02045 | 1 | 1.902985075 |
| TpMuguga_02g02045 | 2 | 0.298507463 |
| TpMuguga_02g02045 | 3 | 0.52238806 |
| TpMuguga_02g02045 | 4 | 0.410447761 |
| TpMuguga_02g00324 | 1 | 0 |
| TpMuguga_02g00324 | 2 | 0 |
| TpMuguga_02g00324 | 3 | 0 |
| TpMuguga_02g02040 | 1 | 10.8 |
| TpMuguga_02g02040 | 2 | 5.4 |
| TpMuguga_02g02040 | 3 | 0 |
| TpMuguga_02g02040 | 4 | 0 |
| TpMuguga_02g02040 | 5 | 1.8 |

| | | |
|---|---|---|
| TpMuguga_02g02040 | 6 | 5.4 |
| TpMuguga_02g02040 | 7 | 12.6 |
| TpMuguga_02g02040 | 8 | 0 |
| TpMuguga_02g00326 | 1 | 0 |
| TpMuguga_02g00326 | 2 | 0 |
| TpMuguga_02g00326 | 3 | 0 |
| TpMuguga_02g00326 | 4 | 0 |
| TpMuguga_02g00326 | 5 | 0 |
| TpMuguga_02g00326 | 6 | 0 |
| TpMuguga_02g00326 | 7 | 0 |
| TpMuguga_02g00326 | 8 | 3.428571429 |
| TpMuguga_02g00326 | 9 | 0 |
| TpMuguga_02g00326 | 10 | 0 |
| TpMuguga_02g00326 | 11 | 0 |
| TpMuguga_02g00328 | 1 | 0.257142857 |
| TpMuguga_02g00329 | 1 | 0 |
| TpMuguga_02g00331 | 1 | 0 |
| TpMuguga_02g00331 | 2 | 0 |
| TpMuguga_02g00333 | 1 | 0 |
| TpMuguga_02g00333 | 2 | 0 |
| TpMuguga_02g00333 | 3 | 0 |
| TpMuguga_02g00334 | 1 | 0 |
| TpMuguga_02g00334 | 2 | 8.461538462 |
| TpMuguga_02g00334 | 3 | 0 |
| TpMuguga_02g00334 | 4 | 2.307692308 |
| TpMuguga_02g00334 | 5 | 0.769230769 |
| TpMuguga_02g00334 | 6 | 0 |
| TpMuguga_02g00334 | 7 | 0 |
| TpMuguga_02g00334 | 8 | 1.538461538 |
| TpMuguga_02g00334 | 9 | 1.538461538 |
| TpMuguga_02g00335 | 1 | 0 |
| TpMuguga_02g00335 | 2 | 0 |
| TpMuguga_02g00335 | 3 | 0 |
| TpMuguga_02g00335 | 4 | 0 |
| TpMuguga_02g00335 | 5 | 3 |
| TpMuguga_02g00336 | 1 | 0 |
| TpMuguga_02g00336 | 2 | 0 |
| TpMuguga_02g00336 | 3 | 0 |
| TpMuguga_02g00336 | 4 | 0.365535248 |
| TpMuguga_02g02550 | 1 | 0 |
| TpMuguga_02g02550 | 2 | 0 |
| TpMuguga_02g02550 | 3 | 0.088105727 |

| | | |
|---|---|---|
| TpMuguga_02g00337 | 1 | 0.255102041 |
| TpMuguga_02g00337 | 2 | 0.193877551 |
| TpMuguga_02g00338 | 1 | 0 |
| TpMuguga_02g00338 | 2 | 0 |
| TpMuguga_02g00339 | 1 | 0 |
| TpMuguga_02g00339 | 2 | 0 |
| TpMuguga_02g00339 | 3 | 0 |
| TpMuguga_02g00339 | 4 | 0 |
| TpMuguga_02g00341 | 1 | 0 |
| TpMuguga_02g00343 | 1 | 0 |
| TpMuguga_02g00343 | 2 | 0 |
| TpMuguga_02g00343 | 3 | 0 |
| TpMuguga_02g00342 | 1 | 0 |
| TpMuguga_02g00342 | 2 | 0 |
| TpMuguga_02g00342 | 3 | 0 |
| TpMuguga_02g00342 | 4 | 0 |
| TpMuguga_02g00342 | 5 | 0 |
| TpMuguga_02g00342 | 6 | 0 |
| TpMuguga_02g00342 | 7 | 0.139534884 |
| TpMuguga_02g02270 | 1 | 0 |
| TpMuguga_02g02270 | 2 | 0 |
| TpMuguga_02g02270 | 3 | 0 |
| TpMuguga_02g00345 | 1 | 0.190713101 |
| TpMuguga_02g00345 | 2 | 0 |
| TpMuguga_02g00345 | 3 | 0 |
| TpMuguga_02g00345 | 4 | 0 |
| TpMuguga_02g00346 | 1 | 0 |
| TpMuguga_02g00346 | 2 | 0 |
| TpMuguga_02g00347 | 1 | 0 |
| TpMuguga_02g00347 | 2 | 0 |
| TpMuguga_02g00348 | 1 | 0.220588235 |
| TpMuguga_02g00348 | 2 | 0 |
| TpMuguga_02g00349 | 1 | 0 |
| TpMuguga_02g00349 | 2 | 0 |
| TpMuguga_02g00349 | 3 | 0 |
| TpMuguga_02g00350 | 1 | 0 |
| TpMuguga_02g00350 | 2 | 0 |
| TpMuguga_02g00351 | 1 | 0 |
| TpMuguga_02g00351 | 2 | 0 |
| TpMuguga_02g00352 | 1 | 12 |
| TpMuguga_02g00352 | 2 | 0 |
| TpMuguga_02g00352 | 3 | 0 |

| | | |
|---|---|---|
| TpMuguga_02g00352 | 4 | 0 |
| TpMuguga_02g00352 | 5 | 0 |
| TpMuguga_02g00353 | 1 | 0 |
| TpMuguga_02g00353 | 2 | 0 |
| TpMuguga_02g00353 | 3 | 0 |
| TpMuguga_02g00354 | 1 | 0 |
| TpMuguga_02g00354 | 2 | 0 |
| TpMuguga_02g00354 | 3 | 0 |
| TpMuguga_02g00354 | 4 | 0 |
| TpMuguga_02g00354 | 5 | 0 |
| TpMuguga_02g00354 | 6 | 0 |
| TpMuguga_02g00356 | 1 | 0 |
| TpMuguga_02g00356 | 2 | 0 |
| TpMuguga_02g00355 | 1 | 0 |
| TpMuguga_02g00355 | 2 | 0 |
| TpMuguga_02g00355 | 3 | 0 |
| TpMuguga_02g00355 | 4 | 0 |
| TpMuguga_02g00355 | 5 | 0 |
| TpMuguga_02g00355 | 6 | 0 |
| TpMuguga_02g00355 | 7 | 0 |
| TpMuguga_02g00355 | 8 | 0 |
| TpMuguga_02g00355 | 9 | 0 |
| TpMuguga_02g00355 | 10 | 0 |
| TpMuguga_02g00358 | 1 | 1 |
| TpMuguga_02g00358 | 2 | 0 |
| TpMuguga_02g00358 | 3 | 0 |
| TpMuguga_02g00358 | 4 | 0 |
| TpMuguga_02g00358 | 5 | 0 |
| TpMuguga_02g00358 | 6 | 0 |
| TpMuguga_02g00358 | 7 | 0 |
| TpMuguga_02g00358 | 8 | 0 |
| TpMuguga_02g00358 | 9 | 0 |
| TpMuguga_02g00358 | 10 | 0 |
| TpMuguga_02g00358 | 11 | 0 |
| TpMuguga_02g00358 | 12 | 0 |
| TpMuguga_02g00357 | 1 | 0 |
| TpMuguga_02g00357 | 2 | 0 |
| TpMuguga_02g00357 | 3 | 1 |
| TpMuguga_02g00359 | 1 | 0 |
| TpMuguga_02g00359 | 2 | 0 |
| TpMuguga_02g00360 | 1 | 0 |
| TpMuguga_02g00360 | 2 | 0 |

| | | |
|---|---|---|
| TpMuguga_02g00362 | 1 | 0 |
| TpMuguga_02g00363 | 1 | 0 |
| TpMuguga_02g00363 | 2 | 0.760869565 |
| TpMuguga_02g00363 | 3 | 0.152173913 |
| TpMuguga_02g00363 | 4 | 0 |
| TpMuguga_02g00363 | 5 | 0 |
| TpMuguga_02g00363 | 6 | 3.5 |
| TpMuguga_02g00364 | 1 | 0 |
| TpMuguga_02g00364 | 2 | 0 |
| TpMuguga_02g00364 | 3 | 0 |
| TpMuguga_02g00364 | 4 | 0 |
| TpMuguga_02g00365 | 1 | 0 |
| TpMuguga_02g00365 | 2 | 0 |
| TpMuguga_02g02160 | 1 | 0 |
| TpMuguga_02g02160 | 2 | 0 |
| TpMuguga_02g00366 | 1 | 0.306569343 |
| TpMuguga_02g00366 | 2 | 0.131386861 |
| TpMuguga_02g02570 | 1 | 0.413943355 |
| TpMuguga_02g02570 | 2 | 0.272331155 |
| TpMuguga_02g02570 | 3 | 0.231481481 |
| TpMuguga_02g02570 | 4 | 0.639978214 |
| TpMuguga_02g00369 | 1 | 0 |
| TpMuguga_02g00369 | 2 | 0 |
| TpMuguga_02g00370 | 1 | 0 |
| TpMuguga_02g00370 | 2 | 0 |
| TpMuguga_02g00370 | 3 | 0 |
| TpMuguga_02g00372 | 1 | 2 |
| TpMuguga_02g00373 | 1 | 1.75 |
| TpMuguga_02g00373 | 2 | 0 |
| TpMuguga_02g00373 | 3 | 0.35 |
| TpMuguga_02g00373 | 4 | 0.116666667 |
| TpMuguga_02g00373 | 5 | 0 |
| TpMuguga_02g00373 | 6 | 0 |
| TpMuguga_02g00374 | 1 | 3.736596737 |
| TpMuguga_02g00374 | 2 | 0.538461538 |
| TpMuguga_02g00374 | 3 | 0.522144522 |
| TpMuguga_02g00374 | 4 | 0.163170163 |
| TpMuguga_02g00374 | 5 | 0.097902098 |
| TpMuguga_02g00374 | 6 | 0.587412587 |
| TpMuguga_02g00375 | 1 | 0 |
| TpMuguga_02g00375 | 2 | 0.071656051 |
| TpMuguga_02g00376 | 1 | 0 |

| | | |
|---|---|---|
| TpMuguga_02g00376 | 2 | 0 |
| TpMuguga_02g00376 | 3 | 0 |
| TpMuguga_02g00376 | 4 | 0 |
| TpMuguga_02g00376 | 5 | 0 |
| TpMuguga_02g00377 | 1 | 0 |
| TpMuguga_02g00378 | 1 | 0 |
| TpMuguga_02g00378 | 2 | 6.944444444 |
| TpMuguga_02g00378 | 3 | 5.555555556 |
| TpMuguga_02g00378 | 4 | 2.777777778 |
| TpMuguga_02g00380 | 1 | 0 |
| TpMuguga_02g00380 | 2 | 0 |
| TpMuguga_02g00380 | 3 | 0 |
| TpMuguga_02g00380 | 4 | 0 |
| TpMuguga_02g00380 | 5 | 0 |
| TpMuguga_02g00380 | 6 | 0 |
| TpMuguga_02g00380 | 7 | 0 |
| TpMuguga_02g00965 | 1 | 0 |
| TpMuguga_02g00965 | 2 | 0 |
| TpMuguga_02g00965 | 3 | 2.666666667 |
| TpMuguga_02g00381 | 1 | 0 |
| TpMuguga_02g00381 | 2 | 0 |
| TpMuguga_02g00381 | 3 | 0 |
| TpMuguga_02g00382 | 1 | 0 |
| TpMuguga_02g00384 | 1 | 0 |
| TpMuguga_02g00384 | 2 | 0 |
| TpMuguga_02g02640 | 1 | 0 |
| TpMuguga_02g02640 | 2 | 0 |
| TpMuguga_02g02640 | 3 | 0 |
| TpMuguga_02g02640 | 4 | 0 |
| TpMuguga_02g02640 | 5 | 0 |
| TpMuguga_02g02640 | 6 | 0 |
| TpMuguga_02g02640 | 7 | 0 |
| TpMuguga_02g02640 | 8 | 0 |
| TpMuguga_02g02640 | 9 | 0 |
| TpMuguga_02g02640 | 10 | 0 |
| TpMuguga_02g02640 | 11 | 0 |
| TpMuguga_02g02640 | 12 | 0 |
| TpMuguga_02g00389 | 1 | 0 |
| TpMuguga_02g00389 | 2 | 0 |
| TpMuguga_02g00389 | 3 | 0 |
| TpMuguga_02g00389 | 4 | 0 |
| TpMuguga_02g00389 | 5 | 0 |

| | | |
|---|---|---|
| TpMuguga_02g00389 | 6 | 0 |
| TpMuguga_02g00389 | 7 | 0 |
| TpMuguga_02g00389 | 8 | 0 |
| TpMuguga_02g00389 | 9 | 0 |
| TpMuguga_02g00388 | 1 | 0.002522068 |
| TpMuguga_02g00392 | 1 | 0 |
| TpMuguga_02g00393 | 1 | 0 |
| TpMuguga_02g00393 | 2 | 0 |
| TpMuguga_02g00393 | 3 | 48 |
| TpMuguga_02g00393 | 4 | 0 |
| TpMuguga_02g00393 | 5 | 0 |
| TpMuguga_02g00393 | 6 | 0 |
| TpMuguga_02g00393 | 7 | 0 |
| TpMuguga_02g00393 | 8 | 0 |
| TpMuguga_02g00393 | 9 | 0 |
| TpMuguga_02g00393 | 10 | 0 |
| TpMuguga_02g00393 | 11 | 0 |
| TpMuguga_02g00394 | 1 | 0 |
| TpMuguga_02g00394 | 2 | 0.185840708 |
| TpMuguga_02g00395 | 1 | 0 |
| TpMuguga_02g00395 | 2 | 0 |
| TpMuguga_02g00395 | 3 | 0 |
| TpMuguga_02g00395 | 4 | 0.146341463 |
| TpMuguga_02g00395 | 5 | 1.609756098 |
| TpMuguga_02g00396 | 1 | 0 |
| TpMuguga_02g00396 | 2 | 0 |
| TpMuguga_02g00397 | 1 | 0.705882353 |
| TpMuguga_02g00397 | 2 | 3.647058824 |
| TpMuguga_02g00397 | 3 | 0 |
| TpMuguga_02g00397 | 4 | 0 |
| TpMuguga_02g00397 | 5 | 0 |
| TpMuguga_02g00399 | 1 | 0 |
| TpMuguga_02g00399 | 2 | 0 |
| TpMuguga_02g00399 | 3 | 0 |
| TpMuguga_02g00399 | 4 | 0 |
| TpMuguga_02g00402 | 1 | 0 |
| TpMuguga_02g00402 | 2 | 0 |
| TpMuguga_02g00402 | 3 | 0 |
| TpMuguga_02g00402 | 4 | 0 |
| TpMuguga_02g00401 | 1 | 0 |
| TpMuguga_02g00401 | 2 | 0 |
| TpMuguga_02g00401 | 3 | 0 |

| | | |
|---|---|---|
| TpMuguga_02g00401 | 4 | 4.390243902 |
| TpMuguga_02g00401 | 5 | 1.463414634 |
| TpMuguga_02g00401 | 6 | 0.731707317 |
| TpMuguga_02g00401 | 7 | 1.951219512 |
| TpMuguga_02g00401 | 8 | 2.926829268 |
| TpMuguga_02g00401 | 9 | 0.894308943 |
| TpMuguga_02g00404 | 1 | 130 |
| TpMuguga_02g00404 | 2 | 0 |
| TpMuguga_02g00404 | 3 | 0 |
| TpMuguga_02g00404 | 4 | 0 |
| TpMuguga_02g00404 | 5 | 0 |
| TpMuguga_02g00404 | 6 | 0 |
| TpMuguga_02g00404 | 7 | 40 |
| TpMuguga_02g00404 | 8 | 0 |
| TpMuguga_02g00404 | 9 | 0 |
| TpMuguga_02g00405 | 1 | 0 |
| TpMuguga_02g00405 | 2 | 13 |
| TpMuguga_02g00405 | 3 | 0 |
| TpMuguga_02g00405 | 4 | 0 |
| TpMuguga_02g00405 | 5 | 6.5 |
| TpMuguga_02g00405 | 6 | 19.5 |
| TpMuguga_02g00405 | 7 | 0 |
| TpMuguga_02g00405 | 8 | 0 |
| TpMuguga_02g00405 | 9 | 0 |
| TpMuguga_02g00405 | 10 | 0 |
| TpMuguga_02g00405 | 11 | 0 |
| TpMuguga_02g00405 | 12 | 0 |
| TpMuguga_02g00408 | 1 | 0 |
| TpMuguga_02g00408 | 2 | 0 |
| TpMuguga_02g02560 | 1 | 0 |
| TpMuguga_02g02560 | 2 | 0 |
| TpMuguga_02g02560 | 3 | 0 |
| TpMuguga_02g02560 | 4 | 0 |
| TpMuguga_02g00412 | 1 | 0 |
| TpMuguga_02g00412 | 2 | 2.011210762 |
| TpMuguga_02g00413 | 1 | 0 |
| TpMuguga_02g00413 | 2 | 0 |
| TpMuguga_02g00413 | 3 | 0 |
| TpMuguga_02g00413 | 4 | 0 |
| TpMuguga_02g00413 | 5 | 0 |
| TpMuguga_02g00413 | 6 | 0 |
| TpMuguga_02g00413 | 7 | 0 |

| | | |
|---|---|---|
| TpMuguga_02g00413 | 8 | 0 |
| TpMuguga_02g00413 | 9 | 4.8125 |
| TpMuguga_02g00413 | 10 | 5.729166667 |
| TpMuguga_02g00414 | 1 | 0 |
| TpMuguga_02g00416 | 1 | 0 |
| TpMuguga_02g00417 | 1 | 1.578947368 |
| TpMuguga_02g00418 | 1 | 0.470588235 |
| TpMuguga_02g00419 | 1 | 0 |
| TpMuguga_02g00422 | 1 | 0 |
| TpMuguga_02g00423 | 1 | 0 |
| TpMuguga_02g00423 | 2 | 0 |
| TpMuguga_02g00425 | 1 | 0 |
| TpMuguga_02g00425 | 2 | 0.793650794 |
| TpMuguga_02g00425 | 3 | 1.19047619 |
| TpMuguga_02g00425 | 4 | 0.873015873 |
| TpMuguga_02g00426 | 1 | 0 |
| TpMuguga_02g00428 | 1 | 1.575757576 |
| TpMuguga_02g00429 | 1 | 0 |
| TpMuguga_02g00429 | 2 | 0 |
| TpMuguga_02g00430 | 1 | 0.244897959 |
| TpMuguga_02g00430 | 2 | 0 |
| TpMuguga_02g00431 | 1 | 0 |
| TpMuguga_02g00431 | 2 | 0 |
| TpMuguga_02g00431 | 3 | 0 |
| TpMuguga_02g00431 | 4 | 0 |
| TpMuguga_02g00431 | 5 | 0 |
| TpMuguga_02g00433 | 1 | 1.2 |
| TpMuguga_02g00432 | 1 | 0 |
| TpMuguga_02g00432 | 2 | 0 |
| TpMuguga_02g00434 | 1 | 0 |
| TpMuguga_02g00434 | 2 | 0 |
| TpMuguga_02g00434 | 3 | 0 |
| TpMuguga_02g00434 | 4 | 0 |
| TpMuguga_02g00435 | 1 | 0 |
| TpMuguga_02g00435 | 2 | 0 |
| TpMuguga_02g00435 | 3 | 0 |
| TpMuguga_02g00435 | 4 | 0 |
| TpMuguga_02g00435 | 5 | 0 |
| TpMuguga_02g00435 | 6 | 0 |
| TpMuguga_02g00435 | 7 | 0 |
| TpMuguga_02g00435 | 8 | 0 |
| TpMuguga_02g00435 | 9 | 0 |

| | | |
|---|---|---|
| TpMuguga_02g00435 | 10 | 0 |
| TpMuguga_02g00435 | 11 | 0 |
| TpMuguga_02g00439 | 1 | 1.833333333 |
| TpMuguga_02g00441 | 1 | 0 |
| TpMuguga_02g00441 | 2 | 0 |
| TpMuguga_02g00441 | 3 | 4 |
| TpMuguga_02g00442 | 1 | 0.285714286 |
| TpMuguga_02g00443 | 1 | 0 |
| TpMuguga_02g00443 | 2 | 0 |
| TpMuguga_02g00443 | 3 | 0 |
| TpMuguga_02g00443 | 4 | 0 |
| TpMuguga_02g00443 | 5 | 0 |
| TpMuguga_02g00443 | 6 | 0 |
| TpMuguga_02g00443 | 7 | 0 |
| TpMuguga_02g00443 | 8 | 0 |
| TpMuguga_02g00443 | 9 | 0 |
| TpMuguga_02g00444 | 1 | 0 |
| TpMuguga_02g00444 | 2 | 0 |
| TpMuguga_02g00444 | 3 | 0 |
| TpMuguga_02g00444 | 4 | 0 |
| TpMuguga_02g00444 | 5 | 0 |
| TpMuguga_02g00444 | 6 | 0 |
| TpMuguga_02g00444 | 7 | 6.486486486 |
| TpMuguga_02g00445 | 1 | 2.28 |
| TpMuguga_02g00445 | 2 | 0 |
| TpMuguga_02g00446 | 1 | 0 |
| TpMuguga_02g00446 | 2 | 0 |
| TpMuguga_02g00447 | 1 | 0 |
| TpMuguga_02g00447 | 2 | 0 |
| TpMuguga_02g00447 | 3 | 5.333333333 |
| TpMuguga_02g00447 | 4 | 0 |
| TpMuguga_02g00447 | 5 | 0 |
| TpMuguga_02g00447 | 6 | 1.333333333 |
| TpMuguga_02g00447 | 7 | 0 |
| TpMuguga_02g02115 | 1 | 0.3 |
| TpMuguga_02g02115 | 2 | 1.25 |
| TpMuguga_02g02115 | 3 | 0.9 |
| TpMuguga_02g02115 | 4 | 1.35 |
| TpMuguga_02g02115 | 5 | 0.9 |
| TpMuguga_02g02115 | 6 | 0.55 |
| TpMuguga_02g02115 | 7 | 0.9 |
| TpMuguga_02g00448 | 1 | 0 |

| | | |
|---|---|---|
| TpMuguga_02g00448 | 2 | 0 |
| TpMuguga_02g00448 | 3 | 1 |
| TpMuguga_02g00449 | 1 | 0.048951049 |
| TpMuguga_02g00451 | 1 | 0 |
| TpMuguga_02g02660 | 1 | 16.88888889 |
| TpMuguga_02g02660 | 2 | 0.888888889 |
| TpMuguga_02g02660 | 3 | 1.777777778 |
| TpMuguga_02g02660 | 4 | 0 |
| TpMuguga_02g02660 | 5 | 0 |
| TpMuguga_02g02660 | 6 | 0 |
| TpMuguga_02g02660 | 7 | 0 |
| TpMuguga_02g02510 | 1 | 0 |
| TpMuguga_02g02510 | 2 | 0 |
| TpMuguga_02g02510 | 3 | 0.081447964 |
| TpMuguga_02g02510 | 4 | 0 |
| TpMuguga_02g02510 | 5 | 0.21719457 |
| TpMuguga_02g02120 | 1 | 0 |
| TpMuguga_02g02120 | 2 | 0 |
| TpMuguga_02g02120 | 3 | 0 |
| TpMuguga_02g02120 | 4 | 0 |
| TpMuguga_02g02135 | 1 | 8 |
| TpMuguga_02g02135 | 2 | 0 |
| TpMuguga_02g02135 | 3 | 0 |
| TpMuguga_02g02135 | 4 | 0 |
| TpMuguga_02g02135 | 5 | 0 |
| TpMuguga_02g02135 | 6 | 0 |
| TpMuguga_02g02135 | 7 | 0 |
| TpMuguga_02g00460 | 1 | 0 |
| TpMuguga_02g00460 | 2 | 0 |
| TpMuguga_02g00460 | 3 | 0 |
| TpMuguga_02g02110 | 1 | 2.459749553 |
| TpMuguga_02g02110 | 2 | 1.708407871 |
| TpMuguga_02g02110 | 3 | 0.134168157 |
| TpMuguga_02g02110 | 4 | 0.026833631 |
| TpMuguga_02g02105 | 1 | 4 |
| TpMuguga_02g02105 | 2 | 0 |
| TpMuguga_02g02105 | 3 | 8 |
| TpMuguga_02g02105 | 4 | 0 |
| TpMuguga_02g00462 | 1 | 0 |
| TpMuguga_02g00462 | 2 | 0 |
| TpMuguga_02g00463 | 1 | 0 |
| TpMuguga_02g00463 | 2 | 0 |

| | | |
|---|---|---|
| TpMuguga_02g00463 | 3 | 0.003416856 |
| TpMuguga_02g00463 | 4 | 0.167425968 |
| TpMuguga_02g00463 | 5 | 5.812072893 |
| TpMuguga_02g00464 | 1 | 0.079314041 |
| TpMuguga_02g00467 | 1 | 0 |
| TpMuguga_02g00467 | 2 | 0 |
| TpMuguga_02g00468 | 1 | 0 |
| TpMuguga_02g00468 | 2 | 0 |
| TpMuguga_02g00468 | 3 | 0 |
| TpMuguga_02g02390 | 1 | 0 |
| TpMuguga_02g02390 | 2 | 0 |
| TpMuguga_02g00470 | 1 | 0 |
| TpMuguga_02g00469 | 1 | 2.619047619 |
| TpMuguga_02g00469 | 2 | 0 |
| TpMuguga_02g00469 | 3 | 0.523809524 |
| TpMuguga_02g00469 | 4 | 0 |
| TpMuguga_02g00469 | 5 | 0 |
| TpMuguga_02g00469 | 6 | 0 |
| TpMuguga_02g00469 | 7 | 5.761904762 |
| TpMuguga_02g00469 | 8 | 5.238095238 |
| TpMuguga_02g00469 | 9 | 1.047619048 |
| TpMuguga_02g00469 | 10 | 0 |
| TpMuguga_02g00471 | 1 | 0 |
| TpMuguga_02g00471 | 2 | 0 |
| TpMuguga_02g00471 | 3 | 0 |
| TpMuguga_02g00471 | 4 | 0 |
| TpMuguga_02g00473 | 1 | 0 |
| TpMuguga_02g00474 | 1 | 0 |
| TpMuguga_02g00474 | 2 | 0 |
| TpMuguga_02g00474 | 3 | 0 |
| TpMuguga_02g00475 | 1 | 0.833333333 |
| TpMuguga_02g00475 | 2 | 2.5 |
| TpMuguga_02g00475 | 3 | 0.833333333 |
| TpMuguga_02g00475 | 4 | 3.888888889 |
| TpMuguga_02g00475 | 5 | 1.388888889 |
| TpMuguga_02g00475 | 6 | 3.055555556 |
| TpMuguga_02g00475 | 7 | 4.444444444 |
| TpMuguga_02g00475 | 8 | 2.777777778 |
| TpMuguga_02g00475 | 9 | 0.277777778 |
| TpMuguga_02g00476 | 1 | 6 |
| TpMuguga_02g00476 | 2 | 9 |
| TpMuguga_02g00476 | 3 | 27 |

| | | |
|---|---|---|
| TpMuguga_02g00476 | 4 | 0 |
| TpMuguga_02g00476 | 5 | 0 |
| TpMuguga_02g00476 | 6 | 0 |
| TpMuguga_02g00476 | 7 | 0 |
| TpMuguga_02g00476 | 8 | 0 |
| TpMuguga_02g00478 | 1 | 0 |
| TpMuguga_02g00478 | 2 | 0 |
| TpMuguga_02g00478 | 3 | 0 |
| TpMuguga_02g00478 | 4 | 0 |
| TpMuguga_02g00478 | 5 | 0 |
| TpMuguga_02g00478 | 6 | 0 |
| TpMuguga_02g02350 | 1 | 0 |
| TpMuguga_02g02350 | 2 | 0 |
| TpMuguga_02g00480 | 1 | 3 |
| TpMuguga_02g00480 | 2 | 2.454545455 |
| TpMuguga_02g00481 | 1 | 0.599250936 |
| TpMuguga_02g00481 | 2 | 0.262172285 |
| TpMuguga_02g00481 | 3 | 0.337078652 |
| TpMuguga_02g00481 | 4 | 0 |
| TpMuguga_02g00481 | 5 | 0.411985019 |
| TpMuguga_02g00481 | 6 | 0.037453184 |
| TpMuguga_02g00481 | 7 | 0 |
| TpMuguga_02g00481 | 8 | 0.112359551 |
| TpMuguga_02g00481 | 9 | 0 |
| TpMuguga_02g00482 | 1 | 0 |
| TpMuguga_02g00483 | 1 | 0.093959732 |
| TpMuguga_02g00484 | 1 | 0 |
| TpMuguga_02g00485 | 1 | 0 |
| TpMuguga_02g00485 | 2 | 0 |
| TpMuguga_02g00486 | 1 | 0 |
| TpMuguga_02g00486 | 2 | 0 |
| TpMuguga_02g00486 | 3 | 2.909090909 |
| TpMuguga_02g00488 | 1 | 0 |
| TpMuguga_02g00488 | 2 | 4.666666667 |
| TpMuguga_02g00488 | 3 | 0.777777778 |
| TpMuguga_02g00488 | 4 | 2.333333333 |
| TpMuguga_02g00488 | 5 | 0 |
| TpMuguga_02g00488 | 6 | 1.555555556 |
| TpMuguga_02g00489 | 1 | 0 |
| TpMuguga_02g00490 | 1 | 0 |
| TpMuguga_02g00490 | 2 | 0 |
| TpMuguga_02g00490 | 3 | 0 |

| | | |
|---|---|---|
| TpMuguga_02g00490 | 4 | 0 |
| TpMuguga_02g00493 | 1 | 0.307692308 |
| TpMuguga_02g00494 | 1 | 0 |
| TpMuguga_02g00494 | 2 | 2.347826087 |
| TpMuguga_02g00494 | 3 | 2.739130435 |
| TpMuguga_02g00494 | 4 | 0.782608696 |
| TpMuguga_02g00494 | 5 | 0 |
| TpMuguga_02g00494 | 6 | 0.391304348 |
| TpMuguga_02g00494 | 7 | 0.391304348 |
| TpMuguga_02g00494 | 8 | 0 |
| TpMuguga_02g00495 | 1 | 0 |
| TpMuguga_02g00495 | 2 | 0 |
| TpMuguga_02g00495 | 3 | 0 |
| TpMuguga_02g00496 | 1 | 0 |
| TpMuguga_02g00498 | 1 | 0 |
| TpMuguga_02g00498 | 2 | 0 |
| TpMuguga_02g00499 | 1 | 0 |
| TpMuguga_02g00499 | 2 | 0 |
| TpMuguga_02g00499 | 3 | 0 |
| TpMuguga_02g00501 | 1 | 1.375 |
| TpMuguga_02g00502 | 1 | 0 |
| TpMuguga_02g00505 | 1 | 0 |
| TpMuguga_02g00505 | 2 | 0 |
| TpMuguga_02g00505 | 3 | 0 |
| TpMuguga_02g00505 | 4 | 0 |
| TpMuguga_02g00504 | 1 | 0 |
| TpMuguga_02g00504 | 2 | 0 |
| TpMuguga_02g00508 | 1 | 0.021709634 |
| TpMuguga_02g00508 | 2 | 0 |
| TpMuguga_02g00508 | 3 | 0 |
| TpMuguga_02g00511 | 1 | 0.207920792 |
| TpMuguga_02g00511 | 2 | 0.01980198 |
| TpMuguga_02g00511 | 3 | 0.00990099 |
| TpMuguga_02g00511 | 4 | 0.623762376 |
| TpMuguga_02g00511 | 5 | 0.653465347 |
| TpMuguga_02g00513 | 1 | 0.638297872 |
| TpMuguga_02g00513 | 2 | 1.042553191 |
| TpMuguga_02g00514 | 1 | 0 |
| TpMuguga_02g00516 | 1 | 0 |
| TpMuguga_02g00516 | 2 | 0 |
| TpMuguga_02g00516 | 3 | 0 |
| TpMuguga_02g00516 | 4 | 0 |

| | | |
|---|---|---|
| TpMuguga_02g00516 | 5 | 0 |
| TpMuguga_02g00516 | 6 | 0 |
| TpMuguga_02g00516 | 7 | 11.42857143 |
| TpMuguga_02g00517 | 1 | 0 |
| TpMuguga_02g00517 | 2 | 0 |
| TpMuguga_02g00517 | 3 | 0 |
| TpMuguga_02g00517 | 4 | 0 |
| TpMuguga_02g00517 | 5 | 0 |
| TpMuguga_02g00517 | 6 | 0 |
| TpMuguga_02g00517 | 7 | 0 |
| TpMuguga_02g00518 | 1 | 0 |
| TpMuguga_02g00518 | 2 | 2.5 |
| TpMuguga_02g00518 | 3 | 0 |
| TpMuguga_02g00518 | 4 | 0 |
| TpMuguga_02g00518 | 5 | 0 |
| TpMuguga_02g00518 | 6 | 0 |
| TpMuguga_02g00519 | 1 | 4.045454545 |
| TpMuguga_02g00520 | 1 | 0.555555556 |
| TpMuguga_02g00520 | 2 | 1.111111111 |
| TpMuguga_02g00520 | 3 | 2.222222222 |
| TpMuguga_02g00520 | 4 | 0 |
| TpMuguga_02g00520 | 5 | 2.777777778 |
| TpMuguga_02g00520 | 6 | 0.555555556 |
| TpMuguga_02g00520 | 7 | 1.666666667 |
| TpMuguga_02g00520 | 8 | 0 |
| TpMuguga_02g00520 | 9 | 0 |
| TpMuguga_02g02415 | 1 | 0.5 |
| TpMuguga_02g02415 | 2 | 0 |
| TpMuguga_02g02415 | 3 | 0 |
| TpMuguga_02g02415 | 4 | 0 |
| TpMuguga_02g02415 | 5 | 2 |
| TpMuguga_02g02415 | 6 | 0 |
| TpMuguga_02g00524 | 1 | 0 |
| TpMuguga_02g00525 | 1 | 0 |
| TpMuguga_02g00525 | 2 | 3.483870968 |
| TpMuguga_02g00525 | 3 | 1.35483871 |
| TpMuguga_02g00525 | 4 | 0 |
| TpMuguga_02g00525 | 5 | 0.580645161 |
| TpMuguga_02g00528 | 1 | 0 |
| TpMuguga_02g00528 | 2 | 0 |
| TpMuguga_02g00529 | 1 | 0 |
| TpMuguga_02g00529 | 2 | 0 |

| | | |
|---|---|---|
| TpMuguga_02g00529 | 3 | 0 |
| TpMuguga_02g00529 | 4 | 0 |
| TpMuguga_02g00530 | 1 | 0 |
| TpMuguga_02g00530 | 2 | 0 |
| TpMuguga_02g00530 | 3 | 1.064516129 |
| TpMuguga_02g00530 | 4 | 4.064516129 |
| TpMuguga_02g00530 | 5 | 6.870967742 |
| TpMuguga_02g00532 | 1 | 0 |
| TpMuguga_02g00532 | 2 | 0 |
| TpMuguga_02g00532 | 3 | 0 |
| TpMuguga_02g00532 | 4 | 0 |
| TpMuguga_02g00532 | 5 | 0 |
| TpMuguga_02g00532 | 6 | 0 |
| TpMuguga_02g00531 | 1 | 0 |
| TpMuguga_02g00531 | 2 | 0.004301075 |
| TpMuguga_02g00531 | 3 | 0.4 |
| TpMuguga_02g00533 | 1 | 0 |
| TpMuguga_02g00533 | 2 | 1.666666667 |
| TpMuguga_02g00533 | 3 | 0 |
| TpMuguga_02g00533 | 4 | 0 |
| TpMuguga_02g00536 | 1 | 4.734246575 |
| TpMuguga_02g00536 | 2 | 4.266666667 |
| TpMuguga_02g00536 | 3 | 5.640182648 |
| TpMuguga_02g00536 | 4 | 2.147945205 |
| TpMuguga_02g00536 | 5 | 1.300456621 |
| TpMuguga_02g00536 | 6 | 9.380821918 |
| TpMuguga_02g00536 | 7 | 0.175342466 |
| TpMuguga_02g00536 | 8 | 0 |
| TpMuguga_02g00536 | 9 | 0 |
| TpMuguga_02g00536 | 10 | 0 |
| TpMuguga_02g00536 | 11 | 0.043835616 |
| TpMuguga_02g00536 | 12 | 0.321461187 |
| TpMuguga_02g00536 | 13 | 0.087671233 |
| TpMuguga_02g00536 | 14 | 0.81826484 |
| TpMuguga_02g00536 | 15 | 2.191780822 |
| TpMuguga_02g00537 | 1 | 2.666666667 |
| TpMuguga_02g00537 | 2 | 0 |
| TpMuguga_02g00538 | 1 | 0.053571429 |
| TpMuguga_02g00538 | 2 | 0 |
| TpMuguga_02g00540 | 1 | 6.6 |
| TpMuguga_02g00540 | 2 | 0 |
| TpMuguga_02g00539 | 1 | 0.260869565 |

| | | |
|---|---|---|
| TpMuguga_02g00539 | 2 | 3.739130435 |
| TpMuguga_02g00539 | 3 | 1.304347826 |
| TpMuguga_02g02655 | 1 | 0 |
| TpMuguga_02g00541 | 1 | 0 |
| TpMuguga_02g00541 | 2 | 0 |
| TpMuguga_02g00543 | 1 | 0.25 |
| TpMuguga_02g00545 | 1 | 0 |
| TpMuguga_02g00545 | 2 | 24 |
| TpMuguga_02g00546 | 1 | 0 |
| TpMuguga_02g00546 | 2 | 0 |
| TpMuguga_02g02490 | 1 | 0.274285714 |
| TpMuguga_02g02490 | 2 | 0 |
| TpMuguga_02g02490 | 3 | 0.091428571 |
| TpMuguga_02g00548 | 1 | 2.149253731 |
| TpMuguga_02g00548 | 2 | 0.179104478 |
| TpMuguga_02g00548 | 3 | 0 |
| TpMuguga_02g00548 | 4 | 0 |
| TpMuguga_02g00548 | 5 | 0.134328358 |
| TpMuguga_02g00550 | 1 | 0 |
| TpMuguga_02g02425 | 1 | 0 |
| TpMuguga_02g02425 | 2 | 0 |
| TpMuguga_02g02425 | 3 | 0 |
| TpMuguga_02g02305 | 1 | 0.028248588 |
| TpMuguga_02g02305 | 2 | 0.451977401 |
| TpMuguga_02g02305 | 3 | 0.93220339 |
| TpMuguga_02g02305 | 4 | 0.423728814 |
| TpMuguga_02g00556 | 1 | 0.048076923 |
| TpMuguga_02g00556 | 2 | 0 |
| TpMuguga_02g00556 | 3 | 0 |
| TpMuguga_02g00556 | 4 | 0.060096154 |
| TpMuguga_02g00555 | 1 | 0.372093023 |
| TpMuguga_02g00555 | 2 | 0 |
| TpMuguga_02g00555 | 3 | 0 |
| TpMuguga_02g00557 | 1 | 1.333333333 |
| TpMuguga_02g00558 | 1 | 0 |
| TpMuguga_02g00558 | 2 | 0 |
| TpMuguga_02g00558 | 3 | 0 |
| TpMuguga_02g00558 | 4 | 0 |
| TpMuguga_02g00558 | 5 | 0 |
| TpMuguga_02g00558 | 6 | 0.275590551 |
| TpMuguga_02g00559 | 1 | 0.317880795 |
| TpMuguga_02g00559 | 2 | 0.139072848 |

| | | |
|---|---|---|
| TpMuguga_02g00560 | 1 | 0.418181818 |
| TpMuguga_02g00561 | 1 | 0.838095238 |
| TpMuguga_02g00562 | 1 | 0.246575342 |
| TpMuguga_02g00562 | 2 | 0.054794521 |
| TpMuguga_02g00562 | 3 | 0.301369863 |
| TpMuguga_02g00562 | 4 | 0.136986301 |
| TpMuguga_02g00562 | 5 | 0.630136986 |
| TpMuguga_02g00563 | 1 | 1 |
| TpMuguga_02g00563 | 2 | 3.5 |
| TpMuguga_02g00563 | 3 | 4.5 |
| TpMuguga_02g00563 | 4 | 0 |
| TpMuguga_02g02170 | 1 | 1.353383459 |
| TpMuguga_02g02170 | 2 | 2.932330827 |
| TpMuguga_02g02170 | 3 | 0 |
| TpMuguga_02g02170 | 4 | 0 |
| TpMuguga_02g02170 | 5 | 0 |
| TpMuguga_02g02170 | 6 | 0 |
| TpMuguga_02g02170 | 7 | 0 |
| TpMuguga_02g02170 | 8 | 0 |
| TpMuguga_02g02170 | 9 | 2.781954887 |
| TpMuguga_02g02280 | 1 | 0 |
| TpMuguga_02g02280 | 2 | 0 |
| TpMuguga_02g00571 | 1 | 0 |
| TpMuguga_02g00571 | 2 | 0 |
| TpMuguga_02g02100 | 1 | 0 |
| TpMuguga_02g02100 | 2 | 0 |
| TpMuguga_02g02100 | 3 | 2.142857143 |
| TpMuguga_02g02100 | 4 | 0 |
| TpMuguga_02g00574 | 1 | 0 |
| TpMuguga_02g00574 | 2 | 0 |
| TpMuguga_02g00574 | 3 | 0 |
| TpMuguga_02g00574 | 4 | 0 |
| TpMuguga_02g00574 | 5 | 0.030017153 |
| TpMuguga_02g00574 | 6 | 0 |
| TpMuguga_02g00575 | 1 | 0.006284916 |
| TpMuguga_02g00575 | 2 | 0 |
| TpMuguga_02g00577 | 1 | 0 |
| TpMuguga_02g00577 | 2 | 0 |
| TpMuguga_02g00577 | 3 | 0 |
| TpMuguga_02g00577 | 4 | 0 |
| TpMuguga_02g00579 | 1 | 0 |
| TpMuguga_02g00580 | 1 | 0.923076923 |

| | | |
|---|---|---|
| TpMuguga_02g00581 | 1 | 0 |
| TpMuguga_02g00581 | 2 | 0 |
| TpMuguga_02g00581 | 3 | 0.564102564 |
| TpMuguga_02g00581 | 4 | 6.299145299 |
| TpMuguga_02g00581 | 5 | 0 |
| TpMuguga_02g00581 | 6 | 0 |
| TpMuguga_02g00581 | 7 | 0 |
| TpMuguga_02g00581 | 8 | 0 |
| TpMuguga_02g00581 | 9 | 0.188034188 |
| TpMuguga_02g00581 | 10 | 0 |
| TpMuguga_02g00582 | 1 | 0.014285714 |
| TpMuguga_02g00583 | 1 | 0 |
| TpMuguga_02g00583 | 2 | 0 |
| TpMuguga_02g00583 | 3 | 0 |
| TpMuguga_02g00583 | 4 | 0 |
| TpMuguga_02g00585 | 1 | 0 |
| TpMuguga_02g00585 | 2 | 1.913580247 |
| TpMuguga_02g00585 | 3 | 2.222222222 |
| TpMuguga_02g00585 | 4 | 1.111111111 |
| TpMuguga_02g00584 | 1 | 4.851612903 |
| TpMuguga_02g00584 | 2 | 0.051612903 |
| TpMuguga_02g00584 | 3 | 0 |
| TpMuguga_02g00587 | 1 | 2.43902439 |
| TpMuguga_02g00587 | 2 | 0.975609756 |
| TpMuguga_02g00587 | 3 | 0.780487805 |
| TpMuguga_02g02240 | 1 | 0 |
| TpMuguga_02g02240 | 2 | 0 |
| TpMuguga_02g02240 | 3 | 0 |
| TpMuguga_02g02240 | 4 | 0 |
| TpMuguga_02g02240 | 5 | 1.636363636 |
| TpMuguga_02g00596 | 1 | 0 |
| TpMuguga_02g00596 | 2 | 0 |
| TpMuguga_02g00596 | 3 | 0 |
| TpMuguga_02g00596 | 4 | 0 |
| TpMuguga_02g00596 | 5 | 0 |
| TpMuguga_02g00596 | 6 | 0 |
| TpMuguga_02g00596 | 7 | 0 |
| TpMuguga_02g00596 | 8 | 0 |
| TpMuguga_02g00596 | 9 | 0 |
| TpMuguga_02g00597 | 1 | 0.11827957 |
| TpMuguga_02g00597 | 2 | 0.053763441 |
| TpMuguga_02g00595 | 1 | 0 |

| | | |
|---|---|---|
| TpMuguga_02g00595 | 2 | 0.130612245 |
| TpMuguga_02g00595 | 3 | 0 |
| TpMuguga_02g00595 | 4 | 0 |
| TpMuguga_02g00595 | 5 | 0 |
| TpMuguga_02g00595 | 6 | 0.424489796 |
| TpMuguga_02g00595 | 7 | 0.751020408 |
| TpMuguga_02g00598 | 1 | 0 |
| TpMuguga_02g00598 | 2 | 0 |
| TpMuguga_02g00598 | 3 | 0 |
| TpMuguga_02g00598 | 4 | 10 |
| TpMuguga_02g00598 | 5 | 0 |
| TpMuguga_02g00598 | 6 | 0 |
| TpMuguga_02g00598 | 7 | 0 |
| TpMuguga_02g00598 | 8 | 0 |
| TpMuguga_02g00598 | 9 | 0 |
| TpMuguga_02g00599 | 1 | 8.666666667 |
| TpMuguga_02g00600 | 1 | 1.090909091 |
| TpMuguga_02g00601 | 1 | 0 |
| TpMuguga_02g00601 | 2 | 0 |
| TpMuguga_02g00602 | 1 | 0 |
| TpMuguga_02g00602 | 2 | 3 |
| TpMuguga_02g00603 | 1 | 0 |
| TpMuguga_02g00603 | 2 | 0 |
| TpMuguga_02g00603 | 3 | 0 |
| TpMuguga_02g00603 | 4 | 0 |
| TpMuguga_02g00604 | 1 | 0.264150943 |
| TpMuguga_02g00604 | 2 | 0 |
| TpMuguga_02g00604 | 3 | 0 |
| TpMuguga_02g00605 | 1 | 0 |
| TpMuguga_02g00605 | 2 | 4.470588235 |
| TpMuguga_02g00605 | 3 | 0 |
| TpMuguga_02g00607 | 1 | 0.625 |
| TpMuguga_02g00607 | 2 | 1.875 |
| TpMuguga_02g00607 | 3 | 4.375 |
| TpMuguga_02g00607 | 4 | 1.25 |
| TpMuguga_02g00609 | 1 | 3.203389831 |
| TpMuguga_02g00609 | 2 | 0.711864407 |
| TpMuguga_02g00609 | 3 | 1.542372881 |
| TpMuguga_02g00609 | 4 | 0.118644068 |
| TpMuguga_02g00609 | 5 | 0.474576271 |
| TpMuguga_02g00609 | 6 | 1.423728814 |
| TpMuguga_02g00610 | 1 | 0.816326531 |

| | | |
|---|---|---|
| TpMuguga_02g00611 | 1 | 0.306306306 |
| TpMuguga_02g00613 | 1 | 0 |
| TpMuguga_02g00613 | 2 | 1.428571429 |
| TpMuguga_02g00613 | 3 | 0 |
| TpMuguga_02g00613 | 4 | 0 |
| TpMuguga_02g00612 | 1 | 0 |
| TpMuguga_02g00612 | 2 | 0 |
| TpMuguga_02g00612 | 3 | 0 |
| TpMuguga_02g00612 | 4 | 0 |
| TpMuguga_02g00612 | 5 | 0 |
| TpMuguga_02g02030 | 1 | 1.603098387 |
| TpMuguga_02g02030 | 2 | 0.000452981 |
| TpMuguga_02g02030 | 3 | 0.005435767 |
| TpMuguga_02g02030 | 4 | 0 |
| TpMuguga_02g00614 | 1 | 0.283950617 |
| TpMuguga_02g02615 | 1 | 0 |
| TpMuguga_02g02615 | 2 | 0 |
| TpMuguga_02g00617 | 1 | 0 |
| TpMuguga_02g00619 | 1 | 0 |
| TpMuguga_02g00619 | 2 | 0 |
| TpMuguga_02g00619 | 3 | 0 |
| TpMuguga_02g02470 | 1 | 0 |
| TpMuguga_02g02470 | 2 | 0 |
| TpMuguga_02g02470 | 3 | 0.381679389 |
| TpMuguga_02g02470 | 4 | 0 |
| TpMuguga_02g02470 | 5 | 0.152671756 |
| TpMuguga_02g02470 | 6 | 0.038167939 |
| TpMuguga_02g02470 | 7 | 1.06870229 |
| TpMuguga_02g02470 | 8 | 1.259541985 |
| TpMuguga_02g02470 | 9 | 1.641221374 |
| TpMuguga_02g02460 | 1 | 0 |
| TpMuguga_02g02460 | 2 | 0 |
| TpMuguga_02g00624 | 1 | 0 |
| TpMuguga_02g00625 | 1 | 0 |
| TpMuguga_02g00625 | 2 | 0.421052632 |
| TpMuguga_02g00625 | 3 | 0.842105263 |
| TpMuguga_02g00625 | 4 | 0 |
| TpMuguga_02g00625 | 5 | 2.315789474 |
| TpMuguga_02g00625 | 6 | 1.684210526 |
| TpMuguga_02g00625 | 7 | 0.421052632 |
| TpMuguga_02g00627 | 1 | 0.058252427 |
| TpMuguga_02g00627 | 2 | 1.106796117 |

| | | |
|---|---|---|
| TpMuguga_02g00627 | 3 | 0.349514563 |
| TpMuguga_02g00627 | 4 | 1.689320388 |
| TpMuguga_02g00627 | 5 | 0 |
| TpMuguga_02g00626 | 1 | 0 |
| TpMuguga_02g00629 | 1 | 1.56 |
| TpMuguga_02g00629 | 2 | 0 |
| TpMuguga_02g00630 | 1 | 0 |
| TpMuguga_02g00630 | 2 | 0 |
| TpMuguga_02g00630 | 3 | 0 |
| TpMuguga_02g00630 | 4 | 0 |
| TpMuguga_02g00630 | 5 | 0 |
| TpMuguga_02g00631 | 1 | 0 |
| TpMuguga_02g00631 | 2 | 0 |
| TpMuguga_02g00631 | 3 | 1.4 |
| TpMuguga_02g00631 | 4 | 0 |
| TpMuguga_02g00631 | 5 | 9.8 |
| TpMuguga_02g00631 | 6 | 19.6 |
| TpMuguga_02g00632 | 1 | 8 |
| TpMuguga_02g00632 | 2 | 3.555555556 |
| TpMuguga_02g00632 | 3 | 0 |
| TpMuguga_02g00632 | 4 | 0 |
| TpMuguga_02g00632 | 5 | 2.666666667 |
| TpMuguga_02g00632 | 6 | 0 |
| TpMuguga_02g00632 | 7 | 0 |
| TpMuguga_02g00633 | 1 | 0 |
| TpMuguga_02g02245 | 1 | 0 |
| TpMuguga_02g02245 | 2 | 4.444444444 |
| TpMuguga_02g02245 | 3 | 0 |
| TpMuguga_02g02245 | 4 | 0 |
| TpMuguga_02g02245 | 5 | 0 |
| TpMuguga_02g02245 | 6 | 0 |
| TpMuguga_02g02245 | 7 | 0 |
| TpMuguga_02g02245 | 8 | 0 |
| TpMuguga_02g02245 | 9 | 5.555555556 |
| TpMuguga_02g00636 | 1 | 0 |
| TpMuguga_02g00636 | 2 | 0 |
| TpMuguga_02g00636 | 3 | 0 |
| TpMuguga_02g00638 | 1 | 0 |
| TpMuguga_02g00638 | 2 | 0 |
| TpMuguga_02g00638 | 3 | 0 |
| TpMuguga_02g00638 | 4 | 0 |
| TpMuguga_02g00638 | 5 | 5.142857143 |

| | | |
|---|---|---|
| TpMuguga_02g00638 | 6 | 2.285714286 |
| TpMuguga_02g00638 | 7 | 0 |
| TpMuguga_02g00639 | 1 | 0 |
| TpMuguga_02g00639 | 2 | 0 |
| TpMuguga_02g00639 | 3 | 0 |
| TpMuguga_02g00639 | 4 | 0 |
| TpMuguga_02g00639 | 5 | 0 |
| TpMuguga_02g00639 | 6 | 3.333333333 |
| TpMuguga_02g00639 | 7 | 10.66666667 |
| TpMuguga_02g00639 | 8 | 3.666666667 |
| TpMuguga_02g00639 | 9 | 6.333333333 |
| TpMuguga_02g02230 | 1 | 2.608695652 |
| TpMuguga_02g02230 | 2 | 0.347826087 |
| TpMuguga_02g02230 | 3 | 4.347826087 |
| TpMuguga_02g02620 | 1 | 1.105263158 |
| TpMuguga_02g02620 | 2 | 0 |
| TpMuguga_02g00642 | 1 | 0.735159817 |
| TpMuguga_02g00642 | 2 | 0.383561644 |
| TpMuguga_02g00642 | 3 | 0.99086758 |
| TpMuguga_02g00642 | 4 | 0.287671233 |
| TpMuguga_02g00642 | 5 | 0.191780822 |
| TpMuguga_02g00642 | 6 | 0.255707763 |
| TpMuguga_02g02330 | 1 | 0 |
| TpMuguga_02g02330 | 2 | 1.5 |
| TpMuguga_02g00645 | 1 | 0 |
| TpMuguga_02g00645 | 2 | 0.48 |
| TpMuguga_02g00645 | 3 | 0 |
| TpMuguga_02g00647 | 1 | 0 |
| TpMuguga_02g00647 | 2 | 0 |
| TpMuguga_02g00646 | 1 | 0 |
| TpMuguga_02g00646 | 2 | 0 |
| TpMuguga_02g00646 | 3 | 0 |
| TpMuguga_02g00646 | 4 | 0 |
| TpMuguga_02g00646 | 5 | 0 |
| TpMuguga_02g00648 | 1 | 0 |
| TpMuguga_02g00648 | 2 | 0 |
| TpMuguga_02g00648 | 3 | 0 |
| TpMuguga_02g00648 | 4 | 6.571428571 |
| TpMuguga_02g00649 | 1 | 1.454545455 |
| TpMuguga_02g00649 | 2 | 1.454545455 |
| TpMuguga_02g00649 | 3 | 1.272727273 |
| TpMuguga_02g00649 | 4 | 0 |

| | | |
|---|---|---|
| TpMuguga_02g00649 | 5 | 0 |
| TpMuguga_02g00649 | 6 | 1.272727273 |
| TpMuguga_02g00649 | 7 | 0 |
| TpMuguga_02g00649 | 8 | 0 |
| TpMuguga_02g00649 | 9 | 0 |
| TpMuguga_02g00649 | 10 | 0 |
| TpMuguga_02g00649 | 11 | 0 |
| TpMuguga_02g00649 | 12 | 0.181818182 |
| TpMuguga_02g00649 | 13 | 0 |
| TpMuguga_02g00652 | 1 | 0.542986425 |
| TpMuguga_02g00652 | 2 | 0.20361991 |
| TpMuguga_02g00653 | 1 | 1.627697842 |
| TpMuguga_02g00653 | 2 | 1.48381295 |
| TpMuguga_02g00653 | 3 | 2.059352518 |
| TpMuguga_02g00653 | 4 | 3.714028777 |
| TpMuguga_02g00654 | 1 | 0 |
| TpMuguga_02g00655 | 1 | 1.166666667 |
| TpMuguga_02g00656 | 1 | 0.666666667 |
| TpMuguga_02g00657 | 1 | 0.642857143 |
| TpMuguga_02g00658 | 1 | 1.333333333 |
| TpMuguga_02g00659 | 1 | 0 |
| TpMuguga_02g00660 | 1 | 0 |
| TpMuguga_02g00661 | 1 | 0 |
| TpMuguga_02g00661 | 2 | 0 |
| TpMuguga_02g00661 | 3 | 0 |
| TpMuguga_02g00661 | 4 | 0 |
| TpMuguga_02g00661 | 5 | 0 |
| TpMuguga_02g00661 | 6 | 0 |
| TpMuguga_02g00661 | 7 | 0 |
| TpMuguga_02g00661 | 8 | 0 |
| TpMuguga_02g00661 | 9 | 0 |
| TpMuguga_02g00661 | 10 | 0 |
| TpMuguga_02g00661 | 11 | 0 |
| TpMuguga_02g00661 | 12 | 0 |
| TpMuguga_02g00662 | 1 | 0 |
| TpMuguga_02g00662 | 2 | 0 |
| TpMuguga_02g00662 | 3 | 0 |
| TpMuguga_02g00662 | 4 | 0 |
| TpMuguga_02g00662 | 5 | 0 |
| TpMuguga_02g00662 | 6 | 0 |
| TpMuguga_02g00662 | 7 | 0 |
| TpMuguga_02g00663 | 1 | 0 |

| | | |
|---|---|---|
| TpMuguga_02g00663 | 2 | 0 |
| TpMuguga_02g00663 | 3 | 0 |
| TpMuguga_02g00663 | 4 | 0 |
| TpMuguga_02g00664 | 1 | 0 |
| TpMuguga_02g00664 | 2 | 0 |
| TpMuguga_02g00665 | 1 | 0.666666667 |
| TpMuguga_02g00666 | 1 | 0.454545455 |
| TpMuguga_02g00666 | 2 | 1 |
| TpMuguga_02g00666 | 3 | 3.454545455 |
| TpMuguga_02g00667 | 1 | 383.5 |
| TpMuguga_02g00667 | 2 | 32.5 |
| TpMuguga_02g00667 | 3 | 0 |
| TpMuguga_02g00667 | 4 | 0 |
| TpMuguga_02g00667 | 5 | 0 |
| TpMuguga_02g00667 | 6 | 0 |
| TpMuguga_02g00667 | 7 | 0 |
| TpMuguga_02g00667 | 8 | 0 |
| TpMuguga_02g00667 | 9 | 0 |
| TpMuguga_02g00667 | 10 | 0 |
| TpMuguga_02g00667 | 11 | 26 |
| TpMuguga_02g00667 | 12 | 0 |
| TpMuguga_02g02605 | 1 | 0 |
| TpMuguga_02g02605 | 2 | 0 |
| TpMuguga_02g02600 | 1 | 24 |
| TpMuguga_02g02600 | 2 | 0 |
| TpMuguga_02g02600 | 3 | 0 |
| TpMuguga_02g02600 | 4 | 0 |
| TpMuguga_02g02600 | 5 | 0 |
| TpMuguga_02g00669 | 1 | 0 |
| TpMuguga_02g00669 | 2 | 0 |
| TpMuguga_02g00669 | 3 | 0 |
| TpMuguga_02g00669 | 4 | 0 |
| TpMuguga_02g00669 | 5 | 0 |
| TpMuguga_02g00669 | 6 | 0 |
| TpMuguga_02g00670 | 1 | 0 |
| TpMuguga_02g00670 | 2 | 0 |
| TpMuguga_02g00670 | 3 | 20 |
| TpMuguga_02g00671 | 1 | 4.363636364 |
| TpMuguga_02g00671 | 2 | 0 |
| TpMuguga_02g00671 | 3 | 0 |
| TpMuguga_02g00671 | 4 | 0 |
| TpMuguga_02g00671 | 5 | 0 |

| | | |
|---|---|---|
| TpMuguga_02g00671 | 6 | 0 |
| TpMuguga_02g00671 | 7 | 0 |
| TpMuguga_02g00673 | 1 | 0 |
| TpMuguga_02g00675 | 1 | 42 |
| TpMuguga_02g00675 | 2 | 0 |
| TpMuguga_02g00675 | 3 | 0 |
| TpMuguga_02g00675 | 4 | 0 |
| TpMuguga_02g00675 | 5 | 0 |
| TpMuguga_02g00677 | 1 | 0 |
| TpMuguga_02g00677 | 2 | 0 |
| TpMuguga_02g00677 | 3 | 0 |
| TpMuguga_02g00677 | 4 | 0 |
| TpMuguga_02g00677 | 5 | 4.444444444 |
| TpMuguga_02g00677 | 6 | 0 |
| TpMuguga_02g00677 | 7 | 0 |
| TpMuguga_02g00678 | 1 | 0 |
| TpMuguga_02g00678 | 2 | 0 |
| TpMuguga_02g00678 | 3 | 0 |
| TpMuguga_02g00678 | 4 | 0 |
| TpMuguga_02g00678 | 5 | 0 |
| TpMuguga_02g00678 | 6 | 0 |
| TpMuguga_02g00679 | 1 | 5 |
| TpMuguga_02g00679 | 2 | 0 |
| TpMuguga_02g00681 | 1 | 0 |
| TpMuguga_02g00681 | 2 | 0.266666667 |
| TpMuguga_02g00681 | 3 | 0.533333333 |
| TpMuguga_02g00680 | 1 | 2.777777778 |
| TpMuguga_02g00680 | 2 | 2.777777778 |
| TpMuguga_02g00680 | 3 | 2.777777778 |
| TpMuguga_02g00680 | 4 | 1.666666667 |
| TpMuguga_02g00683 | 1 | 0 |
| TpMuguga_02g00683 | 2 | 0 |
| TpMuguga_02g00683 | 3 | 0 |
| TpMuguga_02g00683 | 4 | 0 |
| TpMuguga_02g00683 | 5 | 0 |
| TpMuguga_02g00683 | 6 | 0 |
| TpMuguga_02g00682 | 1 | 0.05655527 |
| TpMuguga_02g00682 | 2 | 0.025706941 |
| TpMuguga_02g00682 | 3 | 0.105398458 |
| TpMuguga_02g00682 | 4 | 0.051413882 |
| TpMuguga_02g00682 | 5 | 2.010282776 |
| TpMuguga_02g00684 | 1 | 0 |

| | | |
|---|---|---|
| TpMuguga_02g00684 | 2 | 0 |
| TpMuguga_02g00684 | 3 | 0 |
| TpMuguga_02g00684 | 4 | 0 |
| TpMuguga_02g00684 | 5 | 0 |
| TpMuguga_02g00684 | 6 | 0 |
| TpMuguga_02g00685 | 1 | 0 |
| TpMuguga_02g00685 | 2 | 0 |
| TpMuguga_02g00685 | 3 | 0 |
| TpMuguga_02g00686 | 1 | 7 |
| TpMuguga_02g00686 | 2 | 0 |
| TpMuguga_02g00686 | 3 | 0 |
| TpMuguga_02g00686 | 4 | 0 |
| TpMuguga_02g00686 | 5 | 0 |
| TpMuguga_02g00686 | 6 | 0 |
| TpMuguga_02g00688 | 1 | 0 |
| TpMuguga_02g00688 | 2 | 0 |
| TpMuguga_02g00688 | 3 | 0 |
| TpMuguga_02g00688 | 4 | 0 |
| TpMuguga_02g00688 | 5 | 0 |
| TpMuguga_02g00688 | 6 | 0 |
| TpMuguga_02g00688 | 7 | 0 |
| TpMuguga_02g00688 | 8 | 0 |
| TpMuguga_02g00688 | 9 | 0 |
| TpMuguga_02g00688 | 10 | 0 |
| TpMuguga_02g00688 | 11 | 0 |
| TpMuguga_02g00688 | 12 | 2.926829268 |
| TpMuguga_02g00688 | 13 | 0 |
| TpMuguga_02g00688 | 14 | 9.512195122 |
| TpMuguga_02g00689 | 1 | 3 |
| TpMuguga_02g00689 | 2 | 0 |
| TpMuguga_02g00689 | 3 | 0 |
| TpMuguga_02g00689 | 4 | 0 |
| TpMuguga_02g00689 | 5 | 0 |
| TpMuguga_02g00689 | 6 | 0 |
| TpMuguga_02g00689 | 7 | 0 |
| TpMuguga_02g00689 | 8 | 0 |
| TpMuguga_02g00689 | 9 | 0 |
| TpMuguga_02g00689 | 10 | 0 |
| TpMuguga_02g00689 | 11 | 0 |
| TpMuguga_02g00690 | 1 | 0 |
| TpMuguga_02g00690 | 2 | 0 |
| TpMuguga_02g00690 | 3 | 0 |

| | | |
|---|---|---|
| TpMuguga_02g02650 | 1 | 0 |
| TpMuguga_02g02650 | 2 | 0 |
| TpMuguga_02g02185 | 1 | 0.921052632 |
| TpMuguga_02g02185 | 2 | 0.184210526 |
| TpMuguga_02g02185 | 3 | 0 |
| TpMuguga_02g02185 | 4 | 0 |
| TpMuguga_02g02185 | 5 | 0 |
| TpMuguga_02g02185 | 6 | 0 |
| TpMuguga_02g02260 | 1 | 0 |
| TpMuguga_02g02260 | 2 | 0 |
| TpMuguga_02g02260 | 3 | 0 |
| TpMuguga_02g00696 | 1 | 0 |
| TpMuguga_02g00696 | 2 | 0 |
| TpMuguga_02g00697 | 1 | 0 |
| TpMuguga_02g00698 | 1 | 0.025817556 |
| TpMuguga_02g00698 | 2 | 0.004302926 |
| TpMuguga_02g00698 | 3 | 0.595524957 |
| TpMuguga_02g00699 | 1 | 0 |
| TpMuguga_02g00699 | 2 | 0 |
| TpMuguga_02g00699 | 3 | 0.193548387 |
| TpMuguga_02g00699 | 4 | 0 |
| TpMuguga_02g00699 | 5 | 0 |
| TpMuguga_02g00699 | 6 | 0 |
| TpMuguga_02g00699 | 7 | 0 |
| TpMuguga_02g00700 | 1 | 0 |
| TpMuguga_02g00701 | 1 | 0 |
| TpMuguga_02g00702 | 1 | 0 |
| TpMuguga_02g00702 | 2 | 0 |
| TpMuguga_02g00702 | 3 | 1.736842105 |
| TpMuguga_02g00702 | 4 | 0 |
| TpMuguga_02g00702 | 5 | 7.526315789 |
| TpMuguga_02g00702 | 6 | 2.894736842 |
| TpMuguga_02g00702 | 7 | 0 |
| TpMuguga_02g00702 | 8 | 0 |
| TpMuguga_02g00702 | 9 | 0 |
| TpMuguga_02g00702 | 10 | 9.842105263 |
| TpMuguga_02g02050 | 1 | 0 |
| TpMuguga_02g00705 | 1 | 0.782608696 |
| TpMuguga_02g00706 | 1 | 0.169014085 |
| TpMuguga_02g00706 | 2 | 0 |
| TpMuguga_02g00962 | 1 | 0 |
| TpMuguga_02g00962 | 2 | 0 |

| | | |
|---|---|---|
| TpMuguga_02g00708 | 1 | 0 |
| TpMuguga_02g00708 | 2 | 0 |
| TpMuguga_02g00708 | 3 | 0 |
| TpMuguga_02g00708 | 4 | 0 |
| TpMuguga_02g00708 | 5 | 0 |
| TpMuguga_02g00708 | 6 | 0 |
| TpMuguga_02g00708 | 7 | 0 |
| TpMuguga_02g00708 | 8 | 0 |
| TpMuguga_02g00708 | 9 | 0 |
| TpMuguga_02g00708 | 10 | 0 |
| TpMuguga_02g02020 | 1 | 0.205882353 |
| TpMuguga_02g02020 | 2 | 0.117647059 |
| TpMuguga_02g02020 | 3 | 0 |
| TpMuguga_02g02020 | 4 | 0 |
| TpMuguga_02g02020 | 5 | 0.441176471 |
| TpMuguga_02g02020 | 6 | 0.529411765 |
| TpMuguga_02g00709 | 1 | 0 |
| TpMuguga_02g00709 | 2 | 0 |
| TpMuguga_02g00709 | 3 | 0 |
| TpMuguga_02g00709 | 4 | 0 |
| TpMuguga_02g02565 | 1 | 0 |
| TpMuguga_02g02565 | 2 | 0 |
| TpMuguga_02g02565 | 3 | 0 |
| TpMuguga_02g02565 | 4 | 10 |
| TpMuguga_02g00714 | 1 | 0 |
| TpMuguga_02g00715 | 1 | 0 |
| TpMuguga_02g00715 | 2 | 0 |
| TpMuguga_02g00715 | 3 | 0 |
| TpMuguga_02g00715 | 4 | 0 |
| TpMuguga_02g00716 | 1 | 0.337662338 |
| TpMuguga_02g00717 | 1 | 6 |
| TpMuguga_02g00717 | 2 | 0 |
| TpMuguga_02g00717 | 3 | 6 |
| TpMuguga_02g00720 | 1 | 0 |
| TpMuguga_02g00720 | 2 | 3 |
| TpMuguga_02g00722 | 1 | 0 |
| TpMuguga_02g00722 | 2 | 0 |
| TpMuguga_02g00723 | 1 | 0 |
| TpMuguga_02g00724 | 1 | 0 |
| TpMuguga_02g00724 | 2 | 0.110769231 |
| TpMuguga_02g00724 | 3 | 0.32 |
| TpMuguga_02g00725 | 1 | 0 |

| | | |
|---|---|---|
| TpMuguga_02g00725 | 2 | 0 |
| TpMuguga_02g00725 | 3 | 0 |
| TpMuguga_02g00725 | 4 | 0 |
| TpMuguga_02g00725 | 5 | 3.6 |
| TpMuguga_02g00725 | 6 | 0 |
| TpMuguga_02g00725 | 7 | 0 |
| TpMuguga_02g00725 | 8 | 0 |
| TpMuguga_02g00726 | 1 | 0 |
| TpMuguga_02g00726 | 2 | 0 |
| TpMuguga_02g00726 | 3 | 0 |
| TpMuguga_02g00726 | 4 | 0 |
| TpMuguga_02g00726 | 5 | 0 |
| TpMuguga_02g00726 | 6 | 0 |
| TpMuguga_02g00726 | 7 | 0 |
| TpMuguga_02g00726 | 8 | 0 |
| TpMuguga_02g00728 | 1 | 0 |
| TpMuguga_02g00728 | 2 | 0 |
| TpMuguga_02g00728 | 3 | 0 |
| TpMuguga_02g00728 | 4 | 0 |
| TpMuguga_02g00728 | 5 | 0 |
| TpMuguga_02g00728 | 6 | 0 |
| TpMuguga_02g00728 | 7 | 0 |
| TpMuguga_02g00729 | 1 | 0 |
| TpMuguga_02g00729 | 2 | 0 |
| TpMuguga_02g00730 | 1 | 0 |
| TpMuguga_02g00730 | 2 | 1.780701754 |
| TpMuguga_02g00730 | 3 | 0 |
| TpMuguga_02g00730 | 4 | 0 |
| TpMuguga_02g00730 | 5 | 0 |
| TpMuguga_02g00730 | 6 | 0 |
| TpMuguga_02g00732 | 1 | 0 |
| TpMuguga_02g02365 | 1 | 0.768089054 |
| TpMuguga_02g02235 | 1 | 0 |
| TpMuguga_02g02235 | 2 | 0 |
| TpMuguga_02g02235 | 3 | 0 |
| TpMuguga_02g02235 | 4 | 0 |
| TpMuguga_02g02235 | 5 | 0 |
| TpMuguga_02g02235 | 6 | 0 |
| TpMuguga_02g02235 | 7 | 0 |
| TpMuguga_02g02235 | 8 | 0 |
| TpMuguga_02g02235 | 9 | 0 |
| TpMuguga_02g00735 | 1 | 0.381231672 |

| | | |
|---|---|---|
| TpMuguga_02g00735 | 2 | 0.82111437 |
| TpMuguga_02g00735 | 3 | 2.390029326 |
| TpMuguga_02g00735 | 4 | 0.674486804 |
| TpMuguga_02g00736 | 1 | 7.05 |
| TpMuguga_02g00736 | 2 | 1.65 |
| TpMuguga_02g00736 | 3 | 5.7 |
| TpMuguga_02g00736 | 4 | 0.6 |
| TpMuguga_02g00736 | 5 | 0 |
| TpMuguga_02g00737 | 1 | 0 |
| TpMuguga_02g00737 | 2 | 0 |
| TpMuguga_02g00737 | 3 | 0 |
| TpMuguga_02g00737 | 4 | 0 |
| TpMuguga_02g00738 | 1 | 4.117647059 |
| TpMuguga_02g00738 | 2 | 0 |
| TpMuguga_02g00738 | 3 | 0 |
| TpMuguga_02g00738 | 4 | 0 |
| TpMuguga_02g00738 | 5 | 0 |
| TpMuguga_02g00738 | 6 | 0 |
| TpMuguga_02g00738 | 7 | 1.176470588 |
| TpMuguga_02g00738 | 8 | 0 |
| TpMuguga_02g00738 | 9 | 0 |
| TpMuguga_02g02090 | 1 | 5.789473684 |
| TpMuguga_02g02090 | 2 | 0 |
| TpMuguga_02g02090 | 3 | 0 |
| TpMuguga_02g02090 | 4 | 0 |
| TpMuguga_02g00740 | 1 | 0 |
| TpMuguga_02g00740 | 2 | 0 |
| TpMuguga_02g00740 | 3 | 0 |
| TpMuguga_02g00740 | 4 | 0.04454343 |
| TpMuguga_02g00740 | 5 | 0 |
| TpMuguga_02g00741 | 1 | 0.124804992 |
| TpMuguga_02g00741 | 2 | 0.132605304 |
| TpMuguga_02g00741 | 3 | 0 |
| TpMuguga_02g00741 | 4 | 0.210608424 |
| TpMuguga_02g00742 | 1 | 4.457142857 |
| TpMuguga_02g00742 | 2 | 0 |
| TpMuguga_02g00742 | 3 | 0.228571429 |
| TpMuguga_02g00743 | 1 | 0 |
| TpMuguga_02g00743 | 2 | 0 |
| TpMuguga_02g00743 | 3 | 12.5 |
| TpMuguga_02g00743 | 4 | 0 |
| TpMuguga_02g00744 | 1 | 0 |

| | | |
|---|---|---|
| TpMuguga_02g00744 | 2 | 0 |
| TpMuguga_02g00744 | 3 | 0 |
| TpMuguga_02g00744 | 4 | 0 |
| TpMuguga_02g00747 | 1 | 0 |
| TpMuguga_02g00747 | 2 | 0 |
| TpMuguga_02g00747 | 3 | 0 |
| TpMuguga_02g00745 | 1 | 0 |
| TpMuguga_02g00745 | 2 | 0 |
| TpMuguga_02g00746 | 1 | 0 |
| TpMuguga_02g00746 | 2 | 0 |
| TpMuguga_02g00749 | 1 | 0 |
| TpMuguga_02g00749 | 2 | 0 |
| TpMuguga_02g00749 | 3 | 0 |
| TpMuguga_02g00749 | 4 | 0 |
| TpMuguga_02g00749 | 5 | 0 |
| TpMuguga_02g00750 | 1 | 0 |
| TpMuguga_02g00750 | 2 | 0 |
| TpMuguga_02g00750 | 3 | 0 |
| TpMuguga_02g00750 | 4 | 0 |
| TpMuguga_02g00750 | 5 | 0 |
| TpMuguga_02g00750 | 6 | 0 |
| TpMuguga_02g00754 | 1 | 0 |
| TpMuguga_02g00754 | 2 | 0.48 |
| TpMuguga_02g00754 | 3 | 2.08 |
| TpMuguga_02g00755 | 1 | 0.404312668 |
| TpMuguga_02g00755 | 2 | 0.663072776 |
| TpMuguga_02g02635 | 1 | 0.509433962 |
| TpMuguga_02g02635 | 2 | 0.679245283 |
| TpMuguga_02g02635 | 3 | 1.528301887 |
| TpMuguga_02g02635 | 4 | 1.613207547 |
| TpMuguga_02g02635 | 5 | 1.698113208 |
| TpMuguga_02g02635 | 6 | 3.311320755 |
| TpMuguga_02g02635 | 7 | 3.820754717 |
| TpMuguga_02g02635 | 8 | 2.462264151 |
| TpMuguga_02g00759 | 1 | 0.148468708 |
| TpMuguga_02g00761 | 1 | 0 |
| TpMuguga_02g00762 | 1 | 0 |
| TpMuguga_02g00762 | 2 | 0 |
| TpMuguga_02g00762 | 3 | 0 |
| TpMuguga_02g00763 | 1 | 0.25 |
| TpMuguga_02g00763 | 2 | 0 |
| TpMuguga_02g00764 | 1 | 5.333333333 |

| | | |
|---|---|---|
| TpMuguga_02g00764 | 2 | 0 |
| TpMuguga_02g00764 | 3 | 0 |
| TpMuguga_02g00764 | 4 | 0 |
| TpMuguga_02g00764 | 5 | 0 |
| TpMuguga_02g00764 | 6 | 0 |
| TpMuguga_02g00764 | 7 | 0 |
| TpMuguga_02g00766 | 1 | 0 |
| TpMuguga_02g00766 | 2 | 5.4 |
| TpMuguga_02g00766 | 3 | 0 |
| TpMuguga_02g00766 | 4 | 0 |
| TpMuguga_02g00766 | 5 | 0 |
| TpMuguga_02g00766 | 6 | 0 |
| TpMuguga_02g00766 | 7 | 0 |
| TpMuguga_02g00766 | 8 | 0 |
| TpMuguga_02g00768 | 1 | 6 |
| TpMuguga_02g00768 | 2 | 0 |
| TpMuguga_02g00768 | 3 | 0 |
| TpMuguga_02g00768 | 4 | 0 |
| TpMuguga_02g00768 | 5 | 0 |
| TpMuguga_02g00767 | 1 | 0.18018018 |
| TpMuguga_02g00767 | 2 | 0 |
| TpMuguga_02g00767 | 3 | 0 |
| TpMuguga_02g00769 | 1 | 0 |
| TpMuguga_02g00770 | 1 | 1.363636364 |
| TpMuguga_02g00770 | 2 | 0 |
| TpMuguga_02g00771 | 1 | 0 |
| TpMuguga_02g00771 | 2 | 0 |
| TpMuguga_02g00771 | 3 | 2.5 |
| TpMuguga_02g00771 | 4 | 3 |
| TpMuguga_02g00771 | 5 | 0 |
| TpMuguga_02g00771 | 6 | 0 |
| TpMuguga_02g00771 | 7 | 0 |
| TpMuguga_02g00771 | 8 | 0 |
| TpMuguga_02g00771 | 9 | 0 |
| TpMuguga_02g00771 | 10 | 0 |
| TpMuguga_02g00772 | 1 | 0 |
| TpMuguga_02g00772 | 2 | 0 |
| TpMuguga_02g00772 | 3 | 0 |
| TpMuguga_02g00774 | 1 | 0 |
| TpMuguga_02g02630 | 1 | 0 |
| TpMuguga_02g02630 | 2 | 6.4 |
| TpMuguga_02g02630 | 3 | 0 |

| | | |
|---|---|---|
| TpMuguga_02g02540 | 1 | 0 |
| TpMuguga_02g02540 | 2 | 0 |
| TpMuguga_02g02540 | 3 | 2.578947368 |
| TpMuguga_02g02540 | 4 | 0 |
| TpMuguga_02g02540 | 5 | 0 |
| TpMuguga_02g02540 | 6 | 0 |
| TpMuguga_02g00777 | 1 | 0 |
| TpMuguga_02g00777 | 2 | 0 |
| TpMuguga_02g00777 | 3 | 5.4 |
| TpMuguga_02g00777 | 4 | 14.4 |
| TpMuguga_02g00777 | 5 | 5.4 |
| TpMuguga_02g00777 | 6 | 0 |
| TpMuguga_02g00777 | 7 | 0 |
| TpMuguga_02g00777 | 8 | 3.6 |
| TpMuguga_02g02275 | 1 | 0 |
| TpMuguga_02g02275 | 2 | 0 |
| TpMuguga_02g02275 | 3 | 0 |
| TpMuguga_02g00779 | 1 | 0 |
| TpMuguga_02g00779 | 2 | 0 |
| TpMuguga_02g00779 | 3 | 0 |
| TpMuguga_02g00780 | 1 | 0.88 |
| TpMuguga_02g00781 | 1 | 0 |
| TpMuguga_02g00781 | 2 | 0 |
| TpMuguga_02g00781 | 3 | 0 |
| TpMuguga_02g00781 | 4 | 0 |
| TpMuguga_02g00781 | 5 | 0 |
| TpMuguga_02g00783 | 1 | 0 |
| TpMuguga_02g00784 | 1 | 0.546583851 |
| TpMuguga_02g00784 | 2 | 0.360248447 |
| TpMuguga_02g00784 | 3 | 0.782608696 |
| TpMuguga_02g00786 | 1 | 0.931506849 |
| TpMuguga_02g00787 | 1 | 0 |
| TpMuguga_02g00790 | 1 | 0 |
| TpMuguga_02g00790 | 2 | 0 |
| TpMuguga_02g00790 | 3 | 3.230769231 |
| TpMuguga_02g00790 | 4 | 0.461538462 |
| TpMuguga_02g00790 | 5 | 3.692307692 |
| TpMuguga_02g00791 | 1 | 0 |
| TpMuguga_02g02625 | 1 | 0 |
| TpMuguga_02g02625 | 2 | 0 |
| TpMuguga_02g00792 | 1 | 1.612903226 |
| TpMuguga_02g00795 | 1 | 0.175 |

| | | |
|---|---|---|
| TpMuguga_02g00795 | 2 | 0 |
| TpMuguga_02g00795 | 3 | 0 |
| TpMuguga_02g00795 | 4 | 0 |
| TpMuguga_02g00795 | 5 | 4.2 |
| TpMuguga_02g00795 | 6 | 0.175 |
| TpMuguga_02g00796 | 1 | 0 |
| TpMuguga_02g00796 | 2 | 0 |
| TpMuguga_02g02025 | 1 | 0 |
| TpMuguga_02g00798 | 1 | 0 |
| TpMuguga_02g00800 | 1 | 0 |
| TpMuguga_02g00800 | 2 | 0 |
| TpMuguga_02g00799 | 1 | 0 |
| TpMuguga_02g00799 | 2 | 1.090909091 |
| TpMuguga_02g00799 | 3 | 0 |
| TpMuguga_02g00799 | 4 | 0 |
| TpMuguga_02g00799 | 5 | 0 |
| TpMuguga_02g00801 | 1 | 0 |
| TpMuguga_02g00801 | 2 | 0 |
| TpMuguga_02g00802 | 1 | 1.923076923 |
| TpMuguga_02g00804 | 1 | 0 |
| TpMuguga_02g00806 | 1 | 1.12 |
| TpMuguga_02g00807 | 1 | 0.577487765 |
| TpMuguga_02g00807 | 2 | 0.527848986 |
| TpMuguga_02g00808 | 1 | 0 |
| TpMuguga_02g00808 | 2 | 0 |
| TpMuguga_02g00808 | 3 | 0 |
| TpMuguga_02g00808 | 4 | 0 |
| TpMuguga_02g00808 | 5 | 0 |
| TpMuguga_02g00808 | 6 | 0 |
| TpMuguga_02g00808 | 7 | 0 |
| TpMuguga_02g00810 | 1 | 0 |
| TpMuguga_02g00811 | 1 | 0 |
| TpMuguga_02g00813 | 1 | 0 |
| TpMuguga_02g00817 | 1 | 4 |
| TpMuguga_02g00817 | 2 | 0 |
| TpMuguga_02g00817 | 3 | 0 |
| TpMuguga_02g00817 | 4 | 0 |
| TpMuguga_02g00817 | 5 | 0 |
| TpMuguga_02g00820 | 1 | 0 |
| TpMuguga_02g00820 | 2 | 0 |
| TpMuguga_02g00821 | 1 | 0 |
| TpMuguga_02g00821 | 2 | 0 |

| | | |
|---|---|---|
| TpMuguga_02g00821 | 3 | 0 |
| TpMuguga_02g00821 | 4 | 0 |
| TpMuguga_02g00821 | 5 | 0 |
| TpMuguga_02g00821 | 6 | 0 |
| TpMuguga_02g00821 | 7 | 0 |
| TpMuguga_02g00821 | 8 | 0 |
| TpMuguga_02g00821 | 9 | 0 |
| TpMuguga_02g00821 | 10 | 0 |
| TpMuguga_02g00821 | 11 | 0.857142857 |
| TpMuguga_02g02430 | 1 | 0.375 |
| TpMuguga_02g02430 | 2 | 0 |
| TpMuguga_02g02430 | 3 | 0.5625 |
| TpMuguga_02g02430 | 4 | 2.625 |
| TpMuguga_02g02430 | 5 | 0.9375 |
| TpMuguga_02g02430 | 6 | 4.125 |
| TpMuguga_02g02430 | 7 | 0.5625 |
| TpMuguga_02g02430 | 8 | 4.5 |
| TpMuguga_02g00824 | 1 | 0 |
| TpMuguga_02g00824 | 2 | 0 |
| TpMuguga_02g00824 | 3 | 0 |
| TpMuguga_02g00824 | 4 | 0 |
| TpMuguga_02g00824 | 5 | 0 |
| TpMuguga_02g00825 | 1 | 0 |
| TpMuguga_02g00825 | 2 | 0 |
| TpMuguga_02g00825 | 3 | 0 |
| TpMuguga_02g00825 | 4 | 0 |
| TpMuguga_02g00825 | 5 | 0 |
| TpMuguga_02g00825 | 6 | 2.333333333 |
| TpMuguga_02g00826 | 1 | 0 |
| TpMuguga_02g00826 | 2 | 0 |
| TpMuguga_02g00826 | 3 | 0 |
| TpMuguga_02g00826 | 4 | 0 |
| TpMuguga_02g00826 | 5 | 0 |
| TpMuguga_02g00827 | 1 | 15 |
| TpMuguga_02g00827 | 2 | 0 |
| TpMuguga_02g00827 | 3 | 0 |
| TpMuguga_02g00827 | 4 | 0 |
| TpMuguga_02g02545 | 1 | 0.341463415 |
| TpMuguga_02g02545 | 2 | 0.170731707 |
| TpMuguga_02g02545 | 3 | 8.365853659 |
| TpMuguga_02g02545 | 4 | 0 |
| TpMuguga_02g02545 | 5 | 1.536585366 |

| | | |
|---|---|---|
| TpMuguga_02g02545 | 6 | 0 |
| TpMuguga_02g00830 | 1 | 0 |
| TpMuguga_02g00830 | 2 | 0 |
| TpMuguga_02g00830 | 3 | 0 |
| TpMuguga_02g00830 | 4 | 0 |
| TpMuguga_02g00830 | 5 | 0 |
| TpMuguga_02g00830 | 6 | 0 |
| TpMuguga_02g00830 | 7 | 0 |
| TpMuguga_02g00831 | 1 | 0 |
| TpMuguga_02g00831 | 2 | 0 |
| TpMuguga_02g00831 | 3 | 0 |
| TpMuguga_02g00831 | 4 | 0.933333333 |
| TpMuguga_02g00831 | 5 | 4.9 |
| TpMuguga_02g00831 | 6 | 0.466666667 |
| TpMuguga_02g00832 | 1 | 0 |
| TpMuguga_02g00832 | 2 | 0 |
| TpMuguga_02g00832 | 3 | 0 |
| TpMuguga_02g00833 | 1 | 3.75 |
| TpMuguga_02g00833 | 2 | 1.25 |
| TpMuguga_02g00833 | 3 | 0 |
| TpMuguga_02g00833 | 4 | 0 |
| TpMuguga_02g00833 | 5 | 0 |
| TpMuguga_02g00833 | 6 | 0 |
| TpMuguga_02g00833 | 7 | 0 |
| TpMuguga_02g00833 | 8 | 8.75 |
| TpMuguga_02g00833 | 9 | 0 |
| TpMuguga_02g00834 | 1 | 0 |
| TpMuguga_02g00834 | 2 | 0 |
| TpMuguga_02g00834 | 3 | 0 |
| TpMuguga_02g00836 | 1 | 0.120481928 |
| TpMuguga_02g00836 | 2 | 1.204819277 |
| TpMuguga_02g00836 | 3 | 0.301204819 |
| TpMuguga_02g00836 | 4 | 0.120481928 |
| TpMuguga_02g00835 | 1 | 0 |
| TpMuguga_02g00835 | 2 | 0 |
| TpMuguga_02g00835 | 3 | 0 |
| TpMuguga_02g00835 | 4 | 0.909090909 |
| TpMuguga_02g00835 | 5 | 0.545454545 |
| TpMuguga_02g00837 | 1 | 0 |
| TpMuguga_02g00837 | 2 | 0 |
| TpMuguga_02g00837 | 3 | 0 |
| TpMuguga_02g00837 | 4 | 0 |

| | | |
|---|---|---|
| TpMuguga_02g00837 | 5 | 0 |
| TpMuguga_02g00837 | 6 | 9 |
| TpMuguga_02g00837 | 7 | 0 |
| TpMuguga_02g00837 | 8 | 0 |
| TpMuguga_02g00838 | 1 | 0 |
| TpMuguga_02g00838 | 2 | 0.888888889 |
| TpMuguga_02g00838 | 3 | 0 |
| TpMuguga_02g00838 | 4 | 0 |
| TpMuguga_02g00838 | 5 | 0 |
| TpMuguga_02g00838 | 6 | 0 |
| TpMuguga_02g00838 | 7 | 0 |
| TpMuguga_02g00839 | 1 | 0 |
| TpMuguga_02g00839 | 2 | 0 |
| TpMuguga_02g00839 | 3 | 0 |
| TpMuguga_02g00839 | 4 | 0 |
| TpMuguga_02g00839 | 5 | 0 |
| TpMuguga_02g00839 | 6 | 0 |
| TpMuguga_02g00843 | 1 | 0.129032258 |
| TpMuguga_02g00843 | 2 | 0 |
| TpMuguga_02g00843 | 3 | 0 |
| TpMuguga_02g00843 | 4 | 0 |
| TpMuguga_02g00843 | 5 | 0 |
| TpMuguga_02g00843 | 6 | 0 |
| TpMuguga_02g00843 | 7 | 0.258064516 |
| TpMuguga_02g00842 | 1 | 0 |
| TpMuguga_02g00842 | 2 | 0 |
| TpMuguga_02g00842 | 3 | 0 |
| TpMuguga_02g00844 | 1 | 0 |
| TpMuguga_02g00844 | 2 | 0 |
| TpMuguga_02g00844 | 3 | 0 |
| TpMuguga_02g00844 | 4 | 0 |
| TpMuguga_02g00844 | 5 | 0 |
| TpMuguga_02g00844 | 6 | 0 |
| TpMuguga_02g00844 | 7 | 0.021917808 |
| TpMuguga_02g00847 | 1 | 0 |
| TpMuguga_02g00847 | 2 | 0 |
| TpMuguga_02g00847 | 3 | 0 |
| TpMuguga_02g02150 | 1 | 2.411273486 |
| TpMuguga_02g02150 | 2 | 0.56993737 |
| TpMuguga_02g02150 | 3 | 0 |
| TpMuguga_02g02150 | 4 | 0.029227557 |
| TpMuguga_02g02150 | 5 | 0 |

| | | |
|---|---|---|
| TpMuguga_02g02150 | 6 | 0 |
| TpMuguga_02g00848 | 1 | 0 |
| TpMuguga_02g00848 | 2 | 0 |
| TpMuguga_02g00848 | 3 | 0 |
| TpMuguga_02g00848 | 4 | 0 |
| TpMuguga_02g00848 | 5 | 0 |
| TpMuguga_02g00848 | 6 | 0 |
| TpMuguga_02g00848 | 7 | 0 |
| TpMuguga_02g00848 | 8 | 0 |
| TpMuguga_02g00848 | 9 | 2.166666667 |
| TpMuguga_02g00848 | 10 | 0.393939394 |
| TpMuguga_02g00848 | 11 | 10.63636364 |
| TpMuguga_02g00848 | 12 | 4.333333333 |
| TpMuguga_02g00851 | 1 | 0 |
| TpMuguga_02g00851 | 2 | 0 |
| TpMuguga_02g00851 | 3 | 0 |
| TpMuguga_02g00853 | 1 | 0 |
| TpMuguga_02g00853 | 2 | 0 |
| TpMuguga_02g00853 | 3 | 2.666666667 |
| TpMuguga_02g00853 | 4 | 0 |
| TpMuguga_02g00853 | 5 | 0 |
| TpMuguga_02g00854 | 1 | 0 |
| TpMuguga_02g00861 | 1 | 0.475247525 |
| TpMuguga_02g00861 | 2 | 0 |
| TpMuguga_02g00861 | 3 | 0 |
| TpMuguga_02g02505 | 1 | 0.205479452 |
| TpMuguga_02g00864 | 1 | 0.326530612 |
| TpMuguga_02g00867 | 1 | 0 |
| TpMuguga_02g00867 | 2 | 0 |
| TpMuguga_02g00867 | 3 | 0 |
| TpMuguga_02g00868 | 1 | 0.4 |
| TpMuguga_02g00868 | 2 | 0 |
| TpMuguga_02g02525 | 1 | 0 |
| TpMuguga_02g00872 | 1 | 0 |
| TpMuguga_02g00873 | 1 | 0.151515152 |
| TpMuguga_02g00873 | 2 | 0 |
| TpMuguga_02g00874 | 1 | 0.13592233 |
| TpMuguga_02g00880 | 1 | 0 |
| TpMuguga_02g00880 | 2 | 0 |
| TpMuguga_02g00880 | 3 | 0 |
| TpMuguga_02g00881 | 1 | 0 |
| TpMuguga_02g00881 | 2 | 0.727272727 |

| | | |
|---|---|---|
| TpMuguga_02g00881 | 3 | 0.727272727 |
| TpMuguga_02g00881 | 4 | 0.181818182 |
| TpMuguga_02g00881 | 5 | 0 |
| TpMuguga_02g00881 | 6 | 1.272727273 |
| TpMuguga_02g00881 | 7 | 1.272727273 |
| TpMuguga_02g00882 | 1 | 0 |
| TpMuguga_02g00882 | 2 | 0 |
| TpMuguga_02g00883 | 1 | 3.991935484 |
| TpMuguga_02g00883 | 2 | 0 |
| TpMuguga_02g00883 | 3 | 0 |
| TpMuguga_02g00883 | 4 | 0 |
| TpMuguga_02g00883 | 5 | 0 |
| TpMuguga_02g00883 | 6 | 0 |
| TpMuguga_02g00883 | 7 | 0 |
| TpMuguga_02g00883 | 8 | 0 |
| TpMuguga_02g00883 | 9 | 0 |
| TpMuguga_02g00883 | 10 | 0 |
| TpMuguga_02g00885 | 1 | 0 |
| TpMuguga_02g00887 | 1 | 0 |
| TpMuguga_02g00887 | 2 | 0.578313253 |
| TpMuguga_02g00887 | 3 | 1.686746988 |
| TpMuguga_02g00886 | 1 | 0.125 |
| TpMuguga_02g00886 | 2 | 2.25 |
| TpMuguga_02g00886 | 3 | 0 |
| TpMuguga_02g00886 | 4 | 0 |
| TpMuguga_02g00886 | 5 | 0 |
| TpMuguga_02g00891 | 1 | 0.230769231 |
| TpMuguga_02g00891 | 2 | 0 |
| TpMuguga_02g00897 | 1 | 0 |
| TpMuguga_02g00897 | 2 | 0.030456853 |
| TpMuguga_02g00900 | 1 | 0.115384615 |
| TpMuguga_02g00901 | 1 | 0 |
| TpMuguga_02g00901 | 2 | 0 |
| TpMuguga_02g00901 | 3 | 0 |
| TpMuguga_02g00901 | 4 | 0 |
| TpMuguga_02g00902 | 1 | 0 |
| TpMuguga_02g00902 | 2 | 0 |
| TpMuguga_02g00904 | 1 | 2.143902439 |
| TpMuguga_02g00904 | 2 | 0.080487805 |
| TpMuguga_02g00903 | 1 | 0.163265306 |
| TpMuguga_02g00905 | 1 | 0 |
| TpMuguga_02g02075 | 1 | 0 |

| | | |
|---|---|---|
| TpMuguga_02g02075 | 2 | 0 |
| TpMuguga_02g02075 | 3 | 0 |
| TpMuguga_02g00906 | 1 | 0 |
| TpMuguga_02g00906 | 2 | 0 |
| TpMuguga_02g00906 | 3 | 0 |
| TpMuguga_02g00906 | 4 | 3.9375 |
| TpMuguga_02g00906 | 5 | 0 |
| TpMuguga_02g00906 | 6 | 0 |
| TpMuguga_02g00906 | 7 | 0 |
| TpMuguga_02g00906 | 8 | 0 |
| TpMuguga_02g00908 | 1 | 0.904977376 |
| TpMuguga_02g00912 | 1 | 0 |
| TpMuguga_02g00912 | 2 | 0 |
| TpMuguga_02g00913 | 1 | 0 |
| TpMuguga_02g00913 | 2 | 0 |
| TpMuguga_02g00913 | 3 | 0 |
| TpMuguga_02g00913 | 4 | 0 |
| TpMuguga_02g00916 | 1 | 0.32 |
| TpMuguga_02g00917 | 1 | 4.5 |
| TpMuguga_02g00917 | 2 | 0 |
| TpMuguga_02g00919 | 1 | 0 |
| TpMuguga_02g00920 | 1 | 0 |
| TpMuguga_02g00920 | 2 | 0 |
| TpMuguga_02g00920 | 3 | 0 |
| TpMuguga_02g00924 | 1 | 0 |
| TpMuguga_02g00926 | 1 | 1.6 |
| TpMuguga_02g00926 | 2 | 0 |
| TpMuguga_02g00926 | 3 | 4.4 |
| TpMuguga_02g00927 | 1 | 0 |
| TpMuguga_02g00927 | 2 | 6 |
| TpMuguga_02g00927 | 3 | 0 |
| TpMuguga_02g00927 | 4 | 0 |
| TpMuguga_02g00927 | 5 | 0 |
| TpMuguga_02g00928 | 1 | 5.555555556 |
| TpMuguga_02g00928 | 2 | 0 |
| TpMuguga_02g00928 | 3 | 0 |
| TpMuguga_02g00928 | 4 | 0 |
| TpMuguga_02g00929 | 1 | 0 |
| TpMuguga_02g00929 | 2 | 1.647058824 |
| TpMuguga_02g00929 | 3 | 0.705882353 |
| TpMuguga_02g00930 | 1 | 0 |
| TpMuguga_02g00930 | 2 | 0 |

| | | |
|---|---|---|
| TpMuguga_02g00930 | 3 | 0 |
| TpMuguga_02g00930 | 4 | 0 |
| TpMuguga_02g00930 | 5 | 0 |
| TpMuguga_02g02015 | 1 | 3.2 |
| TpMuguga_02g02015 | 2 | 0 |
| TpMuguga_02g02015 | 3 | 0 |
| TpMuguga_02g00931 | 1 | 0 |
| TpMuguga_02g00931 | 2 | 0 |
| TpMuguga_02g00931 | 3 | 0 |
| TpMuguga_02g00931 | 4 | 0.651356994 |
| TpMuguga_02g00931 | 5 | 0.964509395 |
| TpMuguga_02g00932 | 1 | 0 |
| TpMuguga_02g00933 | 1 | 0 |
| TpMuguga_02g00933 | 2 | 0 |
| TpMuguga_02g00934 | 1 | 0 |
| TpMuguga_02g00935 | 1 | 0 |
| TpMuguga_02g00935 | 2 | 0 |
| TpMuguga_02g00935 | 3 | 0 |
| TpMuguga_02g00937 | 1 | 0 |
| TpMuguga_02g00938 | 1 | 0 |
| TpMuguga_02g00938 | 2 | 0 |
| TpMuguga_02g00938 | 3 | 0 |
| TpMuguga_02g00938 | 4 | 0 |
| TpMuguga_02g00938 | 5 | 0 |
| TpMuguga_02g00938 | 6 | 0 |
| TpMuguga_02g00941 | 1 | 0.121412804 |
| TpMuguga_02g00941 | 2 | 0 |
| TpMuguga_02g00941 | 3 | 0.072847682 |
| TpMuguga_02g00941 | 4 | 0 |
| TpMuguga_02g00941 | 5 | 0 |
| TpMuguga_02g00941 | 6 | 0 |
| TpMuguga_02g00941 | 7 | 0 |
| TpMuguga_02g00941 | 8 | 0 |
| TpMuguga_02g00941 | 9 | 0 |
| TpMuguga_02g00941 | 10 | 0 |
| TpMuguga_02g00942 | 1 | 0 |
| TpMuguga_02g00944 | 1 | 0 |
| TpMuguga_02g00944 | 2 | 0 |
| TpMuguga_02g00944 | 3 | 0 |
| TpMuguga_02g00944 | 4 | 0 |
| TpMuguga_02g00944 | 5 | 0 |
| TpMuguga_02g00944 | 6 | 0 |

| | | |
|---|---|---|
| TpMuguga_02g00944 | 7 | 0 |
| TpMuguga_02g00964 | 1 | 0 |
| TpMuguga_02g00964 | 2 | 0 |
| TpMuguga_02g00945 | 1 | 7 |
| TpMuguga_02g00945 | 2 | 0 |
| TpMuguga_02g00945 | 3 | 0 |
| TpMuguga_02g00945 | 4 | 4 |
| TpMuguga_02g00945 | 5 | 0 |
| TpMuguga_02g00945 | 6 | 0 |
| TpMuguga_02g00948 | 1 | 0.695718654 |
| TpMuguga_02g00948 | 2 | 0 |
| TpMuguga_02g00948 | 3 | 0 |
| TpMuguga_02g00948 | 4 | 0 |
| TpMuguga_02g00948 | 5 | 0 |
| TpMuguga_02g00948 | 6 | 0 |
| TpMuguga_02g00949 | 1 | 27 |
| TpMuguga_02g00949 | 2 | 0 |
| TpMuguga_04g02000 | 1 | 0 |
| TpMuguga_04g02000 | 2 | 0.51228978 |
| TpMuguga_04g00003 | 1 | 0 |
| TpMuguga_04g00008 | 1 | 0 |
| TpMuguga_04g00008 | 2 | 0 |
| TpMuguga_04g00008 | 3 | 0 |
| TpMuguga_04g02555 | 1 | 0 |
| TpMuguga_04g02555 | 2 | 0 |
| TpMuguga_04g02555 | 3 | 0 |
| TpMuguga_04g02555 | 4 | 0 |
| TpMuguga_04g02555 | 5 | 0 |
| TpMuguga_04g02555 | 6 | 0 |
| TpMuguga_04g02555 | 7 | 0 |
| TpMuguga_04g02555 | 8 | 0 |
| TpMuguga_04g02555 | 9 | 0 |
| TpMuguga_04g02560 | 1 | 3.5 |
| TpMuguga_04g02560 | 2 | 1.5 |
| TpMuguga_04g02560 | 3 | 1.166666667 |
| TpMuguga_04g02560 | 4 | 0.833333333 |
| TpMuguga_04g02560 | 5 | 1.333333333 |
| TpMuguga_04g00026 | 1 | 0 |
| TpMuguga_04g00026 | 2 | 0 |
| TpMuguga_04g00026 | 3 | 0 |
| TpMuguga_04g00026 | 4 | 0 |
| TpMuguga_04g00026 | 5 | 0 |

| | | |
|---|---|---|
| TpMuguga_04g00026 | 6 | 0 |
| TpMuguga_04g00026 | 7 | 0 |
| TpMuguga_04g00026 | 8 | 0 |
| TpMuguga_04g00026 | 9 | 0 |
| TpMuguga_04g00027 | 1 | 0 |
| TpMuguga_04g00027 | 2 | 0 |
| TpMuguga_04g00027 | 3 | 0 |
| TpMuguga_04g00027 | 4 | 0 |
| TpMuguga_04g00027 | 5 | 0 |
| TpMuguga_04g00027 | 6 | 0 |
| TpMuguga_04g02425 | 1 | 0 |
| TpMuguga_04g02425 | 2 | 0 |
| TpMuguga_04g00031 | 1 | 0 |
| TpMuguga_04g00031 | 2 | 1 |
| TpMuguga_04g00031 | 3 | 6.666666667 |
| TpMuguga_04g00031 | 4 | 1.666666667 |
| TpMuguga_04g00031 | 5 | 1.333333333 |
| TpMuguga_04g00031 | 6 | 0.333333333 |
| TpMuguga_04g00032 | 1 | 0 |
| TpMuguga_04g00032 | 2 | 0 |
| TpMuguga_04g00032 | 3 | 0 |
| TpMuguga_04g00032 | 4 | 0.857142857 |
| TpMuguga_04g00032 | 5 | 0 |
| TpMuguga_04g00034 | 1 | 0 |
| TpMuguga_04g00034 | 2 | 0.225806452 |
| TpMuguga_04g00034 | 3 | 0 |
| TpMuguga_04g00034 | 4 | 0.225806452 |
| TpMuguga_04g00034 | 5 | 0.225806452 |
| TpMuguga_04g00034 | 6 | 0 |
| TpMuguga_04g00033 | 1 | 0.572504708 |
| TpMuguga_04g00033 | 2 | 0.165725047 |
| TpMuguga_04g00033 | 3 | 0 |
| TpMuguga_04g00033 | 4 | 0 |
| TpMuguga_04g00033 | 5 | 0 |
| TpMuguga_04g00033 | 6 | 0 |
| TpMuguga_04g00033 | 7 | 0 |
| TpMuguga_04g00035 | 1 | 3.818181818 |
| TpMuguga_04g00035 | 2 | 0 |
| TpMuguga_04g00036 | 1 | 0.788461538 |
| TpMuguga_04g00037 | 1 | 0 |
| TpMuguga_04g00037 | 2 | 0 |
| TpMuguga_04g00037 | 3 | 0 |

| | | |
|---|---|---|
| TpMuguga_04g00038 | 1 | 0.212121212 |
| TpMuguga_04g00038 | 2 | 5.393939394 |
| TpMuguga_04g00038 | 3 | 0.757575758 |
| TpMuguga_04g00038 | 4 | 1.575757576 |
| TpMuguga_04g00038 | 5 | 0 |
| TpMuguga_04g00038 | 6 | 0.181818182 |
| TpMuguga_04g00039 | 1 | 0.569732938 |
| TpMuguga_04g00039 | 2 | 0.338278932 |
| TpMuguga_04g00039 | 3 | 0.213649852 |
| TpMuguga_04g00039 | 4 | 0.605341246 |
| TpMuguga_04g00039 | 5 | 0.462908012 |
| TpMuguga_04g00040 | 1 | 0 |
| TpMuguga_04g00040 | 2 | 0 |
| TpMuguga_04g00040 | 3 | 0 |
| TpMuguga_04g00040 | 4 | 12.6 |
| TpMuguga_04g00040 | 5 | 0 |
| TpMuguga_04g00040 | 6 | 7 |
| TpMuguga_04g00041 | 1 | 0 |
| TpMuguga_04g00041 | 2 | 0.666666667 |
| TpMuguga_04g00041 | 3 | 0 |
| TpMuguga_04g02645 | 1 | 0 |
| TpMuguga_04g02645 | 2 | 0 |
| TpMuguga_04g02645 | 3 | 0 |
| TpMuguga_04g02645 | 4 | 0 |
| TpMuguga_04g02645 | 5 | 0 |
| TpMuguga_04g02645 | 6 | 33 |
| TpMuguga_04g02645 | 7 | 0 |
| TpMuguga_04g02645 | 8 | 0 |
| TpMuguga_04g02645 | 9 | 0 |
| TpMuguga_04g02645 | 10 | 0 |
| TpMuguga_04g02640 | 1 | 0 |
| TpMuguga_04g02640 | 2 | 0 |
| TpMuguga_04g02640 | 3 | 0 |
| TpMuguga_04g02640 | 4 | 0 |
| TpMuguga_04g02640 | 5 | 0 |
| TpMuguga_04g00044 | 1 | 0 |
| TpMuguga_04g00044 | 2 | 0 |
| TpMuguga_04g00045 | 1 | 0 |
| TpMuguga_04g00046 | 1 | 2 |
| TpMuguga_04g00049 | 1 | 0 |
| TpMuguga_04g00049 | 2 | 0 |
| TpMuguga_04g00050 | 1 | 0 |

| | | |
|---|---|---|
| TpMuguga_04g00050 | 2 | 0 |
| TpMuguga_04g00051 | 1 | 0.489130435 |
| TpMuguga_04g00051 | 2 | 0.22826087 |
| TpMuguga_04g00052 | 1 | 3 |
| TpMuguga_04g00052 | 2 | 6 |
| TpMuguga_04g00053 | 1 | 0 |
| TpMuguga_04g00055 | 1 | 0.029727498 |
| TpMuguga_04g00055 | 2 | 0.002477291 |
| TpMuguga_04g00057 | 1 | 0 |
| TpMuguga_04g00057 | 2 | 0 |
| TpMuguga_04g00057 | 3 | 0 |
| TpMuguga_04g00057 | 4 | 0 |
| TpMuguga_04g00059 | 1 | 0 |
| TpMuguga_04g00059 | 2 | 0 |
| TpMuguga_04g00059 | 3 | 0 |
| TpMuguga_04g00059 | 4 | 0 |
| TpMuguga_04g00059 | 5 | 0 |
| TpMuguga_04g00060 | 1 | 0 |
| TpMuguga_04g00060 | 2 | 0 |
| TpMuguga_04g00060 | 3 | 2.315789474 |
| TpMuguga_04g00060 | 4 | 2.736842105 |
| TpMuguga_04g00060 | 5 | 4.210526316 |
| TpMuguga_04g00061 | 1 | 0 |
| TpMuguga_04g00061 | 2 | 0 |
| TpMuguga_04g00061 | 3 | 0 |
| TpMuguga_04g00061 | 4 | 0 |
| TpMuguga_04g00064 | 1 | 0 |
| TpMuguga_04g00063 | 1 | 0 |
| TpMuguga_04g00063 | 2 | 0 |
| TpMuguga_04g00063 | 3 | 0 |
| TpMuguga_04g00063 | 4 | 0 |
| TpMuguga_04g00063 | 5 | 0.15037594 |
| TpMuguga_04g00063 | 6 | 0.30075188 |
| TpMuguga_04g00063 | 7 | 0.15037594 |
| TpMuguga_04g00063 | 8 | 0 |
| TpMuguga_04g00063 | 9 | 0 |
| TpMuguga_04g00065 | 1 | 0 |
| TpMuguga_04g00066 | 1 | 0 |
| TpMuguga_04g00066 | 2 | 0 |
| TpMuguga_04g00066 | 3 | 0 |
| TpMuguga_04g00066 | 4 | 0 |
| TpMuguga_04g00066 | 5 | 0 |

| | | |
|---|---|---|
| TpMuguga_04g00066 | 6 | 0 |
| TpMuguga_04g02400 | 1 | 0 |
| TpMuguga_04g02400 | 2 | 0 |
| TpMuguga_04g02395 | 1 | 0 |
| TpMuguga_04g02395 | 2 | 0 |
| TpMuguga_04g02395 | 3 | 0 |
| TpMuguga_04g02395 | 4 | 0 |
| TpMuguga_04g02395 | 5 | 0 |
| TpMuguga_04g02395 | 6 | 0 |
| TpMuguga_04g02395 | 7 | 0 |
| TpMuguga_04g02395 | 8 | 0 |
| TpMuguga_04g02395 | 9 | 0 |
| TpMuguga_04g02395 | 10 | 0 |
| TpMuguga_04g02395 | 11 | 0 |
| TpMuguga_04g02395 | 12 | 10.5625 |
| TpMuguga_04g00068 | 1 | 3.214285714 |
| TpMuguga_04g00068 | 2 | 2.142857143 |
| TpMuguga_04g00069 | 1 | 0 |
| TpMuguga_04g00069 | 2 | 0 |
| TpMuguga_04g00069 | 3 | 0 |
| TpMuguga_04g00070 | 1 | 0 |
| TpMuguga_04g00071 | 1 | 0 |
| TpMuguga_04g00074 | 1 | 0 |
| TpMuguga_04g00074 | 2 | 0 |
| TpMuguga_04g00074 | 3 | 0 |
| TpMuguga_04g00074 | 4 | 33 |
| TpMuguga_04g00074 | 5 | 0 |
| TpMuguga_04g00075 | 1 | 0 |
| TpMuguga_04g00076 | 1 | 0 |
| TpMuguga_04g00076 | 2 | 0.170940171 |
| TpMuguga_04g00076 | 3 | 1.504273504 |
| TpMuguga_04g02500 | 1 | 18.66666667 |
| TpMuguga_04g02500 | 2 | 0 |
| TpMuguga_04g02500 | 3 | 0 |
| TpMuguga_04g02500 | 4 | 0 |
| TpMuguga_04g02500 | 5 | 0 |
| TpMuguga_04g02500 | 6 | 0 |
| TpMuguga_04g02500 | 7 | 0 |
| TpMuguga_04g00087 | 1 | 0 |
| TpMuguga_04g00088 | 1 | 0 |
| TpMuguga_04g00088 | 2 | 0 |
| TpMuguga_04g00088 | 3 | 0 |

| | | |
|---|---|---|
| TpMuguga_04g00088 | 4 | 0 |
| TpMuguga_04g00089 | 1 | 0.492873218 |
| TpMuguga_04g00089 | 2 | 0.945236309 |
| TpMuguga_04g00089 | 3 | 0.425356339 |
| TpMuguga_04g00089 | 4 | 0.236309077 |
| TpMuguga_04g00089 | 5 | 0.175543886 |
| TpMuguga_04g00089 | 6 | 0.040510128 |
| TpMuguga_04g00089 | 7 | 1.492123031 |
| TpMuguga_04g00089 | 8 | 0.432108027 |
| TpMuguga_04g02455 | 1 | 0.021603184 |
| TpMuguga_04g02450 | 1 | 0.34375 |
| TpMuguga_04g02450 | 2 | 0 |
| TpMuguga_04g02450 | 3 | 0 |
| TpMuguga_04g02450 | 4 | 0 |
| TpMuguga_04g02450 | 5 | 0 |
| TpMuguga_04g02450 | 6 | 0 |
| TpMuguga_04g02450 | 7 | 0 |
| TpMuguga_04g02450 | 8 | 0 |
| TpMuguga_04g02450 | 9 | 1.03125 |
| TpMuguga_04g02450 | 10 | 6.53125 |
| TpMuguga_04g00092 | 1 | 0.458333333 |
| TpMuguga_04g00092 | 2 | 0.5 |
| TpMuguga_04g00093 | 1 | 1.028571429 |
| TpMuguga_04g00093 | 2 | 0 |
| TpMuguga_04g00093 | 3 | 0 |
| TpMuguga_04g00093 | 4 | 0 |
| TpMuguga_04g00093 | 5 | 2.228571429 |
| TpMuguga_04g00094 | 1 | 0 |
| TpMuguga_04g00104 | 1 | 0 |
| TpMuguga_04g00105 | 1 | 0 |
| TpMuguga_04g00105 | 2 | 0 |
| TpMuguga_04g00106 | 1 | 2.100238663 |
| TpMuguga_04g00106 | 2 | 0.935560859 |
| TpMuguga_04g00106 | 3 | 0.801909308 |
| TpMuguga_04g00106 | 4 | 0.439140811 |
| TpMuguga_04g00106 | 5 | 0.210023866 |
| TpMuguga_04g00106 | 6 | 0.076372315 |
| TpMuguga_04g00106 | 7 | 0 |
| TpMuguga_04g00107 | 1 | 0 |
| TpMuguga_04g00107 | 2 | 0.099447514 |
| TpMuguga_04g00107 | 3 | 0 |
| TpMuguga_04g00107 | 4 | 0 |

| | | |
|---|---|---|
| TpMuguga_04g00107 | 5 | 0.049723757 |
| TpMuguga_04g00107 | 6 | 0.198895028 |
| TpMuguga_04g00107 | 7 | 1.889502762 |
| TpMuguga_04g00107 | 8 | 2.08839779 |
| TpMuguga_04g00108 | 1 | 0.029535865 |
| TpMuguga_04g00108 | 2 | 0.139240506 |
| TpMuguga_04g00109 | 1 | 0 |
| TpMuguga_04g00110 | 1 | 0 |
| TpMuguga_04g00111 | 1 | 6.080742459 |
| TpMuguga_04g00111 | 2 | 0.555452436 |
| TpMuguga_04g00111 | 3 | 0.338283063 |
| TpMuguga_04g00111 | 4 | 0.008352668 |
| TpMuguga_04g00111 | 5 | 0.004176334 |
| TpMuguga_04g00111 | 6 | 0.004176334 |
| TpMuguga_04g00111 | 7 | 0.045939675 |
| TpMuguga_04g00111 | 8 | 0 |
| TpMuguga_04g00112 | 1 | 0.263157895 |
| TpMuguga_04g00112 | 2 | 0.026315789 |
| TpMuguga_04g00113 | 1 | 0.575342466 |
| TpMuguga_04g00114 | 1 | 0 |
| TpMuguga_04g00114 | 2 | 0 |
| TpMuguga_04g00117 | 1 | 0 |
| TpMuguga_04g02385 | 1 | 0.207407407 |
| TpMuguga_04g02385 | 2 | 0.607407407 |
| TpMuguga_04g02385 | 3 | 0.4 |
| TpMuguga_04g00118 | 1 | 0 |
| TpMuguga_04g00118 | 2 | 1.212121212 |
| TpMuguga_04g00118 | 3 | 0 |
| TpMuguga_04g00118 | 4 | 0.363636364 |
| TpMuguga_04g00118 | 5 | 0 |
| TpMuguga_04g00118 | 6 | 0 |
| TpMuguga_04g00118 | 7 | 0 |
| TpMuguga_04g00121 | 1 | 0 |
| TpMuguga_04g00121 | 2 | 0 |
| TpMuguga_04g00122 | 1 | 0 |
| TpMuguga_04g00122 | 2 | 0.222222222 |
| TpMuguga_04g00123 | 1 | 0 |
| TpMuguga_04g02590 | 1 | 9.333333333 |
| TpMuguga_04g02590 | 2 | 0 |
| TpMuguga_04g02590 | 3 | 1.333333333 |
| TpMuguga_04g00125 | 1 | 0 |
| TpMuguga_04g00125 | 2 | 0 |

| | | |
|---|---|---|
| TpMuguga_04g00126 | 1 | 0 |
| TpMuguga_04g02480 | 1 | 0 |
| TpMuguga_04g02480 | 2 | 0 |
| TpMuguga_04g00129 | 1 | 0 |
| TpMuguga_04g00129 | 2 | 0.718954248 |
| TpMuguga_04g00129 | 3 | 2.22875817 |
| TpMuguga_04g00129 | 4 | 6.614379085 |
| TpMuguga_04g00129 | 5 | 2.947712418 |
| TpMuguga_04g00129 | 6 | 1.869281046 |
| TpMuguga_04g00129 | 7 | 0.431372549 |
| TpMuguga_04g00129 | 8 | 0.14379085 |
| TpMuguga_04g00129 | 9 | 1.294117647 |
| TpMuguga_04g00129 | 10 | 0.287581699 |
| TpMuguga_04g02405 | 1 | 2.8 |
| TpMuguga_04g02405 | 2 | 10.4 |
| TpMuguga_04g02405 | 3 | 5.2 |
| TpMuguga_04g02405 | 4 | 12 |
| TpMuguga_04g02405 | 5 | 3.6 |
| TpMuguga_04g02405 | 6 | 4.4 |
| TpMuguga_04g02405 | 7 | 0 |
| TpMuguga_04g02405 | 8 | 3.2 |
| TpMuguga_04g02405 | 9 | 2.8 |
| TpMuguga_04g02020 | 1 | 0 |
| TpMuguga_04g02020 | 2 | 0 |
| TpMuguga_04g02020 | 3 | 0 |
| TpMuguga_04g02020 | 4 | 0 |
| TpMuguga_04g02020 | 5 | 0 |
| TpMuguga_04g02020 | 6 | 0 |
| TpMuguga_04g02020 | 7 | 0 |
| TpMuguga_04g02020 | 8 | 0 |
| TpMuguga_04g02020 | 9 | 0 |
| TpMuguga_04g00134 | 1 | 6.517241379 |
| TpMuguga_04g00134 | 2 | 9.155172414 |
| TpMuguga_04g00134 | 3 | 3.413793103 |
| TpMuguga_04g00134 | 4 | 0 |
| TpMuguga_04g00134 | 5 | 0 |
| TpMuguga_04g00134 | 6 | 0 |
| TpMuguga_04g00134 | 7 | 0 |
| TpMuguga_04g00134 | 8 | 0 |
| TpMuguga_04g00135 | 1 | 0 |
| TpMuguga_04g00135 | 2 | 0 |
| TpMuguga_04g00135 | 3 | 0 |

| | | |
|---|---|---|
| TpMuguga_04g00135 | 4 | 0 |
| TpMuguga_04g00135 | 5 | 0 |
| TpMuguga_04g00135 | 6 | 0 |
| TpMuguga_04g00135 | 7 | 0 |
| TpMuguga_04g00136 | 1 | 0 |
| TpMuguga_04g00137 | 1 | 0.875 |
| TpMuguga_04g00137 | 2 | 0 |
| TpMuguga_04g00137 | 3 | 0 |
| TpMuguga_04g00137 | 4 | 0 |
| TpMuguga_04g00137 | 5 | 0 |
| TpMuguga_04g00137 | 6 | 2.625 |
| TpMuguga_04g00138 | 1 | 0.046601942 |
| TpMuguga_04g00138 | 2 | 0 |
| TpMuguga_04g00138 | 3 | 0 |
| TpMuguga_04g00138 | 4 | 0.011650485 |
| TpMuguga_04g00138 | 5 | 1.817475728 |
| TpMuguga_04g00139 | 1 | 0.371162791 |
| TpMuguga_04g00139 | 2 | 0 |
| TpMuguga_04g00139 | 3 | 0 |
| TpMuguga_04g00139 | 4 | 0 |
| TpMuguga_04g00139 | 5 | 0 |
| TpMuguga_04g00141 | 1 | 1.333333333 |
| TpMuguga_04g00141 | 2 | 3 |
| TpMuguga_04g00141 | 3 | 0 |
| TpMuguga_04g00141 | 4 | 0 |
| TpMuguga_04g00141 | 5 | 0 |
| TpMuguga_04g00141 | 6 | 0.333333333 |
| TpMuguga_04g00141 | 7 | 0 |
| TpMuguga_04g00141 | 8 | 0 |
| TpMuguga_04g00141 | 9 | 0 |
| TpMuguga_04g00141 | 10 | 0 |
| TpMuguga_04g00142 | 1 | 0 |
| TpMuguga_04g00143 | 1 | 0.833333333 |
| TpMuguga_04g00143 | 2 | 0 |
| TpMuguga_04g00143 | 3 | 0 |
| TpMuguga_04g00143 | 4 | 0.694444444 |
| TpMuguga_04g02025 | 1 | 0 |
| TpMuguga_04g00147 | 1 | 1.043478261 |
| TpMuguga_04g00147 | 2 | 0 |
| TpMuguga_04g00147 | 3 | 0 |
| TpMuguga_04g00148 | 1 | 0.242424242 |
| TpMuguga_04g00149 | 1 | 0 |

| | | |
|---|---|---|
| TpMuguga_04g00149 | 2 | 0 |
| TpMuguga_04g00149 | 3 | 0 |
| TpMuguga_04g00151 | 1 | 0 |
| TpMuguga_04g00151 | 2 | 0 |
| TpMuguga_04g00151 | 3 | 0 |
| TpMuguga_04g00152 | 1 | 0 |
| TpMuguga_04g00152 | 2 | 12 |
| TpMuguga_04g00152 | 3 | 0 |
| TpMuguga_04g00152 | 4 | 0 |
| TpMuguga_04g00152 | 5 | 0 |
| TpMuguga_04g00153 | 1 | 0 |
| TpMuguga_04g00155 | 1 | 0 |
| TpMuguga_04g00155 | 2 | 0 |
| TpMuguga_04g00155 | 3 | 0 |
| TpMuguga_04g00156 | 1 | 0 |
| TpMuguga_04g02695 | 1 | 0.116883117 |
| TpMuguga_04g02695 | 2 | 0 |
| TpMuguga_04g02690 | 1 | 0 |
| TpMuguga_04g02690 | 2 | 24 |
| TpMuguga_04g02690 | 3 | 0 |
| TpMuguga_04g02150 | 1 | 0 |
| TpMuguga_04g02290 | 1 | 0 |
| TpMuguga_04g02290 | 2 | 0 |
| TpMuguga_04g00161 | 1 | 2.454545455 |
| TpMuguga_04g00161 | 2 | 0.772727273 |
| TpMuguga_04g00161 | 3 | 0.181818182 |
| TpMuguga_04g00162 | 1 | 0.382978723 |
| TpMuguga_04g00162 | 2 | 5.808510638 |
| TpMuguga_04g00163 | 1 | 0.989473684 |
| TpMuguga_04g00166 | 1 | 2.480769231 |
| TpMuguga_04g00166 | 2 | 0.173076923 |
| TpMuguga_04g00166 | 3 | 0.75 |
| TpMuguga_04g00166 | 4 | 0 |
| TpMuguga_04g00166 | 5 | 0.403846154 |
| TpMuguga_04g00167 | 1 | 0 |
| TpMuguga_04g00167 | 2 | 0 |
| TpMuguga_04g00167 | 3 | 0 |
| TpMuguga_04g00167 | 4 | 0 |
| TpMuguga_04g00167 | 5 | 0 |
| TpMuguga_04g00167 | 6 | 0 |
| TpMuguga_04g00167 | 7 | 0 |
| TpMuguga_04g00167 | 8 | 0 |

| | | |
|---|---|---|
| TpMuguga_04g00167 | 9 | 0 |
| TpMuguga_04g02380 | 1 | 0 |
| TpMuguga_04g02380 | 2 | 0 |
| TpMuguga_04g00169 | 1 | 0.75 |
| TpMuguga_04g00169 | 2 | 0.490384615 |
| TpMuguga_04g00168 | 1 | 0.059701493 |
| TpMuguga_04g00168 | 2 | 0 |
| TpMuguga_04g00168 | 3 | 0.07960199 |
| TpMuguga_04g00170 | 1 | 0.203389831 |
| TpMuguga_04g00170 | 2 | 0 |
| TpMuguga_04g00170 | 3 | 0 |
| TpMuguga_04g00170 | 4 | 0 |
| TpMuguga_04g00170 | 5 | 0.745762712 |
| TpMuguga_04g00170 | 6 | 0 |
| TpMuguga_04g00170 | 7 | 1.288135593 |
| TpMuguga_04g02210 | 1 | 1.363636364 |
| TpMuguga_04g02210 | 2 | 0.909090909 |
| TpMuguga_04g02210 | 3 | 0 |
| TpMuguga_04g02210 | 4 | 0 |
| TpMuguga_04g00171 | 1 | 1.813333333 |
| TpMuguga_04g00171 | 2 | 0.906666667 |
| TpMuguga_04g00171 | 3 | 0.266666667 |
| TpMuguga_04g00172 | 1 | 0 |
| TpMuguga_04g00172 | 2 | 0 |
| TpMuguga_04g00173 | 1 | 0 |
| TpMuguga_04g00173 | 2 | 0 |
| TpMuguga_04g00174 | 1 | 0 |
| TpMuguga_04g00174 | 2 | 0 |
| TpMuguga_04g00174 | 3 | 0 |
| TpMuguga_04g00174 | 4 | 0 |
| TpMuguga_04g00174 | 5 | 0 |
| TpMuguga_04g00174 | 6 | 0 |
| TpMuguga_04g00174 | 7 | 0.4375 |
| TpMuguga_04g00174 | 8 | 0 |
| TpMuguga_04g00174 | 9 | 1.3125 |
| TpMuguga_04g00174 | 10 | 0 |
| TpMuguga_04g00174 | 11 | 0 |
| TpMuguga_04g00174 | 12 | 0 |
| TpMuguga_04g00174 | 13 | 17.9375 |
| TpMuguga_04g00175 | 1 | 0 |
| TpMuguga_04g00175 | 2 | 0 |
| TpMuguga_04g00177 | 1 | 0 |

| | | |
|---|---|---|
| TpMuguga_04g00177 | 2 | 0 |
| TpMuguga_04g00177 | 3 | 0 |
| TpMuguga_04g00178 | 1 | 0 |
| TpMuguga_04g00179 | 1 | 0 |
| TpMuguga_04g00182 | 1 | 0 |
| TpMuguga_04g00182 | 2 | 0 |
| TpMuguga_04g00182 | 3 | 0 |
| TpMuguga_04g00183 | 1 | 9 |
| TpMuguga_04g00183 | 2 | 4.5 |
| TpMuguga_04g00183 | 3 | 4.5 |
| TpMuguga_04g00183 | 4 | 0 |
| TpMuguga_04g00183 | 5 | 0 |
| TpMuguga_04g00183 | 6 | 0 |
| TpMuguga_04g00183 | 7 | 0 |
| TpMuguga_04g00183 | 8 | 0 |
| TpMuguga_04g00183 | 9 | 0 |
| TpMuguga_04g00183 | 10 | 0 |
| TpMuguga_04g00183 | 11 | 0 |
| TpMuguga_04g00185 | 1 | 0 |
| TpMuguga_04g00187 | 1 | 0 |
| TpMuguga_04g00187 | 2 | 0.041666667 |
| TpMuguga_04g00193 | 1 | 0 |
| TpMuguga_04g00193 | 2 | 0 |
| TpMuguga_04g00193 | 3 | 30 |
| TpMuguga_04g00194 | 1 | 0 |
| TpMuguga_04g00194 | 2 | 0 |
| TpMuguga_04g00194 | 3 | 0 |
| TpMuguga_04g00194 | 4 | 0 |
| TpMuguga_04g00195 | 1 | 0 |
| TpMuguga_04g00195 | 2 | 0 |
| TpMuguga_04g02065 | 1 | 0 |
| TpMuguga_04g02065 | 2 | 0 |
| TpMuguga_04g02230 | 1 | 0 |
| TpMuguga_04g02230 | 2 | 0 |
| TpMuguga_04g02230 | 3 | 0 |
| TpMuguga_04g02230 | 4 | 0 |
| TpMuguga_04g02230 | 5 | 0 |
| TpMuguga_04g02230 | 6 | 5.677419355 |
| TpMuguga_04g02230 | 7 | 13.16129032 |
| TpMuguga_04g02570 | 1 | 0.317180617 |
| TpMuguga_04g02570 | 2 | 1.797356828 |
| TpMuguga_04g02570 | 3 | 0.634361233 |

| | | |
|---|---|---|
| TpMuguga_04g02570 | 4 | 0.237885463 |
| TpMuguga_04g02570 | 5 | 0.343612335 |
| TpMuguga_04g00200 | 1 | 3 |
| TpMuguga_04g00200 | 2 | 0 |
| TpMuguga_04g00201 | 1 | 1.178571429 |
| TpMuguga_04g00201 | 2 | 2.892857143 |
| TpMuguga_04g02275 | 1 | 0.375 |
| TpMuguga_04g02275 | 2 | 0.9375 |
| TpMuguga_04g00202 | 1 | 0.6 |
| TpMuguga_04g00202 | 2 | 7.8 |
| TpMuguga_04g02625 | 1 | 5.142857143 |
| TpMuguga_04g02625 | 2 | 0 |
| TpMuguga_04g00206 | 1 | 0 |
| TpMuguga_04g00206 | 2 | 0 |
| TpMuguga_04g00206 | 3 | 0 |
| TpMuguga_04g00208 | 1 | 0 |
| TpMuguga_04g00208 | 2 | 0 |
| TpMuguga_04g00208 | 3 | 0 |
| TpMuguga_04g00208 | 4 | 0 |
| TpMuguga_04g00207 | 1 | 0.034482759 |
| TpMuguga_04g00209 | 1 | 0 |
| TpMuguga_04g00209 | 2 | 0 |
| TpMuguga_04g00209 | 3 | 0 |
| TpMuguga_04g00209 | 4 | 0 |
| TpMuguga_04g00209 | 5 | 0 |
| TpMuguga_04g00209 | 6 | 0 |
| TpMuguga_04g00209 | 7 | 0 |
| TpMuguga_04g00209 | 8 | 0 |
| TpMuguga_04g00209 | 9 | 0 |
| TpMuguga_04g00209 | 10 | 0 |
| TpMuguga_04g00209 | 11 | 0 |
| TpMuguga_04g00209 | 12 | 0 |
| TpMuguga_04g00209 | 13 | 0 |
| TpMuguga_04g00209 | 14 | 0 |
| TpMuguga_04g00209 | 15 | 0 |
| TpMuguga_04g00209 | 16 | 0 |
| TpMuguga_04g00209 | 17 | 0 |
| TpMuguga_04g00209 | 18 | 0 |
| TpMuguga_04g00209 | 19 | 0 |
| TpMuguga_04g00210 | 1 | 6.940068493 |
| TpMuguga_04g00210 | 2 | 0.455479452 |
| TpMuguga_04g00210 | 3 | 0.095890411 |

| | | |
|---|---|---|
| TpMuguga_04g00210 | 4 | 0 |
| TpMuguga_04g00210 | 5 | 0.047945205 |
| TpMuguga_04g00210 | 6 | 0.011986301 |
| TpMuguga_04g02260 | 1 | 0 |
| TpMuguga_04g02260 | 2 | 0 |
| TpMuguga_04g02260 | 3 | 0 |
| TpMuguga_04g02260 | 4 | 0 |
| TpMuguga_04g00213 | 1 | 0 |
| TpMuguga_04g00215 | 1 | 3.428571429 |
| TpMuguga_04g00215 | 2 | 0 |
| TpMuguga_04g00215 | 3 | 2.285714286 |
| TpMuguga_04g00215 | 4 | 2.285714286 |
| TpMuguga_04g00215 | 5 | 0 |
| TpMuguga_04g00215 | 6 | 8 |
| TpMuguga_04g00215 | 7 | 3.428571429 |
| TpMuguga_04g00216 | 1 | 0 |
| TpMuguga_04g00217 | 1 | 0.38961039 |
| TpMuguga_04g00217 | 2 | 0.194805195 |
| TpMuguga_04g00217 | 3 | 0 |
| TpMuguga_04g00217 | 4 | 0 |
| TpMuguga_04g02175 | 1 | 0 |
| TpMuguga_04g02175 | 2 | 0 |
| TpMuguga_04g02175 | 3 | 0 |
| TpMuguga_04g00218 | 1 | 0 |
| TpMuguga_04g00218 | 2 | 0 |
| TpMuguga_04g00218 | 3 | 0 |
| TpMuguga_04g00219 | 1 | 0 |
| TpMuguga_04g00219 | 2 | 0 |
| TpMuguga_04g00219 | 3 | 0 |
| TpMuguga_04g00220 | 1 | 0 |
| TpMuguga_04g00220 | 2 | 0 |
| TpMuguga_04g00221 | 1 | 0 |
| TpMuguga_04g02255 | 1 | 0 |
| TpMuguga_04g00225 | 1 | 0.016177958 |
| TpMuguga_04g00225 | 2 | 0 |
| TpMuguga_04g00225 | 3 | 0 |
| TpMuguga_04g00227 | 1 | 0 |
| TpMuguga_04g00227 | 2 | 0 |
| TpMuguga_04g00227 | 3 | 0 |
| TpMuguga_04g00228 | 1 | 0 |
| TpMuguga_04g00228 | 2 | 0 |
| TpMuguga_04g00228 | 3 | 0 |

| | | |
|---|---|---|
| TpMuguga_04g00228 | 4 | 0 |
| TpMuguga_04g00228 | 5 | 0 |
| TpMuguga_04g00228 | 6 | 0 |
| TpMuguga_04g00228 | 7 | 9 |
| TpMuguga_04g00228 | 8 | 0 |
| TpMuguga_04g00229 | 1 | 0 |
| TpMuguga_04g00229 | 2 | 0 |
| TpMuguga_04g00229 | 3 | 2.352941176 |
| TpMuguga_04g00229 | 4 | 1.411764706 |
| TpMuguga_04g00229 | 5 | 0.941176471 |
| TpMuguga_04g00229 | 6 | 4.705882353 |
| TpMuguga_04g00229 | 7 | 15.05882353 |
| TpMuguga_04g00231 | 1 | 0.14254386 |
| TpMuguga_04g00231 | 2 | 0 |
| TpMuguga_04g00231 | 3 | 0 |
| TpMuguga_04g00231 | 4 | 0 |
| TpMuguga_04g00231 | 5 | 0.313596491 |
| TpMuguga_04g00231 | 6 | 0 |
| TpMuguga_04g00231 | 7 | 0 |
| TpMuguga_04g00231 | 8 | 0 |
| TpMuguga_04g00231 | 9 | 0 |
| TpMuguga_04g00231 | 10 | 0 |
| TpMuguga_04g00231 | 11 | 0 |
| TpMuguga_04g00231 | 12 | 0 |
| TpMuguga_04g00230 | 1 | 1.578947368 |
| TpMuguga_04g00230 | 2 | 0.526315789 |
| TpMuguga_04g00230 | 3 | 0 |
| TpMuguga_04g00230 | 4 | 0 |
| TpMuguga_04g00232 | 1 | 0 |
| TpMuguga_04g00232 | 2 | 0 |
| TpMuguga_04g00232 | 3 | 0 |
| TpMuguga_04g00232 | 4 | 0 |
| TpMuguga_04g00234 | 1 | 0 |
| TpMuguga_04g00234 | 2 | 2.748538012 |
| TpMuguga_04g00235 | 1 | 0 |
| TpMuguga_04g00235 | 2 | 0 |
| TpMuguga_04g00235 | 3 | 0 |
| TpMuguga_04g00235 | 4 | 0 |
| TpMuguga_04g00235 | 5 | 0 |
| TpMuguga_04g00235 | 6 | 5.333333333 |
| TpMuguga_04g00235 | 7 | 0 |
| TpMuguga_04g00237 | 1 | 0 |

| | | |
|---|---|---|
| TpMuguga_04g00237 | 2 | 0 |
| TpMuguga_04g00237 | 3 | 0 |
| TpMuguga_04g00236 | 1 | 0.146143437 |
| TpMuguga_04g00236 | 2 | 0 |
| TpMuguga_04g00236 | 3 | 0.056833559 |
| TpMuguga_04g00236 | 4 | 0.01623816 |
| TpMuguga_04g00236 | 5 | 0.430311231 |
| TpMuguga_04g00238 | 1 | 0 |
| TpMuguga_04g00239 | 1 | 5.25 |
| TpMuguga_04g00239 | 2 | 0.375 |
| TpMuguga_04g00239 | 3 | 0.625 |
| TpMuguga_04g00239 | 4 | 0 |
| TpMuguga_04g00239 | 5 | 2.375 |
| TpMuguga_04g00239 | 6 | 0.75 |
| TpMuguga_04g00240 | 1 | 0 |
| TpMuguga_04g00240 | 2 | 0 |
| TpMuguga_04g00240 | 3 | 0 |
| TpMuguga_04g00240 | 4 | 0 |
| TpMuguga_04g00241 | 1 | 0 |
| TpMuguga_04g00242 | 1 | 0.288461538 |
| TpMuguga_04g00242 | 2 | 0 |
| TpMuguga_04g00242 | 3 | 0 |
| TpMuguga_04g00242 | 4 | 1.057692308 |
| TpMuguga_04g00243 | 1 | 0 |
| TpMuguga_04g00243 | 2 | 0 |
| TpMuguga_04g00243 | 3 | 2.285714286 |
| TpMuguga_04g00244 | 1 | 0.586956522 |
| TpMuguga_04g00244 | 2 | 0.065217391 |
| TpMuguga_04g00244 | 3 | 1.913043478 |
| TpMuguga_04g00244 | 4 | 0.565217391 |
| TpMuguga_04g00244 | 5 | 0.043478261 |
| TpMuguga_04g00248 | 1 | 0 |
| TpMuguga_04g00248 | 2 | 0 |
| TpMuguga_04g00248 | 3 | 0 |
| TpMuguga_04g00248 | 4 | 0 |
| TpMuguga_04g00248 | 5 | 0 |
| TpMuguga_04g00248 | 6 | 0 |
| TpMuguga_04g00248 | 7 | 0 |
| TpMuguga_04g00248 | 8 | 1.304347826 |
| TpMuguga_04g00248 | 9 | 0 |
| TpMuguga_04g00248 | 10 | 4.173913043 |
| TpMuguga_04g00248 | 11 | 6.260869565 |

| | | |
|---|---|---|
| TpMuguga_04g02165 | 1 | 0 |
| TpMuguga_04g02165 | 2 | 0 |
| TpMuguga_04g02165 | 3 | 0 |
| TpMuguga_04g02165 | 4 | 0 |
| TpMuguga_04g02165 | 5 | 0 |
| TpMuguga_04g02165 | 6 | 0 |
| TpMuguga_04g02470 | 1 | 0 |
| TpMuguga_04g02470 | 2 | 0 |
| TpMuguga_04g02465 | 1 | 0 |
| TpMuguga_04g02465 | 2 | 0 |
| TpMuguga_04g02465 | 3 | 0 |
| TpMuguga_04g02465 | 4 | 0 |
| TpMuguga_04g00254 | 1 | 5.333333333 |
| TpMuguga_04g00254 | 2 | 0 |
| TpMuguga_04g00254 | 3 | 0 |
| TpMuguga_04g00253 | 1 | 3.018867925 |
| TpMuguga_04g00253 | 2 | 0 |
| TpMuguga_04g00253 | 3 | 0 |
| TpMuguga_04g00257 | 1 | 0 |
| TpMuguga_04g00257 | 2 | 0 |
| TpMuguga_04g00257 | 3 | 0 |
| TpMuguga_04g00257 | 4 | 0.008856089 |
| TpMuguga_04g00257 | 5 | 0 |
| TpMuguga_04g00257 | 6 | 0 |
| TpMuguga_04g00257 | 7 | 1.966051661 |
| TpMuguga_04g00258 | 1 | 0.414058739 |
| TpMuguga_04g00258 | 2 | 0 |
| TpMuguga_04g00258 | 3 | 0.002888782 |
| TpMuguga_04g00259 | 1 | 0 |
| TpMuguga_04g00259 | 2 | 11 |
| TpMuguga_04g00259 | 3 | 0 |
| TpMuguga_04g00259 | 4 | 0 |
| TpMuguga_04g00259 | 5 | 0 |
| TpMuguga_04g00259 | 6 | 0 |
| TpMuguga_04g00259 | 7 | 0 |
| TpMuguga_04g00259 | 8 | 0 |
| TpMuguga_04g00259 | 9 | 0 |
| TpMuguga_04g00259 | 10 | 11 |
| TpMuguga_04g00260 | 1 | 1.363636364 |
| TpMuguga_04g00260 | 2 | 0 |
| TpMuguga_04g00260 | 3 | 0.545454545 |
| TpMuguga_04g00260 | 4 | 0 |

| | | |
|---|---|---|
| TpMuguga_04g00260 | 5 | 0.545454545 |
| TpMuguga_04g00261 | 1 | 0 |
| TpMuguga_04g00261 | 2 | 0 |
| TpMuguga_04g00261 | 3 | 0 |
| TpMuguga_04g00261 | 4 | 0 |
| TpMuguga_04g00261 | 5 | 0 |
| TpMuguga_04g00261 | 6 | 0 |
| TpMuguga_04g00261 | 7 | 0 |
| TpMuguga_04g00261 | 8 | 0 |
| TpMuguga_04g00261 | 9 | 0 |
| TpMuguga_04g00262 | 1 | 0 |
| TpMuguga_04g00263 | 1 | 0 |
| TpMuguga_04g00263 | 2 | 0 |
| TpMuguga_04g00264 | 1 | 0.227272727 |
| TpMuguga_04g00264 | 2 | 0 |
| TpMuguga_04g00264 | 3 | 0.757575758 |
| TpMuguga_04g00264 | 4 | 0.378787879 |
| TpMuguga_04g00265 | 1 | 0 |
| TpMuguga_04g00265 | 2 | 0.341463415 |
| TpMuguga_04g00265 | 3 | 0.081300813 |
| TpMuguga_04g00265 | 4 | 0 |
| TpMuguga_04g00265 | 5 | 0.162601626 |
| TpMuguga_04g00265 | 6 | 0.130081301 |
| TpMuguga_04g00265 | 7 | 1.447154472 |
| TpMuguga_04g00265 | 8 | 0 |
| TpMuguga_04g00265 | 9 | 0.829268293 |
| TpMuguga_04g00265 | 10 | 0.37398374 |
| TpMuguga_04g00265 | 11 | 0.081300813 |
| TpMuguga_04g00266 | 1 | 0 |
| TpMuguga_04g00267 | 1 | 0 |
| TpMuguga_04g00267 | 2 | 2.625 |
| TpMuguga_04g00267 | 3 | 0 |
| TpMuguga_04g00267 | 4 | 0 |
| TpMuguga_04g00267 | 5 | 0 |
| TpMuguga_04g00267 | 6 | 1.75 |
| TpMuguga_04g00268 | 1 | 13.75 |
| TpMuguga_04g00268 | 2 | 0 |
| TpMuguga_04g00268 | 3 | 0 |
| TpMuguga_04g00268 | 4 | 0 |
| TpMuguga_04g00268 | 5 | 0 |
| TpMuguga_04g00268 | 6 | 0 |
| TpMuguga_04g00268 | 7 | 0 |

| | | |
|---|---|---|
| TpMuguga_04g00268 | 8 | 0 |
| TpMuguga_04g00268 | 9 | 0 |
| TpMuguga_04g00268 | 10 | 0 |
| TpMuguga_04g00269 | 1 | 0.684179802 |
| TpMuguga_04g00269 | 2 | 0.721541156 |
| TpMuguga_04g00269 | 3 | 2.300058377 |
| TpMuguga_04g00271 | 1 | 0 |
| TpMuguga_04g00272 | 1 | 0 |
| TpMuguga_04g00272 | 2 | 0 |
| TpMuguga_04g00272 | 3 | 0 |
| TpMuguga_04g00272 | 4 | 0 |
| TpMuguga_04g00272 | 5 | 0 |
| TpMuguga_04g00272 | 6 | 0 |
| TpMuguga_04g00272 | 7 | 0 |
| TpMuguga_04g00272 | 8 | 0 |
| TpMuguga_04g00273 | 1 | 0 |
| TpMuguga_04g00276 | 1 | 0 |
| TpMuguga_04g00277 | 1 | 0 |
| TpMuguga_04g00277 | 2 | 0 |
| TpMuguga_04g00277 | 3 | 0 |
| TpMuguga_04g00278 | 1 | 0 |
| TpMuguga_04g00278 | 2 | 0 |
| TpMuguga_04g00278 | 3 | 0 |
| TpMuguga_04g00278 | 4 | 0 |
| TpMuguga_04g00279 | 1 | 1.791666667 |
| TpMuguga_04g00279 | 2 | 0.916666667 |
| TpMuguga_04g00279 | 3 | 1.791666667 |
| TpMuguga_04g00279 | 4 | 3.458333333 |
| TpMuguga_04g00280 | 1 | 0 |
| TpMuguga_04g00280 | 2 | 0.647058824 |
| TpMuguga_04g00280 | 3 | 0.566176471 |
| TpMuguga_04g00280 | 4 | 0.161764706 |
| TpMuguga_04g00280 | 5 | 0 |
| TpMuguga_04g00280 | 6 | 0 |
| TpMuguga_04g00280 | 7 | 0.485294118 |
| TpMuguga_04g00280 | 8 | 2.102941176 |
| TpMuguga_04g00280 | 9 | 1.051470588 |
| TpMuguga_04g00280 | 10 | 1.698529412 |
| TpMuguga_04g00281 | 1 | 0 |
| TpMuguga_04g00281 | 2 | 0 |
| TpMuguga_04g00281 | 3 | 0 |
| TpMuguga_04g00281 | 4 | 0 |

| | | |
|---|---|---|
| TpMuguga_04g00281 | 5 | 1.157894737 |
| TpMuguga_04g00281 | 6 | 0 |
| TpMuguga_04g00281 | 7 | 0 |
| TpMuguga_04g00281 | 8 | 0 |
| TpMuguga_04g00281 | 9 | 0 |
| TpMuguga_04g00281 | 10 | 3.473684211 |
| TpMuguga_04g00282 | 1 | 0 |
| TpMuguga_04g00283 | 1 | 3.694444444 |
| TpMuguga_04g00283 | 2 | 0 |
| TpMuguga_04g00283 | 3 | 0 |
| TpMuguga_04g00283 | 4 | 0.777777778 |
| TpMuguga_04g00283 | 5 | 2.916666667 |
| TpMuguga_04g00283 | 6 | 0 |
| TpMuguga_04g00284 | 1 | 3.402439024 |
| TpMuguga_04g00284 | 2 | 0.219512195 |
| TpMuguga_04g00284 | 3 | 0.109756098 |
| TpMuguga_04g00284 | 4 | 1.426829268 |
| TpMuguga_04g00284 | 5 | 0.987804878 |
| TpMuguga_04g00284 | 6 | 0 |
| TpMuguga_04g00284 | 7 | 0.43902439 |
| TpMuguga_04g00284 | 8 | 0.987804878 |
| TpMuguga_04g00285 | 1 | 13.14285714 |
| TpMuguga_04g00285 | 2 | 0 |
| TpMuguga_04g00285 | 3 | 0 |
| TpMuguga_04g00285 | 4 | 0 |
| TpMuguga_04g00285 | 5 | 0 |
| TpMuguga_04g00285 | 6 | 0 |
| TpMuguga_04g00285 | 7 | 0.571428571 |
| TpMuguga_04g00285 | 8 | 0 |
| TpMuguga_04g00285 | 9 | 0 |
| TpMuguga_04g00285 | 10 | 0 |
| TpMuguga_04g00285 | 11 | 0 |
| TpMuguga_04g00286 | 1 | 0 |
| TpMuguga_04g00286 | 2 | 0 |
| TpMuguga_04g00286 | 3 | 0 |
| TpMuguga_04g00286 | 4 | 0 |
| TpMuguga_04g00286 | 5 | 0 |
| TpMuguga_04g00286 | 6 | 0 |
| TpMuguga_04g00286 | 7 | 0 |
| TpMuguga_04g00287 | 1 | 0.358778626 |
| TpMuguga_04g00288 | 1 | 0 |
| TpMuguga_04g00288 | 2 | 0 |

| | | |
|---|---|---|
| TpMuguga_04g00288 | 3 | 1.411764706 |
| TpMuguga_04g00289 | 1 | 0 |
| TpMuguga_04g00289 | 2 | 0 |
| TpMuguga_04g00289 | 3 | 0 |
| TpMuguga_04g00289 | 4 | 0 |
| TpMuguga_04g00289 | 5 | 0 |
| TpMuguga_04g00289 | 6 | 0 |
| TpMuguga_04g00289 | 7 | 0 |
| TpMuguga_04g00290 | 1 | 0.005488474 |
| TpMuguga_04g00290 | 2 | 0 |
| TpMuguga_04g00290 | 3 | 0 |
| TpMuguga_04g00290 | 4 | 0.00034303 |
| TpMuguga_04g00291 | 1 | 0.827713816 |
| TpMuguga_04g00291 | 2 | 0 |
| TpMuguga_04g00293 | 1 | 0.197368421 |
| TpMuguga_04g00293 | 2 | 0.006578947 |
| TpMuguga_04g00292 | 1 | 0 |
| TpMuguga_04g00292 | 2 | 0 |
| TpMuguga_04g00292 | 3 | 0 |
| TpMuguga_04g00292 | 4 | 0 |
| TpMuguga_04g00292 | 5 | 0 |
| TpMuguga_04g00292 | 6 | 0 |
| TpMuguga_04g00292 | 7 | 0.64957265 |
| TpMuguga_04g00295 | 1 | 0 |
| TpMuguga_04g00295 | 2 | 0 |
| TpMuguga_04g00295 | 3 | 0 |
| TpMuguga_04g00296 | 1 | 0 |
| TpMuguga_04g00296 | 2 | 24 |
| TpMuguga_04g00296 | 3 | 0 |
| TpMuguga_04g00298 | 1 | 0 |
| TpMuguga_04g00298 | 2 | 0 |
| TpMuguga_04g00299 | 1 | 4.49382716 |
| TpMuguga_04g00299 | 2 | 7 |
| TpMuguga_04g00299 | 3 | 2.24691358 |
| TpMuguga_04g00299 | 4 | 1.814814815 |
| TpMuguga_04g00299 | 5 | 3.543209877 |
| TpMuguga_04g00299 | 6 | 0 |
| TpMuguga_04g00301 | 1 | 0 |
| TpMuguga_04g00301 | 2 | 0 |
| TpMuguga_04g00301 | 3 | 0 |
| TpMuguga_04g00302 | 1 | 0.285714286 |
| TpMuguga_04g00302 | 2 | 0 |

| | | |
|---|---|---|
| TpMuguga_04g00302 | 3 | 0 |
| TpMuguga_04g00302 | 4 | 0 |
| TpMuguga_04g00303 | 1 | 0 |
| TpMuguga_04g00303 | 2 | 0 |
| TpMuguga_04g00303 | 3 | 0 |
| TpMuguga_04g00304 | 1 | 0.145631068 |
| TpMuguga_04g00304 | 2 | 0 |
| TpMuguga_04g00305 | 1 | 0.301204819 |
| TpMuguga_04g00305 | 2 | 0 |
| TpMuguga_04g00305 | 3 | 0 |
| TpMuguga_04g00305 | 4 | 0.542168675 |
| TpMuguga_04g02550 | 1 | 0 |
| TpMuguga_04g02550 | 2 | 4 |
| TpMuguga_04g00307 | 1 | 0 |
| TpMuguga_04g00307 | 2 | 0 |
| TpMuguga_04g00308 | 1 | 0 |
| TpMuguga_04g00308 | 2 | 0 |
| TpMuguga_04g00309 | 1 | 0.096774194 |
| TpMuguga_04g00309 | 2 | 1.741935484 |
| TpMuguga_04g02205 | 1 | 0 |
| TpMuguga_04g02205 | 2 | 0 |
| TpMuguga_04g02205 | 3 | 0 |
| TpMuguga_04g02205 | 4 | 0 |
| TpMuguga_04g02205 | 5 | 0 |
| TpMuguga_04g02205 | 6 | 0 |
| TpMuguga_04g02205 | 7 | 0 |
| TpMuguga_04g02205 | 8 | 0 |
| TpMuguga_04g02205 | 9 | 0 |
| TpMuguga_04g02205 | 10 | 0 |
| TpMuguga_04g02205 | 11 | 0 |
| TpMuguga_04g02205 | 12 | 0 |
| TpMuguga_04g02205 | 13 | 0 |
| TpMuguga_04g00313 | 1 | 0 |
| TpMuguga_04g00313 | 2 | 0 |
| TpMuguga_04g00315 | 1 | 0 |
| TpMuguga_04g00315 | 2 | 0 |
| TpMuguga_04g00315 | 3 | 0 |
| TpMuguga_04g00315 | 4 | 0 |
| TpMuguga_04g00315 | 5 | 7 |
| TpMuguga_04g00315 | 6 | 0 |
| TpMuguga_04g00317 | 1 | 3.333333333 |
| TpMuguga_04g00317 | 2 | 0 |

| | | |
|---|---|---|
| TpMuguga_04g00317 | 3 | 0.833333333 |
| TpMuguga_04g00317 | 4 | 0 |
| TpMuguga_04g02295 | 1 | 0 |
| TpMuguga_04g02295 | 2 | 0 |
| TpMuguga_04g02295 | 3 | 0 |
| TpMuguga_04g02295 | 4 | 2.459459459 |
| TpMuguga_04g02295 | 5 | 0 |
| TpMuguga_04g02295 | 6 | 5.486486486 |
| TpMuguga_04g00320 | 1 | 0.856067733 |
| TpMuguga_04g00322 | 1 | 0 |
| TpMuguga_04g00322 | 2 | 0 |
| TpMuguga_04g00323 | 1 | 0 |
| TpMuguga_04g00323 | 2 | 0 |
| TpMuguga_04g00323 | 3 | 0 |
| TpMuguga_04g00323 | 4 | 0 |
| TpMuguga_04g00323 | 5 | 0 |
| TpMuguga_04g00323 | 6 | 0 |
| TpMuguga_04g00323 | 7 | 0 |
| TpMuguga_04g00323 | 8 | 0 |
| TpMuguga_04g00323 | 9 | 0 |
| TpMuguga_04g00323 | 10 | 0 |
| TpMuguga_04g00324 | 1 | 0 |
| TpMuguga_04g00324 | 2 | 0 |
| TpMuguga_04g00324 | 3 | 0 |
| TpMuguga_04g00324 | 4 | 0 |
| TpMuguga_04g00324 | 5 | 0 |
| TpMuguga_04g00324 | 6 | 0 |
| TpMuguga_04g00325 | 1 | 0.531914894 |
| TpMuguga_04g00325 | 2 | 2.446808511 |
| TpMuguga_04g00325 | 3 | 0.319148936 |
| TpMuguga_04g00325 | 4 | 1.276595745 |
| TpMuguga_04g00326 | 1 | 0.03591954 |
| TpMuguga_04g00326 | 2 | 0.014367816 |
| TpMuguga_04g00326 | 3 | 0 |
| TpMuguga_04g00326 | 4 | 0 |
| TpMuguga_04g00327 | 1 | 0 |
| TpMuguga_04g00327 | 2 | 0 |
| TpMuguga_04g00327 | 3 | 4 |
| TpMuguga_04g00329 | 1 | 0 |
| TpMuguga_04g00329 | 2 | 0 |
| TpMuguga_04g00329 | 3 | 0 |
| TpMuguga_04g00329 | 4 | 0 |

| | | |
|---|---|---|
| TpMuguga_04g00329 | 5 | 0.540540541 |
| TpMuguga_04g00329 | 6 | 0.108108108 |
| TpMuguga_04g00329 | 7 | 0.864864865 |
| TpMuguga_04g02195 | 1 | 0 |
| TpMuguga_04g02195 | 2 | 0 |
| TpMuguga_04g02715 | 1 | 0 |
| TpMuguga_04g02715 | 2 | 0 |
| TpMuguga_04g02715 | 3 | 0 |
| TpMuguga_04g02715 | 4 | 0 |
| TpMuguga_04g02080 | 1 | 3.2 |
| TpMuguga_04g02080 | 2 | 2.273684211 |
| TpMuguga_04g02080 | 3 | 0.084210526 |
| TpMuguga_04g02080 | 4 | 0.421052632 |
| TpMuguga_04g02080 | 5 | 0 |
| TpMuguga_04g02080 | 6 | 0.084210526 |
| TpMuguga_04g02080 | 7 | 0 |
| TpMuguga_04g00334 | 1 | 0 |
| TpMuguga_04g00334 | 2 | 10 |
| TpMuguga_04g00334 | 3 | 0 |
| TpMuguga_04g00334 | 4 | 0 |
| TpMuguga_04g00334 | 5 | 0 |
| TpMuguga_04g00334 | 6 | 0 |
| TpMuguga_04g00334 | 7 | 0 |
| TpMuguga_04g00334 | 8 | 0 |
| TpMuguga_04g00334 | 9 | 0 |
| TpMuguga_04g02045 | 1 | 0 |
| TpMuguga_04g02045 | 2 | 0 |
| TpMuguga_04g02045 | 3 | 0 |
| TpMuguga_04g02045 | 4 | 0 |
| TpMuguga_04g02045 | 5 | 0 |
| TpMuguga_04g02240 | 1 | 0 |
| TpMuguga_04g02240 | 2 | 0 |
| TpMuguga_04g02240 | 3 | 0 |
| TpMuguga_04g00337 | 1 | 0 |
| TpMuguga_04g00337 | 2 | 0 |
| TpMuguga_04g00337 | 3 | 0 |
| TpMuguga_04g00337 | 4 | 0 |
| TpMuguga_04g00337 | 5 | 0 |
| TpMuguga_04g00337 | 6 | 0 |
| TpMuguga_04g00337 | 7 | 0 |
| TpMuguga_04g00337 | 8 | 3.09375 |
| TpMuguga_04g00337 | 9 | 0 |

| | | |
|---|---|---|
| TpMuguga_04g00337 | 10 | 0.34375 |
| TpMuguga_04g00338 | 1 | 0 |
| TpMuguga_04g00338 | 2 | 0 |
| TpMuguga_04g00338 | 3 | 27 |
| TpMuguga_04g00338 | 4 | 0 |
| TpMuguga_04g00338 | 5 | 0 |
| TpMuguga_04g00339 | 1 | 0 |
| TpMuguga_04g00340 | 1 | 0 |
| TpMuguga_04g00340 | 2 | 0 |
| TpMuguga_04g00340 | 3 | 0.039553753 |
| TpMuguga_04g00340 | 4 | 0 |
| TpMuguga_04g00340 | 5 | 0 |
| TpMuguga_04g00340 | 6 | 0 |
| TpMuguga_04g00340 | 7 | 0 |
| TpMuguga_04g00340 | 8 | 0 |
| TpMuguga_04g00340 | 9 | 0 |
| TpMuguga_04g00340 | 10 | 0 |
| TpMuguga_04g00340 | 11 | 0 |
| TpMuguga_04g00340 | 12 | 0 |
| TpMuguga_04g00342 | 1 | 0 |
| TpMuguga_04g00342 | 2 | 0 |
| TpMuguga_04g00342 | 3 | 0 |
| TpMuguga_04g00342 | 4 | 0 |
| TpMuguga_04g00342 | 5 | 0 |
| TpMuguga_04g00342 | 6 | 0 |
| TpMuguga_04g00343 | 1 | 0 |
| TpMuguga_04g00345 | 1 | 0 |
| TpMuguga_04g00345 | 2 | 0 |
| TpMuguga_04g00345 | 3 | 2.2 |
| TpMuguga_04g00345 | 4 | 13.2 |
| TpMuguga_04g00345 | 5 | 0 |
| TpMuguga_04g00345 | 6 | 0 |
| TpMuguga_04g00345 | 7 | 0 |
| TpMuguga_04g00345 | 8 | 0 |
| TpMuguga_04g00345 | 9 | 0 |
| TpMuguga_04g00345 | 10 | 0 |
| TpMuguga_04g00347 | 1 | 2.678571429 |
| TpMuguga_04g00347 | 2 | 0.214285714 |
| TpMuguga_04g00348 | 1 | 0.507936508 |
| TpMuguga_04g00348 | 2 | 0 |
| TpMuguga_04g00348 | 3 | 1.015873016 |
| TpMuguga_04g00348 | 4 | 0 |

| | | |
|---|---|---|
| TpMuguga_04g00348 | 5 | 0.126984127 |
| TpMuguga_04g00348 | 6 | 5.333333333 |
| TpMuguga_04g00348 | 7 | 0 |
| TpMuguga_04g00349 | 1 | 0 |
| TpMuguga_04g00349 | 2 | 0.003237992 |
| TpMuguga_04g00349 | 3 | 0 |
| TpMuguga_04g00349 | 4 | 0 |
| TpMuguga_04g00349 | 5 | 0.093901781 |
| TpMuguga_04g00350 | 1 | 0 |
| TpMuguga_04g00350 | 2 | 0 |
| TpMuguga_04g00350 | 3 | 0 |
| TpMuguga_04g00351 | 1 | 0 |
| TpMuguga_04g00351 | 2 | 0 |
| TpMuguga_04g00351 | 3 | 0 |
| TpMuguga_04g00351 | 4 | 0 |
| TpMuguga_04g00351 | 5 | 0 |
| TpMuguga_04g02060 | 1 | 0 |
| TpMuguga_04g02060 | 2 | 0 |
| TpMuguga_04g02060 | 3 | 0 |
| TpMuguga_04g02060 | 4 | 0 |
| TpMuguga_04g00354 | 1 | 0.426666667 |
| TpMuguga_04g00357 | 1 | 0 |
| TpMuguga_04g00357 | 2 | 0 |
| TpMuguga_04g00358 | 1 | 1.339534884 |
| TpMuguga_04g00358 | 2 | 0.711627907 |
| TpMuguga_04g00361 | 1 | 0 |
| TpMuguga_04g00361 | 2 | 17.5 |
| TpMuguga_04g00361 | 3 | 3.5 |
| TpMuguga_04g00361 | 4 | 17.5 |
| TpMuguga_04g00361 | 5 | 0 |
| TpMuguga_04g00361 | 6 | 0 |
| TpMuguga_04g00360 | 1 | 0 |
| TpMuguga_04g00360 | 2 | 0 |
| TpMuguga_04g00360 | 3 | 0.65625 |
| TpMuguga_04g00360 | 4 | 0 |
| TpMuguga_04g00360 | 5 | 0 |
| TpMuguga_04g00360 | 6 | 0 |
| TpMuguga_04g00363 | 1 | 0 |
| TpMuguga_04g00363 | 2 | 0 |
| TpMuguga_04g00363 | 3 | 0 |
| TpMuguga_04g00364 | 1 | 0 |
| TpMuguga_04g00364 | 2 | 0 |

| | | |
|---|---|---|
| TpMuguga_04g00364 | 3 | 10 |
| TpMuguga_04g00364 | 4 | 0 |
| TpMuguga_04g00364 | 5 | 0 |
| TpMuguga_04g00364 | 6 | 0 |
| TpMuguga_04g00364 | 7 | 0 |
| TpMuguga_04g00364 | 8 | 0 |
| TpMuguga_04g00364 | 9 | 0 |
| TpMuguga_04g00365 | 1 | 0 |
| TpMuguga_04g00366 | 1 | 0 |
| TpMuguga_04g00366 | 2 | 0 |
| TpMuguga_04g00366 | 3 | 0 |
| TpMuguga_04g00366 | 4 | 0 |
| TpMuguga_04g00366 | 5 | 2.25 |
| TpMuguga_04g00367 | 1 | 8 |
| TpMuguga_04g00367 | 2 | 0 |
| TpMuguga_04g00367 | 3 | 0 |
| TpMuguga_04g00367 | 4 | 0 |
| TpMuguga_04g00367 | 5 | 0 |
| TpMuguga_04g00367 | 6 | 0 |
| TpMuguga_04g00367 | 7 | 0 |
| TpMuguga_04g00368 | 1 | 4.375 |
| TpMuguga_04g00368 | 2 | 0 |
| TpMuguga_04g00368 | 3 | 0 |
| TpMuguga_04g00368 | 4 | 0 |
| TpMuguga_04g00369 | 1 | 0 |
| TpMuguga_04g00369 | 2 | 0 |
| TpMuguga_04g00371 | 1 | 0 |
| TpMuguga_04g00371 | 2 | 0 |
| TpMuguga_04g00372 | 1 | 0 |
| TpMuguga_04g00373 | 1 | 0 |
| TpMuguga_04g00373 | 2 | 1.125 |
| TpMuguga_04g00374 | 1 | 1 |
| TpMuguga_04g00374 | 2 | 0 |
| TpMuguga_04g00374 | 3 | 0 |
| TpMuguga_04g00374 | 4 | 0.333333333 |
| TpMuguga_04g00374 | 5 | 0.666666667 |
| TpMuguga_04g00374 | 6 | 0 |
| TpMuguga_04g00374 | 7 | 2 |
| TpMuguga_04g00374 | 8 | 2 |
| TpMuguga_04g02675 | 1 | 0 |
| TpMuguga_04g02675 | 2 | 0 |
| TpMuguga_04g02675 | 3 | 0 |

| | | |
|---|---|---|
| TpMuguga_04g02675 | 4 | 0 |
| TpMuguga_04g02675 | 5 | 0 |
| TpMuguga_04g02675 | 6 | 0 |
| TpMuguga_04g02675 | 7 | 0 |
| TpMuguga_04g02675 | 8 | 0 |
| TpMuguga_04g00379 | 1 | 0.652173913 |
| TpMuguga_04g00379 | 2 | 0.652173913 |
| TpMuguga_04g00380 | 1 | 0 |
| TpMuguga_04g00381 | 1 | 0 |
| TpMuguga_04g00381 | 2 | 0 |
| TpMuguga_04g00381 | 3 | 0 |
| TpMuguga_04g00381 | 4 | 0.033238367 |
| TpMuguga_04g00384 | 1 | 0 |
| TpMuguga_04g00386 | 1 | 0 |
| TpMuguga_04g00386 | 2 | 0.137142857 |
| TpMuguga_04g00386 | 3 | 0 |
| TpMuguga_04g00386 | 4 | 0 |
| TpMuguga_04g00386 | 5 | 0 |
| TpMuguga_04g00388 | 1 | 0 |
| TpMuguga_04g00388 | 2 | 9 |
| TpMuguga_04g00388 | 3 | 0 |
| TpMuguga_04g00387 | 1 | 0 |
| TpMuguga_04g00387 | 2 | 0 |
| TpMuguga_04g00387 | 3 | 0 |
| TpMuguga_04g00387 | 4 | 0 |
| TpMuguga_04g00389 | 1 | 0 |
| TpMuguga_04g00389 | 2 | 0 |
| TpMuguga_04g00389 | 3 | 0 |
| TpMuguga_04g00389 | 4 | 0 |
| TpMuguga_04g00389 | 5 | 0 |
| TpMuguga_04g00389 | 6 | 0 |
| TpMuguga_04g00389 | 7 | 0 |
| TpMuguga_04g00389 | 8 | 0 |
| TpMuguga_04g00390 | 1 | 0.769230769 |
| TpMuguga_04g00390 | 2 | 9.615384615 |
| TpMuguga_04g00390 | 3 | 0 |
| TpMuguga_04g00390 | 4 | 0 |
| TpMuguga_04g00393 | 1 | 0 |
| TpMuguga_04g00394 | 1 | 0.290322581 |
| TpMuguga_04g00394 | 2 | 0 |
| TpMuguga_04g00395 | 1 | 0 |
| TpMuguga_04g00395 | 2 | 0 |

| | | |
|---|---|---|
| TpMuguga_04g00395 | 3 | 0 |
| TpMuguga_04g00395 | 4 | 0 |
| TpMuguga_04g00395 | 5 | 0 |
| TpMuguga_04g00395 | 6 | 0 |
| TpMuguga_04g00395 | 7 | 0 |
| TpMuguga_04g00395 | 8 | 0 |
| TpMuguga_04g00395 | 9 | 0 |
| TpMuguga_04g00395 | 10 | 0 |
| TpMuguga_04g00396 | 1 | 1.333333333 |
| TpMuguga_04g00396 | 2 | 0 |
| TpMuguga_04g00396 | 3 | 0 |
| TpMuguga_04g00397 | 1 | 0 |
| TpMuguga_04g00397 | 2 | 0 |
| TpMuguga_04g00398 | 1 | 0 |
| TpMuguga_04g00398 | 2 | 0 |
| TpMuguga_04g00398 | 3 | 0 |
| TpMuguga_04g00398 | 4 | 0 |
| TpMuguga_04g00399 | 1 | 0.409090909 |
| TpMuguga_04g00399 | 2 | 0.409090909 |
| TpMuguga_04g00401 | 1 | 6.909090909 |
| TpMuguga_04g00401 | 2 | 3.454545455 |
| TpMuguga_04g00401 | 3 | 0 |
| TpMuguga_04g00401 | 4 | 0.545454545 |
| TpMuguga_04g00401 | 5 | 1.090909091 |
| TpMuguga_04g00401 | 6 | 1.272727273 |
| TpMuguga_04g00401 | 7 | 0 |
| TpMuguga_04g00402 | 1 | 0.039473684 |
| TpMuguga_04g00402 | 2 | 0 |
| TpMuguga_04g00402 | 3 | 0 |
| TpMuguga_04g00402 | 4 | 0 |
| TpMuguga_04g00402 | 5 | 0 |
| TpMuguga_04g00402 | 6 | 0 |
| TpMuguga_04g00402 | 7 | 0.947368421 |
| TpMuguga_04g00402 | 8 | 0.710526316 |
| TpMuguga_04g00405 | 1 | 1.75 |
| TpMuguga_04g00410 | 1 | 0 |
| TpMuguga_04g00410 | 2 | 0 |
| TpMuguga_04g00410 | 3 | 0 |
| TpMuguga_04g00410 | 4 | 0 |
| TpMuguga_04g00410 | 5 | 0 |
| TpMuguga_04g00411 | 1 | 16 |
| TpMuguga_04g00411 | 2 | 0 |

| | | |
|---|---|---|
| TpMuguga_04g00411 | 3 | 0 |
| TpMuguga_04g00412 | 1 | 0 |
| TpMuguga_04g00412 | 2 | 0 |
| TpMuguga_04g00412 | 3 | 0 |
| TpMuguga_04g00412 | 4 | 0 |
| TpMuguga_04g00413 | 1 | 0 |
| TpMuguga_04g00415 | 1 | 0 |
| TpMuguga_04g00415 | 2 | 0.130434783 |
| TpMuguga_04g00416 | 1 | 0 |
| TpMuguga_04g00416 | 2 | 0 |
| TpMuguga_04g00416 | 3 | 0.094594595 |
| TpMuguga_04g00416 | 4 | 0.094594595 |
| TpMuguga_04g00416 | 5 | 0 |
| TpMuguga_04g00416 | 6 | 0 |
| TpMuguga_04g00416 | 7 | 0 |
| TpMuguga_04g00416 | 8 | 0 |
| TpMuguga_04g00416 | 9 | 0 |
| TpMuguga_04g00416 | 10 | 0 |
| TpMuguga_04g00416 | 11 | 0 |
| TpMuguga_04g00416 | 12 | 0 |
| TpMuguga_04g00416 | 13 | 0 |
| TpMuguga_04g02055 | 1 | 1.5 |
| TpMuguga_04g02535 | 1 | 2.375 |
| TpMuguga_04g02535 | 2 | 0 |
| TpMuguga_04g02535 | 3 | 0 |
| TpMuguga_04g02535 | 4 | 0 |
| TpMuguga_04g02535 | 5 | 1.1875 |
| TpMuguga_04g02535 | 6 | 0 |
| TpMuguga_04g02535 | 7 | 0 |
| TpMuguga_04g02535 | 8 | 0 |
| TpMuguga_04g02535 | 9 | 0 |
| TpMuguga_04g02535 | 10 | 0 |
| TpMuguga_04g02535 | 11 | 2.375 |
| TpMuguga_04g02535 | 12 | 8.3125 |
| TpMuguga_04g02535 | 13 | 0 |
| TpMuguga_04g02535 | 14 | 0 |
| TpMuguga_04g02535 | 15 | 2.375 |
| TpMuguga_04g02535 | 16 | 0 |
| TpMuguga_04g02535 | 17 | 0 |
| TpMuguga_04g02535 | 18 | 0 |
| TpMuguga_04g00420 | 1 | 0 |
| TpMuguga_04g00420 | 2 | 0 |

| | | |
|---|---|---|
| TpMuguga_04g00420 | 3 | 0 |
| TpMuguga_04g00420 | 4 | 0 |
| TpMuguga_04g00420 | 5 | 0 |
| TpMuguga_04g00420 | 6 | 0 |
| TpMuguga_04g00420 | 7 | 0 |
| TpMuguga_04g00421 | 1 | 0 |
| TpMuguga_04g00422 | 1 | 0 |
| TpMuguga_04g00422 | 2 | 0 |
| TpMuguga_04g00422 | 3 | 0 |
| TpMuguga_04g00422 | 4 | 0 |
| TpMuguga_04g00422 | 5 | 0 |
| TpMuguga_04g00422 | 6 | 0 |
| TpMuguga_04g00422 | 7 | 0 |
| TpMuguga_04g00422 | 8 | 0 |
| TpMuguga_04g00422 | 9 | 0 |
| TpMuguga_04g00422 | 10 | 3.5 |
| TpMuguga_04g00423 | 1 | 0 |
| TpMuguga_04g00423 | 2 | 0.400911162 |
| TpMuguga_04g00423 | 3 | 0.036446469 |
| TpMuguga_04g00423 | 4 | 0 |
| TpMuguga_04g00423 | 5 | 0 |
| TpMuguga_04g00423 | 6 | 0.036446469 |
| TpMuguga_04g00423 | 7 | 0 |
| TpMuguga_04g00423 | 8 | 0.072892938 |
| TpMuguga_04g00423 | 9 | 0.218678815 |
| TpMuguga_04g00423 | 10 | 0 |
| TpMuguga_04g00423 | 11 | 0 |
| TpMuguga_04g00423 | 12 | 0.036446469 |
| TpMuguga_04g00423 | 13 | 1.603644647 |
| TpMuguga_04g00423 | 14 | 10.46013667 |
| TpMuguga_04g00423 | 15 | 6.815489749 |
| TpMuguga_04g00424 | 1 | 0 |
| TpMuguga_04g00424 | 2 | 0 |
| TpMuguga_04g00424 | 3 | 0 |
| TpMuguga_04g00424 | 4 | 4.666666667 |
| TpMuguga_04g00424 | 5 | 0 |
| TpMuguga_04g00424 | 6 | 0 |
| TpMuguga_04g00424 | 7 | 0 |
| TpMuguga_04g00424 | 8 | 0 |
| TpMuguga_04g00424 | 9 | 9.333333333 |
| TpMuguga_04g00424 | 10 | 0 |
| TpMuguga_04g00424 | 11 | 0 |

| | | |
|---|---|---|
| TpMuguga_04g00424 | 12 | 14 |
| TpMuguga_04g00424 | 13 | 0 |
| TpMuguga_04g00429 | 1 | 4.663299663 |
| TpMuguga_04g00429 | 2 | 0.084175084 |
| TpMuguga_04g00429 | 3 | 0 |
| TpMuguga_04g00429 | 4 | 0 |
| TpMuguga_04g00428 | 1 | 0 |
| TpMuguga_04g00430 | 1 | 0 |
| TpMuguga_04g00430 | 2 | 0 |
| TpMuguga_04g00430 | 3 | 6 |
| TpMuguga_04g00430 | 4 | 0 |
| TpMuguga_04g00430 | 5 | 9 |
| TpMuguga_04g00431 | 1 | 0 |
| TpMuguga_04g00431 | 2 | 0 |
| TpMuguga_04g00431 | 3 | 0 |
| TpMuguga_04g00431 | 4 | 6 |
| TpMuguga_04g00431 | 5 | 3.6 |
| TpMuguga_04g00432 | 1 | 0 |
| TpMuguga_04g00432 | 2 | 0 |
| TpMuguga_04g00432 | 3 | 0 |
| TpMuguga_04g00432 | 4 | 0 |
| TpMuguga_04g00433 | 1 | 0 |
| TpMuguga_04g00433 | 2 | 0 |
| TpMuguga_04g00434 | 1 | 0 |
| TpMuguga_04g00434 | 2 | 0 |
| TpMuguga_04g00434 | 3 | 0 |
| TpMuguga_04g00434 | 4 | 0 |
| TpMuguga_04g00434 | 5 | 0 |
| TpMuguga_04g00434 | 6 | 7 |
| TpMuguga_04g00435 | 1 | 0 |
| TpMuguga_04g00435 | 2 | 0 |
| TpMuguga_04g00436 | 1 | 0 |
| TpMuguga_04g00436 | 2 | 0 |
| TpMuguga_04g00436 | 3 | 0 |
| TpMuguga_04g00436 | 4 | 0 |
| TpMuguga_04g00439 | 1 | 0 |
| TpMuguga_04g00439 | 2 | 0 |
| TpMuguga_04g00439 | 3 | 0 |
| TpMuguga_04g00439 | 4 | 0 |
| TpMuguga_04g00439 | 5 | 0 |
| TpMuguga_04g00439 | 6 | 0.014748133 |
| TpMuguga_04g00438 | 1 | 4 |

| | | |
|---|---|---|
| TpMuguga_04g00438 | 2 | 0 |
| TpMuguga_04g00438 | 3 | 0 |
| TpMuguga_04g00438 | 4 | 0 |
| TpMuguga_04g00438 | 5 | 0 |
| TpMuguga_04g00438 | 6 | 0 |
| TpMuguga_04g00438 | 7 | 0 |
| TpMuguga_04g00438 | 8 | 0 |
| TpMuguga_04g00438 | 9 | 0 |
| TpMuguga_04g00438 | 10 | 12 |
| TpMuguga_04g00438 | 11 | 6.666666667 |
| TpMuguga_04g00442 | 1 | 0.627906977 |
| TpMuguga_04g00442 | 2 | 0.069767442 |
| TpMuguga_04g00443 | 1 | 0 |
| TpMuguga_04g00443 | 2 | 0 |
| TpMuguga_04g00443 | 3 | 0 |
| TpMuguga_04g00443 | 4 | 0 |
| TpMuguga_04g00443 | 5 | 0 |
| TpMuguga_04g00443 | 6 | 0.549618321 |
| TpMuguga_04g00444 | 1 | 7.789473684 |
| TpMuguga_04g00444 | 2 | 0.571428571 |
| TpMuguga_04g00444 | 3 | 0 |
| TpMuguga_04g00444 | 4 | 0.070175439 |
| TpMuguga_04g00444 | 5 | 0 |
| TpMuguga_04g00444 | 6 | 0 |
| TpMuguga_04g00444 | 7 | 0 |
| TpMuguga_04g00445 | 1 | 0 |
| TpMuguga_04g02575 | 1 | 0.069444444 |
| TpMuguga_04g02575 | 2 | 0.138888889 |
| TpMuguga_04g00448 | 1 | 0.285714286 |
| TpMuguga_04g00448 | 2 | 0 |
| TpMuguga_04g00448 | 3 | 0 |
| TpMuguga_04g00448 | 4 | 0 |
| TpMuguga_04g00448 | 5 | 0 |
| TpMuguga_04g00448 | 6 | 0 |
| TpMuguga_04g00448 | 7 | 0 |
| TpMuguga_04g00449 | 1 | 0 |
| TpMuguga_04g00449 | 2 | 0 |
| TpMuguga_04g00450 | 1 | 0 |
| TpMuguga_04g00450 | 2 | 0 |
| TpMuguga_04g00450 | 3 | 0 |
| TpMuguga_04g00450 | 4 | 0 |
| TpMuguga_04g00450 | 5 | 0 |

| | | |
|---|---|---|
| TpMuguga_04g00450 | 6 | 0.205882353 |
| TpMuguga_04g02505 | 1 | 0 |
| TpMuguga_04g02505 | 2 | 0 |
| TpMuguga_04g02510 | 1 | 0 |
| TpMuguga_04g02510 | 2 | 0 |
| TpMuguga_04g02510 | 3 | 1.390243902 |
| TpMuguga_04g02510 | 4 | 0 |
| TpMuguga_04g02510 | 5 | 4.682926829 |
| TpMuguga_04g00454 | 1 | 0 |
| TpMuguga_04g00454 | 2 | 0 |
| TpMuguga_04g00454 | 3 | 0 |
| TpMuguga_04g00454 | 4 | 0 |
| TpMuguga_04g00454 | 5 | 0 |
| TpMuguga_04g00454 | 6 | 0 |
| TpMuguga_04g00454 | 7 | 0 |
| TpMuguga_04g00454 | 8 | 0 |
| TpMuguga_04g00454 | 9 | 0 |
| TpMuguga_04g00453 | 1 | 0 |
| TpMuguga_04g00453 | 2 | 0 |
| TpMuguga_04g00453 | 3 | 0 |
| TpMuguga_04g00453 | 4 | 0 |
| TpMuguga_04g00453 | 5 | 0 |
| TpMuguga_04g00453 | 6 | 0 |
| TpMuguga_04g00453 | 7 | 0 |
| TpMuguga_04g00453 | 8 | 27 |
| TpMuguga_04g00455 | 1 | 0.048780488 |
| TpMuguga_04g00456 | 1 | 0 |
| TpMuguga_04g00456 | 2 | 1.285714286 |
| TpMuguga_04g00457 | 1 | 2 |
| TpMuguga_04g00457 | 2 | 0 |
| TpMuguga_04g00457 | 3 | 0 |
| TpMuguga_04g00458 | 1 | 0 |
| TpMuguga_04g00459 | 1 | 0 |
| TpMuguga_04g00459 | 2 | 0 |
| TpMuguga_04g00459 | 3 | 3.230769231 |
| TpMuguga_04g00459 | 4 | 3.692307692 |
| TpMuguga_04g00459 | 5 | 0 |
| TpMuguga_04g00460 | 1 | 0.248618785 |
| TpMuguga_04g00460 | 2 | 0.016574586 |
| TpMuguga_04g00460 | 3 | 0.182320442 |
| TpMuguga_04g00460 | 4 | 0.745856354 |
| TpMuguga_04g00460 | 5 | 1.906077348 |

| | | |
|---|---|---|
| TpMuguga_04g02440 | 1 | 0 |
| TpMuguga_04g02445 | 1 | 2 |
| TpMuguga_04g00462 | 1 | 0 |
| TpMuguga_04g00462 | 2 | 0 |
| TpMuguga_04g00462 | 3 | 0 |
| TpMuguga_04g00462 | 4 | 0 |
| TpMuguga_04g00462 | 5 | 2.129032258 |
| TpMuguga_04g00463 | 1 | 0 |
| TpMuguga_04g00463 | 2 | 0 |
| TpMuguga_04g00463 | 3 | 0 |
| TpMuguga_04g00463 | 4 | 0 |
| TpMuguga_04g00463 | 5 | 0 |
| TpMuguga_04g00463 | 6 | 0 |
| TpMuguga_04g00463 | 7 | 0 |
| TpMuguga_04g00463 | 8 | 0 |
| TpMuguga_04g00463 | 9 | 0 |
| TpMuguga_04g00463 | 10 | 0 |
| TpMuguga_04g00463 | 11 | 0 |
| TpMuguga_04g00463 | 12 | 0 |
| TpMuguga_04g00463 | 13 | 0 |
| TpMuguga_04g00463 | 14 | 0 |
| TpMuguga_04g00463 | 15 | 0 |
| TpMuguga_04g00464 | 1 | 0.807692308 |
| TpMuguga_04g00464 | 2 | 1.096153846 |
| TpMuguga_04g00465 | 1 | 0 |
| TpMuguga_04g00466 | 1 | 0 |
| TpMuguga_04g02580 | 1 | 5 |
| TpMuguga_04g02580 | 2 | 2.222222222 |
| TpMuguga_04g02580 | 3 | 1.666666667 |
| TpMuguga_04g02580 | 4 | 1.111111111 |
| TpMuguga_04g02580 | 5 | 0.555555556 |
| TpMuguga_04g02580 | 6 | 0.555555556 |
| TpMuguga_04g02580 | 7 | 0 |
| TpMuguga_04g02580 | 8 | 0 |
| TpMuguga_04g02580 | 9 | 2.222222222 |
| TpMuguga_04g00469 | 1 | 1.8 |
| TpMuguga_04g00469 | 2 | 12 |
| TpMuguga_04g00470 | 1 | 0 |
| TpMuguga_04g00471 | 1 | 0 |
| TpMuguga_04g00472 | 1 | 0.138888889 |
| TpMuguga_04g00472 | 2 | 0.416666667 |
| TpMuguga_04g00472 | 3 | 0 |

| | | |
|---|---|---|
| TpMuguga_04g00472 | 4 | 0 |
| TpMuguga_04g00472 | 5 | 0.138888889 |
| TpMuguga_04g00472 | 6 | 0 |
| TpMuguga_04g00472 | 7 | 0 |
| TpMuguga_04g00472 | 8 | 0 |
| TpMuguga_04g00472 | 9 | 4.027777778 |
| TpMuguga_04g00473 | 1 | 0 |
| TpMuguga_04g00473 | 2 | 0 |
| TpMuguga_04g00473 | 3 | 0 |
| TpMuguga_04g00473 | 4 | 0 |
| TpMuguga_04g00473 | 5 | 0 |
| TpMuguga_04g00473 | 6 | 0 |
| TpMuguga_04g00473 | 7 | 0 |
| TpMuguga_04g00473 | 8 | 0 |
| TpMuguga_04g00473 | 9 | 0 |
| TpMuguga_04g00473 | 10 | 0 |
| TpMuguga_04g00473 | 11 | 0 |
| TpMuguga_04g00473 | 12 | 30 |
| TpMuguga_04g00473 | 13 | 45 |
| TpMuguga_04g00473 | 14 | 0 |
| TpMuguga_04g00474 | 1 | 0 |
| TpMuguga_04g00474 | 2 | 0 |
| TpMuguga_04g00474 | 3 | 0 |
| TpMuguga_04g00474 | 4 | 1.111111111 |
| TpMuguga_04g00474 | 5 | 0 |
| TpMuguga_04g00474 | 6 | 0 |
| TpMuguga_04g00474 | 7 | 8.888888889 |
| TpMuguga_04g00474 | 8 | 0 |
| TpMuguga_04g00474 | 9 | 0 |
| TpMuguga_04g00475 | 1 | 0 |
| TpMuguga_04g00475 | 2 | 0 |
| TpMuguga_04g00475 | 3 | 0 |
| TpMuguga_04g00475 | 4 | 0 |
| TpMuguga_04g00475 | 5 | 0 |
| TpMuguga_04g00475 | 6 | 0 |
| TpMuguga_04g00475 | 7 | 0.516923077 |
| TpMuguga_04g00476 | 1 | 0.044609665 |
| TpMuguga_04g00477 | 1 | 0.872727273 |
| TpMuguga_04g00477 | 2 | 0.872727273 |
| TpMuguga_04g00477 | 3 | 0 |
| TpMuguga_04g00477 | 4 | 0 |
| TpMuguga_04g00477 | 5 | 0 |

| | | |
|---|---|---|
| TpMuguga_04g00477 | 6 | 0 |
| TpMuguga_04g00477 | 7 | 0 |
| TpMuguga_04g00479 | 1 | 0 |
| TpMuguga_04g00479 | 2 | 0 |
| TpMuguga_04g00479 | 3 | 0 |
| TpMuguga_04g00479 | 4 | 0 |
| TpMuguga_04g00480 | 1 | 0 |
| TpMuguga_04g00480 | 2 | 0 |
| TpMuguga_04g00480 | 3 | 0 |
| TpMuguga_04g00481 | 1 | 0 |
| TpMuguga_04g00481 | 2 | 0 |
| TpMuguga_04g02095 | 1 | 0 |
| TpMuguga_04g00482 | 1 | 0 |
| TpMuguga_04g00483 | 1 | 0 |
| TpMuguga_04g00484 | 1 | 0.170731707 |
| TpMuguga_04g00484 | 2 | 1.682926829 |
| TpMuguga_04g00484 | 3 | 0 |
| TpMuguga_04g00485 | 1 | 0 |
| TpMuguga_04g00485 | 2 | 0 |
| TpMuguga_04g00485 | 3 | 0 |
| TpMuguga_04g00485 | 4 | 0 |
| TpMuguga_04g00485 | 5 | 0 |
| TpMuguga_04g00485 | 6 | 0 |
| TpMuguga_04g00485 | 7 | 0 |
| TpMuguga_04g00485 | 8 | 0 |
| TpMuguga_04g00485 | 9 | 0 |
| TpMuguga_04g00485 | 10 | 0 |
| TpMuguga_04g00486 | 1 | 0 |
| TpMuguga_04g00486 | 2 | 0 |
| TpMuguga_04g00486 | 3 | 0 |
| TpMuguga_04g00487 | 1 | 0 |
| TpMuguga_04g00487 | 2 | 0 |
| TpMuguga_04g00487 | 3 | 0 |
| TpMuguga_04g02215 | 1 | 0 |
| TpMuguga_04g02215 | 2 | 0 |
| TpMuguga_04g02215 | 3 | 0 |
| TpMuguga_04g02215 | 4 | 0 |
| TpMuguga_04g02215 | 5 | 0 |
| TpMuguga_04g02215 | 6 | 0 |
| TpMuguga_04g02215 | 7 | 0 |
| TpMuguga_04g02215 | 8 | 0 |
| TpMuguga_04g02215 | 9 | 0 |

| | | |
|---|---|---|
| TpMuguga_04g02215 | 10 | 0 |
| TpMuguga_04g02215 | 11 | 0 |
| TpMuguga_04g02215 | 12 | 0 |
| TpMuguga_04g02215 | 13 | 0 |
| TpMuguga_04g00492 | 1 | 0 |
| TpMuguga_04g00491 | 1 | 0 |
| TpMuguga_04g00491 | 2 | 0 |
| TpMuguga_04g00491 | 3 | 0 |
| TpMuguga_04g00491 | 4 | 0 |
| TpMuguga_04g00491 | 5 | 0 |
| TpMuguga_04g00491 | 6 | 0 |
| TpMuguga_04g00494 | 1 | 0 |
| TpMuguga_04g00494 | 2 | 0 |
| TpMuguga_04g00494 | 3 | 0 |
| TpMuguga_04g00494 | 4 | 0 |
| TpMuguga_04g00494 | 5 | 0 |
| TpMuguga_04g00495 | 1 | 4.8 |
| TpMuguga_04g00495 | 2 | 0 |
| TpMuguga_04g00495 | 3 | 0 |
| TpMuguga_04g00495 | 4 | 0 |
| TpMuguga_04g00495 | 5 | 0 |
| TpMuguga_04g00495 | 6 | 14.4 |
| TpMuguga_04g00495 | 7 | 0 |
| TpMuguga_04g00496 | 1 | 0 |
| TpMuguga_04g00496 | 2 | 0 |
| TpMuguga_04g00496 | 3 | 0 |
| TpMuguga_04g00496 | 4 | 0 |
| TpMuguga_04g00496 | 5 | 0 |
| TpMuguga_04g00496 | 6 | 0 |
| TpMuguga_04g00496 | 7 | 0 |
| TpMuguga_04g00497 | 1 | 0 |
| TpMuguga_04g00497 | 2 | 0 |
| TpMuguga_04g00497 | 3 | 0 |
| TpMuguga_04g00497 | 4 | 0 |
| TpMuguga_04g00497 | 5 | 0 |
| TpMuguga_04g00497 | 6 | 0 |
| TpMuguga_04g00497 | 7 | 0 |
| TpMuguga_04g00499 | 1 | 0 |
| TpMuguga_04g00499 | 2 | 0 |
| TpMuguga_04g00499 | 3 | 0 |
| TpMuguga_04g00499 | 4 | 0.943262411 |
| TpMuguga_04g00499 | 5 | 1.489361702 |

| | | |
|---|---|---|
| TpMuguga_04g00499 | 6 | 0.496453901 |
| TpMuguga_04g00500 | 1 | 0 |
| TpMuguga_04g02225 | 1 | 0.469798658 |
| TpMuguga_04g02225 | 2 | 0 |
| TpMuguga_04g02225 | 3 | 0 |
| TpMuguga_04g02225 | 4 | 0.33557047 |
| TpMuguga_04g00502 | 1 | 0 |
| TpMuguga_04g00502 | 2 | 0 |
| TpMuguga_04g00502 | 3 | 3.652173913 |
| TpMuguga_04g00502 | 4 | 0 |
| TpMuguga_04g00502 | 5 | 0 |
| TpMuguga_04g00502 | 6 | 4.695652174 |
| TpMuguga_04g00502 | 7 | 1.565217391 |
| TpMuguga_04g00502 | 8 | 1.565217391 |
| TpMuguga_04g00502 | 9 | 3.652173913 |
| TpMuguga_04g00502 | 10 | 0 |
| TpMuguga_04g00502 | 11 | 2.608695652 |
| TpMuguga_04g00504 | 1 | 0 |
| TpMuguga_04g00504 | 2 | 0 |
| TpMuguga_04g00504 | 3 | 0 |
| TpMuguga_04g00504 | 4 | 0 |
| TpMuguga_04g00504 | 5 | 0 |
| TpMuguga_04g00504 | 6 | 0 |
| TpMuguga_04g00504 | 7 | 0 |
| TpMuguga_04g00504 | 8 | 0 |
| TpMuguga_04g00505 | 1 | 0 |
| TpMuguga_04g00505 | 2 | 0 |
| TpMuguga_04g00506 | 1 | 10.83333333 |
| TpMuguga_04g00506 | 2 | 13 |
| TpMuguga_04g00506 | 3 | 2.166666667 |
| TpMuguga_04g00506 | 4 | 4.333333333 |
| TpMuguga_04g00506 | 5 | 13 |
| TpMuguga_04g00506 | 6 | 0 |
| TpMuguga_04g00506 | 7 | 0 |
| TpMuguga_04g00506 | 8 | 0 |
| TpMuguga_04g00506 | 9 | 21.66666667 |
| TpMuguga_04g00506 | 10 | 0 |
| TpMuguga_04g00506 | 11 | 0 |
| TpMuguga_04g00506 | 12 | 0 |
| TpMuguga_04g00508 | 1 | 0.909090909 |
| TpMuguga_04g00508 | 2 | 0 |
| TpMuguga_04g00508 | 3 | 1.363636364 |

| | | |
|---|---|---|
| TpMuguga_04g00508 | 4 | 2.272727273 |
| TpMuguga_04g02520 | 1 | 0.544554455 |
| TpMuguga_04g02520 | 2 | 0 |
| TpMuguga_04g02520 | 3 | 0 |
| TpMuguga_04g02520 | 4 | 0.435643564 |
| TpMuguga_04g02520 | 5 | 0.108910891 |
| TpMuguga_04g02520 | 6 | 0 |
| TpMuguga_04g02520 | 7 | 0.653465347 |
| TpMuguga_04g02520 | 8 | 0.108910891 |
| TpMuguga_04g02520 | 9 | 0 |
| TpMuguga_04g02520 | 10 | 0.326732673 |
| TpMuguga_04g02525 | 1 | 0 |
| TpMuguga_04g02525 | 2 | 0 |
| TpMuguga_04g02525 | 3 | 0 |
| TpMuguga_04g02525 | 4 | 0 |
| TpMuguga_04g02525 | 5 | 0 |
| TpMuguga_04g02525 | 6 | 0 |
| TpMuguga_04g02525 | 7 | 11.25 |
| TpMuguga_04g02525 | 8 | 0 |
| TpMuguga_04g02525 | 9 | 0 |
| TpMuguga_04g02525 | 10 | 22.5 |
| TpMuguga_04g02525 | 11 | 0 |
| TpMuguga_04g02525 | 12 | 0 |
| TpMuguga_04g02525 | 13 | 0 |
| TpMuguga_04g02525 | 14 | 0 |
| TpMuguga_04g00514 | 1 | 0 |
| TpMuguga_04g00514 | 2 | 0.365853659 |
| TpMuguga_04g00514 | 3 | 0.43902439 |
| TpMuguga_04g00514 | 4 | 0.073170732 |
| TpMuguga_04g00514 | 5 | 1.024390244 |
| TpMuguga_04g00513 | 1 | 0 |
| TpMuguga_04g00517 | 1 | 0.312233519 |
| TpMuguga_04g00517 | 2 | 0.084945884 |
| TpMuguga_04g00517 | 3 | 0 |
| TpMuguga_04g00517 | 4 | 0 |
| TpMuguga_04g00517 | 5 | 0 |
| TpMuguga_04g00517 | 6 | 0 |
| TpMuguga_04g02410 | 1 | 0 |
| TpMuguga_04g02410 | 2 | 0.306122449 |
| TpMuguga_04g02410 | 3 | 0 |
| TpMuguga_04g02410 | 4 | 0.306122449 |
| TpMuguga_04g02410 | 5 | 0 |

| | | |
|---|---|---|
| TpMuguga_04g02410 | 6 | 0.204081633 |
| TpMuguga_04g02410 | 7 | 0.051020408 |
| TpMuguga_04g02410 | 8 | 1.147959184 |
| TpMuguga_04g02410 | 9 | 2.780612245 |
| TpMuguga_04g00518 | 1 | 0 |
| TpMuguga_04g00518 | 2 | 0 |
| TpMuguga_04g00518 | 3 | 0 |
| TpMuguga_04g00518 | 4 | 0 |
| TpMuguga_04g02540 | 1 | 0 |
| TpMuguga_04g02540 | 2 | 0 |
| TpMuguga_04g02540 | 3 | 0 |
| TpMuguga_04g02540 | 4 | 0 |
| TpMuguga_04g02540 | 5 | 0 |
| TpMuguga_04g02540 | 6 | 0 |
| TpMuguga_04g02540 | 7 | 0 |
| TpMuguga_04g02540 | 8 | 0 |
| TpMuguga_04g02540 | 9 | 0 |
| TpMuguga_04g00521 | 1 | 0 |
| TpMuguga_04g00521 | 2 | 4.5 |
| TpMuguga_04g02485 | 1 | 0 |
| TpMuguga_04g02485 | 2 | 0 |
| TpMuguga_04g02485 | 3 | 0 |
| TpMuguga_04g02485 | 4 | 0 |
| TpMuguga_04g00524 | 1 | 0 |
| TpMuguga_04g00524 | 2 | 0 |
| TpMuguga_04g00525 | 1 | 0 |
| TpMuguga_04g00525 | 2 | 0 |
| TpMuguga_04g00525 | 3 | 0 |
| TpMuguga_04g00525 | 4 | 0 |
| TpMuguga_04g00525 | 5 | 0 |
| TpMuguga_04g00525 | 6 | 0 |
| TpMuguga_04g00525 | 7 | 0 |
| TpMuguga_04g00525 | 8 | 0 |
| TpMuguga_04g00527 | 1 | 1.875 |
| TpMuguga_04g00527 | 2 | 1.875 |
| TpMuguga_04g00527 | 3 | 0 |
| TpMuguga_04g00527 | 4 | 0 |
| TpMuguga_04g00529 | 1 | 2 |
| TpMuguga_04g00528 | 1 | 0 |
| TpMuguga_04g00528 | 2 | 0 |
| TpMuguga_04g00528 | 3 | 0 |
| TpMuguga_04g00528 | 4 | 0 |

| | | |
|---|---|---|
| TpMuguga_04g00528 | 5 | 0 |
| TpMuguga_04g00528 | 6 | 0 |
| TpMuguga_04g00528 | 7 | 0 |
| TpMuguga_04g00531 | 1 | 0 |
| TpMuguga_04g00531 | 2 | 0 |
| TpMuguga_04g00534 | 1 | 0 |
| TpMuguga_04g00534 | 2 | 1.68 |
| TpMuguga_04g00533 | 1 | 0 |
| TpMuguga_04g00533 | 2 | 0 |
| TpMuguga_04g00533 | 3 | 0 |
| TpMuguga_04g00533 | 4 | 0 |
| TpMuguga_04g00533 | 5 | 0 |
| TpMuguga_04g00533 | 6 | 0 |
| TpMuguga_04g00533 | 7 | 0 |
| TpMuguga_04g00535 | 1 | 0 |
| TpMuguga_04g00537 | 1 | 0 |
| TpMuguga_04g00538 | 1 | 0.875 |
| TpMuguga_04g00538 | 2 | 0 |
| TpMuguga_04g00539 | 1 | 0 |
| TpMuguga_04g00539 | 2 | 0 |
| TpMuguga_04g00540 | 1 | 0 |
| TpMuguga_04g00540 | 2 | 0 |
| TpMuguga_04g00540 | 3 | 0 |
| TpMuguga_04g00540 | 4 | 0 |
| TpMuguga_04g00540 | 5 | 0 |
| TpMuguga_04g00541 | 1 | 0 |
| TpMuguga_04g00541 | 2 | 0 |
| TpMuguga_04g00541 | 3 | 0 |
| TpMuguga_04g00541 | 4 | 0 |
| TpMuguga_04g00541 | 5 | 0 |
| TpMuguga_04g00541 | 6 | 0 |
| TpMuguga_04g00542 | 1 | 1.285470085 |
| TpMuguga_04g00543 | 1 | 0 |
| TpMuguga_04g00543 | 2 | 0.876404494 |
| TpMuguga_04g00543 | 3 | 2.02247191 |
| TpMuguga_04g00543 | 4 | 3.842696629 |
| TpMuguga_04g00543 | 5 | 4.247191011 |
| TpMuguga_04g00544 | 1 | 0.625 |
| TpMuguga_04g00544 | 2 | 0 |
| TpMuguga_04g00544 | 3 | 0 |
| TpMuguga_04g00544 | 4 | 0 |
| TpMuguga_04g00545 | 1 | 0 |

| | | |
|---|---|---|
| TpMuguga_04g00545 | 2 | 32 |
| TpMuguga_04g00545 | 3 | 13.33333333 |
| TpMuguga_04g00547 | 1 | 0.189090909 |
| TpMuguga_04g00547 | 2 | 0 |
| TpMuguga_04g00547 | 3 | 0 |
| TpMuguga_04g00546 | 1 | 0 |
| TpMuguga_04g00546 | 2 | 0 |
| TpMuguga_04g00548 | 1 | 0.064880113 |
| TpMuguga_04g00550 | 1 | 0 |
| TpMuguga_04g00550 | 2 | 0 |
| TpMuguga_04g00550 | 3 | 0 |
| TpMuguga_04g00550 | 4 | 0 |
| TpMuguga_04g00550 | 5 | 0 |
| TpMuguga_04g00550 | 6 | 0 |
| TpMuguga_04g00550 | 7 | 0 |
| TpMuguga_04g00550 | 8 | 0.392857143 |
| TpMuguga_04g00550 | 9 | 0 |
| TpMuguga_04g00550 | 10 | 8.642857143 |
| TpMuguga_04g00549 | 1 | 0.133333333 |
| TpMuguga_04g00549 | 2 | 0 |
| TpMuguga_04g00549 | 3 | 0 |
| TpMuguga_04g00549 | 4 | 0.577777778 |
| TpMuguga_04g00549 | 5 | 0.177777778 |
| TpMuguga_04g00551 | 1 | 0.071684588 |
| TpMuguga_04g00551 | 2 | 0 |
| TpMuguga_04g00551 | 3 | 0.896057348 |
| TpMuguga_04g00551 | 4 | 0.663082437 |
| TpMuguga_04g00552 | 1 | 0 |
| TpMuguga_04g00552 | 2 | 0 |
| TpMuguga_04g00552 | 3 | 0 |
| TpMuguga_04g00552 | 4 | 5 |
| TpMuguga_04g02365 | 1 | 0 |
| TpMuguga_04g02365 | 2 | 0 |
| TpMuguga_04g02365 | 3 | 0 |
| TpMuguga_04g02365 | 4 | 0 |
| TpMuguga_04g02365 | 5 | 0 |
| TpMuguga_04g02365 | 6 | 0 |
| TpMuguga_04g02365 | 7 | 0.103448276 |
| TpMuguga_04g02365 | 8 | 0.362068966 |
| TpMuguga_04g00553 | 1 | 0.592592593 |
| TpMuguga_04g00556 | 1 | 1.287939988 |
| TpMuguga_04g00556 | 2 | 0.173110214 |

| | | |
|---|---|---|
| TpMuguga_04g00556 | 3 | 0.657818811 |
| TpMuguga_04g00556 | 4 | 1.010963647 |
| TpMuguga_04g00556 | 5 | 0.609347952 |
| TpMuguga_04g00556 | 6 | 1.260242354 |
| TpMuguga_04g00556 | 7 | 1.710328909 |
| TpMuguga_04g00556 | 8 | 0.581650317 |
| TpMuguga_04g00556 | 9 | 0.353144836 |
| TpMuguga_04g00556 | 10 | 0.422388921 |
| TpMuguga_04g00556 | 11 | 0.879399885 |
| TpMuguga_04g00557 | 1 | 0.057142857 |
| TpMuguga_04g00557 | 2 | 0 |
| TpMuguga_04g00557 | 3 | 0.228571429 |
| TpMuguga_04g00557 | 4 | 0 |
| TpMuguga_04g00557 | 5 | 0.133333333 |
| TpMuguga_04g00558 | 1 | 0 |
| TpMuguga_04g00558 | 2 | 0 |
| TpMuguga_04g00558 | 3 | 0 |
| TpMuguga_04g00558 | 4 | 0 |
| TpMuguga_04g00558 | 5 | 0 |
| TpMuguga_04g00558 | 6 | 0 |
| TpMuguga_04g00558 | 7 | 1.454545455 |
| TpMuguga_04g00559 | 1 | 0 |
| TpMuguga_04g00559 | 2 | 0 |
| TpMuguga_04g00559 | 3 | 0 |
| TpMuguga_04g00559 | 4 | 1.589830508 |
| TpMuguga_04g00559 | 5 | 1.589830508 |
| TpMuguga_04g00559 | 6 | 8.566101695 |
| TpMuguga_04g00560 | 1 | 18 |
| TpMuguga_04g00560 | 2 | 0 |
| TpMuguga_04g00560 | 3 | 0 |
| TpMuguga_04g00560 | 4 | 0 |
| TpMuguga_04g00560 | 5 | 0 |
| TpMuguga_04g00561 | 1 | 1.565217391 |
| TpMuguga_04g00561 | 2 | 0 |
| TpMuguga_04g00561 | 3 | 0 |
| TpMuguga_04g00561 | 4 | 0 |
| TpMuguga_04g00561 | 5 | 0 |
| TpMuguga_04g00561 | 6 | 0 |
| TpMuguga_04g00561 | 7 | 0.869565217 |
| TpMuguga_04g00562 | 1 | 0 |
| TpMuguga_04g00562 | 2 | 0 |
| TpMuguga_04g00563 | 1 | 0.333333333 |

| | | |
|---|---|---|
| TpMuguga_04g00563 | 2 | 0.114583333 |
| TpMuguga_04g00566 | 1 | 0 |
| TpMuguga_04g00566 | 2 | 0 |
| TpMuguga_04g00566 | 3 | 0 |
| TpMuguga_04g00566 | 4 | 0 |
| TpMuguga_04g00566 | 5 | 0 |
| TpMuguga_04g00566 | 6 | 4.375 |
| TpMuguga_04g02710 | 1 | 0 |
| TpMuguga_04g02710 | 2 | 0 |
| TpMuguga_04g02710 | 3 | 0 |
| TpMuguga_04g02710 | 4 | 10 |
| TpMuguga_04g02705 | 1 | 0 |
| TpMuguga_04g02705 | 2 | 0 |
| TpMuguga_04g00568 | 1 | 2.976744186 |
| TpMuguga_04g00568 | 2 | 2.139534884 |
| TpMuguga_04g00568 | 3 | 2.88372093 |
| TpMuguga_04g00571 | 1 | 0 |
| TpMuguga_04g00571 | 2 | 0 |
| TpMuguga_04g00571 | 3 | 0 |
| TpMuguga_04g00571 | 4 | 2.055 |
| TpMuguga_04g00571 | 5 | 1.515 |
| TpMuguga_04g00574 | 1 | 0 |
| TpMuguga_04g00574 | 2 | 0 |
| TpMuguga_04g00574 | 3 | 0 |
| TpMuguga_04g00574 | 4 | 0 |
| TpMuguga_04g00574 | 5 | 0 |
| TpMuguga_04g00574 | 6 | 0 |
| TpMuguga_04g00574 | 7 | 0 |
| TpMuguga_04g00574 | 8 | 0 |
| TpMuguga_04g00574 | 9 | 0 |
| TpMuguga_04g00574 | 10 | 0 |
| TpMuguga_04g00574 | 11 | 0 |
| TpMuguga_04g00574 | 12 | 0 |
| TpMuguga_04g00574 | 13 | 0 |
| TpMuguga_04g00577 | 1 | 14.24015009 |
| TpMuguga_04g00577 | 2 | 0.041275797 |
| TpMuguga_04g00577 | 3 | 0 |
| TpMuguga_04g00577 | 4 | 0.041275797 |
| TpMuguga_04g00577 | 5 | 0 |
| TpMuguga_04g00577 | 6 | 0.268292683 |
| TpMuguga_04g00577 | 7 | 0 |
| TpMuguga_04g00577 | 8 | 0 |

| | | |
|---|---|---|
| TpMuguga_04g00577 | 9 | 0 |
| TpMuguga_04g00577 | 10 | 0 |
| TpMuguga_04g00576 | 1 | 0.01188825 |
| TpMuguga_04g00578 | 1 | 0.010126582 |
| TpMuguga_04g00578 | 2 | 0.005063291 |
| TpMuguga_04g00578 | 3 | 0.030379747 |
| TpMuguga_04g00578 | 4 | 0 |
| TpMuguga_04g00578 | 5 | 0.470886076 |
| TpMuguga_04g00579 | 1 | 0 |
| TpMuguga_04g00581 | 1 | 0 |
| TpMuguga_04g00582 | 1 | 0 |
| TpMuguga_04g00583 | 1 | 1.471698113 |
| TpMuguga_04g00584 | 1 | 3.6 |
| TpMuguga_04g00585 | 1 | 1.883161512 |
| TpMuguga_04g02130 | 1 | 0 |
| TpMuguga_04g02130 | 2 | 0 |
| TpMuguga_04g02130 | 3 | 0 |
| TpMuguga_04g02130 | 4 | 0 |
| TpMuguga_04g02270 | 1 | 0 |
| TpMuguga_04g02270 | 2 | 0 |
| TpMuguga_04g02270 | 3 | 0 |
| TpMuguga_04g02270 | 4 | 0.014705882 |
| TpMuguga_04g02270 | 5 | 0.068627451 |
| TpMuguga_04g02155 | 1 | 0 |
| TpMuguga_04g02155 | 2 | 0 |
| TpMuguga_04g00592 | 1 | 0 |
| TpMuguga_04g00592 | 2 | 0 |
| TpMuguga_04g00592 | 3 | 0 |
| TpMuguga_04g00592 | 4 | 0 |
| TpMuguga_04g00592 | 5 | 0 |
| TpMuguga_04g00592 | 6 | 0 |
| TpMuguga_04g00592 | 7 | 0 |
| TpMuguga_04g00592 | 8 | 0 |
| TpMuguga_04g00592 | 9 | 0 |
| TpMuguga_04g00594 | 1 | 0 |
| TpMuguga_04g00594 | 2 | 0 |
| TpMuguga_04g00594 | 3 | 0 |
| TpMuguga_04g00594 | 4 | 0 |
| TpMuguga_04g00594 | 5 | 0 |
| TpMuguga_04g00594 | 6 | 0 |
| TpMuguga_04g00595 | 1 | 1.724137931 |
| TpMuguga_04g00595 | 2 | 0 |

| | | |
|---|---|---|
| TpMuguga_04g00595 | 3 | 0 |
| TpMuguga_04g00595 | 4 | 0 |
| TpMuguga_04g00596 | 1 | 0 |
| TpMuguga_04g00596 | 2 | 0 |
| TpMuguga_04g00597 | 1 | 0.3 |
| TpMuguga_04g00598 | 1 | 0 |
| TpMuguga_04g00598 | 2 | 0 |
| TpMuguga_04g00598 | 3 | 0 |
| TpMuguga_04g00598 | 4 | 0 |
| TpMuguga_04g00598 | 5 | 0 |
| TpMuguga_04g02650 | 1 | 3.428571429 |
| TpMuguga_04g02650 | 2 | 0 |
| TpMuguga_04g02655 | 1 | 0 |
| TpMuguga_04g02655 | 2 | 0 |
| TpMuguga_04g00604 | 1 | 0 |
| TpMuguga_04g00604 | 2 | 2 |
| TpMuguga_04g00604 | 3 | 10 |
| TpMuguga_04g00603 | 1 | 0 |
| TpMuguga_04g00605 | 1 | 0 |
| TpMuguga_04g00605 | 2 | 0 |
| TpMuguga_04g00605 | 3 | 0 |
| TpMuguga_04g00605 | 4 | 0 |
| TpMuguga_04g00606 | 1 | 3.571428571 |
| TpMuguga_04g00606 | 2 | 0 |
| TpMuguga_04g00606 | 3 | 0 |
| TpMuguga_04g00606 | 4 | 0 |
| TpMuguga_04g00608 | 1 | 0 |
| TpMuguga_04g00608 | 2 | 0 |
| TpMuguga_04g00608 | 3 | 0 |
| TpMuguga_04g00608 | 4 | 0 |
| TpMuguga_04g00608 | 5 | 0 |
| TpMuguga_04g00608 | 6 | 0 |
| TpMuguga_04g00608 | 7 | 0 |
| TpMuguga_04g00608 | 8 | 0 |
| TpMuguga_04g00608 | 9 | 0 |
| TpMuguga_04g00607 | 1 | 0 |
| TpMuguga_04g00607 | 2 | 3 |
| TpMuguga_04g00611 | 1 | 0.060098119 |
| TpMuguga_04g00611 | 2 | 0 |
| TpMuguga_04g00610 | 1 | 0.265895954 |
| TpMuguga_04g02220 | 1 | 0 |
| TpMuguga_04g02220 | 2 | 1.695652174 |

| | | |
|---|---|---|
| TpMuguga_04g00614 | 1 | 0 |
| TpMuguga_04g00614 | 2 | 0 |
| TpMuguga_04g00614 | 3 | 0 |
| TpMuguga_04g00614 | 4 | 0 |
| TpMuguga_04g00614 | 5 | 0 |
| TpMuguga_04g00614 | 6 | 0 |
| TpMuguga_04g00616 | 1 | 0 |
| TpMuguga_04g00616 | 2 | 0 |
| TpMuguga_04g00615 | 1 | 0 |
| TpMuguga_04g00615 | 2 | 0 |
| TpMuguga_04g00615 | 3 | 0 |
| TpMuguga_04g00617 | 1 | 0 |
| TpMuguga_04g00617 | 2 | 0 |
| TpMuguga_04g00618 | 1 | 0 |
| TpMuguga_04g00618 | 2 | 0 |
| TpMuguga_04g00618 | 3 | 0 |
| TpMuguga_04g00620 | 1 | 0 |
| TpMuguga_04g00620 | 2 | 0.5 |
| TpMuguga_04g00620 | 3 | 0 |
| TpMuguga_04g00620 | 4 | 0 |
| TpMuguga_04g00620 | 5 | 1 |
| TpMuguga_04g00621 | 1 | 5.210526316 |
| TpMuguga_04g00621 | 2 | 0 |
| TpMuguga_04g00621 | 3 | 0 |
| TpMuguga_04g00621 | 4 | 0 |
| TpMuguga_04g00621 | 5 | 0 |
| TpMuguga_04g00621 | 6 | 0 |
| TpMuguga_04g00621 | 7 | 0 |
| TpMuguga_04g00621 | 8 | 0.144736842 |
| TpMuguga_04g00621 | 9 | 0 |
| TpMuguga_04g00621 | 10 | 0 |
| TpMuguga_04g00622 | 1 | 6.545454545 |
| TpMuguga_04g00622 | 2 | 0.727272727 |
| TpMuguga_04g00622 | 3 | 0 |
| TpMuguga_04g00622 | 4 | 0 |
| TpMuguga_04g00622 | 5 | 0 |
| TpMuguga_04g00622 | 6 | 0.727272727 |
| TpMuguga_04g00622 | 7 | 0 |
| TpMuguga_04g02075 | 1 | 0 |
| TpMuguga_04g02075 | 2 | 0 |
| TpMuguga_04g02075 | 3 | 1.372434018 |
| TpMuguga_04g00623 | 1 | 0.511627907 |

| | | |
|---|---|---|
| TpMuguga_04g00624 | 1 | 0 |
| TpMuguga_04g00624 | 2 | 5 |
| TpMuguga_04g00624 | 3 | 2.5 |
| TpMuguga_04g00624 | 4 | 0 |
| TpMuguga_04g00625 | 1 | 3.5 |
| TpMuguga_04g00625 | 2 | 0 |
| TpMuguga_04g00625 | 3 | 0 |
| TpMuguga_04g00625 | 4 | 10.5 |
| TpMuguga_04g00625 | 5 | 0 |
| TpMuguga_04g00625 | 6 | 0 |
| TpMuguga_04g00626 | 1 | 0 |
| TpMuguga_04g00629 | 1 | 1.2 |
| TpMuguga_04g02545 | 1 | 0 |
| TpMuguga_04g02545 | 2 | 0 |
| TpMuguga_04g02545 | 3 | 0 |
| TpMuguga_04g02545 | 4 | 0 |
| TpMuguga_04g02545 | 5 | 0 |
| TpMuguga_04g02545 | 6 | 0 |
| TpMuguga_04g02545 | 7 | 0 |
| TpMuguga_04g02545 | 8 | 0 |
| TpMuguga_04g02545 | 9 | 0 |
| TpMuguga_04g02545 | 10 | 0 |
| TpMuguga_04g02545 | 11 | 0 |
| TpMuguga_04g02545 | 12 | 0 |
| TpMuguga_04g02545 | 13 | 0 |
| TpMuguga_04g02545 | 14 | 0 |
| TpMuguga_04g02185 | 1 | 0 |
| TpMuguga_04g02185 | 2 | 0 |
| TpMuguga_04g02185 | 3 | 0 |
| TpMuguga_04g02185 | 4 | 0 |
| TpMuguga_04g02185 | 5 | 0 |
| TpMuguga_04g02185 | 6 | 0 |
| TpMuguga_04g02185 | 7 | 0 |
| TpMuguga_04g02185 | 8 | 0 |
| TpMuguga_04g02185 | 9 | 0 |
| TpMuguga_04g00634 | 1 | 9 |
| TpMuguga_04g00634 | 2 | 1.25 |
| TpMuguga_04g00634 | 3 | 2.25 |
| TpMuguga_04g00633 | 1 | 1.5 |
| TpMuguga_04g00633 | 2 | 0 |
| TpMuguga_04g00636 | 1 | 0 |
| TpMuguga_04g00637 | 1 | 0 |

| | | |
|---|---|---|
| TpMuguga_04g00637 | 2 | 0.077170418 |
| TpMuguga_04g00637 | 3 | 0.115755627 |
| TpMuguga_04g00638 | 1 | 0.014563107 |
| TpMuguga_04g00641 | 1 | 0.918367347 |
| TpMuguga_04g00641 | 2 | 0.006122449 |
| TpMuguga_04g00640 | 1 | 0.041880151 |
| TpMuguga_04g00640 | 2 | 0.118511918 |
| TpMuguga_04g00640 | 3 | 0.327021608 |
| TpMuguga_04g00642 | 1 | 0.413043478 |
| TpMuguga_04g00643 | 1 | 0 |
| TpMuguga_04g00643 | 2 | 0.090909091 |
| TpMuguga_04g00644 | 1 | 0 |
| TpMuguga_04g00644 | 2 | 0 |
| TpMuguga_04g00644 | 3 | 0 |
| TpMuguga_04g00644 | 4 | 0 |
| TpMuguga_04g00644 | 5 | 0 |
| TpMuguga_04g00644 | 6 | 0 |
| TpMuguga_04g00644 | 7 | 0 |
| TpMuguga_04g00644 | 8 | 0 |
| TpMuguga_04g00644 | 9 | 0 |
| TpMuguga_04g00645 | 1 | 0 |
| TpMuguga_04g00645 | 2 | 0 |
| TpMuguga_04g00645 | 3 | 0 |
| TpMuguga_04g00645 | 4 | 0 |
| TpMuguga_04g00646 | 1 | 0 |
| TpMuguga_04g00646 | 2 | 0 |
| TpMuguga_04g00646 | 3 | 0 |
| TpMuguga_04g00647 | 1 | 0 |
| TpMuguga_04g00648 | 1 | 0 |
| TpMuguga_04g00648 | 2 | 0 |
| TpMuguga_04g00648 | 3 | 0 |
| TpMuguga_04g00648 | 4 | 0 |
| TpMuguga_04g00648 | 5 | 0 |
| TpMuguga_04g00648 | 6 | 0 |
| TpMuguga_04g00648 | 7 | 0 |
| TpMuguga_04g00648 | 8 | 0 |
| TpMuguga_04g00648 | 9 | 0 |
| TpMuguga_04g00648 | 10 | 0 |
| TpMuguga_04g00649 | 1 | 0 |
| TpMuguga_04g00924 | 1 | 0 |
| TpMuguga_04g00924 | 2 | 0 |
| TpMuguga_04g00924 | 3 | 0 |

| | | |
|---|---|---|
| TpMuguga_04g00924 | 4 | 0 |
| TpMuguga_04g00924 | 5 | 0 |
| TpMuguga_04g00925 | 1 | 5 |
| TpMuguga_04g00925 | 2 | 0 |
| TpMuguga_04g00925 | 3 | 0 |
| TpMuguga_04g00925 | 4 | 1.363636364 |
| TpMuguga_04g00651 | 1 | 0 |
| TpMuguga_04g00652 | 1 | 0.033039648 |
| TpMuguga_04g00652 | 2 | 0 |
| TpMuguga_04g02370 | 1 | 0 |
| TpMuguga_04g02370 | 2 | 0 |
| TpMuguga_04g02370 | 3 | 3.5 |
| TpMuguga_04g02370 | 4 | 0 |
| TpMuguga_04g02370 | 5 | 0 |
| TpMuguga_04g02370 | 6 | 0 |
| TpMuguga_04g00654 | 1 | 0 |
| TpMuguga_04g00658 | 1 | 0 |
| TpMuguga_04g00658 | 2 | 0 |
| TpMuguga_04g00658 | 3 | 0 |
| TpMuguga_04g00658 | 4 | 0 |
| TpMuguga_04g00658 | 5 | 52 |
| TpMuguga_04g00658 | 6 | 0 |
| TpMuguga_04g00658 | 7 | 13 |
| TpMuguga_04g00658 | 8 | 13 |
| TpMuguga_04g00658 | 9 | 0 |
| TpMuguga_04g00658 | 10 | 0 |
| TpMuguga_04g00658 | 11 | 0 |
| TpMuguga_04g00658 | 12 | 0 |
| TpMuguga_04g00660 | 1 | 0 |
| TpMuguga_04g00660 | 2 | 0 |
| TpMuguga_04g00660 | 3 | 0 |
| TpMuguga_04g00660 | 4 | 0 |
| TpMuguga_04g00660 | 5 | 0 |
| TpMuguga_04g00660 | 6 | 0 |
| TpMuguga_04g00661 | 1 | 0 |
| TpMuguga_04g00661 | 2 | 0 |
| TpMuguga_04g00661 | 3 | 0 |
| TpMuguga_04g00661 | 4 | 0 |
| TpMuguga_04g00661 | 5 | 0 |
| TpMuguga_04g00661 | 6 | 7.438596491 |
| TpMuguga_04g00661 | 7 | 1.122807018 |
| TpMuguga_04g00662 | 1 | 0 |

| | | |
|---|---|---|
| TpMuguga_04g00662 | 2 | 0 |
| TpMuguga_04g00662 | 3 | 0 |
| TpMuguga_04g00663 | 1 | 0.714285714 |
| TpMuguga_04g00663 | 2 | 0.428571429 |
| TpMuguga_04g00663 | 3 | 0 |
| TpMuguga_04g00663 | 4 | 0 |
| TpMuguga_04g00663 | 5 | 0.857142857 |
| TpMuguga_04g00664 | 1 | 0 |
| TpMuguga_04g00664 | 2 | 0 |
| TpMuguga_04g00664 | 3 | 2.666666667 |
| TpMuguga_04g00665 | 1 | 0 |
| TpMuguga_04g00665 | 2 | 0 |
| TpMuguga_04g00665 | 3 | 0 |
| TpMuguga_04g00665 | 4 | 0 |
| TpMuguga_04g00665 | 5 | 0 |
| TpMuguga_04g00665 | 6 | 0 |
| TpMuguga_04g00665 | 7 | 0 |
| TpMuguga_04g00665 | 8 | 0 |
| TpMuguga_04g00665 | 9 | 0 |
| TpMuguga_04g00665 | 10 | 0 |
| TpMuguga_04g00665 | 11 | 0 |
| TpMuguga_04g00665 | 12 | 0 |
| TpMuguga_04g00665 | 13 | 0 |
| TpMuguga_04g00665 | 14 | 0 |
| TpMuguga_04g00665 | 15 | 0 |
| TpMuguga_04g02090 | 1 | 0 |
| TpMuguga_04g02090 | 2 | 0 |
| TpMuguga_04g02670 | 1 | 0 |
| TpMuguga_04g02670 | 2 | 0 |
| TpMuguga_04g02670 | 3 | 0 |
| TpMuguga_04g02670 | 4 | 0 |
| TpMuguga_04g02285 | 1 | 3.733333333 |
| TpMuguga_04g02285 | 2 | 2.133333333 |
| TpMuguga_04g02285 | 3 | 0 |
| TpMuguga_04g00668 | 1 | 0 |
| TpMuguga_04g00668 | 2 | 0 |
| TpMuguga_04g00668 | 3 | 0 |
| TpMuguga_04g00668 | 4 | 0 |
| TpMuguga_04g00668 | 5 | 0 |
| TpMuguga_04g00668 | 6 | 0 |
| TpMuguga_04g00668 | 7 | 0 |
| TpMuguga_04g00668 | 8 | 0 |

| | | |
|---|---|---|
| TpMuguga_04g00668 | 9 | 0 |
| TpMuguga_04g00669 | 1 | 2 |
| TpMuguga_04g00671 | 1 | 0 |
| TpMuguga_04g00671 | 2 | 0 |
| TpMuguga_04g00671 | 3 | 0 |
| TpMuguga_04g00671 | 4 | 0 |
| TpMuguga_04g00671 | 5 | 0 |
| TpMuguga_04g00671 | 6 | 0 |
| TpMuguga_04g00671 | 7 | 0 |
| TpMuguga_04g00671 | 8 | 0 |
| TpMuguga_04g00671 | 9 | 0 |
| TpMuguga_04g00671 | 10 | 0 |
| TpMuguga_04g00671 | 11 | 0 |
| TpMuguga_04g00671 | 12 | 0 |
| TpMuguga_04g00671 | 13 | 0 |
| TpMuguga_04g00670 | 1 | 0 |
| TpMuguga_04g02160 | 1 | 0 |
| TpMuguga_04g02160 | 2 | 16.25 |
| TpMuguga_04g02160 | 3 | 0 |
| TpMuguga_04g02160 | 4 | 3.75 |
| TpMuguga_04g00672 | 1 | 0 |
| TpMuguga_04g00672 | 2 | 0 |
| TpMuguga_04g00672 | 3 | 0 |
| TpMuguga_04g00672 | 4 | 0 |
| TpMuguga_04g00672 | 5 | 0 |
| TpMuguga_04g00672 | 6 | 0 |
| TpMuguga_04g00672 | 7 | 0 |
| TpMuguga_04g00676 | 1 | 0 |
| TpMuguga_04g00676 | 2 | 0 |
| TpMuguga_04g00676 | 3 | 0 |
| TpMuguga_04g00676 | 4 | 0 |
| TpMuguga_04g00676 | 5 | 0 |
| TpMuguga_04g00676 | 6 | 0 |
| TpMuguga_04g00679 | 1 | 0 |
| TpMuguga_04g00679 | 2 | 0 |
| TpMuguga_04g00679 | 3 | 0.131147541 |
| TpMuguga_04g00678 | 1 | 0.438709677 |
| TpMuguga_04g00680 | 1 | 0.333333333 |
| TpMuguga_04g00681 | 1 | 0 |
| TpMuguga_04g00681 | 2 | 0 |
| TpMuguga_04g00681 | 3 | 0 |
| TpMuguga_04g00683 | 1 | 0.575539568 |

| | | |
|---|---|---|
| TpMuguga_04g00683 | 2 | 1.007194245 |
| TpMuguga_04g00683 | 3 | 0 |
| TpMuguga_04g00683 | 4 | 0.323741007 |
| TpMuguga_04g02245 | 1 | 0 |
| TpMuguga_04g02630 | 1 | 0 |
| TpMuguga_04g02630 | 2 | 0 |
| TpMuguga_04g02630 | 3 | 0 |
| TpMuguga_04g02630 | 4 | 0 |
| TpMuguga_04g02630 | 5 | 0 |
| TpMuguga_04g02630 | 6 | 0 |
| TpMuguga_04g02630 | 7 | 0 |
| TpMuguga_04g02630 | 8 | 0 |
| TpMuguga_04g02630 | 9 | 0 |
| TpMuguga_04g02630 | 10 | 0 |
| TpMuguga_04g02630 | 11 | 0 |
| TpMuguga_04g02630 | 12 | 0 |
| TpMuguga_04g00688 | 1 | 5.6875 |
| TpMuguga_04g00688 | 2 | 5.6875 |
| TpMuguga_04g00688 | 3 | 5.6875 |
| TpMuguga_04g00688 | 4 | 0 |
| TpMuguga_04g00688 | 5 | 0 |
| TpMuguga_04g00688 | 6 | 0 |
| TpMuguga_04g00687 | 1 | 0 |
| TpMuguga_04g00687 | 2 | 0 |
| TpMuguga_04g00687 | 3 | 0 |
| TpMuguga_04g00687 | 4 | 0 |
| TpMuguga_04g00687 | 5 | 0.211764706 |
| TpMuguga_04g00687 | 6 | 0 |
| TpMuguga_04g00687 | 7 | 8.047058824 |
| TpMuguga_04g00687 | 8 | 16.83529412 |
| TpMuguga_04g00689 | 1 | 0.172413793 |
| TpMuguga_04g00690 | 1 | 0.939393939 |
| TpMuguga_04g00690 | 2 | 1.515151515 |
| TpMuguga_04g00690 | 3 | 2.515151515 |
| TpMuguga_04g00690 | 4 | 1.545454545 |
| TpMuguga_04g00690 | 5 | 0.787878788 |
| TpMuguga_04g00691 | 1 | 0 |
| TpMuguga_04g02620 | 1 | 0.545454545 |
| TpMuguga_04g02620 | 2 | 0 |
| TpMuguga_04g02250 | 1 | 0 |
| TpMuguga_04g02250 | 2 | 0 |
| TpMuguga_04g02250 | 3 | 0 |

| | | |
|---|---|---|
| TpMuguga_04g02250 | 4 | 0 |
| TpMuguga_04g02250 | 5 | 0 |
| TpMuguga_04g00693 | 1 | 0.580645161 |
| TpMuguga_04g02040 | 1 | 0 |
| TpMuguga_04g02040 | 2 | 0 |
| TpMuguga_04g02040 | 3 | 0 |
| TpMuguga_04g02040 | 4 | 0 |
| TpMuguga_04g00696 | 1 | 0 |
| TpMuguga_04g00696 | 2 | 0 |
| TpMuguga_04g00696 | 3 | 0 |
| TpMuguga_04g00696 | 4 | 0 |
| TpMuguga_04g00696 | 5 | 0 |
| TpMuguga_04g00697 | 1 | 0 |
| TpMuguga_04g00697 | 2 | 0 |
| TpMuguga_04g00697 | 3 | 0 |
| TpMuguga_04g00697 | 4 | 0 |
| TpMuguga_04g00697 | 5 | 0 |
| TpMuguga_04g00697 | 6 | 0 |
| TpMuguga_04g00697 | 7 | 0 |
| TpMuguga_04g00697 | 8 | 0 |
| TpMuguga_04g02530 | 1 | 6.857142857 |
| TpMuguga_04g02530 | 2 | 0 |
| TpMuguga_04g02530 | 3 | 2.285714286 |
| TpMuguga_04g02530 | 4 | 0 |
| TpMuguga_04g02530 | 5 | 0 |
| TpMuguga_04g02530 | 6 | 5.714285714 |
| TpMuguga_04g02530 | 7 | 0 |
| TpMuguga_04g00700 | 1 | 0.4 |
| TpMuguga_04g00702 | 1 | 0 |
| TpMuguga_04g00703 | 1 | 0 |
| TpMuguga_04g00705 | 1 | 0 |
| TpMuguga_04g00705 | 2 | 0 |
| TpMuguga_04g00706 | 1 | 0 |
| TpMuguga_04g00706 | 2 | 0 |
| TpMuguga_04g00706 | 3 | 0 |
| TpMuguga_04g00706 | 4 | 0 |
| TpMuguga_04g00706 | 5 | 0 |
| TpMuguga_04g00706 | 6 | 0 |
| TpMuguga_04g00706 | 7 | 0 |
| TpMuguga_04g00706 | 8 | 0 |
| TpMuguga_04g00706 | 9 | 0 |
| TpMuguga_04g00706 | 10 | 0 |

| | | |
|---|---|---|
| TpMuguga_04g00706 | 11 | 0 |
| TpMuguga_04g00706 | 12 | 0 |
| TpMuguga_04g00706 | 13 | 0 |
| TpMuguga_04g00706 | 14 | 0 |
| TpMuguga_04g00706 | 15 | 0 |
| TpMuguga_04g00708 | 1 | 0 |
| TpMuguga_04g00708 | 2 | 0 |
| TpMuguga_04g02495 | 1 | 0 |
| TpMuguga_04g02495 | 2 | 0 |
| TpMuguga_04g02495 | 3 | 0 |
| TpMuguga_04g02495 | 4 | 3.714285714 |
| TpMuguga_04g02495 | 5 | 0 |
| TpMuguga_04g02495 | 6 | 0 |
| TpMuguga_04g02495 | 7 | 0 |
| TpMuguga_04g02495 | 8 | 0 |
| TpMuguga_04g02495 | 9 | 0 |
| TpMuguga_04g02495 | 10 | 0 |
| TpMuguga_04g02495 | 11 | 0 |
| TpMuguga_04g02495 | 12 | 0 |
| TpMuguga_04g00711 | 1 | 0 |
| TpMuguga_04g00713 | 1 | 0 |
| TpMuguga_04g00712 | 1 | 0.361111111 |
| TpMuguga_04g00712 | 2 | 0.277777778 |
| TpMuguga_04g00714 | 1 | 0 |
| TpMuguga_04g00714 | 2 | 0 |
| TpMuguga_04g00714 | 3 | 0 |
| TpMuguga_04g00714 | 4 | 0 |
| TpMuguga_04g00714 | 5 | 0 |
| TpMuguga_04g00714 | 6 | 0 |
| TpMuguga_04g00714 | 7 | 0 |
| TpMuguga_04g00715 | 1 | 0.006430868 |
| TpMuguga_04g00715 | 2 | 0.012861736 |
| TpMuguga_04g00717 | 1 | 0 |
| TpMuguga_04g00717 | 2 | 0 |
| TpMuguga_04g00716 | 1 | 0 |
| TpMuguga_04g00716 | 2 | 0 |
| TpMuguga_04g00716 | 3 | 0 |
| TpMuguga_04g00719 | 1 | 0 |
| TpMuguga_04g00719 | 2 | 0 |
| TpMuguga_04g00719 | 3 | 3.818181818 |
| TpMuguga_04g00719 | 4 | 1.272727273 |
| TpMuguga_04g00719 | 5 | 1.272727273 |

| | | |
|---|---|---|
| TpMuguga_04g00719 | 6 | 5.727272727 |
| TpMuguga_04g00720 | 1 | 0.254545455 |
| TpMuguga_04g00722 | 1 | 0.076433121 |
| TpMuguga_04g00722 | 2 | 0 |
| TpMuguga_04g00722 | 3 | 0 |
| TpMuguga_04g00722 | 4 | 0 |
| TpMuguga_04g00722 | 5 | 0 |
| TpMuguga_04g00723 | 1 | 1.43442623 |
| TpMuguga_04g00723 | 2 | 0.286885246 |
| TpMuguga_04g00723 | 3 | 0.327868852 |
| TpMuguga_04g00723 | 4 | 0.327868852 |
| TpMuguga_04g00724 | 1 | 0 |
| TpMuguga_04g00724 | 2 | 0.033857316 |
| TpMuguga_04g00724 | 3 | 2.471584039 |
| TpMuguga_04g00727 | 1 | 0.120171674 |
| TpMuguga_04g00726 | 1 | 0 |
| TpMuguga_04g00726 | 2 | 0 |
| TpMuguga_04g00728 | 1 | 0 |
| TpMuguga_04g00728 | 2 | 0.164705882 |
| TpMuguga_04g00728 | 3 | 0.658823529 |
| TpMuguga_04g00728 | 4 | 1.729411765 |
| TpMuguga_04g00728 | 5 | 1.070588235 |
| TpMuguga_04g00728 | 6 | 0 |
| TpMuguga_04g02660 | 1 | 0 |
| TpMuguga_04g02660 | 2 | 0 |
| TpMuguga_04g02660 | 3 | 0 |
| TpMuguga_04g02660 | 4 | 0 |
| TpMuguga_04g00732 | 1 | 1.376470588 |
| TpMuguga_04g00732 | 2 | 0.282352941 |
| TpMuguga_04g00733 | 1 | 1.238095238 |
| TpMuguga_04g00734 | 1 | 11.48717949 |
| TpMuguga_04g00734 | 2 | 0 |
| TpMuguga_04g00734 | 3 | 0 |
| TpMuguga_04g00734 | 4 | 0 |
| TpMuguga_04g00734 | 5 | 0 |
| TpMuguga_04g00734 | 6 | 0.41025641 |
| TpMuguga_04g00734 | 7 | 0 |
| TpMuguga_04g00737 | 1 | 0 |
| TpMuguga_04g00737 | 2 | 0.001433306 |
| TpMuguga_04g00737 | 3 | 0 |
| TpMuguga_04g00736 | 1 | 0.056198347 |
| TpMuguga_04g00738 | 1 | 0 |

| | | |
|---|---|---|
| TpMuguga_04g00738 | 2 | 0 |
| TpMuguga_04g00738 | 3 | 0 |
| TpMuguga_04g00738 | 4 | 2.571428571 |
| TpMuguga_04g00738 | 5 | 0 |
| TpMuguga_04g00738 | 6 | 0 |
| TpMuguga_04g00738 | 7 | 0 |
| TpMuguga_04g00738 | 8 | 0 |
| TpMuguga_04g00738 | 9 | 0 |
| TpMuguga_04g00738 | 10 | 6 |
| TpMuguga_04g00738 | 11 | 0 |
| TpMuguga_04g00740 | 1 | 0 |
| TpMuguga_04g00740 | 2 | 0 |
| TpMuguga_04g00740 | 3 | 0 |
| TpMuguga_04g00740 | 4 | 0 |
| TpMuguga_04g00740 | 5 | 0 |
| TpMuguga_04g00740 | 6 | 0 |
| TpMuguga_04g00740 | 7 | 0 |
| TpMuguga_04g02355 | 1 | 0 |
| TpMuguga_04g02355 | 2 | 0 |
| TpMuguga_04g02355 | 3 | 0 |
| TpMuguga_04g02355 | 4 | 0 |
| TpMuguga_04g00743 | 1 | 0 |
| TpMuguga_04g00744 | 1 | 2.181818182 |
| TpMuguga_04g00744 | 2 | 0 |
| TpMuguga_04g00744 | 3 | 0 |
| TpMuguga_04g00744 | 4 | 0 |
| TpMuguga_04g00744 | 5 | 0 |
| TpMuguga_04g00744 | 6 | 0 |
| TpMuguga_04g00744 | 7 | 0 |
| TpMuguga_04g00744 | 8 | 0 |
| TpMuguga_04g00744 | 9 | 0 |
| TpMuguga_04g00744 | 10 | 0 |
| TpMuguga_04g00744 | 11 | 0 |
| TpMuguga_04g02280 | 1 | 0 |
| TpMuguga_04g02280 | 2 | 0 |
| TpMuguga_04g02280 | 3 | 0 |
| TpMuguga_04g02280 | 4 | 0 |
| TpMuguga_04g02280 | 5 | 0 |
| TpMuguga_04g02280 | 6 | 0 |
| TpMuguga_04g02280 | 7 | 0 |
| TpMuguga_04g02280 | 8 | 0 |
| TpMuguga_04g02280 | 9 | 0 |

| | | |
|---|---|---|
| TpMuguga_04g02280 | 10 | 0 |
| TpMuguga_04g00746 | 1 | 0.334036568 |
| TpMuguga_04g00746 | 2 | 0.091420534 |
| TpMuguga_04g00746 | 3 | 0.042194093 |
| TpMuguga_04g00746 | 4 | 0.003516174 |
| TpMuguga_04g02665 | 1 | 0 |
| TpMuguga_04g02665 | 2 | 0 |
| TpMuguga_04g00748 | 1 | 0.128205128 |
| TpMuguga_04g00749 | 1 | 4.215568862 |
| TpMuguga_04g00750 | 1 | 0 |
| TpMuguga_04g00751 | 1 | 0 |
| TpMuguga_04g00751 | 2 | 0 |
| TpMuguga_04g00751 | 3 | 0 |
| TpMuguga_04g00751 | 4 | 0 |
| TpMuguga_04g00751 | 5 | 0 |
| TpMuguga_04g00751 | 6 | 0 |
| TpMuguga_04g00751 | 7 | 0 |
| TpMuguga_04g00751 | 8 | 0 |
| TpMuguga_04g00751 | 9 | 0 |
| TpMuguga_04g00751 | 10 | 0 |
| TpMuguga_04g00751 | 11 | 0 |
| TpMuguga_04g00751 | 12 | 0 |
| TpMuguga_04g00751 | 13 | 0 |
| TpMuguga_04g00751 | 14 | 0 |
| TpMuguga_04g00753 | 1 | 1.6 |
| TpMuguga_04g00753 | 2 | 1.3 |
| TpMuguga_04g00753 | 3 | 1.05 |
| TpMuguga_04g00753 | 4 | 0.3 |
| TpMuguga_04g00754 | 1 | 0 |
| TpMuguga_04g00754 | 2 | 0 |
| TpMuguga_04g00754 | 3 | 0 |
| TpMuguga_04g00754 | 4 | 0 |
| TpMuguga_04g00755 | 1 | 0 |
| TpMuguga_04g00755 | 2 | 0 |
| TpMuguga_04g00755 | 3 | 0 |
| TpMuguga_04g00755 | 4 | 0 |
| TpMuguga_04g00755 | 5 | 0 |
| TpMuguga_04g00755 | 6 | 0 |
| TpMuguga_04g00756 | 1 | 0 |
| TpMuguga_04g00756 | 2 | 0 |
| TpMuguga_04g00757 | 1 | 0 |
| TpMuguga_04g00757 | 2 | 0.267857143 |

| | | |
|---|---|---|
| TpMuguga_04g00757 | 3 | 0 |
| TpMuguga_04g00757 | 4 | 2.1875 |
| TpMuguga_04g00759 | 1 | 10.36486486 |
| TpMuguga_04g00759 | 2 | 2.810810811 |
| TpMuguga_04g00759 | 3 | 0 |
| TpMuguga_04g00759 | 4 | 0 |
| TpMuguga_04g00759 | 5 | 0 |
| TpMuguga_04g00759 | 6 | 0 |
| TpMuguga_04g00759 | 7 | 0 |
| TpMuguga_04g00759 | 8 | 0 |
| TpMuguga_04g00759 | 9 | 0 |
| TpMuguga_04g00759 | 10 | 0 |
| TpMuguga_04g00759 | 11 | 0 |
| TpMuguga_04g00759 | 12 | 0 |
| TpMuguga_04g00758 | 1 | 0.035683942 |
| TpMuguga_04g00758 | 2 | 0 |
| TpMuguga_04g00758 | 3 | 0 |
| TpMuguga_04g00758 | 4 | 0 |
| TpMuguga_04g00758 | 5 | 0 |
| TpMuguga_04g00758 | 6 | 0 |
| TpMuguga_04g00758 | 7 | 0 |
| TpMuguga_04g00758 | 8 | 0 |
| TpMuguga_04g00758 | 9 | 0 |
| TpMuguga_04g00758 | 10 | 0 |
| TpMuguga_04g00758 | 11 | 0 |
| TpMuguga_04g00758 | 12 | 0 |
| TpMuguga_04g00758 | 13 | 0.832625319 |
| TpMuguga_04g02070 | 1 | 0 |
| TpMuguga_04g02070 | 2 | 0 |
| TpMuguga_04g02070 | 3 | 0 |
| TpMuguga_04g02070 | 4 | 0 |
| TpMuguga_04g02070 | 5 | 0 |
| TpMuguga_04g02070 | 6 | 0 |
| TpMuguga_04g02070 | 7 | 0 |
| TpMuguga_04g02070 | 8 | 0 |
| TpMuguga_04g02070 | 9 | 0 |
| TpMuguga_04g02070 | 10 | 0 |
| TpMuguga_04g02310 | 1 | 0 |
| TpMuguga_04g02310 | 2 | 8 |
| TpMuguga_04g02310 | 3 | 0 |
| TpMuguga_04g02310 | 4 | 0 |
| TpMuguga_04g02310 | 5 | 4 |

| | | |
|---|---|---|
| TpMuguga_04g00764 | 1 | 0 |
| TpMuguga_04g00764 | 2 | 0 |
| TpMuguga_04g00765 | 1 | 0 |
| TpMuguga_04g00765 | 2 | 0 |
| TpMuguga_04g00765 | 3 | 0 |
| TpMuguga_04g00765 | 4 | 0 |
| TpMuguga_04g00765 | 5 | 0 |
| TpMuguga_04g00765 | 6 | 0 |
| TpMuguga_04g00765 | 7 | 0 |
| TpMuguga_04g00765 | 8 | 0 |
| TpMuguga_04g00765 | 9 | 0 |
| TpMuguga_04g00765 | 10 | 0 |
| TpMuguga_04g00765 | 11 | 0 |
| TpMuguga_04g00765 | 12 | 0 |
| TpMuguga_04g00766 | 1 | 0 |
| TpMuguga_04g00766 | 2 | 1.227272727 |
| TpMuguga_04g00766 | 3 | 0 |
| TpMuguga_04g00766 | 4 | 0 |
| TpMuguga_04g00766 | 5 | 0 |
| TpMuguga_04g00767 | 1 | 0 |
| TpMuguga_04g00767 | 2 | 0 |
| TpMuguga_04g00767 | 3 | 0 |
| TpMuguga_04g00767 | 4 | 0 |
| TpMuguga_04g00767 | 5 | 0 |
| TpMuguga_04g00767 | 6 | 0 |
| TpMuguga_04g02325 | 1 | 0 |
| TpMuguga_04g02325 | 2 | 0 |
| TpMuguga_04g02325 | 3 | 0 |
| TpMuguga_04g02325 | 4 | 9.333333333 |
| TpMuguga_04g02325 | 5 | 0 |
| TpMuguga_04g02325 | 6 | 0 |
| TpMuguga_04g00770 | 1 | 0 |
| TpMuguga_04g00770 | 2 | 0 |
| TpMuguga_04g00770 | 3 | 0 |
| TpMuguga_04g00770 | 4 | 0 |
| TpMuguga_04g00770 | 5 | 1.333333333 |
| TpMuguga_04g00770 | 6 | 8.666666667 |
| TpMuguga_04g00770 | 7 | 10 |
| TpMuguga_04g02635 | 1 | 0 |
| TpMuguga_04g02635 | 2 | 0 |
| TpMuguga_04g02635 | 3 | 0 |
| TpMuguga_04g02635 | 4 | 0 |

| | | |
|---|---|---|
| TpMuguga_04g02635 | 5 | 4 |
| TpMuguga_04g00771 | 1 | 3 |
| TpMuguga_04g00771 | 2 | 0 |
| TpMuguga_04g00772 | 1 | 28.75 |
| TpMuguga_04g00772 | 2 | 0 |
| TpMuguga_04g00772 | 3 | 0 |
| TpMuguga_04g00772 | 4 | 0 |
| TpMuguga_04g00772 | 5 | 0 |
| TpMuguga_04g00772 | 6 | 0 |
| TpMuguga_04g00772 | 7 | 0 |
| TpMuguga_04g00772 | 8 | 2.5 |
| TpMuguga_04g00772 | 9 | 0 |
| TpMuguga_04g00774 | 1 | 0 |
| TpMuguga_04g00774 | 2 | 12 |
| TpMuguga_04g00774 | 3 | 0 |
| TpMuguga_04g00774 | 4 | 0 |
| TpMuguga_04g00774 | 5 | 0 |
| TpMuguga_04g00774 | 6 | 0 |
| TpMuguga_04g00774 | 7 | 0 |
| TpMuguga_04g00774 | 8 | 0 |
| TpMuguga_04g00774 | 9 | 0 |
| TpMuguga_04g00774 | 10 | 0 |
| TpMuguga_04g00774 | 11 | 0 |
| TpMuguga_04g00775 | 1 | 0 |
| TpMuguga_04g00775 | 2 | 0 |
| TpMuguga_04g00776 | 1 | 0 |
| TpMuguga_04g00776 | 2 | 0 |
| TpMuguga_04g00776 | 3 | 0 |
| TpMuguga_04g00776 | 4 | 0 |
| TpMuguga_04g00776 | 5 | 0 |
| TpMuguga_04g00776 | 6 | 0 |
| TpMuguga_04g00776 | 7 | 0 |
| TpMuguga_04g00776 | 8 | 0 |
| TpMuguga_04g02460 | 1 | 0 |
| TpMuguga_04g02460 | 2 | 0 |
| TpMuguga_04g02460 | 3 | 0 |
| TpMuguga_04g02460 | 4 | 0 |
| TpMuguga_04g02460 | 5 | 0 |
| TpMuguga_04g02460 | 6 | 0 |
| TpMuguga_04g02460 | 7 | 0 |
| TpMuguga_04g02460 | 8 | 0 |
| TpMuguga_04g02460 | 9 | 0 |

| | | |
|---|---|---|
| TpMuguga_04g02460 | 10 | 0 |
| TpMuguga_04g02460 | 11 | 0.028382214 |
| TpMuguga_04g02460 | 12 | 0.028382214 |
| TpMuguga_04g02460 | 13 | 0.283822138 |
| TpMuguga_04g02460 | 14 | 11.82119205 |
| TpMuguga_04g00780 | 1 | 0 |
| TpMuguga_04g00780 | 2 | 0 |
| TpMuguga_04g00780 | 3 | 0 |
| TpMuguga_04g00780 | 4 | 0 |
| TpMuguga_04g00780 | 5 | 0 |
| TpMuguga_04g00780 | 6 | 32.5 |
| TpMuguga_04g00780 | 7 | 0 |
| TpMuguga_04g00780 | 8 | 0 |
| TpMuguga_04g00780 | 9 | 0 |
| TpMuguga_04g00780 | 10 | 0 |
| TpMuguga_04g00780 | 11 | 0 |
| TpMuguga_04g00780 | 12 | 0 |
| TpMuguga_04g02110 | 1 | 0 |
| TpMuguga_04g02110 | 2 | 0 |
| TpMuguga_04g02110 | 3 | 0 |
| TpMuguga_04g00781 | 1 | 3.461538462 |
| TpMuguga_04g00781 | 2 | 0 |
| TpMuguga_04g00781 | 3 | 0 |
| TpMuguga_04g00781 | 4 | 0 |
| TpMuguga_04g00781 | 5 | 0 |
| TpMuguga_04g00781 | 6 | 0 |
| TpMuguga_04g00781 | 7 | 0 |
| TpMuguga_04g00781 | 8 | 0 |
| TpMuguga_04g00782 | 1 | 0 |
| TpMuguga_04g00783 | 1 | 0 |
| TpMuguga_04g00783 | 2 | 0 |
| TpMuguga_04g00783 | 3 | 0 |
| TpMuguga_04g00783 | 4 | 0 |
| TpMuguga_04g00783 | 5 | 0 |
| TpMuguga_04g00783 | 6 | 0 |
| TpMuguga_04g00786 | 1 | 0.010989011 |
| TpMuguga_04g00787 | 1 | 0 |
| TpMuguga_04g00787 | 2 | 6.1875 |
| TpMuguga_04g00787 | 3 | 0 |
| TpMuguga_04g00787 | 4 | 0 |
| TpMuguga_04g00787 | 5 | 0 |
| TpMuguga_04g00787 | 6 | 1.125 |

| | | |
|---|---|---|
| TpMuguga_04g00787 | 7 | 0 |
| TpMuguga_04g00787 | 8 | 0 |
| TpMuguga_04g00788 | 1 | 0 |
| TpMuguga_04g00788 | 2 | 1.407506702 |
| TpMuguga_04g00788 | 3 | 1.849865952 |
| TpMuguga_04g00788 | 4 | 1.970509383 |
| TpMuguga_04g00788 | 5 | 2.010723861 |
| TpMuguga_04g00788 | 6 | 0.361930295 |
| TpMuguga_04g00788 | 7 | 0.804289544 |
| TpMuguga_04g00788 | 8 | 3.337801609 |
| TpMuguga_04g00788 | 9 | 4.101876676 |
| TpMuguga_04g00788 | 10 | 4.021447721 |
| TpMuguga_04g00788 | 11 | 2.935656836 |
| TpMuguga_04g00788 | 12 | 2.131367292 |
| TpMuguga_04g00788 | 13 | 2.45308311 |
| TpMuguga_04g00788 | 14 | 5.348525469 |
| TpMuguga_04g00789 | 1 | 0 |
| TpMuguga_04g00789 | 2 | 0 |
| TpMuguga_04g00789 | 3 | 0 |
| TpMuguga_04g00789 | 4 | 0 |
| TpMuguga_04g00789 | 5 | 0 |
| TpMuguga_04g00789 | 6 | 8 |
| TpMuguga_04g00789 | 7 | 0 |
| TpMuguga_04g00791 | 1 | 0 |
| TpMuguga_04g00791 | 2 | 0 |
| TpMuguga_04g00791 | 3 | 0.954545455 |
| TpMuguga_04g00791 | 4 | 0 |
| TpMuguga_04g00791 | 5 | 0 |
| TpMuguga_04g00791 | 6 | 0 |
| TpMuguga_04g02610 | 1 | 0 |
| TpMuguga_04g02610 | 2 | 2.173913043 |
| TpMuguga_04g02610 | 3 | 0 |
| TpMuguga_04g02610 | 4 | 0 |
| TpMuguga_04g02610 | 5 | 0.434782609 |
| TpMuguga_04g02610 | 6 | 0.869565217 |
| TpMuguga_04g02610 | 7 | 0 |
| TpMuguga_04g02610 | 8 | 0 |
| TpMuguga_04g02610 | 9 | 0.869565217 |
| TpMuguga_04g00926 | 1 | 0 |
| TpMuguga_04g00926 | 2 | 0 |
| TpMuguga_04g00926 | 3 | 15 |
| TpMuguga_04g00926 | 4 | 6.666666667 |

| | | |
|---|---|---|
| TpMuguga_04g00794 | 1 | 0.432432432 |
| TpMuguga_04g00794 | 2 | 0.540540541 |
| TpMuguga_04g00794 | 3 | 0.810810811 |
| TpMuguga_04g00796 | 1 | 0 |
| TpMuguga_04g00796 | 2 | 0 |
| TpMuguga_04g00796 | 3 | 0 |
| TpMuguga_04g00796 | 4 | 0 |
| TpMuguga_04g00796 | 5 | 0 |
| TpMuguga_04g00796 | 6 | 0 |
| TpMuguga_04g00796 | 7 | 0 |
| TpMuguga_04g00796 | 8 | 0 |
| TpMuguga_04g00796 | 9 | 0 |
| TpMuguga_04g00796 | 10 | 0 |
| TpMuguga_04g00796 | 11 | 0 |
| TpMuguga_04g00796 | 12 | 0 |
| TpMuguga_04g00796 | 13 | 0 |
| TpMuguga_04g00796 | 14 | 0 |
| TpMuguga_04g00796 | 15 | 0 |
| TpMuguga_04g00796 | 16 | 0 |
| TpMuguga_04g00796 | 17 | 0 |
| TpMuguga_04g00796 | 18 | 0 |
| TpMuguga_04g00796 | 19 | 0 |
| TpMuguga_04g00796 | 20 | 0 |
| TpMuguga_04g00796 | 21 | 0 |
| TpMuguga_04g00796 | 22 | 0 |
| TpMuguga_04g00796 | 23 | 0 |
| TpMuguga_04g00796 | 24 | 0 |
| TpMuguga_04g00796 | 25 | 0 |
| TpMuguga_04g00796 | 26 | 0 |
| TpMuguga_04g00795 | 1 | 0 |
| TpMuguga_04g00795 | 2 | 0 |
| TpMuguga_04g00795 | 3 | 0 |
| TpMuguga_04g00795 | 4 | 0 |
| TpMuguga_04g00795 | 5 | 0 |
| TpMuguga_04g00795 | 6 | 0 |
| TpMuguga_04g00797 | 1 | 0 |
| TpMuguga_04g00797 | 2 | 0 |
| TpMuguga_04g00797 | 3 | 0 |
| TpMuguga_04g00797 | 4 | 0 |
| TpMuguga_04g00797 | 5 | 0 |
| TpMuguga_04g00797 | 6 | 0 |
| TpMuguga_04g00797 | 7 | 0 |

| | | |
|---|---|---|
| TpMuguga_04g00797 | 8 | 0 |
| TpMuguga_04g00797 | 9 | 0 |
| TpMuguga_04g00798 | 1 | 0 |
| TpMuguga_04g00798 | 2 | 0.571428571 |
| TpMuguga_04g00798 | 3 | 4.428571429 |
| TpMuguga_04g00799 | 1 | 0 |
| TpMuguga_04g00800 | 1 | 0 |
| TpMuguga_04g00800 | 2 | 0 |
| TpMuguga_04g00800 | 3 | 0 |
| TpMuguga_04g00800 | 4 | 0 |
| TpMuguga_04g00800 | 5 | 0 |
| TpMuguga_04g00800 | 6 | 0 |
| TpMuguga_04g00800 | 7 | 0 |
| TpMuguga_04g00800 | 8 | 4.404255319 |
| TpMuguga_04g00801 | 1 | 0 |
| TpMuguga_04g00802 | 1 | 0 |
| TpMuguga_04g00802 | 2 | 0 |
| TpMuguga_04g00802 | 3 | 0 |
| TpMuguga_04g00802 | 4 | 0 |
| TpMuguga_04g00803 | 1 | 0.409090909 |
| TpMuguga_04g00803 | 2 | 0 |
| TpMuguga_04g00803 | 3 | 0 |
| TpMuguga_04g00803 | 4 | 0 |
| TpMuguga_04g00803 | 5 | 0.204545455 |
| TpMuguga_04g00803 | 6 | 0 |
| TpMuguga_04g00803 | 7 | 0 |
| TpMuguga_04g00803 | 8 | 0 |
| TpMuguga_04g02605 | 1 | 0 |
| TpMuguga_04g02605 | 2 | 0 |
| TpMuguga_04g02605 | 3 | 0 |
| TpMuguga_04g02605 | 4 | 0 |
| TpMuguga_04g02605 | 5 | 0 |
| TpMuguga_04g02605 | 6 | 0 |
| TpMuguga_04g02605 | 7 | 0 |
| TpMuguga_04g02605 | 8 | 0 |
| TpMuguga_04g02605 | 9 | 0 |
| TpMuguga_04g00806 | 1 | 0 |
| TpMuguga_04g00807 | 1 | 0.567567568 |
| TpMuguga_04g00807 | 2 | 0 |
| TpMuguga_04g00807 | 3 | 1.513513514 |
| TpMuguga_04g00807 | 4 | 3.972972973 |
| TpMuguga_04g00807 | 5 | 0.189189189 |

| | | |
|---|---|---|
| TpMuguga_04g00807 | 6 | 0 |
| TpMuguga_04g00808 | 1 | 0 |
| TpMuguga_04g00809 | 1 | 0.049886621 |
| TpMuguga_04g00809 | 2 | 0 |
| TpMuguga_04g00809 | 3 | 0 |
| TpMuguga_04g00809 | 4 | 0 |
| TpMuguga_04g00809 | 5 | 0 |
| TpMuguga_04g00809 | 6 | 0.199546485 |
| TpMuguga_04g00809 | 7 | 0 |
| TpMuguga_04g00809 | 8 | 0 |
| TpMuguga_04g00809 | 9 | 0 |
| TpMuguga_04g00809 | 10 | 0.074829932 |
| TpMuguga_04g00810 | 1 | 0 |
| TpMuguga_04g00810 | 2 | 0 |
| TpMuguga_04g00810 | 3 | 0 |
| TpMuguga_04g00810 | 4 | 0 |
| TpMuguga_04g00810 | 5 | 0 |
| TpMuguga_04g00810 | 6 | 0 |
| TpMuguga_04g00810 | 7 | 0 |
| TpMuguga_04g00810 | 8 | 13.2 |
| TpMuguga_04g00810 | 9 | 0 |
| TpMuguga_04g00810 | 10 | 13.2 |
| TpMuguga_04g02235 | 1 | 0 |
| TpMuguga_04g02235 | 2 | 0 |
| TpMuguga_04g02235 | 3 | 0 |
| TpMuguga_04g02235 | 4 | 0 |
| TpMuguga_04g02235 | 5 | 0 |
| TpMuguga_04g02125 | 1 | 0 |
| TpMuguga_04g02125 | 2 | 0 |
| TpMuguga_04g02125 | 3 | 0.114649682 |
| TpMuguga_04g02125 | 4 | 0 |
| TpMuguga_04g02125 | 5 | 2.178343949 |
| TpMuguga_04g02125 | 6 | 0.573248408 |
| TpMuguga_04g02125 | 7 | 0 |
| TpMuguga_04g02125 | 8 | 0.114649682 |
| TpMuguga_04g02125 | 9 | 1.146496815 |
| TpMuguga_04g02125 | 10 | 0 |
| TpMuguga_04g02125 | 11 | 0 |
| TpMuguga_04g02125 | 12 | 0 |
| TpMuguga_04g02125 | 13 | 0.114649682 |
| TpMuguga_04g02125 | 14 | 0 |
| TpMuguga_04g02125 | 15 | 0 |

| | | |
|---|---|---|
| TpMuguga_04g02125 | 16 | 0 |
| TpMuguga_04g02125 | 17 | 11.92356688 |
| TpMuguga_04g00815 | 1 | 2.134529148 |
| TpMuguga_04g00815 | 2 | 0.188340807 |
| TpMuguga_04g00815 | 3 | 1.820627803 |
| TpMuguga_04g00815 | 4 | 0.439461883 |
| TpMuguga_04g00815 | 5 | 0.094170404 |
| TpMuguga_04g00815 | 6 | 0.408071749 |
| TpMuguga_04g00816 | 1 | 0 |
| TpMuguga_04g00816 | 2 | 0 |
| TpMuguga_04g00816 | 3 | 0 |
| TpMuguga_04g00816 | 4 | 0 |
| TpMuguga_04g00816 | 5 | 0 |
| TpMuguga_04g00816 | 6 | 0 |
| TpMuguga_04g00816 | 7 | 0 |
| TpMuguga_04g00816 | 8 | 0 |
| TpMuguga_04g00816 | 9 | 0 |
| TpMuguga_04g00816 | 10 | 0 |
| TpMuguga_04g00816 | 11 | 0 |
| TpMuguga_04g00817 | 1 | 0 |
| TpMuguga_04g00817 | 2 | 0 |
| TpMuguga_04g00817 | 3 | 0 |
| TpMuguga_04g00818 | 1 | 0 |
| TpMuguga_04g00818 | 2 | 0 |
| TpMuguga_04g00818 | 3 | 0 |
| TpMuguga_04g00818 | 4 | 0 |
| TpMuguga_04g00818 | 5 | 0.571428571 |
| TpMuguga_04g00818 | 6 | 0 |
| TpMuguga_04g00818 | 7 | 0 |
| TpMuguga_04g00821 | 1 | 0 |
| TpMuguga_04g00821 | 2 | 0 |
| TpMuguga_04g00821 | 3 | 0 |
| TpMuguga_04g00821 | 4 | 0 |
| TpMuguga_04g00821 | 5 | 0 |
| TpMuguga_04g00822 | 1 | 0 |
| TpMuguga_04g00823 | 1 | 0 |
| TpMuguga_04g00823 | 2 | 0 |
| TpMuguga_04g00823 | 3 | 0 |
| TpMuguga_04g00823 | 4 | 0 |
| TpMuguga_04g00823 | 5 | 0 |
| TpMuguga_04g00823 | 6 | 0 |
| TpMuguga_04g00823 | 7 | 0 |

| | | |
|---|---|---|
| TpMuguga_04g00824 | 1 | 0 |
| TpMuguga_04g00824 | 2 | 0.9 |
| TpMuguga_04g00825 | 1 | 0.734693878 |
| TpMuguga_04g00825 | 2 | 0 |
| TpMuguga_04g00825 | 3 | 0 |
| TpMuguga_04g00827 | 1 | 0.463917526 |
| TpMuguga_04g00827 | 2 | 0.216494845 |
| TpMuguga_04g00826 | 1 | 0 |
| TpMuguga_04g00826 | 2 | 0 |
| TpMuguga_04g00826 | 3 | 0 |
| TpMuguga_04g00826 | 4 | 0 |
| TpMuguga_04g00826 | 5 | 0 |
| TpMuguga_04g00826 | 6 | 0 |
| TpMuguga_04g00826 | 7 | 0 |
| TpMuguga_04g00828 | 1 | 0.058139535 |
| TpMuguga_04g00828 | 2 | 0.697674419 |
| TpMuguga_04g00828 | 3 | 3.430232558 |
| TpMuguga_04g00828 | 4 | 1.395348837 |
| TpMuguga_04g00829 | 1 | 0.523364486 |
| TpMuguga_04g00829 | 2 | 0 |
| TpMuguga_04g00829 | 3 | 0 |
| TpMuguga_04g00829 | 4 | 0 |
| TpMuguga_04g00829 | 5 | 0 |
| TpMuguga_04g00829 | 6 | 0 |
| TpMuguga_04g00829 | 7 | 0 |
| TpMuguga_04g00832 | 1 | 3.689655172 |
| TpMuguga_04g00832 | 2 | 3.413793103 |
| TpMuguga_04g00832 | 3 | 0.379310345 |
| TpMuguga_04g00831 | 1 | 0 |
| TpMuguga_04g00831 | 2 | 0 |
| TpMuguga_04g00834 | 1 | 0 |
| TpMuguga_04g00834 | 2 | 1.108433735 |
| TpMuguga_04g00834 | 3 | 0 |
| TpMuguga_04g00836 | 1 | 0 |
| TpMuguga_04g00838 | 1 | 0 |
| TpMuguga_04g00838 | 2 | 0 |
| TpMuguga_04g00838 | 3 | 0 |
| TpMuguga_04g00838 | 4 | 0 |
| TpMuguga_04g00838 | 5 | 0 |
| TpMuguga_04g00838 | 6 | 0 |
| TpMuguga_04g00838 | 7 | 0 |
| TpMuguga_04g00837 | 1 | 0 |

| | | |
|---|---|---|
| TpMuguga_04g00840 | 1 | 0 |
| TpMuguga_04g00840 | 2 | 0 |
| TpMuguga_04g00841 | 1 | 0 |
| TpMuguga_04g00841 | 2 | 0 |
| TpMuguga_04g00842 | 1 | 0 |
| TpMuguga_04g00842 | 2 | 0 |
| TpMuguga_04g00842 | 3 | 0 |
| TpMuguga_04g00842 | 4 | 0 |
| TpMuguga_04g00842 | 5 | 12 |
| TpMuguga_04g02475 | 1 | 0 |
| TpMuguga_04g02475 | 2 | 0 |
| TpMuguga_04g02475 | 3 | 0 |
| TpMuguga_04g02475 | 4 | 0 |
| TpMuguga_04g02475 | 5 | 0 |
| TpMuguga_04g02475 | 6 | 0 |
| TpMuguga_04g02475 | 7 | 0 |
| TpMuguga_04g02335 | 1 | 0 |
| TpMuguga_04g02335 | 2 | 0 |
| TpMuguga_04g02335 | 3 | 0 |
| TpMuguga_04g02335 | 4 | 5 |
| TpMuguga_04g00844 | 1 | 2 |
| TpMuguga_04g00845 | 1 | 0 |
| TpMuguga_04g00845 | 2 | 0 |
| TpMuguga_04g00845 | 3 | 0 |
| TpMuguga_04g00847 | 1 | 0.846153846 |
| TpMuguga_04g00847 | 2 | 0 |
| TpMuguga_04g00847 | 3 | 0 |
| TpMuguga_04g00847 | 4 | 0 |
| TpMuguga_04g00847 | 5 | 0 |
| TpMuguga_04g00847 | 6 | 0 |
| TpMuguga_04g00847 | 7 | 0 |
| TpMuguga_04g00847 | 8 | 0 |
| TpMuguga_04g00847 | 9 | 0 |
| TpMuguga_04g00847 | 10 | 0 |
| TpMuguga_04g00848 | 1 | 0 |
| TpMuguga_04g00848 | 2 | 0 |
| TpMuguga_04g00848 | 3 | 0 |
| TpMuguga_04g00848 | 4 | 0 |
| TpMuguga_04g00848 | 5 | 0 |
| TpMuguga_04g00848 | 6 | 0 |
| TpMuguga_04g00851 | 1 | 1.220183486 |
| TpMuguga_04g00851 | 2 | 0.256880734 |

| | | |
|---|---|---|
| TpMuguga_04g00851 | 3 | 0.642201835 |
| TpMuguga_04g00851 | 4 | 0.385321101 |
| TpMuguga_04g00851 | 5 | 1.220183486 |
| TpMuguga_04g00851 | 6 | 2.119266055 |
| TpMuguga_04g00850 | 1 | 0 |
| TpMuguga_04g00850 | 2 | 0 |
| TpMuguga_04g00850 | 3 | 0 |
| TpMuguga_04g00850 | 4 | 0 |
| TpMuguga_04g00850 | 5 | 0 |
| TpMuguga_04g00850 | 6 | 0 |
| TpMuguga_04g00850 | 7 | 0 |
| TpMuguga_04g00850 | 8 | 0 |
| TpMuguga_04g00850 | 9 | 0 |
| TpMuguga_04g00850 | 10 | 3.723076923 |
| TpMuguga_04g02585 | 1 | 1.180327869 |
| TpMuguga_04g02585 | 2 | 0 |
| TpMuguga_04g02585 | 3 | 0 |
| TpMuguga_04g02585 | 4 | 0 |
| TpMuguga_04g02585 | 5 | 0 |
| TpMuguga_04g02585 | 6 | 0 |
| TpMuguga_04g02585 | 7 | 0 |
| TpMuguga_04g02585 | 8 | 0 |
| TpMuguga_04g00855 | 1 | 0.495 |
| TpMuguga_04g00855 | 2 | 0.765 |
| TpMuguga_04g00855 | 3 | 0.765 |
| TpMuguga_04g00855 | 4 | 1.17 |
| TpMuguga_04g00855 | 5 | 1.98 |
| TpMuguga_04g00855 | 6 | 1.08 |
| TpMuguga_04g00855 | 7 | 0.9 |
| TpMuguga_04g00855 | 8 | 1.845 |
| TpMuguga_04g02265 | 1 | 0 |
| TpMuguga_04g02265 | 2 | 0 |
| TpMuguga_04g02265 | 3 | 0 |
| TpMuguga_04g02265 | 4 | 0 |
| TpMuguga_04g02265 | 5 | 0 |
| TpMuguga_04g02265 | 6 | 0 |
| TpMuguga_04g02265 | 7 | 0 |
| TpMuguga_04g02265 | 8 | 0 |
| TpMuguga_04g02265 | 9 | 0 |
| TpMuguga_04g00859 | 1 | 1.379310345 |
| TpMuguga_04g00859 | 2 | 0.689655172 |
| TpMuguga_04g00859 | 3 | 0 |

| | | |
|---|---|---|
| TpMuguga_04g00859 | 4 | 0 |
| TpMuguga_04g00859 | 5 | 0 |
| TpMuguga_04g00859 | 6 | 0 |
| TpMuguga_04g00859 | 7 | 0 |
| TpMuguga_04g00859 | 8 | 0.689655172 |
| TpMuguga_04g00859 | 9 | 0 |
| TpMuguga_04g02340 | 1 | 0 |
| TpMuguga_04g02340 | 2 | 0 |
| TpMuguga_04g02415 | 1 | 0 |
| TpMuguga_04g02415 | 2 | 0 |
| TpMuguga_04g02415 | 3 | 0 |
| TpMuguga_04g02415 | 4 | 0 |
| TpMuguga_04g02415 | 5 | 0 |
| TpMuguga_04g00866 | 1 | 0.312056738 |
| TpMuguga_04g00866 | 2 | 0 |
| TpMuguga_04g00866 | 3 | 0 |
| TpMuguga_04g00869 | 1 | 0 |
| TpMuguga_04g00869 | 2 | 0 |
| TpMuguga_04g00869 | 3 | 9 |
| TpMuguga_04g00869 | 4 | 0 |
| TpMuguga_04g00869 | 5 | 14.4 |
| TpMuguga_04g00869 | 6 | 1.8 |
| TpMuguga_04g00869 | 7 | 21.6 |
| TpMuguga_04g00869 | 8 | 0 |
| TpMuguga_04g00870 | 1 | 0.333333333 |
| TpMuguga_04g00870 | 2 | 2 |
| TpMuguga_04g00870 | 3 | 3.333333333 |
| TpMuguga_04g00871 | 1 | 0 |
| TpMuguga_04g00871 | 2 | 0 |
| TpMuguga_04g00871 | 3 | 2.6 |
| TpMuguga_04g00871 | 4 | 0 |
| TpMuguga_04g00871 | 5 | 0 |
| TpMuguga_04g00871 | 6 | 0 |
| TpMuguga_04g00871 | 7 | 0 |
| TpMuguga_04g00871 | 8 | 0 |
| TpMuguga_04g00871 | 9 | 0 |
| TpMuguga_04g00871 | 10 | 0 |
| TpMuguga_04g00871 | 11 | 0 |
| TpMuguga_04g00871 | 12 | 0 |
| TpMuguga_04g00873 | 1 | 0 |
| TpMuguga_04g00873 | 2 | 0 |
| TpMuguga_04g00873 | 3 | 0 |

| | | |
|---|---|---|
| TpMuguga_04g00873 | 4 | 0 |
| TpMuguga_04g00873 | 5 | 0 |
| TpMuguga_04g00873 | 6 | 0 |
| TpMuguga_04g00873 | 7 | 0 |
| TpMuguga_04g00873 | 8 | 0 |
| TpMuguga_04g00873 | 9 | 0 |
| TpMuguga_04g00873 | 10 | 0 |
| TpMuguga_04g00873 | 11 | 0 |
| TpMuguga_04g02320 | 1 | 0 |
| TpMuguga_04g02320 | 2 | 0 |
| TpMuguga_04g02320 | 3 | 0 |
| TpMuguga_04g02435 | 1 | 0 |
| TpMuguga_04g02435 | 2 | 0 |
| TpMuguga_04g02435 | 3 | 0 |
| TpMuguga_04g02435 | 4 | 0 |
| TpMuguga_04g02435 | 5 | 0 |
| TpMuguga_04g02435 | 6 | 0 |
| TpMuguga_04g00878 | 1 | 0 |
| TpMuguga_04g00878 | 2 | 1.018867925 |
| TpMuguga_04g00878 | 3 | 2.490566038 |
| TpMuguga_04g00878 | 4 | 0.905660377 |
| TpMuguga_04g00878 | 5 | 0 |
| TpMuguga_04g00880 | 1 | 0 |
| TpMuguga_04g00879 | 1 | 0 |
| TpMuguga_04g00879 | 2 | 0 |
| TpMuguga_04g00881 | 1 | 0 |
| TpMuguga_04g00881 | 2 | 0 |
| TpMuguga_04g00881 | 3 | 0 |
| TpMuguga_04g00881 | 4 | 1.384615385 |
| TpMuguga_04g00881 | 5 | 4.153846154 |
| TpMuguga_04g00881 | 6 | 0 |
| TpMuguga_04g00881 | 7 | 0.692307692 |
| TpMuguga_04g00881 | 8 | 0 |
| TpMuguga_04g00882 | 1 | 0 |
| TpMuguga_04g00882 | 2 | 0 |
| TpMuguga_04g00882 | 3 | 0 |
| TpMuguga_04g00882 | 4 | 0 |
| TpMuguga_04g00882 | 5 | 0 |
| TpMuguga_04g00882 | 6 | 0 |
| TpMuguga_04g00882 | 7 | 0 |
| TpMuguga_04g00883 | 1 | 0 |
| TpMuguga_04g00883 | 2 | 0 |

| | | |
|---|---|---|
| TpMuguga_04g00884 | 1 | 0 |
| TpMuguga_04g00884 | 2 | 0 |
| TpMuguga_04g00884 | 3 | 0 |
| TpMuguga_04g00884 | 4 | 0 |
| TpMuguga_04g00884 | 5 | 0 |
| TpMuguga_04g00884 | 6 | 3.80952381 |
| TpMuguga_04g00884 | 7 | 3.047619048 |
| TpMuguga_04g00886 | 1 | 0 |
| TpMuguga_04g02680 | 1 | 2.25 |
| TpMuguga_04g02680 | 2 | 0 |
| TpMuguga_04g02680 | 3 | 0 |
| TpMuguga_04g02680 | 4 | 0 |
| TpMuguga_04g02680 | 5 | 0 |
| TpMuguga_04g02680 | 6 | 0 |
| TpMuguga_04g02680 | 7 | 0 |
| TpMuguga_04g02680 | 8 | 0 |
| TpMuguga_04g02680 | 9 | 0 |
| TpMuguga_04g02680 | 10 | 0 |
| TpMuguga_04g02680 | 11 | 0 |
| TpMuguga_04g00891 | 1 | 0 |
| TpMuguga_04g00892 | 1 | 1.041666667 |
| TpMuguga_04g00892 | 2 | 0.958333333 |
| TpMuguga_04g00893 | 1 | 0 |
| TpMuguga_04g00893 | 2 | 0 |
| TpMuguga_04g00893 | 3 | 0.063241107 |
| TpMuguga_04g00894 | 1 | 0 |
| TpMuguga_04g00894 | 2 | 0 |
| TpMuguga_04g00894 | 3 | 0 |
| TpMuguga_04g00894 | 4 | 0 |
| TpMuguga_04g02035 | 1 | 0 |
| TpMuguga_04g02035 | 2 | 0 |
| TpMuguga_04g02035 | 3 | 20 |
| TpMuguga_04g02030 | 1 | 0 |
| TpMuguga_04g02030 | 2 | 0 |
| TpMuguga_04g02030 | 3 | 0 |
| TpMuguga_04g02030 | 4 | 0 |
| TpMuguga_04g02030 | 5 | 0 |
| TpMuguga_04g00896 | 1 | 0 |
| TpMuguga_04g00898 | 1 | 0 |
| TpMuguga_04g00898 | 2 | 0 |
| TpMuguga_04g00898 | 3 | 0 |
| TpMuguga_04g00898 | 4 | 0 |

| | | |
|---|---|---|
| TpMuguga_04g00898 | 5 | 0 |
| TpMuguga_04g00899 | 1 | 0.222222222 |
| TpMuguga_04g02420 | 1 | 0.406779661 |
| TpMuguga_04g02420 | 2 | 0.06779661 |
| TpMuguga_04g02420 | 3 | 0.406779661 |
| TpMuguga_04g00901 | 1 | 0 |
| TpMuguga_04g00904 | 1 | 0.456140351 |
| TpMuguga_04g00908 | 1 | 0.142857143 |
| TpMuguga_04g00908 | 2 | 0 |
| TpMuguga_04g00908 | 3 | 1.428571429 |
| TpMuguga_04g00909 | 1 | 0 |
| TpMuguga_04g00909 | 2 | 0 |
| TpMuguga_04g02305 | 1 | 0.785714286 |
| TpMuguga_04g02600 | 1 | 0 |
| TpMuguga_04g02600 | 2 | 0 |
| TpMuguga_04g00911 | 1 | 0 |
| TpMuguga_04g00911 | 2 | 0 |
| TpMuguga_04g00911 | 3 | 2 |
| TpMuguga_04g00911 | 4 | 0 |
| TpMuguga_04g00911 | 5 | 0.333333333 |
| TpMuguga_04g00912 | 1 | 0 |
| TpMuguga_04g00912 | 2 | 0 |
| TpMuguga_04g00913 | 1 | 0 |
| TpMuguga_04g00913 | 2 | 0 |
| TpMuguga_04g00913 | 3 | 0 |
| TpMuguga_04g00922 | 1 | 1.644444444 |
| TpMuguga_03g00012 | 1 | 0.05027933 |
| TpMuguga_03g00012 | 2 | 0 |
| TpMuguga_03g00012 | 3 | 0 |
| TpMuguga_03g00012 | 4 | 0 |
| TpMuguga_03g00012 | 5 | 0 |
| TpMuguga_03g00012 | 6 | 0.05027933 |
| TpMuguga_03g00012 | 7 | 0 |
| TpMuguga_03g00012 | 8 | 3.569832402 |
| TpMuguga_03g00011 | 1 | 1.512465374 |
| TpMuguga_03g00013 | 1 | 4.661157025 |
| TpMuguga_03g00013 | 2 | 1.090909091 |
| TpMuguga_03g00013 | 3 | 1.140495868 |
| TpMuguga_03g00013 | 4 | 0.099173554 |
| TpMuguga_03g00013 | 5 | 0 |
| TpMuguga_03g02250 | 1 | 0 |
| TpMuguga_03g00014 | 1 | 0 |

| | | |
|---|---|---|
| TpMuguga_03g00014 | 2 | 3.6 |
| TpMuguga_03g00014 | 3 | 0 |
| TpMuguga_03g00014 | 4 | 0 |
| TpMuguga_03g00014 | 5 | 0 |
| TpMuguga_03g00015 | 1 | 0.714285714 |
| TpMuguga_03g00015 | 2 | 1.142857143 |
| TpMuguga_03g00015 | 3 | 0.142857143 |
| TpMuguga_03g00015 | 4 | 0 |
| TpMuguga_03g02460 | 1 | 0.098360656 |
| TpMuguga_03g02460 | 2 | 0.540983607 |
| TpMuguga_03g02465 | 1 | 1.6125 |
| TpMuguga_03g02465 | 2 | 2.8875 |
| TpMuguga_03g02465 | 3 | 0.9375 |
| TpMuguga_03g02465 | 4 | 3.4125 |
| TpMuguga_03g02465 | 5 | 0 |
| TpMuguga_03g00018 | 1 | 0 |
| TpMuguga_03g00018 | 2 | 0 |
| TpMuguga_03g00020 | 1 | 0 |
| TpMuguga_03g00020 | 2 | 0 |
| TpMuguga_03g00020 | 3 | 0 |
| TpMuguga_03g00020 | 4 | 0 |
| TpMuguga_03g00020 | 5 | 0 |
| TpMuguga_03g00020 | 6 | 0 |
| TpMuguga_03g02475 | 1 | 0 |
| TpMuguga_03g02475 | 2 | 0 |
| TpMuguga_03g02475 | 3 | 0 |
| TpMuguga_03g02475 | 4 | 0 |
| TpMuguga_03g02475 | 5 | 0 |
| TpMuguga_03g02475 | 6 | 0 |
| TpMuguga_03g02475 | 7 | 0 |
| TpMuguga_03g02475 | 8 | 0 |
| TpMuguga_03g02475 | 9 | 0 |
| TpMuguga_03g02475 | 10 | 0 |
| TpMuguga_03g02475 | 11 | 0 |
| TpMuguga_03g02435 | 1 | 0.37037037 |
| TpMuguga_03g00024 | 1 | 0.026490066 |
| TpMuguga_03g00024 | 2 | 0 |
| TpMuguga_03g00024 | 3 | 0 |
| TpMuguga_03g00024 | 4 | 0 |
| TpMuguga_03g00024 | 5 | 0 |
| TpMuguga_03g00024 | 6 | 0 |
| TpMuguga_03g00025 | 1 | 0 |

| | | |
|---|---|---|
| TpMuguga_03g00025 | 2 | 0 |
| TpMuguga_03g00025 | 3 | 0 |
| TpMuguga_03g00026 | 1 | 0.919104991 |
| TpMuguga_03g00026 | 2 | 0.309810671 |
| TpMuguga_03g00027 | 1 | 1.420074349 |
| TpMuguga_03g00027 | 2 | 4.780669145 |
| TpMuguga_03g00027 | 3 | 0.936802974 |
| TpMuguga_03g02355 | 1 | 0 |
| TpMuguga_03g02355 | 2 | 0 |
| TpMuguga_03g02355 | 3 | 0 |
| TpMuguga_03g02355 | 4 | 0 |
| TpMuguga_03g02355 | 5 | 0 |
| TpMuguga_03g02355 | 6 | 0 |
| TpMuguga_03g02355 | 7 | 0 |
| TpMuguga_03g02355 | 8 | 0 |
| TpMuguga_03g02355 | 9 | 0 |
| TpMuguga_03g00030 | 1 | 0 |
| TpMuguga_03g00033 | 1 | 1.008 |
| TpMuguga_03g00033 | 2 | 0.792 |
| TpMuguga_03g00033 | 3 | 0.216 |
| TpMuguga_03g00033 | 4 | 0 |
| TpMuguga_03g00033 | 5 | 0 |
| TpMuguga_03g00033 | 6 | 0.936 |
| TpMuguga_03g00033 | 7 | 0 |
| TpMuguga_03g00033 | 8 | 0 |
| TpMuguga_03g02325 | 1 | 0 |
| TpMuguga_03g02325 | 2 | 0 |
| TpMuguga_03g02325 | 3 | 0 |
| TpMuguga_03g02325 | 4 | 0 |
| TpMuguga_03g02325 | 5 | 0 |
| TpMuguga_03g02325 | 6 | 0 |
| TpMuguga_03g02325 | 7 | 0 |
| TpMuguga_03g00036 | 1 | 0 |
| TpMuguga_03g00036 | 2 | 0 |
| TpMuguga_03g00036 | 3 | 0 |
| TpMuguga_03g00036 | 4 | 0 |
| TpMuguga_03g00037 | 1 | 0 |
| TpMuguga_03g00037 | 2 | 0 |
| TpMuguga_03g00037 | 3 | 0.358208955 |
| TpMuguga_03g00037 | 4 | 0.268656716 |
| TpMuguga_03g00037 | 5 | 1.567164179 |
| TpMuguga_03g00041 | 1 | 0 |

| | | |
|---|---|---|
| TpMuguga_03g00041 | 2 | 0.25 |
| TpMuguga_03g00041 | 3 | 0.6875 |
| TpMuguga_03g00041 | 4 | 1.0625 |
| TpMuguga_03g00041 | 5 | 0 |
| TpMuguga_03g00043 | 1 | 0 |
| TpMuguga_03g00043 | 2 | 0 |
| TpMuguga_03g00045 | 1 | 0.307692308 |
| TpMuguga_03g00046 | 1 | 5.043478261 |
| TpMuguga_03g00046 | 2 | 0 |
| TpMuguga_03g00046 | 3 | 0 |
| TpMuguga_03g00047 | 1 | 1.263157895 |
| TpMuguga_03g00047 | 2 | 0 |
| TpMuguga_03g00048 | 1 | 0.547368421 |
| TpMuguga_03g00048 | 2 | 4.378947368 |
| TpMuguga_03g00048 | 3 | 0.136842105 |
| TpMuguga_03g00048 | 4 | 0 |
| TpMuguga_03g00048 | 5 | 0 |
| TpMuguga_03g00048 | 6 | 0 |
| TpMuguga_03g00048 | 7 | 0 |
| TpMuguga_03g00048 | 8 | 0.273684211 |
| TpMuguga_03g00048 | 9 | 0 |
| TpMuguga_03g00048 | 10 | 0 |
| TpMuguga_03g00048 | 11 | 0 |
| TpMuguga_03g00048 | 12 | 0.684210526 |
| TpMuguga_03g00050 | 1 | 0 |
| TpMuguga_03g00050 | 2 | 0.6 |
| TpMuguga_03g00051 | 1 | 0 |
| TpMuguga_03g00052 | 1 | 0.153846154 |
| TpMuguga_03g00053 | 1 | 0 |
| TpMuguga_03g00053 | 2 | 0 |
| TpMuguga_03g00053 | 3 | 0 |
| TpMuguga_03g00053 | 4 | 0 |
| TpMuguga_03g00053 | 5 | 0 |
| TpMuguga_03g00053 | 6 | 0 |
| TpMuguga_03g00054 | 1 | 0 |
| TpMuguga_03g00055 | 1 | 0 |
| TpMuguga_03g00055 | 2 | 0 |
| TpMuguga_03g00055 | 3 | 0 |
| TpMuguga_03g00056 | 1 | 0 |
| TpMuguga_03g00056 | 2 | 0 |
| TpMuguga_03g00056 | 3 | 0 |
| TpMuguga_03g00056 | 4 | 0 |

| | | |
|---|---|---|
| TpMuguga_03g00056 | 5 | 0 |
| TpMuguga_03g00056 | 6 | 0 |
| TpMuguga_03g00056 | 7 | 0 |
| TpMuguga_03g00056 | 8 | 0 |
| TpMuguga_03g00058 | 1 | 0 |
| TpMuguga_03g00058 | 2 | 0 |
| TpMuguga_03g00058 | 3 | 0 |
| TpMuguga_03g00058 | 4 | 0 |
| TpMuguga_03g00058 | 5 | 5.294117647 |
| TpMuguga_03g02520 | 1 | 0 |
| TpMuguga_03g02520 | 2 | 0 |
| TpMuguga_03g02520 | 3 | 0 |
| TpMuguga_03g02520 | 4 | 0 |
| TpMuguga_03g02520 | 5 | 0 |
| TpMuguga_03g02520 | 6 | 0 |
| TpMuguga_03g02520 | 7 | 0 |
| TpMuguga_03g02215 | 1 | 0 |
| TpMuguga_03g02215 | 2 | 0 |
| TpMuguga_03g02285 | 1 | 0 |
| TpMuguga_03g02285 | 2 | 0.04 |
| TpMuguga_03g02285 | 3 | 0.08 |
| TpMuguga_03g02285 | 4 | 0 |
| TpMuguga_03g02285 | 5 | 0 |
| TpMuguga_03g02285 | 6 | 0 |
| TpMuguga_03g02285 | 7 | 0 |
| TpMuguga_03g00065 | 1 | 0 |
| TpMuguga_03g00065 | 2 | 0.545454545 |
| TpMuguga_03g00065 | 3 | 0 |
| TpMuguga_03g00065 | 4 | 0 |
| TpMuguga_03g00065 | 5 | 0.545454545 |
| TpMuguga_03g00065 | 6 | 1.090909091 |
| TpMuguga_03g00065 | 7 | 0.363636364 |
| TpMuguga_03g00065 | 8 | 0 |
| TpMuguga_03g00065 | 9 | 0 |
| TpMuguga_03g00065 | 10 | 0 |
| TpMuguga_03g00065 | 11 | 4.909090909 |
| TpMuguga_03g02675 | 1 | 0 |
| TpMuguga_03g02675 | 2 | 0 |
| TpMuguga_03g02675 | 3 | 0 |
| TpMuguga_03g02675 | 4 | 0 |
| TpMuguga_03g02675 | 5 | 0 |
| TpMuguga_03g02675 | 6 | 0 |

| | | |
|---|---|---|
| TpMuguga_03g00067 | 1 | 0 |
| TpMuguga_03g00067 | 2 | 0 |
| TpMuguga_03g00068 | 1 | 0 |
| TpMuguga_03g00068 | 2 | 0 |
| TpMuguga_03g00068 | 3 | 0 |
| TpMuguga_03g00068 | 4 | 0 |
| TpMuguga_03g00069 | 1 | 0 |
| TpMuguga_03g00069 | 2 | 0 |
| TpMuguga_03g00069 | 3 | 0 |
| TpMuguga_03g00069 | 4 | 0 |
| TpMuguga_03g00071 | 1 | 0 |
| TpMuguga_03g00071 | 2 | 0 |
| TpMuguga_03g00071 | 3 | 0 |
| TpMuguga_03g00071 | 4 | 0 |
| TpMuguga_03g00071 | 5 | 0 |
| TpMuguga_03g00071 | 6 | 0 |
| TpMuguga_03g00071 | 7 | 0 |
| TpMuguga_03g00071 | 8 | 0 |
| TpMuguga_03g00071 | 9 | 0.155844156 |
| TpMuguga_03g00071 | 10 | 0 |
| TpMuguga_03g00071 | 11 | 0 |
| TpMuguga_03g00074 | 1 | 0 |
| TpMuguga_03g00074 | 2 | 0 |
| TpMuguga_03g00074 | 3 | 0 |
| TpMuguga_03g00075 | 1 | 0 |
| TpMuguga_03g00075 | 2 | 0 |
| TpMuguga_03g00075 | 3 | 0 |
| TpMuguga_03g00075 | 4 | 0 |
| TpMuguga_03g00075 | 5 | 0 |
| TpMuguga_03g00076 | 1 | 0 |
| TpMuguga_03g00077 | 1 | 0 |
| TpMuguga_03g00077 | 2 | 0 |
| TpMuguga_03g00077 | 3 | 0 |
| TpMuguga_03g00078 | 1 | 2.5 |
| TpMuguga_03g00078 | 2 | 1.634615385 |
| TpMuguga_03g00078 | 3 | 0 |
| TpMuguga_03g00078 | 4 | 0.384615385 |
| TpMuguga_03g00078 | 5 | 0.769230769 |
| TpMuguga_03g00078 | 6 | 0 |
| TpMuguga_03g00078 | 7 | 0 |
| TpMuguga_03g00078 | 8 | 0.480769231 |
| TpMuguga_03g00078 | 9 | 1.057692308 |

| | | |
|---|---|---|
| TpMuguga_03g00079 | 1 | 0 |
| TpMuguga_03g00079 | 2 | 0 |
| TpMuguga_03g00079 | 3 | 0 |
| TpMuguga_03g00082 | 1 | 4 |
| TpMuguga_03g00082 | 2 | 0 |
| TpMuguga_03g00082 | 3 | 0 |
| TpMuguga_03g00082 | 4 | 0 |
| TpMuguga_03g00084 | 1 | 1.615384615 |
| TpMuguga_03g00084 | 2 | 0 |
| TpMuguga_03g00085 | 1 | 0 |
| TpMuguga_03g00087 | 1 | 0 |
| TpMuguga_03g00087 | 2 | 0 |
| TpMuguga_03g00087 | 3 | 2.5 |
| TpMuguga_03g00087 | 4 | 10.5 |
| TpMuguga_03g00088 | 1 | 6 |
| TpMuguga_03g00088 | 2 | 0 |
| TpMuguga_03g00088 | 3 | 0 |
| TpMuguga_03g00088 | 4 | 0 |
| TpMuguga_03g00088 | 5 | 0 |
| TpMuguga_03g00090 | 1 | 0.11827957 |
| TpMuguga_03g00092 | 1 | 0 |
| TpMuguga_03g00092 | 2 | 0 |
| TpMuguga_03g00092 | 3 | 0 |
| TpMuguga_03g00092 | 4 | 0 |
| TpMuguga_03g00092 | 5 | 14 |
| TpMuguga_03g00092 | 6 | 0 |
| TpMuguga_03g00092 | 7 | 0 |
| TpMuguga_03g00092 | 8 | 0 |
| TpMuguga_03g00092 | 9 | 0 |
| TpMuguga_03g00092 | 10 | 0 |
| TpMuguga_03g00092 | 11 | 0 |
| TpMuguga_03g00092 | 12 | 0 |
| TpMuguga_03g00092 | 13 | 0 |
| TpMuguga_03g00093 | 1 | 0 |
| TpMuguga_03g00093 | 2 | 0 |
| TpMuguga_03g02175 | 1 | 0 |
| TpMuguga_03g02175 | 2 | 3.692307692 |
| TpMuguga_03g02175 | 3 | 0 |
| TpMuguga_03g02175 | 4 | 0 |
| TpMuguga_03g02175 | 5 | 0 |
| TpMuguga_03g02175 | 6 | 0 |
| TpMuguga_03g02175 | 7 | 0 |

| | | |
|---|---|---|
| TpMuguga_03g02175 | 8 | 0 |
| TpMuguga_03g02175 | 9 | 0 |
| TpMuguga_03g02175 | 10 | 10.15384615 |
| TpMuguga_03g02175 | 11 | 5.538461538 |
| TpMuguga_03g02315 | 1 | 0 |
| TpMuguga_03g02315 | 2 | 1.1 |
| TpMuguga_03g02315 | 3 | 0.8 |
| TpMuguga_03g00098 | 1 | 0 |
| TpMuguga_03g00100 | 1 | 0 |
| TpMuguga_03g00100 | 2 | 0 |
| TpMuguga_03g00100 | 3 | 0 |
| TpMuguga_03g00101 | 1 | 2.086956522 |
| TpMuguga_03g00101 | 2 | 2.739130435 |
| TpMuguga_03g00101 | 3 | 0 |
| TpMuguga_03g00101 | 4 | 0.260869565 |
| TpMuguga_03g00101 | 5 | 0 |
| TpMuguga_03g00102 | 1 | 0 |
| TpMuguga_03g00102 | 2 | 0.5 |
| TpMuguga_03g00105 | 1 | 0 |
| TpMuguga_03g00106 | 1 | 0 |
| TpMuguga_03g00106 | 2 | 0.725274725 |
| TpMuguga_03g00107 | 1 | 0 |
| TpMuguga_03g00107 | 2 | 0 |
| TpMuguga_03g00108 | 1 | 0 |
| TpMuguga_03g00111 | 1 | 0 |
| TpMuguga_03g00111 | 2 | 0 |
| TpMuguga_03g00111 | 3 | 0 |
| TpMuguga_03g00110 | 1 | 0 |
| TpMuguga_03g00110 | 2 | 0 |
| TpMuguga_03g00110 | 3 | 0.208877285 |
| TpMuguga_03g00110 | 4 | 0 |
| TpMuguga_03g00110 | 5 | 0.287206266 |
| TpMuguga_03g00110 | 6 | 0.078328982 |
| TpMuguga_03g00110 | 7 | 0.104438642 |
| TpMuguga_03g00110 | 8 | 0.130548303 |
| TpMuguga_03g00110 | 9 | 0.052219321 |
| TpMuguga_03g00109 | 1 | 0.07 |
| TpMuguga_03g00112 | 1 | 2 |
| TpMuguga_03g00112 | 2 | 1.25 |
| TpMuguga_03g00112 | 3 | 1.5 |
| TpMuguga_03g00112 | 4 | 0.75 |
| TpMuguga_03g00112 | 5 | 0 |

| | | |
|---|---|---|
| TpMuguga_03g00114 | 1 | 0 |
| TpMuguga_03g02495 | 1 | 1.003782368 |
| TpMuguga_03g02495 | 2 | 0.007273785 |
| TpMuguga_03g02495 | 3 | 0.027640384 |
| TpMuguga_03g02495 | 4 | 0.001454757 |
| TpMuguga_03g00115 | 1 | 0.026666667 |
| TpMuguga_03g00115 | 2 | 2.648888889 |
| TpMuguga_03g00118 | 1 | 0 |
| TpMuguga_03g02565 | 1 | 0 |
| TpMuguga_03g02565 | 2 | 0 |
| TpMuguga_03g02565 | 3 | 0 |
| TpMuguga_03g00123 | 1 | 1.12244898 |
| TpMuguga_03g00123 | 2 | 2.448979592 |
| TpMuguga_03g00123 | 3 | 1.836734694 |
| TpMuguga_03g00123 | 4 | 0 |
| TpMuguga_03g00124 | 1 | 0 |
| TpMuguga_03g00124 | 2 | 0 |
| TpMuguga_03g00124 | 3 | 0 |
| TpMuguga_03g00124 | 4 | 0 |
| TpMuguga_03g00124 | 5 | 0 |
| TpMuguga_03g00124 | 6 | 0 |
| TpMuguga_03g00124 | 7 | 0 |
| TpMuguga_03g00124 | 8 | 0 |
| TpMuguga_03g00124 | 9 | 0 |
| TpMuguga_03g00125 | 1 | 0 |
| TpMuguga_03g00128 | 1 | 0.45 |
| TpMuguga_03g00128 | 2 | 0 |
| TpMuguga_03g00128 | 3 | 0 |
| TpMuguga_03g00128 | 4 | 0 |
| TpMuguga_03g00128 | 5 | 1.575 |
| TpMuguga_03g00128 | 6 | 2.8125 |
| TpMuguga_03g00128 | 7 | 0.5625 |
| TpMuguga_03g00128 | 8 | 0 |
| TpMuguga_03g00129 | 1 | 0.242424242 |
| TpMuguga_03g00129 | 2 | 3.878787879 |
| TpMuguga_03g00129 | 3 | 0 |
| TpMuguga_03g00130 | 1 | 0.923076923 |
| TpMuguga_03g00130 | 2 | 0 |
| TpMuguga_03g00130 | 3 | 0 |
| TpMuguga_03g00131 | 1 | 1.222222222 |
| TpMuguga_03g00131 | 2 | 2.444444444 |
| TpMuguga_03g00131 | 3 | 0.111111111 |

| | | |
|---|---|---|
| TpMuguga_03g00131 | 4 | 0 |
| TpMuguga_03g00132 | 1 | 4 |
| TpMuguga_03g00132 | 2 | 0 |
| TpMuguga_03g00132 | 3 | 0.679245283 |
| TpMuguga_03g00137 | 1 | 0 |
| TpMuguga_03g00137 | 2 | 0 |
| TpMuguga_03g02415 | 1 | 0 |
| TpMuguga_03g02415 | 2 | 0 |
| TpMuguga_03g02415 | 3 | 0 |
| TpMuguga_03g02415 | 4 | 0 |
| TpMuguga_03g02415 | 5 | 0 |
| TpMuguga_03g02415 | 6 | 0 |
| TpMuguga_03g02415 | 7 | 77.33333333 |
| TpMuguga_03g00141 | 1 | 0 |
| TpMuguga_03g00141 | 2 | 0 |
| TpMuguga_03g00141 | 3 | 5.185185185 |
| TpMuguga_03g00141 | 4 | 0 |
| TpMuguga_03g00141 | 5 | 0 |
| TpMuguga_03g00141 | 6 | 0 |
| TpMuguga_03g00141 | 7 | 0 |
| TpMuguga_03g00141 | 8 | 0 |
| TpMuguga_03g00141 | 9 | 0 |
| TpMuguga_03g00142 | 1 | 2.8 |
| TpMuguga_03g00142 | 2 | 0 |
| TpMuguga_03g00142 | 3 | 0 |
| TpMuguga_03g00142 | 4 | 0 |
| TpMuguga_03g00142 | 5 | 0 |
| TpMuguga_03g00142 | 6 | 0 |
| TpMuguga_03g00143 | 1 | 0.698412698 |
| TpMuguga_03g00144 | 1 | 0 |
| TpMuguga_03g00145 | 1 | 0 |
| TpMuguga_03g00145 | 2 | 0 |
| TpMuguga_03g00145 | 3 | 0 |
| TpMuguga_03g00145 | 4 | 0 |
| TpMuguga_03g00145 | 5 | 0 |
| TpMuguga_03g00145 | 6 | 2.571428571 |
| TpMuguga_03g00146 | 1 | 0 |
| TpMuguga_03g00146 | 2 | 0 |
| TpMuguga_03g00146 | 3 | 0 |
| TpMuguga_03g00146 | 4 | 0 |
| TpMuguga_03g00146 | 5 | 0 |
| TpMuguga_03g00150 | 1 | 0 |

| | | |
|---|---|---|
| TpMuguga_03g00150 | 2 | 9.866666667 |
| TpMuguga_03g00150 | 3 | 4.266666667 |
| TpMuguga_03g00155 | 1 | 7.060344828 |
| TpMuguga_03g00155 | 2 | 0 |
| TpMuguga_03g00155 | 3 | 0 |
| TpMuguga_03g00155 | 4 | 0 |
| TpMuguga_03g00155 | 5 | 0 |
| TpMuguga_03g00155 | 6 | 0 |
| TpMuguga_03g00154 | 1 | 0 |
| TpMuguga_03g00154 | 2 | 0 |
| TpMuguga_03g02685 | 1 | 0.019966722 |
| TpMuguga_03g02685 | 2 | 0 |
| TpMuguga_03g02685 | 3 | 0 |
| TpMuguga_03g02685 | 4 | 0 |
| TpMuguga_03g02685 | 5 | 0.059900166 |
| TpMuguga_03g02685 | 6 | 0 |
| TpMuguga_03g02685 | 7 | 0 |
| TpMuguga_03g02685 | 8 | 0 |
| TpMuguga_03g02685 | 9 | 0 |
| TpMuguga_03g02685 | 10 | 0 |
| TpMuguga_03g02685 | 11 | 0 |
| TpMuguga_03g00158 | 1 | 0 |
| TpMuguga_03g00158 | 2 | 0.010989011 |
| TpMuguga_03g00158 | 3 | 0.054945055 |
| TpMuguga_03g00158 | 4 | 0 |
| TpMuguga_03g00158 | 5 | 0.054945055 |
| TpMuguga_03g00160 | 1 | 0 |
| TpMuguga_03g00161 | 1 | 0 |
| TpMuguga_03g00161 | 2 | 0.238938053 |
| TpMuguga_03g00162 | 1 | 0 |
| TpMuguga_03g00162 | 2 | 0 |
| TpMuguga_03g00162 | 3 | 0 |
| TpMuguga_03g00162 | 4 | 0.007798284 |
| TpMuguga_03g00163 | 1 | 0.440677966 |
| TpMuguga_03g00164 | 1 | 0 |
| TpMuguga_03g02275 | 1 | 0 |
| TpMuguga_03g00165 | 1 | 0 |
| TpMuguga_03g00165 | 2 | 0 |
| TpMuguga_03g00165 | 3 | 0 |
| TpMuguga_03g00165 | 4 | 0 |
| TpMuguga_03g00165 | 5 | 0 |
| TpMuguga_03g00165 | 6 | 0 |

| | | |
|---|---|---|
| TpMuguga_03g00165 | 7 | 0.87804878 |
| TpMuguga_03g00165 | 8 | 0 |
| TpMuguga_03g00166 | 1 | 0.019512195 |
| TpMuguga_03g00166 | 2 | 5.248780488 |
| TpMuguga_03g00166 | 3 | 7.12195122 |
| TpMuguga_03g00167 | 1 | 1.227722772 |
| TpMuguga_03g00167 | 2 | 0.475247525 |
| TpMuguga_03g00167 | 3 | 2.059405941 |
| TpMuguga_03g00168 | 1 | 0 |
| TpMuguga_03g00168 | 2 | 0 |
| TpMuguga_03g00170 | 1 | 0 |
| TpMuguga_03g00170 | 2 | 0 |
| TpMuguga_03g00170 | 3 | 0 |
| TpMuguga_03g00171 | 1 | 0 |
| TpMuguga_03g00171 | 2 | 0 |
| TpMuguga_03g00171 | 3 | 0 |
| TpMuguga_03g00171 | 4 | 0.5 |
| TpMuguga_03g00171 | 5 | 0 |
| TpMuguga_03g00171 | 6 | 0 |
| TpMuguga_03g00171 | 7 | 0 |
| TpMuguga_03g00171 | 8 | 0 |
| TpMuguga_03g00171 | 9 | 8.5 |
| TpMuguga_03g00171 | 10 | 0 |
| TpMuguga_03g00171 | 11 | 0 |
| TpMuguga_03g00172 | 1 | 0 |
| TpMuguga_03g00172 | 2 | 0 |
| TpMuguga_03g00172 | 3 | 0 |
| TpMuguga_03g00172 | 4 | 0 |
| TpMuguga_03g00172 | 5 | 0 |
| TpMuguga_03g00172 | 6 | 0 |
| TpMuguga_03g00172 | 7 | 2.448979592 |
| TpMuguga_03g00172 | 8 | 2.448979592 |
| TpMuguga_03g00172 | 9 | 0.244897959 |
| TpMuguga_03g00172 | 10 | 0 |
| TpMuguga_03g00172 | 11 | 0 |
| TpMuguga_03g02570 | 1 | 0.610169492 |
| TpMuguga_03g02570 | 2 | 0 |
| TpMuguga_03g02570 | 3 | 0 |
| TpMuguga_03g02570 | 4 | 0 |
| TpMuguga_03g02570 | 5 | 0.610169492 |
| TpMuguga_03g02570 | 6 | 0.762711864 |
| TpMuguga_03g02570 | 7 | 2.440677966 |

| | | |
|---|---|---|
| TpMuguga_03g02570 | 8 | 16.62711864 |
| TpMuguga_03g00175 | 1 | 1.150684932 |
| TpMuguga_03g00176 | 1 | 0 |
| TpMuguga_03g00179 | 1 | 3.123595506 |
| TpMuguga_03g00179 | 2 | 2.719101124 |
| TpMuguga_03g00179 | 3 | 2.741573034 |
| TpMuguga_03g00179 | 4 | 0.08988764 |
| TpMuguga_03g00179 | 5 | 0.08988764 |
| TpMuguga_03g02605 | 1 | 0 |
| TpMuguga_03g02605 | 2 | 0 |
| TpMuguga_03g02605 | 3 | 0 |
| TpMuguga_03g02605 | 4 | 0 |
| TpMuguga_03g02595 | 1 | 0 |
| TpMuguga_03g02595 | 2 | 0 |
| TpMuguga_03g00182 | 1 | 0 |
| TpMuguga_03g00182 | 2 | 0 |
| TpMuguga_03g00183 | 1 | 0.482142857 |
| TpMuguga_03g00183 | 2 | 0.482142857 |
| TpMuguga_03g00183 | 3 | 1.285714286 |
| TpMuguga_03g00183 | 4 | 0 |
| TpMuguga_03g00183 | 5 | 0 |
| TpMuguga_03g00183 | 6 | 0 |
| TpMuguga_03g00183 | 7 | 0 |
| TpMuguga_03g00183 | 8 | 0 |
| TpMuguga_03g00184 | 1 | 0 |
| TpMuguga_03g00184 | 2 | 0 |
| TpMuguga_03g00184 | 3 | 0.294117647 |
| TpMuguga_03g00184 | 4 | 0 |
| TpMuguga_03g00186 | 1 | 0 |
| TpMuguga_03g00186 | 2 | 0 |
| TpMuguga_03g00186 | 3 | 0 |
| TpMuguga_03g00186 | 4 | 32 |
| TpMuguga_03g00186 | 5 | 0 |
| TpMuguga_03g00186 | 6 | 0 |
| TpMuguga_03g00186 | 7 | 32 |
| TpMuguga_03g00187 | 1 | 0.228426396 |
| TpMuguga_03g00187 | 2 | 2.258883249 |
| TpMuguga_03g00187 | 3 | 1.598984772 |
| TpMuguga_03g00187 | 4 | 1.675126904 |
| TpMuguga_03g02380 | 1 | 0 |
| TpMuguga_03g02380 | 2 | 0 |
| TpMuguga_03g02380 | 3 | 0 |

| | | |
|---|---|---|
| TpMuguga_03g02380 | 4 | 0 |
| TpMuguga_03g02380 | 5 | 12 |
| TpMuguga_03g02380 | 6 | 2 |
| TpMuguga_03g02380 | 7 | 6 |
| TpMuguga_03g00192 | 1 | 0 |
| TpMuguga_03g00192 | 2 | 0 |
| TpMuguga_03g00192 | 3 | 0 |
| TpMuguga_03g00192 | 4 | 0 |
| TpMuguga_03g00192 | 5 | 0 |
| TpMuguga_03g00195 | 1 | 0 |
| TpMuguga_03g00195 | 2 | 0 |
| TpMuguga_03g00196 | 1 | 0.275862069 |
| TpMuguga_03g00197 | 1 | 0.157894737 |
| TpMuguga_03g02545 | 1 | 0.002737851 |
| TpMuguga_03g02545 | 2 | 0 |
| TpMuguga_03g02545 | 3 | 0.004106776 |
| TpMuguga_03g02545 | 4 | 0.010951403 |
| TpMuguga_03g02545 | 5 | 0.013689254 |
| TpMuguga_03g00200 | 1 | 0 |
| TpMuguga_03g00200 | 2 | 1.285714286 |
| TpMuguga_03g00201 | 1 | 0 |
| TpMuguga_03g00201 | 2 | 0 |
| TpMuguga_03g00202 | 1 | 0 |
| TpMuguga_03g00202 | 2 | 2.727272727 |
| TpMuguga_03g00202 | 3 | 0 |
| TpMuguga_03g00202 | 4 | 0 |
| TpMuguga_03g00202 | 5 | 0 |
| TpMuguga_03g00202 | 6 | 0 |
| TpMuguga_03g00202 | 7 | 0 |
| TpMuguga_03g00202 | 8 | 0 |
| TpMuguga_03g00202 | 9 | 2.727272727 |
| TpMuguga_03g00202 | 10 | 4.545454545 |
| TpMuguga_03g00202 | 11 | 0 |
| TpMuguga_03g00202 | 12 | 0 |
| TpMuguga_03g00202 | 13 | 0 |
| TpMuguga_03g00202 | 14 | 0 |
| TpMuguga_03g00202 | 15 | 0 |
| TpMuguga_03g00202 | 16 | 0 |
| TpMuguga_03g00202 | 17 | 0 |
| TpMuguga_03g00202 | 18 | 0 |
| TpMuguga_03g00202 | 19 | 1.818181818 |
| TpMuguga_03g00203 | 1 | 0 |

| | | |
|---|---|---|
| TpMuguga_03g00203 | 2 | 0 |
| TpMuguga_03g00203 | 3 | 0 |
| TpMuguga_03g00203 | 4 | 0 |
| TpMuguga_03g00204 | 1 | 0 |
| TpMuguga_03g00204 | 2 | 0.259259259 |
| TpMuguga_03g00204 | 3 | 1.296296296 |
| TpMuguga_03g00204 | 4 | 5.962962963 |
| TpMuguga_03g00204 | 5 | 0 |
| TpMuguga_03g00204 | 6 | 0 |
| TpMuguga_03g00205 | 1 | 0 |
| TpMuguga_03g00205 | 2 | 0 |
| TpMuguga_03g00205 | 3 | 0 |
| TpMuguga_03g00205 | 4 | 0 |
| TpMuguga_03g00205 | 5 | 0 |
| TpMuguga_03g00205 | 6 | 0 |
| TpMuguga_03g00205 | 7 | 0 |
| TpMuguga_03g00206 | 1 | 0 |
| TpMuguga_03g00206 | 2 | 0 |
| TpMuguga_03g00206 | 3 | 0 |
| TpMuguga_03g00206 | 4 | 0 |
| TpMuguga_03g00206 | 5 | 0 |
| TpMuguga_03g00208 | 1 | 1.272727273 |
| TpMuguga_03g00209 | 1 | 0 |
| TpMuguga_03g00209 | 2 | 0 |
| TpMuguga_03g00210 | 1 | 0 |
| TpMuguga_03g00210 | 2 | 0 |
| TpMuguga_03g00210 | 3 | 0 |
| TpMuguga_03g00210 | 4 | 0 |
| TpMuguga_03g00210 | 5 | 1 |
| TpMuguga_03g00210 | 6 | 0 |
| TpMuguga_03g00210 | 7 | 9 |
| TpMuguga_03g00211 | 1 | 1 |
| TpMuguga_03g00212 | 1 | 0 |
| TpMuguga_03g00212 | 2 | 0 |
| TpMuguga_03g00212 | 3 | 0 |
| TpMuguga_03g00212 | 4 | 0 |
| TpMuguga_03g00212 | 5 | 0 |
| TpMuguga_03g00212 | 6 | 0 |
| TpMuguga_03g00214 | 1 | 0 |
| TpMuguga_03g00214 | 2 | 4.875 |
| TpMuguga_03g00215 | 1 | 0 |
| TpMuguga_03g00215 | 2 | 0 |

| | | |
|---|---|---|
| TpMuguga_03g00215 | 3 | 0 |
| TpMuguga_03g00215 | 4 | 0 |
| TpMuguga_03g00216 | 1 | 0 |
| TpMuguga_03g00216 | 2 | 0 |
| TpMuguga_03g00216 | 3 | 0 |
| TpMuguga_03g00216 | 4 | 0 |
| TpMuguga_03g00216 | 5 | 0 |
| TpMuguga_03g00216 | 6 | 0 |
| TpMuguga_03g00216 | 7 | 0 |
| TpMuguga_03g00216 | 8 | 0 |
| TpMuguga_03g00216 | 9 | 0 |
| TpMuguga_03g00220 | 1 | 0 |
| TpMuguga_03g00220 | 2 | 0 |
| TpMuguga_03g00220 | 3 | 0 |
| TpMuguga_03g00220 | 4 | 0 |
| TpMuguga_03g00220 | 5 | 0 |
| TpMuguga_03g00220 | 6 | 0 |
| TpMuguga_03g00220 | 7 | 0 |
| TpMuguga_03g02560 | 1 | 0 |
| TpMuguga_03g02560 | 2 | 0 |
| TpMuguga_03g00223 | 1 | 0 |
| TpMuguga_03g00223 | 2 | 0 |
| TpMuguga_03g00223 | 3 | 0 |
| TpMuguga_03g00224 | 1 | 0 |
| TpMuguga_03g00224 | 2 | 0 |
| TpMuguga_03g00224 | 3 | 0 |
| TpMuguga_03g00224 | 4 | 0 |
| TpMuguga_03g00224 | 5 | 0 |
| TpMuguga_03g00225 | 1 | 0 |
| TpMuguga_03g00225 | 2 | 0.002881844 |
| TpMuguga_03g00225 | 3 | 0.005763689 |
| TpMuguga_03g00225 | 4 | 0.34870317 |
| TpMuguga_03g00225 | 5 | 10.47262248 |
| TpMuguga_03g00226 | 1 | 0.756880734 |
| TpMuguga_03g00226 | 2 | 0.114678899 |
| TpMuguga_03g00226 | 3 | 0 |
| TpMuguga_03g00226 | 4 | 0.194954128 |
| TpMuguga_03g00227 | 1 | 0.875 |
| TpMuguga_03g00227 | 2 | 1.8125 |
| TpMuguga_03g00227 | 3 | 0.5 |
| TpMuguga_03g00227 | 4 | 1.0625 |
| TpMuguga_03g00228 | 1 | 2 |

| | | |
|---|---|---|
| TpMuguga_03g00228 | 2 | 4 |
| TpMuguga_03g00230 | 1 | 0 |
| TpMuguga_03g00230 | 2 | 0 |
| TpMuguga_03g00230 | 3 | 0 |
| TpMuguga_03g00230 | 4 | 0 |
| TpMuguga_03g00231 | 1 | 0 |
| TpMuguga_03g00233 | 1 | 0 |
| TpMuguga_03g00233 | 2 | 1.428571429 |
| TpMuguga_03g00233 | 3 | 5.714285714 |
| TpMuguga_03g00233 | 4 | 0 |
| TpMuguga_03g00233 | 5 | 0.714285714 |
| TpMuguga_03g00233 | 6 | 0 |
| TpMuguga_03g00233 | 7 | 0 |
| TpMuguga_03g00233 | 8 | 0 |
| TpMuguga_03g00233 | 9 | 0 |
| TpMuguga_03g00235 | 1 | 0 |
| TpMuguga_03g00235 | 2 | 0 |
| TpMuguga_03g00235 | 3 | 0 |
| TpMuguga_03g00235 | 4 | 0 |
| TpMuguga_03g00236 | 1 | 0 |
| TpMuguga_03g00236 | 2 | 0 |
| TpMuguga_03g00236 | 3 | 0 |
| TpMuguga_03g00236 | 4 | 0 |
| TpMuguga_03g00236 | 5 | 2.5 |
| TpMuguga_03g00236 | 6 | 4 |
| TpMuguga_03g00236 | 7 | 6.5 |
| TpMuguga_03g00236 | 8 | 3 |
| TpMuguga_03g00236 | 9 | 0 |
| TpMuguga_03g00236 | 10 | 0 |
| TpMuguga_03g00236 | 11 | 0 |
| TpMuguga_03g00236 | 12 | 0 |
| TpMuguga_03g00238 | 1 | 0 |
| TpMuguga_03g00238 | 2 | 0 |
| TpMuguga_03g00238 | 3 | 0 |
| TpMuguga_03g00238 | 4 | 0 |
| TpMuguga_03g00238 | 5 | 0 |
| TpMuguga_03g02240 | 1 | 0 |
| TpMuguga_03g00239 | 1 | 8 |
| TpMuguga_03g00239 | 2 | 0 |
| TpMuguga_03g00239 | 3 | 0 |
| TpMuguga_03g02665 | 1 | 0 |
| TpMuguga_03g02410 | 1 | 0 |

| | | |
|---|---|---|
| TpMuguga_03g02410 | 2 | 0 |
| TpMuguga_03g02410 | 3 | 0 |
| TpMuguga_03g02410 | 4 | 0 |
| TpMuguga_03g00243 | 1 | 0 |
| TpMuguga_03g00244 | 1 | 0 |
| TpMuguga_03g00244 | 2 | 0 |
| TpMuguga_03g00244 | 3 | 3 |
| TpMuguga_03g00248 | 1 | 0 |
| TpMuguga_03g00248 | 2 | 0 |
| TpMuguga_03g00250 | 1 | 0 |
| TpMuguga_03g00250 | 2 | 0 |
| TpMuguga_03g00250 | 3 | 0 |
| TpMuguga_03g00250 | 4 | 0 |
| TpMuguga_03g00250 | 5 | 0 |
| TpMuguga_03g00250 | 6 | 0 |
| TpMuguga_03g00250 | 7 | 0 |
| TpMuguga_03g00249 | 1 | 0 |
| TpMuguga_03g00249 | 2 | 0 |
| TpMuguga_03g00252 | 1 | 0 |
| TpMuguga_03g00254 | 1 | 0 |
| TpMuguga_03g00257 | 1 | 1.25 |
| TpMuguga_03g02370 | 1 | 0 |
| TpMuguga_03g02370 | 2 | 0 |
| TpMuguga_03g02370 | 3 | 0 |
| TpMuguga_03g02370 | 4 | 0 |
| TpMuguga_03g02370 | 5 | 0 |
| TpMuguga_03g02370 | 6 | 0 |
| TpMuguga_03g02370 | 7 | 0 |
| TpMuguga_03g02370 | 8 | 0 |
| TpMuguga_03g00258 | 1 | 0 |
| TpMuguga_03g00258 | 2 | 0 |
| TpMuguga_03g00258 | 3 | 0 |
| TpMuguga_03g02375 | 1 | 0 |
| TpMuguga_03g02375 | 2 | 0 |
| TpMuguga_03g00261 | 1 | 0.923076923 |
| TpMuguga_03g00261 | 2 | 0.461538462 |
| TpMuguga_03g00261 | 3 | 0.923076923 |
| TpMuguga_03g00261 | 4 | 0.923076923 |
| TpMuguga_03g00261 | 5 | 0.923076923 |
| TpMuguga_03g00262 | 1 | 2.194915254 |
| TpMuguga_03g00262 | 2 | 1.661016949 |
| TpMuguga_03g00262 | 3 | 0.059322034 |

| | | |
|---|---|---|
| TpMuguga_03g00262 | 4 | 0.711864407 |
| TpMuguga_03g00262 | 5 | 1.720338983 |
| TpMuguga_03g00262 | 6 | 0.949152542 |
| TpMuguga_03g00264 | 1 | 0 |
| TpMuguga_03g00264 | 2 | 0 |
| TpMuguga_03g00264 | 3 | 0 |
| TpMuguga_03g00265 | 1 | 0 |
| TpMuguga_03g00265 | 2 | 0 |
| TpMuguga_03g00265 | 3 | 0 |
| TpMuguga_03g00265 | 4 | 0 |
| TpMuguga_03g00265 | 5 | 0 |
| TpMuguga_03g00267 | 1 | 0 |
| TpMuguga_03g00267 | 2 | 0 |
| TpMuguga_03g00269 | 1 | 0 |
| TpMuguga_03g00269 | 2 | 0 |
| TpMuguga_03g00269 | 3 | 0 |
| TpMuguga_03g00269 | 4 | 0 |
| TpMuguga_03g00269 | 5 | 0 |
| TpMuguga_03g00269 | 6 | 0 |
| TpMuguga_03g00270 | 1 | 0 |
| TpMuguga_03g00270 | 2 | 0 |
| TpMuguga_03g00270 | 3 | 0 |
| TpMuguga_03g00270 | 4 | 0 |
| TpMuguga_03g02310 | 1 | 0 |
| TpMuguga_03g02310 | 2 | 0 |
| TpMuguga_03g02310 | 3 | 0 |
| TpMuguga_03g00271 | 1 | 0.411764706 |
| TpMuguga_03g00272 | 1 | 0 |
| TpMuguga_03g02400 | 1 | 0 |
| TpMuguga_03g02400 | 2 | 0 |
| TpMuguga_03g02400 | 3 | 0 |
| TpMuguga_03g02400 | 4 | 0 |
| TpMuguga_03g02400 | 5 | 0 |
| TpMuguga_03g02400 | 6 | 0 |
| TpMuguga_03g02400 | 7 | 0 |
| TpMuguga_03g00275 | 1 | 4.375 |
| TpMuguga_03g00275 | 2 | 0 |
| TpMuguga_03g00275 | 3 | 0 |
| TpMuguga_03g00275 | 4 | 0 |
| TpMuguga_03g00275 | 5 | 0 |
| TpMuguga_03g00275 | 6 | 0.875 |
| TpMuguga_03g00276 | 1 | 0 |

| | | |
|---|---|---|
| TpMuguga_03g00276 | 2 | 0.129310345 |
| TpMuguga_03g00276 | 3 | 0.043103448 |
| TpMuguga_03g00276 | 4 | 3.75 |
| TpMuguga_03g02425 | 1 | 0 |
| TpMuguga_03g02425 | 2 | 6.666666667 |
| TpMuguga_03g02425 | 3 | 2.666666667 |
| TpMuguga_03g00277 | 1 | 5.333333333 |
| TpMuguga_03g00277 | 2 | 0 |
| TpMuguga_03g00277 | 3 | 0 |
| TpMuguga_03g00278 | 1 | 9 |
| TpMuguga_03g00278 | 2 | 0 |
| TpMuguga_03g00279 | 1 | 0 |
| TpMuguga_03g00279 | 2 | 0 |
| TpMuguga_03g00279 | 3 | 0 |
| TpMuguga_03g00280 | 1 | 0.488721805 |
| TpMuguga_03g00282 | 1 | 0 |
| TpMuguga_03g00284 | 1 | 0 |
| TpMuguga_03g00285 | 1 | 0 |
| TpMuguga_03g00286 | 1 | 0 |
| TpMuguga_03g00286 | 2 | 0 |
| TpMuguga_03g00286 | 3 | 0 |
| TpMuguga_03g00287 | 1 | 2.571428571 |
| TpMuguga_03g00287 | 2 | 0 |
| TpMuguga_03g00289 | 1 | 0.038910506 |
| TpMuguga_03g00290 | 1 | 0.251046025 |
| TpMuguga_03g00290 | 2 | 0.10460251 |
| TpMuguga_03g00290 | 3 | 0 |
| TpMuguga_03g00290 | 4 | 0 |
| TpMuguga_03g00291 | 1 | 0 |
| TpMuguga_03g00291 | 2 | 0 |
| TpMuguga_03g00291 | 3 | 0 |
| TpMuguga_03g00291 | 4 | 1.448275862 |
| TpMuguga_03g00291 | 5 | 3.931034483 |
| TpMuguga_03g00292 | 1 | 0 |
| TpMuguga_03g00292 | 2 | 0 |
| TpMuguga_03g00292 | 3 | 0 |
| TpMuguga_03g00292 | 4 | 0 |
| TpMuguga_03g00292 | 5 | 0 |
| TpMuguga_03g00293 | 1 | 0 |
| TpMuguga_03g00294 | 1 | 0.132450331 |
| TpMuguga_03g00294 | 2 | 0.913907285 |
| TpMuguga_03g00294 | 3 | 1.880794702 |

| | | |
|---|---|---|
| TpMuguga_03g00295 | 1 | 0 |
| TpMuguga_03g00299 | 1 | 0 |
| TpMuguga_03g00299 | 2 | 0 |
| TpMuguga_03g00299 | 3 | 0.952380952 |
| TpMuguga_03g00299 | 4 | 0.952380952 |
| TpMuguga_03g00301 | 1 | 5 |
| TpMuguga_03g02350 | 1 | 0.13368984 |
| TpMuguga_03g02350 | 2 | 0.053475936 |
| TpMuguga_03g02350 | 3 | 0 |
| TpMuguga_03g02350 | 4 | 3.493761141 |
| TpMuguga_03g00304 | 1 | 0.235294118 |
| TpMuguga_03g00304 | 2 | 1.882352941 |
| TpMuguga_03g00304 | 3 | 0 |
| TpMuguga_03g00305 | 1 | 0 |
| TpMuguga_03g00305 | 2 | 0 |
| TpMuguga_03g00305 | 3 | 0 |
| TpMuguga_03g02555 | 1 | 0 |
| TpMuguga_03g02555 | 2 | 0 |
| TpMuguga_03g02555 | 3 | 0.128571429 |
| TpMuguga_03g02555 | 4 | 0 |
| TpMuguga_03g02555 | 5 | 0 |
| TpMuguga_03g02555 | 6 | 0 |
| TpMuguga_03g02555 | 7 | 0 |
| TpMuguga_03g02555 | 8 | 0 |
| TpMuguga_03g00308 | 1 | 0 |
| TpMuguga_03g00308 | 2 | 0 |
| TpMuguga_03g00309 | 1 | 19.72727273 |
| TpMuguga_03g00309 | 2 | 0 |
| TpMuguga_03g00309 | 3 | 1.272727273 |
| TpMuguga_03g00309 | 4 | 0 |
| TpMuguga_03g00309 | 5 | 0 |
| TpMuguga_03g00309 | 6 | 1.909090909 |
| TpMuguga_03g00310 | 1 | 1.00862069 |
| TpMuguga_03g00311 | 1 | 0 |
| TpMuguga_03g00311 | 2 | 0 |
| TpMuguga_03g00311 | 3 | 0 |
| TpMuguga_03g00311 | 4 | 0 |
| TpMuguga_03g00311 | 5 | 0 |
| TpMuguga_03g00311 | 6 | 0 |
| TpMuguga_03g00311 | 7 | 0 |
| TpMuguga_03g00311 | 8 | 0 |
| TpMuguga_03g00311 | 9 | 0 |

| | | |
|---|---|---|
| TpMuguga_03g00311 | 10 | 15 |
| TpMuguga_03g00311 | 11 | 7.5 |
| TpMuguga_03g00311 | 12 | 0 |
| TpMuguga_03g00311 | 13 | 0 |
| TpMuguga_03g00311 | 14 | 7.5 |
| TpMuguga_03g02190 | 1 | 0 |
| TpMuguga_03g02190 | 2 | 0 |
| TpMuguga_03g00314 | 1 | 0 |
| TpMuguga_03g00314 | 2 | 0.628158845 |
| TpMuguga_03g02265 | 1 | 0 |
| TpMuguga_03g02265 | 2 | 0 |
| TpMuguga_03g02260 | 1 | 0 |
| TpMuguga_03g02260 | 2 | 0 |
| TpMuguga_03g02260 | 3 | 0 |
| TpMuguga_03g02260 | 4 | 0 |
| TpMuguga_03g02260 | 5 | 0 |
| TpMuguga_03g02260 | 6 | 0 |
| TpMuguga_03g00318 | 1 | 3.714285714 |
| TpMuguga_03g00318 | 2 | 0 |
| TpMuguga_03g00318 | 3 | 0 |
| TpMuguga_03g00318 | 4 | 0 |
| TpMuguga_03g00318 | 5 | 0 |
| TpMuguga_03g00318 | 6 | 0 |
| TpMuguga_03g00318 | 7 | 0 |
| TpMuguga_03g00318 | 8 | 0 |
| TpMuguga_03g00318 | 9 | 0 |
| TpMuguga_03g00318 | 10 | 0 |
| TpMuguga_03g00318 | 11 | 0 |
| TpMuguga_03g00318 | 12 | 0 |
| TpMuguga_03g02245 | 1 | 7.5 |
| TpMuguga_03g02245 | 2 | 2.25 |
| TpMuguga_03g02245 | 3 | 2.25 |
| TpMuguga_03g02245 | 4 | 0.75 |
| TpMuguga_03g02245 | 5 | 0 |
| TpMuguga_03g00320 | 1 | 0.6875 |
| TpMuguga_03g00320 | 2 | 0.5625 |
| TpMuguga_03g02505 | 1 | 4.0625 |
| TpMuguga_03g02505 | 2 | 6.404411765 |
| TpMuguga_03g02505 | 3 | 8.794117647 |
| TpMuguga_03g02505 | 4 | 1.147058824 |
| TpMuguga_03g02505 | 5 | 0 |
| TpMuguga_03g02505 | 6 | 0 |

| | | |
|---|---|---|
| TpMuguga_03g02505 | 7 | 0 |
| TpMuguga_03g02505 | 8 | 0 |
| TpMuguga_03g02505 | 9 | 0 |
| TpMuguga_03g02505 | 10 | 0 |
| TpMuguga_03g02505 | 11 | 0 |
| TpMuguga_03g02505 | 12 | 0 |
| TpMuguga_03g02290 | 1 | 0 |
| TpMuguga_03g02290 | 2 | 0 |
| TpMuguga_03g02290 | 3 | 0 |
| TpMuguga_03g02290 | 4 | 0 |
| TpMuguga_03g02290 | 5 | 0 |
| TpMuguga_03g02290 | 6 | 0 |
| TpMuguga_03g02290 | 7 | 0 |
| TpMuguga_03g02290 | 8 | 0 |
| TpMuguga_03g02290 | 9 | 0 |
| TpMuguga_03g00327 | 1 | 0 |
| TpMuguga_03g00327 | 2 | 0 |
| TpMuguga_03g00327 | 3 | 0 |
| TpMuguga_03g00327 | 4 | 0 |
| TpMuguga_03g00327 | 5 | 0 |
| TpMuguga_03g00329 | 1 | 1.454545455 |
| TpMuguga_03g00330 | 1 | 0 |
| TpMuguga_03g00331 | 1 | 0 |
| TpMuguga_03g00332 | 1 | 0 |
| TpMuguga_03g00333 | 1 | 0 |
| TpMuguga_03g00333 | 2 | 0.714285714 |
| TpMuguga_03g00333 | 3 | 0 |
| TpMuguga_03g00333 | 4 | 2.142857143 |
| TpMuguga_03g00334 | 1 | 0 |
| TpMuguga_03g00334 | 2 | 0 |
| TpMuguga_03g02680 | 1 | 0.233333333 |
| TpMuguga_03g02680 | 2 | 0 |
| TpMuguga_03g02680 | 3 | 0 |
| TpMuguga_03g00344 | 1 | 0 |
| TpMuguga_03g00344 | 2 | 1.838709677 |
| TpMuguga_03g02620 | 1 | 0 |
| TpMuguga_03g02620 | 2 | 0 |
| TpMuguga_03g02620 | 3 | 0 |
| TpMuguga_03g02620 | 4 | 0 |
| TpMuguga_03g02620 | 5 | 0 |
| TpMuguga_03g02620 | 6 | 0 |
| TpMuguga_03g02620 | 7 | 0 |

| | | |
|---|---|---|
| TpMuguga_03g02255 | 1 | 0 |
| TpMuguga_03g02255 | 2 | 0 |
| TpMuguga_03g02255 | 3 | 0 |
| TpMuguga_03g02255 | 4 | 0 |
| TpMuguga_03g02255 | 5 | 0 |
| TpMuguga_03g00346 | 1 | 0 |
| TpMuguga_03g00346 | 2 | 0 |
| TpMuguga_03g00346 | 3 | 0 |
| TpMuguga_03g00346 | 4 | 0 |
| TpMuguga_03g00346 | 5 | 0 |
| TpMuguga_03g00346 | 6 | 0 |
| TpMuguga_03g00346 | 7 | 0 |
| TpMuguga_03g00348 | 1 | 0.034090909 |
| TpMuguga_03g00348 | 2 | 0.272727273 |
| TpMuguga_03g00895 | 1 | 0 |
| TpMuguga_03g00895 | 2 | 0 |
| TpMuguga_03g00349 | 1 | 0.321428571 |
| TpMuguga_03g00349 | 2 | 0 |
| TpMuguga_03g00349 | 3 | 0 |
| TpMuguga_03g00349 | 4 | 0 |
| TpMuguga_03g00349 | 5 | 0 |
| TpMuguga_03g00349 | 6 | 3.857142857 |
| TpMuguga_03g00349 | 7 | 3.535714286 |
| TpMuguga_03g00349 | 8 | 0.321428571 |
| TpMuguga_03g00350 | 1 | 0 |
| TpMuguga_03g00350 | 2 | 0 |
| TpMuguga_03g00350 | 3 | 0 |
| TpMuguga_03g00350 | 4 | 0 |
| TpMuguga_03g02600 | 1 | 0.774193548 |
| TpMuguga_03g00351 | 1 | 0 |
| TpMuguga_03g00351 | 2 | 0 |
| TpMuguga_03g00351 | 3 | 0 |
| TpMuguga_03g00351 | 4 | 0 |
| TpMuguga_03g00351 | 5 | 0 |
| TpMuguga_03g00351 | 6 | 0 |
| TpMuguga_03g00351 | 7 | 0 |
| TpMuguga_03g00351 | 8 | 0 |
| TpMuguga_03g00351 | 9 | 0 |
| TpMuguga_03g02360 | 1 | 0 |
| TpMuguga_03g02360 | 2 | 0 |
| TpMuguga_03g02360 | 3 | 0 |
| TpMuguga_03g02360 | 4 | 0 |

| | | |
|---|---|---|
| TpMuguga_03g02360 | 5 | 0 |
| TpMuguga_03g02360 | 6 | 0 |
| TpMuguga_03g02360 | 7 | 0 |
| TpMuguga_03g02360 | 8 | 0 |
| TpMuguga_03g02360 | 9 | 0 |
| TpMuguga_03g02360 | 10 | 0 |
| TpMuguga_03g02360 | 11 | 4.333333333 |
| TpMuguga_03g02360 | 12 | 13 |
| TpMuguga_03g02470 | 1 | 0 |
| TpMuguga_03g02470 | 2 | 0 |
| TpMuguga_03g02470 | 3 | 2.5 |
| TpMuguga_03g02470 | 4 | 27.5 |
| TpMuguga_03g00357 | 1 | 0 |
| TpMuguga_03g00357 | 2 | 0 |
| TpMuguga_03g00357 | 3 | 0 |
| TpMuguga_03g00357 | 4 | 0 |
| TpMuguga_03g00357 | 5 | 8 |
| TpMuguga_03g00358 | 1 | 0 |
| TpMuguga_03g00358 | 2 | 0 |
| TpMuguga_03g00358 | 3 | 0 |
| TpMuguga_03g00358 | 4 | 0 |
| TpMuguga_03g02235 | 1 | 0 |
| TpMuguga_03g02235 | 2 | 0 |
| TpMuguga_03g02205 | 1 | 0 |
| TpMuguga_03g02205 | 2 | 0 |
| TpMuguga_03g02205 | 3 | 0 |
| TpMuguga_03g02205 | 4 | 0 |
| TpMuguga_03g00362 | 1 | 12.54545455 |
| TpMuguga_03g00362 | 2 | 0.181818182 |
| TpMuguga_03g00362 | 3 | 0 |
| TpMuguga_03g00362 | 4 | 0 |
| TpMuguga_03g00362 | 5 | 0 |
| TpMuguga_03g00362 | 6 | 0 |
| TpMuguga_03g00362 | 7 | 0 |
| TpMuguga_03g00362 | 8 | 0 |
| TpMuguga_03g00362 | 9 | 0 |
| TpMuguga_03g00361 | 1 | 0 |
| TpMuguga_03g00361 | 2 | 0 |
| TpMuguga_03g00361 | 3 | 0 |
| TpMuguga_03g00364 | 1 | 0 |
| TpMuguga_03g00364 | 2 | 0 |
| TpMuguga_03g00366 | 1 | 0 |

| | | |
|---|---|---|
| TpMuguga_03g00366 | 2 | 0 |
| TpMuguga_03g00369 | 1 | 4.655172414 |
| TpMuguga_03g00369 | 2 | 0.310344828 |
| TpMuguga_03g00371 | 1 | 0 |
| TpMuguga_03g00371 | 2 | 0 |
| TpMuguga_03g00372 | 1 | 0 |
| TpMuguga_03g00373 | 1 | 0.100217865 |
| TpMuguga_03g00374 | 1 | 0 |
| TpMuguga_03g00374 | 2 | 0 |
| TpMuguga_03g00374 | 3 | 0 |
| TpMuguga_03g00374 | 4 | 0 |
| TpMuguga_03g00374 | 5 | 0 |
| TpMuguga_03g00374 | 6 | 0 |
| TpMuguga_03g00374 | 7 | 0 |
| TpMuguga_03g00374 | 8 | 0 |
| TpMuguga_03g00375 | 1 | 0.238095238 |
| TpMuguga_03g00375 | 2 | 0 |
| TpMuguga_03g00377 | 1 | 0 |
| TpMuguga_03g00377 | 2 | 2.857142857 |
| TpMuguga_03g00377 | 3 | 0 |
| TpMuguga_03g02330 | 1 | 0 |
| TpMuguga_03g02330 | 2 | 0 |
| TpMuguga_03g02330 | 3 | 0 |
| TpMuguga_03g02330 | 4 | 0 |
| TpMuguga_03g02330 | 5 | 0 |
| TpMuguga_03g02330 | 6 | 0 |
| TpMuguga_03g02330 | 7 | 0 |
| TpMuguga_03g02330 | 8 | 0 |
| TpMuguga_03g02330 | 9 | 0 |
| TpMuguga_03g02330 | 10 | 0 |
| TpMuguga_03g02330 | 11 | 0 |
| TpMuguga_03g02330 | 12 | 0 |
| TpMuguga_03g02330 | 13 | 0 |
| TpMuguga_03g02330 | 14 | 0 |
| TpMuguga_03g02330 | 15 | 0 |
| TpMuguga_03g02330 | 16 | 0 |
| TpMuguga_03g02330 | 17 | 0 |
| TpMuguga_03g02330 | 18 | 0 |
| TpMuguga_03g02330 | 19 | 0 |
| TpMuguga_03g00383 | 1 | 0 |
| TpMuguga_03g00383 | 2 | 0 |
| TpMuguga_03g00383 | 3 | 0 |

| | | |
|---|---|---|
| TpMuguga_03g00383 | 4 | 0 |
| TpMuguga_03g00384 | 1 | 0 |
| TpMuguga_03g00382 | 1 | 0.392156863 |
| TpMuguga_03g00382 | 2 | 0 |
| TpMuguga_03g00382 | 3 | 0 |
| TpMuguga_03g00382 | 4 | 6.960784314 |
| TpMuguga_03g00385 | 1 | 0 |
| TpMuguga_03g00385 | 2 | 0 |
| TpMuguga_03g00385 | 3 | 0 |
| TpMuguga_03g00385 | 4 | 0 |
| TpMuguga_03g00385 | 5 | 0 |
| TpMuguga_03g00386 | 1 | 3 |
| TpMuguga_03g00386 | 2 | 0 |
| TpMuguga_03g00387 | 1 | 0 |
| TpMuguga_03g00387 | 2 | 0 |
| TpMuguga_03g00387 | 3 | 0 |
| TpMuguga_03g00387 | 4 | 0 |
| TpMuguga_03g00387 | 5 | 0 |
| TpMuguga_03g00387 | 6 | 0 |
| TpMuguga_03g00387 | 7 | 0 |
| TpMuguga_03g00387 | 8 | 0 |
| TpMuguga_03g00387 | 9 | 0 |
| TpMuguga_03g00387 | 10 | 9 |
| TpMuguga_03g00387 | 11 | 12 |
| TpMuguga_03g00388 | 1 | 0 |
| TpMuguga_03g00388 | 2 | 0 |
| TpMuguga_03g00388 | 3 | 0 |
| TpMuguga_03g00388 | 4 | 0.196078431 |
| TpMuguga_03g00390 | 1 | 0 |
| TpMuguga_03g00392 | 1 | 0 |
| TpMuguga_03g00392 | 2 | 0 |
| TpMuguga_03g00393 | 1 | 0 |
| TpMuguga_03g00393 | 2 | 1.043478261 |
| TpMuguga_03g00394 | 1 | 0 |
| TpMuguga_03g00394 | 2 | 0 |
| TpMuguga_03g00394 | 3 | 0 |
| TpMuguga_03g00394 | 4 | 0 |
| TpMuguga_03g00394 | 5 | 0 |
| TpMuguga_03g00395 | 1 | 0 |
| TpMuguga_03g00395 | 2 | 0.021126761 |
| TpMuguga_03g00395 | 3 | 0 |
| TpMuguga_03g00395 | 4 | 0.126760563 |

| | | |
|---|---|---|
| TpMuguga_03g00395 | 5 | 0.866197183 |
| TpMuguga_03g00396 | 1 | 0 |
| TpMuguga_03g00396 | 2 | 0 |
| TpMuguga_03g00396 | 3 | 0 |
| TpMuguga_03g00397 | 1 | 0 |
| TpMuguga_03g00398 | 1 | 0 |
| TpMuguga_03g00398 | 2 | 0 |
| TpMuguga_03g00398 | 3 | 0 |
| TpMuguga_03g00398 | 4 | 0 |
| TpMuguga_03g00398 | 5 | 0 |
| TpMuguga_03g00398 | 6 | 0 |
| TpMuguga_03g00398 | 7 | 0 |
| TpMuguga_03g00401 | 1 | 0 |
| TpMuguga_03g02640 | 1 | 0 |
| TpMuguga_03g00404 | 1 | 0.355932203 |
| TpMuguga_03g00404 | 2 | 0 |
| TpMuguga_03g00404 | 3 | 0.118644068 |
| TpMuguga_03g00404 | 4 | 1.06779661 |
| TpMuguga_03g00404 | 5 | 1.542372881 |
| TpMuguga_03g00404 | 6 | 2.728813559 |
| TpMuguga_03g02405 | 1 | 0 |
| TpMuguga_03g02405 | 2 | 0 |
| TpMuguga_03g02405 | 3 | 0 |
| TpMuguga_03g02695 | 1 | 4.666666667 |
| TpMuguga_03g02695 | 2 | 5.833333333 |
| TpMuguga_03g02695 | 3 | 0 |
| TpMuguga_03g02695 | 4 | 2.333333333 |
| TpMuguga_03g02695 | 5 | 0 |
| TpMuguga_03g02695 | 6 | 0 |
| TpMuguga_03g00408 | 1 | 0 |
| TpMuguga_03g02690 | 1 | 0 |
| TpMuguga_03g02690 | 2 | 0 |
| TpMuguga_03g02690 | 3 | 0 |
| TpMuguga_03g02690 | 4 | 0 |
| TpMuguga_03g02690 | 5 | 0 |
| TpMuguga_03g02690 | 6 | 0 |
| TpMuguga_03g02690 | 7 | 0 |
| TpMuguga_03g00411 | 1 | 0 |
| TpMuguga_03g00411 | 2 | 0 |
| TpMuguga_03g00411 | 3 | 0 |
| TpMuguga_03g00413 | 1 | 0.705722071 |
| TpMuguga_03g00413 | 2 | 0.190735695 |

| | | |
|---|---|---|
| TpMuguga_03g00413 | 3 | 0.286103542 |
| TpMuguga_03g00413 | 4 | 0.228882834 |
| TpMuguga_03g00413 | 5 | 0.534059946 |
| TpMuguga_03g00413 | 6 | 0 |
| TpMuguga_03g00412 | 1 | 0.037151703 |
| TpMuguga_03g00412 | 2 | 0.092879257 |
| TpMuguga_03g00412 | 3 | 1.46749226 |
| TpMuguga_03g00415 | 1 | 0 |
| TpMuguga_03g00415 | 2 | 0 |
| TpMuguga_03g00415 | 3 | 0 |
| TpMuguga_03g00415 | 4 | 0 |
| TpMuguga_03g00417 | 1 | 0 |
| TpMuguga_03g00417 | 2 | 0 |
| TpMuguga_03g00417 | 3 | 5 |
| TpMuguga_03g00417 | 4 | 0 |
| TpMuguga_03g00420 | 1 | 5.582037997 |
| TpMuguga_03g00420 | 2 | 0.027633851 |
| TpMuguga_03g00420 | 3 | 0 |
| TpMuguga_03g00418 | 1 | 0.253846154 |
| TpMuguga_03g00418 | 2 | 1.811538462 |
| TpMuguga_03g00419 | 1 | 0.869565217 |
| TpMuguga_03g00419 | 2 | 0.52173913 |
| TpMuguga_03g00419 | 3 | 0.695652174 |
| TpMuguga_03g00421 | 1 | 0 |
| TpMuguga_03g00421 | 2 | 27 |
| TpMuguga_03g00421 | 3 | 4.5 |
| TpMuguga_03g00421 | 4 | 0 |
| TpMuguga_03g00421 | 5 | 0 |
| TpMuguga_03g00423 | 1 | 3.968181818 |
| TpMuguga_03g00423 | 2 | 2.413636364 |
| TpMuguga_03g00423 | 3 | 0.327272727 |
| TpMuguga_03g00423 | 4 | 0 |
| TpMuguga_03g00423 | 5 | 0 |
| TpMuguga_03g00423 | 6 | 0 |
| TpMuguga_03g00423 | 7 | 0.204545455 |
| TpMuguga_03g00423 | 8 | 0.327272727 |
| TpMuguga_03g02645 | 1 | 0 |
| TpMuguga_03g02645 | 2 | 0 |
| TpMuguga_03g02645 | 3 | 0 |
| TpMuguga_03g02645 | 4 | 0 |
| TpMuguga_03g02645 | 5 | 0 |
| TpMuguga_03g02645 | 6 | 0 |

| | | |
|---|---|---|
| TpMuguga_03g02645 | 7 | 0 |
| TpMuguga_03g02645 | 8 | 0 |
| TpMuguga_03g02645 | 9 | 0 |
| TpMuguga_03g02645 | 10 | 0 |
| TpMuguga_03g02645 | 11 | 0 |
| TpMuguga_03g02645 | 12 | 0 |
| TpMuguga_03g02645 | 13 | 0 |
| TpMuguga_03g02645 | 14 | 0 |
| TpMuguga_03g02645 | 15 | 0 |
| TpMuguga_03g00425 | 1 | 3 |
| TpMuguga_03g00425 | 2 | 0 |
| TpMuguga_03g00425 | 3 | 0 |
| TpMuguga_03g02195 | 1 | 2.1875 |
| TpMuguga_03g02195 | 2 | 0 |
| TpMuguga_03g02195 | 3 | 0 |
| TpMuguga_03g02195 | 4 | 0 |
| TpMuguga_03g02195 | 5 | 0 |
| TpMuguga_03g02195 | 6 | 0 |
| TpMuguga_03g00429 | 1 | 3.878787879 |
| TpMuguga_03g00429 | 2 | 1.090909091 |
| TpMuguga_03g00429 | 3 | 0 |
| TpMuguga_03g00430 | 1 | 0 |
| TpMuguga_03g00430 | 2 | 0 |
| TpMuguga_03g00430 | 3 | 1.285714286 |
| TpMuguga_03g00432 | 1 | 0 |
| TpMuguga_03g00432 | 2 | 0 |
| TpMuguga_03g00432 | 3 | 0 |
| TpMuguga_03g00432 | 4 | 0 |
| TpMuguga_03g00432 | 5 | 0 |
| TpMuguga_03g00432 | 6 | 0 |
| TpMuguga_03g00432 | 7 | 0 |
| TpMuguga_03g00432 | 8 | 0 |
| TpMuguga_03g00432 | 9 | 0 |
| TpMuguga_03g00433 | 1 | 0 |
| TpMuguga_03g00433 | 2 | 0 |
| TpMuguga_03g00433 | 3 | 0 |
| TpMuguga_03g00434 | 1 | 0 |
| TpMuguga_03g00434 | 2 | 0 |
| TpMuguga_03g00434 | 3 | 1.111111111 |
| TpMuguga_03g00435 | 1 | 0 |
| TpMuguga_03g00435 | 2 | 0 |
| TpMuguga_03g00435 | 3 | 0 |

| | | |
|---|---|---|
| TpMuguga_03g00436 | 1 | 0 |
| TpMuguga_03g00436 | 2 | 0 |
| TpMuguga_03g00436 | 3 | 0 |
| TpMuguga_03g00436 | 4 | 0 |
| TpMuguga_03g00436 | 5 | 0 |
| TpMuguga_03g00436 | 6 | 0 |
| TpMuguga_03g00436 | 7 | 0 |
| TpMuguga_03g00436 | 8 | 0 |
| TpMuguga_03g00436 | 9 | 0 |
| TpMuguga_03g00437 | 1 | 4 |
| TpMuguga_03g00437 | 2 | 4 |
| TpMuguga_03g00437 | 3 | 0 |
| TpMuguga_03g00438 | 1 | 0 |
| TpMuguga_03g00438 | 2 | 0 |
| TpMuguga_03g00439 | 1 | 0 |
| TpMuguga_03g00439 | 2 | 11.74468085 |
| TpMuguga_03g00442 | 1 | 0.260869565 |
| TpMuguga_03g00444 | 1 | 0 |
| TpMuguga_03g00444 | 2 | 0.342857143 |
| TpMuguga_03g00444 | 3 | 0.8 |
| TpMuguga_03g00443 | 1 | 0 |
| TpMuguga_03g00443 | 2 | 0 |
| TpMuguga_03g00443 | 3 | 0 |
| TpMuguga_03g00443 | 4 | 0 |
| TpMuguga_03g00443 | 5 | 0 |
| TpMuguga_03g00443 | 6 | 0 |
| TpMuguga_03g00443 | 7 | 0 |
| TpMuguga_03g00445 | 1 | 0 |
| TpMuguga_03g00445 | 2 | 0 |
| TpMuguga_03g00445 | 3 | 0 |
| TpMuguga_03g00445 | 4 | 0 |
| TpMuguga_03g00446 | 1 | 0 |
| TpMuguga_03g00446 | 2 | 0 |
| TpMuguga_03g00446 | 3 | 0 |
| TpMuguga_03g00447 | 1 | 3.888888889 |
| TpMuguga_03g00447 | 2 | 1.555555556 |
| TpMuguga_03g00447 | 3 | 5.444444444 |
| TpMuguga_03g00447 | 4 | 1.555555556 |
| TpMuguga_03g00447 | 5 | 2.333333333 |
| TpMuguga_03g00447 | 6 | 0 |
| TpMuguga_03g00448 | 1 | 1.428571429 |
| TpMuguga_03g00448 | 2 | 3.095238095 |

| | | |
|---|---|---|
| TpMuguga_03g00448 | 3 | 1.428571429 |
| TpMuguga_03g00448 | 4 | 0 |
| TpMuguga_03g00449 | 1 | 0 |
| TpMuguga_03g00449 | 2 | 0 |
| TpMuguga_03g00450 | 1 | 0 |
| TpMuguga_03g00450 | 2 | 0 |
| TpMuguga_03g00450 | 3 | 0 |
| TpMuguga_03g00451 | 1 | 0 |
| TpMuguga_03g00451 | 2 | 0.227272727 |
| TpMuguga_03g00451 | 3 | 0 |
| TpMuguga_03g00451 | 4 | 0 |
| TpMuguga_03g00452 | 1 | 0 |
| TpMuguga_03g00452 | 2 | 0 |
| TpMuguga_03g00452 | 3 | 0 |
| TpMuguga_03g00452 | 4 | 0 |
| TpMuguga_03g00452 | 5 | 0 |
| TpMuguga_03g00452 | 6 | 0 |
| TpMuguga_03g02625 | 1 | 0 |
| TpMuguga_03g02625 | 2 | 0 |
| TpMuguga_03g02625 | 3 | 0 |
| TpMuguga_03g02625 | 4 | 0 |
| TpMuguga_03g02625 | 5 | 0 |
| TpMuguga_03g02625 | 6 | 0 |
| TpMuguga_03g00455 | 1 | 0 |
| TpMuguga_03g00455 | 2 | 0 |
| TpMuguga_03g00455 | 3 | 0 |
| TpMuguga_03g00455 | 4 | 0 |
| TpMuguga_03g00455 | 5 | 0 |
| TpMuguga_03g00455 | 6 | 0 |
| TpMuguga_03g00455 | 7 | 0 |
| TpMuguga_03g00456 | 1 | 0 |
| TpMuguga_03g00456 | 2 | 0 |
| TpMuguga_03g00457 | 1 | 0 |
| TpMuguga_03g00457 | 2 | 0 |
| TpMuguga_03g00458 | 1 | 0 |
| TpMuguga_03g00458 | 2 | 0 |
| TpMuguga_03g00458 | 3 | 0 |
| TpMuguga_03g00458 | 4 | 0.5 |
| TpMuguga_03g00460 | 1 | 0 |
| TpMuguga_03g00896 | 1 | 0 |
| TpMuguga_03g00897 | 1 | 0.545454545 |
| TpMuguga_03g00898 | 1 | 0.243055556 |

| | | |
|---|---|---|
| TpMuguga_03g00899 | 1 | 0.904761905 |
| TpMuguga_03g00462 | 1 | 0 |
| TpMuguga_03g00462 | 2 | 0 |
| TpMuguga_03g00462 | 3 | 0 |
| TpMuguga_03g00462 | 4 | 1.280487805 |
| TpMuguga_03g00462 | 5 | 0.170731707 |
| TpMuguga_03g00462 | 6 | 1.62195122 |
| TpMuguga_03g00463 | 1 | 0.293333333 |
| TpMuguga_03g00903 | 1 | 0 |
| TpMuguga_03g00464 | 1 | 1.333333333 |
| TpMuguga_03g00464 | 2 | 10.66666667 |
| TpMuguga_03g00464 | 3 | 6.666666667 |
| TpMuguga_03g00464 | 4 | 0 |
| TpMuguga_03g00464 | 5 | 0 |
| TpMuguga_03g00464 | 6 | 0 |
| TpMuguga_03g00464 | 7 | 0 |
| TpMuguga_03g00465 | 1 | 0 |
| TpMuguga_03g00466 | 1 | 0 |
| TpMuguga_03g00467 | 1 | 0 |
| TpMuguga_03g00467 | 2 | 0 |
| TpMuguga_03g02615 | 1 | 60 |
| TpMuguga_03g02615 | 2 | 0 |
| TpMuguga_03g02615 | 3 | 0 |
| TpMuguga_03g02615 | 4 | 0 |
| TpMuguga_03g02615 | 5 | 0 |
| TpMuguga_03g02170 | 1 | 0 |
| TpMuguga_03g02170 | 2 | 0 |
| TpMuguga_03g00470 | 1 | 0 |
| TpMuguga_03g00470 | 2 | 0 |
| TpMuguga_03g00470 | 3 | 0.324324324 |
| TpMuguga_03g00470 | 4 | 0.486486486 |
| TpMuguga_03g00470 | 5 | 0 |
| TpMuguga_03g00470 | 6 | 0 |
| TpMuguga_03g00470 | 7 | 0 |
| TpMuguga_03g00470 | 8 | 0 |
| TpMuguga_03g00470 | 9 | 0.216216216 |
| TpMuguga_03g00472 | 1 | 0.302620803 |
| TpMuguga_03g00473 | 1 | 0.932038835 |
| TpMuguga_03g00473 | 2 | 1.165048544 |
| TpMuguga_03g00475 | 1 | 0 |
| TpMuguga_03g00476 | 1 | 0 |
| TpMuguga_03g00476 | 2 | 12.83333333 |

| | | |
|---|---|---|
| TpMuguga_03g00476 | 3 | 1.166666667 |
| TpMuguga_03g00476 | 4 | 0 |
| TpMuguga_03g00476 | 5 | 0 |
| TpMuguga_03g00476 | 6 | 0 |
| TpMuguga_03g02575 | 1 | 0 |
| TpMuguga_03g02575 | 2 | 0 |
| TpMuguga_03g00478 | 1 | 0 |
| TpMuguga_03g00479 | 1 | 0.114068441 |
| TpMuguga_03g00479 | 2 | 2.783269962 |
| TpMuguga_03g00479 | 3 | 0.730038023 |
| TpMuguga_03g00479 | 4 | 0.55513308 |
| TpMuguga_03g00479 | 5 | 0.030418251 |
| TpMuguga_03g00480 | 1 | 0 |
| TpMuguga_03g00480 | 2 | 0.570194384 |
| TpMuguga_03g00480 | 3 | 0 |
| TpMuguga_03g00480 | 4 | 0 |
| TpMuguga_03g00480 | 5 | 0 |
| TpMuguga_03g00480 | 6 | 0 |
| TpMuguga_03g00480 | 7 | 0.103671706 |
| TpMuguga_03g00481 | 1 | 0 |
| TpMuguga_03g00481 | 2 | 0 |
| TpMuguga_03g00481 | 3 | 0 |
| TpMuguga_03g00481 | 4 | 0 |
| TpMuguga_03g00481 | 5 | 0 |
| TpMuguga_03g00481 | 6 | 0 |
| TpMuguga_03g00481 | 7 | 0 |
| TpMuguga_03g00481 | 8 | 0 |
| TpMuguga_03g00484 | 1 | 3.125 |
| TpMuguga_03g00484 | 2 | 0.78125 |
| TpMuguga_03g00484 | 3 | 0 |
| TpMuguga_03g00484 | 4 | 0 |
| TpMuguga_03g00484 | 5 | 0.078125 |
| TpMuguga_03g00484 | 6 | 0.390625 |
| TpMuguga_03g00484 | 7 | 0 |
| TpMuguga_03g00484 | 8 | 0 |
| TpMuguga_03g00484 | 9 | 0 |
| TpMuguga_03g00483 | 1 | 0 |
| TpMuguga_03g00483 | 2 | 0 |
| TpMuguga_03g00483 | 3 | 0 |
| TpMuguga_03g00483 | 4 | 0 |
| TpMuguga_03g00483 | 5 | 0 |
| TpMuguga_03g00486 | 1 | 1.5 |

| | | |
|---|---|---|
| TpMuguga_03g00486 | 2 | 0 |
| TpMuguga_03g00487 | 1 | 0.280701754 |
| TpMuguga_03g00487 | 2 | 0.035087719 |
| TpMuguga_03g00487 | 3 | 1.83625731 |
| TpMuguga_03g00488 | 1 | 0 |
| TpMuguga_03g00489 | 1 | 0 |
| TpMuguga_03g00489 | 2 | 0.449197861 |
| TpMuguga_03g00489 | 3 | 0.406417112 |
| TpMuguga_03g00489 | 4 | 0.320855615 |
| TpMuguga_03g00489 | 5 | 0.042780749 |
| TpMuguga_03g00489 | 6 | 0.449197861 |
| TpMuguga_03g00489 | 7 | 1.839572193 |
| TpMuguga_03g00904 | 1 | 0 |
| TpMuguga_03g00904 | 2 | 0 |
| TpMuguga_03g00904 | 3 | 0 |
| TpMuguga_03g00904 | 4 | 1.285714286 |
| TpMuguga_03g00904 | 5 | 19.28571429 |
| TpMuguga_03g00904 | 6 | 0 |
| TpMuguga_03g00904 | 7 | 0 |
| TpMuguga_03g00904 | 8 | 9 |
| TpMuguga_03g00490 | 1 | 0 |
| TpMuguga_03g00490 | 2 | 0 |
| TpMuguga_03g00490 | 3 | 0.28125 |
| TpMuguga_03g00490 | 4 | 3.65625 |
| TpMuguga_03g00490 | 5 | 1.96875 |
| TpMuguga_03g00490 | 6 | 0 |
| TpMuguga_03g00490 | 7 | 0 |
| TpMuguga_03g00490 | 8 | 0 |
| TpMuguga_03g00491 | 1 | 0 |
| TpMuguga_03g00491 | 2 | 0 |
| TpMuguga_03g00491 | 3 | 5 |
| TpMuguga_03g00491 | 4 | 2.5 |
| TpMuguga_03g00491 | 5 | 0 |
| TpMuguga_03g00491 | 6 | 0 |
| TpMuguga_03g00491 | 7 | 0 |
| TpMuguga_03g00491 | 8 | 0 |
| TpMuguga_03g00491 | 9 | 0 |
| TpMuguga_03g02580 | 1 | 0 |
| TpMuguga_03g02580 | 2 | 0 |
| TpMuguga_03g00495 | 1 | 0 |
| TpMuguga_03g00495 | 2 | 0.002516779 |
| TpMuguga_03g00500 | 1 | 0 |

| | | |
|---|---|---|
| TpMuguga_03g00500 | 2 | 0 |
| TpMuguga_03g00500 | 3 | 0 |
| TpMuguga_03g00500 | 4 | 0 |
| TpMuguga_03g00500 | 5 | 0 |
| TpMuguga_03g00501 | 1 | 0 |
| TpMuguga_03g00501 | 2 | 0 |
| TpMuguga_03g00501 | 3 | 0 |
| TpMuguga_03g00501 | 4 | 0 |
| TpMuguga_03g00501 | 5 | 0 |
| TpMuguga_03g00501 | 6 | 0 |
| TpMuguga_03g00501 | 7 | 0 |
| TpMuguga_03g00501 | 8 | 36 |
| TpMuguga_03g00502 | 1 | 0 |
| TpMuguga_03g00502 | 2 | 0 |
| TpMuguga_03g00502 | 3 | 0 |
| TpMuguga_03g00502 | 4 | 0 |
| TpMuguga_03g00502 | 5 | 0 |
| TpMuguga_03g00502 | 6 | 0 |
| TpMuguga_03g00502 | 7 | 0 |
| TpMuguga_03g00502 | 8 | 0 |
| TpMuguga_03g00502 | 9 | 0 |
| TpMuguga_03g00502 | 10 | 0 |
| TpMuguga_03g00502 | 11 | 0 |
| TpMuguga_03g00502 | 12 | 0 |
| TpMuguga_03g00502 | 13 | 0 |
| TpMuguga_03g00503 | 1 | 0.23553719 |
| TpMuguga_03g00505 | 1 | 0 |
| TpMuguga_03g00506 | 1 | 2.727272727 |
| TpMuguga_03g00506 | 2 | 0 |
| TpMuguga_03g00506 | 3 | 0 |
| TpMuguga_03g00506 | 4 | 0 |
| TpMuguga_03g00506 | 5 | 0 |
| TpMuguga_03g00508 | 1 | 0 |
| TpMuguga_03g00508 | 2 | 0 |
| TpMuguga_03g00508 | 3 | 0 |
| TpMuguga_03g00507 | 1 | 0 |
| TpMuguga_03g00507 | 2 | 0 |
| TpMuguga_03g00507 | 3 | 0 |
| TpMuguga_03g00507 | 4 | 0 |
| TpMuguga_03g00507 | 5 | 0 |
| TpMuguga_03g00507 | 6 | 0 |
| TpMuguga_03g00507 | 7 | 0 |

| | | |
|---|---|---|
| TpMuguga_03g00509 | 1 | 0 |
| TpMuguga_03g00509 | 2 | 0 |
| TpMuguga_03g00510 | 1 | 0 |
| TpMuguga_03g00510 | 2 | 0 |
| TpMuguga_03g00511 | 1 | 0 |
| TpMuguga_03g00511 | 2 | 0 |
| TpMuguga_03g00511 | 3 | 0 |
| TpMuguga_03g00511 | 4 | 0 |
| TpMuguga_03g00511 | 5 | 0 |
| TpMuguga_03g00511 | 6 | 0 |
| TpMuguga_03g00511 | 7 | 0 |
| TpMuguga_03g00511 | 8 | 0 |
| TpMuguga_03g00512 | 1 | 0 |
| TpMuguga_03g00513 | 1 | 3.409090909 |
| TpMuguga_03g00513 | 2 | 0 |
| TpMuguga_03g00513 | 3 | 0.113636364 |
| TpMuguga_03g00513 | 4 | 0 |
| TpMuguga_03g00515 | 1 | 0 |
| TpMuguga_03g00515 | 2 | 0 |
| TpMuguga_03g00515 | 3 | 0 |
| TpMuguga_03g00515 | 4 | 0 |
| TpMuguga_03g00514 | 1 | 0 |
| TpMuguga_03g00514 | 2 | 0 |
| TpMuguga_03g00514 | 3 | 0 |
| TpMuguga_03g00514 | 4 | 0 |
| TpMuguga_03g00514 | 5 | 0 |
| TpMuguga_03g00516 | 1 | 24 |
| TpMuguga_03g00516 | 2 | 2 |
| TpMuguga_03g00516 | 3 | 0 |
| TpMuguga_03g00516 | 4 | 0 |
| TpMuguga_03g00516 | 5 | 0 |
| TpMuguga_03g00517 | 1 | 0 |
| TpMuguga_03g00517 | 2 | 0 |
| TpMuguga_03g00517 | 3 | 0 |
| TpMuguga_03g00517 | 4 | 0 |
| TpMuguga_03g00517 | 5 | 0 |
| TpMuguga_03g00517 | 6 | 0 |
| TpMuguga_03g00517 | 7 | 0 |
| TpMuguga_03g00517 | 8 | 0 |
| TpMuguga_03g00517 | 9 | 0 |
| TpMuguga_03g02365 | 1 | 0 |
| TpMuguga_03g00519 | 1 | 0 |

| | | |
|---|---|---|
| TpMuguga_03g00519 | 2 | 0 |
| TpMuguga_03g00521 | 1 | 0 |
| TpMuguga_03g00521 | 2 | 0 |
| TpMuguga_03g00521 | 3 | 0 |
| TpMuguga_03g00521 | 4 | 0 |
| TpMuguga_03g00521 | 5 | 0 |
| TpMuguga_03g00523 | 1 | 0.807692308 |
| TpMuguga_03g00523 | 2 | 0 |
| TpMuguga_03g00523 | 3 | 0 |
| TpMuguga_03g00523 | 4 | 0 |
| TpMuguga_03g00523 | 5 | 0 |
| TpMuguga_03g00523 | 6 | 0.807692308 |
| TpMuguga_03g00522 | 1 | 0.144 |
| TpMuguga_03g00522 | 2 | 0.168 |
| TpMuguga_03g00524 | 1 | 2.903225806 |
| TpMuguga_03g00524 | 2 | 0 |
| TpMuguga_03g00524 | 3 | 0 |
| TpMuguga_03g00524 | 4 | 0 |
| TpMuguga_03g00524 | 5 | 0 |
| TpMuguga_03g00524 | 6 | 0.322580645 |
| TpMuguga_03g00524 | 7 | 1.935483871 |
| TpMuguga_03g00524 | 8 | 0 |
| TpMuguga_03g00524 | 9 | 0 |
| TpMuguga_03g00527 | 1 | 1.020151134 |
| TpMuguga_03g00527 | 2 | 1.734256927 |
| TpMuguga_03g00529 | 1 | 0 |
| TpMuguga_03g00529 | 2 | 0 |
| TpMuguga_03g00529 | 3 | 0 |
| TpMuguga_03g00529 | 4 | 0 |
| TpMuguga_03g00529 | 5 | 0 |
| TpMuguga_03g00529 | 6 | 0 |
| TpMuguga_03g00529 | 7 | 0 |
| TpMuguga_03g00530 | 1 | 0 |
| TpMuguga_03g00530 | 2 | 0 |
| TpMuguga_03g00530 | 3 | 0 |
| TpMuguga_03g00530 | 4 | 0 |
| TpMuguga_03g00530 | 5 | 0 |
| TpMuguga_03g00530 | 6 | 0 |
| TpMuguga_03g00531 | 1 | 0 |
| TpMuguga_03g00531 | 2 | 0 |
| TpMuguga_03g00531 | 3 | 0 |
| TpMuguga_03g00532 | 1 | 0 |

| | | |
|---|---|---|
| TpMuguga_03g00532 | 2 | 0 |
| TpMuguga_03g00532 | 3 | 0 |
| TpMuguga_03g00532 | 4 | 0.71641791 |
| TpMuguga_03g00532 | 5 | 0.71641791 |
| TpMuguga_03g00532 | 6 | 0.71641791 |
| TpMuguga_03g02225 | 1 | 0 |
| TpMuguga_03g02225 | 2 | 0 |
| TpMuguga_03g02225 | 3 | 4.363636364 |
| TpMuguga_03g02225 | 4 | 0 |
| TpMuguga_03g02225 | 5 | 0 |
| TpMuguga_03g02225 | 6 | 0 |
| TpMuguga_03g02225 | 7 | 0 |
| TpMuguga_03g02225 | 8 | 0 |
| TpMuguga_03g02225 | 9 | 0 |
| TpMuguga_03g02225 | 10 | 0 |
| TpMuguga_03g02225 | 11 | 0 |
| TpMuguga_03g02225 | 12 | 0 |
| TpMuguga_03g02225 | 13 | 0 |
| TpMuguga_03g02225 | 14 | 0 |
| TpMuguga_03g02225 | 15 | 7.272727273 |
| TpMuguga_03g00537 | 1 | 0 |
| TpMuguga_03g00537 | 2 | 0 |
| TpMuguga_03g00537 | 3 | 0 |
| TpMuguga_03g00537 | 4 | 0 |
| TpMuguga_03g00539 | 1 | 0 |
| TpMuguga_03g00540 | 1 | 18 |
| TpMuguga_03g00540 | 2 | 0 |
| TpMuguga_03g00541 | 1 | 0 |
| TpMuguga_03g00541 | 2 | 0 |
| TpMuguga_03g00541 | 3 | 0.009222661 |
| TpMuguga_03g00541 | 4 | 0.026350461 |
| TpMuguga_03g00542 | 1 | 0 |
| TpMuguga_03g00542 | 2 | 0 |
| TpMuguga_03g00542 | 3 | 1.266666667 |
| TpMuguga_03g00542 | 4 | 0 |
| TpMuguga_03g00543 | 1 | 2.815884477 |
| TpMuguga_03g00543 | 2 | 0.411552347 |
| TpMuguga_03g00543 | 3 | 3.444043321 |
| TpMuguga_03g00543 | 4 | 0.303249097 |
| TpMuguga_03g00543 | 5 | 0 |
| TpMuguga_03g00544 | 1 | 6 |
| TpMuguga_03g00544 | 2 | 0 |

| | | |
|---|---|---|
| TpMuguga_03g00544 | 3 | 0 |
| TpMuguga_03g00544 | 4 | 3.857142857 |
| TpMuguga_03g00544 | 5 | 2.142857143 |
| TpMuguga_03g00545 | 1 | 0 |
| TpMuguga_03g00545 | 2 | 0 |
| TpMuguga_03g00545 | 3 | 0 |
| TpMuguga_03g00545 | 4 | 2 |
| TpMuguga_03g00545 | 5 | 8.571428571 |
| TpMuguga_03g00546 | 1 | 0 |
| TpMuguga_03g00547 | 1 | 0.533333333 |
| TpMuguga_03g00548 | 1 | 0 |
| TpMuguga_03g00548 | 2 | 0 |
| TpMuguga_03g00548 | 3 | 0 |
| TpMuguga_03g00548 | 4 | 0 |
| TpMuguga_03g00549 | 1 | 0 |
| TpMuguga_03g00549 | 2 | 0 |
| TpMuguga_03g00549 | 3 | 0 |
| TpMuguga_03g00549 | 4 | 0 |
| TpMuguga_03g00549 | 5 | 0 |
| TpMuguga_03g00549 | 6 | 0 |
| TpMuguga_03g00549 | 7 | 0 |
| TpMuguga_03g00549 | 8 | 0 |
| TpMuguga_03g00549 | 9 | 0 |
| TpMuguga_03g00549 | 10 | 0 |
| TpMuguga_03g00549 | 11 | 0 |
| TpMuguga_03g00549 | 12 | 0 |
| TpMuguga_03g00549 | 13 | 0 |
| TpMuguga_03g00549 | 14 | 0 |
| TpMuguga_03g00550 | 1 | 0.025359256 |
| TpMuguga_03g00550 | 2 | 1.78021978 |
| TpMuguga_03g00550 | 3 | 0.446322908 |
| TpMuguga_03g00550 | 4 | 2.429416737 |
| TpMuguga_03g00550 | 5 | 0.136939983 |
| TpMuguga_03g00552 | 1 | 0 |
| TpMuguga_03g00551 | 1 | 0 |
| TpMuguga_03g00551 | 2 | 0 |
| TpMuguga_03g00551 | 3 | 0 |
| TpMuguga_03g00553 | 1 | 0 |
| TpMuguga_03g00553 | 2 | 0 |
| TpMuguga_03g00553 | 3 | 0.198170732 |
| TpMuguga_03g00553 | 4 | 2.136051829 |
| TpMuguga_03g02220 | 1 | 3.333333333 |

| | | |
|---|---|---|
| TpMuguga_03g02220 | 2 | 0.133333333 |
| TpMuguga_03g02230 | 1 | 0 |
| TpMuguga_03g02230 | 2 | 0 |
| TpMuguga_03g02540 | 1 | 0 |
| TpMuguga_03g02540 | 2 | 0 |
| TpMuguga_03g02540 | 3 | 0 |
| TpMuguga_03g02540 | 4 | 0 |
| TpMuguga_03g02540 | 5 | 0 |
| TpMuguga_03g02540 | 6 | 0 |
| TpMuguga_03g02540 | 7 | 0 |
| TpMuguga_03g02540 | 8 | 0 |
| TpMuguga_03g02540 | 9 | 0 |
| TpMuguga_03g02540 | 10 | 0 |
| TpMuguga_03g02540 | 11 | 0 |
| TpMuguga_03g02540 | 12 | 0 |
| TpMuguga_03g02180 | 1 | 6 |
| TpMuguga_03g02180 | 2 | 0 |
| TpMuguga_03g02180 | 3 | 0 |
| TpMuguga_03g02180 | 4 | 0 |
| TpMuguga_03g02180 | 5 | 0 |
| TpMuguga_03g00558 | 1 | 0.7 |
| TpMuguga_03g00558 | 2 | 0.6 |
| TpMuguga_03g00558 | 3 | 0.5 |
| TpMuguga_03g02345 | 1 | 0.615384615 |
| TpMuguga_03g00559 | 1 | 2.866666667 |
| TpMuguga_03g00559 | 2 | 0 |
| TpMuguga_03g00559 | 3 | 0 |
| TpMuguga_03g00560 | 1 | 0 |
| TpMuguga_03g00560 | 2 | 0 |
| TpMuguga_03g00560 | 3 | 0 |
| TpMuguga_03g00560 | 4 | 0 |
| TpMuguga_03g00560 | 5 | 0 |
| TpMuguga_03g00560 | 6 | 0 |
| TpMuguga_03g00560 | 7 | 10.35294118 |
| TpMuguga_03g00561 | 1 | 0.4 |
| TpMuguga_03g00564 | 1 | 0.344988345 |
| TpMuguga_03g00564 | 2 | 0.032634033 |
| TpMuguga_03g00564 | 3 | 0.004662005 |
| TpMuguga_03g02185 | 1 | 0.08183509 |
| TpMuguga_03g02185 | 2 | 0.068815871 |
| TpMuguga_03g00565 | 1 | 0 |
| TpMuguga_03g00565 | 2 | 0.01394214 |

| | | |
|---|---|---|
| TpMuguga_03g00565 | 3 | 0.031369815 |
| TpMuguga_03g00565 | 4 | 0.017427675 |
| TpMuguga_03g00565 | 5 | 0.099337748 |
| TpMuguga_03g00565 | 6 | 0.008713838 |
| TpMuguga_03g00565 | 7 | 0.127222029 |
| TpMuguga_03g00565 | 8 | 0.247472987 |
| TpMuguga_03g00565 | 9 | 6.483095155 |
| TpMuguga_03g00567 | 1 | 0 |
| TpMuguga_03g00567 | 2 | 0 |
| TpMuguga_03g00566 | 1 | 0 |
| TpMuguga_03g02660 | 1 | 0 |
| TpMuguga_03g00569 | 1 | 0 |
| TpMuguga_03g00569 | 2 | 0 |
| TpMuguga_03g00569 | 3 | 2.25 |
| TpMuguga_03g00569 | 4 | 0 |
| TpMuguga_03g00569 | 5 | 1.5 |
| TpMuguga_03g00571 | 1 | 0 |
| TpMuguga_03g00571 | 2 | 0 |
| TpMuguga_03g00571 | 3 | 0 |
| TpMuguga_03g00571 | 4 | 0 |
| TpMuguga_03g00571 | 5 | 0 |
| TpMuguga_03g02500 | 1 | 0.25 |
| TpMuguga_03g02385 | 1 | 9.090909091 |
| TpMuguga_03g02385 | 2 | 4.545454545 |
| TpMuguga_03g02385 | 3 | 0 |
| TpMuguga_03g02385 | 4 | 0 |
| TpMuguga_03g02390 | 1 | 8.181818182 |
| TpMuguga_03g02390 | 2 | 0 |
| TpMuguga_03g02390 | 3 | 0 |
| TpMuguga_03g02390 | 4 | 0 |
| TpMuguga_03g02390 | 5 | 0 |
| TpMuguga_03g02390 | 6 | 0 |
| TpMuguga_03g02390 | 7 | 0 |
| TpMuguga_03g02390 | 8 | 0 |
| TpMuguga_03g02395 | 1 | 0 |
| TpMuguga_03g02395 | 2 | 0 |
| TpMuguga_03g02395 | 3 | 0 |
| TpMuguga_03g02395 | 4 | 0 |
| TpMuguga_03g02395 | 5 | 0 |
| TpMuguga_03g02395 | 6 | 0 |
| TpMuguga_03g02395 | 7 | 0 |
| TpMuguga_03g02395 | 8 | 0 |

| | | |
|---|---|---|
| TpMuguga_03g02395 | 9 | 0 |
| TpMuguga_03g02395 | 10 | 0 |
| TpMuguga_03g02395 | 11 | 0 |
| TpMuguga_03g02395 | 12 | 0 |
| TpMuguga_03g02395 | 13 | 0 |
| TpMuguga_03g02395 | 14 | 0 |
| TpMuguga_03g00577 | 1 | 0 |
| TpMuguga_03g00577 | 2 | 0 |
| TpMuguga_03g00576 | 1 | 0 |
| TpMuguga_03g00576 | 2 | 0.516129032 |
| TpMuguga_03g00576 | 3 | 0.387096774 |
| TpMuguga_03g00576 | 4 | 0.215053763 |
| TpMuguga_03g00576 | 5 | 0 |
| TpMuguga_03g00576 | 6 | 0 |
| TpMuguga_03g00576 | 7 | 0 |
| TpMuguga_03g00576 | 8 | 0 |
| TpMuguga_03g00576 | 9 | 0 |
| TpMuguga_03g00576 | 10 | 0 |
| TpMuguga_03g00576 | 11 | 0.301075269 |
| TpMuguga_03g00578 | 1 | 2.954477834 |
| TpMuguga_03g00578 | 2 | 5.611425171 |
| TpMuguga_03g00578 | 3 | 0.324308242 |
| TpMuguga_03g00578 | 4 | 0.092234454 |
| TpMuguga_03g00578 | 5 | 0.089259149 |
| TpMuguga_03g00578 | 6 | 0.160666468 |
| TpMuguga_03g00578 | 7 | 0 |
| TpMuguga_03g00578 | 8 | 0.157691163 |
| TpMuguga_03g00578 | 9 | 0.348110681 |
| TpMuguga_03g02450 | 1 | 0.630541872 |
| TpMuguga_03g02450 | 2 | 0.866995074 |
| TpMuguga_03g02450 | 3 | 0.197044335 |
| TpMuguga_03g02450 | 4 | 0 |
| TpMuguga_03g02450 | 5 | 0 |
| TpMuguga_03g02450 | 6 | 0.118226601 |
| TpMuguga_03g02450 | 7 | 0.236453202 |
| TpMuguga_03g02530 | 1 | 0.157549234 |
| TpMuguga_03g02530 | 2 | 0.111597374 |
| TpMuguga_03g02530 | 3 | 0.210065646 |
| TpMuguga_03g02530 | 4 | 0.433260394 |
| TpMuguga_03g02530 | 5 | 0.433260394 |
| TpMuguga_03g02530 | 6 | 0.433260394 |
| TpMuguga_03g00585 | 1 | 0 |

| | | |
|---|---|---|
| TpMuguga_03g00585 | 2 | 0 |
| TpMuguga_03g00585 | 3 | 0 |
| TpMuguga_03g00585 | 4 | 0 |
| TpMuguga_03g00585 | 5 | 2.333333333 |
| TpMuguga_03g00585 | 6 | 0 |
| TpMuguga_03g00587 | 1 | 0 |
| TpMuguga_03g00587 | 2 | 0 |
| TpMuguga_03g00587 | 3 | 0 |
| TpMuguga_03g00587 | 4 | 0 |
| TpMuguga_03g00587 | 5 | 0 |
| TpMuguga_03g00587 | 6 | 0 |
| TpMuguga_03g00587 | 7 | 0 |
| TpMuguga_03g00587 | 8 | 0 |
| TpMuguga_03g00587 | 9 | 0 |
| TpMuguga_03g00587 | 10 | 0 |
| TpMuguga_03g00587 | 11 | 0 |
| TpMuguga_03g00588 | 1 | 1.142857143 |
| TpMuguga_03g02510 | 1 | 0 |
| TpMuguga_03g02510 | 2 | 0 |
| TpMuguga_03g02510 | 3 | 0 |
| TpMuguga_03g02510 | 4 | 0 |
| TpMuguga_03g02510 | 5 | 2.222222222 |
| TpMuguga_03g02510 | 6 | 0 |
| TpMuguga_03g02510 | 7 | 0 |
| TpMuguga_03g02510 | 8 | 0 |
| TpMuguga_03g02510 | 9 | 0 |
| TpMuguga_03g00592 | 1 | 0.371779141 |
| TpMuguga_03g00592 | 2 | 0.268711656 |
| TpMuguga_03g00593 | 1 | 0 |
| TpMuguga_03g00593 | 2 | 2.666666667 |
| TpMuguga_03g00593 | 3 | 0 |
| TpMuguga_03g00593 | 4 | 0 |
| TpMuguga_03g00593 | 5 | 0 |
| TpMuguga_03g00594 | 1 | 0 |
| TpMuguga_03g00594 | 2 | 0.125 |
| TpMuguga_03g00594 | 3 | 0 |
| TpMuguga_03g00594 | 4 | 0 |
| TpMuguga_03g00597 | 1 | 0 |
| TpMuguga_03g00597 | 2 | 0.177293935 |
| TpMuguga_03g02635 | 1 | 0 |
| TpMuguga_03g02635 | 2 | 0 |
| TpMuguga_03g02635 | 3 | 0 |

| | | |
|---|---|---|
| TpMuguga_03g02635 | 4 | 1.090909091 |
| TpMuguga_03g02635 | 5 | 0 |
| TpMuguga_03g02635 | 6 | 0 |
| TpMuguga_03g02635 | 7 | 0 |
| TpMuguga_03g02635 | 8 | 0 |
| TpMuguga_03g02635 | 9 | 0 |
| TpMuguga_03g02635 | 10 | 0 |
| TpMuguga_03g02635 | 11 | 0 |
| TpMuguga_03g00600 | 1 | 0.620689655 |
| TpMuguga_03g00601 | 1 | 0 |
| TpMuguga_03g00601 | 2 | 1.764705882 |
| TpMuguga_03g00601 | 3 | 0.352941176 |
| TpMuguga_03g00602 | 1 | 0 |
| TpMuguga_03g00603 | 1 | 0 |
| TpMuguga_03g00604 | 1 | 0.030690537 |
| TpMuguga_03g00604 | 2 | 1.204603581 |
| TpMuguga_03g00605 | 1 | 0.585365854 |
| TpMuguga_03g00606 | 1 | 0.470588235 |
| TpMuguga_03g02630 | 1 | 0.070371588 |
| TpMuguga_03g00607 | 1 | 0.152086137 |
| TpMuguga_03g00608 | 1 | 0 |
| TpMuguga_03g00608 | 2 | 0 |
| TpMuguga_03g00609 | 1 | 0.709090909 |
| TpMuguga_03g00609 | 2 | 0.763636364 |
| TpMuguga_03g02340 | 1 | 0 |
| TpMuguga_03g02335 | 1 | 0 |
| TpMuguga_03g02335 | 2 | 0 |
| TpMuguga_03g02515 | 1 | 0 |
| TpMuguga_03g02515 | 2 | 0 |
| TpMuguga_03g02515 | 3 | 0 |
| TpMuguga_03g02515 | 4 | 0 |
| TpMuguga_03g02515 | 5 | 0.461538462 |
| TpMuguga_03g02515 | 6 | 0 |
| TpMuguga_03g00619 | 1 | 0 |
| TpMuguga_03g00620 | 1 | 0.322580645 |
| TpMuguga_03g00620 | 2 | 0 |
| TpMuguga_03g00620 | 3 | 1.677419355 |
| TpMuguga_03g02165 | 1 | 0 |
| TpMuguga_03g00621 | 1 | 0 |
| TpMuguga_03g02005 | 1 | 1.75 |
| TpMuguga_03g00622 | 1 | 0 |
| TpMuguga_03g00622 | 2 | 0 |

| | | |
|---|---|---|
| TpMuguga_03g02080 | 1 | 0.136752137 |
| TpMuguga_03g02080 | 2 | 0 |
| TpMuguga_03g02080 | 3 | 0 |
| TpMuguga_03g00627 | 1 | 0 |
| TpMuguga_03g00627 | 2 | 0 |
| TpMuguga_03g00627 | 3 | 0 |
| TpMuguga_03g00627 | 4 | 0 |
| TpMuguga_03g00627 | 5 | 0 |
| TpMuguga_03g00627 | 6 | 0 |
| TpMuguga_03g00627 | 7 | 0 |
| TpMuguga_03g02100 | 1 | 0.476190476 |
| TpMuguga_03g02100 | 2 | 0 |
| TpMuguga_03g02100 | 3 | 2.53968254 |
| TpMuguga_03g02100 | 4 | 0.317460317 |
| TpMuguga_03g02120 | 1 | 0.3125 |
| TpMuguga_03g02115 | 1 | 0.754475703 |
| TpMuguga_03g02115 | 2 | 0 |
| TpMuguga_03g02115 | 3 | 0 |
| TpMuguga_03g02115 | 4 | 0 |
| TpMuguga_03g02115 | 5 | 0.012787724 |
| TpMuguga_03g02115 | 6 | 0 |
| TpMuguga_03g02115 | 7 | 0 |
| TpMuguga_03g02115 | 8 | 0 |
| TpMuguga_03g02115 | 9 | 0 |
| TpMuguga_03g00629 | 1 | 1.19047619 |
| TpMuguga_03g02125 | 1 | 0.154639175 |
| TpMuguga_03g02125 | 2 | 0.618556701 |
| TpMuguga_03g00635 | 1 | 0 |
| TpMuguga_03g00636 | 1 | 1.212121212 |
| TpMuguga_03g00640 | 1 | 0 |
| TpMuguga_03g00640 | 2 | 0 |
| TpMuguga_03g00640 | 3 | 0 |
| TpMuguga_03g00640 | 4 | 0 |
| TpMuguga_03g00640 | 5 | 0 |
| TpMuguga_03g00640 | 6 | 0 |
| TpMuguga_03g00640 | 7 | 0 |
| TpMuguga_03g00640 | 8 | 0 |
| TpMuguga_03g00640 | 9 | 0 |
| TpMuguga_03g00640 | 10 | 0 |
| TpMuguga_03g00641 | 1 | 0 |
| TpMuguga_03g00641 | 2 | 0 |
| TpMuguga_03g00644 | 1 | 1.505154639 |

| | | |
|---|---|---|
| TpMuguga_03g00644 | 2 | 0 |
| TpMuguga_03g00644 | 3 | 0 |
| TpMuguga_03g00644 | 4 | 0 |
| TpMuguga_03g00644 | 5 | 0 |
| TpMuguga_03g00644 | 6 | 0 |
| TpMuguga_03g00644 | 7 | 0.020618557 |
| TpMuguga_03g00644 | 8 | 0 |
| TpMuguga_03g00644 | 9 | 0 |
| TpMuguga_03g00644 | 10 | 0 |
| TpMuguga_03g00643 | 1 | 0 |
| TpMuguga_03g00643 | 2 | 0.044345898 |
| TpMuguga_03g00643 | 3 | 0.339985218 |
| TpMuguga_03g00643 | 4 | 5.63932003 |
| TpMuguga_03g02055 | 1 | 0 |
| TpMuguga_03g02055 | 2 | 0 |
| TpMuguga_03g02055 | 3 | 0 |
| TpMuguga_03g02055 | 4 | 0 |
| TpMuguga_03g02055 | 5 | 0 |
| TpMuguga_03g02055 | 6 | 0 |
| TpMuguga_03g02055 | 7 | 0 |
| TpMuguga_03g02055 | 8 | 0 |
| TpMuguga_03g02055 | 9 | 0 |
| TpMuguga_03g02055 | 10 | 0 |
| TpMuguga_03g00650 | 1 | 0.6 |
| TpMuguga_03g00650 | 2 | 6.6 |
| TpMuguga_03g00651 | 1 | 0 |
| TpMuguga_03g00651 | 2 | 0 |
| TpMuguga_03g00651 | 3 | 0 |
| TpMuguga_03g00651 | 4 | 0.416666667 |
| TpMuguga_03g00652 | 1 | 0.002816901 |
| TpMuguga_03g00653 | 1 | 1.208287895 |
| TpMuguga_03g00654 | 1 | 2.628571429 |
| TpMuguga_03g00654 | 2 | 0 |
| TpMuguga_03g00654 | 3 | 0 |
| TpMuguga_03g00654 | 4 | 0 |
| TpMuguga_03g00654 | 5 | 0 |
| TpMuguga_03g00654 | 6 | 0 |
| TpMuguga_03g00654 | 7 | 0 |
| TpMuguga_03g00654 | 8 | 0 |
| TpMuguga_03g00654 | 9 | 0 |
| TpMuguga_03g00654 | 10 | 0 |
| TpMuguga_03g00654 | 11 | 0 |

| | | |
|---|---|---|
| TpMuguga_03g00654 | 12 | 0 |
| TpMuguga_03g00654 | 13 | 0 |
| TpMuguga_03g00654 | 14 | 0 |
| TpMuguga_03g00654 | 15 | 0 |
| TpMuguga_03g00654 | 16 | 0 |
| TpMuguga_03g00654 | 17 | 10.51428571 |
| TpMuguga_03g00654 | 18 | 13.47142857 |
| TpMuguga_03g00654 | 19 | 3.614285714 |
| TpMuguga_03g00654 | 20 | 7.228571429 |
| TpMuguga_03g00654 | 21 | 2.628571429 |
| TpMuguga_03g00654 | 22 | 3.942857143 |
| TpMuguga_03g02075 | 1 | 0 |
| TpMuguga_03g02075 | 2 | 2 |
| TpMuguga_03g00655 | 1 | 0 |
| TpMuguga_03g00655 | 2 | 1.5 |
| TpMuguga_03g00656 | 1 | 0 |
| TpMuguga_03g00656 | 2 | 0 |
| TpMuguga_03g00656 | 3 | 0 |
| TpMuguga_03g00656 | 4 | 0 |
| TpMuguga_03g00656 | 5 | 0 |
| TpMuguga_03g00656 | 6 | 0 |
| TpMuguga_03g00658 | 1 | 0 |
| TpMuguga_03g02140 | 1 | 0 |
| TpMuguga_03g02140 | 2 | 0 |
| TpMuguga_03g02140 | 3 | 0 |
| TpMuguga_03g02140 | 4 | 0 |
| TpMuguga_03g02140 | 5 | 0 |
| TpMuguga_03g02140 | 6 | 0 |
| TpMuguga_03g02140 | 7 | 0 |
| TpMuguga_03g02140 | 8 | 0 |
| TpMuguga_03g02140 | 9 | 0 |
| TpMuguga_03g02140 | 10 | 9 |
| TpMuguga_03g02140 | 11 | 0 |
| TpMuguga_03g02140 | 12 | 0 |
| TpMuguga_03g02140 | 13 | 0 |
| TpMuguga_03g02140 | 14 | 0 |
| TpMuguga_03g02140 | 15 | 0 |
| TpMuguga_03g02140 | 16 | 0 |
| TpMuguga_03g02140 | 17 | 9 |
| TpMuguga_03g00663 | 1 | 0 |
| TpMuguga_03g00663 | 2 | 0 |
| TpMuguga_03g00663 | 3 | 0 |

| | | |
|---|---|---|
| TpMuguga_03g00663 | 4 | 0 |
| TpMuguga_03g00665 | 1 | 0 |
| TpMuguga_03g00666 | 1 | 0 |
| TpMuguga_03g00666 | 2 | 0 |
| TpMuguga_03g00666 | 3 | 4.976076555 |
| TpMuguga_03g00666 | 4 | 2.488038278 |
| TpMuguga_03g02030 | 1 | 0 |
| TpMuguga_03g02030 | 2 | 0.2 |
| TpMuguga_03g02030 | 3 | 3 |
| TpMuguga_03g00671 | 1 | 0 |
| TpMuguga_03g00671 | 2 | 0 |
| TpMuguga_03g00672 | 1 | 2.790697674 |
| TpMuguga_03g00672 | 2 | 0 |
| TpMuguga_03g00672 | 3 | 0 |
| TpMuguga_03g00672 | 4 | 1.046511628 |
| TpMuguga_03g00672 | 5 | 0 |
| TpMuguga_03g00672 | 6 | 0 |
| TpMuguga_03g00672 | 7 | 0 |
| TpMuguga_03g00672 | 8 | 0 |
| TpMuguga_03g00672 | 9 | 0 |
| TpMuguga_03g00672 | 10 | 0 |
| TpMuguga_03g00672 | 11 | 0 |
| TpMuguga_03g00672 | 12 | 0 |
| TpMuguga_03g00672 | 13 | 0 |
| TpMuguga_03g00672 | 14 | 0 |
| TpMuguga_03g00673 | 1 | 0 |
| TpMuguga_03g00673 | 2 | 0 |
| TpMuguga_03g00673 | 3 | 0 |
| TpMuguga_03g00674 | 1 | 2.307692308 |
| TpMuguga_03g00674 | 2 | 0 |
| TpMuguga_03g00674 | 3 | 0 |
| TpMuguga_03g00674 | 4 | 0 |
| TpMuguga_03g00675 | 1 | 0 |
| TpMuguga_03g00676 | 1 | 0 |
| TpMuguga_03g00676 | 2 | 0 |
| TpMuguga_03g00677 | 1 | 0 |
| TpMuguga_03g00678 | 1 | 0 |
| TpMuguga_03g00679 | 1 | 0 |
| TpMuguga_03g00679 | 2 | 0 |
| TpMuguga_03g00680 | 1 | 0.444444444 |
| TpMuguga_03g00681 | 1 | 1.8 |
| TpMuguga_03g00681 | 2 | 0.4 |

| | | |
|---|---|---|
| TpMuguga_03g00682 | 1 | 0 |
| TpMuguga_03g00682 | 2 | 0 |
| TpMuguga_03g00683 | 1 | 0 |
| TpMuguga_03g00683 | 2 | 0 |
| TpMuguga_03g00683 | 3 | 0 |
| TpMuguga_03g00684 | 1 | 0.205263158 |
| TpMuguga_03g00684 | 2 | 0.6 |
| TpMuguga_03g02105 | 1 | 0.967741935 |
| TpMuguga_03g02105 | 2 | 6.290322581 |
| TpMuguga_03g02020 | 1 | 0 |
| TpMuguga_03g02020 | 2 | 0 |
| TpMuguga_03g02020 | 3 | 0 |
| TpMuguga_03g02020 | 4 | 0 |
| TpMuguga_03g02020 | 5 | 0 |
| TpMuguga_03g00689 | 1 | 1.171717172 |
| TpMuguga_03g00689 | 2 | 0.888888889 |
| TpMuguga_03g00689 | 3 | 0 |
| TpMuguga_03g00690 | 1 | 0 |
| TpMuguga_03g00690 | 2 | 4 |
| TpMuguga_03g00690 | 3 | 0 |
| TpMuguga_03g00690 | 4 | 2 |
| TpMuguga_03g00691 | 1 | 0 |
| TpMuguga_03g00692 | 1 | 105 |
| TpMuguga_03g00692 | 2 | 0 |
| TpMuguga_03g00692 | 3 | 0 |
| TpMuguga_03g00692 | 4 | 0 |
| TpMuguga_03g00692 | 5 | 0 |
| TpMuguga_03g00692 | 6 | 0 |
| TpMuguga_03g00692 | 7 | 0 |
| TpMuguga_03g00692 | 8 | 0 |
| TpMuguga_03g00692 | 9 | 0 |
| TpMuguga_03g00692 | 10 | 0 |
| TpMuguga_03g00692 | 11 | 0 |
| TpMuguga_03g00692 | 12 | 0 |
| TpMuguga_03g00692 | 13 | 0 |
| TpMuguga_03g00692 | 14 | 0 |
| TpMuguga_03g00692 | 15 | 0 |
| TpMuguga_03g00692 | 16 | 0 |
| TpMuguga_03g00692 | 17 | 0 |
| TpMuguga_03g00692 | 18 | 0 |
| TpMuguga_03g00692 | 19 | 0 |
| TpMuguga_03g00692 | 20 | 0 |

| | | |
|---|---|---|
| TpMuguga_03g00693 | 1 | 0 |
| TpMuguga_03g00693 | 2 | 0 |
| TpMuguga_03g00693 | 3 | 52 |
| TpMuguga_03g00693 | 4 | 64 |
| TpMuguga_03g00693 | 5 | 0 |
| TpMuguga_03g00693 | 6 | 0 |
| TpMuguga_03g00693 | 7 | 0 |
| TpMuguga_03g02015 | 1 | 0 |
| TpMuguga_03g02015 | 2 | 0 |
| TpMuguga_03g00694 | 1 | 0 |
| TpMuguga_03g00694 | 2 | 0 |
| TpMuguga_03g00694 | 3 | 0 |
| TpMuguga_03g00694 | 4 | 18 |
| TpMuguga_03g00694 | 5 | 0 |
| TpMuguga_03g00695 | 1 | 8.647058824 |
| TpMuguga_03g00695 | 2 | 5.705882353 |
| TpMuguga_03g00695 | 3 | 0 |
| TpMuguga_03g00695 | 4 | 0 |
| TpMuguga_03g02065 | 1 | 0.75 |
| TpMuguga_03g02065 | 2 | 2.25 |
| TpMuguga_03g00698 | 1 | 0 |
| TpMuguga_03g00699 | 1 | 3.5 |
| TpMuguga_03g00699 | 2 | 7 |
| TpMuguga_03g00699 | 3 | 5.25 |
| TpMuguga_03g00699 | 4 | 3.5 |
| TpMuguga_03g00699 | 5 | 3.5 |
| TpMuguga_03g00699 | 6 | 0 |
| TpMuguga_03g00699 | 7 | 1.75 |
| TpMuguga_03g00699 | 8 | 0 |
| TpMuguga_03g00699 | 9 | 0 |
| TpMuguga_03g00699 | 10 | 0 |
| TpMuguga_03g00699 | 11 | 0 |
| TpMuguga_03g00699 | 12 | 0 |
| TpMuguga_03g00699 | 13 | 0 |
| TpMuguga_03g00700 | 1 | 2.315789474 |
| TpMuguga_03g00700 | 2 | 2.526315789 |
| TpMuguga_03g00700 | 3 | 2.105263158 |
| TpMuguga_03g02045 | 1 | 0 |
| TpMuguga_03g02045 | 2 | 0 |
| TpMuguga_03g02045 | 3 | 0 |
| TpMuguga_03g00701 | 1 | 3.970588235 |
| TpMuguga_03g00701 | 2 | 0.735294118 |

| | | |
|---|---|---|
| TpMuguga_03g00701 | 3 | 0 |
| TpMuguga_03g00701 | 4 | 0 |
| TpMuguga_03g00702 | 1 | 0 |
| TpMuguga_03g00702 | 2 | 0 |
| TpMuguga_03g00702 | 3 | 0 |
| TpMuguga_03g00703 | 1 | 0 |
| TpMuguga_03g00703 | 2 | 0 |
| TpMuguga_03g00703 | 3 | 0 |
| TpMuguga_03g00703 | 4 | 0 |
| TpMuguga_03g00703 | 5 | 0 |
| TpMuguga_03g00703 | 6 | 0 |
| TpMuguga_03g00703 | 7 | 0 |
| TpMuguga_03g00703 | 8 | 0 |
| TpMuguga_03g00703 | 9 | 0 |
| TpMuguga_03g00703 | 10 | 0 |
| TpMuguga_03g00703 | 11 | 0 |
| TpMuguga_03g00704 | 1 | 0 |
| TpMuguga_03g00704 | 2 | 1.2 |
| TpMuguga_03g00705 | 1 | 0 |
| TpMuguga_03g00705 | 2 | 5.8125 |
| TpMuguga_03g02050 | 1 | 4 |
| TpMuguga_03g02050 | 2 | 8.666666667 |
| TpMuguga_03g02050 | 3 | 0 |
| TpMuguga_03g02050 | 4 | 0 |
| TpMuguga_03g02050 | 5 | 0.666666667 |
| TpMuguga_03g02050 | 6 | 1.333333333 |
| TpMuguga_03g02050 | 7 | 4.333333333 |
| TpMuguga_03g02050 | 8 | 0 |
| TpMuguga_03g02050 | 9 | 0 |
| TpMuguga_03g02050 | 10 | 0 |
| TpMuguga_03g02050 | 11 | 0 |
| TpMuguga_03g02050 | 12 | 0 |
| TpMuguga_03g02050 | 13 | 0 |
| TpMuguga_03g02050 | 14 | 0 |
| TpMuguga_03g00709 | 1 | 1.833333333 |
| TpMuguga_03g00709 | 2 | 2.566666667 |
| TpMuguga_03g00709 | 3 | 0 |
| TpMuguga_03g00709 | 4 | 0 |
| TpMuguga_03g00709 | 5 | 0 |
| TpMuguga_03g00709 | 6 | 0 |
| TpMuguga_03g00709 | 7 | 0 |
| TpMuguga_03g00709 | 8 | 4.4 |

| | | |
|---|---|---|
| TpMuguga_03g00709 | 9 | 1.466666667 |
| TpMuguga_03g00709 | 10 | 0 |
| TpMuguga_03g00710 | 1 | 2.16 |
| TpMuguga_03g00710 | 2 | 1.92 |
| TpMuguga_03g00710 | 3 | 1.52 |
| TpMuguga_03g00712 | 1 | 4.973811165 |
| TpMuguga_03g00712 | 2 | 0.231564438 |
| TpMuguga_03g00712 | 3 | 0 |
| TpMuguga_03g00712 | 4 | 0 |
| TpMuguga_03g00712 | 5 | 0 |
| TpMuguga_03g00712 | 6 | 0 |
| TpMuguga_03g00713 | 1 | 0 |
| TpMuguga_03g00713 | 2 | 0 |
| TpMuguga_03g00714 | 1 | 0 |
| TpMuguga_03g00714 | 2 | 1.066666667 |
| TpMuguga_03g00714 | 3 | 3.2 |
| TpMuguga_03g00714 | 4 | 9.6 |
| TpMuguga_03g00714 | 5 | 0 |
| TpMuguga_03g00714 | 6 | 0 |
| TpMuguga_03g00714 | 7 | 0 |
| TpMuguga_03g00715 | 1 | 0 |
| TpMuguga_03g00715 | 2 | 0 |
| TpMuguga_03g00715 | 3 | 0 |
| TpMuguga_03g00715 | 4 | 0.222222222 |
| TpMuguga_03g00715 | 5 | 1.777777778 |
| TpMuguga_03g00715 | 6 | 0.055555556 |
| TpMuguga_03g00715 | 7 | 0.055555556 |
| TpMuguga_03g00715 | 8 | 1.277777778 |
| TpMuguga_03g00716 | 1 | 0 |
| TpMuguga_03g02060 | 1 | 0.96969697 |
| TpMuguga_03g02095 | 1 | 0 |
| TpMuguga_03g02095 | 2 | 3.2 |
| TpMuguga_03g02095 | 3 | 0 |
| TpMuguga_03g00720 | 1 | 0.25 |
| TpMuguga_03g00724 | 1 | 0 |
| TpMuguga_03g00724 | 2 | 0 |
| TpMuguga_03g00724 | 3 | 0 |
| TpMuguga_03g00724 | 4 | 9 |
| TpMuguga_03g00724 | 5 | 0 |
| TpMuguga_03g00728 | 1 | 0 |
| TpMuguga_03g00728 | 2 | 0.12195122 |
| TpMuguga_03g00728 | 3 | 0.518292683 |

| | | |
|---|---|---|
| TpMuguga_03g00728 | 4 | 0.426829268 |
| TpMuguga_03g00730 | 1 | 0 |
| TpMuguga_03g00730 | 2 | 0 |
| TpMuguga_03g00730 | 3 | 0 |
| TpMuguga_03g00730 | 4 | 0 |
| TpMuguga_03g00733 | 1 | 0.091086532 |
| TpMuguga_03g00733 | 2 | 0 |
| TpMuguga_03g00733 | 3 | 0 |
| TpMuguga_03g00733 | 4 | 0 |
| TpMuguga_03g00733 | 5 | 0.009108653 |
| TpMuguga_03g00733 | 6 | 0 |
| TpMuguga_03g00732 | 1 | 0 |
| TpMuguga_03g00732 | 2 | 0 |
| TpMuguga_03g00732 | 3 | 0.285714286 |
| TpMuguga_03g00736 | 1 | 2.888888889 |
| TpMuguga_03g00735 | 1 | 0 |
| TpMuguga_03g00735 | 2 | 0 |
| TpMuguga_03g00735 | 3 | 0 |
| TpMuguga_03g00734 | 1 | 1.384615385 |
| TpMuguga_03g00734 | 2 | 0 |
| TpMuguga_03g00734 | 3 | 0 |
| TpMuguga_03g00734 | 4 | 0 |
| TpMuguga_03g00734 | 5 | 0 |
| TpMuguga_03g00734 | 6 | 0 |
| TpMuguga_03g00734 | 7 | 0 |
| TpMuguga_03g00734 | 8 | 0 |
| TpMuguga_03g02155 | 1 | 0 |
| TpMuguga_03g02155 | 2 | 8.368421053 |
| TpMuguga_03g00741 | 1 | 0 |
| TpMuguga_03g00741 | 2 | 0 |
| TpMuguga_03g00742 | 1 | 0 |
| TpMuguga_03g00742 | 2 | 35 |
| TpMuguga_03g00742 | 3 | 45 |
| TpMuguga_03g00742 | 4 | 20 |
| TpMuguga_03g00744 | 1 | 0.757575758 |
| TpMuguga_03g00744 | 2 | 0.303030303 |
| TpMuguga_03g00744 | 3 | 0.606060606 |
| TpMuguga_03g00744 | 4 | 0 |
| TpMuguga_03g00745 | 1 | 0 |
| TpMuguga_03g00745 | 2 | 0 |
| TpMuguga_03g00746 | 1 | 0 |
| TpMuguga_03g00746 | 2 | 0 |

| | | |
|---|---|---|
| TpMuguga_03g00746 | 3 | 4 |
| TpMuguga_03g00747 | 1 | 0 |
| TpMuguga_03g00747 | 2 | 0 |
| TpMuguga_03g00747 | 3 | 0 |
| TpMuguga_03g00747 | 4 | 1.5 |
| TpMuguga_03g00747 | 5 | 0 |
| TpMuguga_03g00748 | 1 | 0 |
| TpMuguga_03g02035 | 1 | 0 |
| TpMuguga_03g02035 | 2 | 0 |
| TpMuguga_03g00752 | 1 | 2.444444444 |
| TpMuguga_03g00754 | 1 | 0 |
| TpMuguga_03g00754 | 2 | 0 |
| TpMuguga_03g00755 | 1 | 0 |
| TpMuguga_03g00756 | 1 | 0 |
| TpMuguga_03g00756 | 2 | 0 |
| TpMuguga_03g00756 | 3 | 0 |
| TpMuguga_03g00756 | 4 | 0 |
| TpMuguga_03g00756 | 5 | 0 |
| TpMuguga_03g00757 | 1 | 0 |
| TpMuguga_03g00757 | 2 | 8.4375 |
| TpMuguga_03g00757 | 3 | 0 |
| TpMuguga_03g00757 | 4 | 0.625 |
| TpMuguga_03g00758 | 1 | 0 |
| TpMuguga_03g00758 | 2 | 0 |
| TpMuguga_03g00758 | 3 | 0 |
| TpMuguga_03g00758 | 4 | 0 |
| TpMuguga_03g00758 | 5 | 0 |
| TpMuguga_03g02145 | 1 | 0 |
| TpMuguga_03g02145 | 2 | 0 |
| TpMuguga_03g00761 | 1 | 0 |
| TpMuguga_03g00761 | 2 | 0 |
| TpMuguga_03g00761 | 3 | 0 |
| TpMuguga_03g00761 | 4 | 0 |
| TpMuguga_03g00762 | 1 | 0 |
| TpMuguga_03g00762 | 2 | 0 |
| TpMuguga_03g00764 | 1 | 0 |
| TpMuguga_03g00764 | 2 | 0 |
| TpMuguga_03g00764 | 3 | 0 |
| TpMuguga_03g00764 | 4 | 0 |
| TpMuguga_03g00764 | 5 | 0.405797101 |
| TpMuguga_03g00764 | 6 | 0 |
| TpMuguga_03g00765 | 1 | 0 |

| | | |
|---|---|---|
| TpMuguga_03g00765 | 2 | 3.555555556 |
| TpMuguga_03g00765 | 3 | 0 |
| TpMuguga_03g00766 | 1 | 1.333333333 |
| TpMuguga_03g00766 | 2 | 2.512820513 |
| TpMuguga_03g00766 | 3 | 0 |
| TpMuguga_03g00767 | 1 | 0 |
| TpMuguga_03g00767 | 2 | 0 |
| TpMuguga_03g00767 | 3 | 0 |
| TpMuguga_03g00767 | 4 | 0 |
| TpMuguga_03g00767 | 5 | 0 |
| TpMuguga_03g00767 | 6 | 0 |
| TpMuguga_03g00767 | 7 | 0 |
| TpMuguga_03g00767 | 8 | 0 |
| TpMuguga_03g00768 | 1 | 0 |
| TpMuguga_03g00768 | 2 | 0 |
| TpMuguga_03g00768 | 3 | 0 |
| TpMuguga_03g00768 | 4 | 0 |
| TpMuguga_03g00768 | 5 | 0 |
| TpMuguga_03g00768 | 6 | 0 |
| TpMuguga_03g00768 | 7 | 0 |
| TpMuguga_03g00768 | 8 | 0 |
| TpMuguga_03g00769 | 1 | 0 |
| TpMuguga_03g00769 | 2 | 0 |
| TpMuguga_03g00769 | 3 | 0 |
| TpMuguga_03g02085 | 1 | 0 |
| TpMuguga_03g02085 | 2 | 0 |
| TpMuguga_03g02085 | 3 | 0 |
| TpMuguga_03g02085 | 4 | 0 |
| TpMuguga_03g02085 | 5 | 0 |
| TpMuguga_03g02085 | 6 | 0 |
| TpMuguga_03g02085 | 7 | 0 |
| TpMuguga_03g02085 | 8 | 0 |
| TpMuguga_03g02085 | 9 | 0 |
| TpMuguga_03g02085 | 10 | 0 |
| TpMuguga_03g02085 | 11 | 0 |
| TpMuguga_03g02085 | 12 | 0 |
| TpMuguga_03g00773 | 1 | 0 |
| TpMuguga_03g00773 | 2 | 0 |
| TpMuguga_03g00773 | 3 | 32 |
| TpMuguga_03g00773 | 4 | 0 |
| TpMuguga_03g00773 | 5 | 0 |
| TpMuguga_03g00773 | 6 | 32 |

| | | |
|---|---|---|
| TpMuguga_03g00773 | 7 | 56 |
| TpMuguga_03g00774 | 1 | 0 |
| TpMuguga_03g00774 | 2 | 0 |
| TpMuguga_03g00774 | 3 | 0 |
| TpMuguga_03g00774 | 4 | 0 |
| TpMuguga_03g00774 | 5 | 7 |
| TpMuguga_03g00774 | 6 | 14 |
| TpMuguga_03g00775 | 1 | 0 |
| TpMuguga_03g00775 | 2 | 0 |
| TpMuguga_03g00775 | 3 | 0.833333333 |
| TpMuguga_03g00775 | 4 | 3.75 |
| TpMuguga_03g00776 | 1 | 0 |
| TpMuguga_03g00776 | 2 | 0 |
| TpMuguga_03g00776 | 3 | 0 |
| TpMuguga_03g00776 | 4 | 0 |
| TpMuguga_03g00776 | 5 | 0 |
| TpMuguga_03g00776 | 6 | 0 |
| TpMuguga_03g00776 | 7 | 0 |
| TpMuguga_03g00776 | 8 | 0 |
| TpMuguga_03g00776 | 9 | 0 |
| TpMuguga_03g00777 | 1 | 0 |
| TpMuguga_03g00777 | 2 | 0 |
| TpMuguga_03g00777 | 3 | 0 |
| TpMuguga_03g00777 | 4 | 0 |
| TpMuguga_03g00777 | 5 | 0 |
| TpMuguga_03g00777 | 6 | 0 |
| TpMuguga_03g00777 | 7 | 0 |
| TpMuguga_03g00777 | 8 | 0 |
| TpMuguga_03g00778 | 1 | 0 |
| TpMuguga_03g00778 | 2 | 0 |
| TpMuguga_03g00778 | 3 | 0 |
| TpMuguga_03g00778 | 4 | 0 |
| TpMuguga_03g00778 | 5 | 0 |
| TpMuguga_03g00778 | 6 | 0 |
| TpMuguga_03g00778 | 7 | 0 |
| TpMuguga_03g00778 | 8 | 0 |
| TpMuguga_03g00779 | 1 | 0.858044164 |
| TpMuguga_03g00780 | 1 | 0 |
| TpMuguga_03g00780 | 2 | 0 |
| TpMuguga_03g00780 | 3 | 0 |
| TpMuguga_03g00780 | 4 | 0 |
| TpMuguga_03g00780 | 5 | 0 |

| | | |
|---|---|---|
| TpMuguga_03g00781 | 1 | 0 |
| TpMuguga_03g00781 | 2 | 0 |
| TpMuguga_03g00781 | 3 | 0 |
| TpMuguga_03g00781 | 4 | 0 |
| TpMuguga_03g00781 | 5 | 0 |
| TpMuguga_03g00782 | 1 | 0 |
| TpMuguga_03g00782 | 2 | 0 |
| TpMuguga_03g00782 | 3 | 0 |
| TpMuguga_03g00782 | 4 | 0 |
| TpMuguga_03g00782 | 5 | 0 |
| TpMuguga_03g00782 | 6 | 0 |
| TpMuguga_03g00782 | 7 | 0 |
| TpMuguga_03g00782 | 8 | 0 |
| TpMuguga_03g00782 | 9 | 0 |
| TpMuguga_03g00782 | 10 | 0 |
| TpMuguga_03g00782 | 11 | 0 |
| TpMuguga_03g02010 | 1 | 0 |
| TpMuguga_03g02010 | 2 | 0 |
| TpMuguga_03g02010 | 3 | 0 |
| TpMuguga_03g02010 | 4 | 0 |
| TpMuguga_03g02010 | 5 | 0.28 |
| TpMuguga_03g02010 | 6 | 0.466666667 |
| TpMuguga_03g00787 | 1 | 0 |
| TpMuguga_03g00787 | 2 | 0 |
| TpMuguga_03g00789 | 1 | 0 |
| TpMuguga_03g00789 | 2 | 0 |
| TpMuguga_03g00789 | 3 | 0 |
| TpMuguga_03g00789 | 4 | 0 |
| TpMuguga_03g00789 | 5 | 0 |
| TpMuguga_03g00789 | 6 | 0 |
| TpMuguga_03g00789 | 7 | 0 |
| TpMuguga_03g00789 | 8 | 0 |
| TpMuguga_03g00789 | 9 | 0 |
| TpMuguga_03g00790 | 1 | 0.5 |
| TpMuguga_03g00791 | 1 | 0.162162162 |
| TpMuguga_03g00791 | 2 | 0.567567568 |
| TpMuguga_03g00792 | 1 | 0.029126214 |
| TpMuguga_03g00792 | 2 | 0 |
| TpMuguga_03g00792 | 3 | 0.029126214 |
| TpMuguga_03g00792 | 4 | 0 |
| TpMuguga_03g00792 | 5 | 0.174757282 |
| TpMuguga_03g00794 | 1 | 0.3 |

| | | |
|---|---|---|
| TpMuguga_03g00794 | 2 | 0 |
| TpMuguga_03g00793 | 1 | 0 |
| TpMuguga_03g00793 | 2 | 0 |
| TpMuguga_03g00793 | 3 | 0 |
| TpMuguga_03g00793 | 4 | 0 |
| TpMuguga_03g00793 | 5 | 0 |
| TpMuguga_03g00795 | 1 | 1.145833333 |
| TpMuguga_03g00795 | 2 | 0.229166667 |
| TpMuguga_03g00795 | 3 | 0 |
| TpMuguga_03g00795 | 4 | 0 |
| TpMuguga_03g00795 | 5 | 0 |
| TpMuguga_03g00795 | 6 | 0 |
| TpMuguga_03g00795 | 7 | 0 |
| TpMuguga_03g00795 | 8 | 1.375 |
| TpMuguga_03g00795 | 9 | 0 |
| TpMuguga_03g00795 | 10 | 0 |
| TpMuguga_03g00798 | 1 | 1.726572529 |
| TpMuguga_03g00798 | 2 | 0.346598203 |
| TpMuguga_03g00798 | 3 | 0.224646983 |
| TpMuguga_03g00798 | 4 | 0.166880616 |
| TpMuguga_03g02160 | 1 | 0 |
| TpMuguga_03g02160 | 2 | 0.726533379 |
| TpMuguga_03g02160 | 3 | 2.399186716 |
| TpMuguga_03g02040 | 1 | 0 |
| TpMuguga_03g02040 | 2 | 10 |
| TpMuguga_03g02040 | 3 | 0 |
| TpMuguga_03g02040 | 4 | 0 |
| TpMuguga_03g02040 | 5 | 0 |
| TpMuguga_03g02040 | 6 | 0 |
| TpMuguga_03g02040 | 7 | 0 |
| TpMuguga_03g02040 | 8 | 0 |
| TpMuguga_03g02040 | 9 | 0 |
| TpMuguga_03g00801 | 1 | 0 |
| TpMuguga_03g00801 | 2 | 0 |
| TpMuguga_03g00801 | 3 | 0 |
| TpMuguga_03g00801 | 4 | 6 |
| TpMuguga_03g00801 | 5 | 0 |
| TpMuguga_03g02025 | 1 | 0.099730458 |
| TpMuguga_03g00802 | 1 | 1.544554455 |
| TpMuguga_03g00802 | 2 | 1.425742574 |
| TpMuguga_03g00802 | 3 | 0.118811881 |
| TpMuguga_03g00802 | 4 | 0.415841584 |

| | | |
|---|---|---|
| TpMuguga_03g00802 | 5 | 1.188118812 |
| TpMuguga_03g00804 | 1 | 0 |
| TpMuguga_03g00805 | 1 | 0 |
| TpMuguga_03g00805 | 2 | 0 |
| TpMuguga_03g00805 | 3 | 0 |
| TpMuguga_03g00805 | 4 | 0 |
| TpMuguga_03g00805 | 5 | 0 |
| TpMuguga_03g00805 | 6 | 0 |
| TpMuguga_03g00805 | 7 | 0 |
| TpMuguga_03g00805 | 8 | 0 |
| TpMuguga_03g00805 | 9 | 0 |
| TpMuguga_03g00805 | 10 | 0 |
| TpMuguga_03g00806 | 1 | 0 |
| TpMuguga_03g00806 | 2 | 0 |
| TpMuguga_03g00806 | 3 | 0 |
| TpMuguga_03g00809 | 1 | 1.6 |
| TpMuguga_03g00809 | 2 | 1.866666667 |
| TpMuguga_03g00809 | 3 | 0.266666667 |
| TpMuguga_03g00808 | 1 | 0 |
| TpMuguga_03g00808 | 2 | 0 |
| TpMuguga_03g00808 | 3 | 0 |
| TpMuguga_03g00808 | 4 | 0 |
| TpMuguga_03g00808 | 5 | 0 |
| TpMuguga_03g00808 | 6 | 0 |
| TpMuguga_03g00808 | 7 | 0 |
| TpMuguga_03g00808 | 8 | 0 |
| TpMuguga_03g00808 | 9 | 0 |
| TpMuguga_03g00808 | 10 | 0 |
| TpMuguga_03g00808 | 11 | 0 |
| TpMuguga_03g00810 | 1 | 0 |
| TpMuguga_03g00810 | 2 | 0 |
| TpMuguga_03g00810 | 3 | 0 |
| TpMuguga_03g00810 | 4 | 0 |
| TpMuguga_03g00812 | 1 | 1.171875 |
| TpMuguga_03g00812 | 2 | 0.651988636 |
| TpMuguga_03g02110 | 1 | 0 |
| TpMuguga_03g02110 | 2 | 0 |
| TpMuguga_03g02110 | 3 | 0 |
| TpMuguga_03g02110 | 4 | 0 |
| TpMuguga_03g02110 | 5 | 0 |
| TpMuguga_03g02110 | 6 | 0 |
| TpMuguga_03g02110 | 7 | 0 |

| | | |
|---|---|---|
| TpMuguga_03g02110 | 8 | 0 |
| TpMuguga_03g02110 | 9 | 0 |
| TpMuguga_03g02110 | 10 | 0 |
| TpMuguga_03g02110 | 11 | 0 |
| TpMuguga_03g02110 | 12 | 0 |
| TpMuguga_03g02110 | 13 | 0 |
| TpMuguga_03g02110 | 14 | 0 |
| TpMuguga_03g02110 | 15 | 0 |
| TpMuguga_03g02110 | 16 | 0 |
| TpMuguga_03g02110 | 17 | 9.523809524 |
| TpMuguga_03g02110 | 18 | 0 |
| TpMuguga_03g02110 | 19 | 0 |
| TpMuguga_03g00817 | 1 | 0.073972603 |
| TpMuguga_03g00817 | 2 | 0.073972603 |
| TpMuguga_03g00816 | 1 | 2.074829932 |
| TpMuguga_03g00816 | 2 | 0.510204082 |
| TpMuguga_03g00816 | 3 | 0 |
| TpMuguga_03g00816 | 4 | 0.408163265 |
| TpMuguga_03g00818 | 1 | 0.25 |
| TpMuguga_03g00818 | 2 | 1.125 |
| TpMuguga_03g00819 | 1 | 0 |
| TpMuguga_03g00819 | 2 | 0 |
| TpMuguga_03g00819 | 3 | 0 |
| TpMuguga_03g00821 | 1 | 13.33333333 |
| TpMuguga_03g00821 | 2 | 0.666666667 |
| TpMuguga_03g00821 | 3 | 0 |
| TpMuguga_03g00821 | 4 | 0 |
| TpMuguga_03g00821 | 5 | 0 |
| TpMuguga_03g00821 | 6 | 0 |
| TpMuguga_03g00821 | 7 | 4.666666667 |
| TpMuguga_03g02090 | 1 | 0 |
| TpMuguga_03g00823 | 1 | 0 |
| TpMuguga_03g00823 | 2 | 0 |
| TpMuguga_03g00823 | 3 | 0 |
| TpMuguga_03g00823 | 4 | 0 |
| TpMuguga_03g00823 | 5 | 0 |
| TpMuguga_03g00823 | 6 | 0 |
| TpMuguga_03g00823 | 7 | 0 |
| TpMuguga_03g00823 | 8 | 0 |
| TpMuguga_03g00823 | 9 | 0 |
| TpMuguga_03g00823 | 10 | 22 |
| TpMuguga_03g00824 | 1 | 0 |

| | | |
|---|---|---|
| TpMuguga_03g00824 | 2 | 0 |
| TpMuguga_03g00825 | 1 | 2.4 |
| TpMuguga_03g00825 | 2 | 0 |
| TpMuguga_03g00825 | 3 | 0 |
| TpMuguga_03g00826 | 1 | 0 |
| TpMuguga_03g00827 | 1 | 0.5 |
| TpMuguga_03g00827 | 2 | 0 |
| TpMuguga_03g00828 | 1 | 0 |
| TpMuguga_03g00828 | 2 | 0 |
| TpMuguga_03g00828 | 3 | 0 |
| TpMuguga_03g00828 | 4 | 0 |
| TpMuguga_03g00830 | 1 | 0 |
| TpMuguga_03g00830 | 2 | 0 |
| TpMuguga_03g00830 | 3 | 3.75 |
| TpMuguga_03g00830 | 4 | 0 |
| TpMuguga_03g00831 | 1 | 4.628571429 |
| TpMuguga_03g00831 | 2 | 0.685714286 |
| TpMuguga_03g00833 | 1 | 3.498891353 |
| TpMuguga_03g00833 | 2 | 0.00443459 |
| TpMuguga_03g00833 | 3 | 0.033259424 |
| TpMuguga_03g00832 | 1 | 0 |
| TpMuguga_03g00832 | 2 | 0.124600639 |
| TpMuguga_03g00836 | 1 | 0 |
| TpMuguga_03g00836 | 2 | 0 |
| TpMuguga_03g00838 | 1 | 0.571428571 |
| TpMuguga_03g00838 | 2 | 0 |
| TpMuguga_03g00838 | 3 | 0 |
| TpMuguga_03g00838 | 4 | 0 |
| TpMuguga_03g00838 | 5 | 0 |
| TpMuguga_03g00838 | 6 | 0.19047619 |
| TpMuguga_03g00838 | 7 | 0.380952381 |
| TpMuguga_03g00838 | 8 | 2.095238095 |
| TpMuguga_03g00838 | 9 | 2.857142857 |
| TpMuguga_03g00838 | 10 | 0 |
| TpMuguga_03g00838 | 11 | 2.666666667 |
| TpMuguga_03g00840 | 1 | 0.882352941 |
| TpMuguga_03g00840 | 2 | 0 |
| TpMuguga_03g00840 | 3 | 0 |
| TpMuguga_03g00840 | 4 | 0 |
| TpMuguga_03g02000 | 1 | 0.033707865 |
| TpMuguga_03g02000 | 2 | 0.451685393 |
| TpMuguga_03g00841 | 1 | 1.125 |

| | | |
|---|---|---|
| TpMuguga_03g00841 | 2 | 9.3375 |
| TpMuguga_03g00841 | 3 | 4.6125 |
| TpMuguga_03g00841 | 4 | 0.1125 |
| TpMuguga_03g00841 | 5 | 3.7125 |
| TpMuguga_03g00841 | 6 | 1.2375 |
| TpMuguga_03g00841 | 7 | 0 |
| TpMuguga_03g00841 | 8 | 1.6875 |
| TpMuguga_03g00843 | 1 | 0.035294118 |
| TpMuguga_03g00843 | 2 | 0.176470588 |
| TpMuguga_03g00843 | 3 | 0.070588235 |
| TpMuguga_03g00843 | 4 | 0.917647059 |
| TpMuguga_03g00843 | 5 | 9.070588235 |
| TpMuguga_03g00842 | 1 | 0 |
| TpMuguga_03g00842 | 2 | 0 |
| TpMuguga_03g00842 | 3 | 0.008417508 |
| TpMuguga_03g00844 | 1 | 6.025641026 |
| TpMuguga_03g00844 | 2 | 1.41025641 |
| TpMuguga_03g00844 | 3 | 0.170940171 |
| TpMuguga_03g00844 | 4 | 0.299145299 |
| TpMuguga_03g02130 | 1 | 2.8 |
| TpMuguga_03g02130 | 2 | 0 |
| TpMuguga_03g02130 | 3 | 0 |
| TpMuguga_03g00846 | 1 | 0 |
| TpMuguga_03g00846 | 2 | 0 |
| TpMuguga_03g02135 | 1 | 0 |
| TpMuguga_03g02135 | 2 | 15.75 |
| TpMuguga_03g02135 | 3 | 0 |
| TpMuguga_03g02135 | 4 | 1.5 |
| TpMuguga_03g02135 | 5 | 35.25 |
| TpMuguga_03g00847 | 1 | 0 |
| TpMuguga_03g00847 | 2 | 0 |
| TpMuguga_03g00847 | 3 | 0 |
| TpMuguga_03g00847 | 4 | 0 |
| TpMuguga_03g00847 | 5 | 0 |
| TpMuguga_03g00847 | 6 | 0 |
| TpMuguga_03g00847 | 7 | 0 |
| TpMuguga_03g00847 | 8 | 19.125 |
| TpMuguga_03g00848 | 1 | 1.145833333 |
| TpMuguga_03g00848 | 2 | 0.729166667 |
| TpMuguga_03g00848 | 3 | 0.3125 |
| TpMuguga_03g00848 | 4 | 0.208333333 |
| TpMuguga_03g00851 | 1 | 0 |

| | | |
|---|---|---|
| TpMuguga_03g00851 | 2 | 0.081871345 |
| TpMuguga_03g00851 | 3 | 1.30994152 |
| TpMuguga_03g00851 | 4 | 0.040935673 |
| TpMuguga_03g00851 | 5 | 2.169590643 |
| TpMuguga_03g00851 | 6 | 2.169590643 |
| TpMuguga_03g02150 | 1 | 0 |
| TpMuguga_03g02150 | 2 | 10 |
| TpMuguga_03g02150 | 3 | 7 |
| TpMuguga_03g02150 | 4 | 0 |
| TpMuguga_03g00852 | 1 | 0 |
| TpMuguga_03g00852 | 2 | 1.304347826 |
| TpMuguga_03g00852 | 3 | 0.652173913 |
| TpMuguga_03g00852 | 4 | 0.108695652 |
| TpMuguga_03g00854 | 1 | 0 |
| TpMuguga_03g00854 | 2 | 0 |
| TpMuguga_03g00854 | 3 | 6.666666667 |
| TpMuguga_03g00853 | 1 | 0.789473684 |
| TpMuguga_03g00855 | 1 | 1.034482759 |
| TpMuguga_03g00859 | 1 | 0 |
| TpMuguga_03g00859 | 2 | 0 |
| TpMuguga_03g00859 | 3 | 10.6875 |
| TpMuguga_03g00859 | 4 | 2.8125 |
| TpMuguga_03g00859 | 5 | 5.0625 |
| TpMuguga_03g00859 | 6 | 5.625 |
| TpMuguga_03g00859 | 7 | 8.4375 |
| TpMuguga_03g00859 | 8 | 0 |
| TpMuguga_03g00858 | 1 | 1.129251701 |
| TpMuguga_03g00858 | 2 | 0.149659864 |
| TpMuguga_03g00858 | 3 | 0 |
| TpMuguga_03g00857 | 1 | 0 |
| TpMuguga_03g00857 | 2 | 0.004008016 |
| TpMuguga_03g00857 | 3 | 0 |
| TpMuguga_03g00857 | 4 | 0.006012024 |
| TpMuguga_03g00857 | 5 | 0.006012024 |
| TpMuguga_03g00857 | 6 | 0.04008016 |
| TpMuguga_03g00857 | 7 | 0.024048096 |
| TpMuguga_03g00857 | 8 | 0.018036072 |
| TpMuguga_03g00857 | 9 | 1.390781563 |
| TpMuguga_03g00929 | 1 | 0.440366972 |
| TpMuguga_03g00929 | 2 | 0 |
| TpMuguga_03g00860 | 1 | 32 |
| TpMuguga_03g00860 | 2 | 0 |

| | | |
|---|---|---|
| TpMuguga_03g00860 | 3 | 0 |
| TpMuguga_03g00868 | 1 | 0 |
| TpMuguga_03g00869 | 1 | 1.166666667 |
| TpMuguga_03g00869 | 2 | 0 |
| TpMuguga_03g00872 | 1 | 0.833333333 |
| TpMuguga_03g00872 | 2 | 2.833333333 |
| TpMuguga_03g00875 | 1 | 0 |
| TpMuguga_03g00875 | 2 | 0.172131148 |
| TpMuguga_03g00876 | 1 | 1 |
| TpMuguga_03g00877 | 1 | 2 |
| TpMuguga_03g00880 | 1 | 0 |
| TpMuguga_03g00880 | 2 | 0 |
| TpMuguga_03g00890 | 1 | 0.620689655 |
| TpMuguga_03g00891 | 1 | 0.526315789 |
